# A three-dimensional histological cell atlas of the developing human brain

**DOI:** 10.1101/2024.12.17.628811

**Authors:** Jaikishan Jayakumar, Mohanasankar Sivaprakasam, Richa Verma, Mihail Bota, Jayaraj Joseph, Supriti Mulay, Jayaraman Kumutha, Chitra Srinivasan, S Suresh, S. Latha, Harish E Kumar, Aparna Bhaduri, Tomasz J. Nowakowski, Prasun K Roy, Stephen Savoia, Samik Banerjee, Daniel Tward, Partha P Mitra

## Abstract

The human brain is believed to contain a full complement of neurons by the time of birth together with a substantial amount of the connectivity architecture, even though a significant amount of growth occurs postnatally. The developmental process leading to this outcome is not well understood in humans in comparison with model organisms. Previous magnetic resonance imaging (MRI) studies give three-dimensional coverage but not cellular resolution. In contrast, sparsely sampled histological or spatial omics analyses have provided cellular resolution but not dense whole brain coverage. To address the unmet need to provide a quantitative spatiotemporal map of developing human brain at cellular resolution, we leveraged tape-transfer assisted serial section histology to obtain contiguous histological series and unbiased imaging with dense coverage. Interleaved 20μ thick Nissl and H&E series and MRI volumes are co-registered into multimodal reference volumes with 60μ isotropic resolution, together with atlas annotations and a stereotactic coordinate system based on skull landmarks. The histological atlas volumes have significantly more contrast and texture than the MRI volumes. We computationally detect cells brain-wide to obtain quantitative characterization of the cytoarchitecture of the developing brain at 13-14 and 20-21 gestational weeks, providing the first comprehensive regional cell counts and characterizing the differential growth of the different brain compartments. Morphological characteristics permit segmentation of cell types from histology. We detected and quantified brain-wide distribution of mitotic figures representing dividing cells, providing an unprecedented spatiotemporal atlas of proliferative dynamics in the developing human brain. Further, we characterized the abundance and distribution of Cajal-Retzius cells, a transient cell population that plays essential roles in organizing glutamatergic cortical neurons into layers. Together, our study provides an unprecedented quantitative window into the developing human brain and the reference volumes and coordinate space should be useful for integrating spatial omics data sets with dense histological context.

## Introduction

Humans are born with disproportionately large brains and neuronal cell and connectivity architectures more closely resembling its mature form compared to other organs of the body. The neonatal brain is believed to have approximately the same number of neurons as the adult brain, with much of the larger-scale connectivity patterns in place. Understanding the complex developmental programs of cell division, migration, differentiation, the growth of neural processes, and establishment of network connectivity is essential for studies of the mechanisms of brain evolution and neuropathology of neurodevelopmental disorders. However, we know very little about the development of human brains compared to that of laboratory animals such as rodents, particularly with detailed cellular resolution. Existing histological atlases are sparsely sampled, lack multimodality and a coordinate system determined from skull landmarks as is the standard for laboratory animals. Elementary facts, such as the number of cells in the growing brain, their spatial distribution and patterns of migration are only partially known based on sparse sampling. A densely sampled multimodal histological dataset at multiple stages subjected to quantitative computational analyses, as presented in the current work, is of outstanding foundational significance to the field.

Apart from the intrinsic challenges arising from limited postmortem sample availability and the impossibility of performing experimental studies, there are significant technical challenges that have impeded progress. Classical studies based on microscopic histological analysis of postmortem brains necessarily had sparse spatial coverage across the brain. Existing neuroanatomical atlases of fetal development lack cellular level resolution (Supplementary information S2). More recent digital histological atlases provide sections through fetal brains with cellular resolution but have large inter-section spacing (Table 1). There are several MRI based developmental atlases (Supplementary information S2). These are three dimensional and permit longitudinal analyses in vivo but do not have cellular resolution. Recent multi-omics efforts are greatly expanding information about the developing brain process at the single cell level, but these have sparse spatial coverage due to comparatively high cost. Consequently, there are no existing data sets that simultaneously have cellular resolution and dense three-dimensional coverage of the developing human brain.

Ongoing studies of transcriptomic cell diversity enabled by widespread adoption of single cell and single nucleus RNA sequencing aim to reveal a comprehensive molecular program of human brain development^1–3^. Advent of spatial transcriptomic datasets from the developing brain^3–5^ will ultimately contextualize transcriptome-based cell types in spatial coordinates of the developing brain, but the throughput of these approaches is currently drastically limited and insufficient to comprehensively survey cell type distributions across entire brain specimens. The theoretically ideal neurodevelopmental data set would require single-cell resolution across entire brains together with histochemical and omics signatures, and dense tracking of the developing neural processes as well as transient scaffolding. We are far from such a goal even in model organisms, thus there is scope for fundamental progress on intermediate milestones especially for the developing human brain where cellular-level data are difficult to obtain.

Here, we present five whole-brain specimens spanning 13-21 gestational weeks that were serially sectioned and processed using a tape-transfer based high throughput histology pipeline and imaged to generate a developmental histological atlas comprising Gigapixel microscopic image data series at 0.5μ in-plane resolution. The high throughput histology pipeline used to generate the data was derived from an original histology pipeline developed for mouse^6,7^ and marmoset^8^ brains and subsequently adapted to larger brains^9^. Utilizing Generative Diffeomorphic Mapping^10^, we assembled five 60μ isotropic multimodal (Nissl, H&E and MRI) data volumes at 13-14 and 20-21 gestational weeks (GW) for the human developing brain, together with volumetric atlas annotations.

We applied machine vision methods to exhaustively segment cell bodies from the Nissl-stained sections to obtain whole-brain cell counts for four brains as well as regional cell counts for two brains based on a series of manually annotated sections. To the best of our knowledge this is the first such detailed set of cell counts in the developing human brain. The quantification demonstrates differential growth and maturation of brain compartments, with brain stem and midbrain structures showing a lower rate of cell division during this time period compared with cortical structures and basal cortical nuclei. Some cell types are distinguishable based on morphological characteristics. We detected and quantified the distribution of Cajal Retzius cells in the outermost layer of cortex based on manually annotated sections, and similarly quantified mitotic figures which indicate dividing cells.

The annotated, multimodal, co-registered data volumes, together with the reference coordinate system which we establish for each age based on skull landmarks, are important outputs of the present work provided as supplemental data sets accompanying the paper. These multimodal volumes provide a reference space and coordinate framework which should be of utility in co-registering multi-omics cell atlases of the developing human brain (https://doi.org/10.5061/dryad.crjdfn3f3).

## Results & Discussion

We utilized tape-transfer based cryosectioning (see methods) to obtain an unprecedented set of histochemically stained 20μ thickness serial sections cut through intact brain specimens (Figure 1). These include two age ranges with a total of 4,228 histological sections, consisting of two brains with ages 20-21 GW (Brain S1: 2,008 sections and Brain S2: 2,209 sections) and three brains with 13-14 GW (Brain S3: 1,153 sections; Brain S4: 997 sections and Brain S5: 1,342 sections). The Nissl, H&E and antibody-stained histological sections imaged using whole-slide light microscopic scanners to obtain the five raw image data volumes presented in this paper (subject metadata are tabulated in supplementary information S1). The tape-transfer based platform for high-throughput serial section histology, established originally to serially section whole mouse brains^6^ and subsequently utilized for processing whole marmoset brains^8^ was adapted to process human fetal brains, with technical advances including methods for cryo-protecting and freezing large brains^9^, and custom-built large format automated stainers and cover slippers. The resulting histological image series outstrip previously available comparable data sets (Table 1) and permitted the assembly of three-dimensional histological atlases using advanced multi-modal, multi-scale registration techniques as detailed below (Figures 2,3).

**Figure 1:**
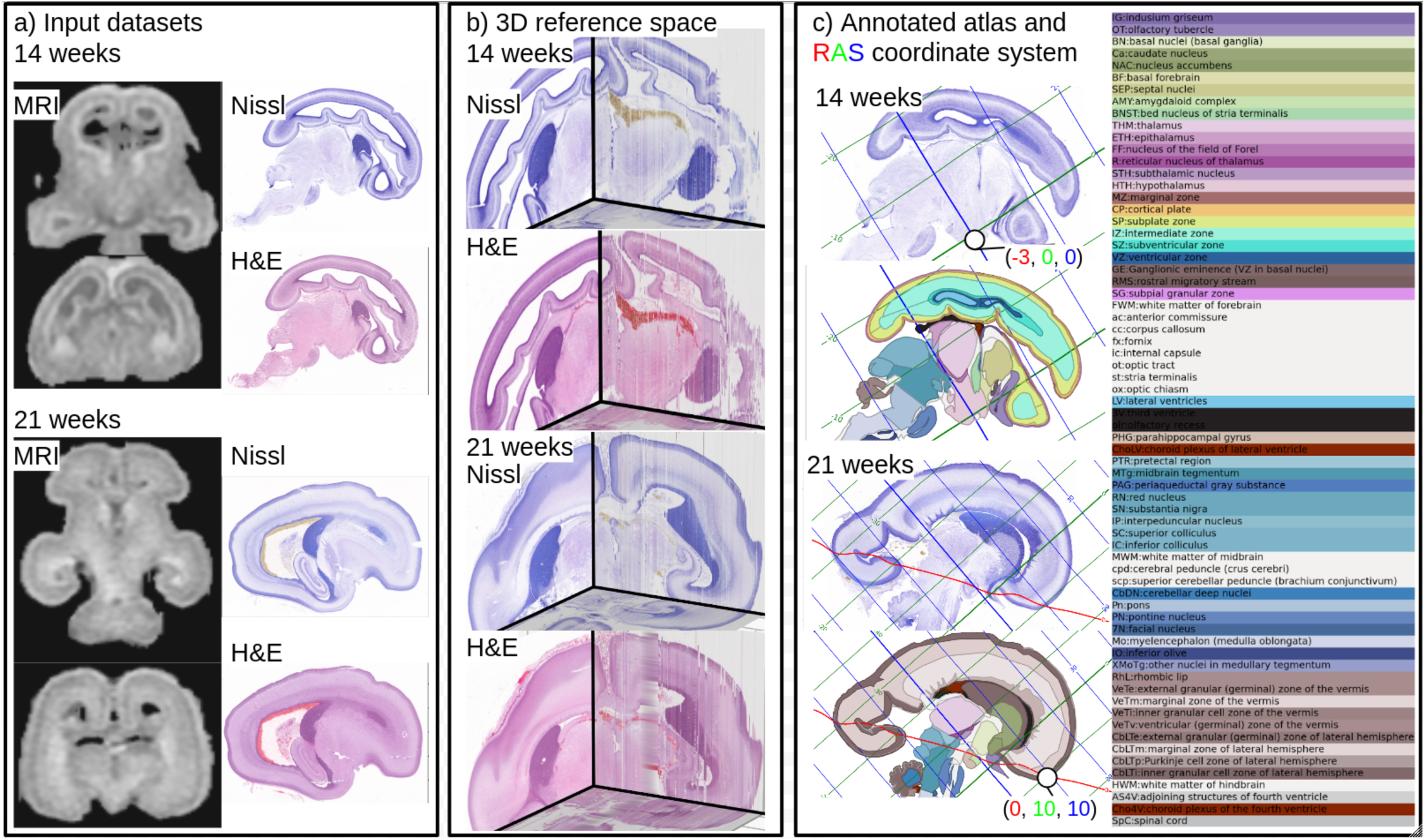
Multimodal reference brain volumes with atlas annotations together with a coordinate system based on skull landmarks (fontanelles) for 14 and 21 gestational week fetal brains constitute a primary output of the paper. The volumes illustrated here are provided as supplementary data sets to this paper. The input data (1a) constitute MRI volumes together with Nissl and H&E stained series (20μ thickness, 60μ spacing, imaged at 0.5μ in-plane resolution), co-registered and reassembled into 3D histological atlases with 60μ isotropic voxels (1b). A subset of sections was manually annotated (1c) to produce a conventional 2D atlas together with coordinate grid lines, and the labels were interpolated to produce 3D annotated volumes, also provided as supplementary data.

**Figure 2:**
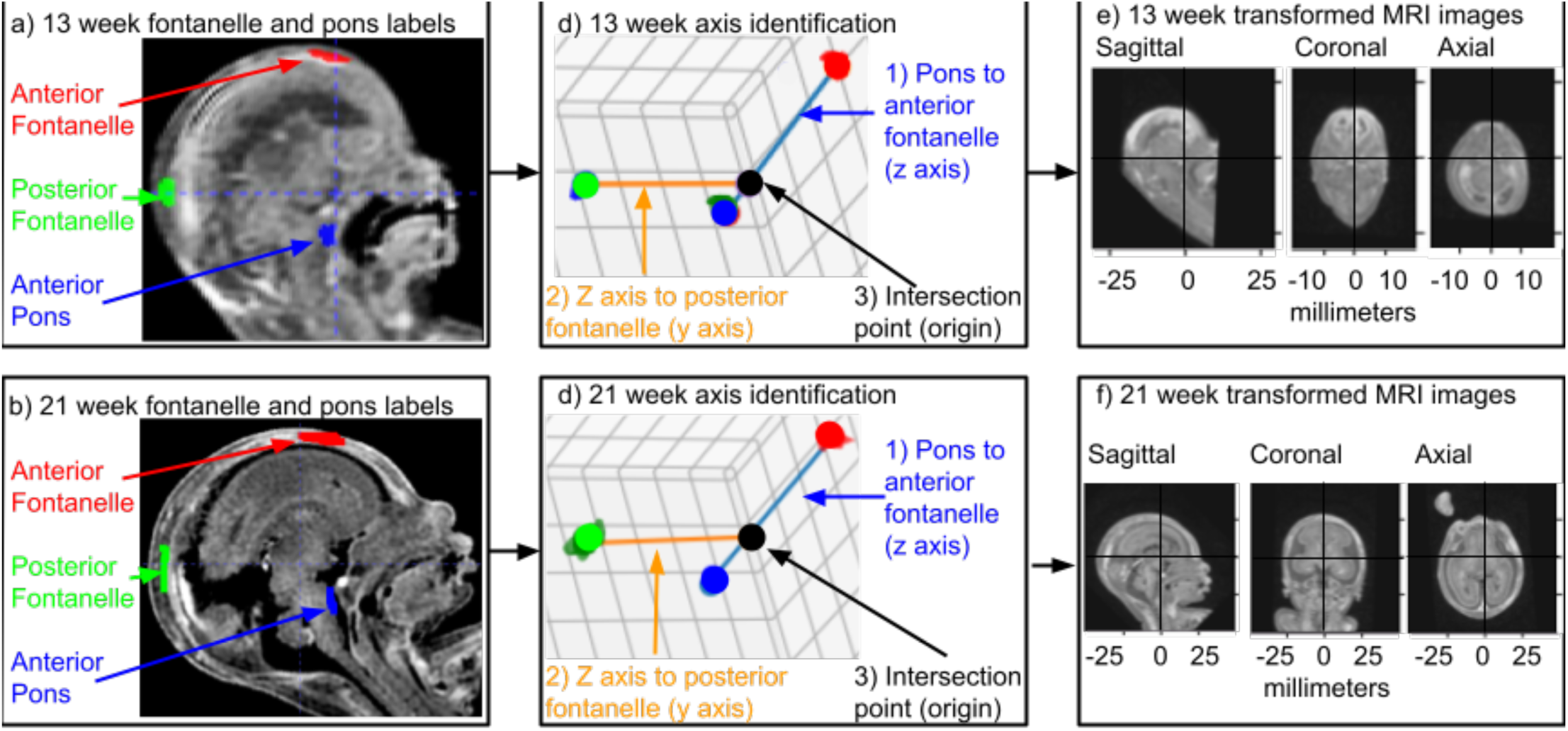
Setup of the reference coordinate system a) Fontanelle and anterior pons are annotated in 3D. b) A line is traced from the anterior pons to the superior fontanelle, identifying the z axis (pointing superior). A second is traced perpendicularly from this line to the posterior fontanelle, identifying the y axis (pointing anterior) and the origin at the intersection of the z and y axes. Finally the x axis is identified as perpendicular to the y and z axes (pointing right).

**Figure 3:**
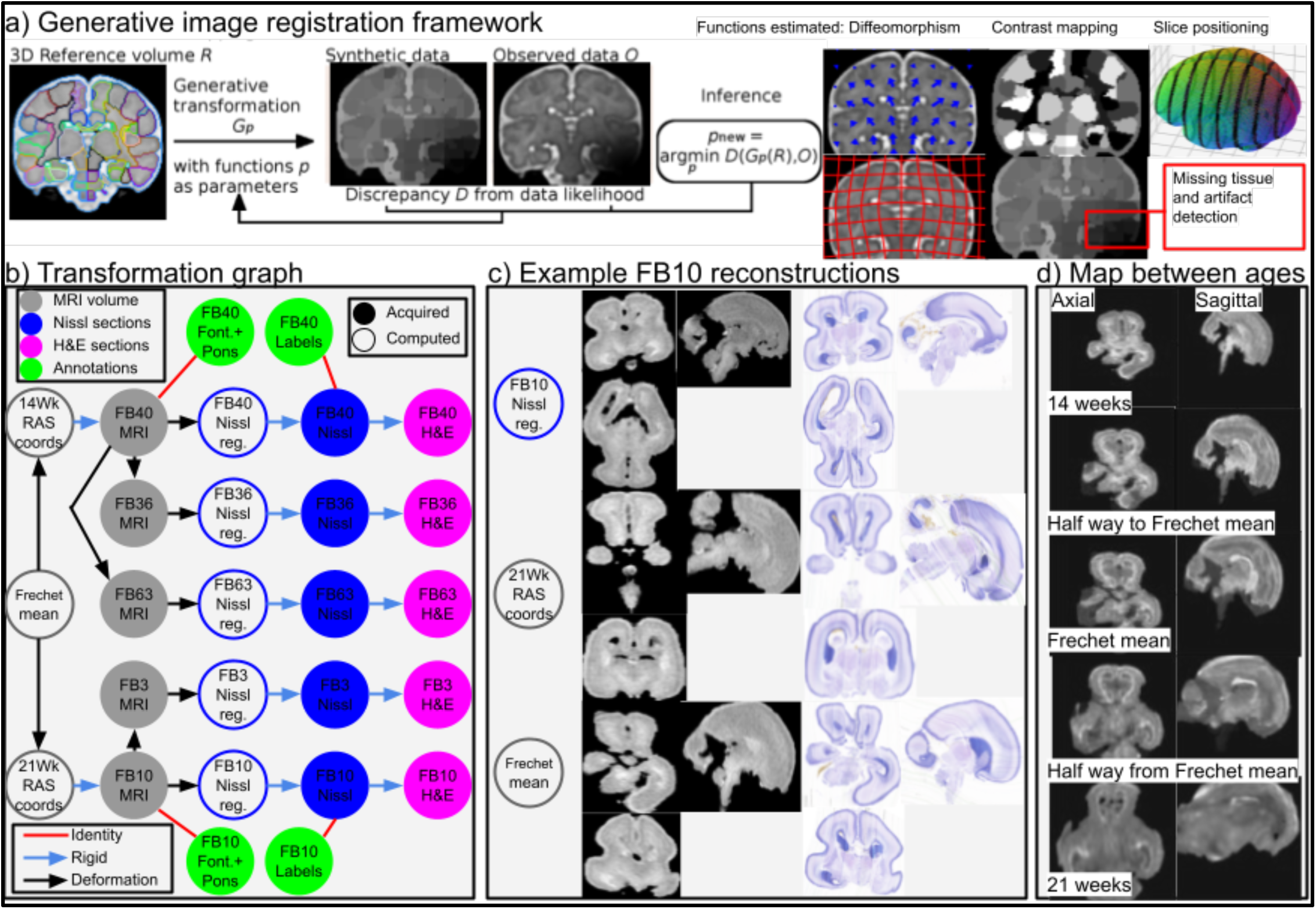
Multimodal registration methodology, morphs from one age to another, graph showing transformation between spaces. a) Our approach to estimating transformations as a generative problem in a maximum a posteriori setting. Data shown in this example is from 2017 Fetal Brain Atlas v3 from the Computational Radiology Lab, Boston Children’s Hospital. b) A directed graph with nodes showing the different spaces where our imaging data is sampled, and edges showing image registrations we compute between these spaces. The left column illustrates three spaces we use to share our 3D volumes. The 3rd column shows reconstructed histology with no interpolation between slices. c) Shows Nissl images and MRI reconstructed in three different spaces from b), to illustrate the differences between them. d) We illustrate our map between 14 week and 21 week MRI datasets.

Three alternating series of 20μ sections were collected for each brain, the first two stained with Thionin (Nissl) and Hematoxylin and Eosin (H&E) and the third utilized for a variety of antibody stains (Figure 4). The serial histological sections of unprecedented completeness permitted us to reassemble the data into 60μ isotropic histological image volumes for Nissl and H&E respectively (Figure 2). These volumes, co-registered to each other, superposed with regional annotations, and the corresponding ex vivo MRI image volumes, all diffeomorphically transformed into a standardized coordinate space for each gestational age (Figure 3), are a main output of the current work and are provided as supplemental data sets together with this paper (https://doi.org/10.5061/dryad.crjdfn3f3). These multimodal atlases and canonical spatial reference spaces (also known as common coordinate frameworks) can in the future be used to integrate and provide histological context to multi-omics data sets being actively collected in several current projects. While the isotropically resampled histology volumes have lower in-plane resolution than the raw in-plane light microscopic data (originally 0.5μ pixels were down sampled by 120x in-plane to generate the isotropic histology atlas volumes) they nevertheless have significantly higher information content than the corresponding MRI image.

**Figure 4.**
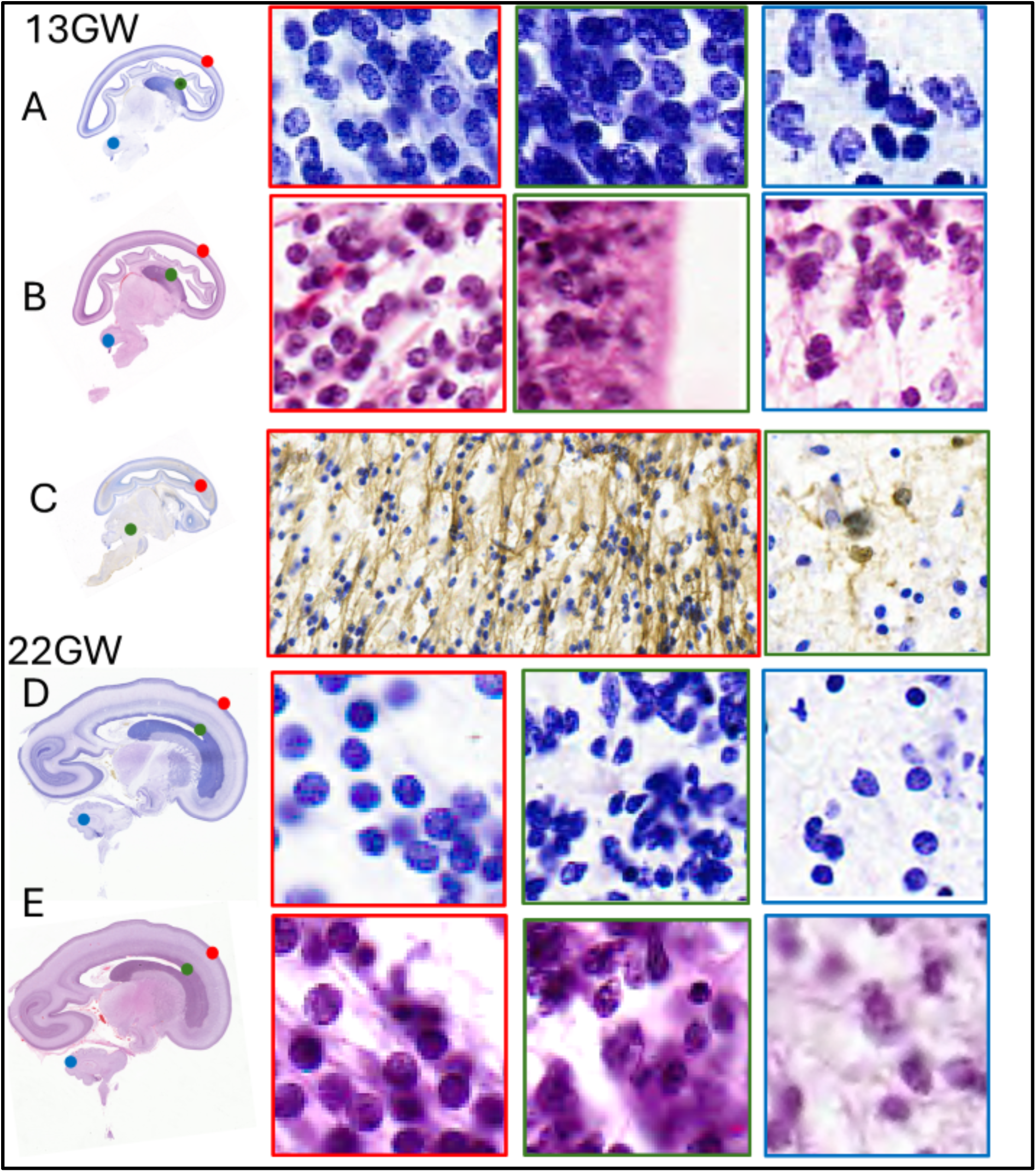
shows examples of cells in the cortical plate of the fetal brain (red boxes, ABDE) for the 13 GW (top two rows) and 22 GW (bottom 2 rows) from a Nissl stained and the adjacent H&E stained sections. The green panels (in ABDE) show examples of the cell in the germinal zone and the blue panels show the examples of cell within the cerebellum. The middle panel C shows an example of a section stained using anti GFAP immunohistochemistry. This panel shows the radial glial fibers in the telencephalon and evidence of astrocytes in the midbrain region

**Table 1.** shows the comparison between the current atlas and the two known reference atlases on fetal brain development.

| Features | Bayer and Altman (2005)<sup>11</sup> | AIBS human developmental atlas<sup>12</sup> | Present atlas |
| --- | --- | --- | --- |
| 2D histological atlas, resolution | 2D print (300dpi)<br>35μ sections | 2D digital (1 μm/pixel)<br>20μ sections | 2D digital (0.5 μm/pixel)<br>20μ sections |
| 3D histological atlas, resolution | None | None | **3D digital volumes**<br>**60μ/voxel** |
| Reference coordinate system | None | None | **Fontanelle based CCF** |
| Number of brains/age, plane of section, total number of sections | 9 whole brains 13.5GW to 24GW (Sagittal, Coronal)<br>88 sections total | 2 Hemibrains 17GW, 23GW<br>1 brainstem 23GW (coronal)<br>168 sections total | 3 whole brains-13-14 GW<br>2 whole brains -21-22GW<br>**~4228 Sagittal sections total** |
| Nissl section spacing | Variable and uneven | 250-1200μ | **60 $\mu$** |
| H&E section spacing | None | None | **60 $\mu$** |
| MRI (ex vivo) | No | No | **Yes** |

### Quantitative assessments of cell numbers in developing human brain

Access to a dense set of histological sections allowed us to make accurate estimates of total cell number across anatomically defined regions of the developing human brain (Table 2), based on automatic detection of cells using machine vision techniques (Figure 5) (see methods). Our sections are sufficiently thin so that the optical imaging plane contains most of the cells in the physical section. Thus, we were able to estimate an appropriate correction factor for cell counts based on a stack of optical sections spanning the physical section. Previous estimates do not span all brain compartments and also utilize statistical sampling approaches^13–17^, whereas we are able to comprehensively detect every cell in serial histological sections in an automated manner (Figure 5; see methods).

**Figure 5.**
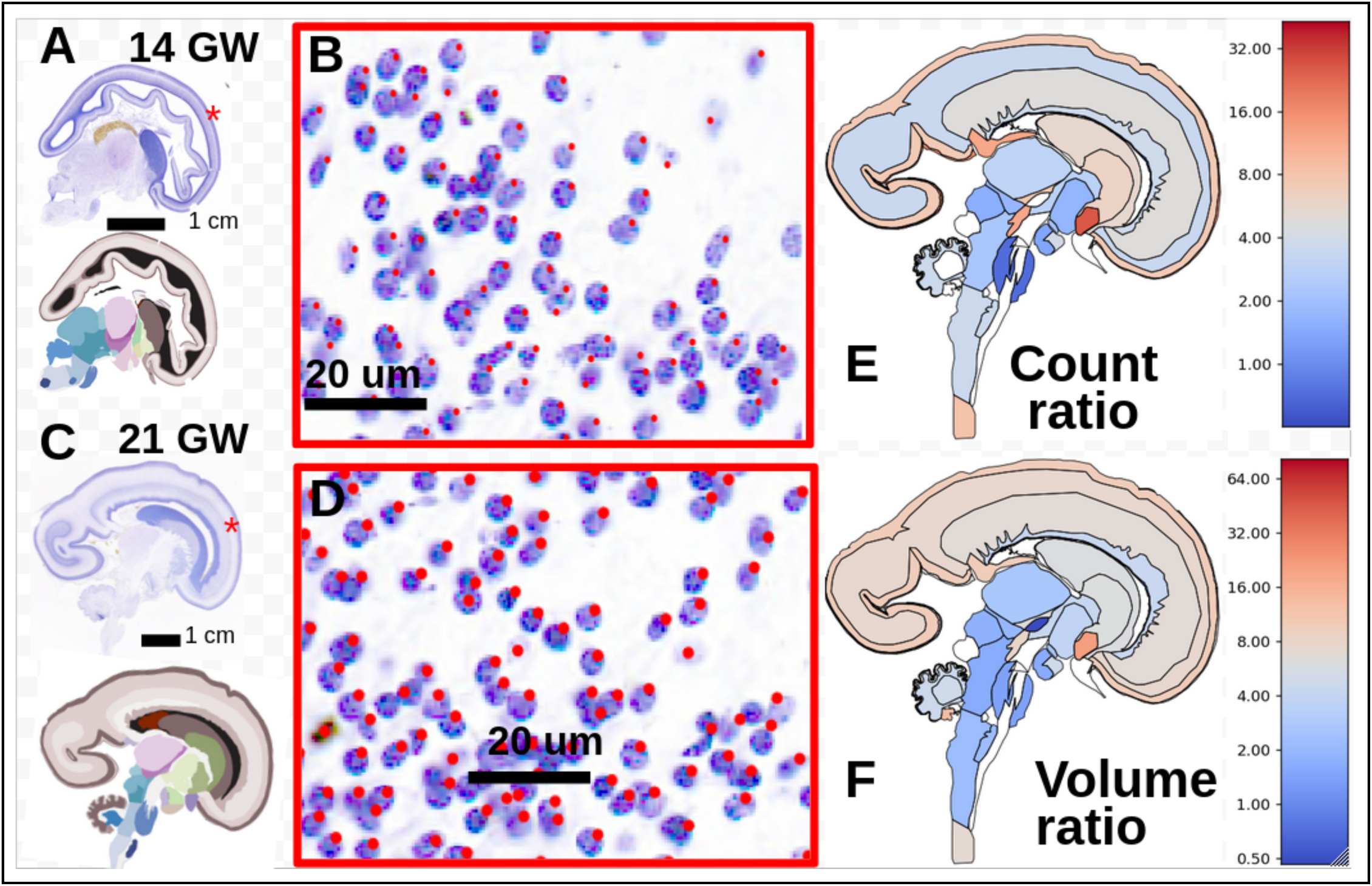
shows the output of the cell detection algorithm for 14 GW brain (A) and 21GW brain (B). Panels B and D show magnified images showing the detected cells for the Nissl-stained sections. E shows a differential regional heat map of the change in cell counts from the 14GW to the 21 GW across the brain. Color is shown on a log scale, centered at the mean across the whole brain. F shows the same as E for structure volumes instead of cell counts.

We quantified both technical and individual variation in whole-brain cell counts across two brains each for the two developmental time points studied. There are multiple potential sources of technical variation, including failure to stain all cells, cells present in the physical section that are not visible in the optical section, and errors in the automated detection of the cells. The first factor is hard to quantify but three of the four brains were well stained, while one brain had lighter stain and lower cell counts. The second component could be estimated by examining a stack of focal planes spanning the physical section. We estimated and applied a stereological correction factor of 1.04 (Supplementary information S3). The third source of technical variation was quantified by manually proofreading several image tiles (see methods). We found that the automated detection performed well, with an average false positive rate of 12% and an average miss rate of 5%. The false positive rate largely arose from single cells that were erroneously split into two. In other words, this means that for every 100 automated detected objects, 88 were true cells, 12 were not cells (arising mostly from splitting single cells), and 5 cells were missed. Thus, our automated counts should reflect the true counts with an error margin of about 5-10%. In summary, the technical variations were estimated to be relatively small, except for light staining which reduced the counts in one of the brains subjected to full counting.

With this estimate of the technical variations (∼5-10%) in mind, it is instructive to look at the total counts for the whole brain, which we estimated to be 2.9x10^9^ (Brain S4; 13 GW 2 days), 3.7 x10^9^ (Brain S5; 14 GW), 10.3 x10^9^ (Brain S1; 20GW 4 days) and 17.4 x10^9^ (Brain S2; 21GW 1 day). Even allowing for the small age differences, the individual variations in cell counts are larger than the technical variations but are nevertheless consistent with each other and with previous estimates in the literature (Supplementary information S4). Notably, adult human brain sizes are estimated to vary up to two-fold^18^ so that this degree of individual variation in the total brain cell counts in prenatal specimens is not unexpected. Since we manually annotated regions for two brains, which also have good stain quality, we have based our regional tabulation and brain cell count estimation to two brains (Brains S5 and S2).

In two well-stained brains which we manually annotated for atlas regions, we found a total of 3.74 x10^9^ and 17.43 x10^9^ cells at 14GW (Brain S5; 7.04cc) and 21GW (Brain S2; 43.88 cc) respectively, a 4.66-fold increase in cell number (6.23-fold increase in volume) over this period. We present a summarized set of regional counts in Table 2 (the full table is provided in supplementary materials). Cell proliferation occurs at different rates across the compartments (Figure 5), with cortical structures growing more rapidly than the subcortical structures, indicating earlier maturation of the subcortical structures compared to the cortical structures. While the total number of cells increase by a factor of 4.7 during this period, the corresponding increases for the thalamus (3.8), hypothalamus (3.3), cerebellum (3.8) and midbrain (2.5) are slower than the average growth rate, while the ratios for the cortical plate (7.3) and marginal zone (15.9) are substantially larger. The number of cells in the ventricular zone shrink during this period by a factor of 0.5.

**Table 2:** Summary counts of select regions in the developing brain (For a comprehensive list see Supplementary information).

| Region | Abbr | Counts<br>GW14 (in<br>millions) | Vol GW14<br>(mm<sup>3</sup>) | Counts<br>GW21(in<br>millions) | Vol<br>GW21<br>(mm<sup>3</sup>) | Count<br>ratio | Vol<br>ratio |
| --- | --- | --- | --- | --- | --- | --- | --- |
| **Developing Neocortex** | NCx | 2317.2 | 4055.9 | 12061.0 | 31248.0 | 5.20 | 7.70 |
| Marginal zone | MZ | 16.6 | 181.5 | 265.0 | 1187.1 | 15.93 | 6.54 |
| Cortical plate | CP | 728.3 | 811.7 | 5312.7 | 7912.5 | 7.30 | 9.75 |
| Subplate zone | SP | 772.8 | 1641.8 | 2654.2 | 12531.4 | 3.43 | 7.63 |
| Intermediate zone | IZ | 799.5 | 1420.9 | 3829.1 | 9617.0 | 4.79 | 6.77 |
| **Germinal zones** | GZ | 810.9 | 721.6 | 2807.3 | 2810.8 | 3.46 | 3.89 |
| Subventricular zone | SZ | 252.4 | 257.1 | 953.7 | 1103.5 | 3.78 | 4.29 |
| Ventricular zone | VZ | 246.0 | 197.4 | 123.8 | 120.9 | 0.50 | 0.61 |
| Ganglionic Eminence | GE | 312.5 | 267.1 | 1729.8 | 1586.4 | 5.54 | 5.94 |
| **Basal Ganglia** | BG | 137.9 | 348.4 | 665.9 | 1590.8 | 4.83 | 4.57 |
| **Allocortex** | ACx | 82.5 | 233.5 | 409.4 | 1243.4 | 4.96 | 5.33 |
| **Thalamus** | THM | 69.9 | 321.0 | 267.0 | 1129.1 | 3.82 | 3.52 |
| **Hypothalamus** | HTM | 18.7 | 77.8 | 62.3 | 240.2 | 3.34 | 3.09 |
| **Cerebellum** | CBX | 73.5 | 206.7 | 276.1 | 1137.6 | 3.76 | 5.50 |
| **Midbrain** | M | 27.1 | 147.8 | 69.7 | 265.4 | 2.58 | 1.80 |
| **Pons** | Pn | 29.8 | 104.5 | 114.1 | 494.9 | 3.83 | 4.74 |
| **Medulla** | Mo | 18.9 | 121.2 | 95.1 | 402.0 | 5.02 | 3.32 |
| **White matter** | wm | 78.7 | 396.9 | 360.1 | 2379.5 | 4.58 | 5.99 |
| **WHOLE BRAIN** |  | 3,733.6 | 6998.8 | 17,365.5 | 43,712.2 | 4.65 | 6.24 |

### Reference coordinate framework and multimodal 3D atlas

Histological atlases for laboratory animals have canonically been presented in coordinate systems defined with reference to landmarks on the skull that provide fiduciary reference points for stereotactic surgery. Following upon previous practices using ultrasonography in fetal brains, we utilized skull landmarks based on apertures in the developing skull, also referred to as fontanelles, as well as the pons, to define analogous stereotactic coordinate systems (Figure 1,2) (see methods). The establishment of a reference coordinate system is of primary importance in our approach, in which we give primacy to reference brains along with associated reference coordinate systems that can provide a scaffold to integrate other data sets and provide an unambiguous methodology for cross-referencing other data sets.

Each imaging dataset was collected in its own coordinate space, and transformations between pairs of spaces were computed until the dataset was spanned by a directed tree (a “transformation graph” ^19^, similar to an “assembly graph” used to reconstruct genomes^20^). Our transformation graph is shown in Figure 3, revealing how imaging and derived data from any space (nodes) can be reconstructed in another, by composing transformations or their inverses (edges). Our image alignment method for computing these transformations builds upon previous work designed to be robust to contrast differences, missing tissue, or other artifacts^19,21^, and are built within the Large Deformation Diffeomorphic Metric Mapping framework^22^ to ensure transformations are smooth and invertible. Our approach seeks to generate a synthetic “target” dataset, using a sequence of shape and intensity/color transforms applied to an “atlas”. Optimal transformations are found by minimizing a measure of discrepancy (here corresponding to maximum a posteriori estimation). The approach is applied in 5 different settings: (i) maps from 3D MRI to 2D Nissl series, (ii) from 2D Nissl series to 2D H&E series, (iii) between two 3D MRI volumes for individuals of the same age, (iv) between two 3D MRI volumes for individuals of different ages, estimating an “average template” halfway between the two (in the sense of a Frechet mean^23^) (v) between annotated sections and their neighbors to interpolate into unannotated sections, as in^24^.

### Atlas plates and nomenclature

Classical neuroanatomical atlases rely on two dimensional images together with compartments defined and named by expert neuroanatomists. The expectation is that such expert judgment will help mitigate the challenges associated with mapping samples drawn from different individual brains through the identification of cytoarchitectonically analogous structures from different brains. However, expert neuroanatomists disagree on how to name and group the different cytoarchitectonic compartments that may be visible in histological sections, leading to conflicting nomenclatures and atlases (Supplementary information, Nomenclature for developing brain compartments). Making atlases for developing brains have the additional challenge of dealing with transient and changing structures and reconciling nomenclatures across developmental stages. Adult compartment nomenclatures, often based on putative function of a brain compartment, may not be appropriate for fetal brains where the underlying evidentiary basis for functional annotations may be missing. In contrast, designating a location in a developing brain using a reference coordinate system provides an unambiguous and neutral reference. A new brain at the same developmental age may be mapped into the reference brain, using the same cross-data set mapping methods described above, thus providing the ability to refer unambiguously to a point on the new brain as well.

The coordinate-system based approach does not preclude traditional atlas annotations but allows multiple such annotations to be applied to the same reference brain, being agnostic to such annotation overlays. In deference to the classical approach, we provide a set of atlas annotations in the present work, in the form of conventional two-dimensional atlas plates (Figure 4; Supplementary information – GW14 and GW21 atlas) together with coordinate axes superposed. Because we include a nonlinear mapping between MRI where we define our right-anterior-superior (RAS) coordinate system, and histology data, this coordinate system appears curved when viewed superimposed on a histology image. In particular, the coordinate grid we include (Fig 1c) corresponds to an isocontour of the nonlinear mapping between registered histology space (Fig. 3b, blue open circles) and the MRI RAS space (Fig 3b, black open circles). Red curves are isocontours of the “R” component, green curves are isocontours of the “A” component, and blue curves are isocontours of the “S” component. We also provide a set of three-dimensional volumetric annotations corresponding to our multimodal reference brains for each gestational age (Fig 1c; Supplemental data).

### Quantification of mitotic figures and Cajal-Retzius cells

Histologically stained sections contain information about cellular morphology (Figure 4) which is used by anatomic pathologists to identify morphological cell types in the clinic. We took advantage of this image-based information, to detect and quantify two types of cells, including Cajal-Retzius cells and mitotic figures (characteristic objects visible in H&E-stained images that indicate cells undergoing mitosis, used *e.g.* by anatomic pathologists to assess cell division in tumors). We manually detected mitotic figures and CR cells in annotated sections to obtain an unprecedented spatial atlas of actively dividing cells in the developing brain (Fig 6 and supplementary information).

**Figure 6:**
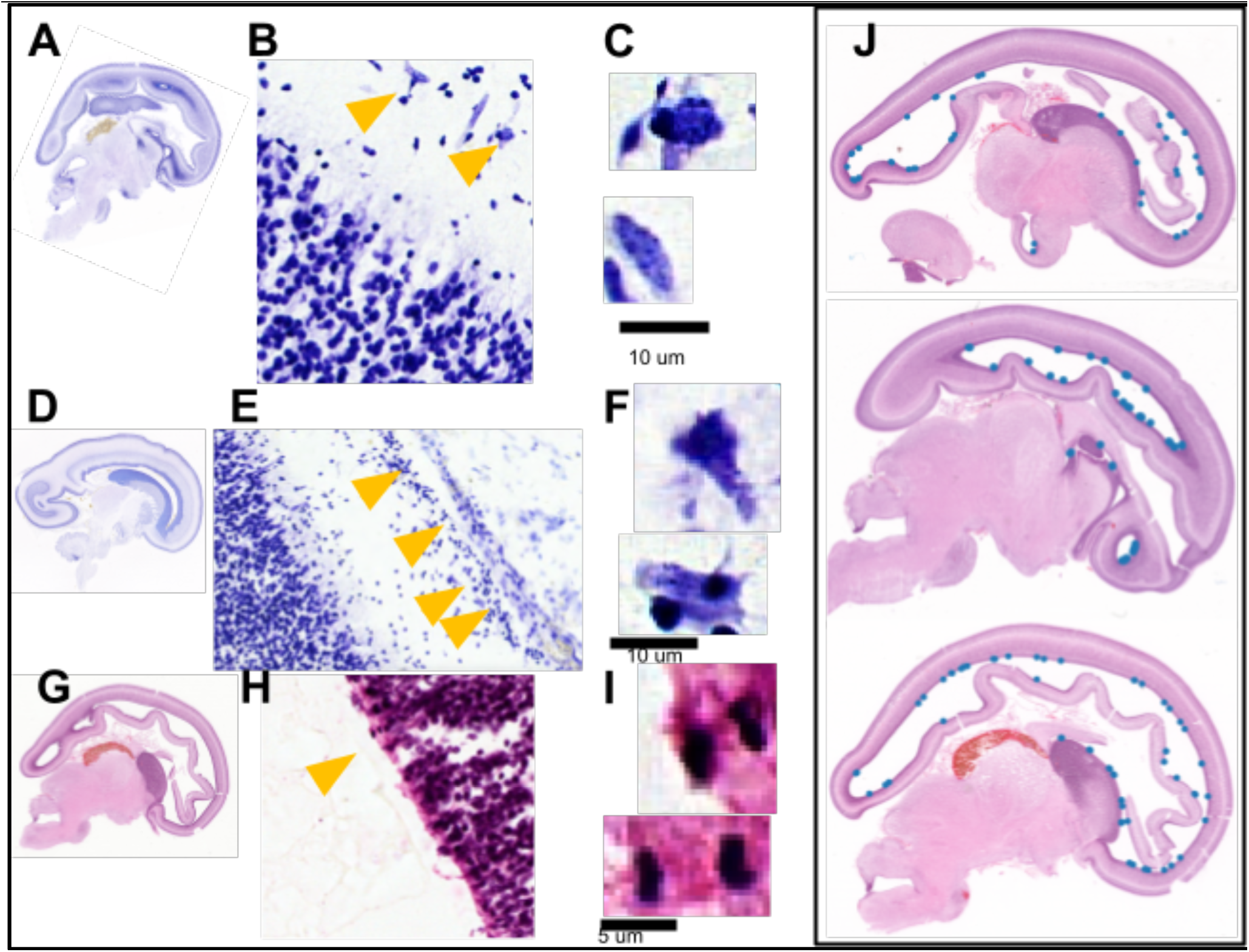
Distribution of Cajal-Retzius cells and Mitotic figures. Panels A and D show the locations of the insets (B,E) which contain the Cajal Retzius cells (yellow arrows), Panels C and F show zoomed in regions with CR cells. Panel G shows the location of the mitotic figures shown in H. I shows the examples of the mitotic figures seen in the ventricular zone of the cortex. J shows the location (blue dots) of mitotic figures on three other H&E slices.

We estimated whole brain counts of CR cells in the two brains with manually annotated sections (GW 14: 2.8 x 10^5, GW21: 8 x 10^5) from counts on the annotated sections, using the spline interpolation method (see methods). Uncertainty in these counts are difficult to estimate due to partial sampling of the subpial granular zone in each section, however we estimated an uncertainty of ∼30% in the GW14 counts and ∼9% in the GW21 counts (see methods). According to these counts, the number of CR cells increases from GW14 to GW21 by a factor of 2.9. By estimating the total surface areas, we found that the number of CR cells per unit area decreased somewhat (GW14: 123.4 cells/mm^2, GW21: 74.7 cells/mm^2). These estimates were consistent with visual examination of the linear spacing between CR cells. These findings are surprising given the well-established attrition of these cells in mice^25^. However, it has been proposed that during human development, transient CR cells are replaced by persisting CR cells by midgestation^26^, offering a possible explanation for the increased number of the overall number of CR cells between GW14 and GW21 specimens.

To obtain these whole brain estimates for the CR cell counts and density we pooled data across sections. The subpial granular zone is the external-most layer of tissue in our brain samples and is very thin. Consequently, it was only partially present in the sections. Consequently, we were not able to quantitatively assess variations in CR cell density across the cortical surface. We anecdotally noted some variation, such as a higher density in the insular region, but these observations of regional variations require follow up studies.

We also estimated a whole brain count of mitotic figures representing dividing cells in the two brains with manually annotated sections, using spline interpolation of total counts across sections (GW14: 64.5 x 10^3 thousand, GW21: 9.0 x 10^3). The estimated number of mitotic figures drops sharply across these ages^27^, also consistent with visual observations of many fewer mitotic figures per section, with the median number of such objects observed per annotated section dropping from 44 to 4.5.

Due to methodological limitations, these estimates should be taken as lower bounds to the true counts. Confirmatory staining for proliferation by the KI-67 antibody indicated that the mitotic figures manually detected in the H&E-stained sections are underestimated, by a factor of about two. We mostly observed the mitotic figures along the ventricular boundaries, although some detects were in the subventricular zone. Further, the mitotic figures can only be reliably detected during the metaphase and anaphase of the mitotic process, leading to a further underestimation of the number of proliferating cells. Nevertheless, these sources of systematic bias apply to both ages, and we expect that the ratio between the two numbers is more reliably estimated, leading to a qualitative conclusion of an order of magnitude reduction in the number of proliferating cells across the age range.

## Conclusions

We have provided the first detailed whole-brain regional counts for human fetal brains at two timepoints in the second trimester. The technical variations in these counts were substantially smaller than individual differences at a given age, as estimated by doing whole brain counts in additional brains. In turn the individual variations at a given age are smaller than the count increases observed across the age range. Thus, the counts paint a biologically meaningful picture of differential growth in cell numbers across brain compartments during the GW 14-21 period. Cell counts in the subcortical regions have not been previously available, and show a significantly slower growth, consistent with an earlier maturation of the subcortical structures.

Morphologies of cells visible in the Nissl-stained sections allowed us to segment and quantify Cajal-Retzius cells. We estimated that the total number of CR cells grew by a factor of about three during the GW 14-21 period, but that the number of CR cells per unit area of the cortical surface decreased modestly. Similarly, we quantified mitotic figures, characteristic of dividing cells, in the H&E-stained sections, and found an approximately one order of magnitude reduction in the total number of mitotic figures from GW 14 to GW 21 indicative of a substantial decrease in the number of dividing cells during that period.

Together with this paper, we are providing several reference brain atlas data sets to the research community. Our data products include conventional histological atlases with regional annotations, as well as isotropic atlas volumes with 60μ isotropic voxels for GW14 and GW21 brains. We established a new reference coordinate space based on skull landmarks and present the histological atlases in this reference coordinate system. The three-dimensional histological atlases in standardized coordinate spaces are unprecedented and illuminate the cytoarchitecture of the developing fetal brain in the second trimester at a level of detail that is not present in previous MRI based studies. The three-dimensional histological atlases together with the associated reference coordinate systems should provide a basis for integrating the currently growing corpora for multi-omics cell typing and spatial transcriptomics.

Our results demonstrate the utility of tape-transfer based dense serial section histology together with computational analysis in the study of developing human brains. Given the significantly lower costs associated with classical histochemical stains compared with antibodies or omics processing, we suggest that this provides a cost-effective but currently under-utilized approach to study human brains. Overall, our findings, as well as the data volumes we provide with this paper, qualitatively and quantitatively advance the state of knowledge about fetal brain development.

## Material & Methods

### IRB Approvals

Postmortem fetal specimens (n=5, GW 13-22 weeks) were obtained from the Departments of Pathology, Mediscan Systems Pvt. Ltd Chennai, India and Saveetha Medical College and Hospital (SMCH) Chennai, India, under the protocol approved by both the Institutes’ Ethics Committee (IEC) and histological processing, was performed according to the approved IEC guidelines, IEC/2021-01/MS/06, of the Indian Institute of Technology, Madras (IITM) at Sudha Gopalakrishnan Brain Centre.

### Sample acquisition

The de-identified whole fetal specimens were collected, after due consent from the next of kin, in accordance with the Declaration of Helsinki, along with the relevant medical history, and investigative records including prenatal ultrasound scans. Specimens were collected in a fixative solution within a few minutes of death, thus keeping the post-mortem interval (PMI) small. Each specimen was injected with 20% formalin into the cranial vault through one of the fontanelles. and after two weeks of whole body immersion fixation, the brain was extracted and immersed in 4% Paraformaldehyde (PFA) for a minimum of two weeks. Prior to extraction, a post-mortem in-skull MRI (1.5T or 3 T) was obtained for all the specimens in this study. Brain specimens obtained in this study were classified as non-pathological based on available medical data including prenatal ultrasonography (USG), postmortem MRI and the routine autopsy examination using standard neuropathologic criteria. In addition, specimens that had significant fixation and extraction damage were excluded from the study. The extracted brain was recorded for specimen weight and dimensions across the dorsal-ventral, anterior-posterior, and right to left lateral axes. The gestational age reported here was calculated as the time from the last menstrual period (LMP) and the prenatal USG assessment.

### Magnetic Resonance Imaging (MRI) Acquisition

Postmortem structural MRI of the fetal brain was performed using T1-weighted and T2-weighted scans in either 3-T GE Medicals, United States, (version DV28.0_R05_2.34.a, Signa Architect) with flex coil (12 mm inner diameter) or a 1.5-T Philips Medical Systems, India, (version 5.3.1/5.3.1.4, with the wrist coil (10mm inner diameter). Custom-designed MRI-compatible containers were designed to place and wrap the finger coil around the specimen for scanning. GW14-15 specimens were placed in the oval-shaped container, while the larger specimens, GW20-22 were placed in a custom-designed rectangular container. In the 3T scanner, T1 MP-RAGE (TE = 2-3 ms, TR = 2400-2600 ms, slice thickness = 1.2 mm, flip angle = 8 deg, overlap = 0.6 mm, and no interleaved space between the slices) and T2W FSE (TE = 125 ms, TR = 3000 ms, slice thickness = 1 mm, overlap = 0.5 mm, flip angle = 90 aͦnd no interleaved space between the slices) in orthogonal planes was utilized for the imaging and analysis. In a 1.5T scanner, T1W 3D TFE (TE = 6 ms, TR = 13 ms, slice thickness = 1.2 mm, overlap = 0.6 mm, and no interleaved space between the slices) and T2W TSE (TE = 150 ms, TR = 3500-4000 ms, slice thickness = 3 mm, no overlap and no interleaved space between the slices) in orthogonal planes for imaging and analysis.

### Cryoprotection and freezing

After extraction and a minimum of two weeks immersed in fixative, the brain specimens were prepared for sectioning by cryoprotecting with increasing gradients of sucrose (10-30% in 0.01M PB), where the first solution was 10% sucrose and 4% PFA in 0.01M PB followed by, 20% and 30% sucrose in 0.01M PB at 4 deg C. The volume of the solution used was approximately five times that of the specimen with the endpoint of each cryoprotection step marked by equilibration with the solution, i.e., when the specimen was completely sunk. The total time taken for the gradient sucrose cryoprotection was 18-20 days and 10-12 days for specimens in GW20-22 and GW14-15 respectively.

Whole brain specimens were embedded in OCT and frozen in an isopentane dry ice bath (-80C) using the custom-designed freezing mold for cryosectioning. Our 3D-printed customized freezing molds were designed using the 3D surface scan (Einscan Pro) of the extracted brain and the postmortem in skull MRI. These molds along with copper cubicles allowed us to prepare a cryoblock of each of the specimens in alignment with the physical resting state of the brain as determined by the postmortem MRI. The temperatures during the freezing process were monitored using thermocouples to ensure that the temperature was maintained between -80°C to -78°C throughout the freezing duration which ranged from 12-16 minutes for the samples with the volumes ranging from 30 to 80cc. Post this rapid freezing, the frozen cryoblock was stored in a -80°C freezer for at least 24 hours to ensure uniform freezing throughout the brain prior to cryosectioning.

### Cryosectioning

Cryosectioning of the frozen blocks was performed in the large cryomacrotome (Leica CM3600, Leica Biosystems, India) for the GW20-22 and the cryostat (CM1520, Leica Biosystems India) for GW14-15 using the modified Tape Transfer Technique (pioneered by ^6^ but adapted for large slides including 75 x 50 mm and 200 x 150 mm slides). Prior to cryosectioning, the block was acclimatized at -20°C and 40% humidity inside the cryostat chamber for a minimum period of 30 mins. Serial sections were collected on either 75 x 50 mm or 200 x 150 mm slides coated 36hrs in advance with a solution comprised of, in the ratio of 20mL -Trimethoxysilyl propyl methacrylate, 5 mL 0.1M acetic acid, 100 mL acetone, followed by a UV curable Optical adhesive (Norland NOA 63, Edmund Optics) spread uniformly on the glass slides. The total volume required to coat one 75 x 50 mm double slide was 40-50 µl. The Tape Transfer technique developed by^6^ was modified to accommodate the larger tissue. The technique uses adhesive tape (Custom Converting Inc, USA), which is rolled on the frozen tissue block and the section is cut along with the tape adhered to it. The tape with the section adhered to it is placed onto a polymer-coated glass slide under ultraviolet (UV) illumination (360nm, 8 x 5W LEDs rendering 1600uW/cm^2, for a maximum time of 8s). Application of UV illumination (for 8s) results in the uncuring of the glue in the tape and the curing of the glue in the glass slide resulting in a smooth hands-free transfer of the sections onto the glass slide, each specimen was sectioned in the sagittal plane as 20 µm thick sections, which were then separated into three series for staining-Nissl, Hematoxylin, and Eosin (H&E) and Immunohistochemistry (IHC).

Along with serial sectioning, custom-designed Blockface Imaging (BFI) using white light and UVC illumination (254nm) was collected^28^. In addition to the BFI, Fluorescent Fiducial markers were placed on the block chuck and were used as a reference for alignment during the registration and reconstruction.

Due to the high-water content present in the developing brains, Nissl and H&E series were dried in a 33°C oven for 24 hours before staining and coverslipping. The IHC slides were stored in 4C until further processing. Our serial sectioning resulted in ∼2000 sections for the GW20-22 and ∼1400 sections for the GW13-14.

### Staining and coverslipping

The mounted sections for Nissl and H&E were dried overnight at 33°C in the dry oven and then stained for Nissl substance using 0.2%Thionine (SRL Chemicals, India) and H&E (Sigma Alrdrich, India) using the Harris’ regressive protocol. The staining for Nissl and H&E was performed using a custom-designed autostainer specifically designed to accommodate the 75 x 50 mm and 200 x 150 mm glass slides. After staining, the slides were once again dried in the 33°C oven for 3 days as the fetal brain has a very high water content. For the Thionin Nissl stain, the slides were stained for Nissl substance (pH:4.0) for 60 seconds, differentiated dehydrated in increasing concentration from 50, 70, 95, and 100% Ethanol and finally cleared in Xylene with two rinses. For H&E staining, we employed the Harris regressive protocol where the slide was stained for Hematoxylin (pH: 2.7), differentiated in distilled water, first in cold water followed by lukewarm in 1% acid alcohol, and then stained with Eosin followed by dehydration in graded ethanol (95% and 100%) and cleared in Xylene, Following drying, the slides were coverslipped with DPX mounting media (Merck) using custom designed automated coverslipper.

### Histological Image Series Acquisition

All stained slides post-drying were scanned using TissueScope LE120 (Huron Digital Pathology, Canada) scanner designed for large format slides at 0.5µm/pixel, in-plane resolution. The scanned images were transferred into the Laboratory Inventory Management System (LIMS) where it underwent Quality Checks (QC). The images were scrutinized based on software issues (stitching, white balance, or focus issues), staining and coverslipping issues (presence of air and water droplets), and the extent of tissue damage (<20% or not). The sections that passed through the quality control checks were included for further analysis.

### Automated cell detection and classification

The nissl-stained sections in the fetal brain contain millions of labeled cells. The morphology of these cells are fairly similar in shape and size. The cells are closer to being circular in shape in the fetal brain before developing into the more diverse somatodendritic morphologies characteristic of the adult brain. We found that classical machine vision techniques provided better performance for the detection of the cells in the Nissl stained sections compared to deep learning based approaches. The deep net based approaches we tested, including semantic segmentation using Mask-RCNN had significantly poorer performance due to a variety of factors, including overlapping cells present in the sections and distributional shifts across sections due to variable staining. The classical MV techniques required no training, were fast to evaluate, and relatively straightforward parameter adjustments allowed for high performance as verified by manual proofreading.

The algorithm for automated cell detection from the Nissl stained sections and the corresponding code together with example data can be found on github (https://github.com/mitragithub/NISSL-Cell-Detection.git). Features of connected components detected in a first pass based on thresholding, specifically the area covered by the component, were used to determine whether the component consisted of a single cell or multiple cells. For the full resolution Nissl images we empirically determined that components with an area between 40 and 75 image pixels were typically single cells, while the larger components with an area greater than 75 pixels were further processed as multiple cells overlapping or touching in the component. We empirically determined the mean area of isolated cells in the image to be 100 pixels. To determine the number of components in an area that crossed our threshold of 75 pixels, we estimated the maximum number of single cells that could fit within the area using a mixture of overlapping Gaussians. Given the estimated number of cells n in a given component, we fit a Gaussian Mixture Model with tied covariance with n components, allowing the Gaussians to be overlapped. The means of the fitted Gaussians were taken to be the centres of cells present in the multi-cell components. The final result includes the union of the centroids of the single cell components and the Gaussian centroids from the multi-cell components. We found this approach to be simple and efficient in determining the cells with partially visible overlapping areas. This approach is highly parallelizable and can be used as a tiled approach (moving window) to compensate for any intensity variation in the image.

To determine performance we manually segmented a set of 8 randomly chosen tiles (1000 x1000 pixels) to obtain ground truth. average false positive rate of 12% (defined as the number of non-cells divided by the number of detected cells) and an average miss rate of 5% (defined as the number of cells not detected, divided by the number of detected cells). The false positives largely arose from single cells that were erroneously split into two or occasionally more parts. Relatively few cells were missed by the detector, reflected in the 5% miss rate.

### Regional cell counts

A subset of Nissl stained sections were manually annotated to delineate the atlas compartment in two brains (median spacing 0.66mm in the GW21 brain and median spacing 0.54mm in the GW14 brain). These sections are the basis of the 2D atlas plates associated with the manuscript. The cell detection algorithm was applied to these sections, and cells counted within each marked region, to obtain a raw (region x section number) data matrix. Stereological considerations based on scanning z-stacks on individual sections gave a correction factor of 1.04, which was applied to the regional counts. Each row of that matrix was then interpolated using the Piecewise Cubic Hermite Interpolating Polynomial method (using the pchip function implementation in MATLAB version R2003a) to a uniform 20 micron section spacing. Each row of the matrix was then summed to obtain the regional counts. A similar process was applied to the areas of the regions in the annotated sections to obtain the total volume of each region. To check for the effect of sub-sampling of sections, we also applied the interpolation method to total cell counts in all Nissl sections without annotations. Lack of annotations meant that cell detects in the ventricular regions were also included in the total counts. The difference between the interpolated total counts based on only the annotated sections, and on all sections, were 1.6% and 4.6% for the GW13 and GW21 brains respectively. Separately, we estimated (see above) that the automated detection algorithm gave counts within 5-10% of the true counts, so that the interpolation error was small compared to the errors due to the automated detection algorithm.

In addition to these estimates of the technical variability, there is also the issue of individual variability. We estimated the individual variations by detecting cells in the Nissl stained sections of one age matched brain each for gestational week. The resulting counts were 2.9B cells for the 13 GW brain and 10.3 cells for the 21 GW brains. We thus estimate the individual variations in total counts to be about 30% and 40% respectively for the 13 GW and 21GW brains. We recognize the limitations of estimating individual variations in cell counts from two brains for each age group, however, this is the first time such exhaustive counting has been undertaken for any fetal human brain and this is a mitigating factor. Our methodology can be generalized to other data sets in the future to obtain a better understanding of individual variability in cell counts for a given gestational age during development.

### Mitotic figure detection, KI staining based validation and quantification

Using the HE stained histology images, we identified the signatures of classical mitosis as defined by^29,30^ in the developing mouse. Examples of the identified mitotic figures are shown in figure 6L. These figures mostly contribute to the metaphase and anaphase of the classical mitotic process. We used a proliferative IHC marker (Ki-67) to validate our observations in the HE stained images as shown in Supplementary 5 We segmented mitotic figures manually in the high resolution digitized H&E stained images using a custom designed web based annotation tool. An atlas of the locations of these mitotic figures is provided as supplementary information. We further estimated whole-brain counts of the mitotic figures using the same spline-based interpolation method as described above for regional cell counts.

### CR cells detection and quantification

The Cajal Retzius cells are a special class of cells found in the surface of the cortex known to be responsible for cortical development^31,32^. We identified the Cajal Retzius cells based on the histological criteria originally proposed by Retzius^33^ for the developing brain in our Nissl stained sections. These cells were identified as cells that are larger than their surrounding cells and having the classic “ carrot-like” morphology. We segmented these cells manually in the digitized images using a custom designed web based annotated tool and estimated the densities of these cells as follows. We quantified total cell counts by estimating the number of cells on the annotated subpial granular zone, which was not present everywhere as it is easily lost during tissue preparation. We estimated the total length of cortex on annotated slices using the cortical plate annotations. From this we estimated the number of cells on each annotated section (count times cortical plate length divided by subpial granular zone length). We converted to counts per unit area (rather than per unit length) by dividing by slice thickness (20 microns), and used spline interpolation to estimate the counts on non-annotated slices. Total counts were estimated by integrating across all slices.

Since the subpial granular zone (SGZ) is only partially preserved in the sections, the uncertainty in the estimation of the CR cell number is expected to be larger than the regional count estimates, but this uncertainty is somewhat difficult to quantify as the un-observed regions of the SGZ may have had different densities of CR cells. Nevertheless, the areal densities ultimately obtained were consistent with visual examination of the linear spacing between the CR cells in the preserved regions of the SGZ.

### Atlas coordinate system

For each fetal brain a neuroanatomist (JJ) manually annotated the anterior fontanelle, the anterior aspect of the pons, and the posterior fontanelle. A robust center of mass was computed for each fontanelle to reduce the effect of any outlier segmented voxels, by locating the point whose sum of absolute value of distances to all voxels was minimum (note the “mean” is the point whose sum of squares of distances to all other voxels is minimum). The procedure is described in Figure 2. The anterior aspect of the pons was identified as the voxel whose distance to the posterior fontanel center of mass was maximized. A line was extended from this aspect to the center of mass of the anterior fontanelle. This line describes our vertical axis (z axis, pointing superior). A second line was extended perpendicular to the first, until it intersected the center of mass of the posterior pons. This line describes our anterior-posterior axis (y axis, pointing anterior). The intersection of these two lines describes our origin. The perpendicular to these two axes was then computed as a left right axis (x axis, pointing right). Given these annotations, a rigid transformation of our MRI was computed so these axes were aligned to a right-anterior-superior (RAS) voxel grid, using a common convention in MRI such as the MNI space atlas^34^.

### Transformation graphs

Each acquired dataset is associated with a space, each with its own origin and axes. Each of these spaces is represented as a node in a transformation graph (Figure 2, filled circles). In addition to these, we create several new spaces: a Nissl registered space (where Nissl sections are are rigidly aligned and stacked into a 3D volume), an RAS space for the 13 week subject, an RAS space for the 21 week subject, and an “frechet mean” RAS space. These spaces are also associated with nodes in the graph (Figure 3, empty circles). A transformation can be computed to link any pair of these spaces, forming a directed edge. The “forward” transformation can be used to transform data in the direction of the arrow, and the “inverse” transformation can be used to transform data in the opposite direction.

Our registration strategy is to link each node in the graph by computing transformations between pairs of similar imaging datasets (which can be registered more accurately than dissimilar datasets). We use three main types of transformations: identity (red arrows) which means these two datasets are in the same space, rigid (blue line) which means we perform only a linear reorientation of axes and shifting of origin, and deformation (black arrows) which means all of shape, scale, and pose may be different between the pair of datasets. These pairwise transformations forms a tree (directed acyclic graph), such that a unique path can be traced from any space to any other space, which can be identified using a depth first search^10^. By composing transformations or their inverses along this path, a transformation can be computed to link any two spaces. This allows us to produce multimodal datasets in the same space.

We manage our growing datasets using a dynamic website built in django, where samples, spaces, images, and transformations correspond to database objects. Transformation graph layouts are computed automatically using classical multidimensional scaling, ensuring images from the same sample are close. These graphs are rendered as svg files, where nodes and edges contain hyperlinks to describe more information about imaging data and transformations.

### Multi-modality image alignment

Our approach to image alignment is to estimate a sequence of spatial and pixel value transformations to synthesize one observed dataset (the target, or fixed image) from another (the template, or moving image) [figure 3a]. The transformations we use are (i) a diffeomorphism modeling shape differences computed by integrating a time dependent dependent velocity field under the Large Deformation Diffeomorphic Metric Mapping (LDDMM) framework^22^, (ii) a 12 parameter affine transformation modeling orientation, scale and shear, (iii) a 3 parameter rigid transformation on each histology slice modeling positioning of tissue onto the microscope, (iv) a polynomial contrast transformation, defined in small blocks, to transform the intensity of one image to match the others. This procedure is detailed in^10,21^. For maps between two MRI scans (iii) are set to identity. For maps between Nissl and H&E sections, (i) and (ii) are set to identity. After applying transformations, we evaluate weighted sum of square error in pixel values between the transformed template and the target. Weights are computed in an Expectation Maximization framework^35^ to minimize the impact of any missing or damaged tissue, or artifacts. Weighted sum of square error is minimized using gradient based methods^22,36^ in pytorch, subject to smoothness as in standard LDDMM.

### Average age atlas estimate

After fixing the RAS coordinate system, we estimate an average MRI image “half way” between the 13 week and 21 week CCF brains. This is a Frechet mean procedure, where (i) an average image is deformed to each MRI using the procedure described above, and (ii) the inverse deformation is applied to each MRI, and a new average image is computed. Steps (i) and (ii) are repeated until convergence, which occurs when regularization cost is balanced against image matching cost. One example of the procedure is described in detail in^37^. In our case we enforce that the scale change from one the average to one MRI, is the inverse of the scale change from the average to the other image.

### Diffeomorphic interpolation

We construct a dense 3D segmented volume from a set of sparse segmentations using a diffeomorphic interpolation technique similar to^24^. We first identify a pair of consecutive labeled slices. In these slices we identify all brain areas that are labeled in both slices, at each level of the ontology from coarse to fine. We then compute a symmetric diffeomorphic alignment (advantages of this symmetry in certain cases are discussed in^38^ from one to the other. The LDDMM framework computes not just a single displacement field, but an entire time dependent trajectory. This trajectory is used to deform labels onto each of the intermediate labeled slices.

## Supporting information

Regional Cell Counts

GW14 atlas plates

GW21 atlas plates

GW14_MitoticFiguresPlates

## Acknowledgements

The experimental work was supported by the office of the Principal Scientific Adviser, Government of India and the Pratiksha trust. Computational work was supported by the H.N.Mahabala Chair professorship at IIT Madras.

P.P.M acknowledges support from the H.N.Mahabala chair professorship at IIT Madras as well as the Crick-Clay professorship at CSHL.

## Author contributions

The tape-transfer based pipeline utilized in the manuscript follows on methodology developed originally in P.P.M.’s laboratory. P.P.M. and M.S. conceived of the study and the experimental pipeline for data collection and implemented it together with J.J., R.V. and J. Joseph., with advice from S. Savoia. Instrumental automation and improvements for larger brain samples was led by J. Joseph. Samples were provided by J.K., C.S., S.S., L.S., H.E.K. together with necessary IRB and institutional permissions. Data collection was supervised by R.V. and J.J. with guidance from M.S., and P.P.M. Cell detection was performed by S.B., and S.M., proofread and collated by J.J., and regional counts computed by P.P.M. Atlas compartments were delineated by M.B., R.V. and J.J. Multimodal brain registration was performed by D.T. The fontanelle-based coordinate system was introduced by P.K.R., marked on the MRIs by J.J. and computationally implemented by D.T. The manuscript was written and figures assembled by P.P.M., D.T., J.J., and further edited by T.J.N., A.B.

## Competing interests

The authors declare no competing interests

## Materials & Correspondence

All correspondence is to be addressed to Partha Mitra at

## Data Availability statement

In addition to the supplementary materials provided with the manuscript, additional data sets associated with the paper are made available via Dryad at DOI: 10.5061/dryad.crjdfn3f3. These include five multimodal reference brains at 60 isotropic resolution, including MR, Nissl and H&E images, and annotated atlas plates with a coordinate system superposed for two brains at 13GW and 21GW.

## Code Availability

Cell detection code used in the manuscript may be found on Github at https://github.com/mitragithub/NISSL-Cell-Detection.git

Code for multimodal brain registration used in the manuscript may be found on Github at https://github.com/twardlab/emlddmm

---

**Table S1:**
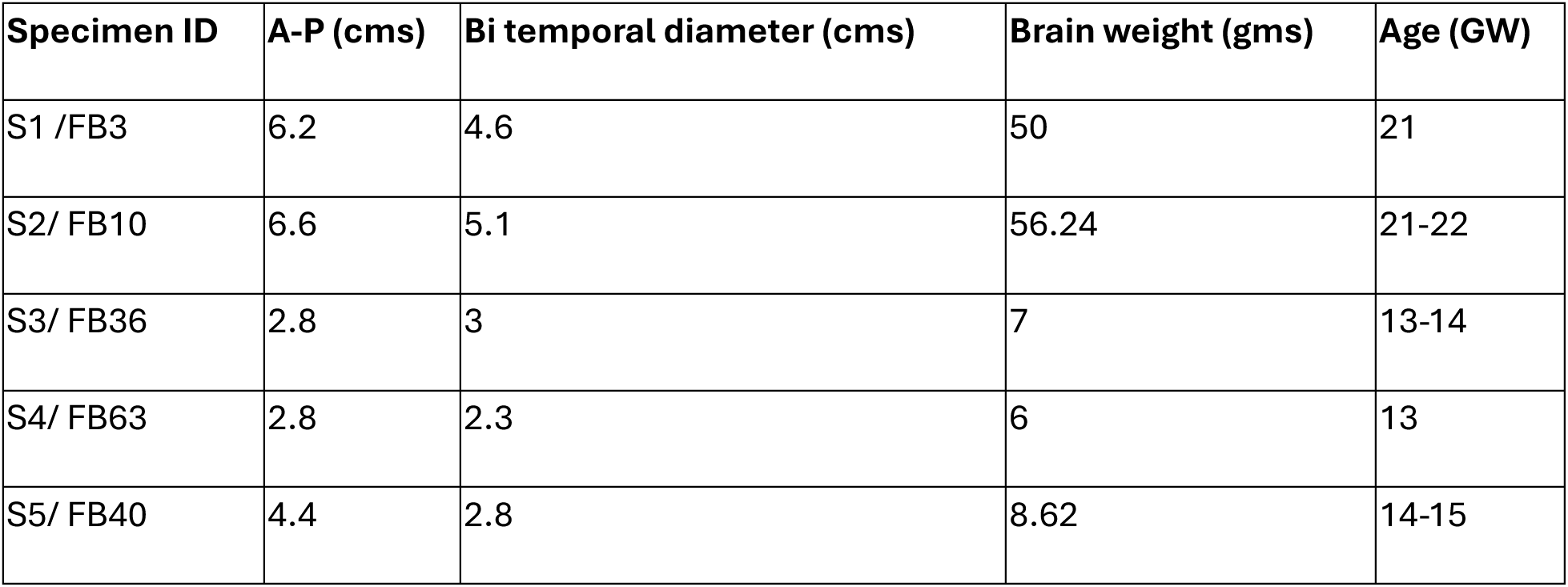
Demographics of donors and input datasets derived from donors.

| **Specimen ID** | **A-P (cms)** | **Bi temporal diameter (cms)** | **Brain weight (gms)** | **Age (GW)** |
| --- | --- | --- | --- | --- |
| S1 /FB3 | 6.2 | 4.6 | 50 | 21 |
| S2/ FB10 | 6.6 | 5.1 | 56.24 | 21-22 |
| S3/ FB36 | 2.8 | 3 | 7 | 13-14 |
| S4/ FB63 | 2.8 | 2.3 | 6 | 13 |
| S5/ FB40 | 4.4 | 2.8 | 8.62 | 14-15 |

## S2: Summary of available atlases and volumes of fetal brain development

**Table A.** Histology based atlases of human brain development.

| Features | Bayer and Altman (2005)<sup>1</sup> | Allen Brain Institute human developmental atlas<sup>2</sup> |
| --- | --- | --- |
| Details of dataset | 2D print atlas (300dpi) | 2D digital (1 um/pixel) available online |
| Number of brains/age | 9 whole brains/ 13.5to 24GW | 2 Hemispheres - 17GW(15PCW), 23GW(21PCW) and one brainstem 23GW(21PCW) |
| Annotated sections | 88 annotations across 9 brains | 168 annotations across the 3 samples |
| Maximum number of annotations per brain | 20 | 81 |
| Section thickness | 35um | 20um |
| Spacing between sections | Variable and uneven | 250-1200um |
| Embedding | Celloidin | OCT |
| Staining | Nissl and Myelin for older brains | Nissl/ ISH/ AchE/ Myelin for older brains |

**Table B.** MRI based data volumes and atlases of human brain development.

| Atlas/Vol | Information |  |  | Atlas Generation method |  |  | N | Method details |
| --- | --- | --- | --- | --- | --- | --- | --- | --- |
|  | Modality | Age (GW) | Resolution | Registration | Atlas construction | Kernel regression |  |  |
| Habas et al., 2010<sup>3</sup> | T2w | 21-24 | 0.5mm | Elastic deformation based on tissue maps | Group-wise | No | 20 | 1.5 T 1x1x3 mm<sup>3</sup> |
| Serag et al., 2011<sup>4</sup> | T2w | 23-37 | 1.18mm | Affine + Bspline free form deformation | Pairwise | Adaptive Gaussian kernel | 80 | 1.5T 1.25x1.25x2.5 mm<sup>3</sup> |
| Zhan et al., 2013<sup>5</sup> | Post-mortem T2w | 15-22 | — | SyN in ANTs | Group-wise | No | 34 | 7T 0.19x0.19x0.5mm<sup>3</sup> or 0.23x0.23x0.5mm<sup>3</sup> |
| Dittrich et al., 2014<sup>6</sup> | Ventricle only | 20 - 30 | 0.78-0.9mm | Semi-supervised learning | — | Gaussian kernel | 32 + 12* | 1.5T 0.78-0.9 |
|  |  |  |  |  |  |  |  | mm<br>(x,y)<br>3-4.4mm<br>slices |
| Wright et al., 2015<sup>7</sup> | Cortical<br>Surface<br>Atlas | 23-37 | — | Surface<br>registration<br>based on<br>sulcal<br>alignment |  | Gaussian<br>kernel | 80 | 1.5T |
| Gholipour et al.,<br>2017<sup>8</sup> | T2w | 21-37 | 0.8mm | Greedy<br>SyN in<br>ANTs | Group-wise | Fixed-<br>width<br>Gaussian<br>kernel | 81 | 3T or<br>1.5T<br>0.9~1.<br>1-2mm<br>slices |
| Xia et al., 2019<sup>9</sup> | Cortical<br>Surface<br>Atlas | 26.5,<br>27,<br>28.2,<br>28.7 |  | Surface<br>registration | — | No | 25 | 1.5T<br>0.5×0.<br>5×2.5<br>mm<sup>3</sup> |
| Li et al., 2021<sup>10</sup> | T2w | 23-36 | 0.8mm | Affine +<br>SyN in<br>ANTs | Pairwise<br>+ group-wise | No | 212 | 1.5T<br>0.74×0<br>.74×4<br>mm<sup>3</sup> |
| Wu et al., 2021<sup>11</sup> | T2w | 22-35 | 0.8mm | SyN in<br>ANTs | Group-wise | Adaptive<br>Gaussian<br>kernel | 735 | 1.5T<br>4mm<br>slice |
| Uus et al., 2021<sup>12</sup> | T2w | 21-36 | 0.7mm | MIRTK<br>atlas<br>generation<br>pipeline | — | Gaussian<br>kernel | 291 | 3T<br>1.1×1.<br>1×2.2<br>mm<sup>3</sup> |
| Xu et al 2022<sup>13</sup> | T2w | 23-38 | 0.8mm | ANTs<br>Rigid+affin<br>e+SyN | Pairwise<br>+group-<br>wise | Adaptive<br>Gaussian<br>kernel | 219 | 3T<br>1.1×1.<br>1×2<br>mm<sup>3</sup> |

\*abnormal brains
Abbreviations: GW = gestational age; SyN = symmetric diffeomorphic deformable registration algorithm; ANTs = advanced normalization tools.

## S3: Stereology correction factor

**Figure 1.**
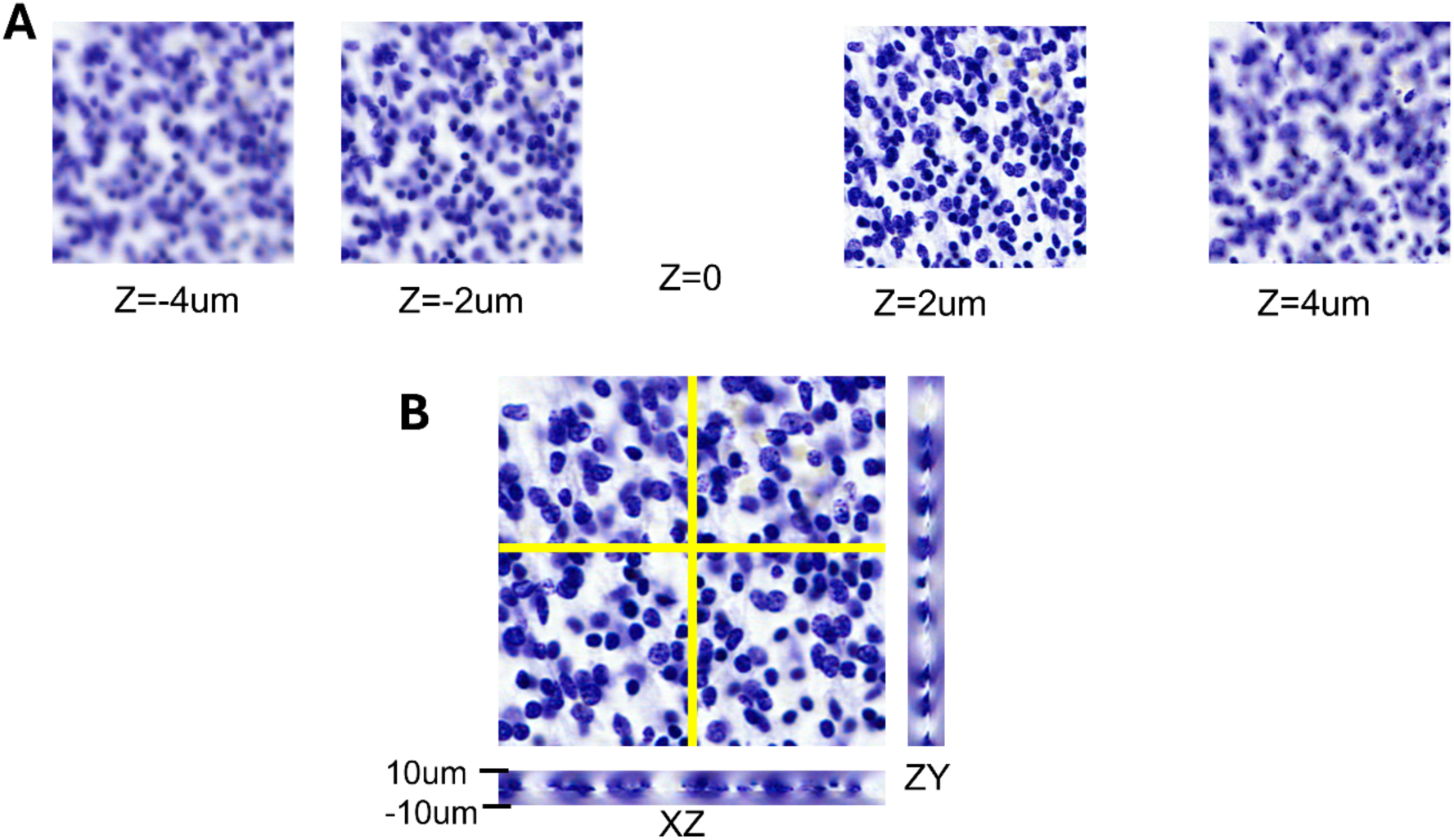
A shows the panel of images obtained as a z-stack (of z-resolution = 1um) of a Thionin-stained section from the 21GW fetal brain. Panel B shows the in-plane focused image (z=0) along with z-cross sections (marked as ZY and XZ) across the x and y planes given by the yellow lines.

We calculated the correction factor and the post-processing effective thickness for computing the cell densities using a z stack of images (1um z-resolution) obtained by our imaging system (Huron LE 120). Figure S xx shows the z stack of images of a thionin-stained section of 20 um thickness. Panel A shows the z stack of images that are focussed between -4um to +4um from an image deemed to be the in-focus plane. Panel B shows the in-plane focussed image and the corresponding z slices in the X and Y axis respectively. This method was performed on several randomly chosen regions with varying cell densities to derive the correction factor.

We calculated the following

1. The thickness of the section (T) was assumed to be 20 um
2. The centres of detected cells within the in-plane focussed sections were marked as mask (A).
3. As one moves through the z-stack of images, any cells that were potentially out of focus were appended to the original detection
4. The correction factor (ƥ or rho) was calculated as a ratio of the total detected cells and the detected cells in the focussed plane.
5. Section densities were calculated as (A x ƥ)/T.

Using this method, we were able to estimate the average correction factor for the whole brain as 1.04. By observing the effective foci of objects within the section, we could estimate the post-processing thickness of sections, which shows a shrinkage following histological processing. In the example shown above, the objects were in focus only in 14 out of 20 sections within the z-stack. The post-processing thickness estimate for the random samples across the tissue sample was between 13-15 um.

## S4: Quantifying regional growth in cell counts in the developing human brain: literature review

Despite the large literature on neurogenesis and brain development, very few studies (table 1) have numerically quantified cell counts in the developing human brain, and all previous studies are based on statistical sampling approaches. Previous studies have methodological challenges including non-uniform and sparse statistical sampling, assumptions and/or stereological correction factors, which add to the sample procurement and fixation issues commonly associated with fetal brains.

Earlier, methods such as estimation of DNA weights as a surrogate of cellular growth showed that there is a rapid exponential growth between 10-18 GW followed by a slower phase of expansion after 20GW^1^. H-Thymidine DNA was used as a marker to show that almost all cortical neurogenesis takes place in the first phase of pregnancy^2,3^. More recently, quantification of cellular counts as a function of developmental time was achieved using histology based methods based on random sampling and the extrapolation of cell counts using the optical fractionator approach, in a very limited number of studies^4–8^. Further, regional cell counts have been confined to the developing cortex^7^.

We provide the first comprehensive cell counts based on machine-vision based detection and counting of cells in closely spaced histological sections across the whole brain, which do not suffer from the sparse sampling limitations of the previous studies. In addition, we also provide the first regional counts in a comprehensive set of compartments including subcortical structures. A comparison between our counting estimates and literature data (Table 2) shows that where these estimates exist, they are generally consistent with our findings.

**Table 1.** Human brain development cell counts.

| Ref | Method | Age /n | Whole brain/<br>parts | Output relevant to this paper |
| --- | --- | --- | --- | --- |
| **Histology based cell counts** |  |  |  |  |
| Samuelsen et al 2003<sup>7</sup> | Cell counts with histology<br>GIEMSA<br>40um paraffin sections | 13- 42 GW<br>(n=22) | Human fetal telencephalon | Random sampling within sections (100 objects per view) x 10 -12 sections per brain<br>Estimated cell counts= 4.5 billion (13 GW)<br>15 billion (21 GW) |
| Larsen et al 2006<sup>5</sup> | Cell counts with histology<br>GIEMSA<br>40um paraffin sections | New-Born<br>(38-42 GW<br>n=10) | Human fetal telencephalon | Random sampling within sections (100 objects per view) x 10 -12 sections per brain<br>The mean total cell number was 32.6 billion cells (coefficient of error=0.04) |
| Samuelsen et al, 2007<sup>6</sup> | Cell counts with histology<br>GIEMSA<br>40um paraffin sections | 18-42 weeks GW | Human fetal telencephalic wall | Estimated cell counts = 15.45 Billion cells (21GW) (same samples as previous paper?) |
| Rabinowicz et al 1996<sup>4</sup> | Cell counts with histology | 17 GW to 17 | Regions -Precentral gyrus | Estimated cell density = 6000 cells/0.001mm<sup>3</sup> (21 GW)<br>4000 cells/0.001mm<sup>3</sup> (17 GW) |
|  | Cresyl violet<br>10-25um<br>Paraffin<br>sections | weeks<br>post<br>birth<br>(n= 9) | -Wernicke's<br>area<br>-Broca's area<br>-Calcarine<br>cortex<br>-Cingulate<br>cortex<br>-Frontal pole |  |
| **Other methods (qualitative no cell counts)** |  |  |  |  |
| Rakic & Sidman, 1973<sup>2</sup> | H-thymidine DNA | <6 – 16GW | Radiological | Shows neuronal differentiation from progenitors happens only in early gestation |
| Dobbing and Sands, 1973<sup>1</sup> | pDNA and Cholesterol | 10GW to 8 years post birth | pDNA (weight per volume) | Qualitative showing two phases of growth : 1st rapid phase between 10-20 GW and a slower growth from 20GW |

**Table 2.** shows the comparison of total and regional cell counts of the fetal brain available in the literature compared to our current study. The literature data is derived from studies using the optical fractionator methods.

| Region | Rabinowicz et al 1996<sup>4</sup> | Samuelsen et al 2000<sup>8</sup> | Samuelsen et al 2003<sup>7</sup> | Samuelsen et al 2007<sup>6</sup> | This paper |
| --- | --- | --- | --- | --- | --- |
| Total | 6000 cells/mm<sup>3</sup><br>(<u>21GW</u>) = 7.5 x10<sup>9</sup> (assuming 50 x 50 x 50 mm) |  | <u>13GW</u><br>3.01 x 10<sup>9</sup><br><u>20 GW</u><br>12.8 x 10<sup>9</sup><br><u>22GW</u>=<br>16.7 x 10<sup>9</sup> | <u>22GW</u><br>16.7 x 10<sup>9</sup> | <u>13GW</u><br>3.7 x 10<sup>9</sup><br><u>21GW</u><br>17.4 x 10<sup>9</sup> |
| MZ/CP | - | <u>22GW</u><br>5.7 x 10<sup>9</sup> | <u>13GW</u><br>0.96 x 10<sup>9</sup><br><u>20GW</u><br>5.87 x 10<sup>9</sup><br><u>22GW</u><br>7.05 x 10<sup>9</sup> | - | <u>13GW</u><br>0.02x 10<sup>9</sup><br>(MZ) and<br>0.73 x 10<sup>9</sup><br>(CP)<br><u>21GW</u><br>0.26 x 10<sup>9</sup><br>(MZ) and<br>5.3 x 10<sup>9</sup><br>(CP) |
| SP | - |  | <u>13GW</u><br>0.66 x 10<sup>9</sup><br><u>20GW</u><br>2.4 x 10<sup>9</sup><br><u>22GW</u><br>1.9 x 10<sup>9</sup> | - | <u>13GW</u><br>0.77 x 10<sup>9</sup><br><u>21GW</u><br>2.6 x 10<sup>9</sup> |
| IZ | - | <u>22GW</u><br>2.2 x 10<sup>9</sup> | <u>13GW</u><br>0.8 x 10<sup>9</sup><br><u>20 GW</u><br>3.06 x 10<sup>9</sup><br><u>22GW</u> | - | <u>13GW</u><br>0.80 x 10<sup>9</sup><br><u>21GW</u><br>3.82 x 10<sup>9</sup> |
|  |  |  | 2.58x 10<sup>9</sup> |  |  |
| VZ/SZ | - | <u>22GW</u><br>5.1 x 10<sup>9</sup> | <u>13GW</u><br>0.57 x 10<sup>9</sup><br><u>20GW</u><br>1.44 x 10<sup>9</sup><br><u>22GW</u><br>4.58 x 10<sup>9</sup> | - | <u>13GW</u><br>0.25 x 10<sup>9</sup><br>(SZ) and<br>0.24 x 10<sup>9</sup><br>(VZ)<br><u>21GW</u><br>0.96 x 10<sup>9</sup><br>(SZ) and<br>0.12 x 10<sup>9</sup><br>(VZ) |

## S5: Evaluation of the detection of mitotic figures in H&E

We evaluated the markings and detections of mitotic figures in the H&E sections by comparing them with adjacent sections stained with immunohistochemistry of the proliferative marker Ki-67. Figure 1 shows two regions from the H&E stained histological section (A) and the adjacent section stained using DAB-based immunohistochemistry of the Ki-67. As shown in the figure, the detection of mitotic figures in the H&E stained sections are limited by the relative contrast of the stain. The Ki-67 antibody stain provided more contrast and more mitotic figures could be detected. Examination of the relative numbers and spacing of the mitotic figures detected in the Ki-67 stained sections, showed a ratio of about two. This implies that the mitotic figures visually detected in the H&E sections may be underestimating the correct number by a factor of two.

**Figure 1.**
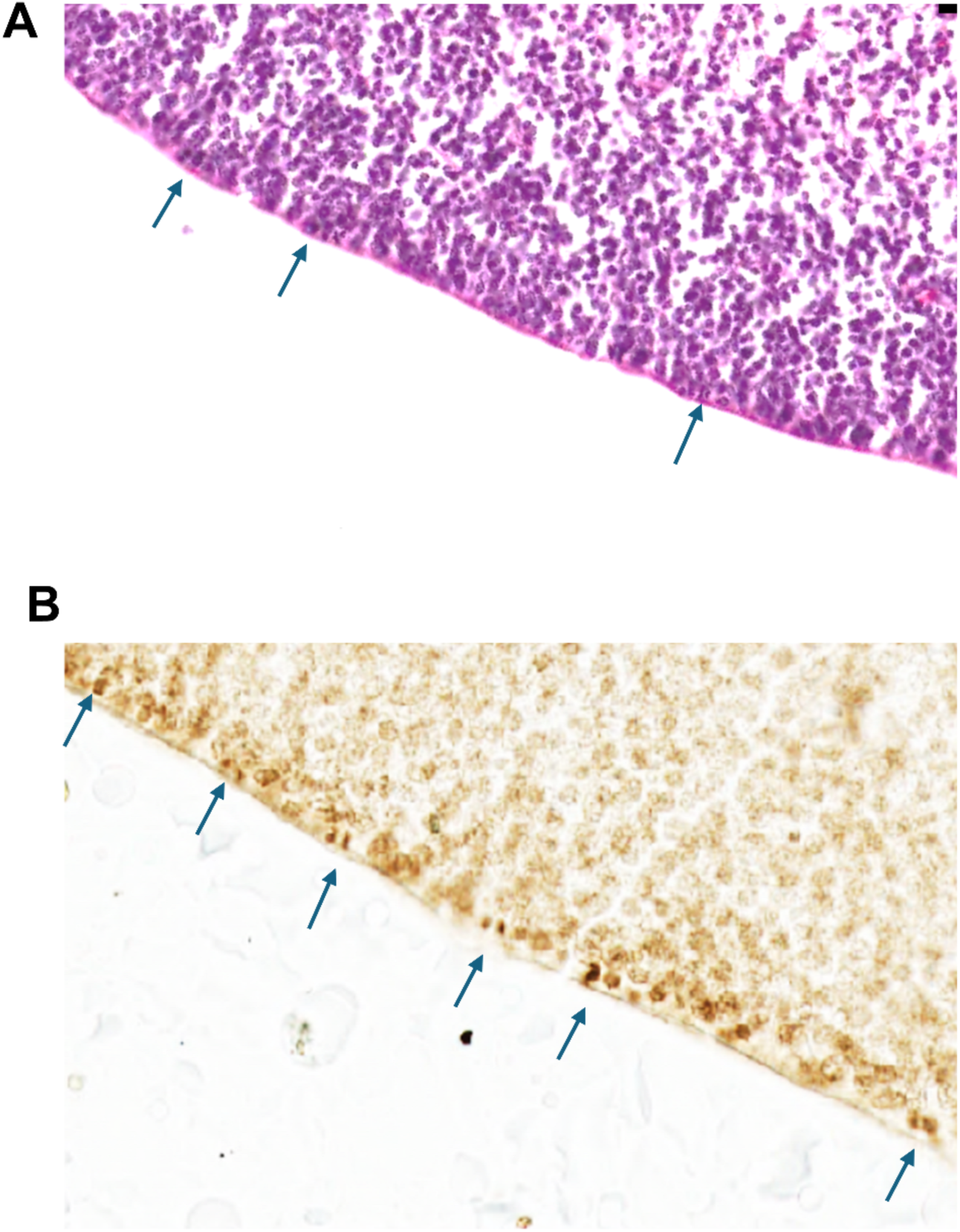
shows detected mitotic figures in the ventricular zone (VZ) of a section stained with H&E (A) and an adjacent section stained for Ki-67, a proliferative marker (B). The arrows show the detected mitotic figures in both staining.

### Nomenclature for developing brain compartments

The existing nomenclature for atlas compartments of the developing human brain is complex and includes names and abbreviations of not only the evolving brain parts (brain regions, fiber tracts and ventricles but also the transitive developmental structures, such as the ganglionic eminence and the migratory streams.

#### Problems associated with atlas nomenclatures and our approach based on coordinate frameworks and a flat list of cytoarch1tectonic regions

Brain atlases have named compartments, with the terminology for mammals being derived from the early work of His^1^. Over time, there has been a proliferation of atlases that differ in compartment boundaries and nomenclatures. Even within a given species, there are multiple atlases with nomenclatures that are not easy to reconcile. In humans, the complexity is higher owing to the larger number of distinct anatomical parcellations. In addition to the challenges that exist for adult atlas nomenclature, development atlases add the extra dimension of time. Adult atlases may be regarded as parcellations of brain space. Developmental atlases conceptually require a parcellation of developmental space-time, considering the formation of new compartments at relevant developmental time points as well as the disappearances of transient structures. Even if a compartment remains trackable throughout development, functional nomenclature appropriate for the adult may not be appropriate for the developing fetus. For these reasons, it is preferable to move towards a common coordinate system-based approach, where a specific location in the brain at a particular age is specified using an appropriate three-dimensional coordinate in a reference brain.

This has been our motivation in focusing on the coordinate framework and reference brain volumes in the current manuscript. Nevertheless, to maintain continuity with the literature we also provide a parcellation and nomenclature for the developmental ages under study in the manuscript. We adopt an approach developed earlier^2^ in which we prioritize a parcellation of the brain into non-overlapping compartments, the union of which gives back the entire brain space. The parcels are chosen to have visibly well-defined boundaries based on cytoarchitectonic differences across boundaries at a given age. We do not focus on how to group those parcels into a hierarchy since that is often the origin of the debates between neuroanatomists. Some high-level groupings are obvious and canonical and may be easily layered on top of the lowest level parcels.

#### Discussion of previous atlases and nomenclatures of the developing human brain

The relevant nomenclature for developing mammalian brains has largely been derived so far from developmental studies in model organisms, mostly rodents and primates. This includes parcellations based on a hierarchical gene nomenclature (The Gene Ontology^3^) with more than 500 directly related terms that are associated with textual definitions, and references and models the development of a neuronal structure to the developmental process (e.g., “telencephalon development is part of forebrain development”;^3^). However, this approach is also limited as it does not fully address the spatial and temporal aspects of the brain development and does not reconcile the problem of different nomenclatures within the same species. Puelles and colleagues^4^ have attempted to reconcile the classical nomenclature with gene expression patterns and those derived from embryonic prosomeres^4,5^. The problem of the developing cortical regions was addressed by the Boulder Committee nomenclature^6,7^, however, this nomenclature leaves out subcortical regions.

The widely accepted print atlas of human brain development by Bayer and Altmann^8^ uses nomenclature derived from adult brain atlases/textbooks supplemented with Bayer and Altmann’s own work on rat brain development^9^. This atlas, however, has conflicting names particularly for the developing cortical zones, referred to as the Stratified transitional fields in their publications. More recently, developmental human brain nomenclature was developed for 15 PCW and 22 PCW brains by the Allen Brain Institute^10^, in an attempt to reconcile the ontology used by the Boulder Committee and the adult brain atlases. These authors provide a hierarchically organized nomenclature that include more than 3000 terms for both the fetus and adult human brain. These terms include names for adult brain regions which are based on “function” which are questionable when applied to fetal brains as there is little experimental basis for such functional designation. Further, several terms are included which are not associated with textual definitions, and not all of the terms are are associated with atlas delineations.

#### Details of our approach

As previously discussed, our focus has been on establishing a reference coordinate system based on skull landmarks, with the idea that atlas parcellations or annotations may be regarded as superposed layers on top of the underlying coordinate system in the reference brains we present. For atlas annotation purposes we have carried out a histology and cytoarchitecture based parcellation of the entire brains involved, while maintaining correspondence with known brain atlases^7,8,10^. Regarding nomenclature, we therefore focus on a flat list of names of these compartments. Table 1 below highlights the marked regions for the plates of histological samples used in this study namely the 13 GW and 21 GW fetal brains.

The list of names we use in this manuscript to designate nonoverlapping compartments include 119 terms presented in the 2D atlas plates as well as in the regional cell count tables. Each term has a name, abbreviation, a textual definition, and a hexadecimal color code used to mark the associated delineations. The terminology included largely follows the nomenclatures proposed by Bayer and Altman^8^, AIBS^10^, and by the Boulder Committee^6,7^. We also included terms proposed by Zecevic^11^, and by Kostovic & Judas^12^. The nomenclature we utilize includes terms and definitions specific only to the human brain, and it is generally not hierarchically organized, although we utilize some generally agreed groupings (eg “thalamus” or “basal ganglia”) to summarize our tables of counts, or to assign residual compartments that are not otherwise specified after individual sub-nuclei have been specified.

**Table 1.** Named brain compartments for the 13GW and 21GW brains relevant to this manuscript.

| 13GW |  |  | 21GW |  |  |
| --- | --- | --- | --- | --- | --- |
| ID | Brain Region | Abbr | id | Brain region | Abbr |
| 10157 | gray matter of forebrain | FGM | 10157 | gray matter of forebrain | FGM |
| 10349 | basal forebrain | BF | 10349 | basal forebrain | BF |
| 11580 | subpial granular zone | SG | 11580 | subpial granular zone | SG |
| 10508 | marginal zone | MZ | 10508 | marginal zone | MZ |
| 10515 | cortical plate | CP | 10515 | cortical plate | CP |
| 10522 | subplate zone | SP | 10506 | subplate zone | SP |
| 10529 | intermediate zone | IZ | 10529 | intermediate zone | IZ |
| 10536 | subventricular zone | SZ | 10522 | subventricular zone | SZ |
| 10542 | ventricular zone | VZ | 10542 | ventricular zone | VZ |
| 10506 | transient structures of forebrain | FTS |  |  |  |
| 10549 | Ganglionic eminence (VZ in basal nuclei) | GE | 10549 | Ganglionic eminence (VZ in basal nuclei) | GE |
| 10294 | hippocampus (hippocampal formation) | HIP | 10294 | hippocampus (hippocampal formation) | HIP |
| 12170 | hippocampal gyrus (formation) | HiF | 12170 | hippocampal gyrus (formation) | HiF |
| 12162 | parahippocampal gyrus | PHG | 12162 | parahippocampal gyrus | PHG |
| 10361 | amygdala | AMY | 10361 | amygdala | AMY |
| 10312 | piriform-entorhinal-amygdaloid area | PEA | 10312 | piriform-entorhinal-amygdaloid area | PEA |
| 10311 | Piriform cortex | Pir | 10311 | Piriform cortex | Pir |
|  |  |  | 10317 | entorhinal cortex | EC |
|  |  |  | 10288 | insular cortex | ICx |
| 10304 | indusium griseum | IG | 10304 | indusium griseum | IG |
| 10350 | septal nuclei | SEP | 10350 | septal nuclei | SEP |
|  |  |  | 13076 | subcallosal gray zone | ScG |
|  |  |  | 11634 | MZ in insular cortex | iMZ |
|  |  |  | 11728 | CP in insular neocortex | iCP |
|  |  |  | 11742 | CP in entorhinal cortex | CPEC |
|  |  |  | 13207 | SG of the allocortex | alSG |
|  |  |  | 10513 | MZ of the allocortex | alMZ |
|  |  |  | 10534 | IZ of the allocortex | alIZ |
|  |  |  | 11638 | MZ in subicular cortex | MZsub |
|  |  |  | 11738 | CP in subicular cortex | CPsub |
|  |  |  | 11787 | SP in the subicular cortex | SPsub |
|  |  |  | 11875 | IZ in subicular cortex | IZsub |
|  |  |  | 13280 | CP in parasubicular cortex | CPpas |
| 13078 | perifornical gray zone | PFG | 13078 | perifornical gray zone | PFG |
| 10331 | basal nuclei | BN | 10331 | basal nuclei | BN |
| 10334 | caudate nucleus | Ca | 10334 | caudate nucleus | Ca |
| 1.46E+08 | Caudate Putamen cell bridges | CaPu | 1.46E+08 | Caudate Putamen cell bridges | CaPu |
|  |  |  | 10335 | head of the caudate | CaH |
| 10338 | Putamen | Pu | 10338 | Putamen | Pu |
| 10333 | Striatum | STR | 10333 | Striatum | STR |
| 10346 | Clastrum | Cla | 10346 | Clastrum | Cla |
| 10339 | nucleus accumbens | NAC | 10339 | nucleus accumbens | NAC |
| 10342 | globus pallidus | GP | 10342 | globus pallidus | GP |
| 10384 | bed nucleus of stria terminalis | BNST | 10384 | bed nucleus of stria terminalis | BNST |
| 10573 | external capsule | extC | 10573 | external capsule | extC |
| 10581 | internal capsule | ic | 10581 | internal capsule | ic |
| 10559 | anterior commissure | ac | 10559 | anterior commissure | ac |
| 10557 | white matter of forebrain | fwm | 10557 | white matter of forebrain | fwm |
| 10591 | stria terminalis | st | 10591 | stria terminalis | st |
| 10576 | fornix | fx | 10576 | fornix | fx |
| 10561 | corpus callosum | cc | 10561 | corpus callosum | cc |
| 12021 | olfactory recess | olr | 12021 | olfactory recess | olr |
| 10567 | posterior commissure | poc | 10567 | posterior commissure | poc |
| 10644 | optic chiasm | oc | 10644 | optic chiasm | oc |
| 10390 | thalamus | THM | 10390 | thalamus | THM |
|  |  |  | 10364 | central medial nucleus of thalamus | CeM |
|  |  |  | 10391 | dorsal thalamus | DTH |
|  |  |  | 10397 | medial nuclear complex of thalamus | MNC |
| 10464 | reticular nucleus of thalamus | R | 10464 | reticular nucleus of thalamus | R |
|  |  |  | 10409 | Pulvinar of the thalamus | Pul |
|  |  |  | 10389 | subthalamus | SubTH |
| 10465 | subthalamic nucleus | STH | 10465 | subthalamic nucleus | STH |
|  |  |  | 12101 | subcommisural organ | SCO |
| 12247 | red nucleus | RN | 12247 | red nucleus | RN |
| 10451 | epithalamus | ETH | 10451 | epithalamus | ETH |
| 10467 | hypothalamus | HTH | 10467 | hypothalamus | HTH |
| 10556 | rostral migratory stream | RMS | 10556 | rostral migratory stream | RMS |
| 10463 | zona incerta | ZI | 10463 | zona incerta | ZI |
| 13034 | substantia innominata | SI | 13034 | substantia innominata | SI |
| 10462 | nucleus of the field of Forel | FF | 10462 | nucleus of the field of Forel | FF |
| 12181 | pretectal region | PTR | 12181 | pretectal region | PTR |
| 12292 | superior colliculus | SC | 12292 | superior colliculus | SC |
| 12305 | inferior colliculus | IC | 12305 | inferior colliculus | IC |
| 12209 | periaqueductal gray | PAG | 12209 | periaqueductal gray | PAG |
| 10650 | white matter of midbrain | mwm | 10650 | white matter of midbrain | mwm |
|  |  |  | 10659 | cerebellum | CBH |
| 13363 | external granular (germinal) zone of lateral hemisphere | CbLTe | 13363 | external granular (germinal) zone of lateral hemisphere | CbLTe |
| 13364 | marginal zone of lateral hemisphere | CbLTm | 13364 | marginal zone of lateral hemisphere | CbLTm |
| 13366 | inner granular cell zone of lateral hemisphere | CbLTi | 13366 | inner granular cell zone of lateral hemisphere | CbLTi |
|  |  |  | 13365 | Purkinje cell zone of lateral hemisphere | CbLTp |
| 13369 | ventricular (germinal) zone of lateral hemisphere | CBITv | 13369 | ventricular (germinal) zone of lateral hemisphere | CBITv |
| 13347 | external granular (germinal) zone of the vermis | VeTe | 13347 | external granular (germinal) zone of the vermis | VeTe |
| 13348 | marginal zone of the vermis | VeTm | 13348 | marginal zone of the vermis | VeTm |
| 13353 | ventricular (germinal) zone of the vermis | VeTv | 13353 | ventricular (germinal) zone of the vermis | VeTv |
|  |  |  | 13349 | Purkinje cell zone of the vermis | VeTp |
| 13350 | inner granular cell zone of the vermis | VeTi | 13350 | inner granular cell zone of the vermis | VeTi |
| 10660 | cerebellar deep nuclei | CbDN | 10660 | cerebellar deep nuclei | CbDN |
| 10664 | Rhombic lip | RhL |  |  |  |
| 10310 | optic tract | ot | 10310 | optic tract | ot |
| 12230 | cerebral peduncle (crus cerebri) | cpd | 12230 | cerebral peduncle (crus cerebri) | cpd |
| 10668 | white matter of hindbrain | hwm | 10668 | white matter of hindbrain | hwm |
|  |  |  | 10652 | surface structures of midbrain | MSS |
| 12462 | superior olivary complex | SOC |  |  |  |
| 10649 | gray matter of midbrain | MGM |  |  |  |
| 12251 | substantia nigra | SN | 12251 | substantia nigra | SN |
| 10661 | Pontine nucleus | PN | 10661 | Pontine nucleus | PN |
| 12406 | pons | Pn | 12406 | pons | Pn |
|  |  |  | 12346 | afferent nuclei of cranial nerves in pons | PnAN |
|  |  |  | 12405 | basilar pons | PnBa |
|  |  |  | 12416 | pontine tegmentum | PnTg |
| 12195 | midbrain tegmentum | MTg | 12195 | midbrain tegmentum | MTg |
| 12645 | other nuclei in medullary tegmentum | XMoTg | 12645 | other nuclei in medullary tegmentum | XMoTg |
| 10662 | myelencephalon (medulla oblongata) | Mo | 10662 | myelencephalon (medulla oblongata) | Mo |
|  |  |  | 12535 | pyramidal part of medulla oblongata | MoPy |
| 12890 | spinal cord | SpC | 12890 | spinal cord | SpC |
| 10460 | pineal body | Pin | 10460 | pineal body | Pin |
| 12270 | interpeduncular nucleus | IP | 12270 | interpeduncular nucleus | IP |
| 12600 | inferior olive | IO | 12600 | inferior olive | IO |
|  |  |  | 12272 | pyramidal decussation | xpy |
|  |  |  | 10654 | gray matter of the hindbrain | HGM |
|  |  |  | 15544 | optic nerve | on |
|  |  |  | 12331 | corticopontine fibers midbrain portion | cpn-m |
|  |  |  | 1.46E+08 | white matter of the spinal cord | swm |
|  |  |  | 12811 | median aperture (foramen of Magendie) | ma4V |
|  |  |  | 12420 | facial nucleus | 7N |
| 1.46E+08 | transient cell zone in external capsule | Tcete |  |  |  |
| 12354 | superior cerebellar peduncle (brachium conjunctivum) | scp |  |  |  |
| 12098 | cavum pellucidum | Pell |  |  |  |
| 2.66E+08 | corticopontine fibers pontine part | cpn |  |  |  |
| 12304 | diagonal band | dib |  |  |  |
| 10590 | stria medullaris of thalamus | smt |  |  |  |
| 10568 | corona radiata | cor |  |  |  |
| 12344 | interpedunculotegmental tract | iptg |  |  |  |

## Notes

https://doi.org/10.5061/dryad.crjdfn3f3

https://github.com/twardlab/emlddmm

https://github.com/mitragithub/NISSL-Cell-Detection.git

### Competing Interest Statement

The authors have declared no competing interest.

