## Supplementary material for "A three-dimensional histological cell atlas of the developing human brain": GW14 atlas plates

### Histological reference atlas and coordinate system for a 14 gestational week fetal brain

December 10, 2024

Corresponding manuscript: Jai Jayakumar, et al. "A three-dimensional histological cell atlas of the developing human brain"

Demographic information: Specimen ID: S5/ FB40 AP (cms): 4.4 Bi temporal diameter (cms): 2.8 Brain weight (gms): 8.62 Age (GW): 14 Sex: M

This atlas includes serial section histology data, with neighoring sections aligned. Sections are 20um thick and collected in a series of three, alternating between Nissl, H&E, and various antibody stains (not shown), so that sections corresponding to a individual stain type are 60 um apart.

We have developed a standard RAS (x=Right, y=Anterior, z=Superior) coordinate system described as shown below. Please refer to figure 2 in the manuscript.

The data below shows this coordinate system as a curvilinear grid, with 10mm spaced isocontours, labeled in mm. Red lines correspond to the left-right axis with right positive. Green lines correspond to the posterior-anterior axis with anterior positive. Blue lines correspond to the inferior-superior axis with superior positive. The x=0, y=0, and z=0 lines are drawn thicker. Note that our sagittal histology sections are nearly perpendicular to the left right axis, and so red lines appear more curved.

Below we show alternating pages with manually drawn annotations with median spacing between annotations of 0.54mm. Corresponding Nissl stained sections on which annotations were drawn are also included, here downsampled to 64 micron in plane resolution. Finally where neighboring tissue sections are available with H&E stain, these sections are also shown also at 64 micron resolution.

The horizontal and vertical axes of the image in each section also labeled with tick marks in mm, with 10 mm spacing to help with understanding scale.

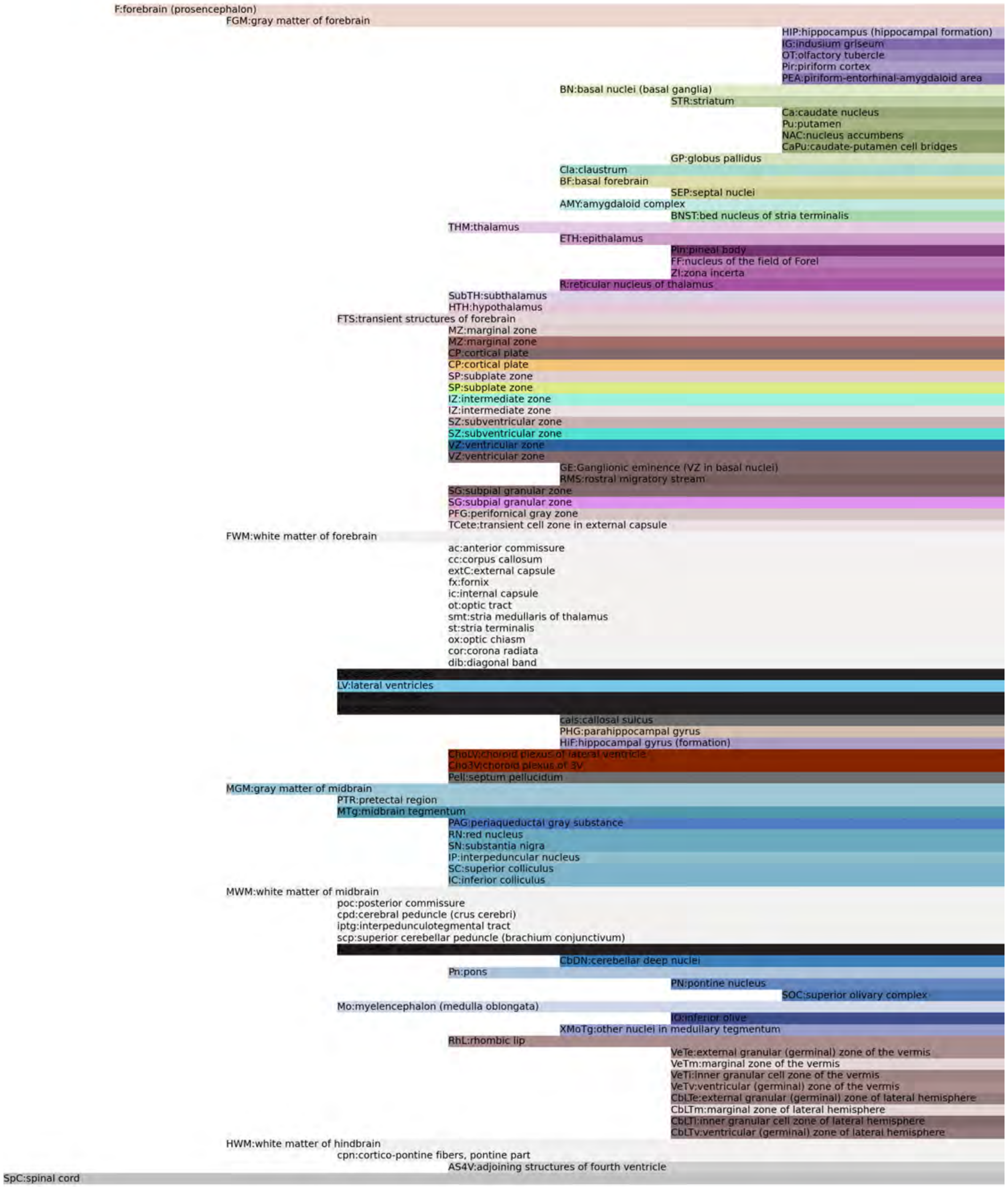
F:forebrain (prosencephalon)
FGM:gray matter of forebrain
HIP:hippocampus (hippocampal formation)
IG:indusium griseum
OT:olfactory tubercle
Pir:piriform cortex
PEA:piriform-entorhinal-amygdaloid area
BN:basal nuclei (basal ganglia)
STR:striatum
Ca:caudate nucleus
Pu:putamen
NAC:nucleus accumbens
CaPu:caudate-putamen cell bridges
GP:globus pallidus
Cla:claustrum
BF:basal forebrain
SEP:septal nuclei
AMY:amygdaloid complex
BNST:bed nucleus of stria terminalis
THM:thalamus
ETH:epithalamus
Pin:pineal body
FF:nucleus of the field of Forel
ZI:zona incerta
R:reticular nucleus of thalamus
SubTH:subthalamus
HTH:hypothalamus
FTS:transient structures of forebrain
MZ:marginal zone
MZ:marginal zone
CP:cortical plate
CP:cortical plate
SP:subplate zone
SP:subplate zone
IZ:intermediate zone
IZ:intermediate zone
SZ:subventricular zone
SZ:subventricular zone
VZ:ventricular zone
VZ:ventricular zone
GE:Ganglionic eminence (VZ in basal nuclei)
RMS:rostral migratory stream
SG:subpial granular zone
SG:subpial granular zone
PFG:perifornical gray zone
TCete:transient cell zone in external capsule
FWM:white matter of forebrain
ac:anterior commissure
cc:corpus callosum
extC:external capsule
fx:fornix
ic:internal capsule
ot:optic tract
smt:stria medullaris of thalamus
st:stria terminalis
ox:optic chiasm
cor:corona radiata
dib:diagonal band
LV:lateral ventricles
cals:callosal sulcus
PHG:parahippocampal gyrus
HiF:hippocampal gyrus (formation)
ChoLV:choroid plexus of lateral ventricle
Cho3V:choroid plexus of 3V
Pell:septum pellucidum
MGM:gray matter of midbrain
PTR:pretectal region
MTg:midbrain tegmentum
PAG:periaqueductal gray substance
RN:red nucleus
SN:substantia nigra
IP:interpeduncular nucleus
SC:superior colliculus
IC:inferior colliculus
MWM:white matter of midbrain
poc:posterior commissure
cpd:cerebral peduncle (crus cerebri)
iptg:interpedunculotegmental tract
scp:superior cerebellar peduncle (brachium conjunctivum)
CbDN:cerebellar deep nuclei
Pn:pons
PN:pontine nucleus
SOC:superior olivary complex
Mo:myelencephalon (medulla oblongata)
IO:inferior olive
XMoTg:other nuclei in medullary tegmentum
RhL:rhombic lip
VeTe:external granular (germinal) zone of the vermis
VeTm:marginal zone of the vermis
VeTi:inner granular cell zone of the vermis
VeTv:ventricular (germinal) zone of the vermis
CbLTe:external granular (germinal) zone of lateral hemisphere
CbLTm:marginal zone of lateral hemisphere
CbLTi:inner granular cell zone of lateral hemisphere
CbLTv:ventricular (germinal) zone of lateral hemisphere
HWM:white matter of hindbrain
cpn:cortico-pontine fibers, pontine part
AS4V:adjoining structures of fourth ventricle
SpC:spinal cord

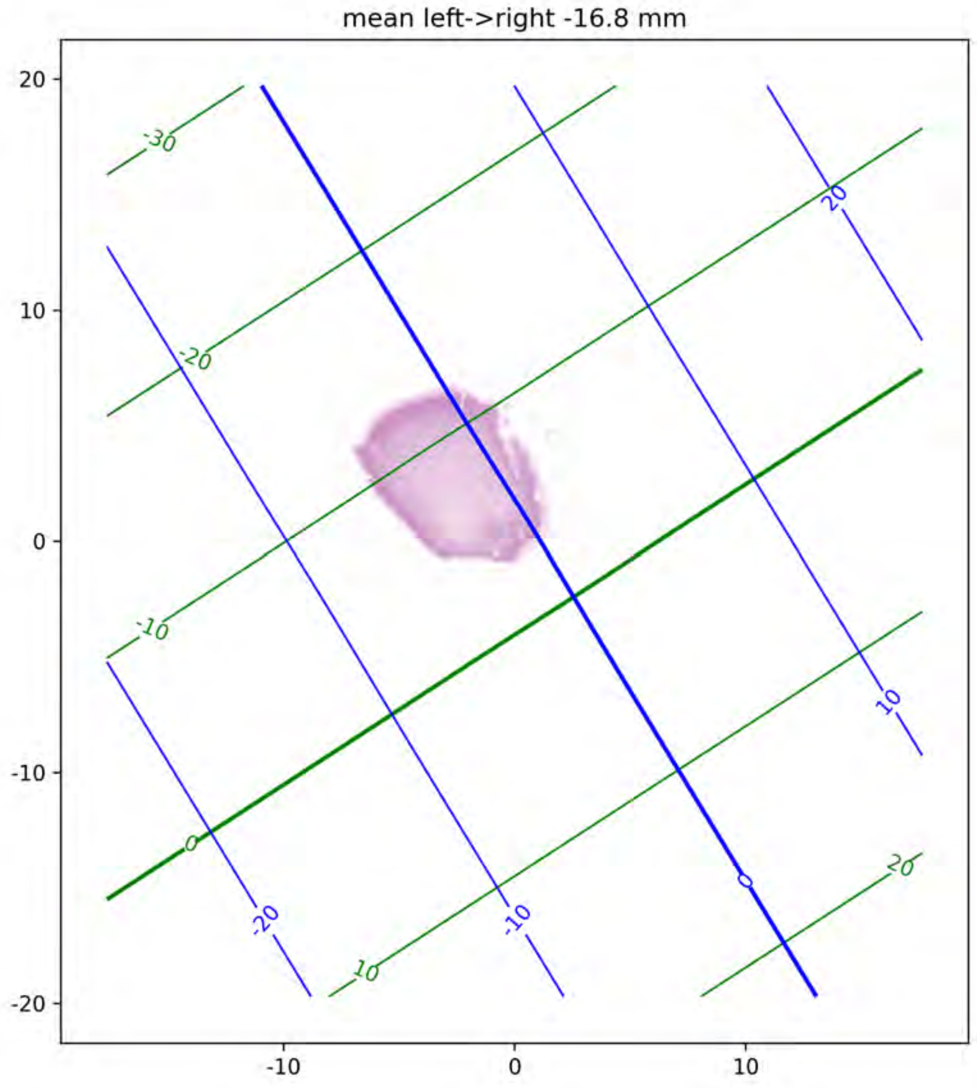

mean left->right -16.8 mm
20
10
0
-10
-20
-10
0
10
-30
-20
-10
0
10
20
20
10
0
-10
-20

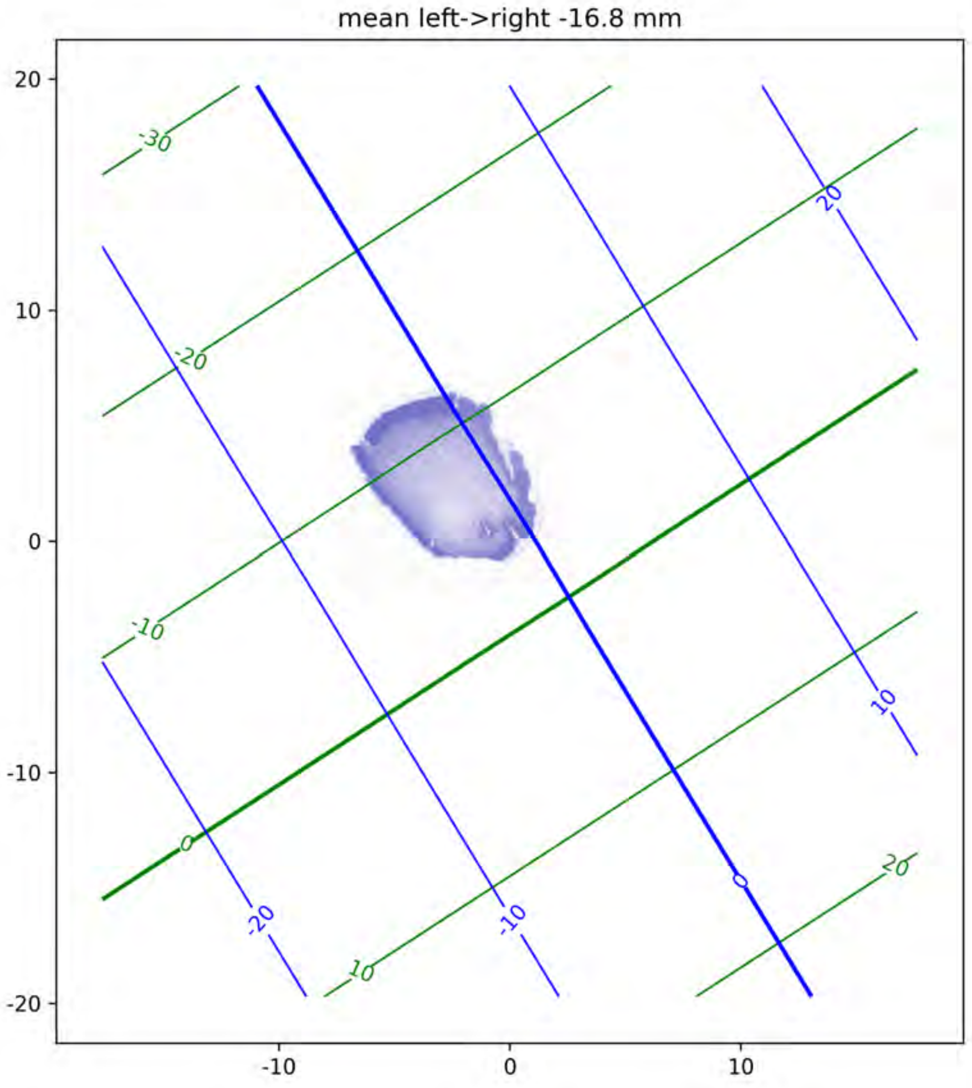
mean left->right -16.8 mm
20
10
0
-10
-20
-10
0
10
-30
-20
-10
0
10
20
20
10
0
-10
-20

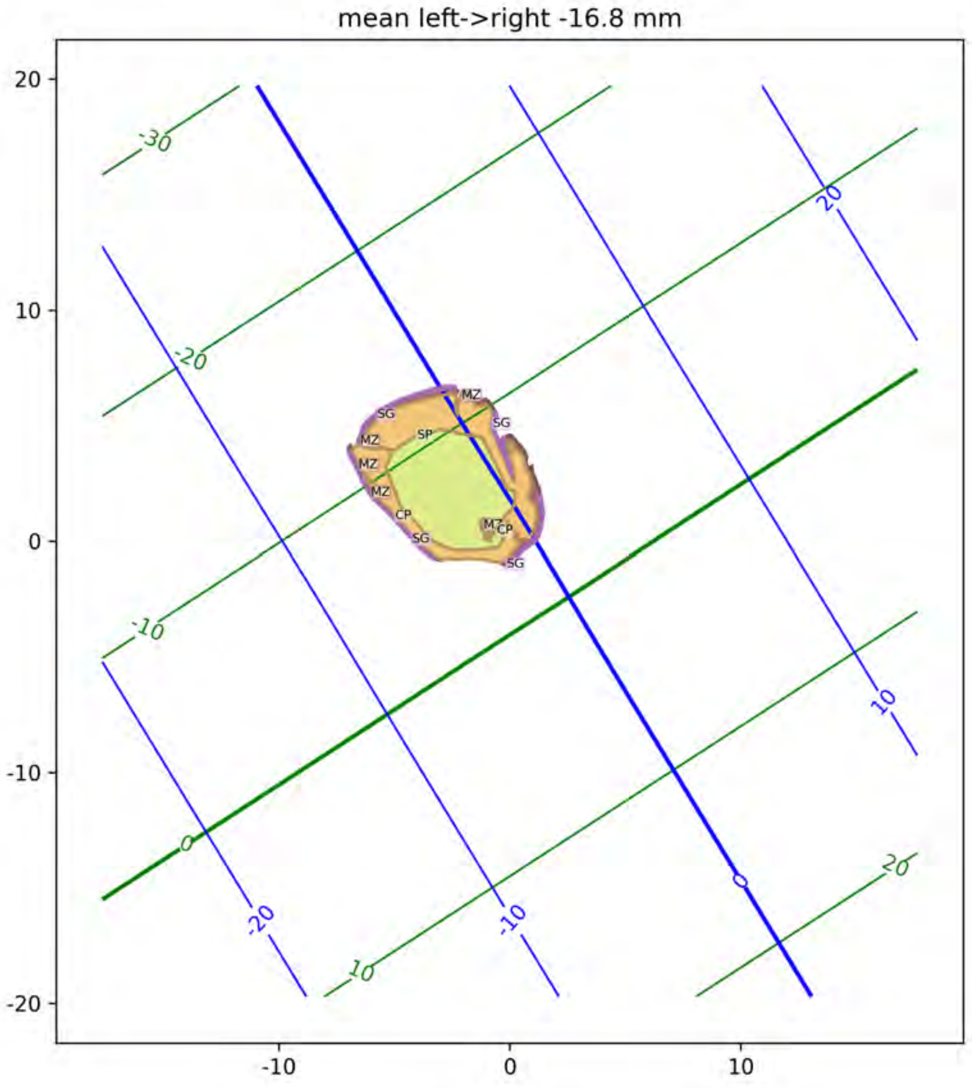
mean left->right -16.8 mm
SG
MZ
SP
MZ
MZ
CP
SG
MZ
CP
SG
MZ
SG

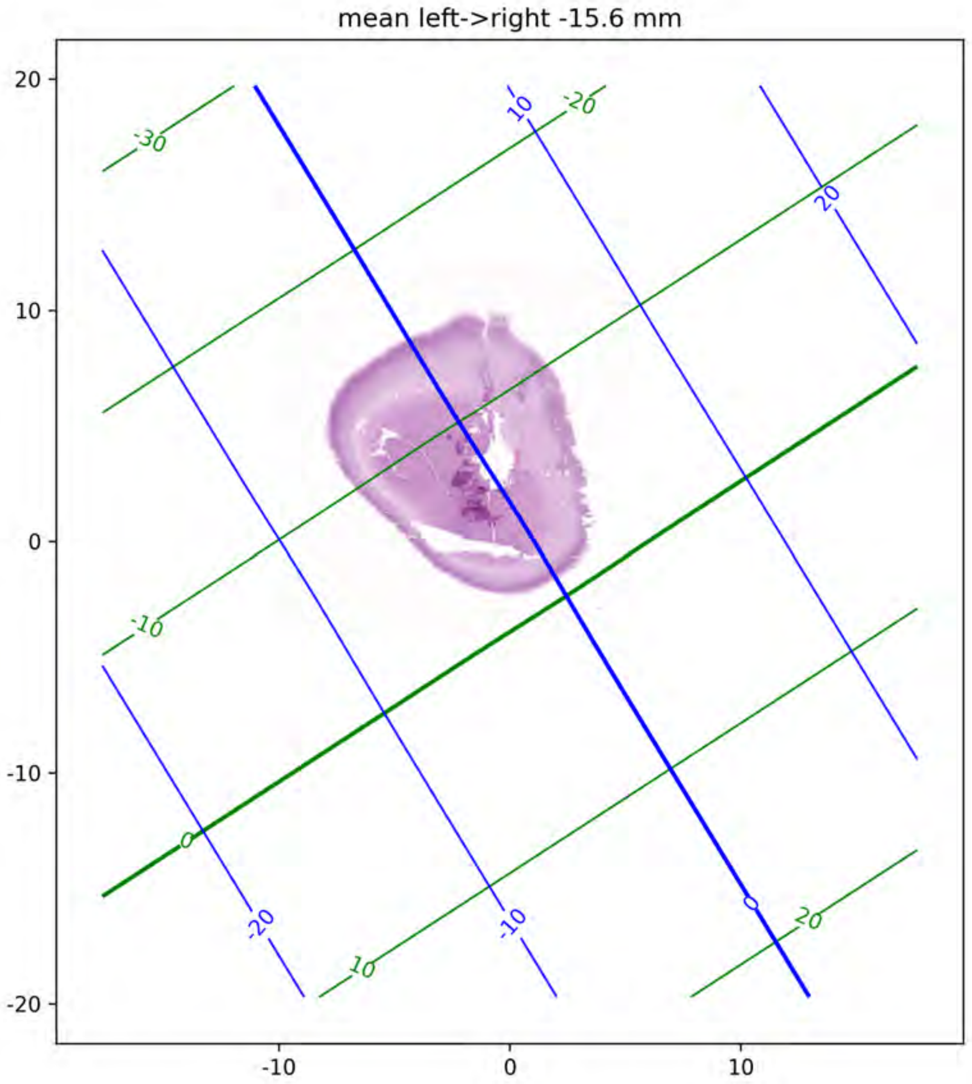
mean left->right -15.6 mm
20
10
0
-10
-20
-10
0
10
-30
-20
-10
0
10
20
10
20
-20
-10
0

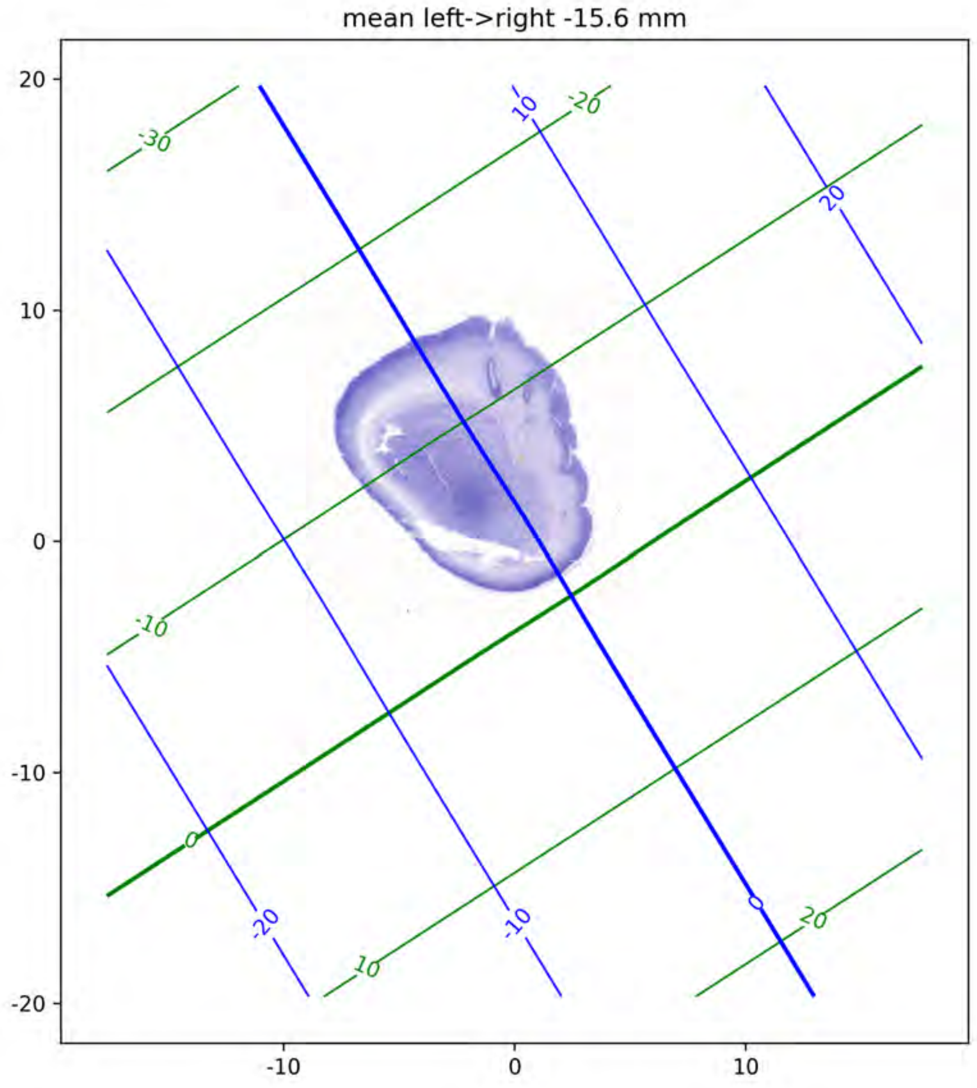
mean left->right -15.6 mm
20
10
0
-10
-20
-10
0
10
-30
-20
-10
0
10
20
10
20
-20
-10
0
20

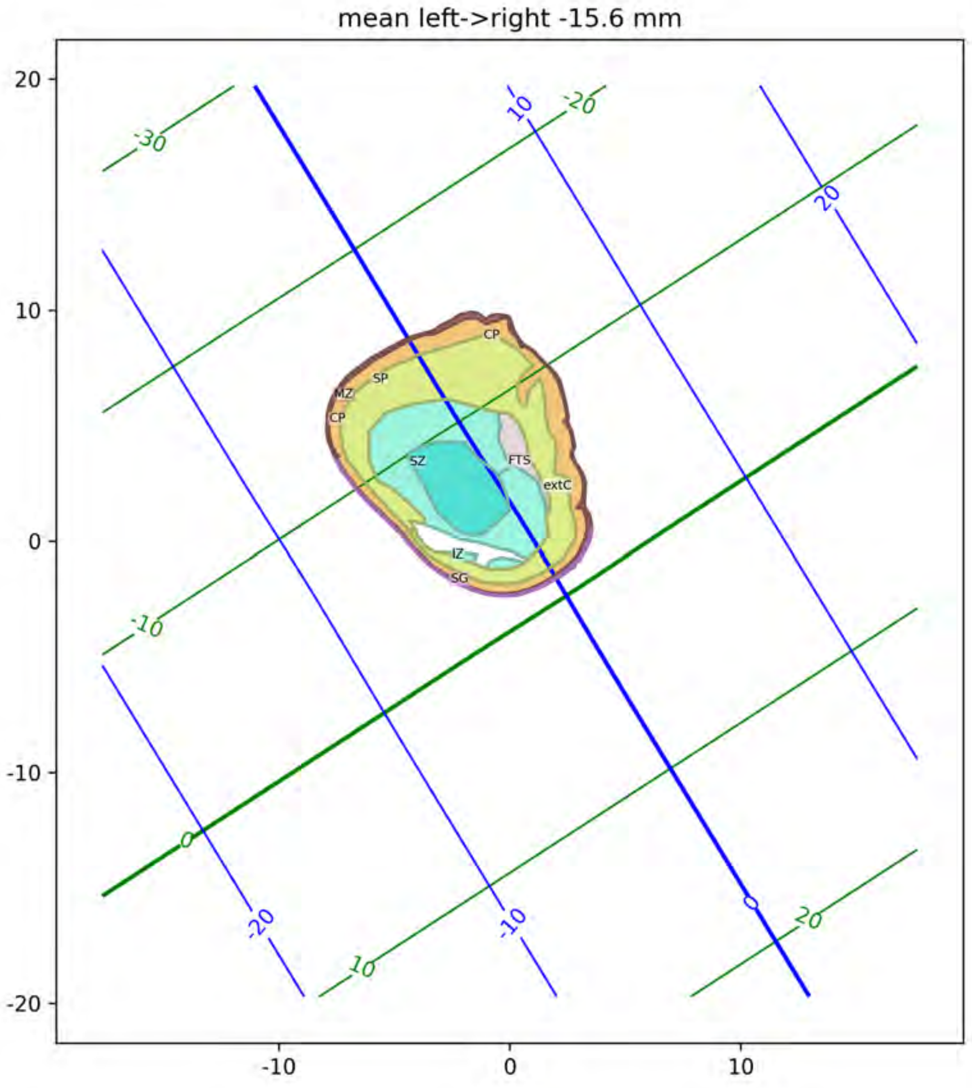
mean left->right -15.6 mm
CP
SP
MZ
CP
SZ
FTS
extC
IZ
SG
20
10
0
-10
-20
-10
0
10
-30
-20
-10
0
10
20
10
20
-20
-10
0

mean left->right -14.8 mm

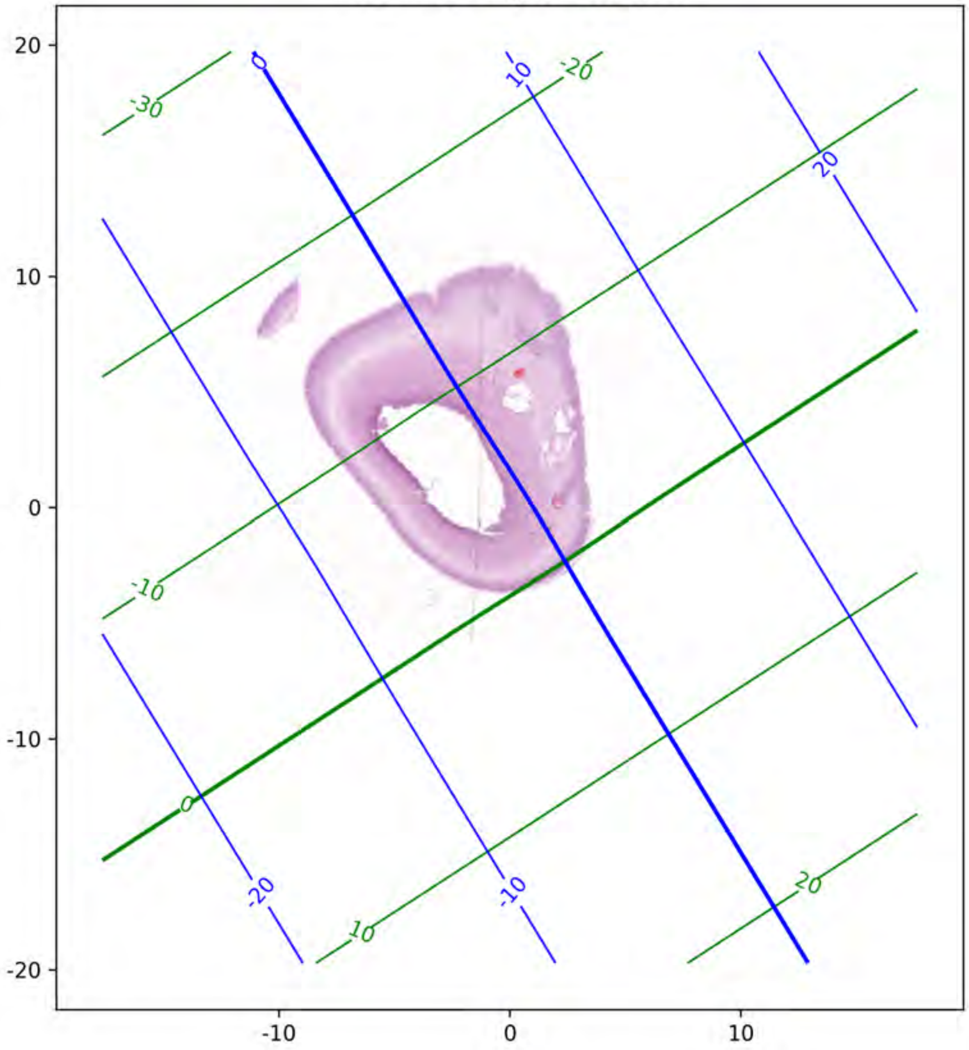

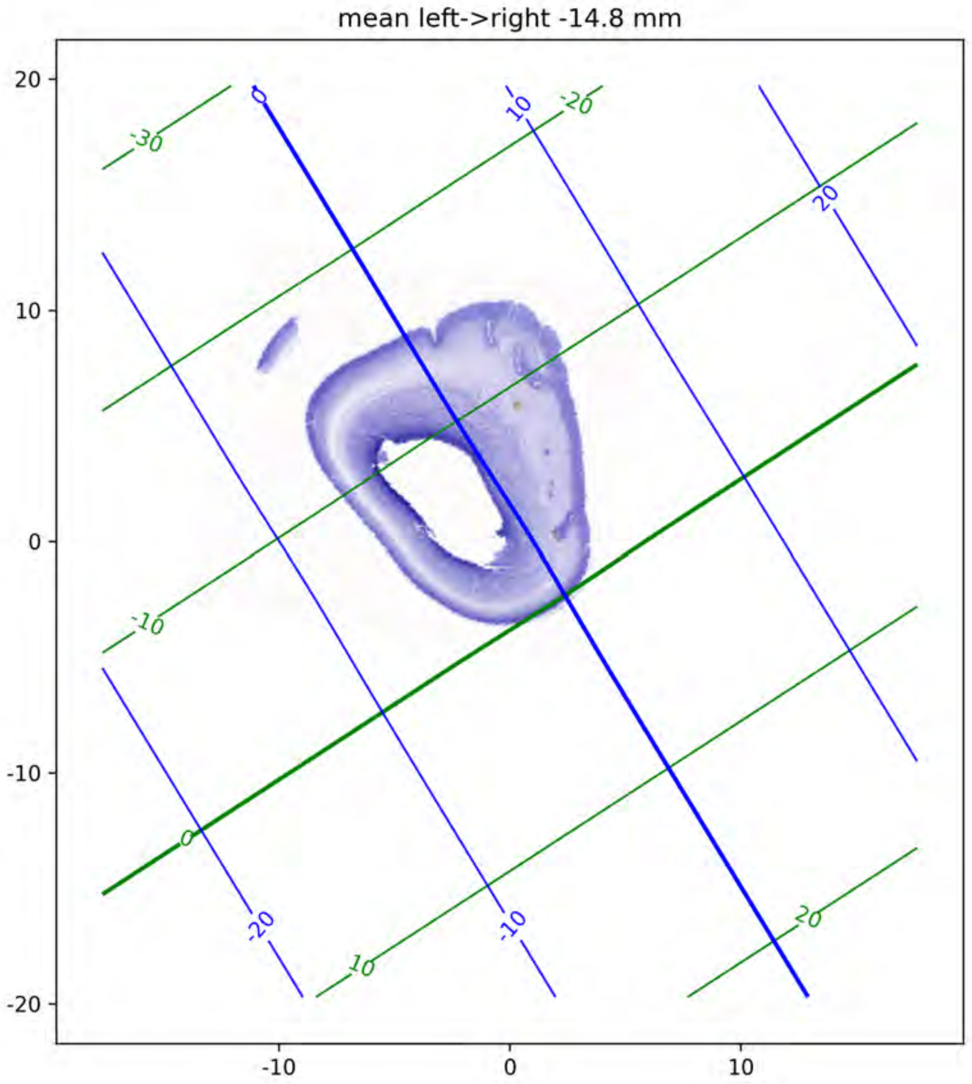

mean left->right -14.8 mm
20
10
0
-10
-20
-10
0
10
-30
-20
-10
0
10
20
0
10
20
-20
-10

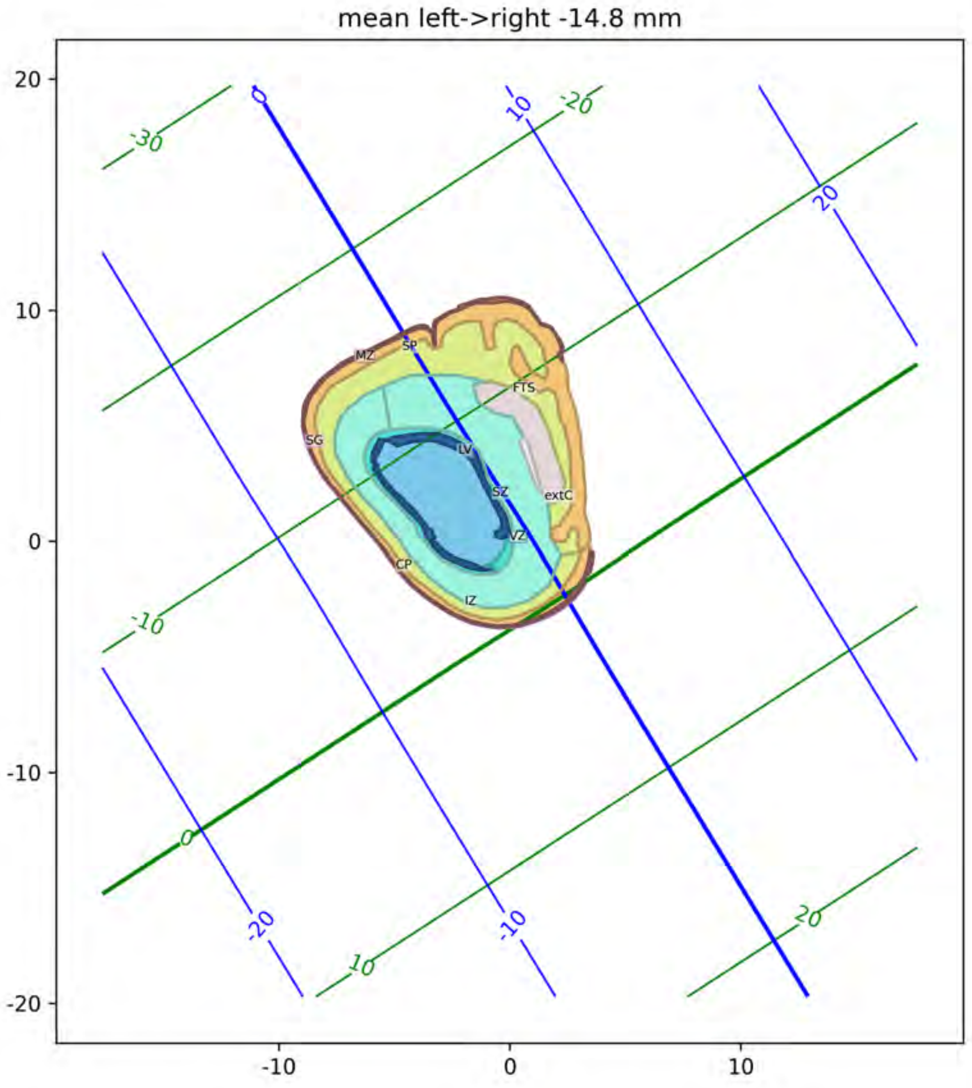

mean left->right -14.8 mm
MZ
SP
FTS
SG
LV
SZ
extC
VZ
CP
IZ
-30
-20
-10
0
10
20
-10
-20
0
10
20

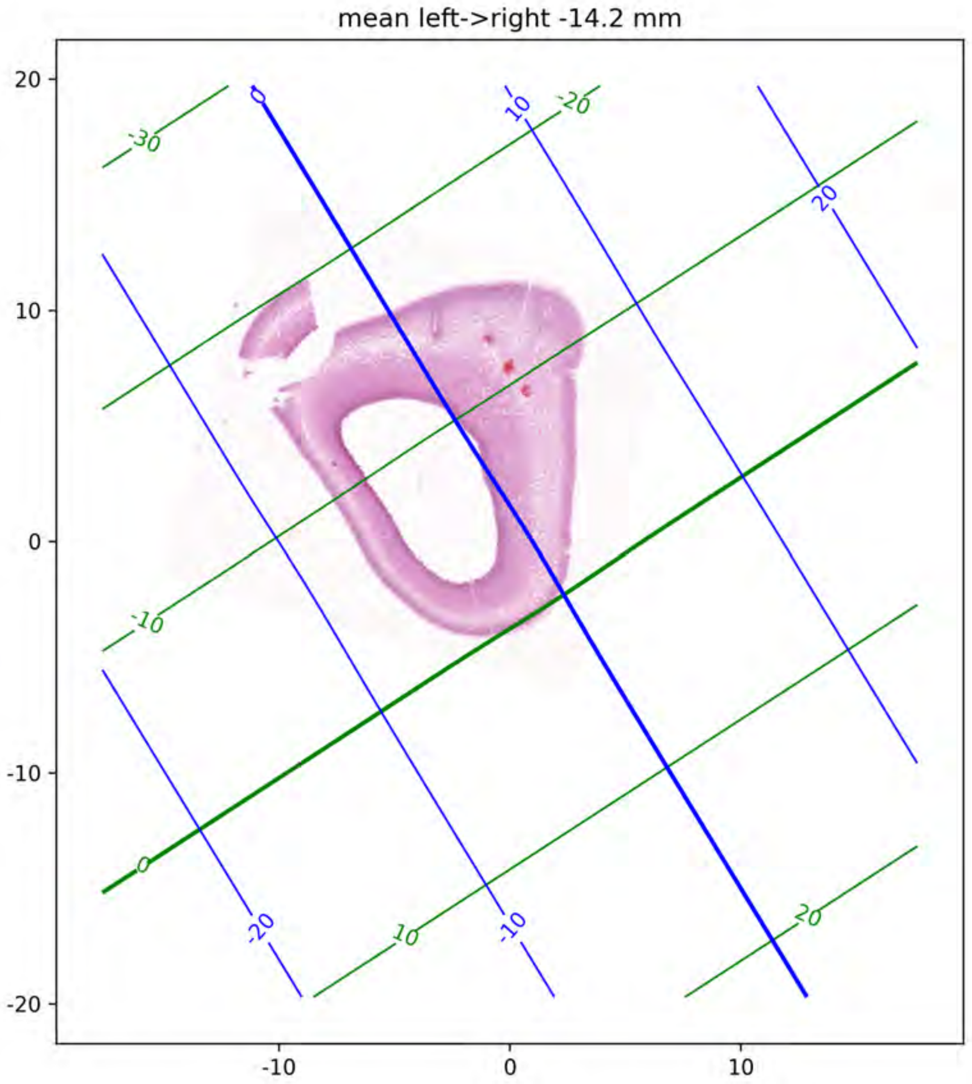
mean left->right -14.2 mm
20
10
0
-10
-20
-10
0
10
-30
-20
-10
0
10
20
0
10
20
-20
-10

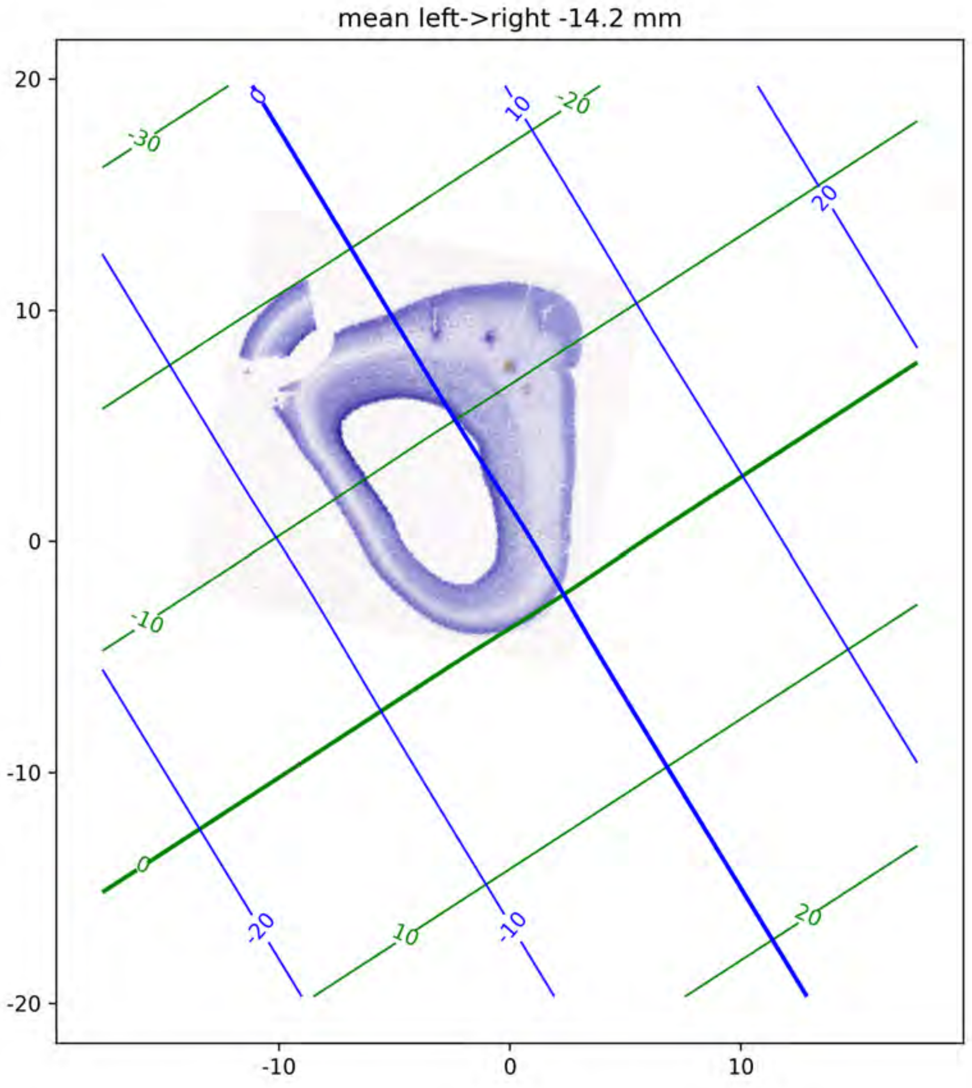

mean left->right -14.2 mm
20
10
0
-10
-20
-10
0
10
-30
-20
-10
0
10
20
0
10
20
-20
-10

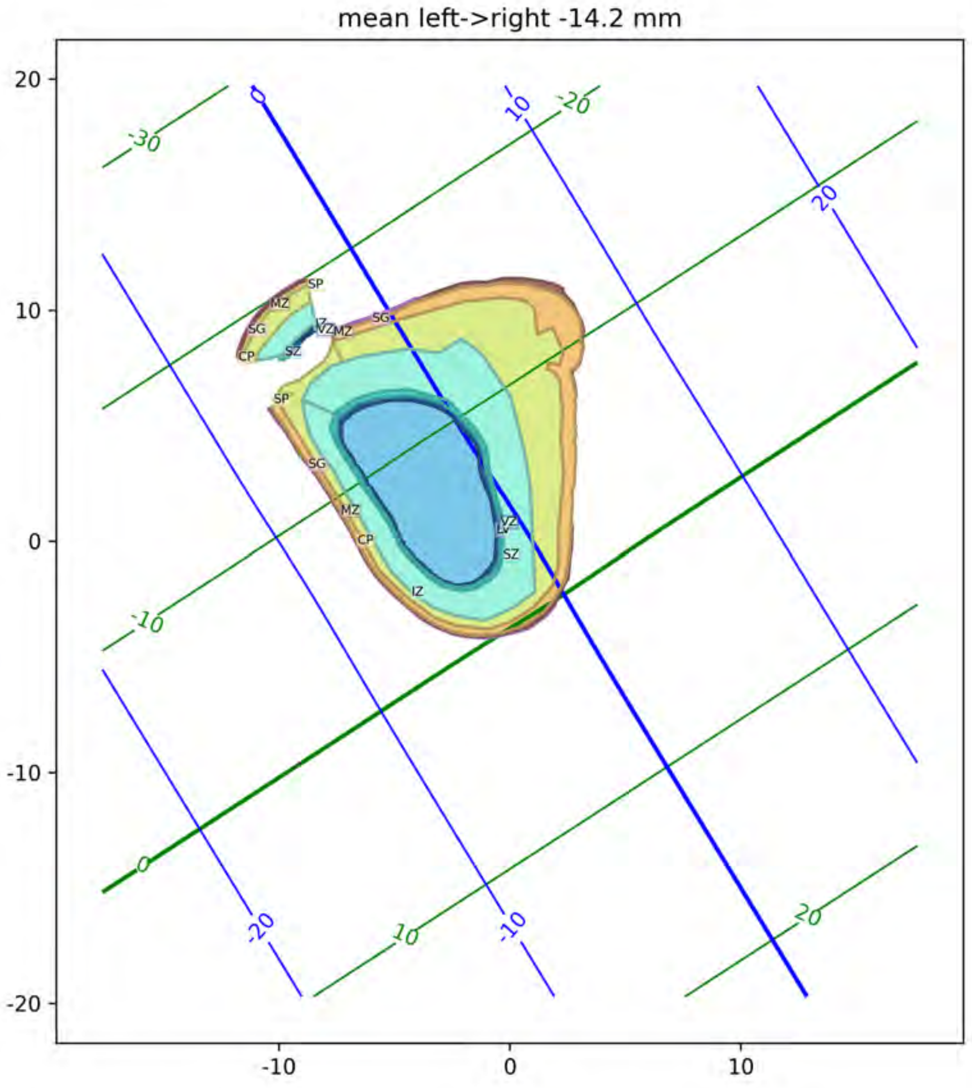
mean left->right -14.2 mm
20
10
0
-10
-20
-10
0
10
-30
-20
-10
0
10
20
0
10
20
-20
-10
SP
MZ
SG
CP
SZ
VZ
MZ
SG
SP
SG
MZ
CP
IZ
VZ
LV
SZ

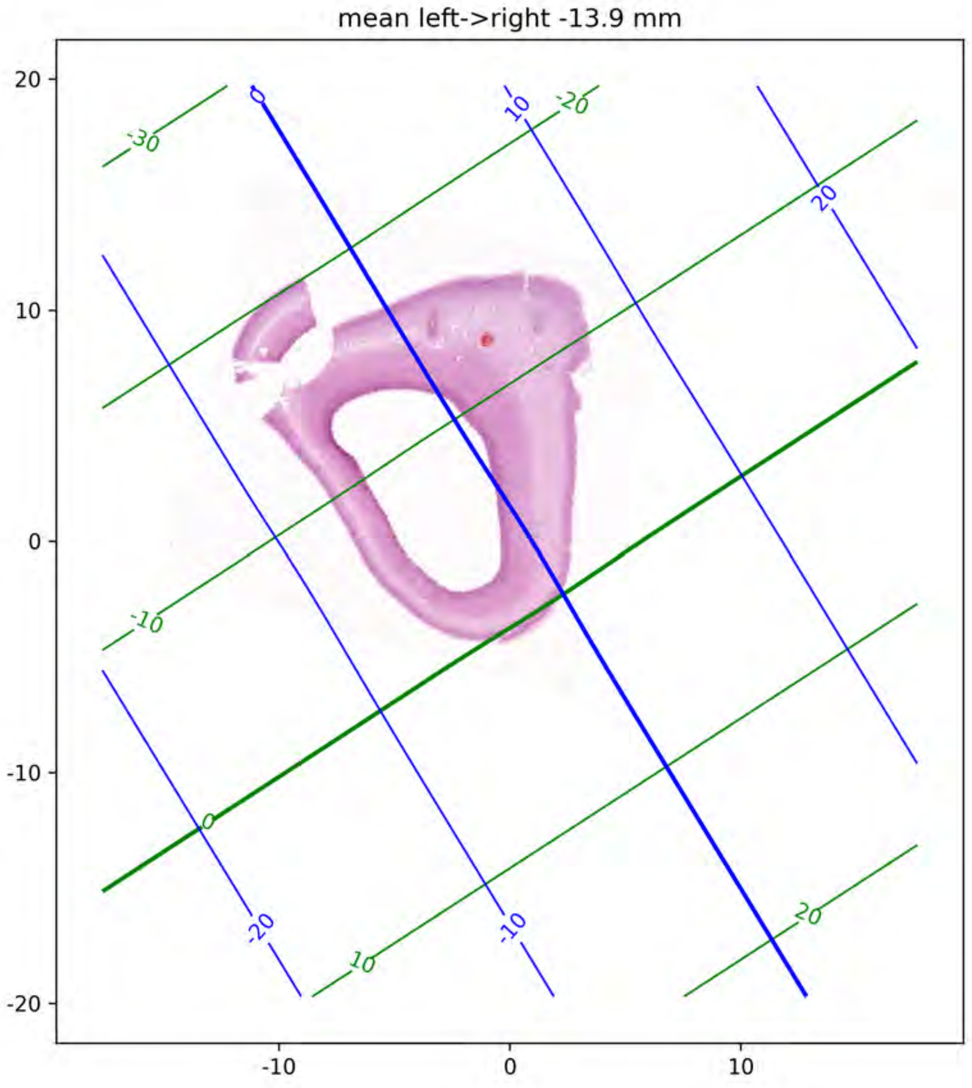
mean left->right -13.9 mm
20
10
0
-10
-20
-10
0
10
-30
-20
-10
0
10
20
0
10
20
-20
-10

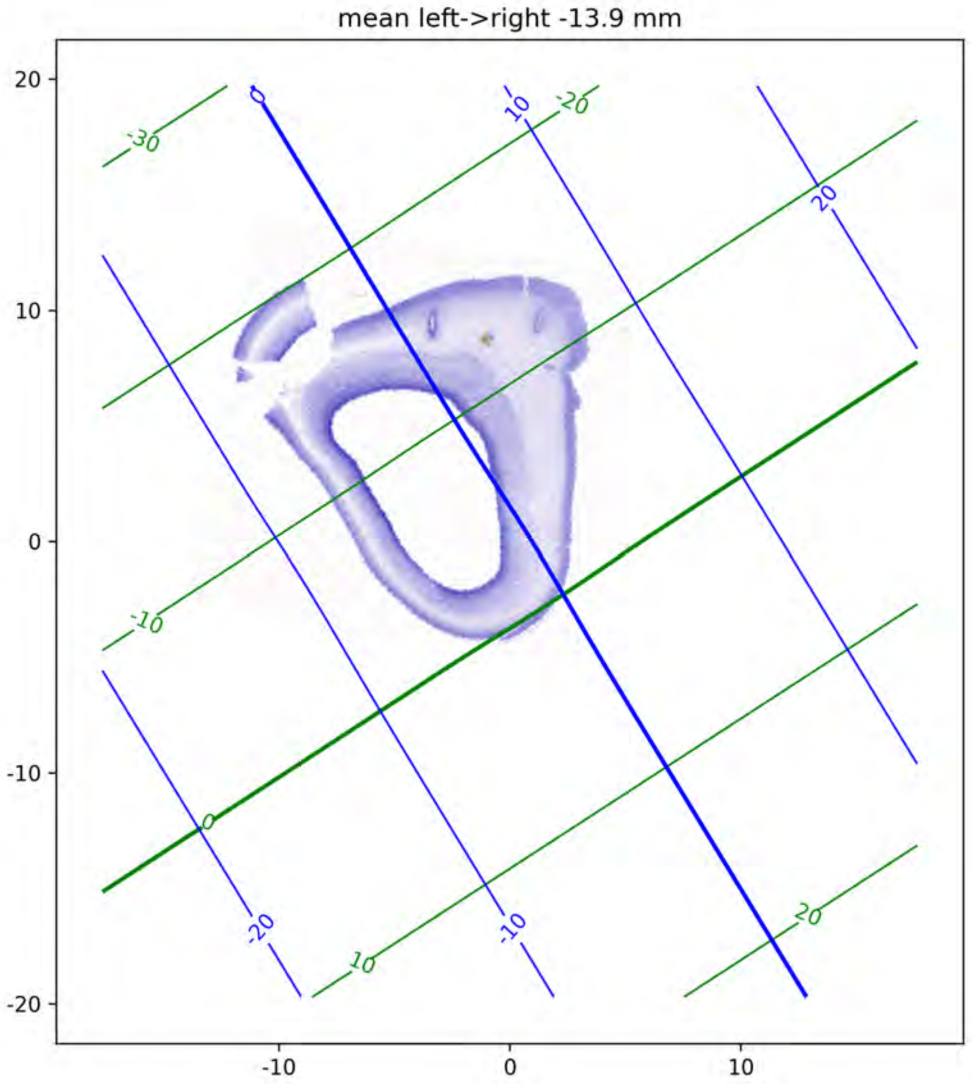

mean left->right -13.9 mm

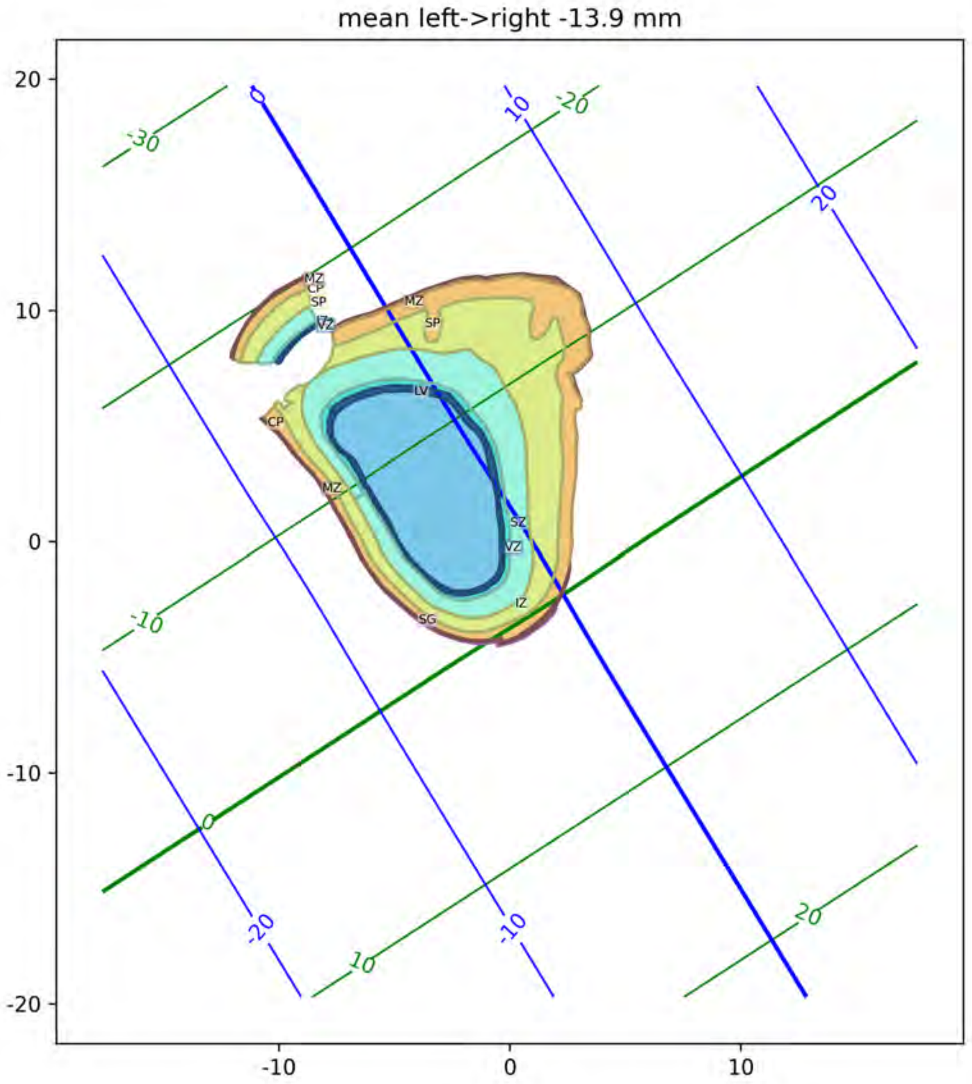
mean left->right -13.9 mm
20
10
0
-10
-20
-10
0
10
-30
-20
-10
0
10
20
0
10
20
-20
-10
MZ
CP
SP
VZ
MZ
SP
LV
CP
MZ
SZ
VZ
IZ
SG

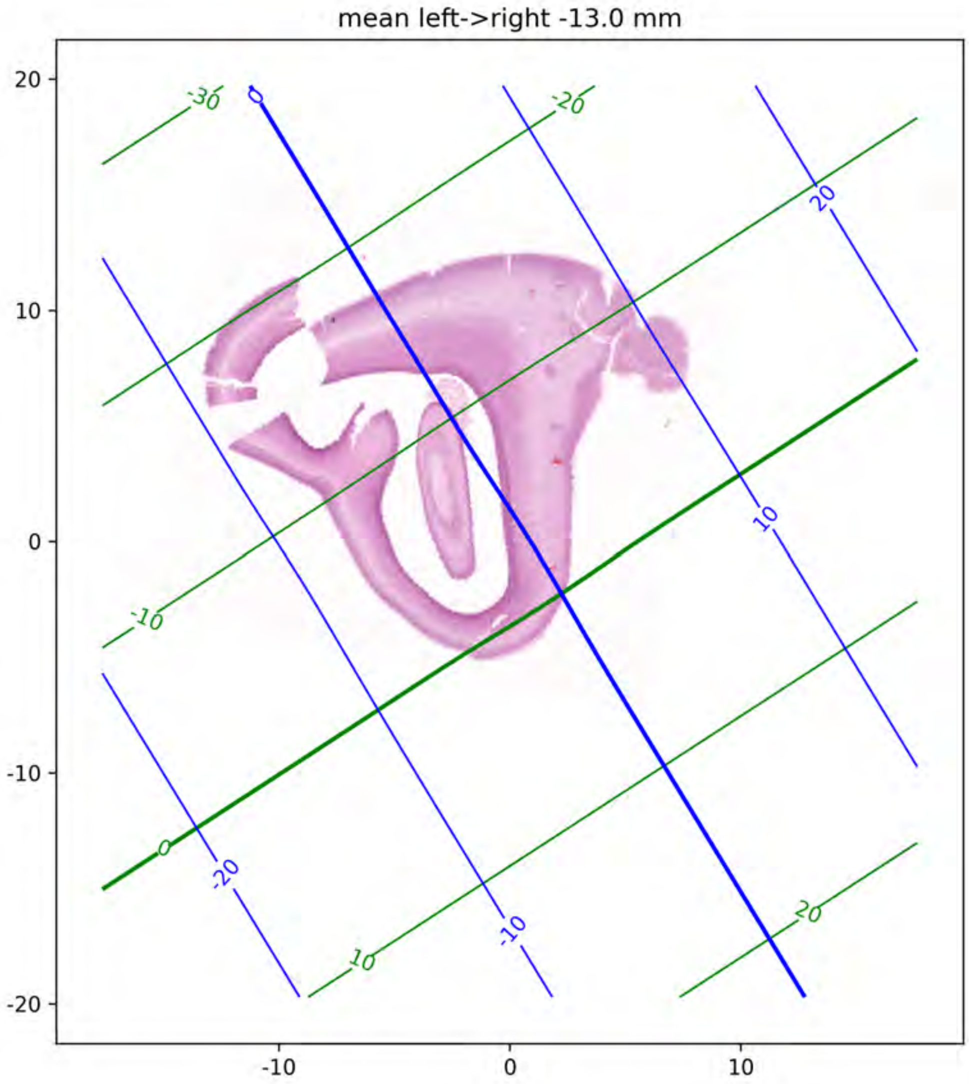
mean left->right -13.0 mm
20
10
0
-10
-20
-10
0
10
-30
-20
-10
0
10
20
0
20
10
-20
-10

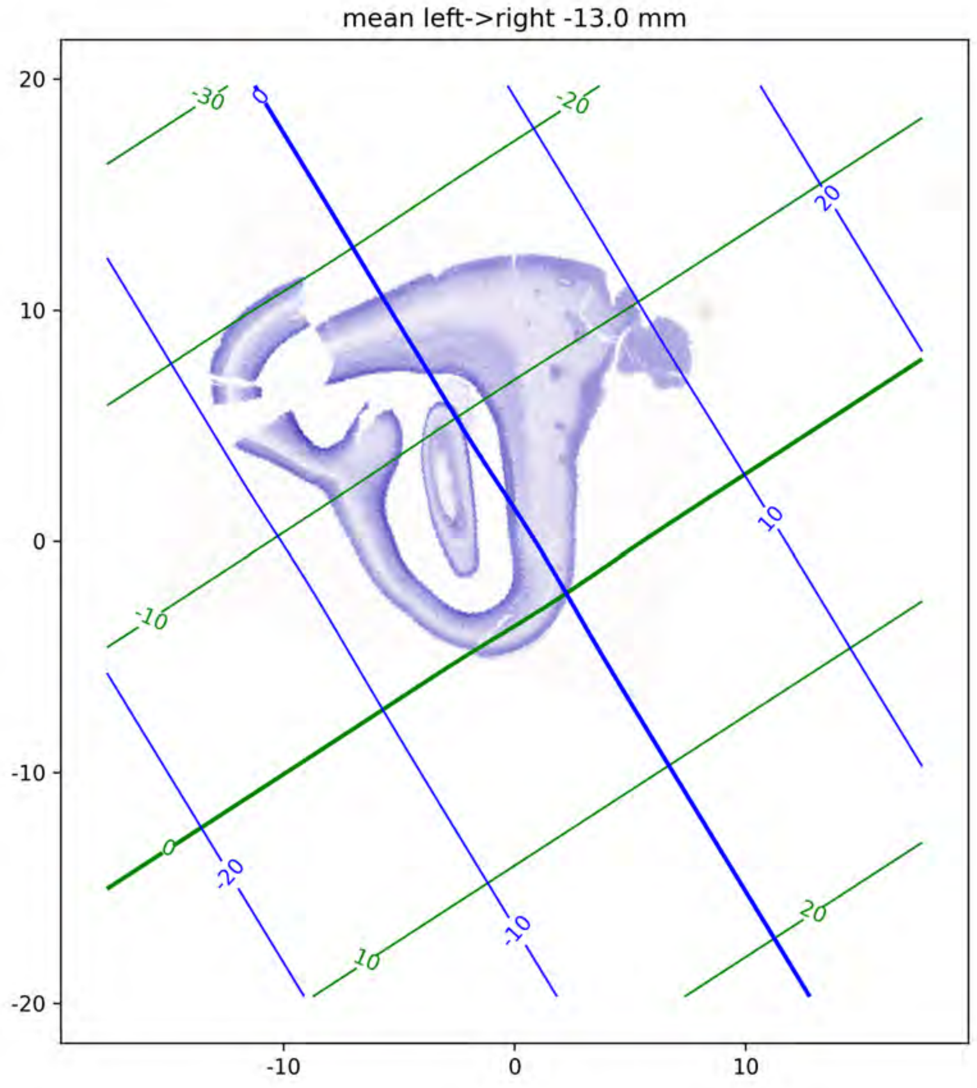
mean left->right -13.0 mm
20
10
0
-10
-20
-10
0
10
-30
-20
-10
0
10
20
0
20
10
-20
-10

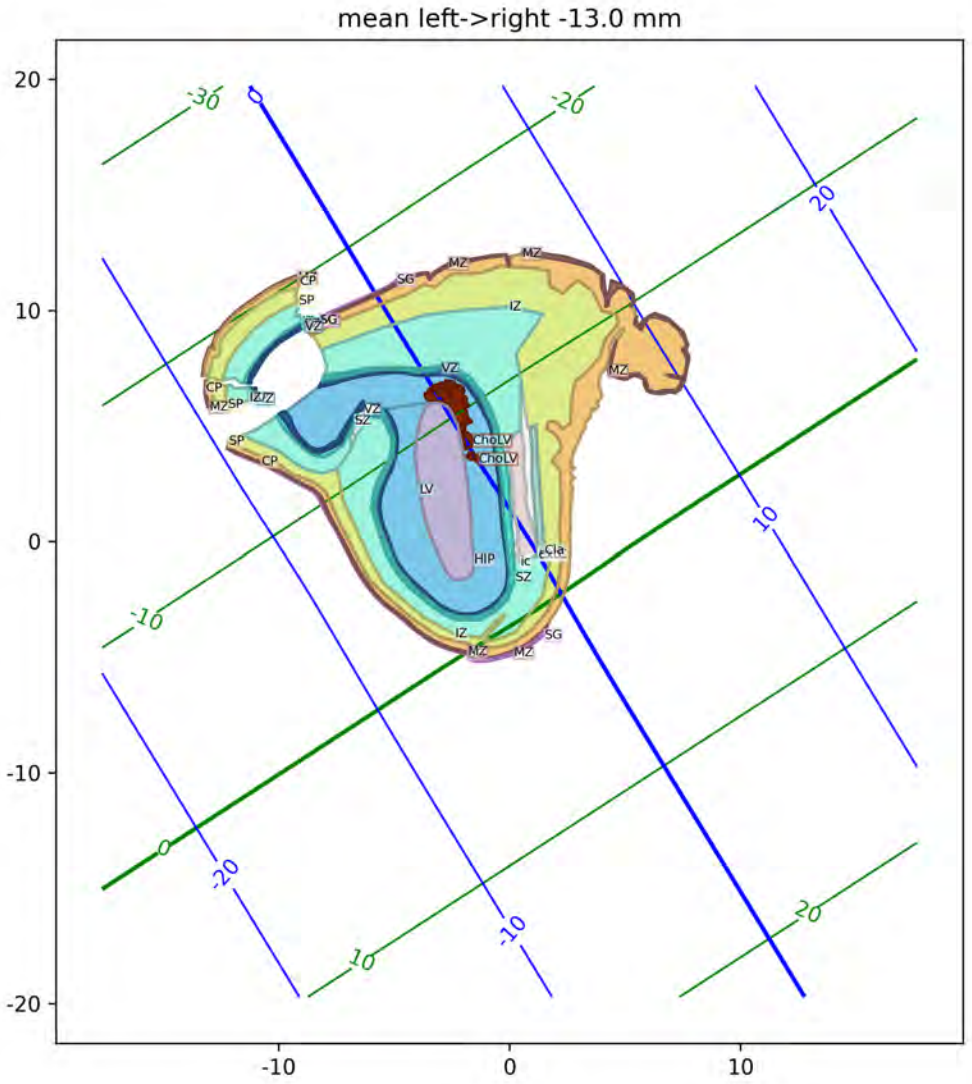
mean left->right -13.0 mm
MZ
SG
CP
SP
IZ
VZ
LV
HIP
ChoLV
SZ
Cla
IC

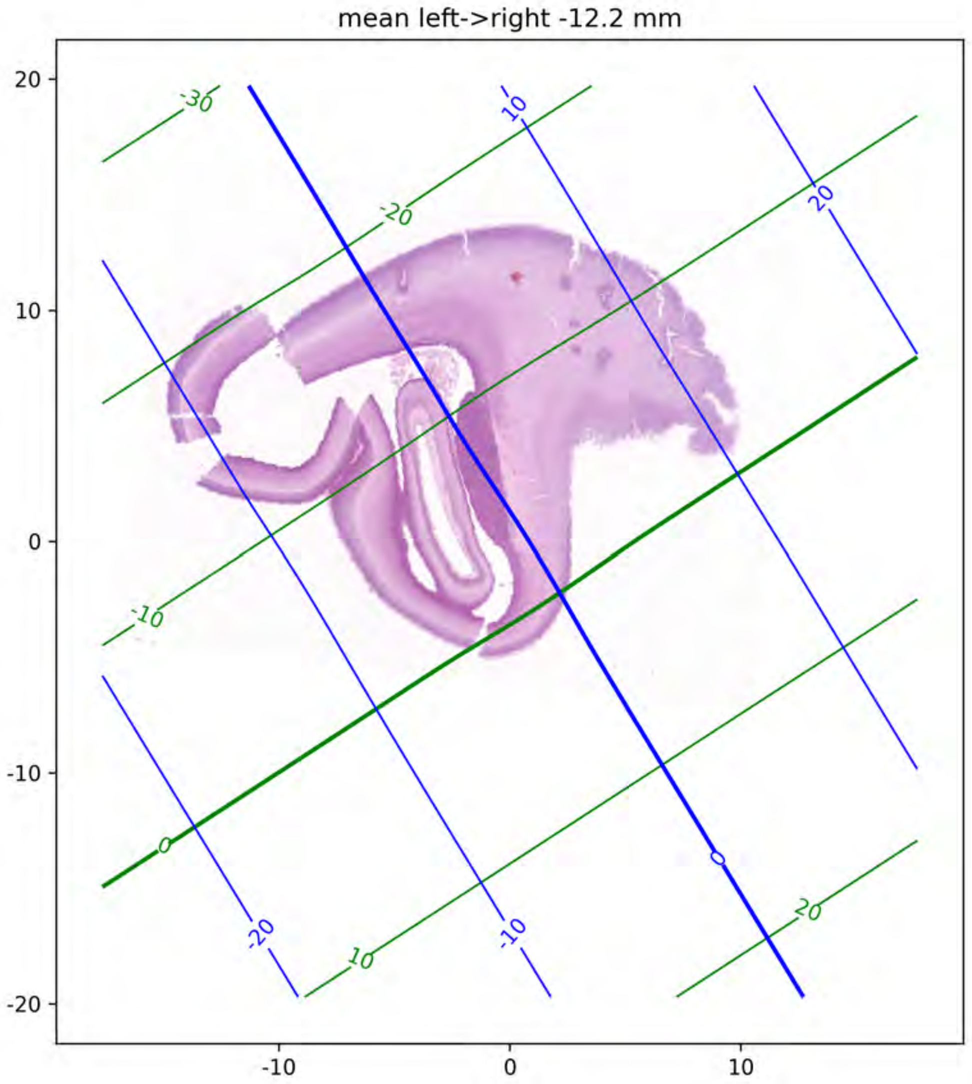

mean left->right -12.2 mm
20
10
0
-10
-20
-10
0
10
-30
-20
-10
0
10
20
10
20
-20
-10

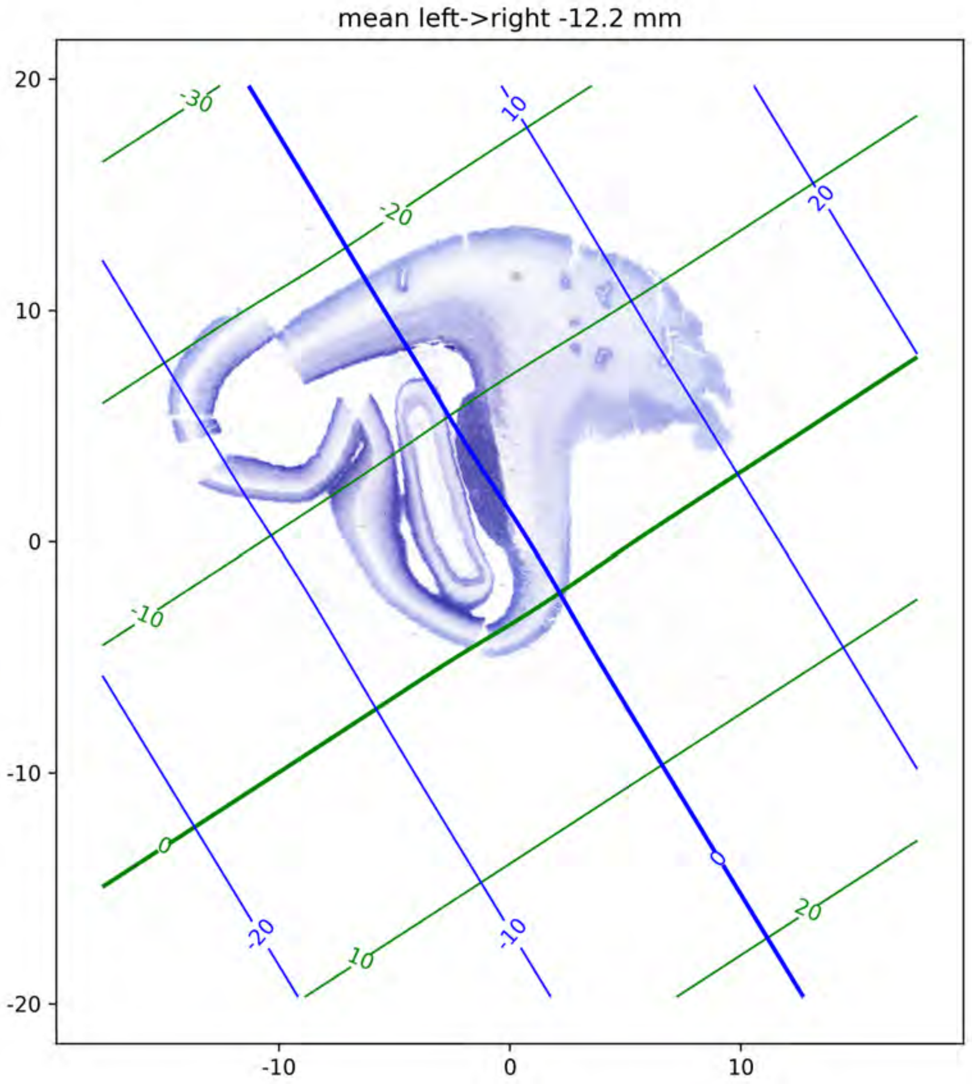
mean left->right -12.2 mm
20
10
0
-10
-20
-10
0
10
-30
-20
-10
0
10
20
10
20
-20
-10
0

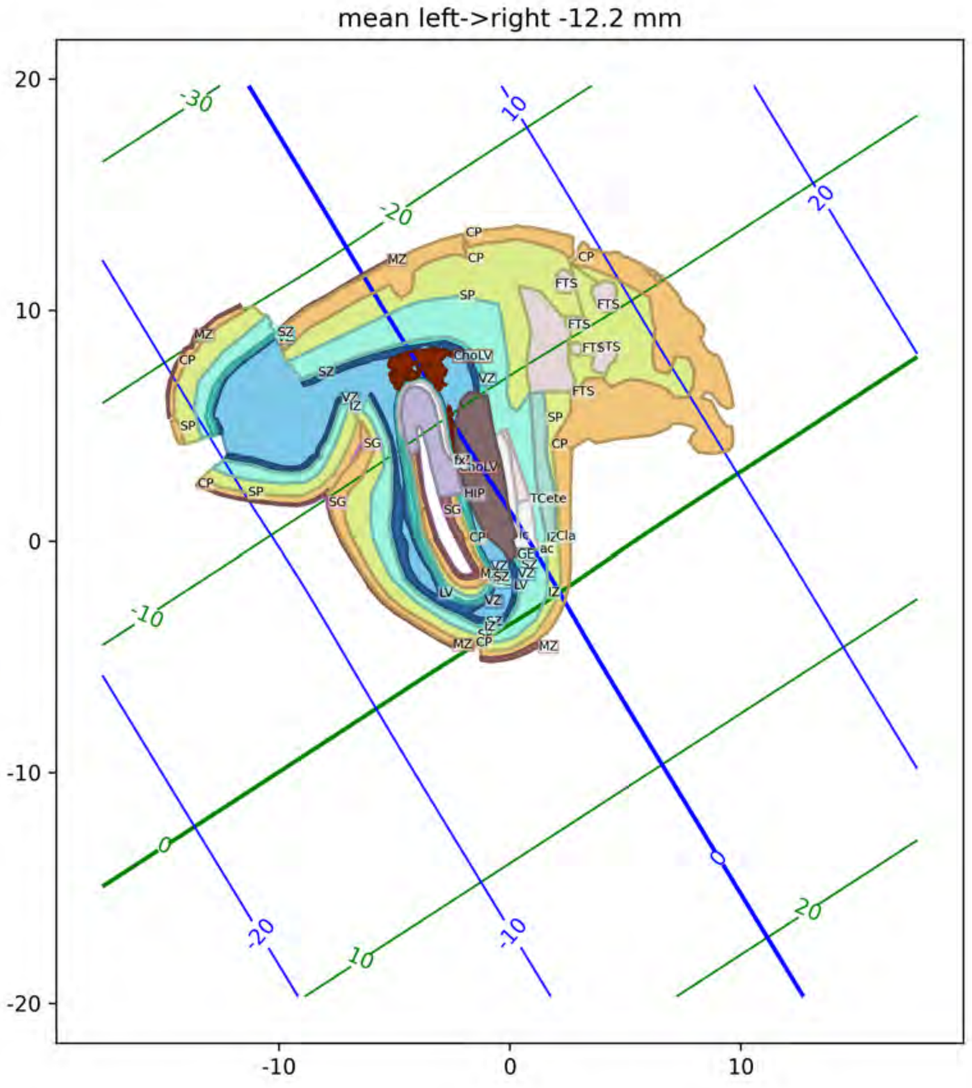
mean left->right -12.2 mm
MZ
CP
SP
SZ
VZ
IZ
SG
FTS
ChoLV
HIP
TCete
Cla
GE
ac
LV
fx

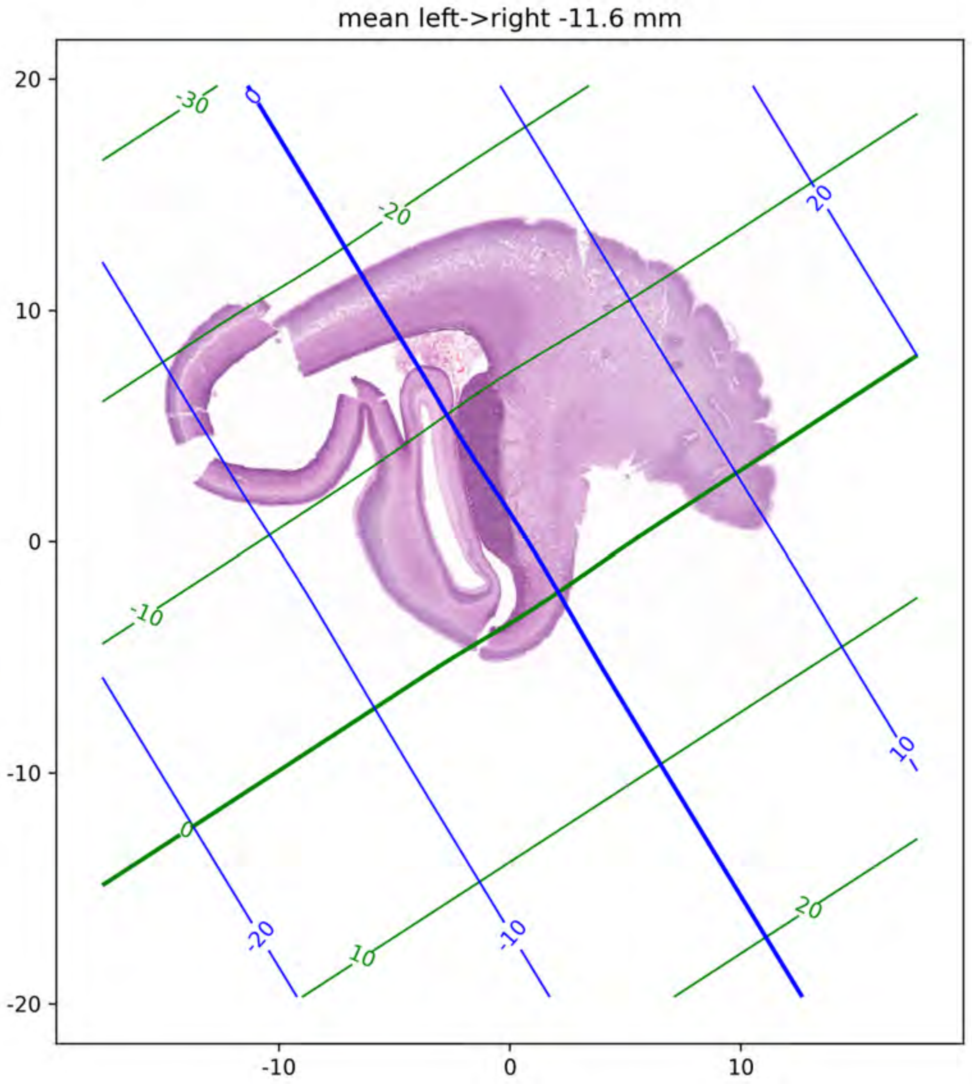
mean left->right -11.6 mm
20
10
0
-10
-20
-10
0
10
-30
-20
-10
0
10
20
0
20
10
-20
-10
20

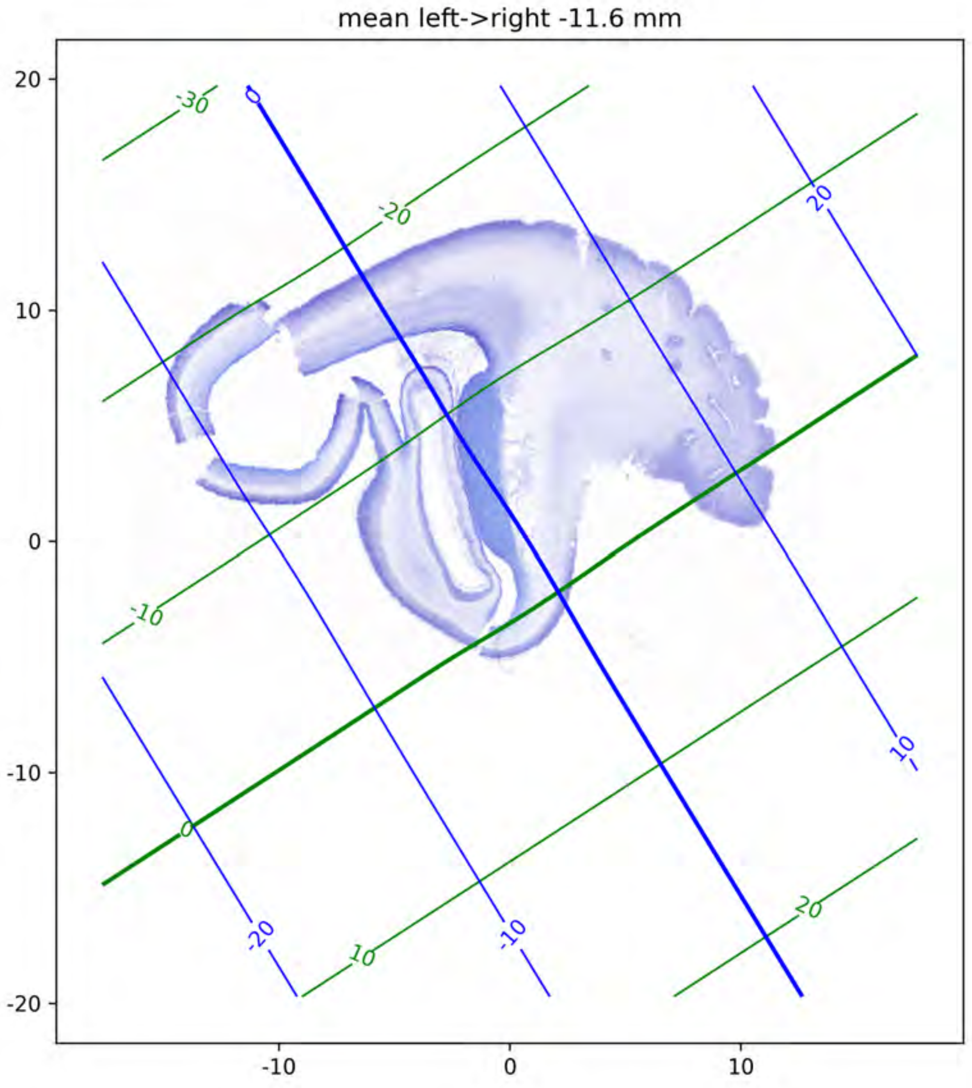
mean left->right -11.6 mm
20
10
0
-10
-20
-10
0
10
-30
-20
-10
0
10
20
0
20
10
-20
-10

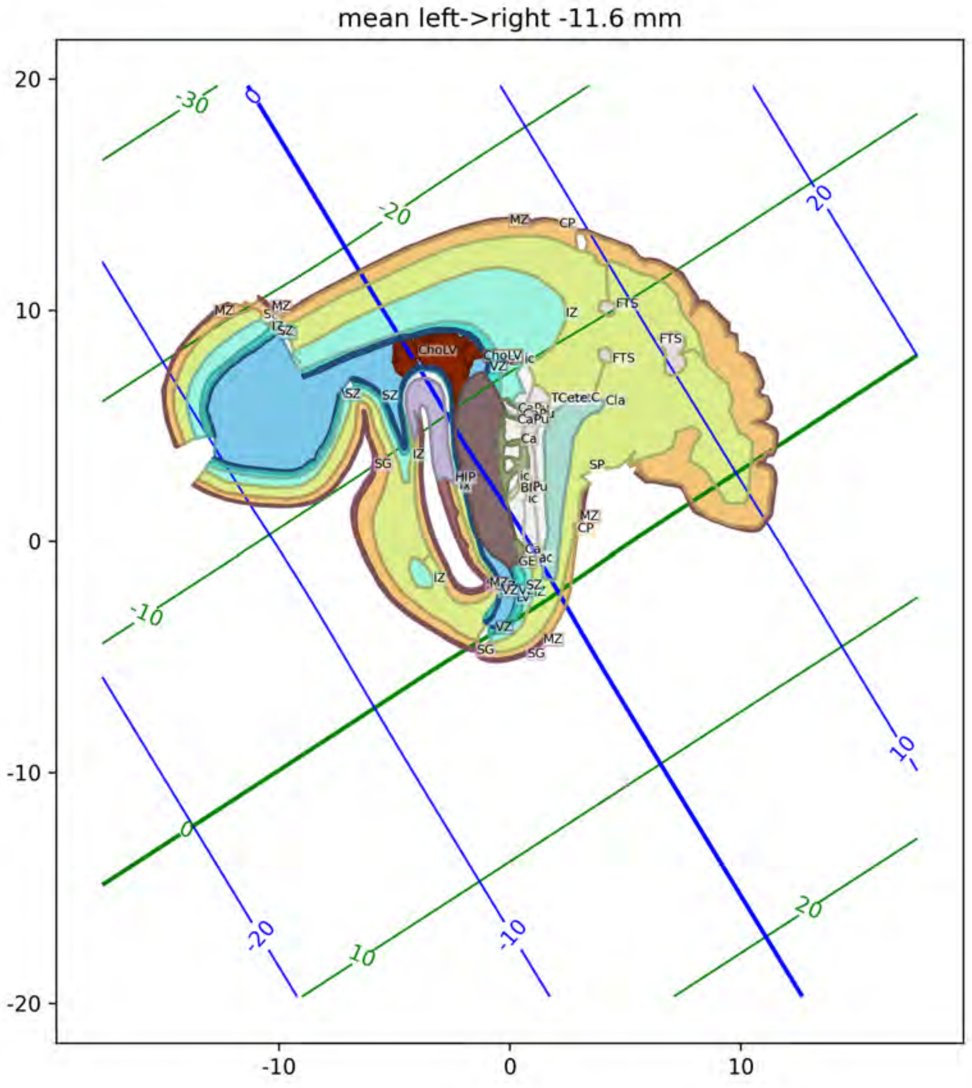
mean left->right -11.6 mm
20
10
0
-10
-20
-10
0
10
-30
-20
-10
0
10
20
MZ
CP
IZ
FTS
FTS
ChoLV
VZ
SZ
SG
HIP
Cla
SP
Ca
GE
ac

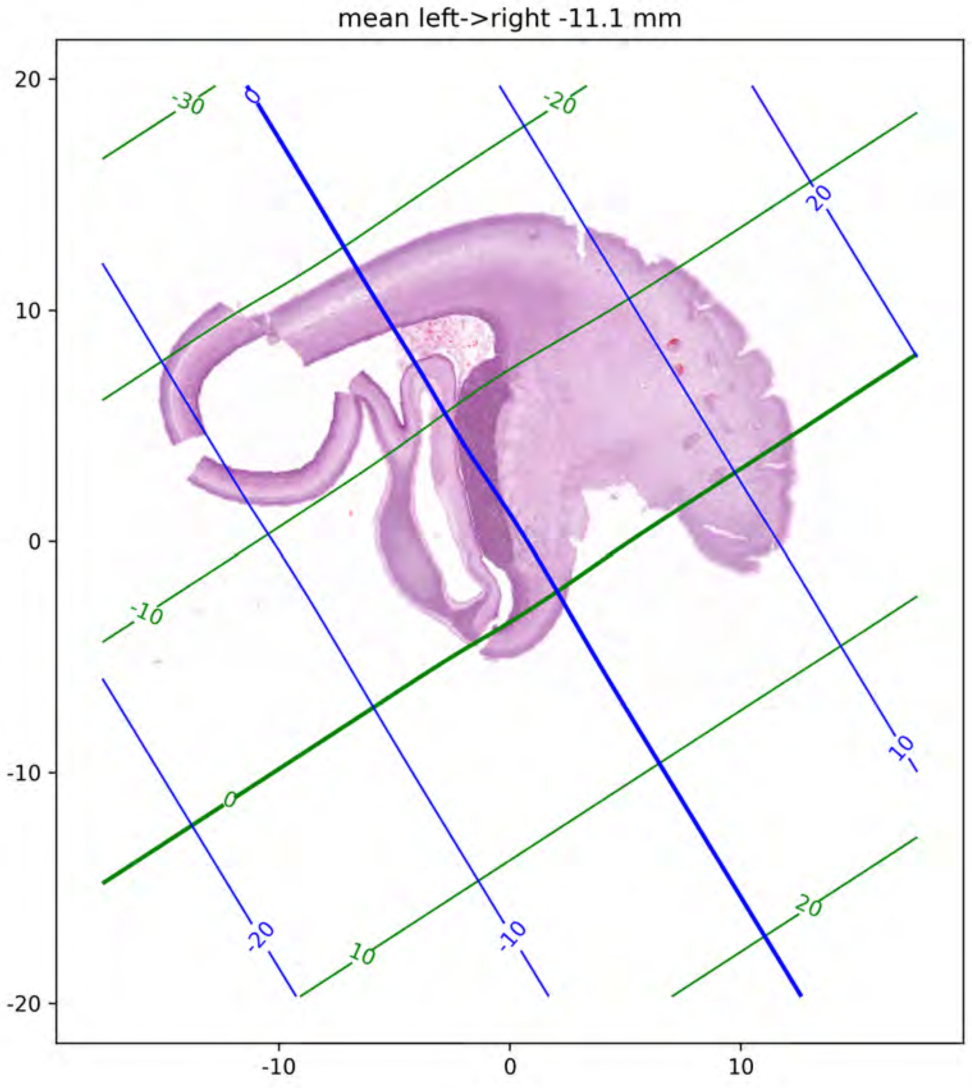
mean left->right -11.1 mm
20
10
0
-10
-20
-10
0
10
-30
-20
-10
0
10
20
0
20
10
-20
-10

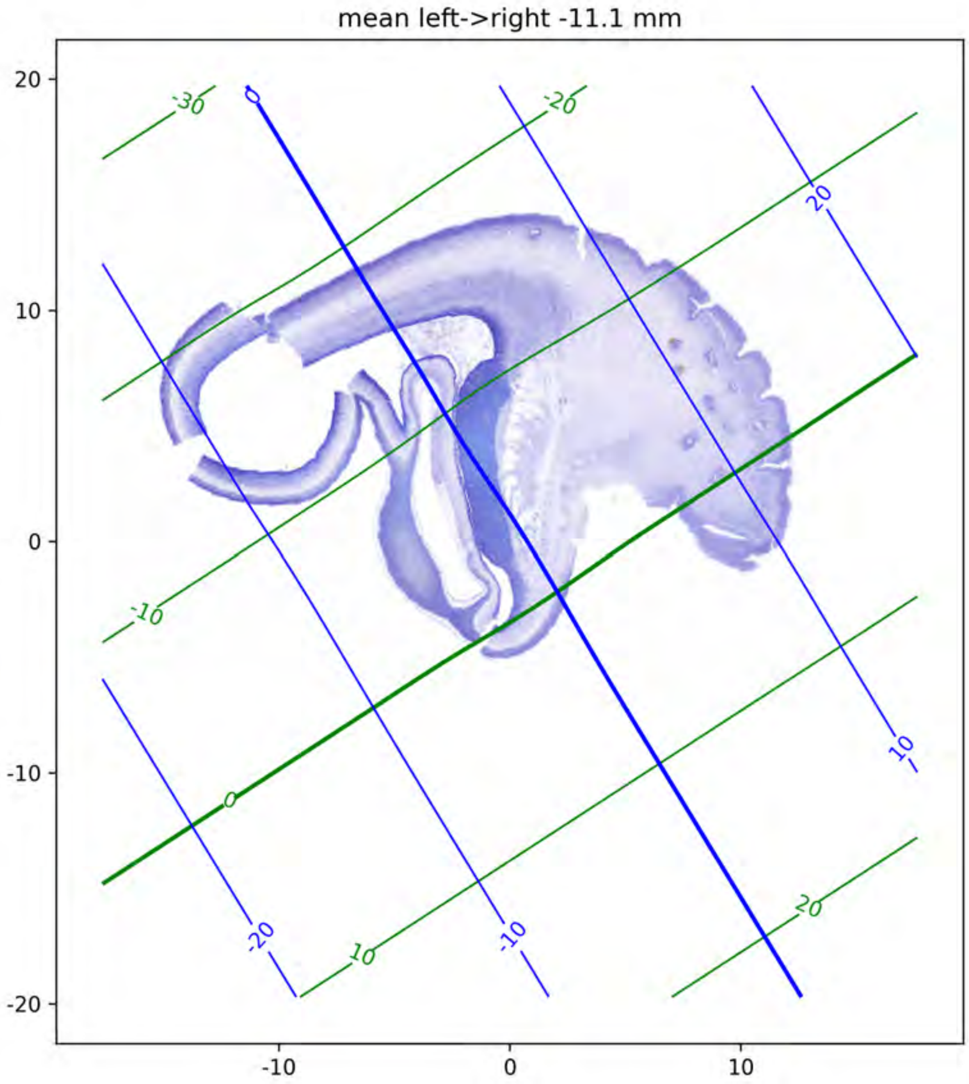
mean left->right -11.1 mm
20
10
0
-10
-20
-10
0
10
-30
-20
-10
0
10
20
0
20
10
-20
-10

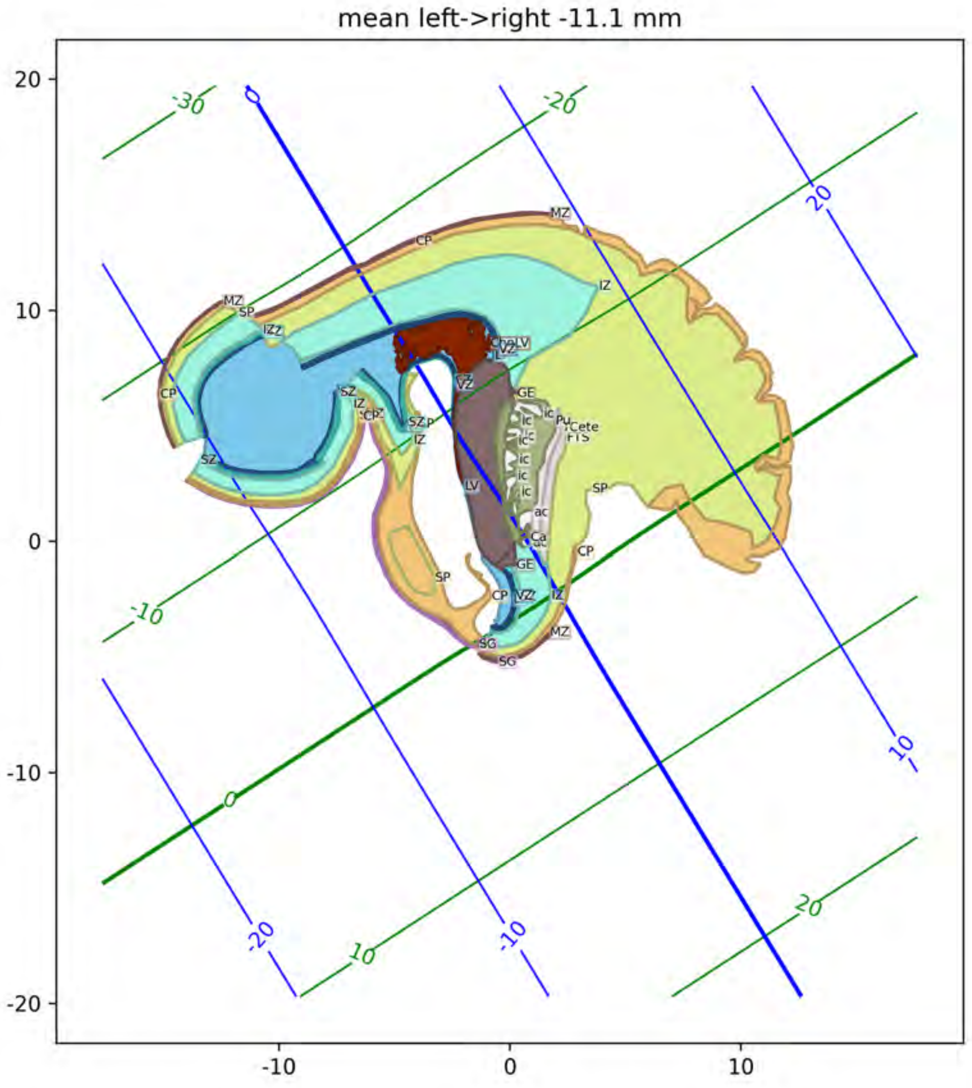
mean left->right -11.1 mm
20
10
0
-10
-20
-10
0
10
-30
-20
-10
0
10
20
MZ
CP
IZ
SP
SZ
VZ
LV
GE
ic
ac
Ca
Pu
SG

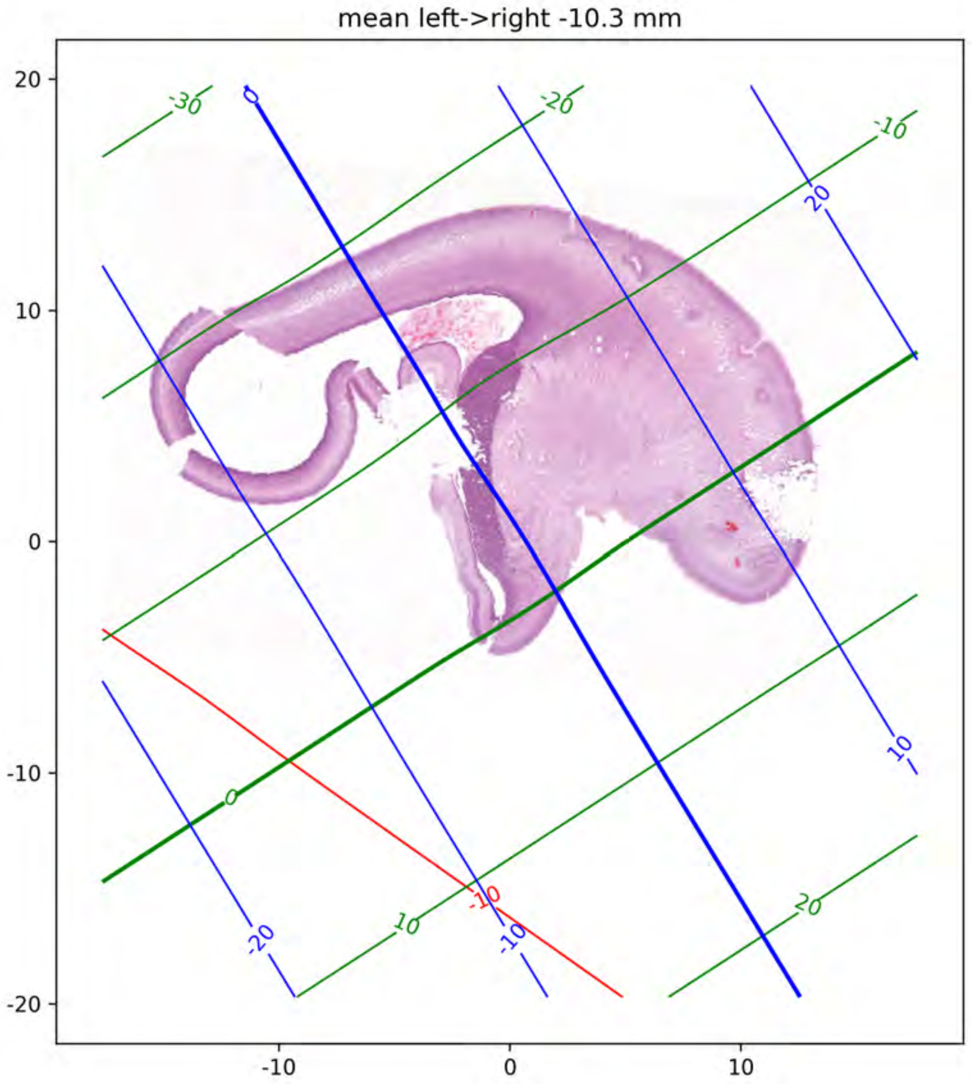
mean left->right -10.3 mm
20
10
0
-10
-20
-10
0
10
-30
-20
-10
0
10
20
0
20
10
-20
-10
-10

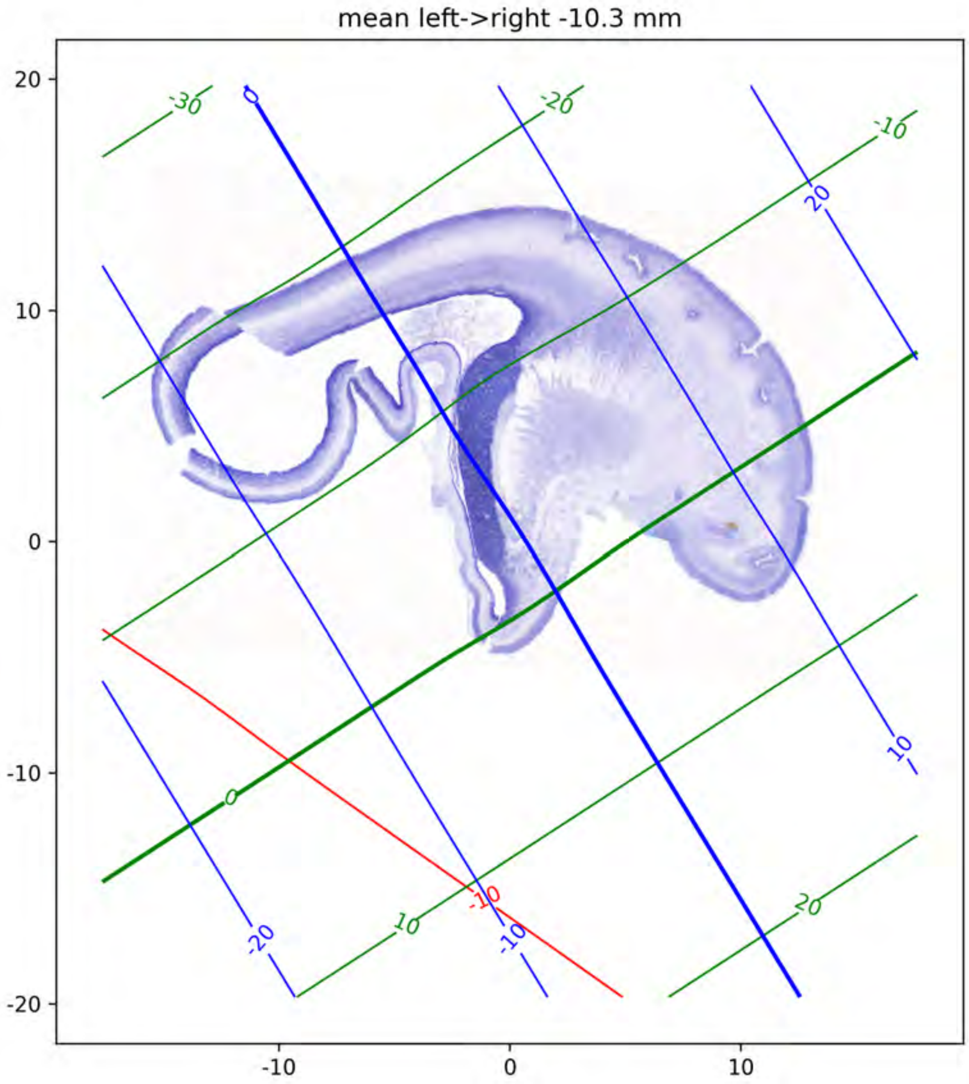
mean left->right -10.3 mm
20
10
0
-10
-20
-10
0
10
-30
-20
-10
0
10
20
0
20
10
-20
-10
-10

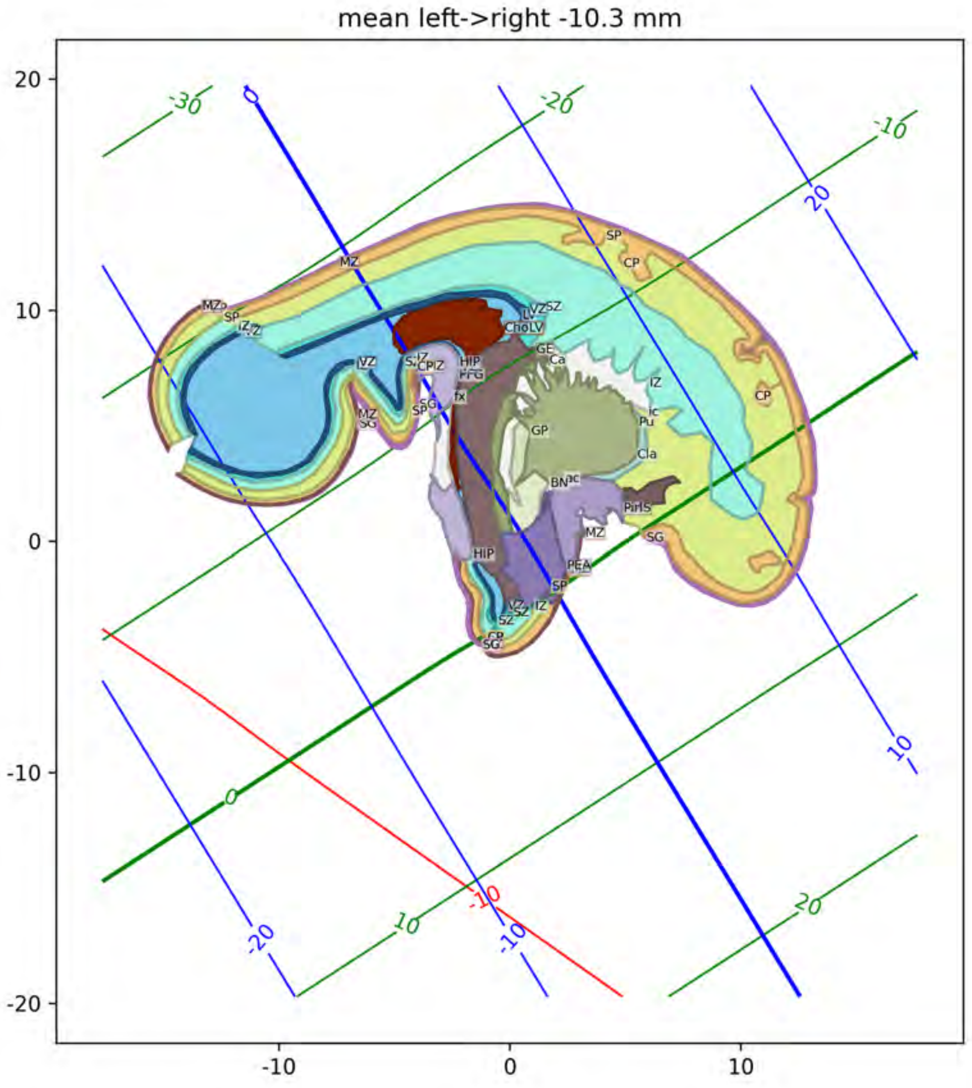
mean left->right -10.3 mm
MZ
SP
IZ
VZ
LV
ChoLV
SZ
GE
Ca
HIP
FFG
fx
CP
SG
GP
IZ
Pu
Cla
BN
PirIS
PEA
CP
HIP

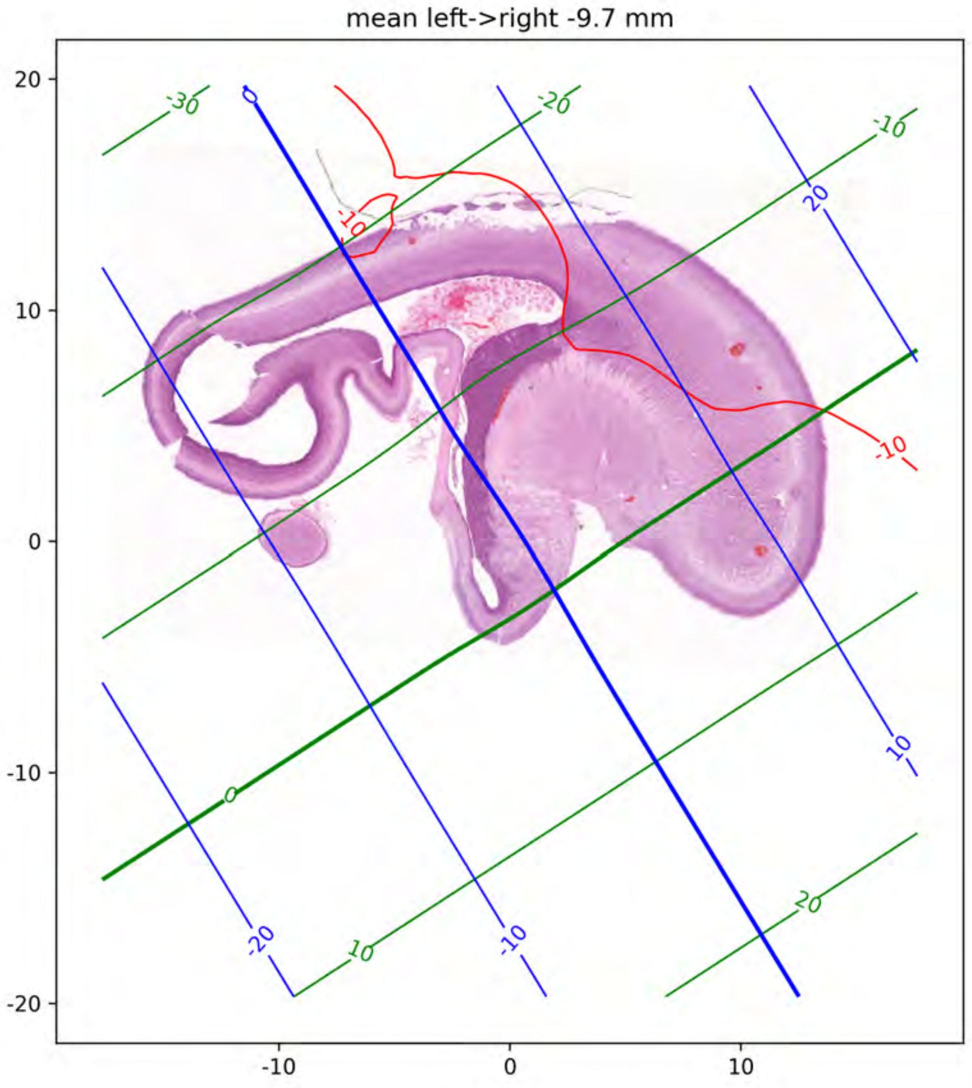
mean left->right -9.7 mm
20
10
0
-10
-20
-10
0
10
-30
-20
-10
0
10
20
0
20
10
-20
-10
-10
-10

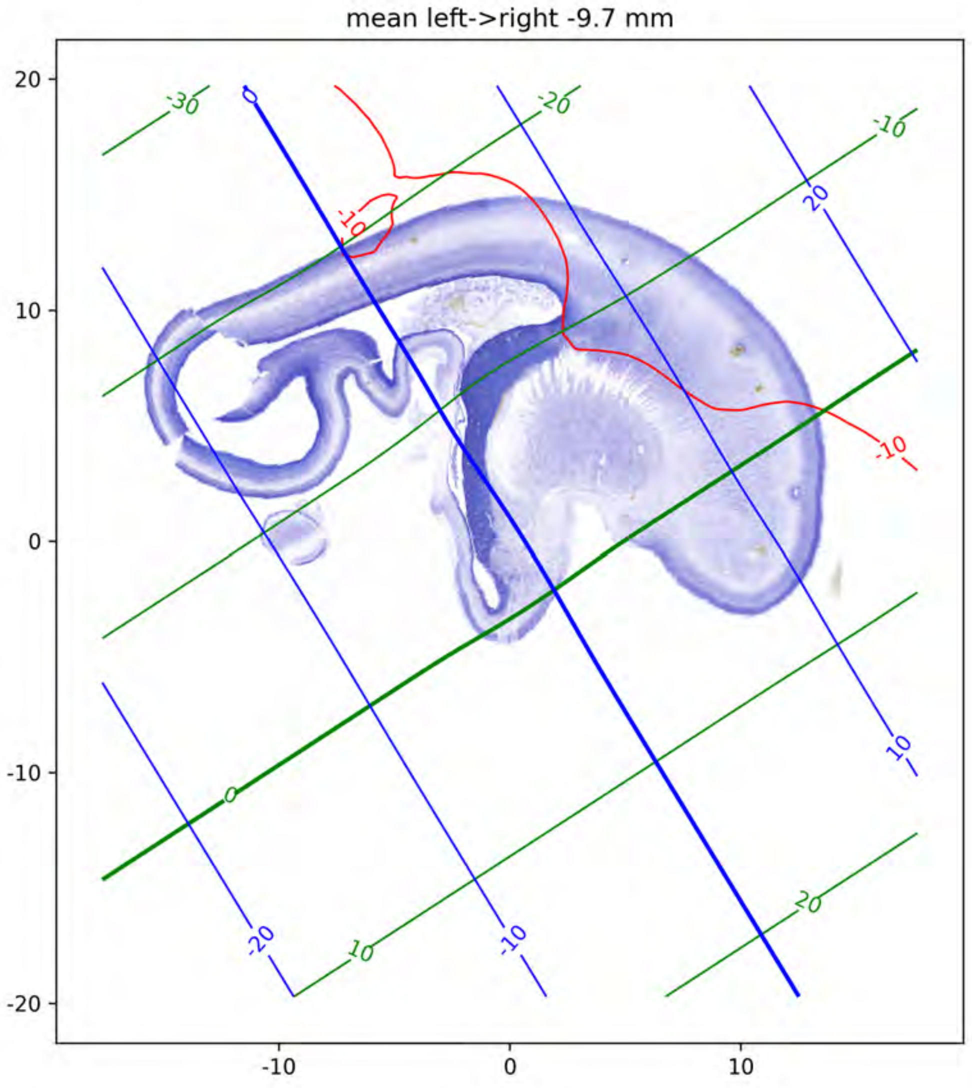
mean left->right -9.7 mm
20
10
0
-10
-20
-10
0
10
-30
-20
-10
20
10
-10
-10
0
-20
-10
10
20
0
20
10

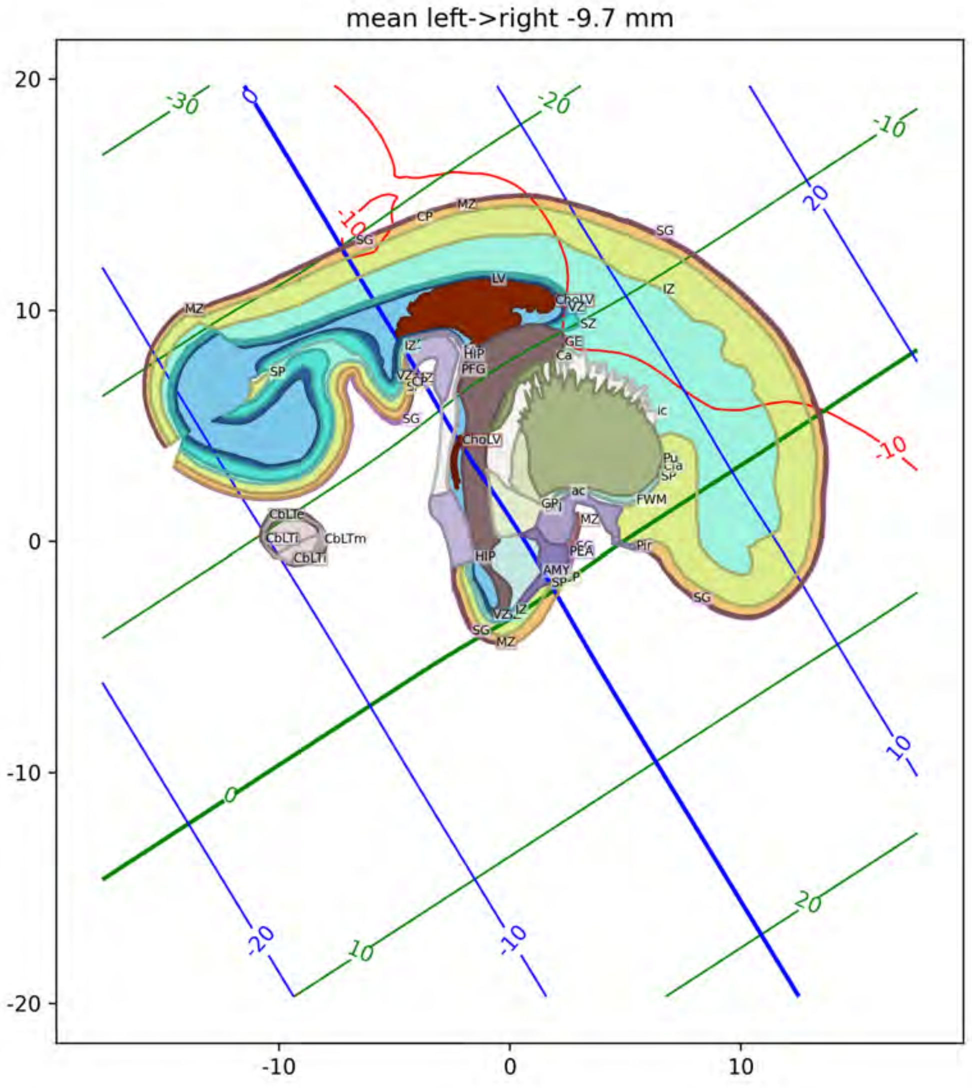
mean left->right -9.7 mm
MZ
CP
SG
LV
ChoLV
VZ
SZ
GE
Ca
IZ
SP
HIP
PFG
ic
Pu
Cla
FWM
ac
GP
PEA
AMY
Pir
CbLTe
CbLTi
CbLTm

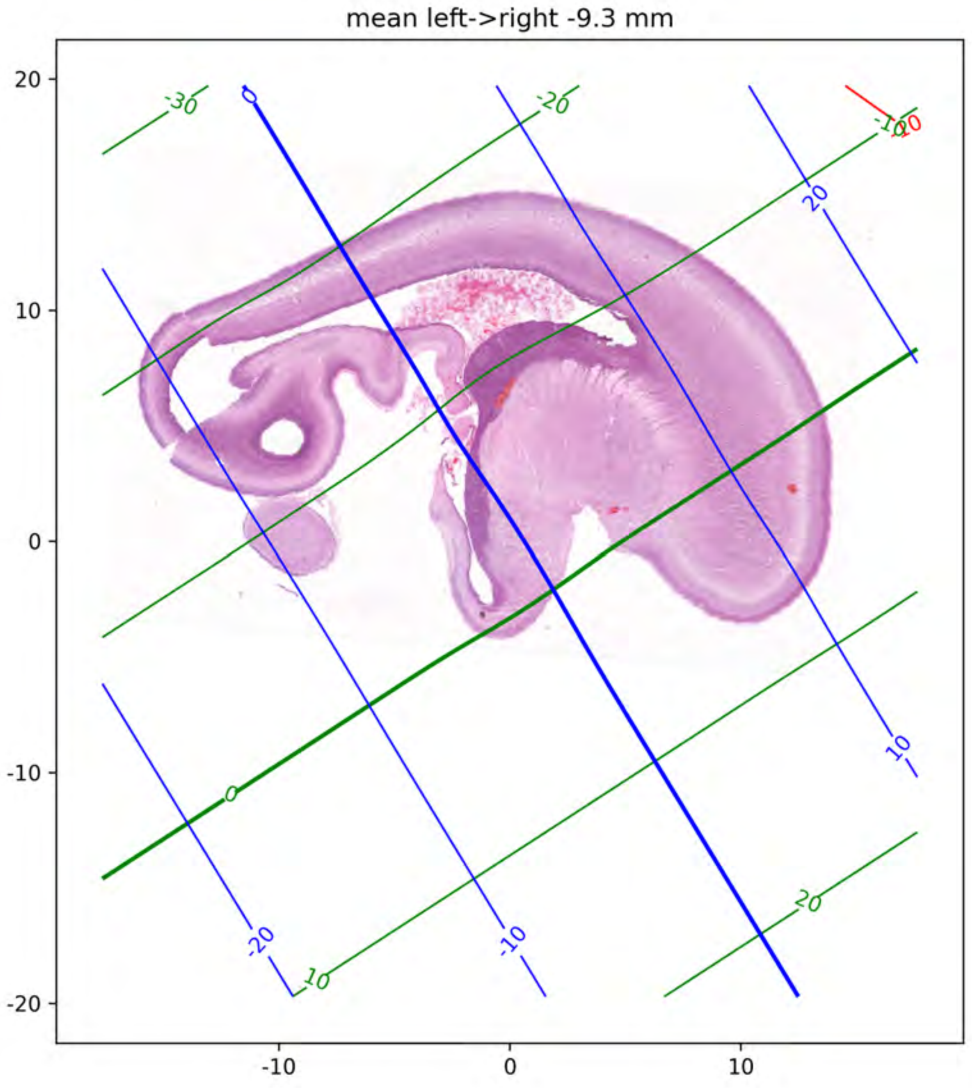
mean left->right -9.3 mm
20
10
0
-10
-20
-10
0
10
-30
-20
-10
0
10
20
0
20
10
-10
-20
-10
0

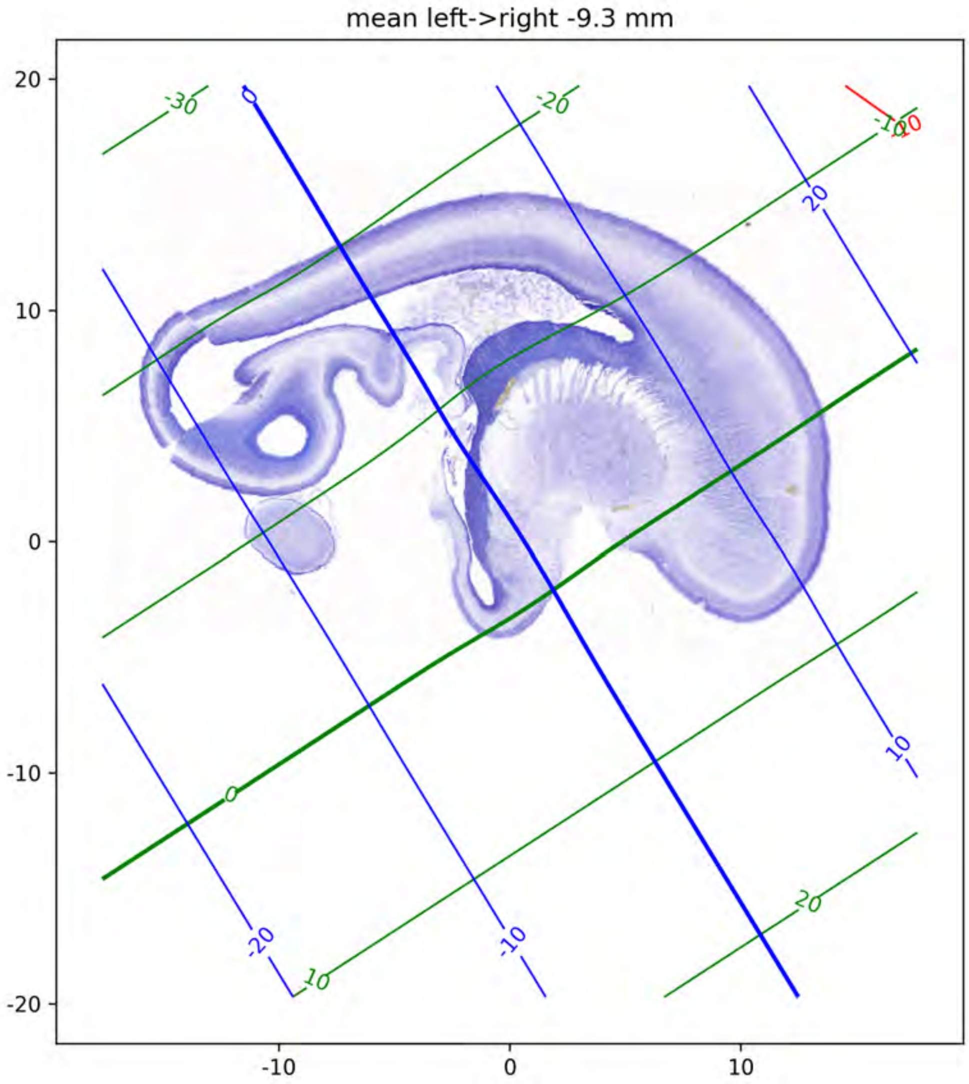
mean left->right -9.3 mm
20
10
0
-10
-20
-10
0
10
-30
-20
-10
0
10
20
0
20
10
-20
-10

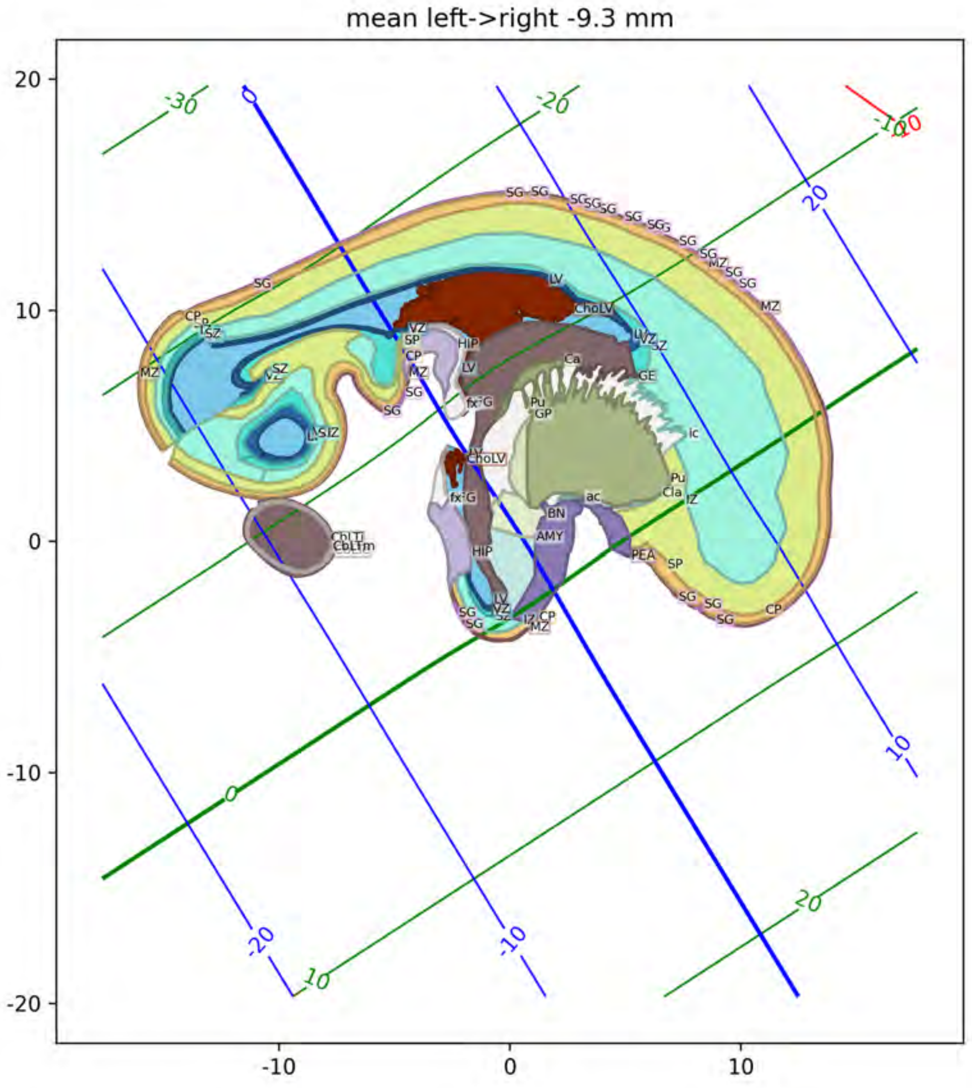
mean left->right -9.3 mm

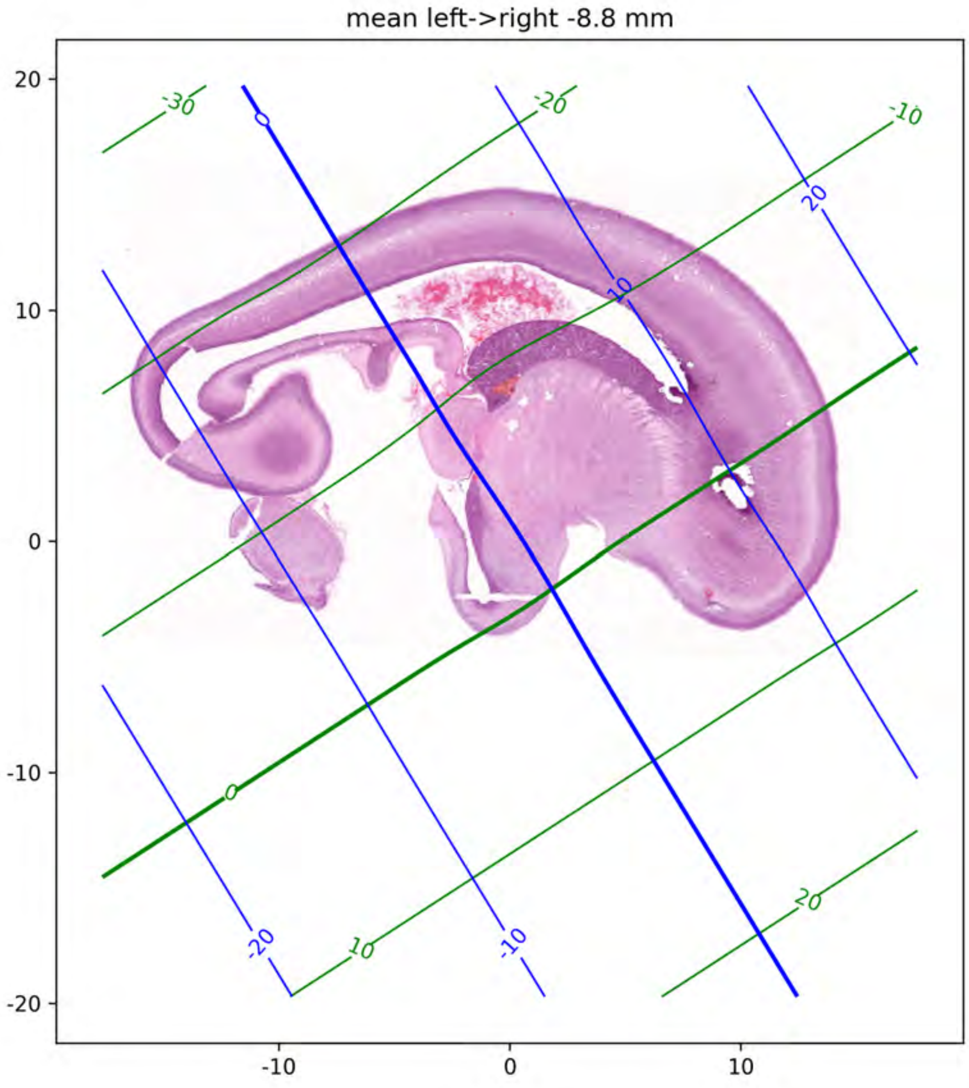
mean left->right -8.8 mm
20
10
0
-10
-20
-10
0
10
-30
-20
-10
0
10
20
0
10
20
-20
-10

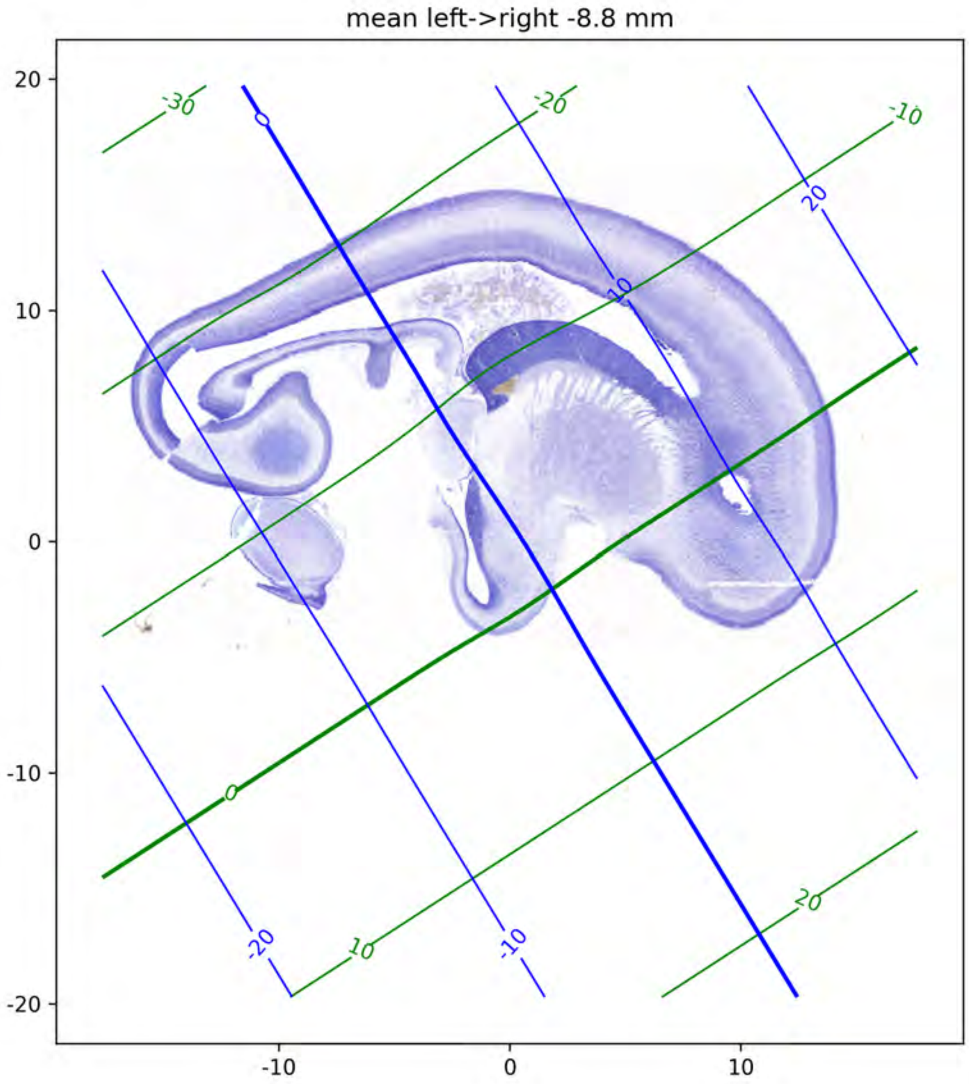
mean left->right -8.8 mm
20
10
0
-10
-20
-10
0
10
-30
-20
-10
0
10
20
0
10
20
-20
-10

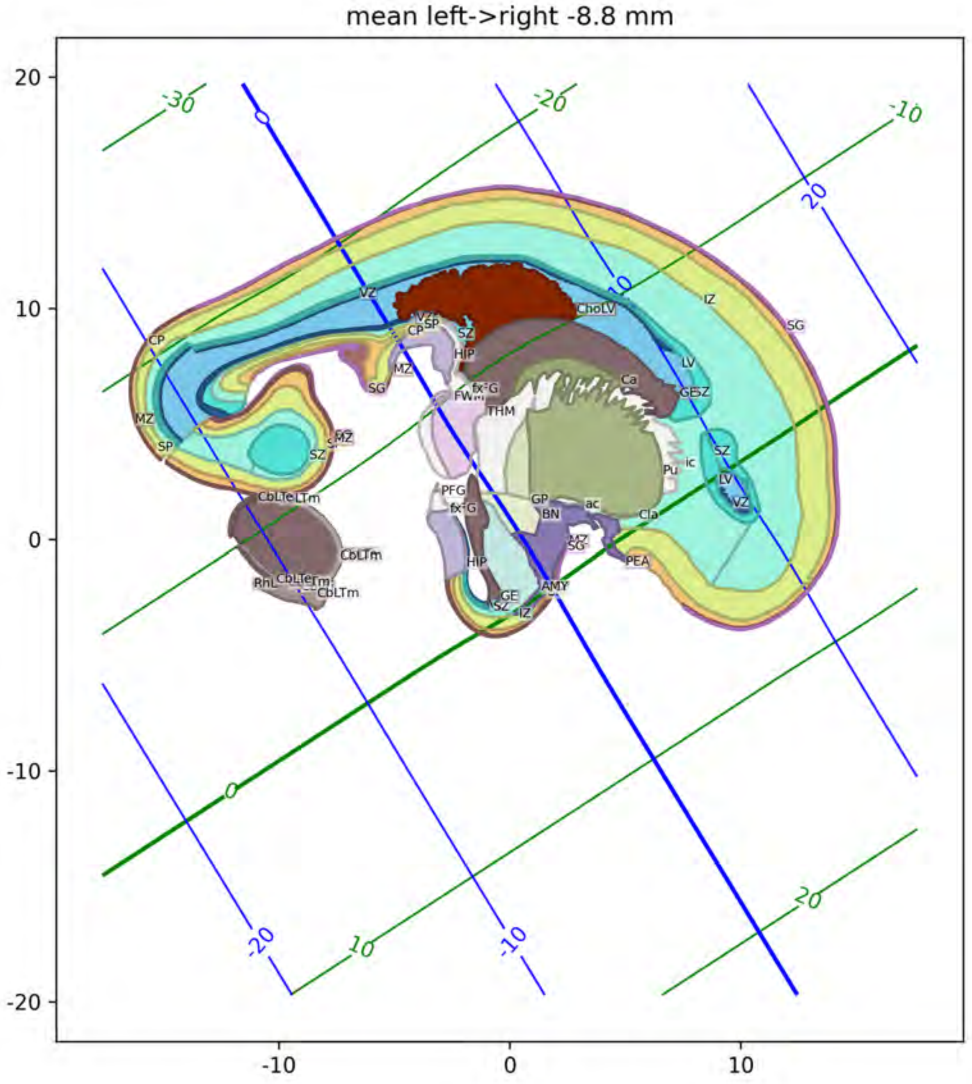

mean left->right -8.8 mm
20
10
0
-10
-20
-10
0
10
-30
-20
-10
0
10
20
-20
-10
0
10
20
VZ
CP
SP
SZ
HIP
MZ
SG
ChoLV
IZ
LV
Ca
GE
SZ
THM
FWM
fx
G
Pu
ic
Cla
GP
BN
ac
PFG
AMY
PEA
CbLTe
LTm
CbLTm
RhL

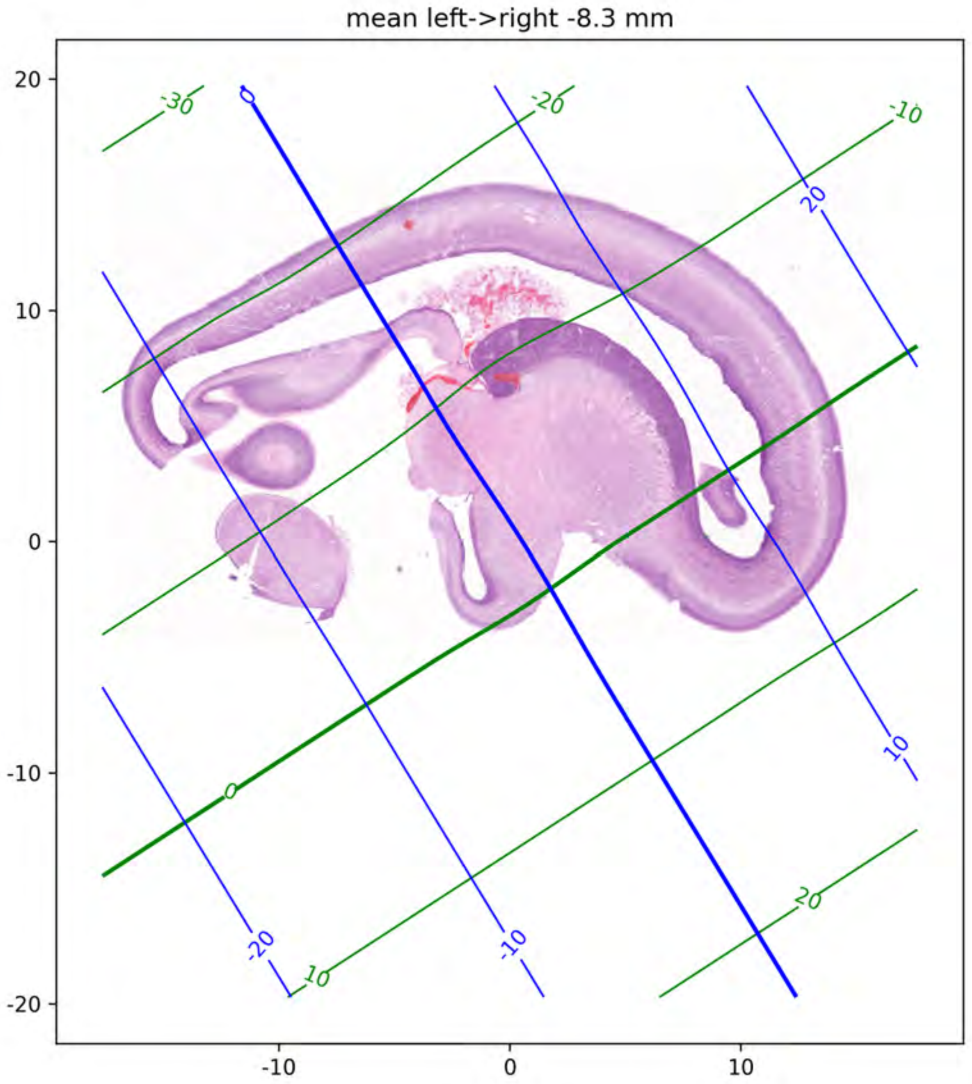
mean left->right -8.3 mm
20
10
0
-10
-20
-10
0
10
-30
-20
-10
0
10
20
0
20
10
-20
-10

mean left->right -8.3 mm

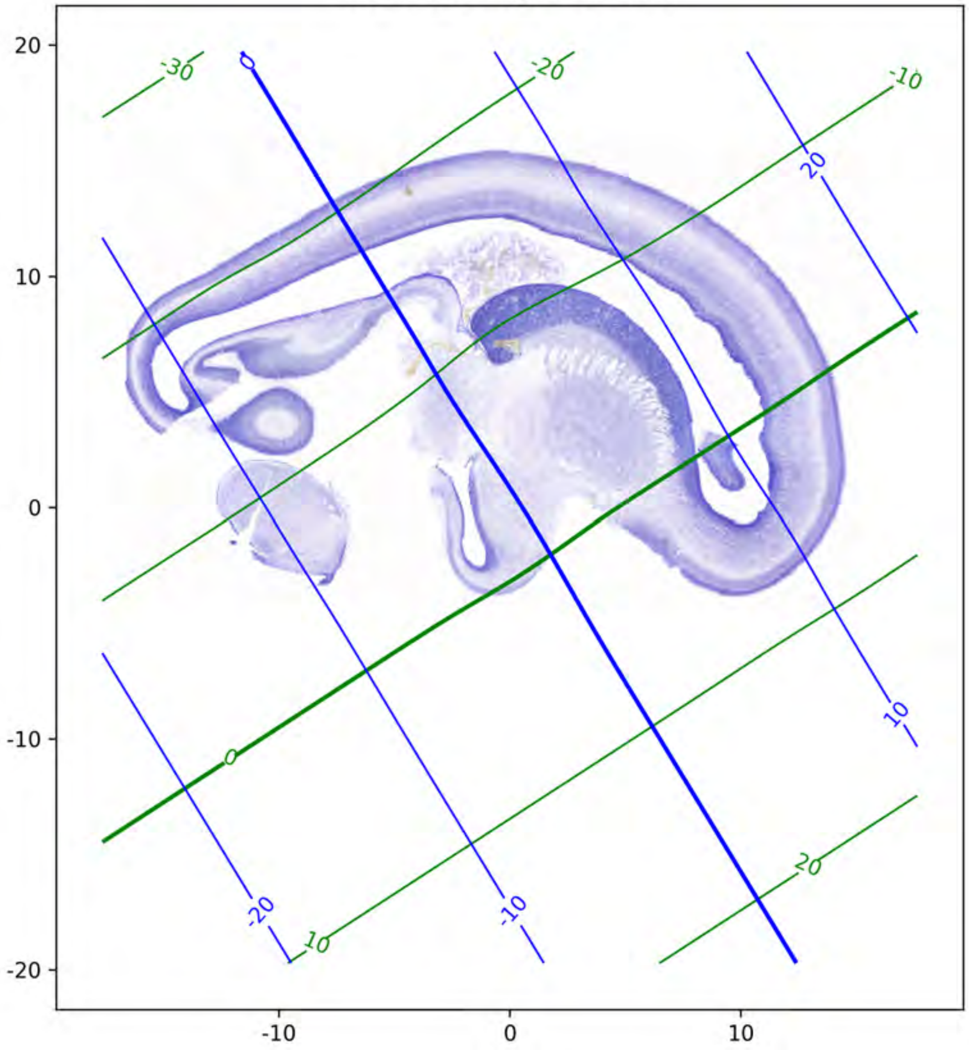

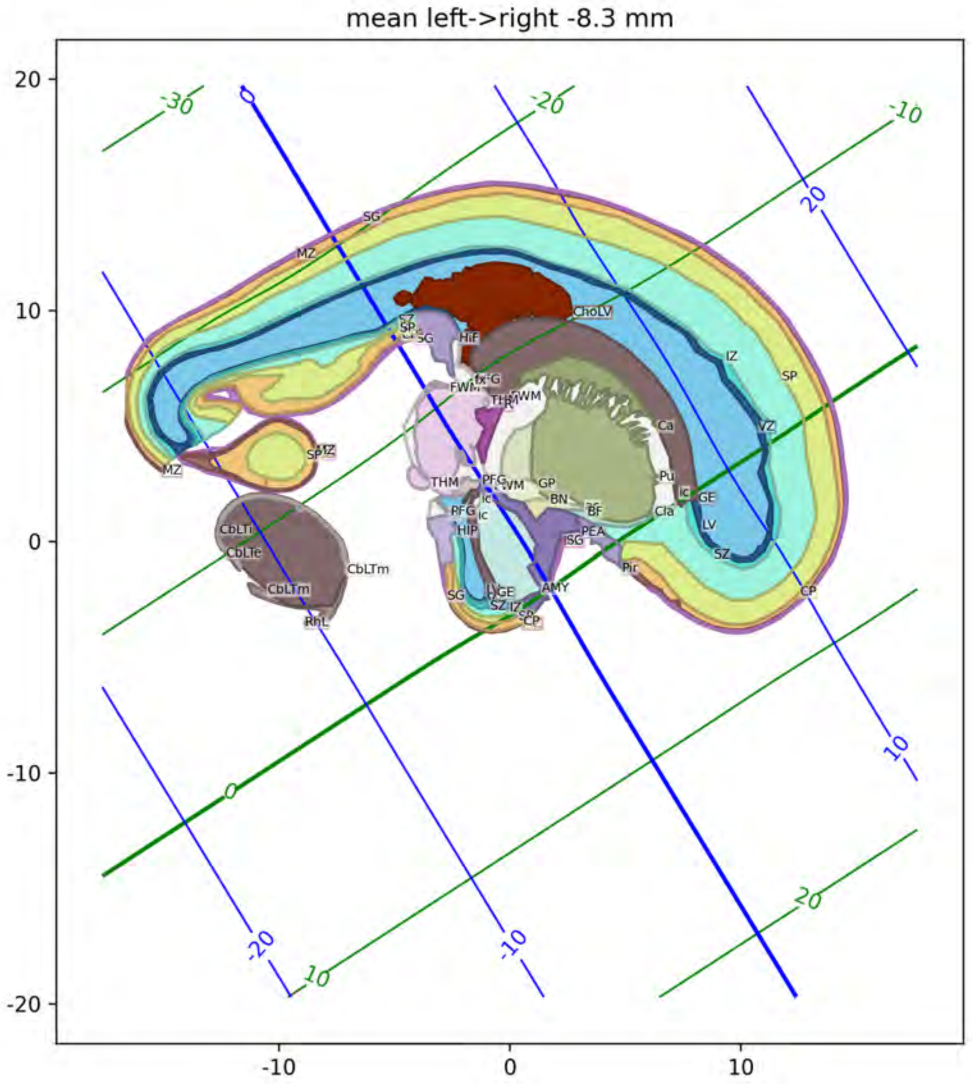
mean left->right -8.3 mm
20
10
0
-10
-20
-10
0
10
-30
-20
-10
20
10
0
-20
-10
10
20
0

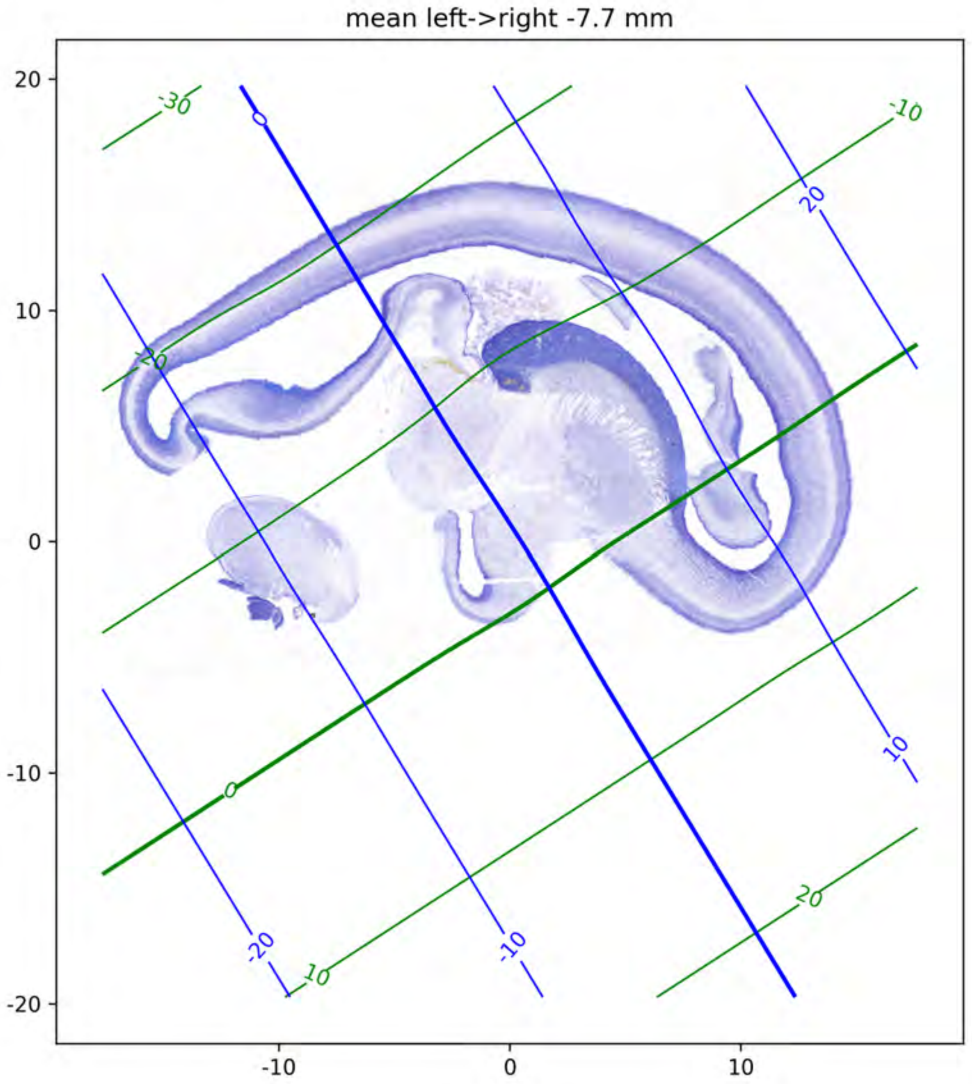
mean left->right -7.7 mm
20
10
0
-10
-20
-10
0
10
-30
-20
-10
0
10
20
0
20
10
-20
-10

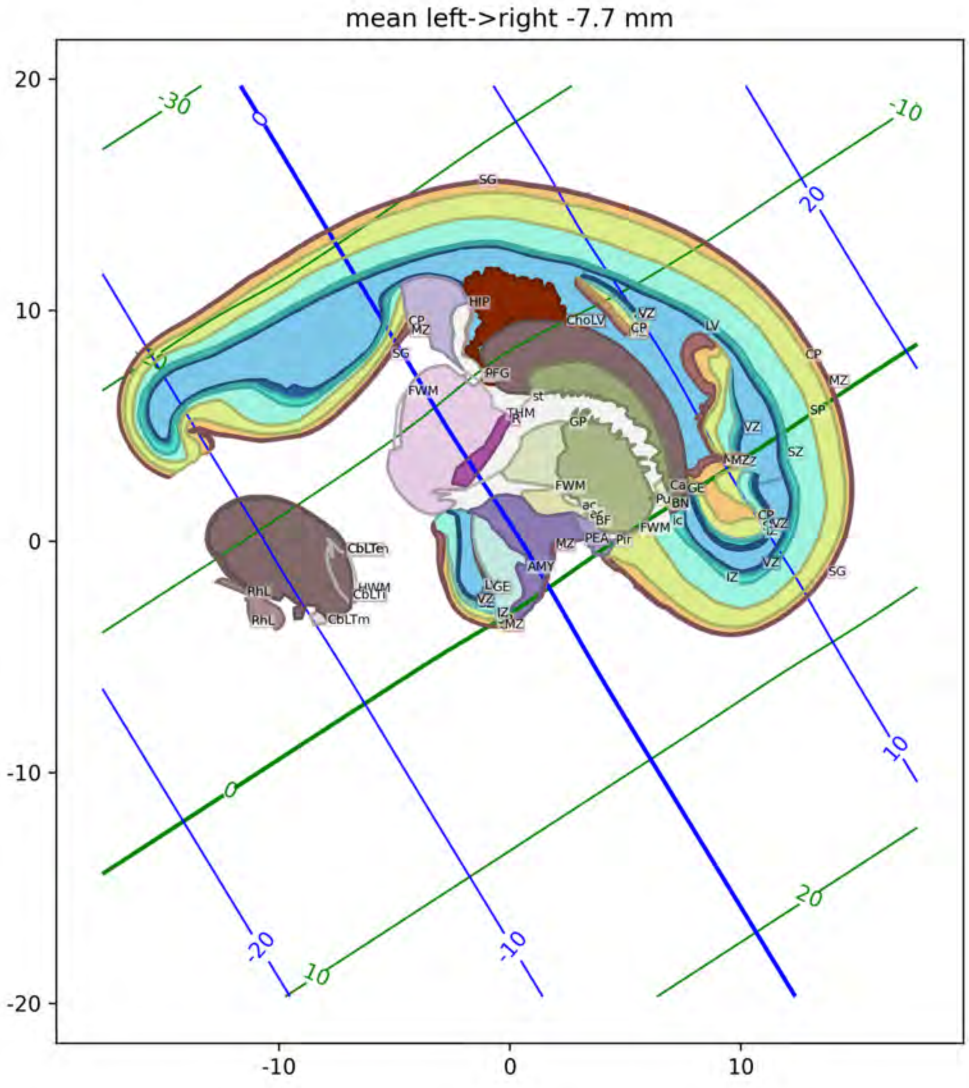
mean left->right -7.7 mm
20
10
0
-10
-20
-10
0
10
-30
-10
20
10
-20
-10
0
10
20
SG
HIP
CP
MZ
ChoLV
VZ
CP
LV
PFG
FWM
st
THM
GP
VZ
MZZ
SZ
SP
MZ
CP
FWM
Ca
GE
Pu
BN
ic
ac
BF
PEA
Pir
FWM
MZ
AMY
LGE
VZ
IZ
MZ
IZ
VZ
SG
CbLTen
HWM
CbLTi
RhL
RhL
CbLTm

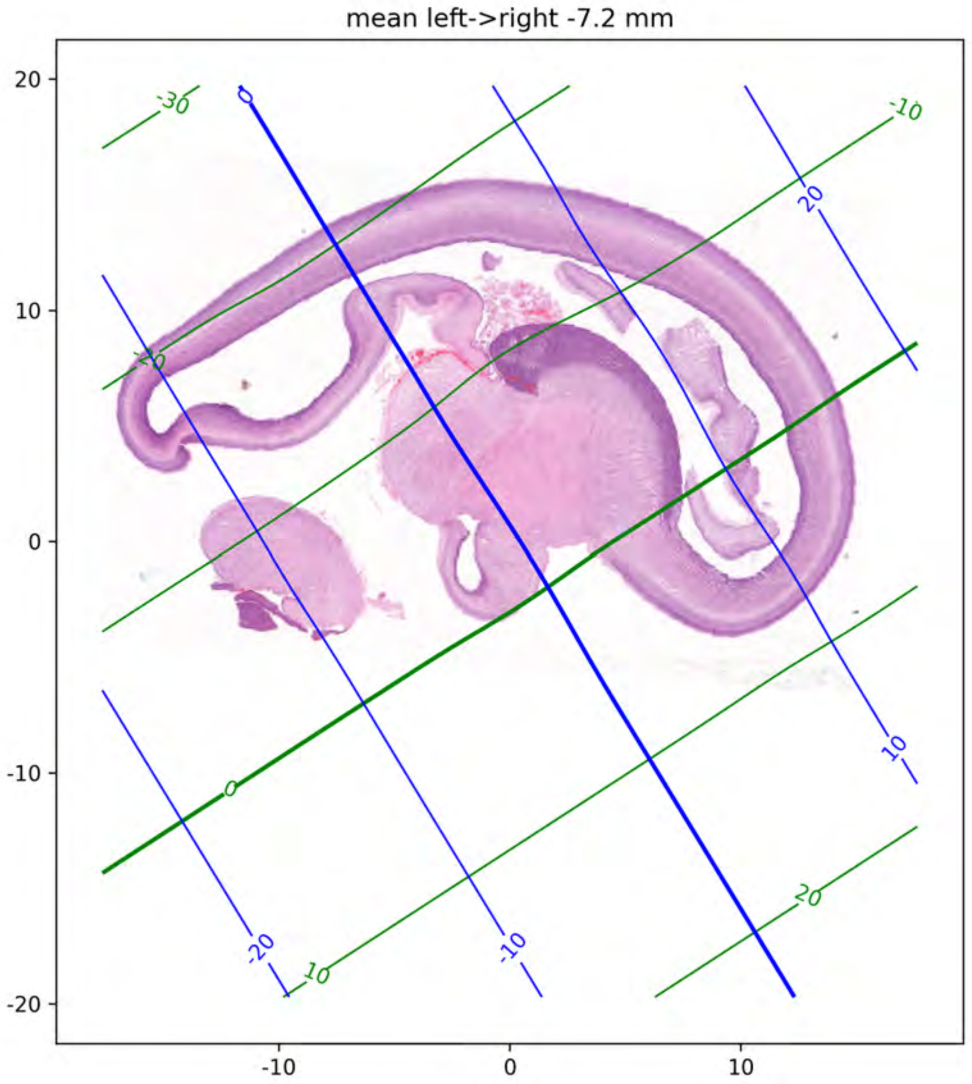
mean left->right -7.2 mm
20
10
0
-10
-20
-10
0
10

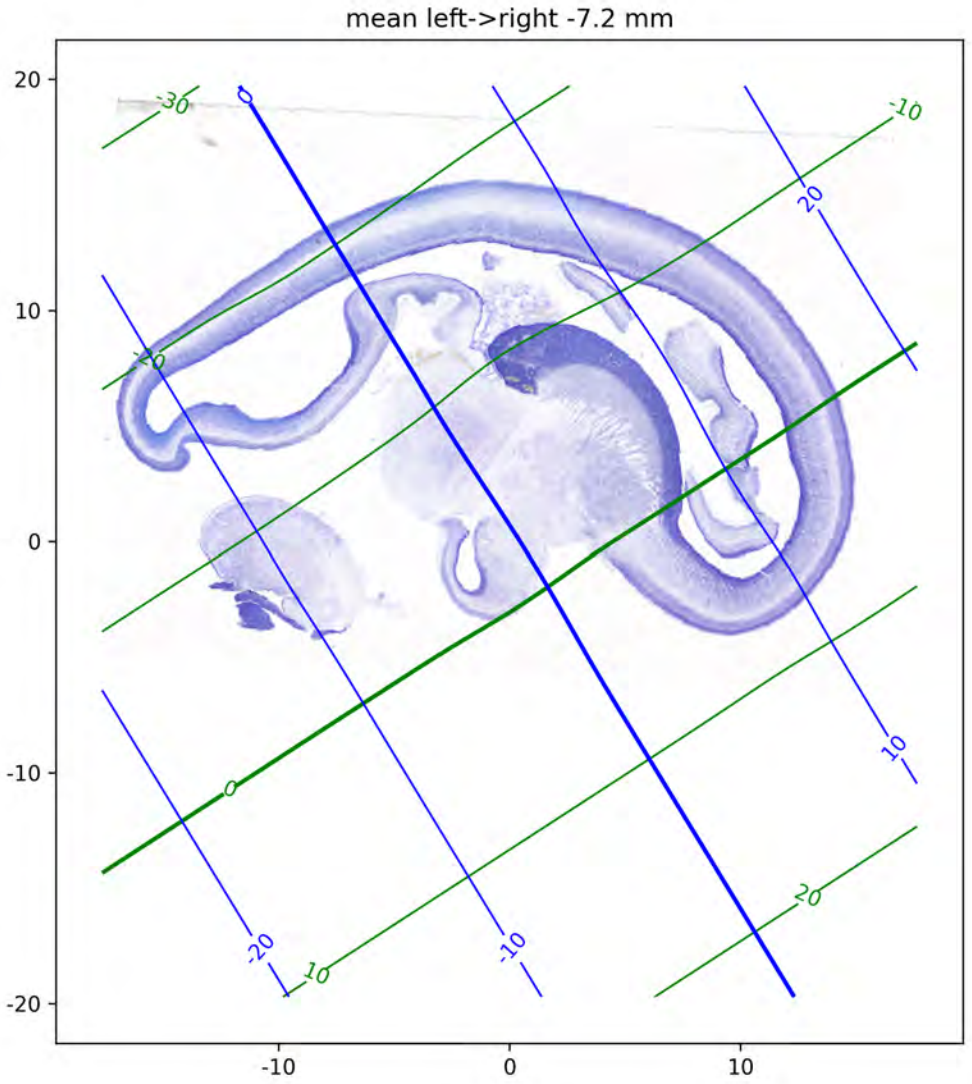
mean left->right -7.2 mm
20
10
0
-10
-20
-10
0
10
-30
-20
-10
0
10
20
0
20
10
-20
-10

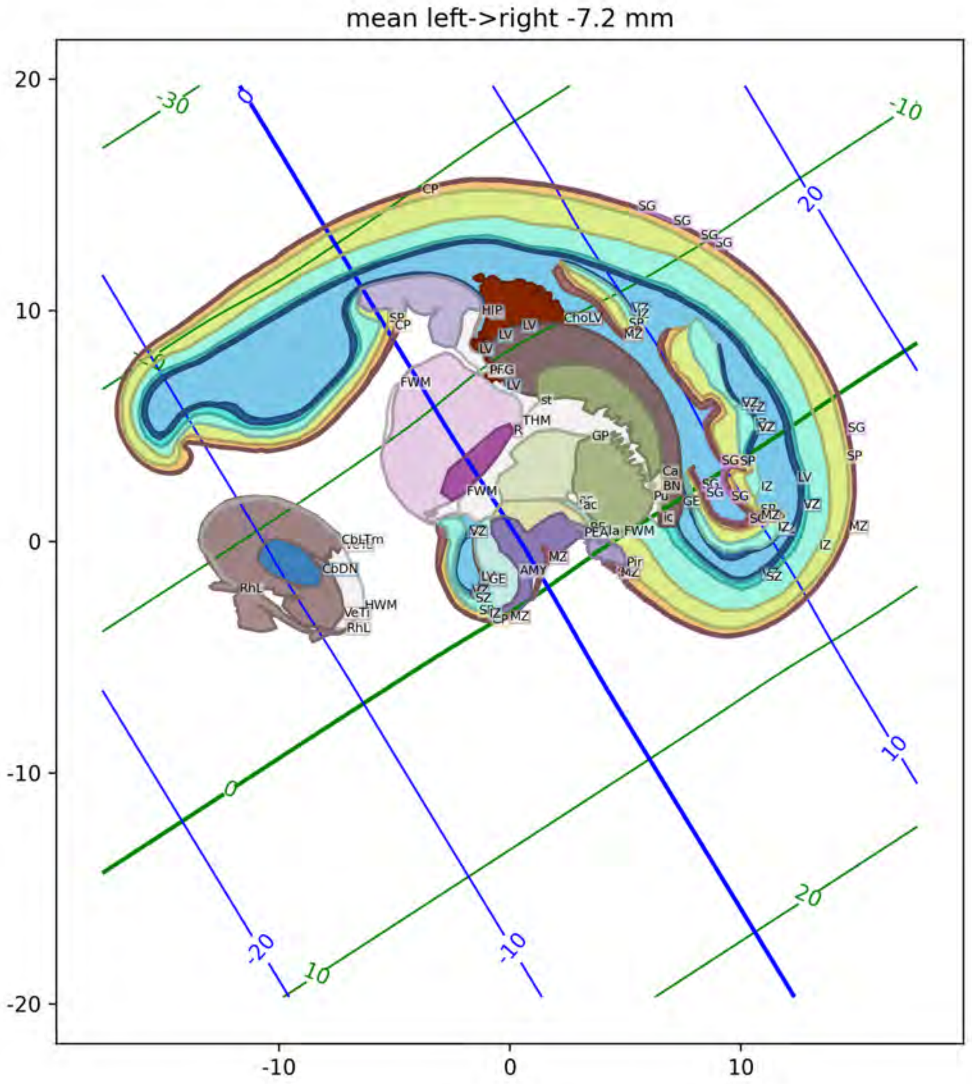
mean left->right -7.2 mm
20
10
0
-10
-20
-10
0
10
-30
-10
-20
-10
0
10
20
CP
SG
HIP
LV
ChoLV
SP
MZ
FWM
PFG
st
THM
GP
Ca
BN
Pu
GE
ic
ac
Ala
AMY
Pir
VZ
IZ
SZ
CbDN
RhL
HWM
VeTi

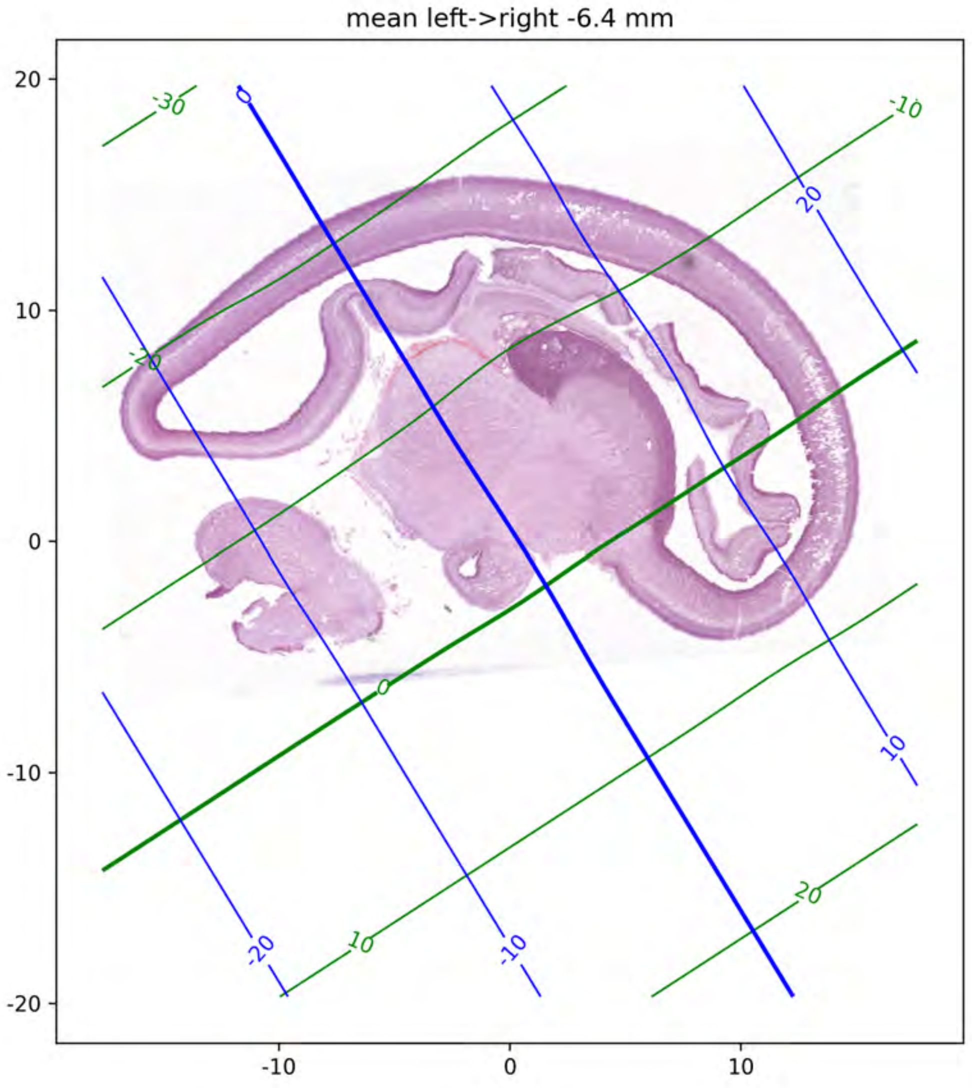
mean left->right -6.4 mm
20
10
0
-10
-20
-10
0
10
-30
-20
-10
0
10
20
-20
-10
0
10
20

mean left->right -6.4 mm

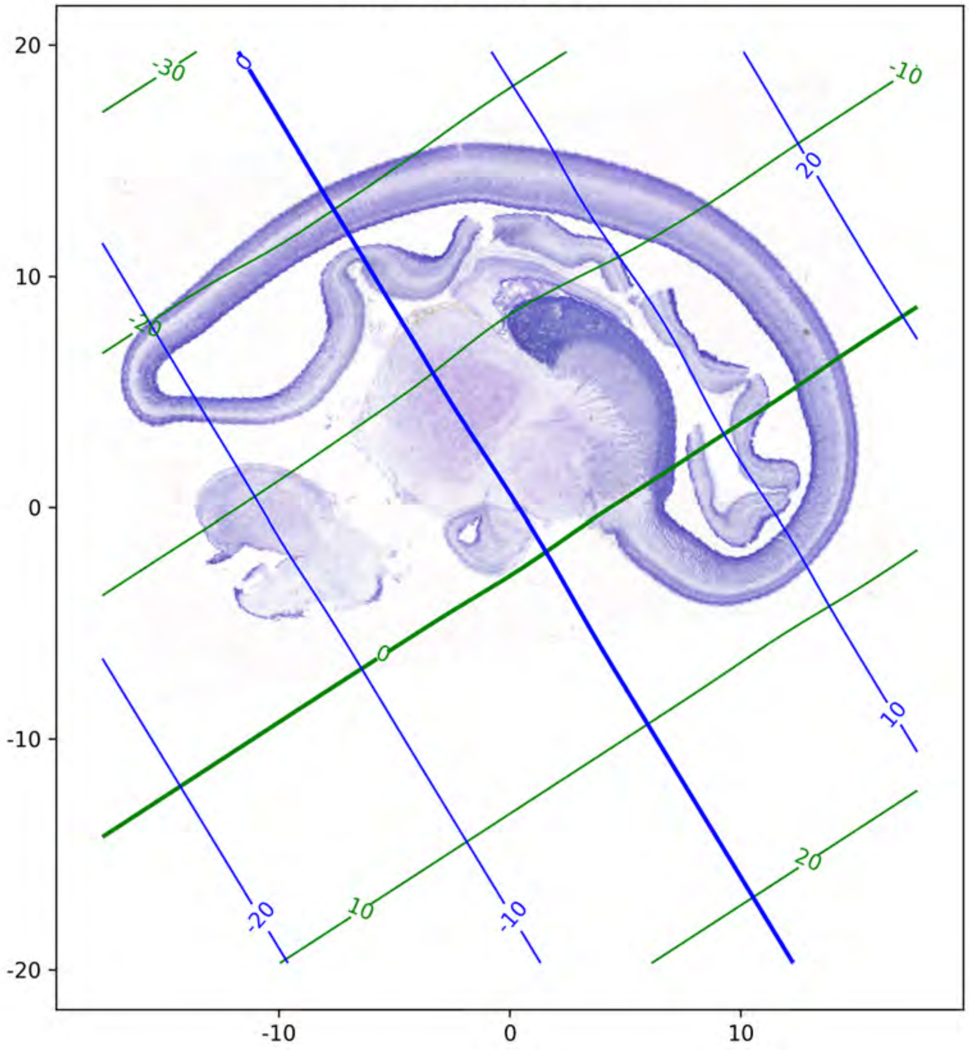

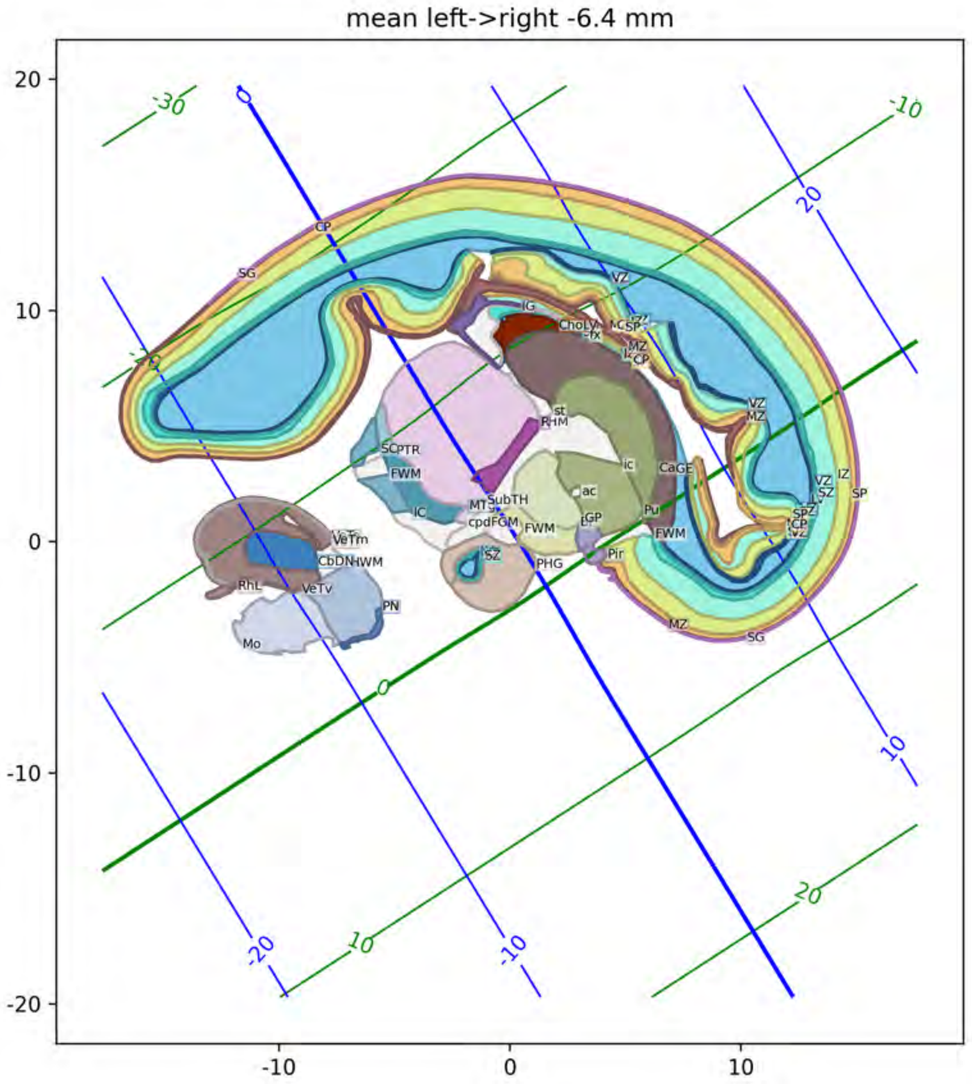

mean left->right -6.4 mm

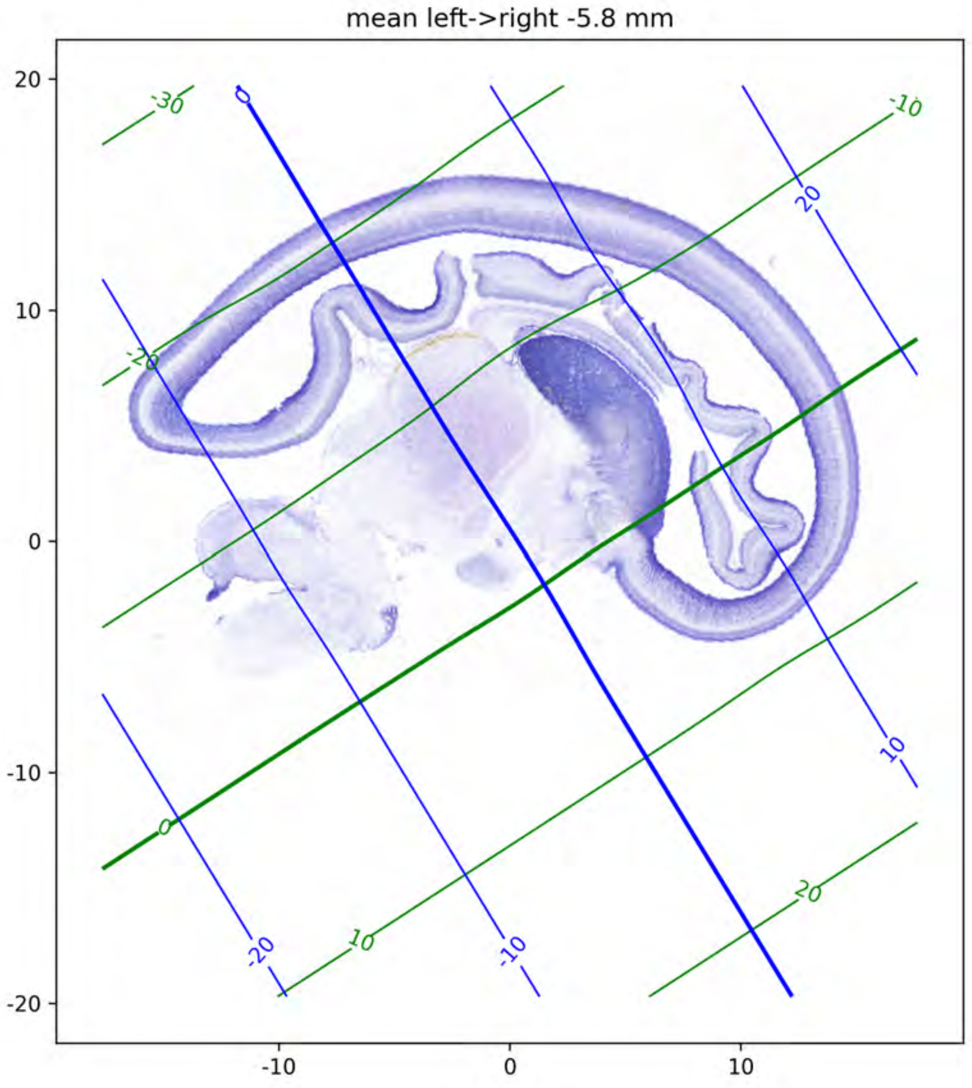

mean left->right -5.8 mm
20
10
0
-10
-20
-10
0
10
-30
-20
-10
0
10
20

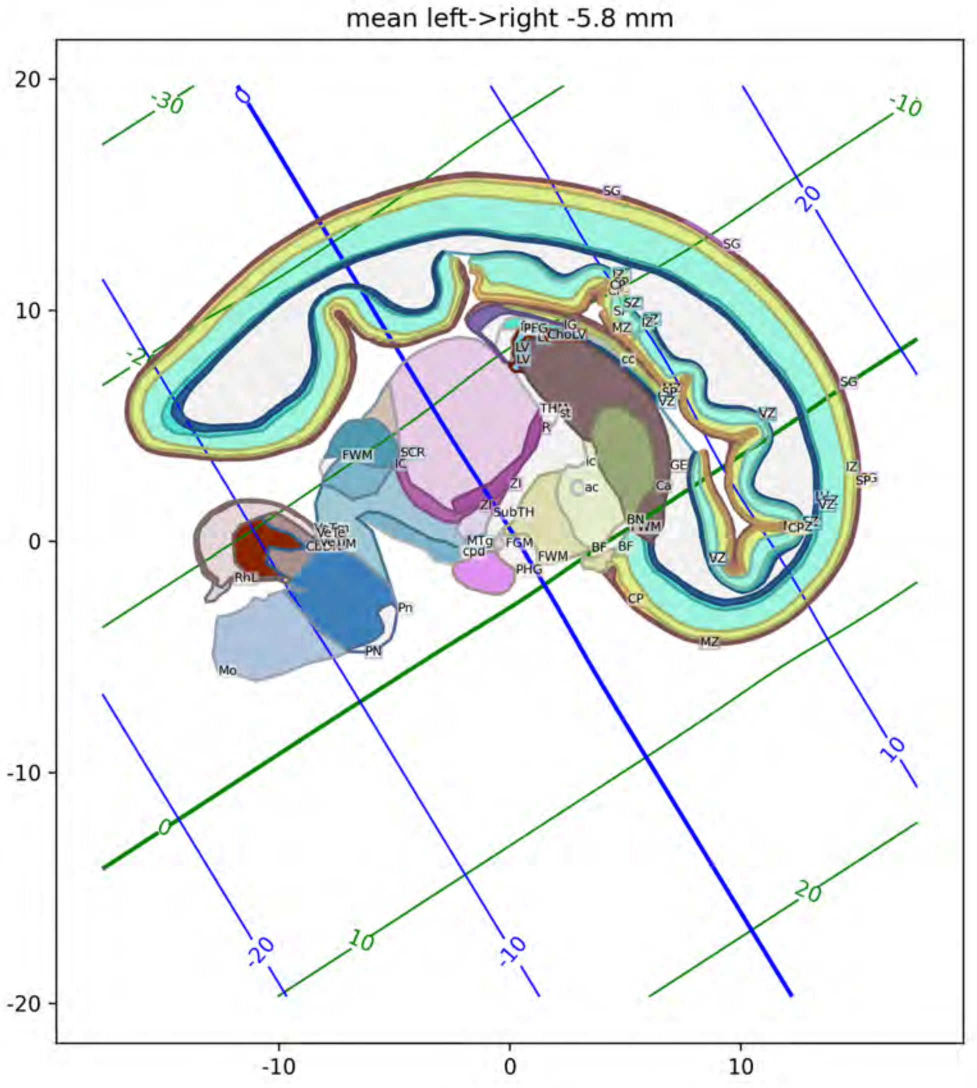
mean left->right -5.8 mm
20
10
0
-10
-20
-10
0
10
SG
SG
SG
IZ
CP
SZ
MZ
IZ
CC
SP
VZ
VZ
IZ
SP
VZ
CP
MZ
CP
FWM
SCR
IC
LV
LV
ChoLV
IG
TH
St
R
ic
ac
GE
Ca
ZI
ZI
SubTH
BN
WM
MTg
cpd
FGM
FWM
BF
BF
PHG
RhL
Pn
PN
Mo
-30
-10
0
20
10
20
-20
10
-10
0

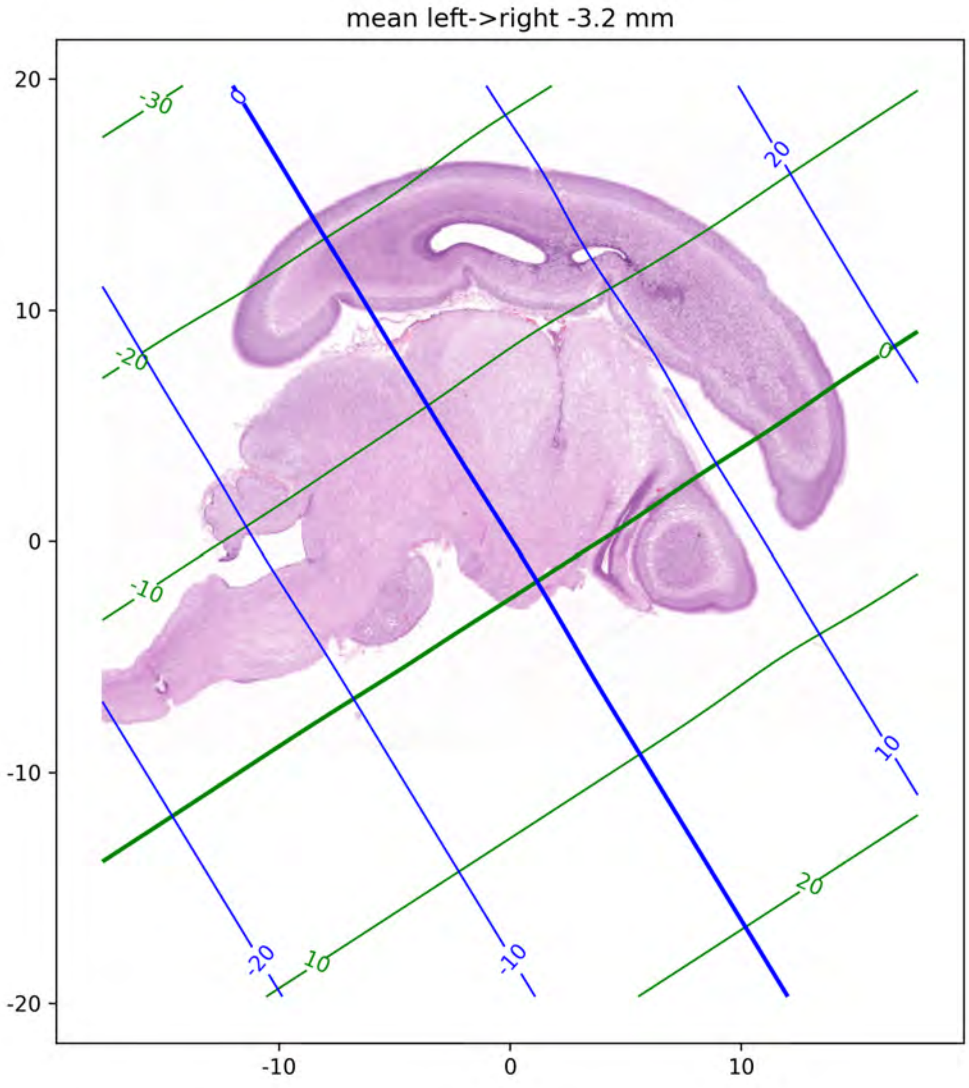

mean left->right -3.2 mm
20
10
0
-10
-20
-10
0
10
-30
-20
-10
0
10
20

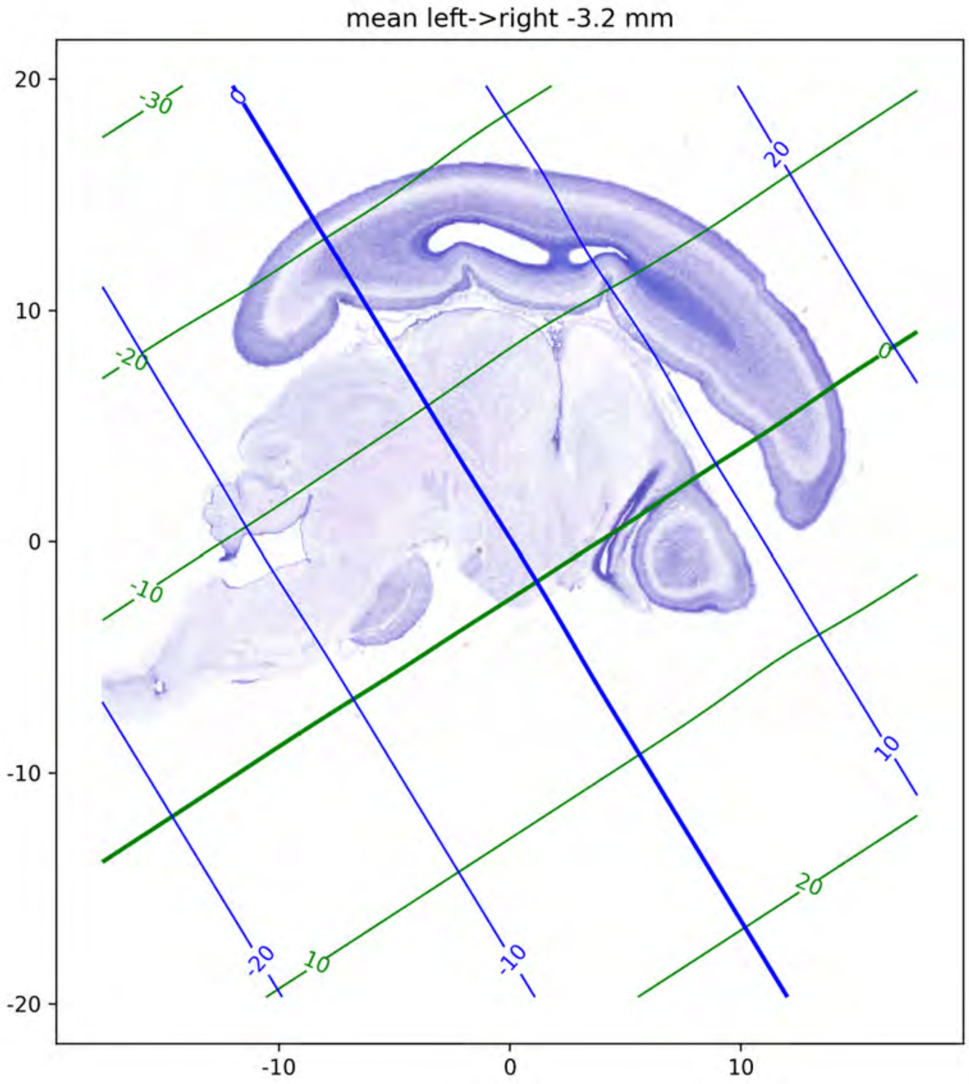
mean left->right -3.2 mm
20
10
0
-10
-20
-10
0
10
-30
-20
-10
0
10
20
0
20
10
-20
-10

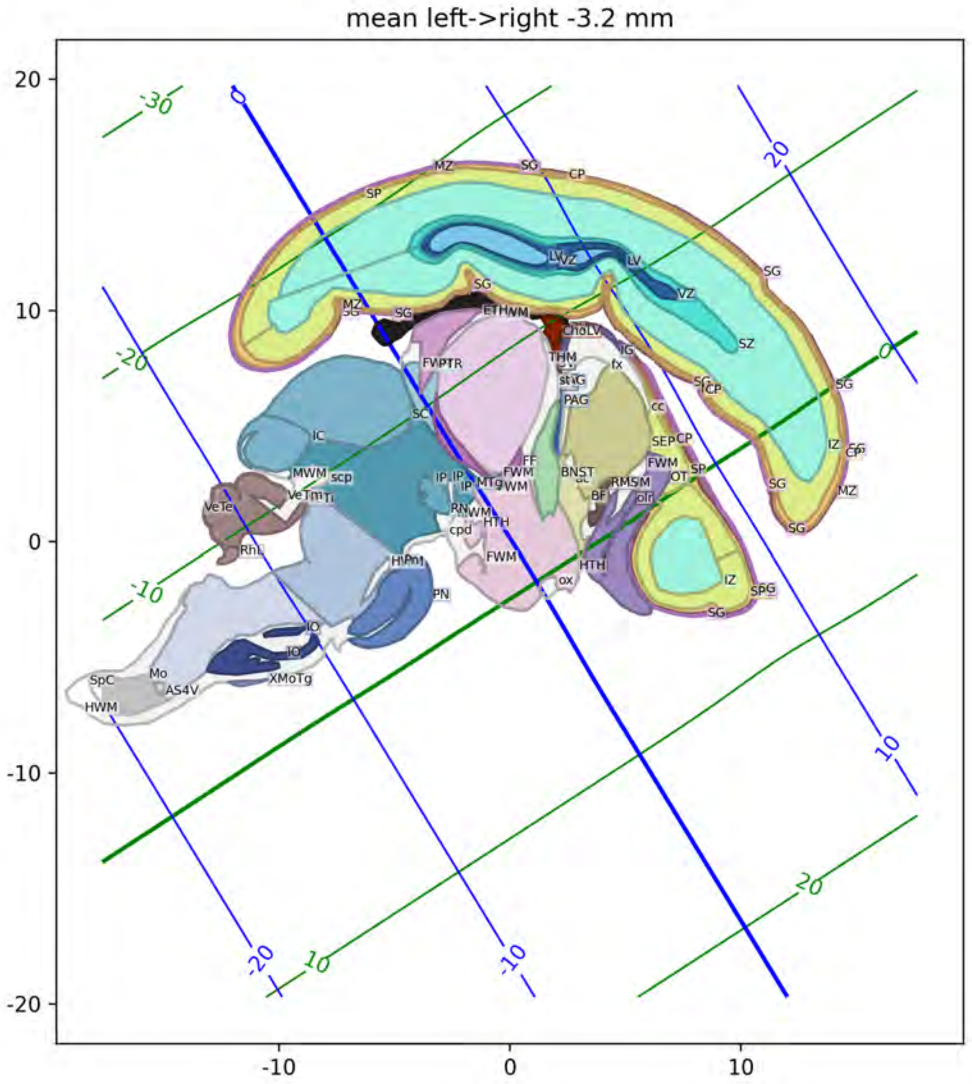
mean left->right -3.2 mm
20
10
0
-10
-20
-10
0
10
MZ
SG
CP
SP
LV
VZ
SZ
IZ
MZ
ETHWM
ChoLV
THM
PAG
fx
cc
SEP
FWM
OT
BNST
RMS
BF
HTH
ox
FVPTR
SC
IC
MWM
scp
VeTe
RhL
IP
MTg
RN
cpd
FWM
PN
IO
SpC
Mo
AS4V
XMoTg
HWM
-30
-20
-10
20

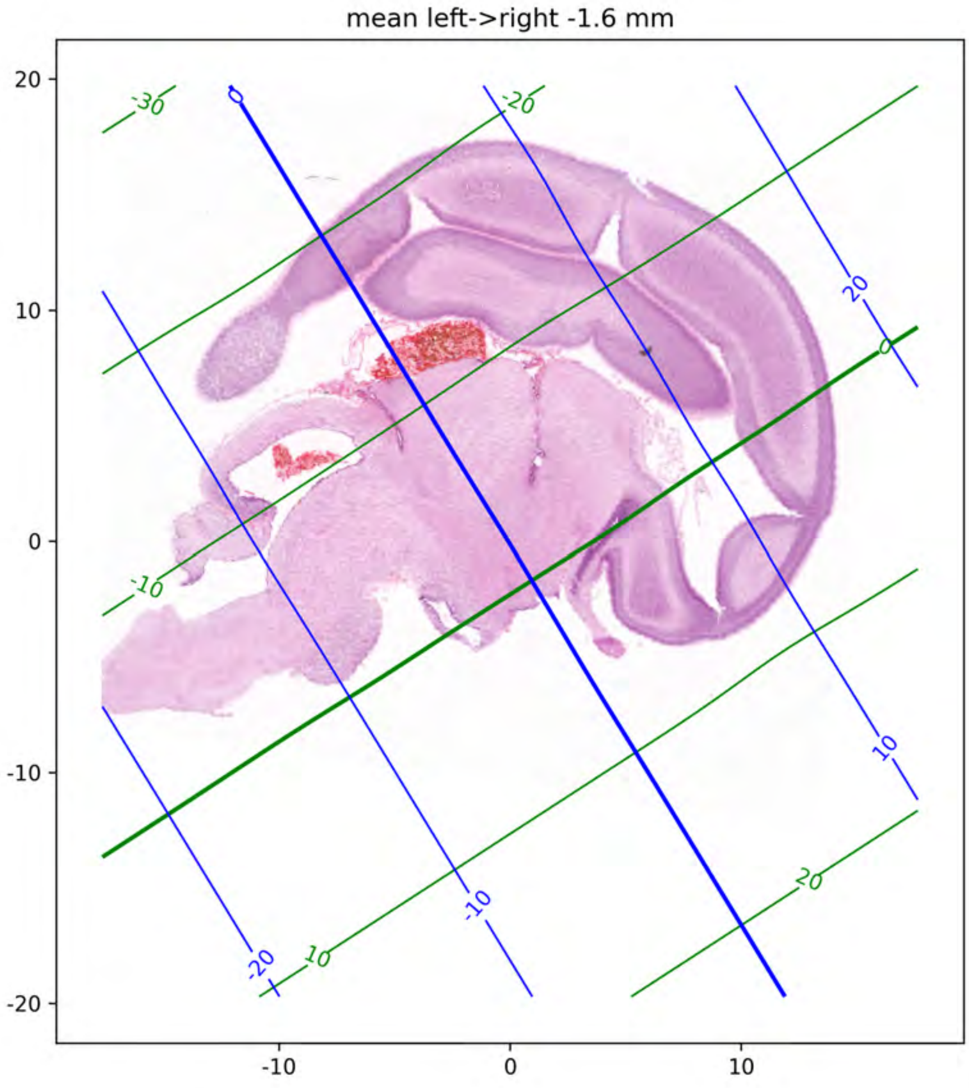
mean left->right -1.6 mm
20
10
0
-10
-20
-10
0
10
-30
-20
-10
0
10
20
0
-20
-10
10
20

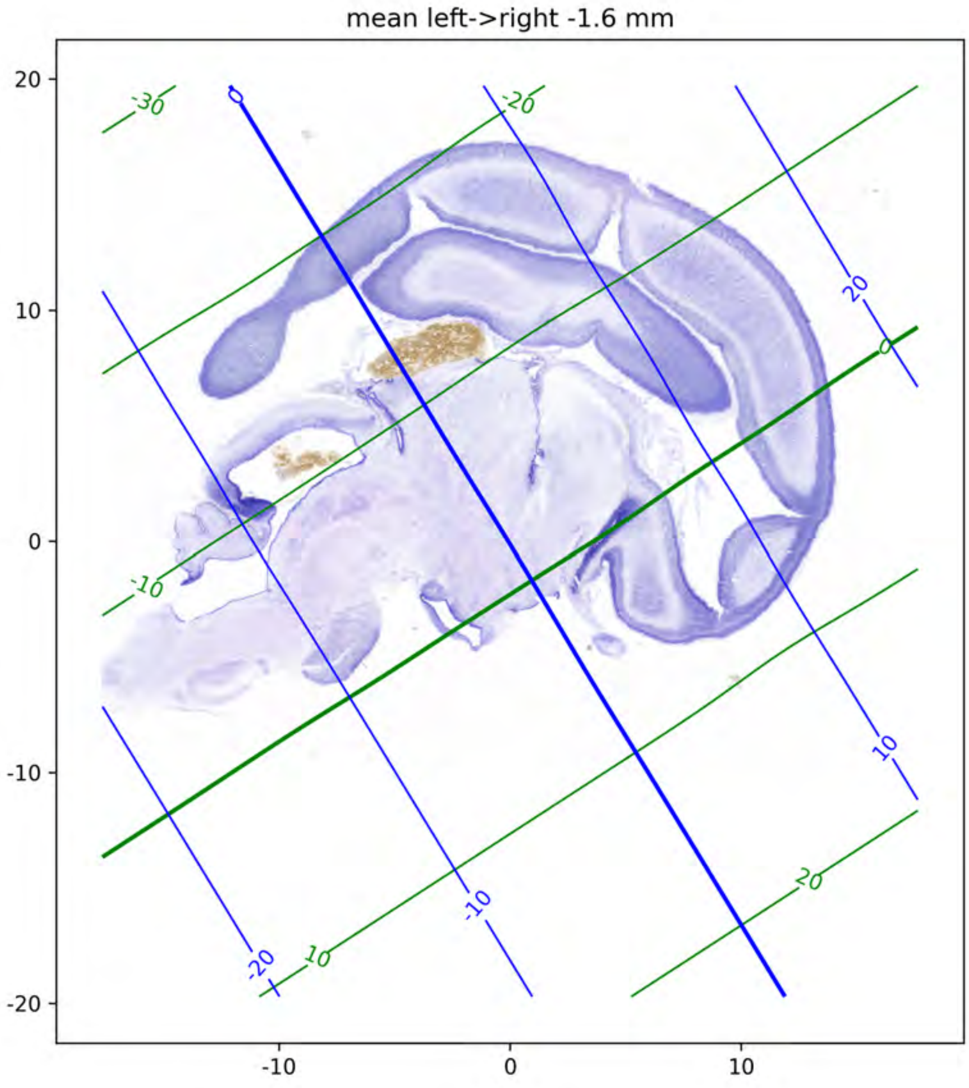
mean left->right -1.6 mm
20
10
0
-10
-20
-10
0
10
-30
-20
-10
0
10
20
0
20
10
-10
-20

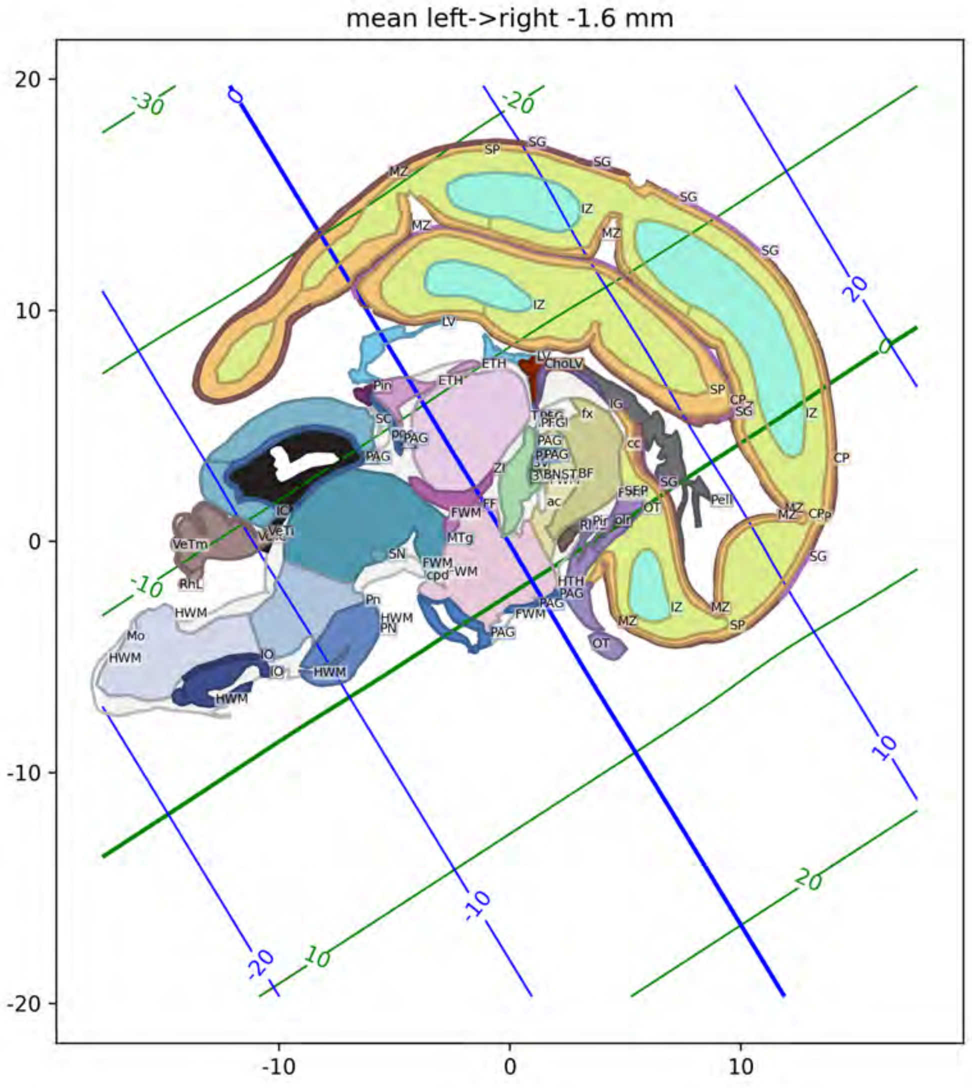
mean left->right -1.6 mm
20
10
0
-10
-20
-10
0
10
-30
-20
-10
0
10
20
0
20
10
-10
-20

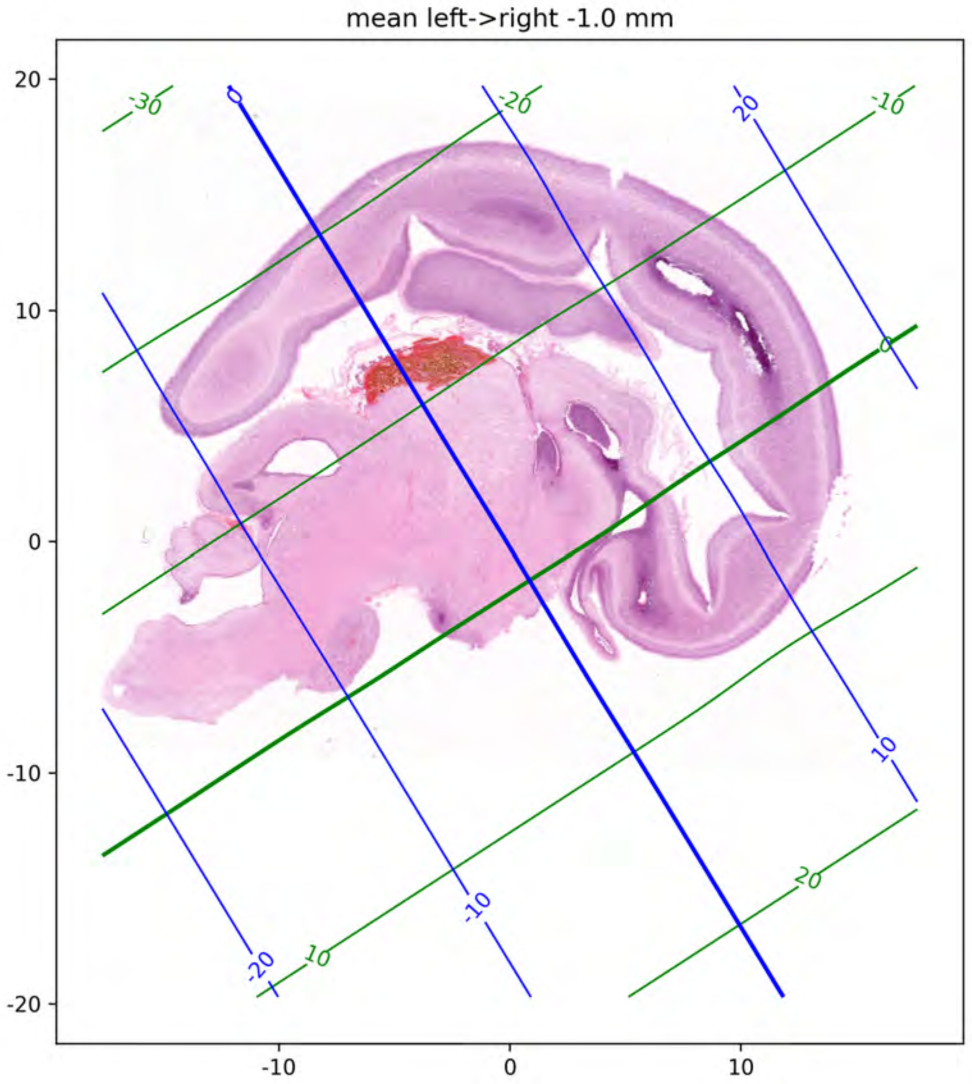
mean left->right -1.0 mm
20
10
0
-10
-20
-10
0
10
-30
-20
-10
0
10
20
0
20
10
-10
-20

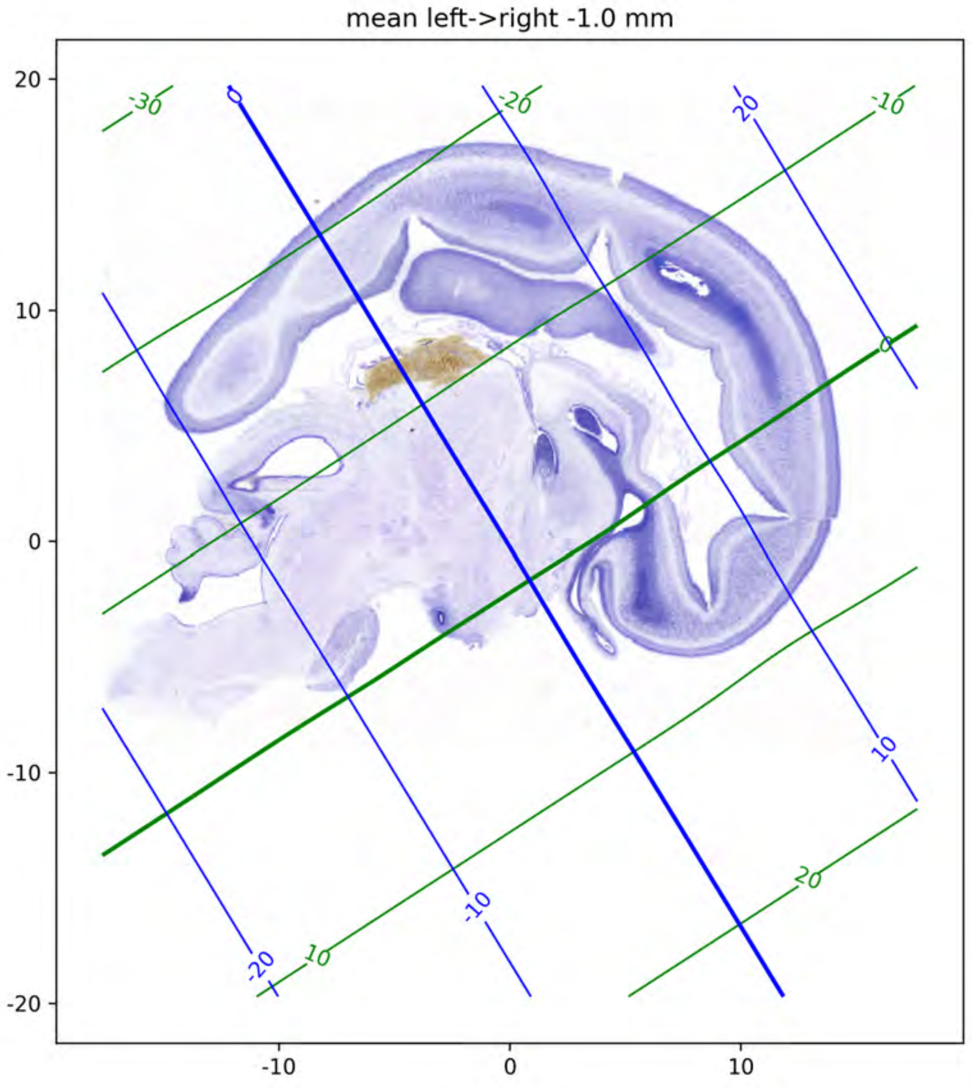
mean left->right -1.0 mm
20
10
0
-10
-20
-10
0
10
-30
-20
-10
0
10
20
0
20
10
-10
-20

mean left->right -1.0 mm

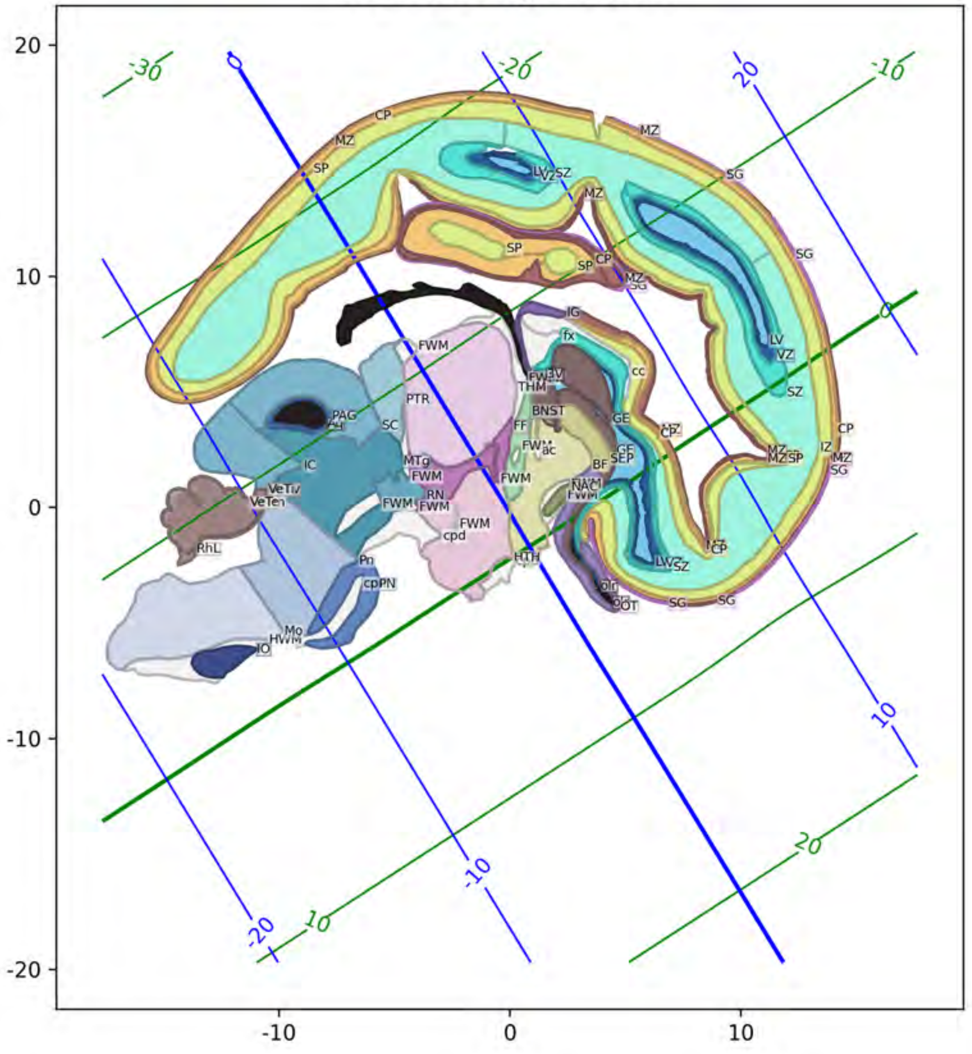

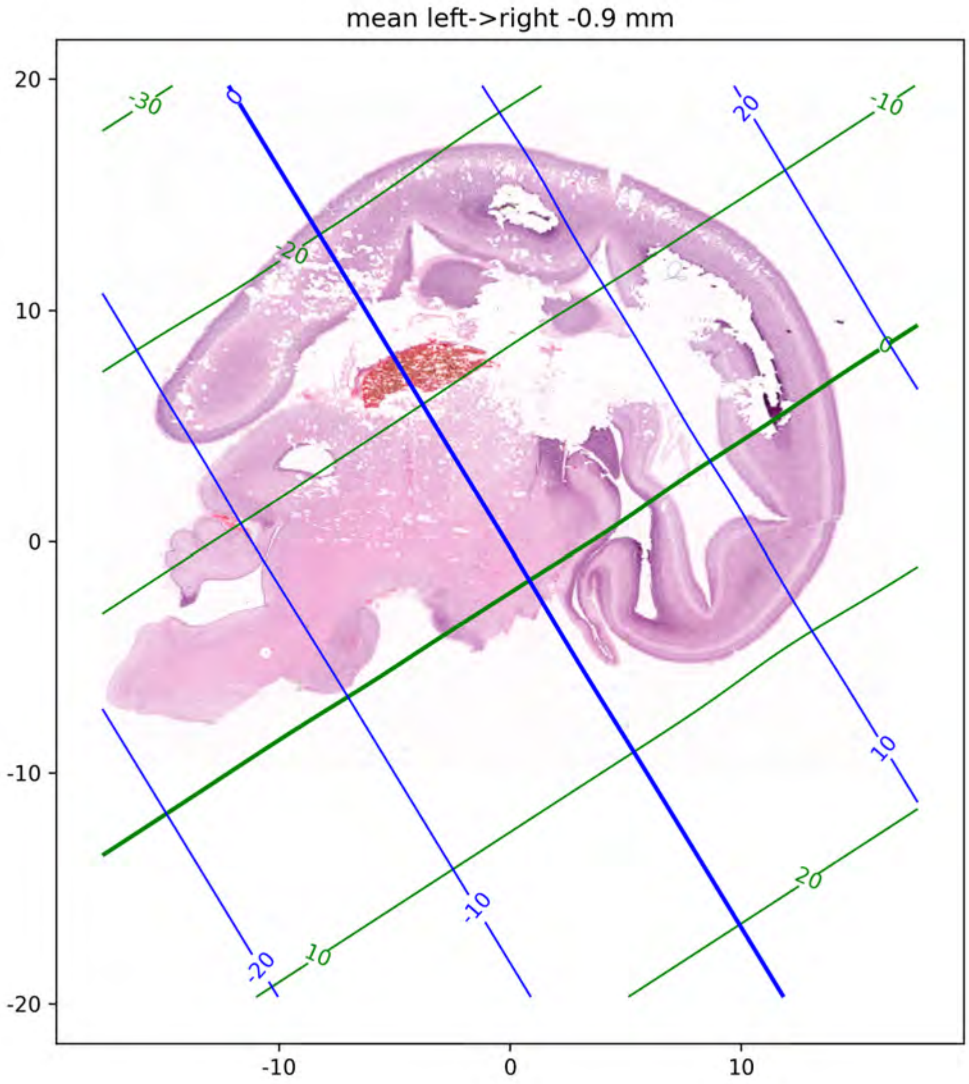

mean left->right -0.9 mm
20
10
0
-10
-20
-10
0
10
-30
-20
-10
0
10
20
0
20
10
-10
-20

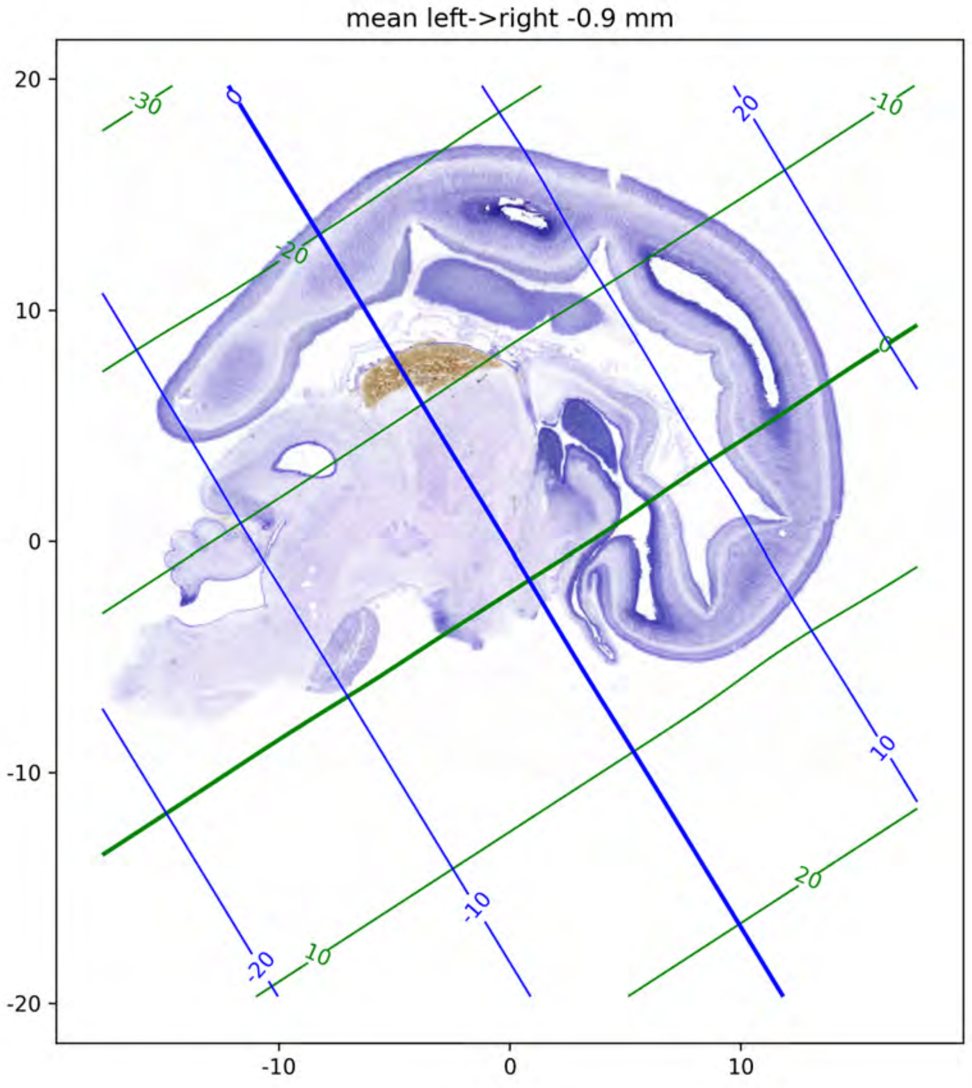

mean left->right -0.9 mm
20
10
0
-10
-20
-10
0
10
-30
-20
-10
0
10
20
0
20
10
-10
-20

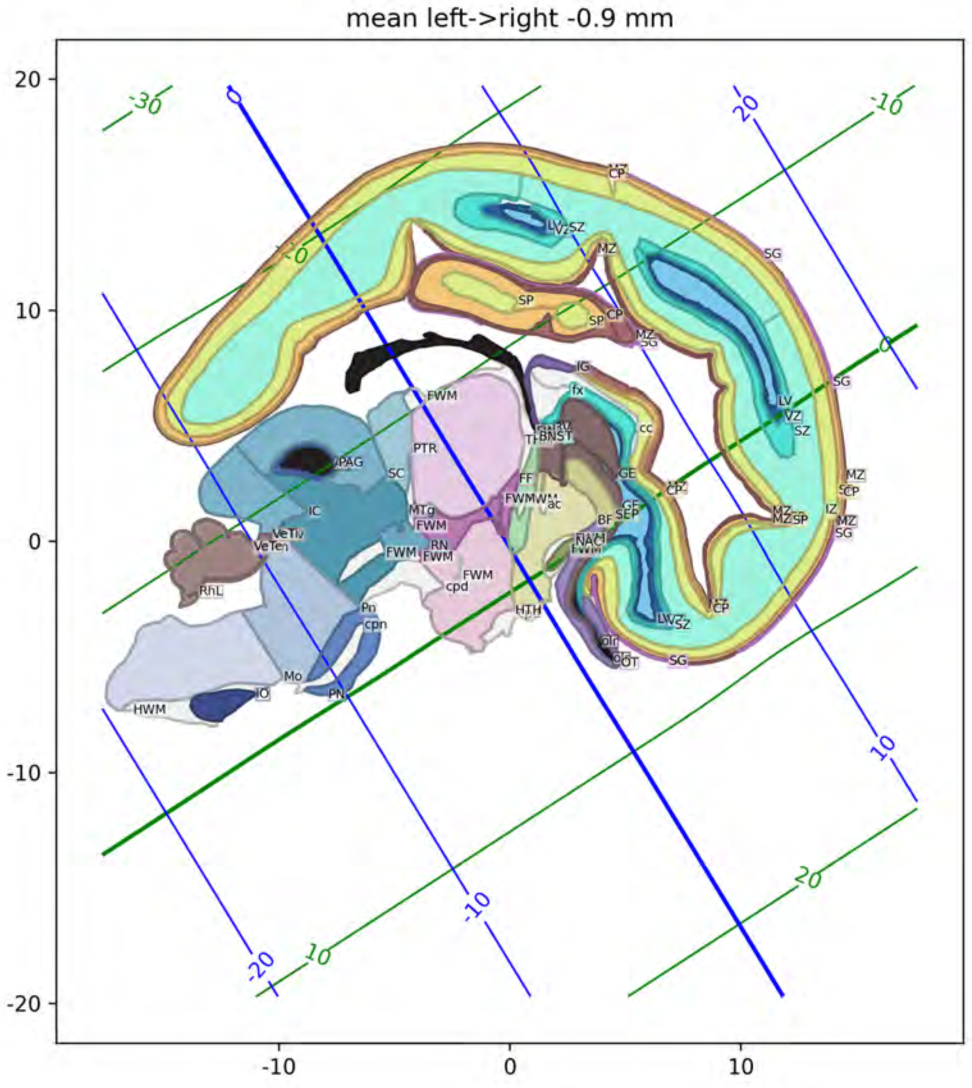
mean left->right -0.9 mm
LV
VZ
SZ
MZ
CP
SG
SP
IG
fx
cc
GE
BNST
TH
FF
FWM
ac
BF
SEP
NAC
HTH
OT
PAG
SC
PTR
IC
MTg
RN
cpd
VeTiv
VeTen
RhL
Pn
cpn
Mo
IO
PN
HWM
IZ

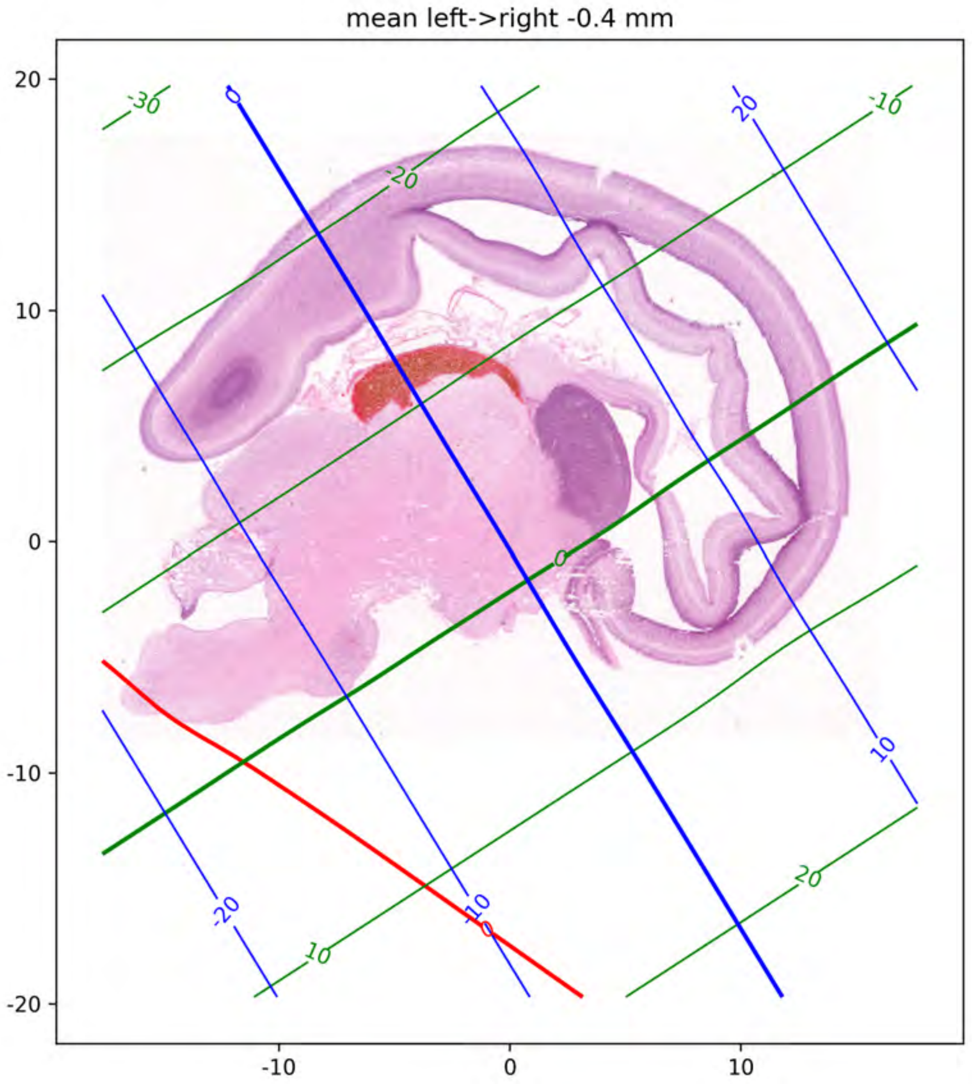
mean left->right -0.4 mm
20
10
0
-10
-20
-10
0
10
-30
-20
-10
0
10
20
0
20
10
-20
-10
0

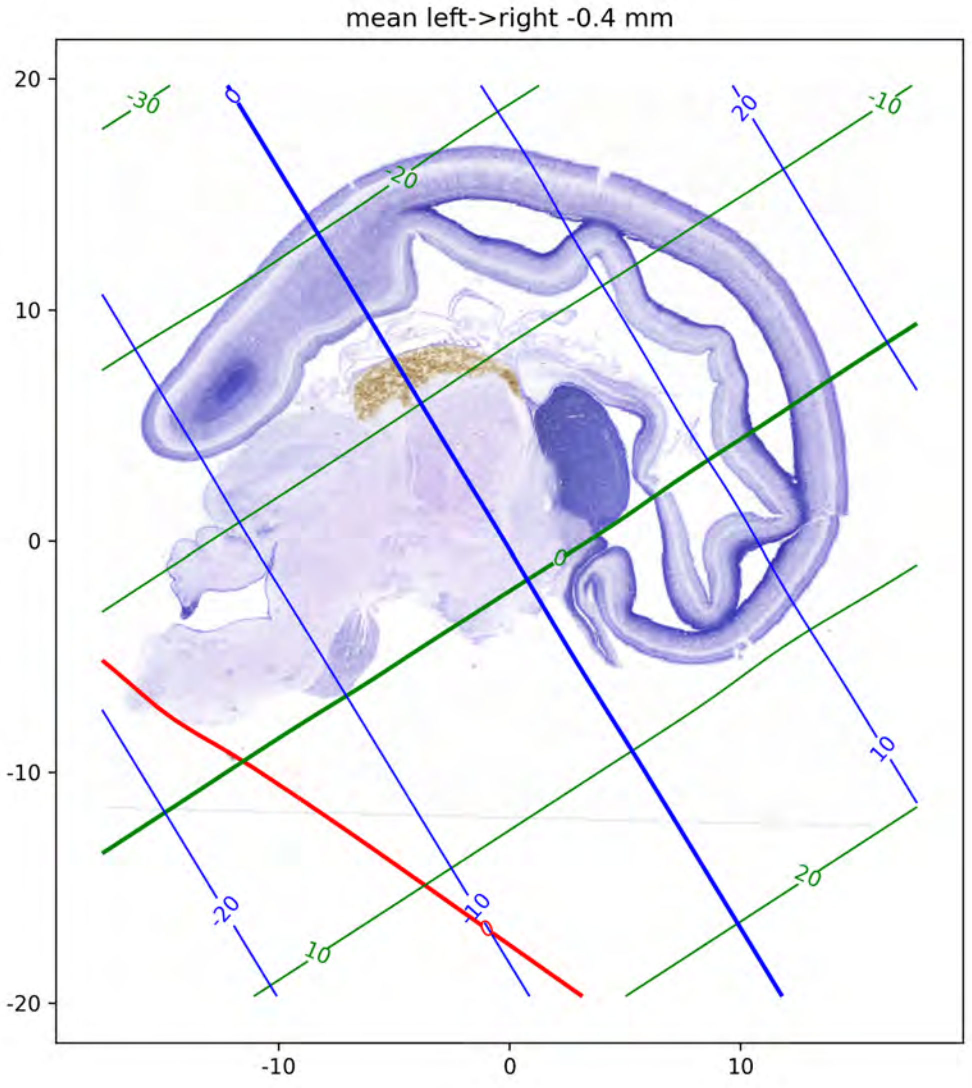
mean left->right -0.4 mm
20
10
0
-10
-20
-10
0
10
-30
-20
-10
0
10
20
0
10
20
-20
-10
10
0

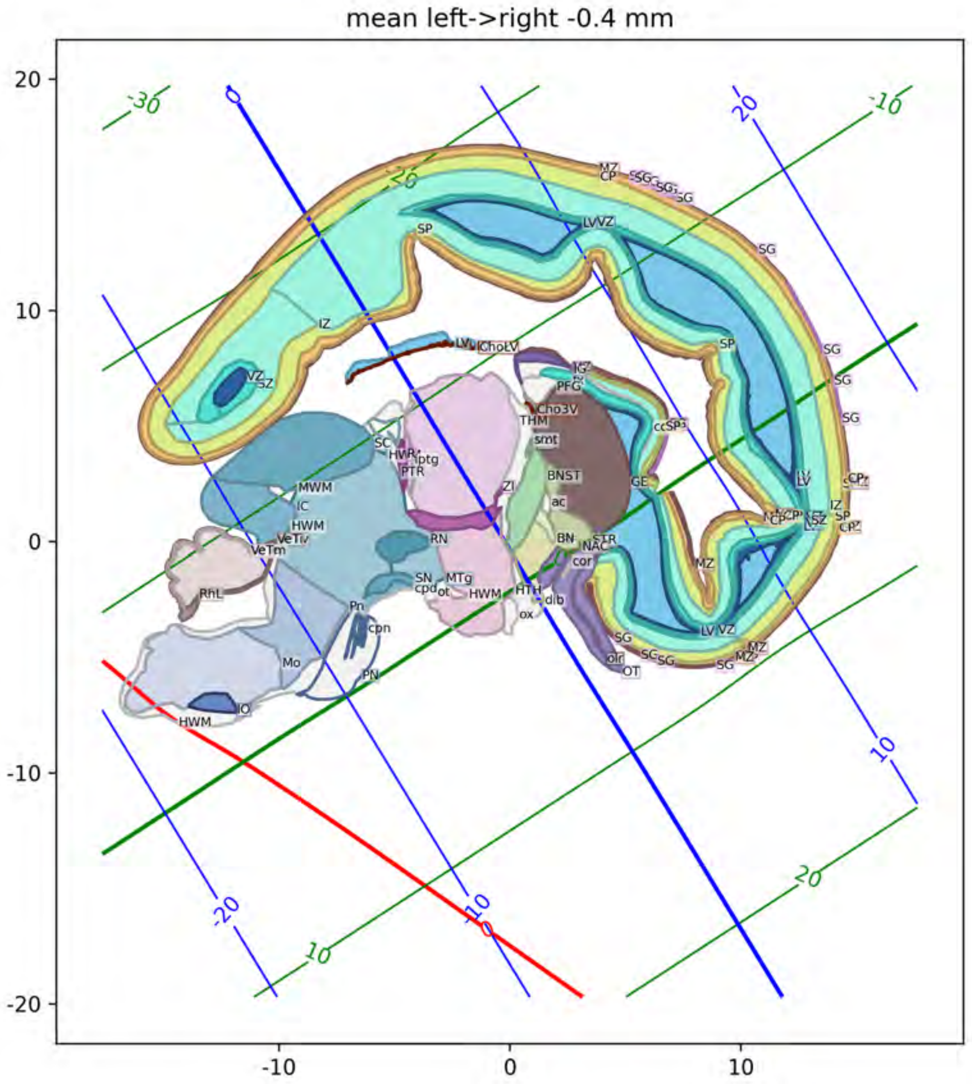
mean left->right -0.4 mm
20
10
0
-10
-20
-10
0
10

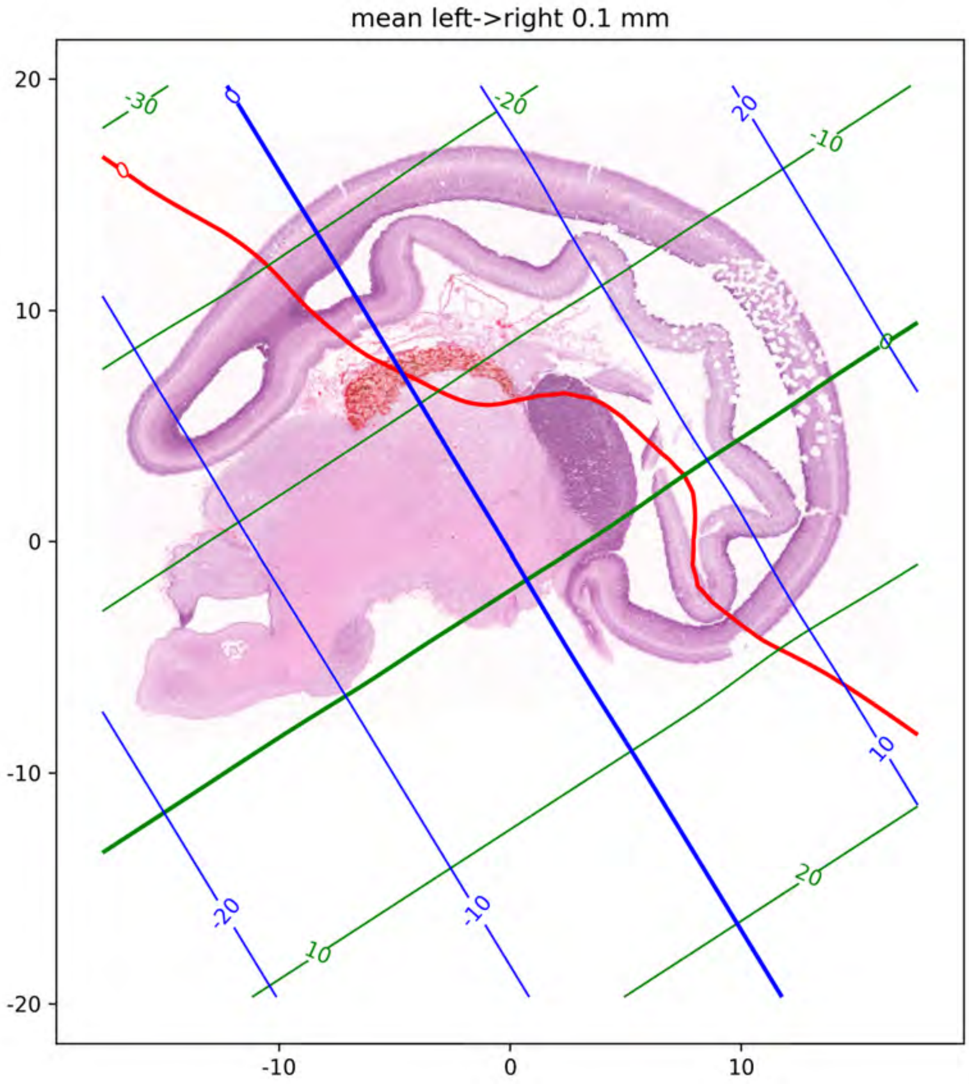
mean left->right 0.1 mm
-30
0
-20
20
-10
0
10
-20
-10
10
20
20
10
0
-10
-20
-10
0
10

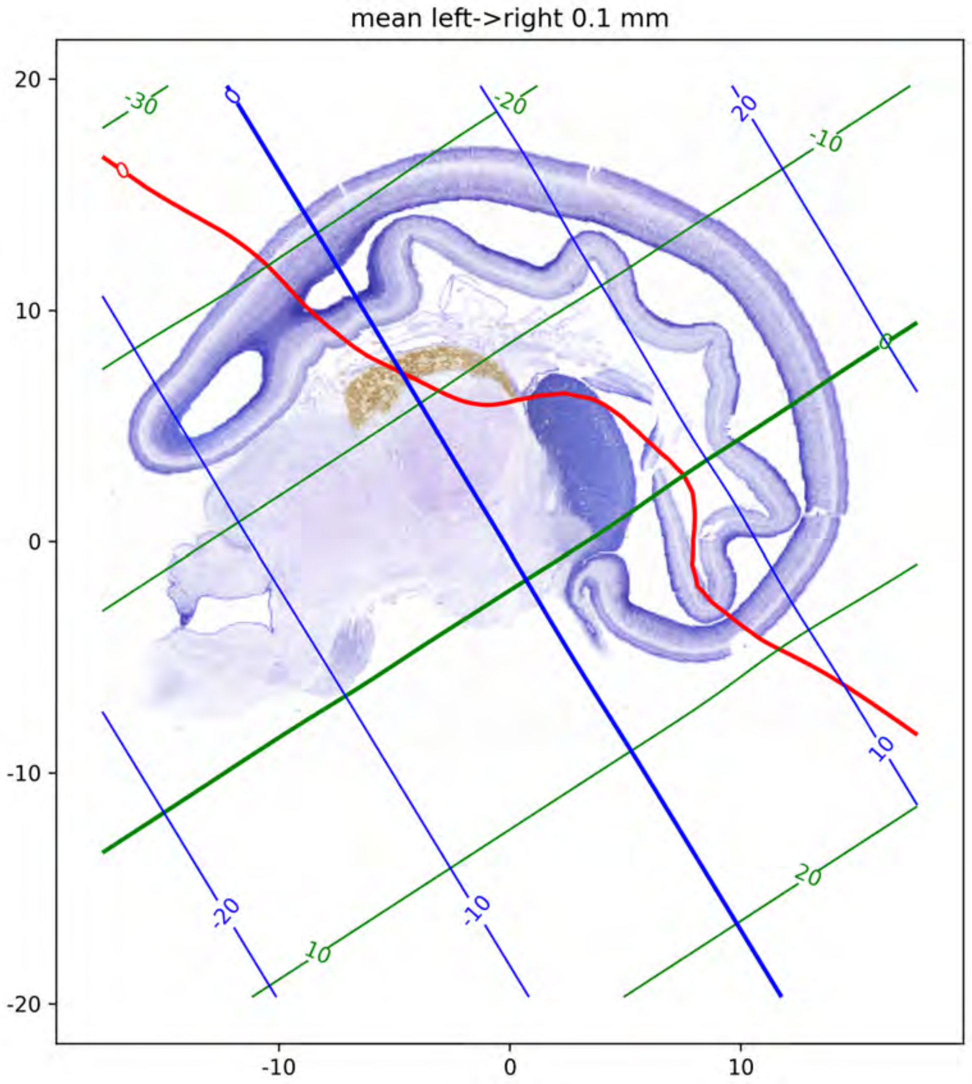
mean left->right 0.1 mm
20
10
0
-10
-20
-10
0
10
-30
-20
-10
0
10
20
0
-20
-10
10
20

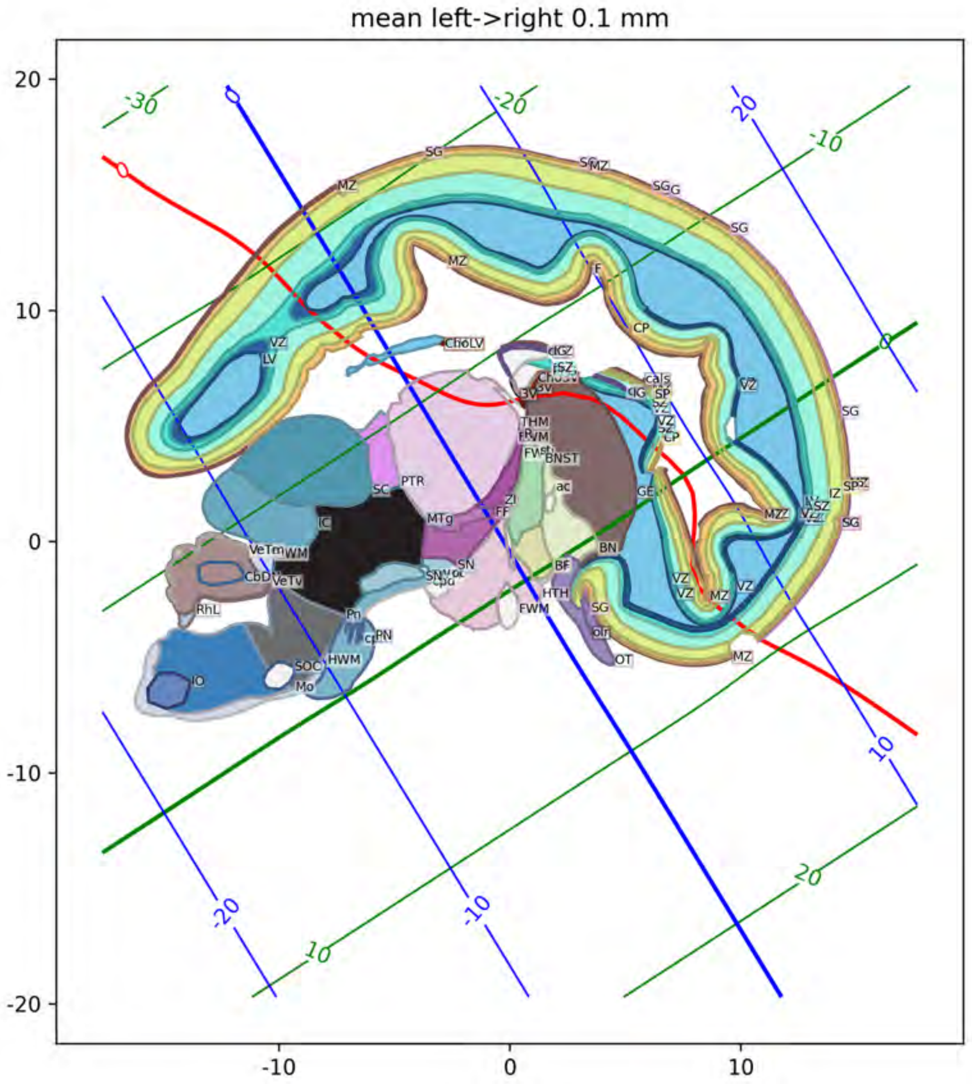
mean left->right 0.1 mm
20
10
0
-10
-20
-10
0
10
-30
-20
-10
0
10
20
SG
MZ
VZ
LV
CP
ChoLV
THM
BNST
ac
GE
BN
BF
HTH
FWM
OT
SC
PTR
MTg
ZI
FF
SN
IC
VeTm
WM
CbD
VeTv
RhL
Pn
PN
HWM
SOC
IO
Mo

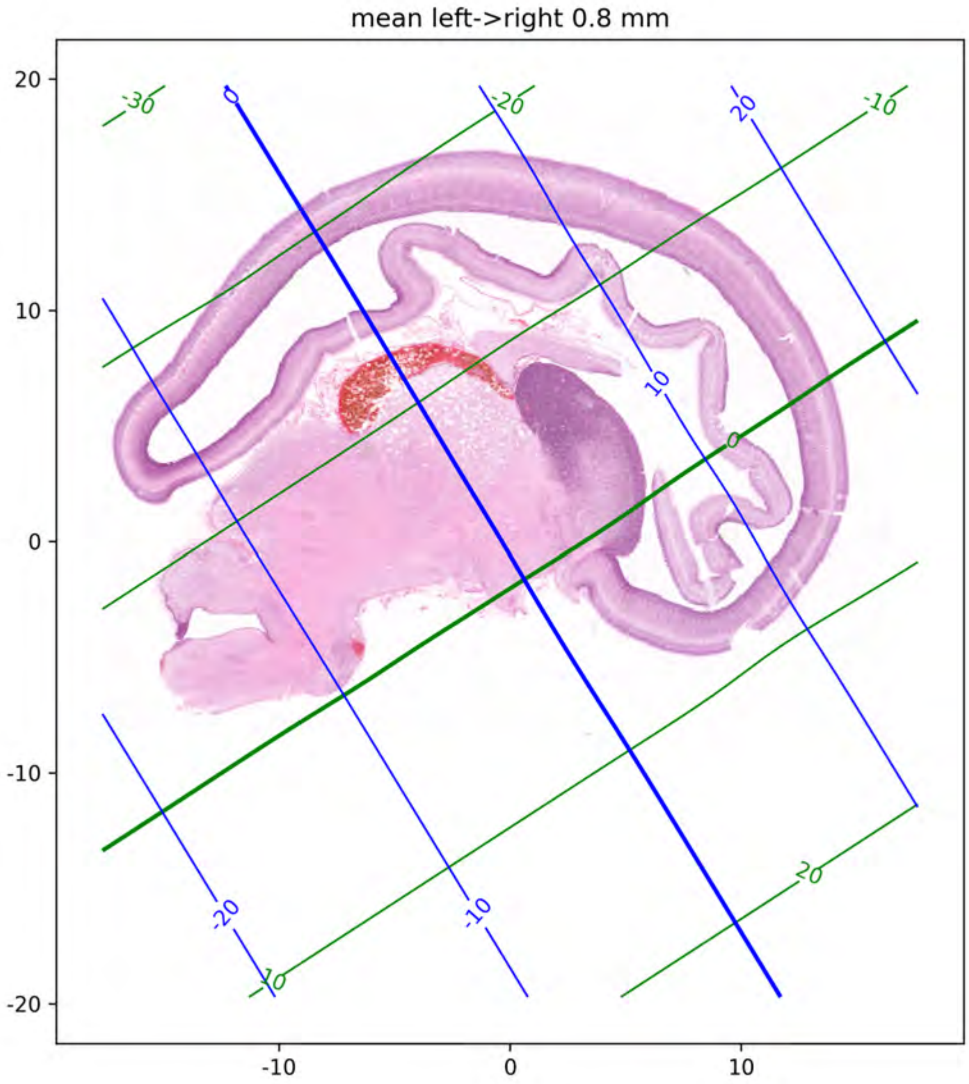
mean left->right 0.8 mm
20
10
0
-10
-20
-10
0
10
-30
-20
-10
0
10
20
0
10
20
-20
-10

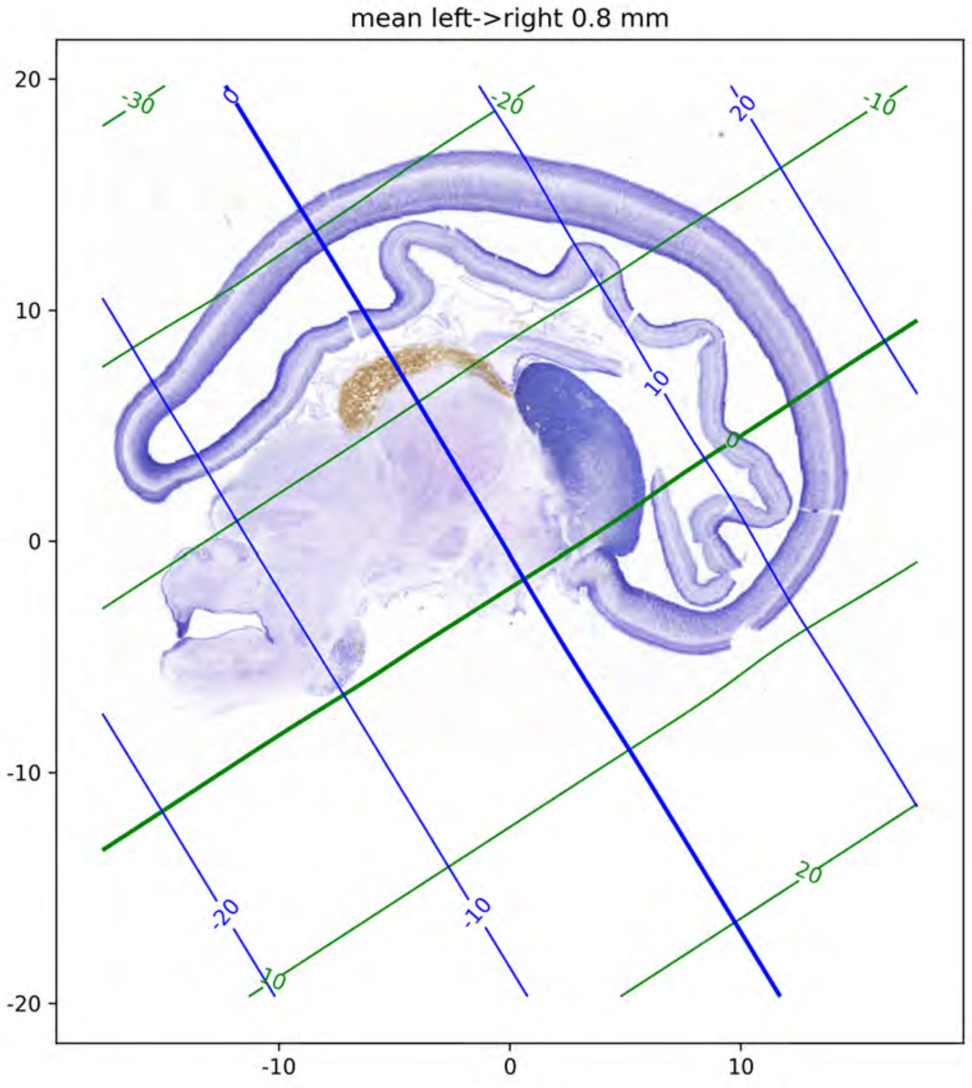

mean left->right 0.8 mm
20
10
0
-10
-20
-10
0
10
-30
-20
-10
0
10
20
20
-20
-10

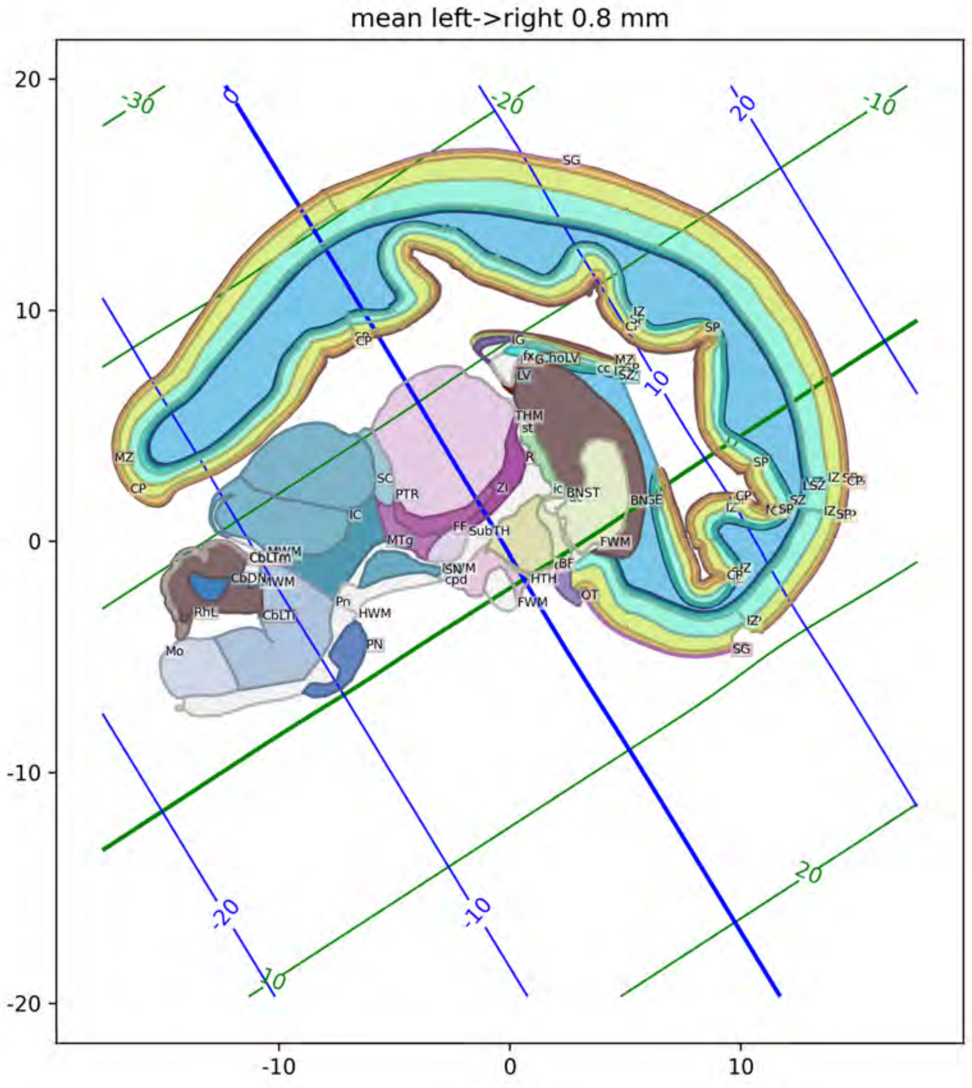

mean left->right 0.8 mm
20
10
0
-10
-20
-10
0
10
-30
-20
-10
20
10
SG
SP
CP
MZ
IG
fx
LV
cc
THM
st
R
Zi
SC
PTR
IC
MTg
FF
SubTH
BNST
FWM
HTH
OT
Pn
HWM
PN
Mo
RhL
CbLTi
CbDN
Tm

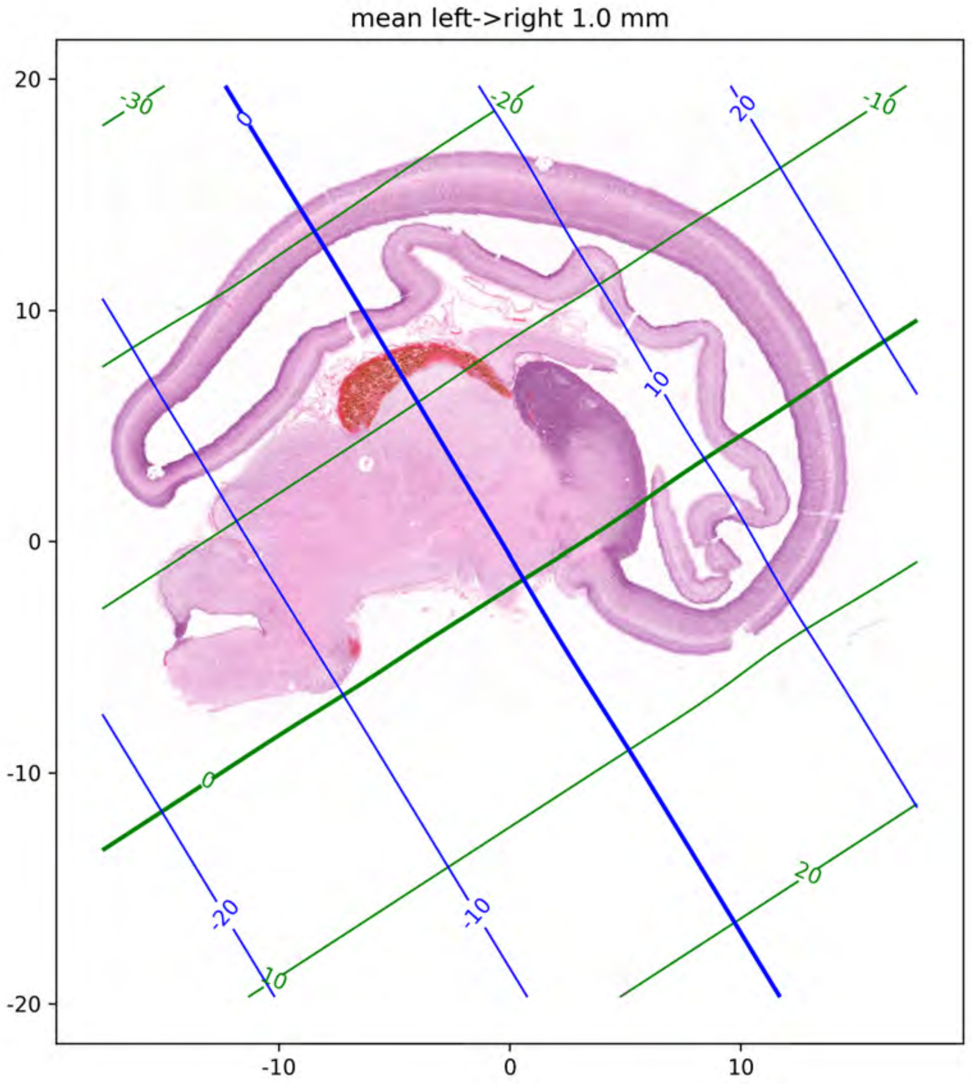

mean left->right 1.0 mm
20
10
0
-10
-20
-10
0
10
-30
-20
-10
0
10
20
0
10
20
-20
-10

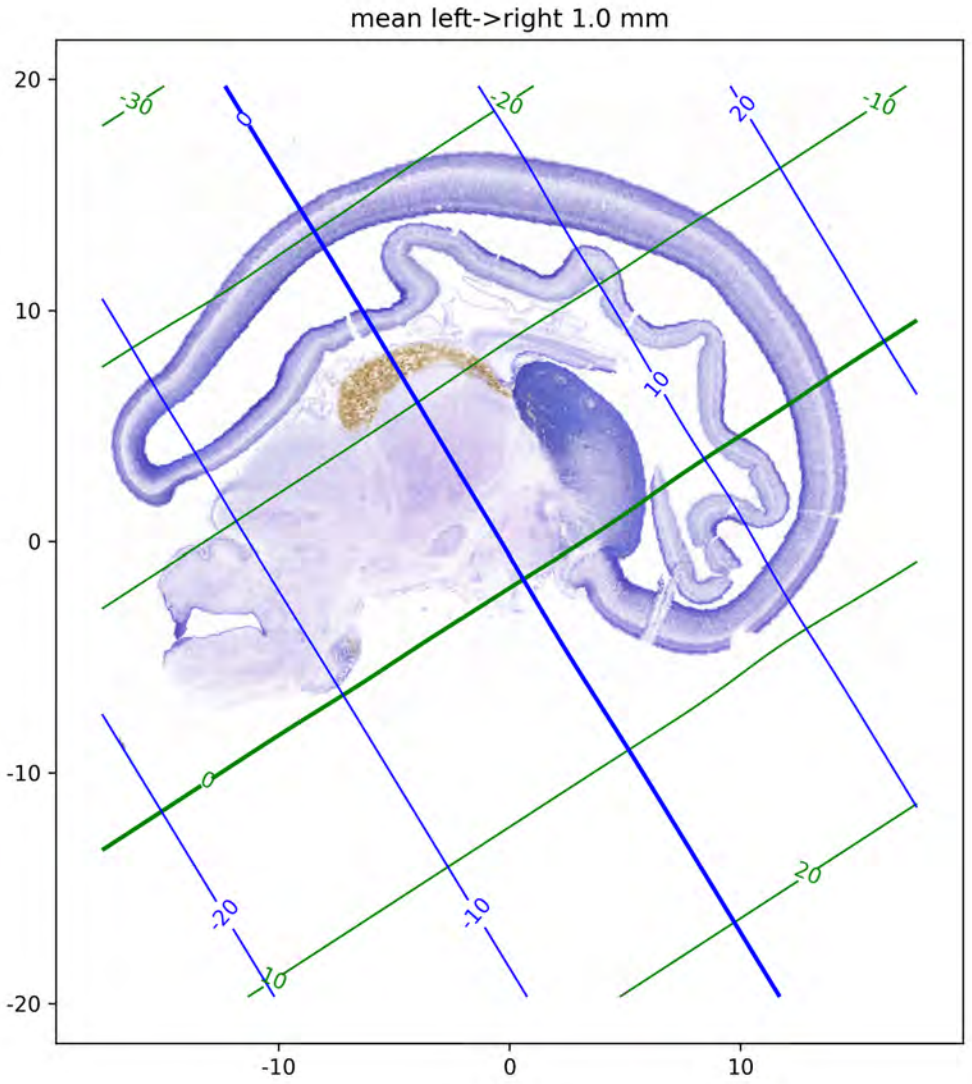

mean left->right 1.0 mm
20
10
0
-10
-20
-10
0
10
-30
-20
-10
0
10
20
0
20
10
-20
-10

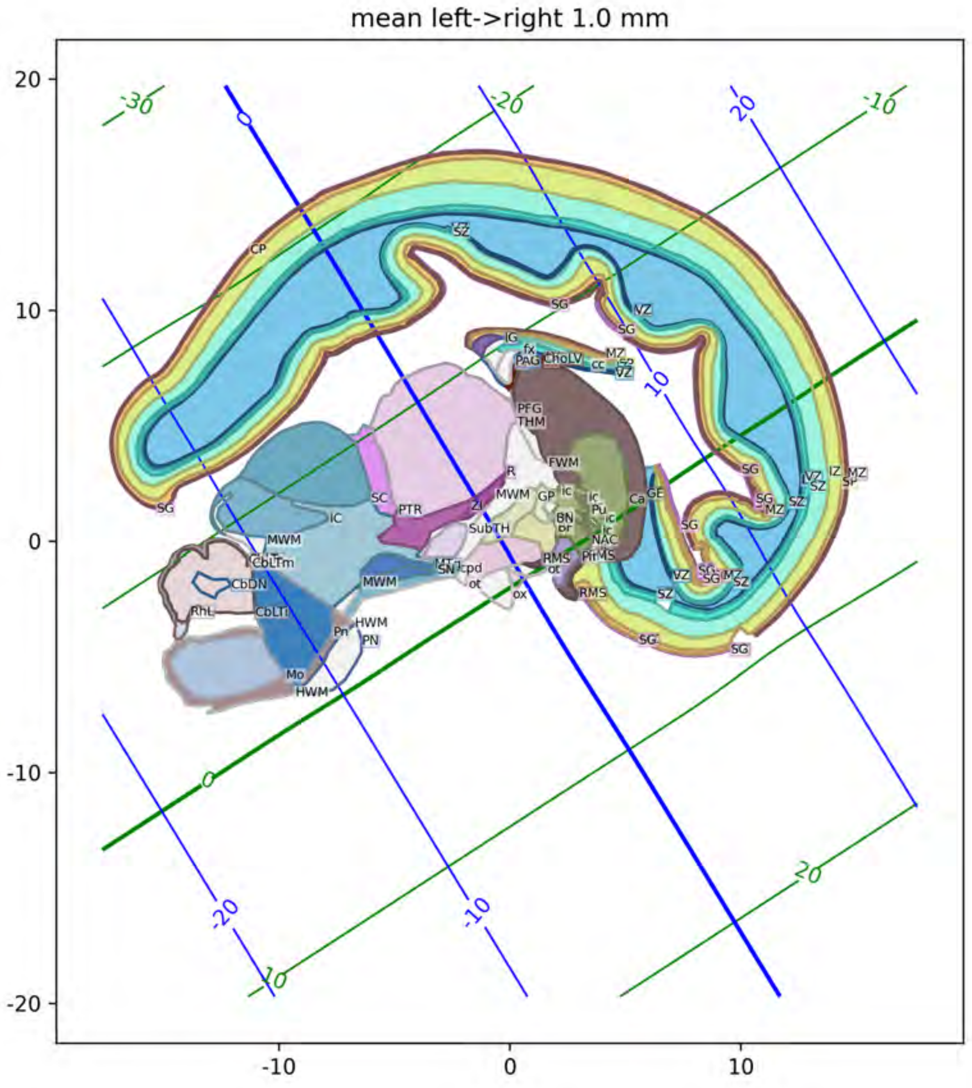
mean left->right 1.0 mm
CP
SZ
SG
VZ
IG
fx
PAG
ChoLV
cc
MZ
PFG
THM
FWM
R
MWM
GP
BN
Pu
NAC
Ca
GE
SC
PTR
ZI
SubTH
IC
MWM
CbLTm
CbDN
RhL
CbLTi
SN
cpd
ot
ox
RMS
HWM
Pn
PN
Mo
IZ
SP

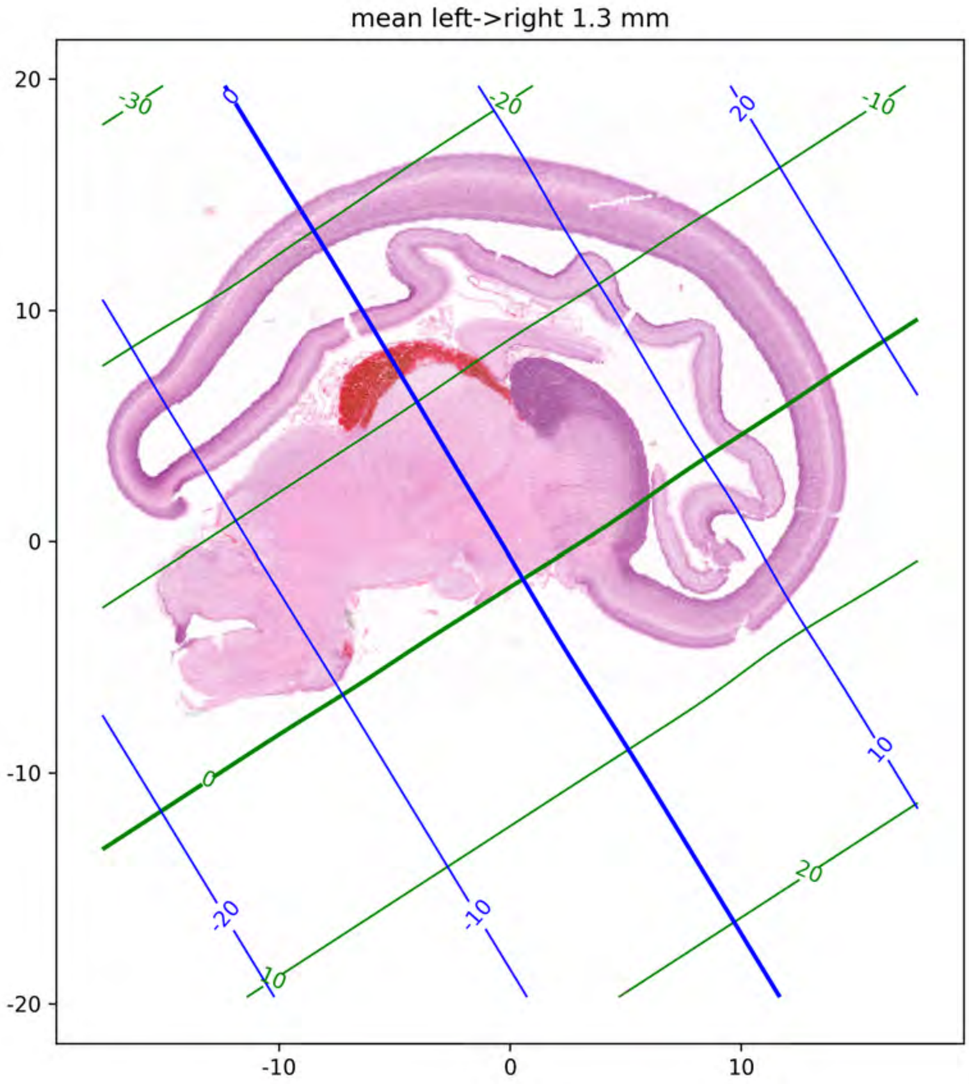
mean left->right 1.3 mm
20
10
0
-10
-20
-10
0
10
-30
-20
-10
0
10
20
0
-20
20
10
-20
-10

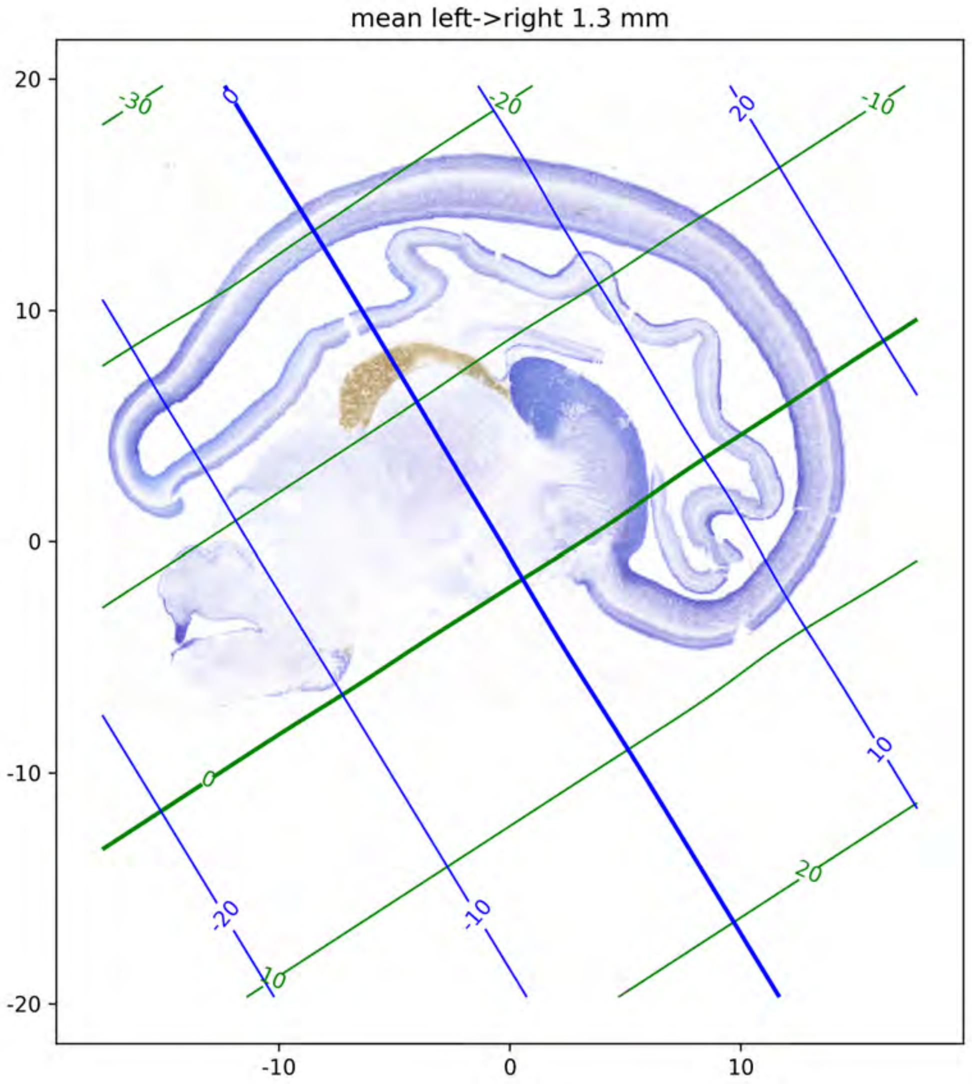

mean left->right 1.3 mm
20
10
0
-10
-20
-10
0
10
-30
-20
-10
0
10
20
0
-20
20
-10
10
-20

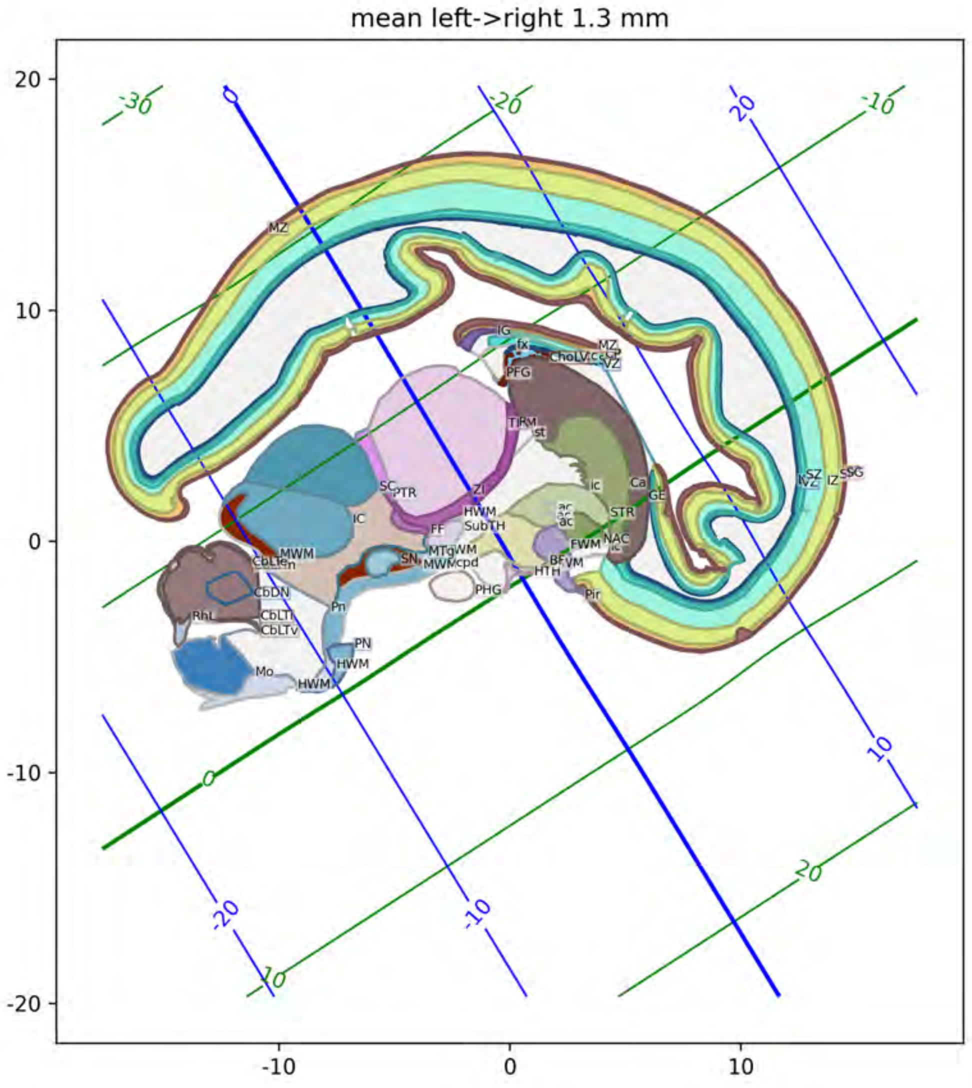

mean left->right 1.3 mm

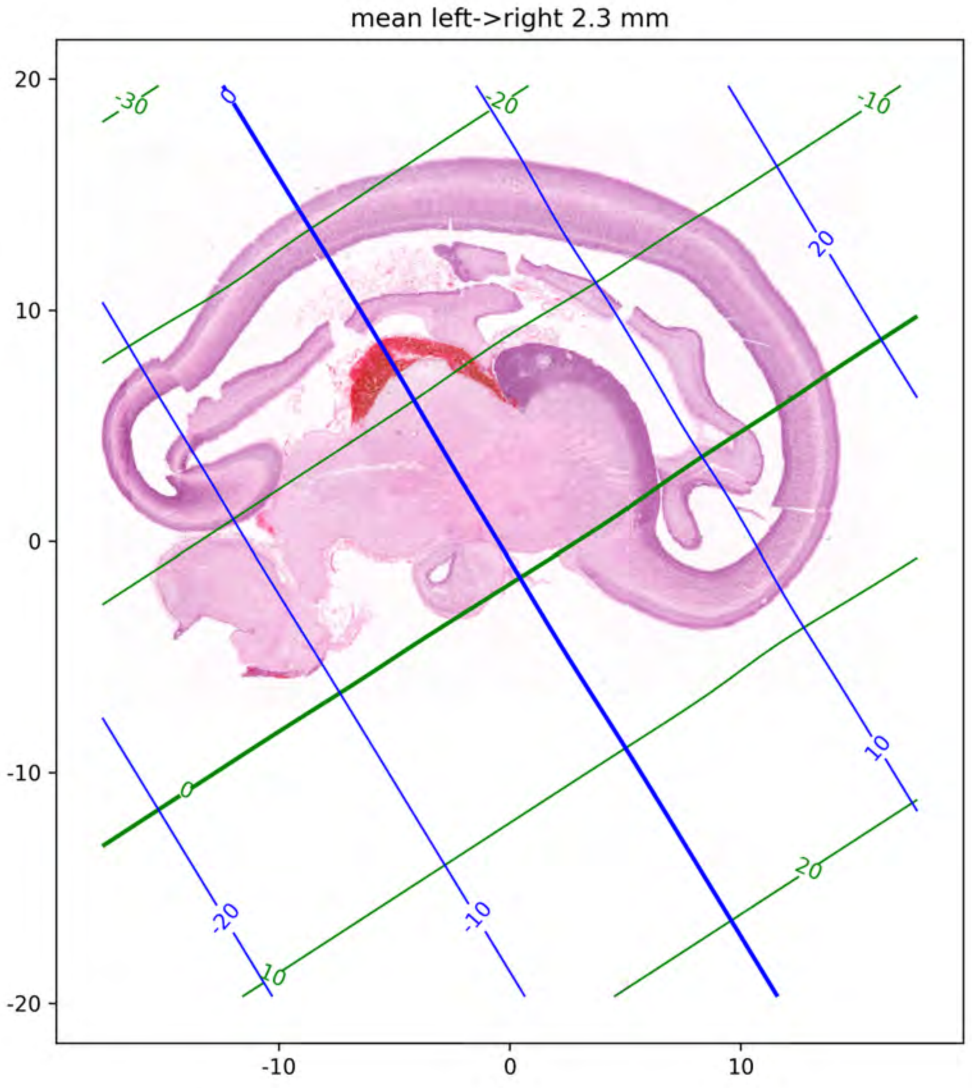
mean left->right 2.3 mm
20
10
0
-10
-20
-10
0
10
-30
-20
-10
0
10
20
0
20
10
-10
-20

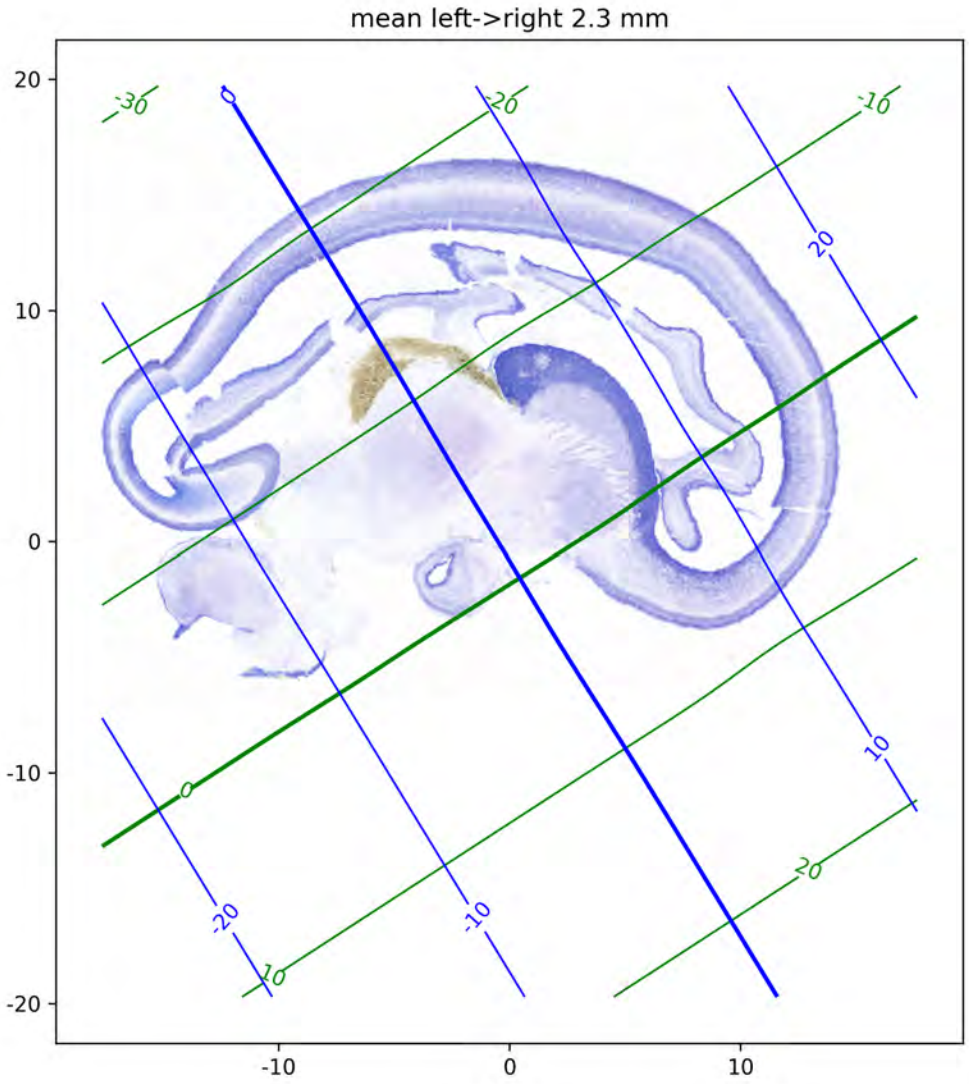
mean left->right 2.3 mm
20
10
0
-10
-20
-10
0
10
-30
-20
-10
0
10
20
0
20
10
-20
-10

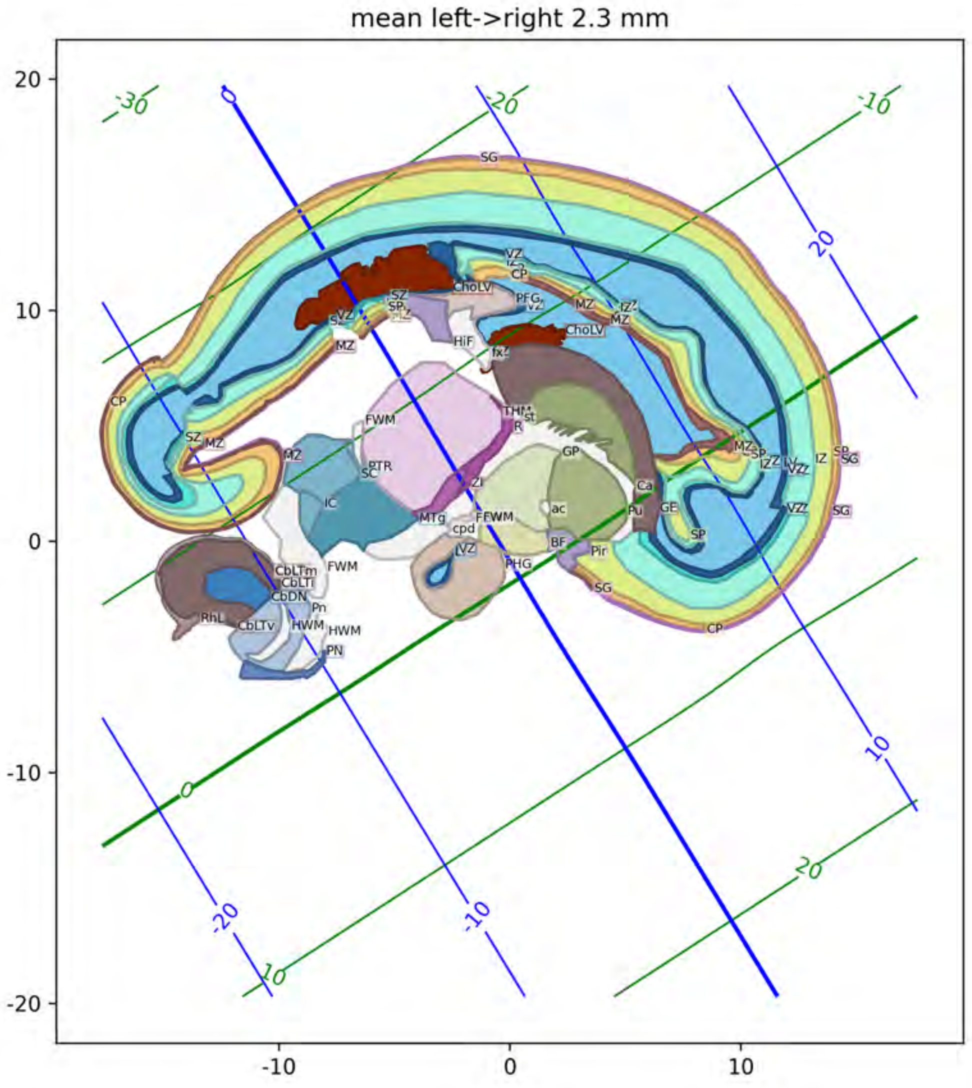
mean left->right 2.3 mm

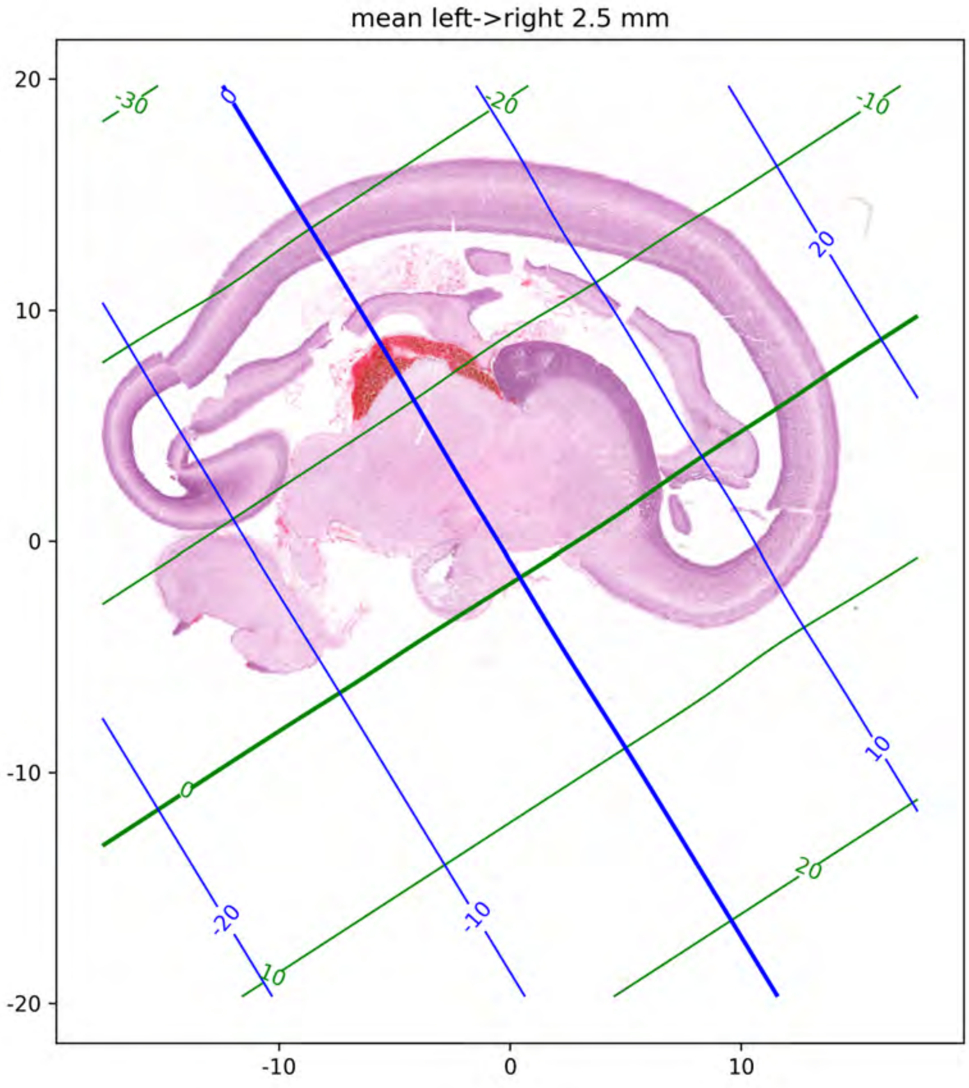

mean left->right 2.5 mm
20
10
0
-10
-20
-10
0
10
-30
-20
-10
0
10
20

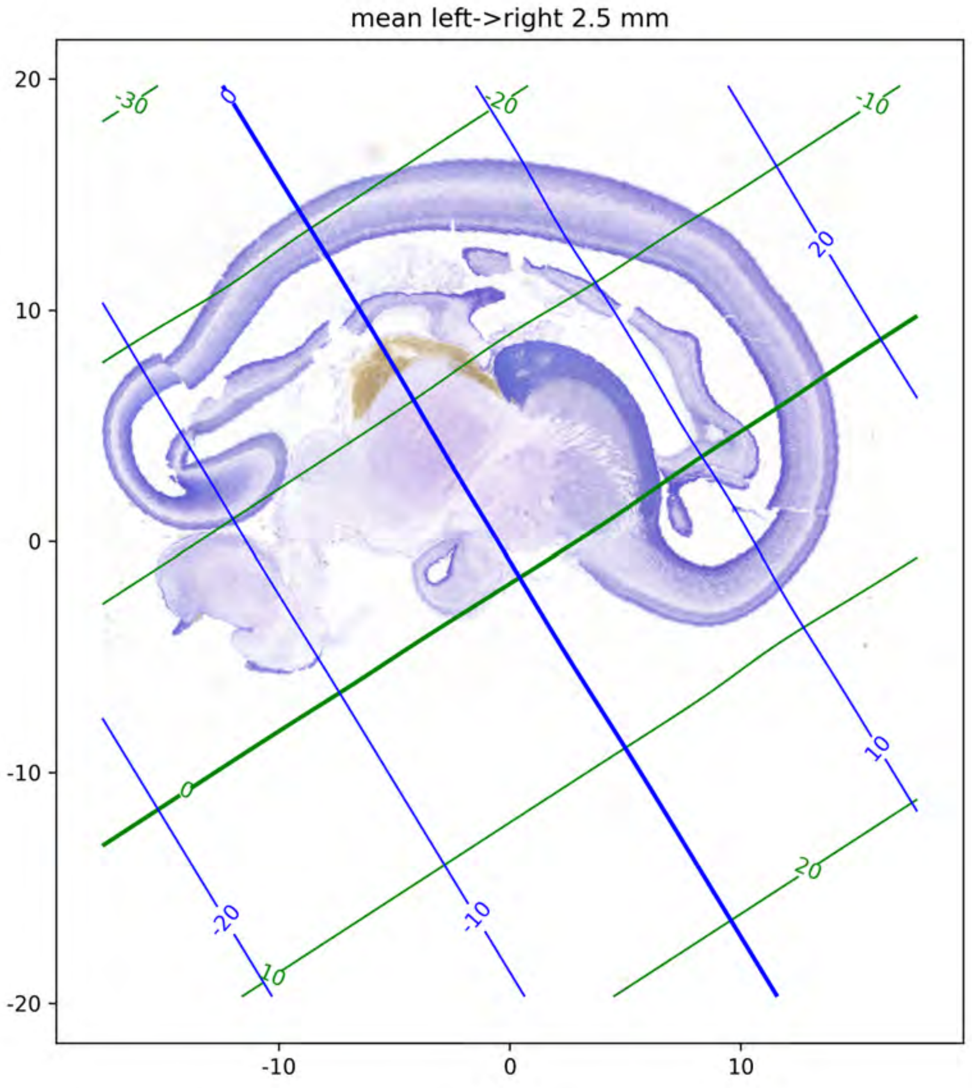
mean left->right 2.5 mm
20
10
0
-10
-20
-10
0
10
-30
-20
-10
0
10
20

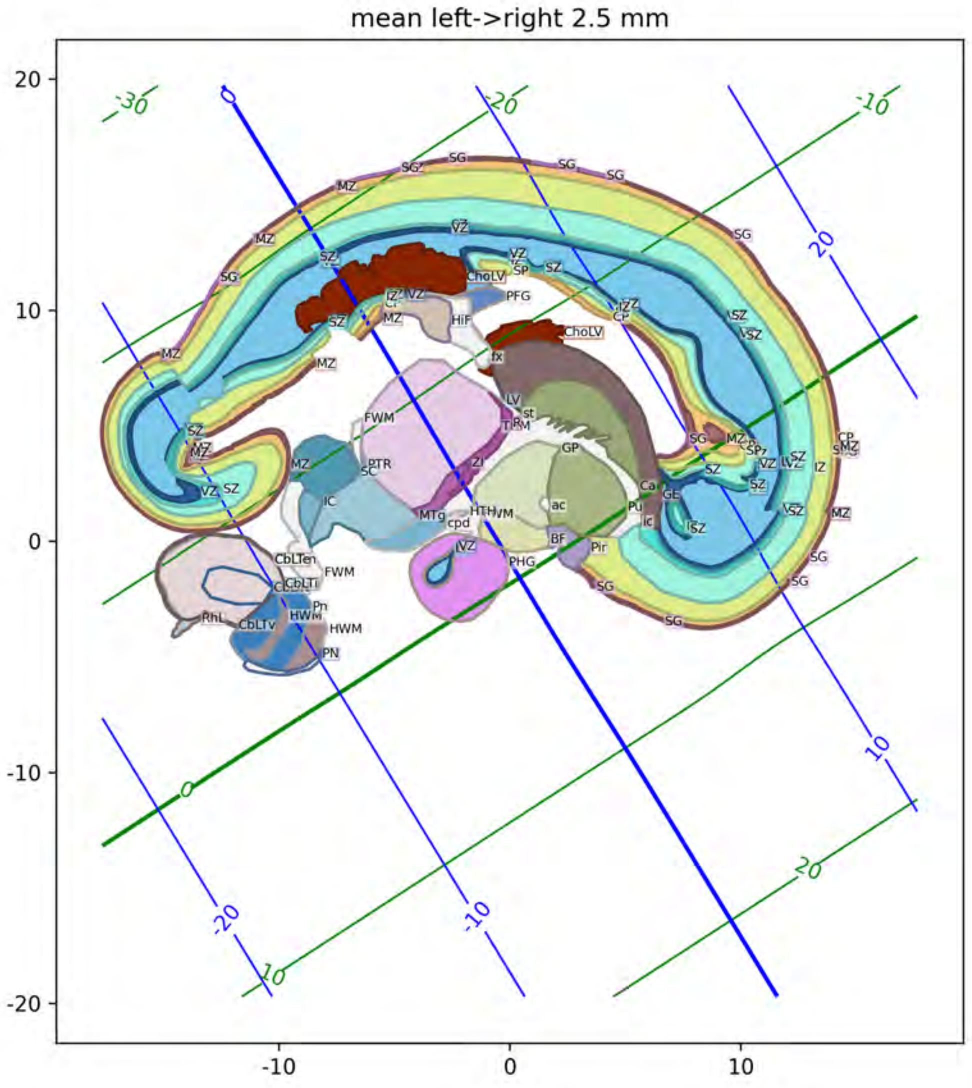
mean left->right 2.5 mm

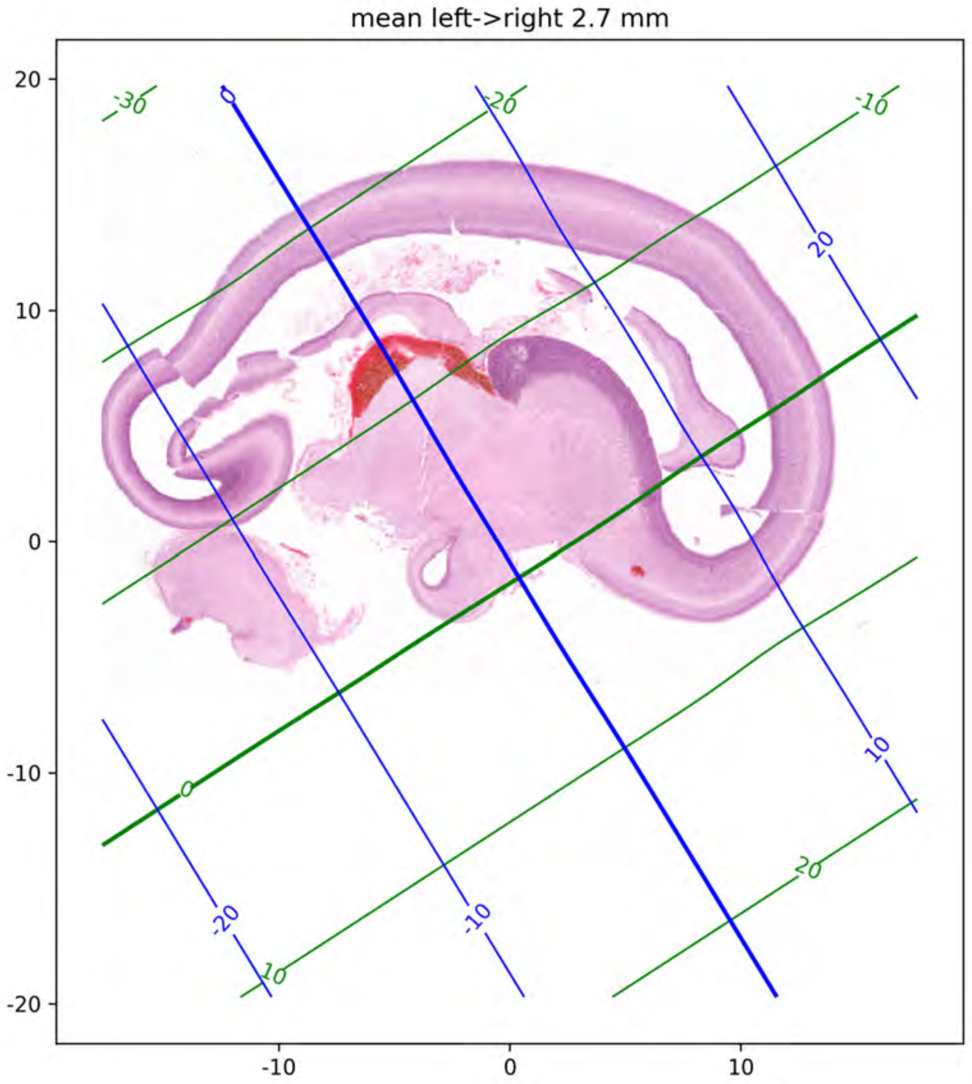
mean left->right 2.7 mm
20
10
0
-10
-20
-10
0
10
-30
-20
-10
0
10
20
0
-20
-10
20
10
-20
-10
0
10
20

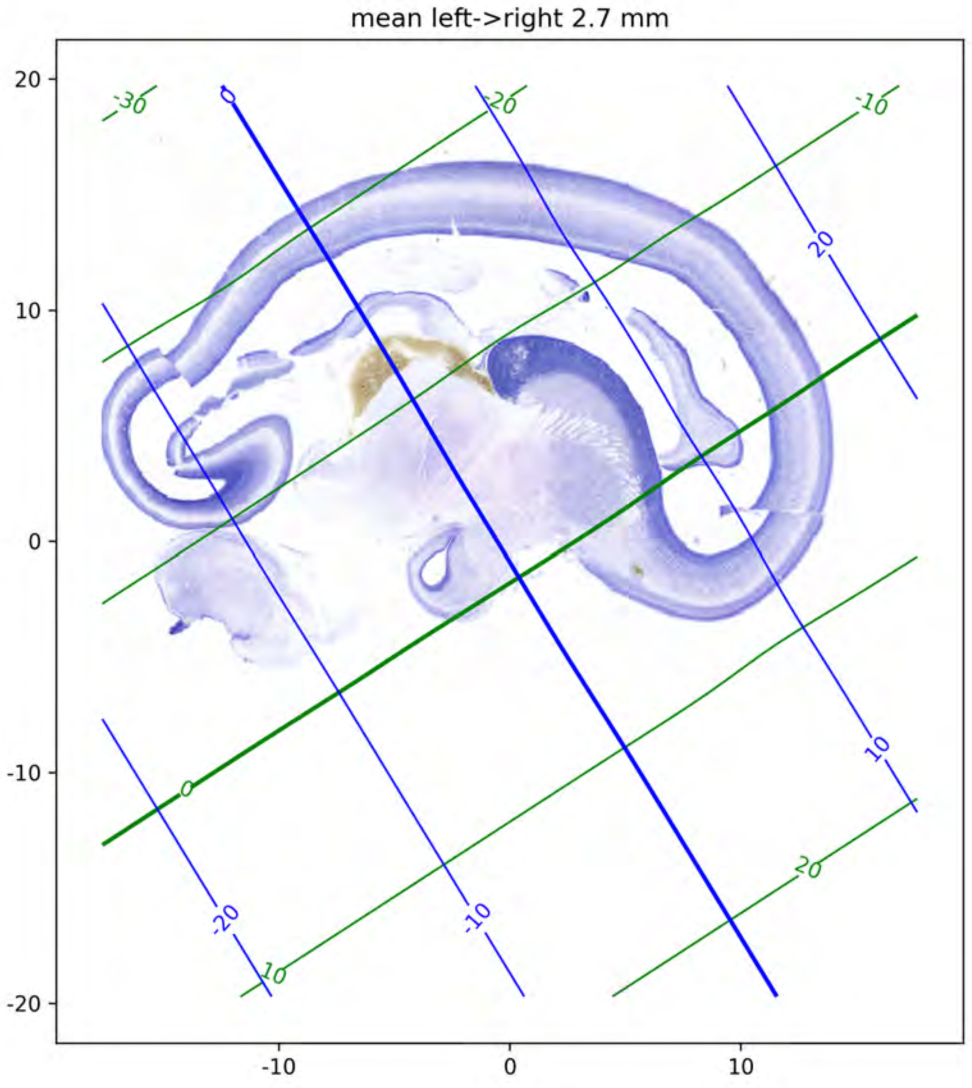

mean left->right 2.7 mm
20
10
0
-10
-20
-10
0
10
-30
-20
-10
0
10
20
0
20
10
-10
-20

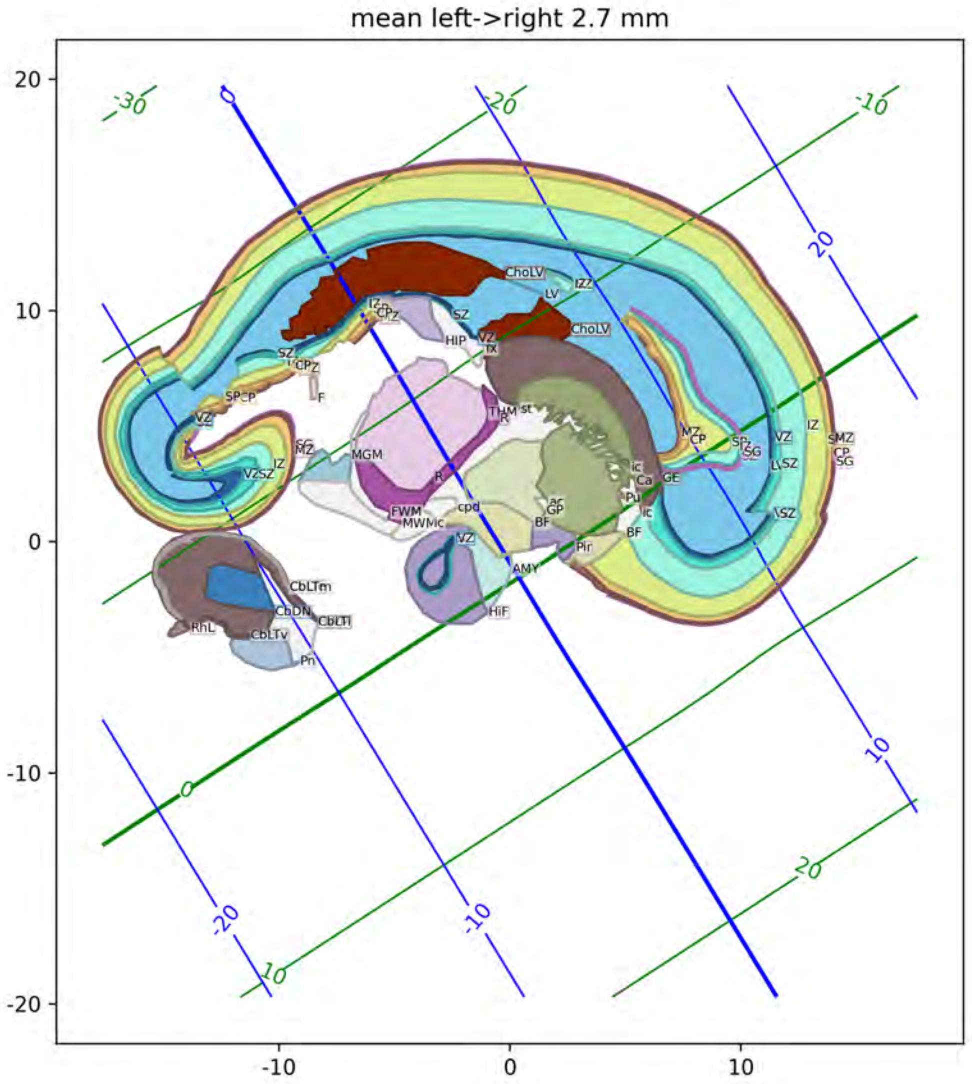
mean left->right 2.7 mm

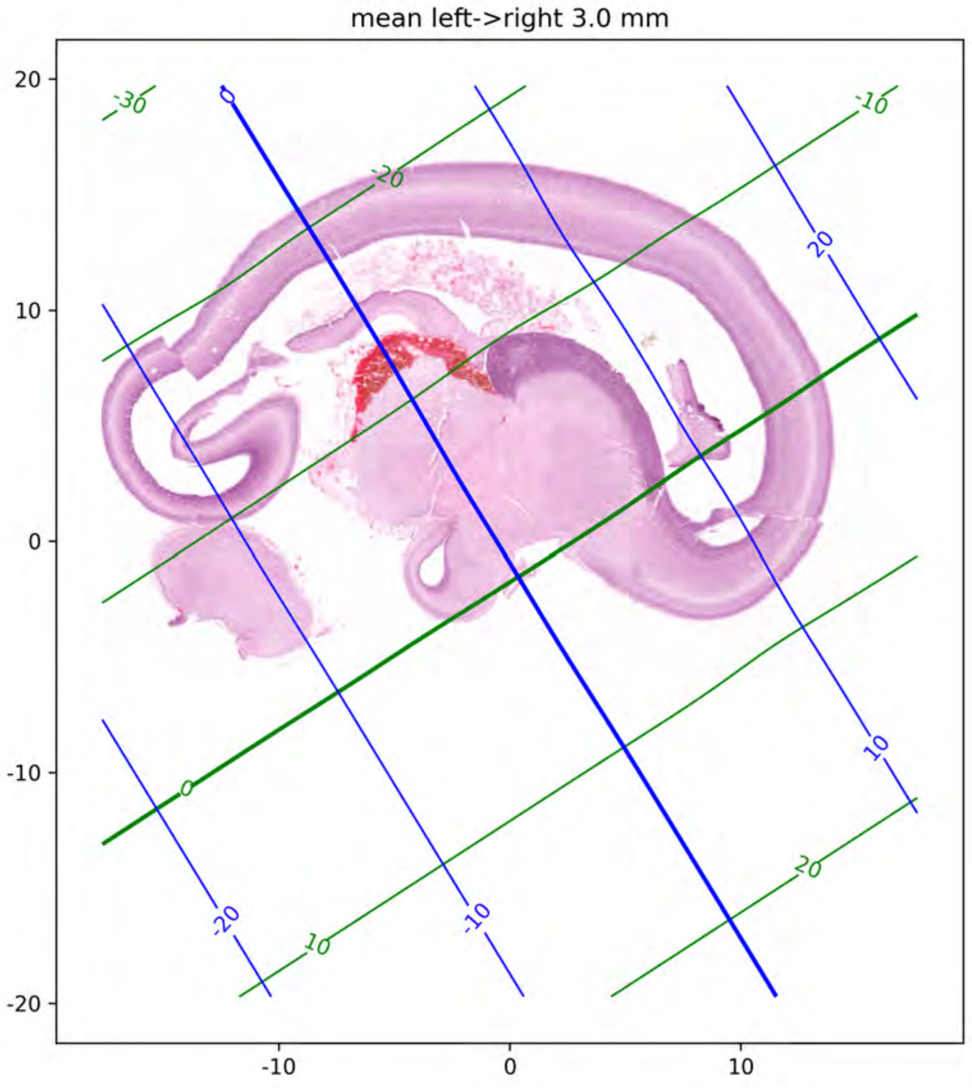
mean left->right 3.0 mm
20
10
0
-10
-20
-10
0
10
-30
-20
-10
0
10
20
0
20
10
-20
-10

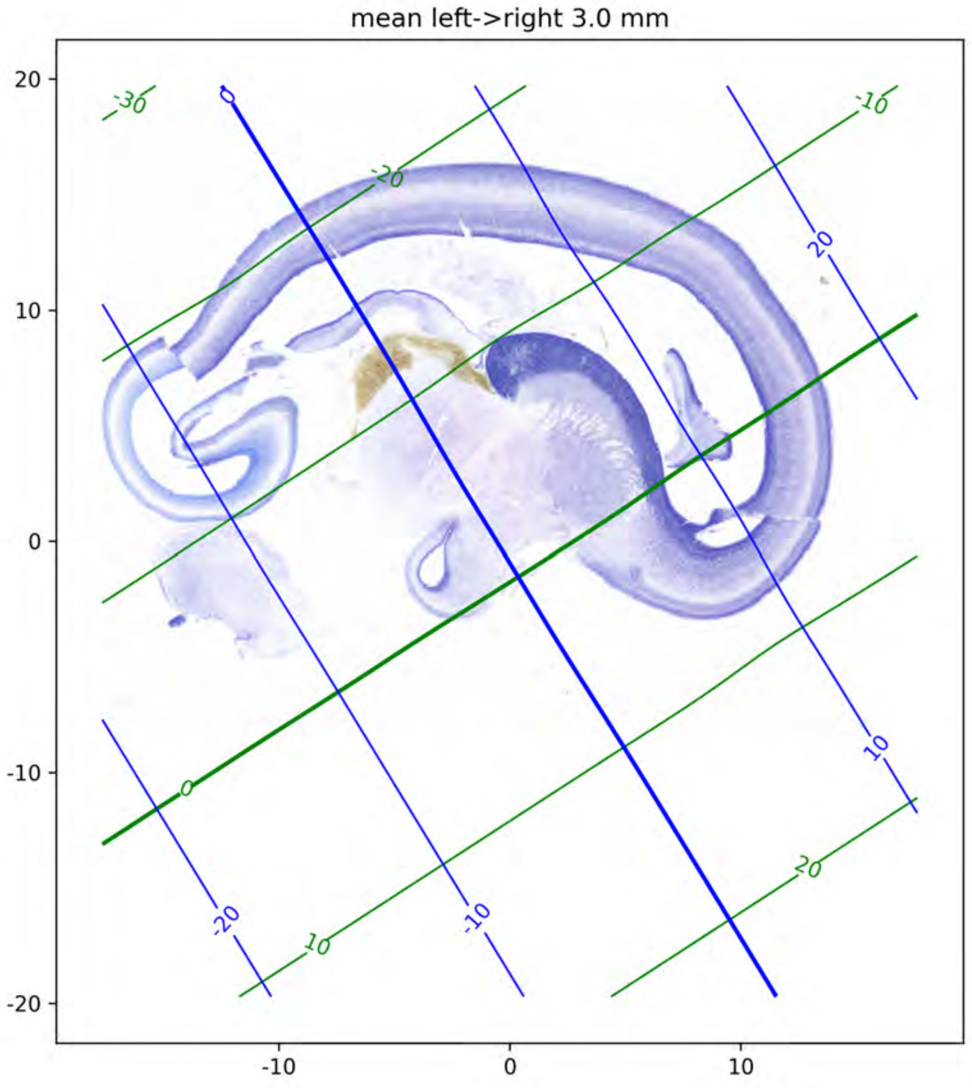
mean left->right 3.0 mm
20
10
0
-10
-20
-10
0
10
-30
-20
-10
0
10
20
0
20
10
-20
-10

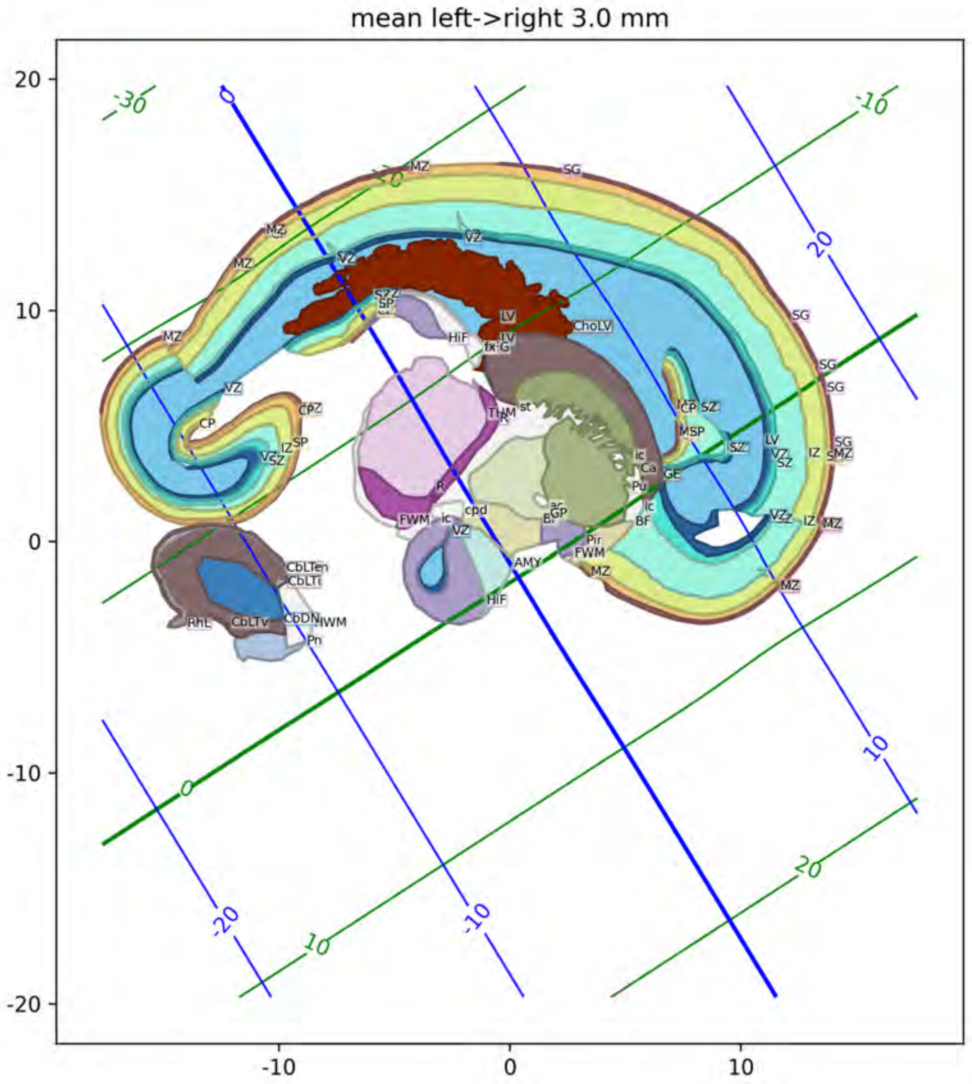
mean left->right 3.0 mm
20
10
0
-10
-20
-10
0
10
-30
-10
-20
-10
10
20
20
MZ
SG
VZ
LV
CholV
HiF
CP
SP
SZ
IZ
MSP
GE
AMY
Pir
FWM
BF
GP
cpd
CbLTen
CbLTi
CbLTv
CbDN
IWM
RhL
Pn

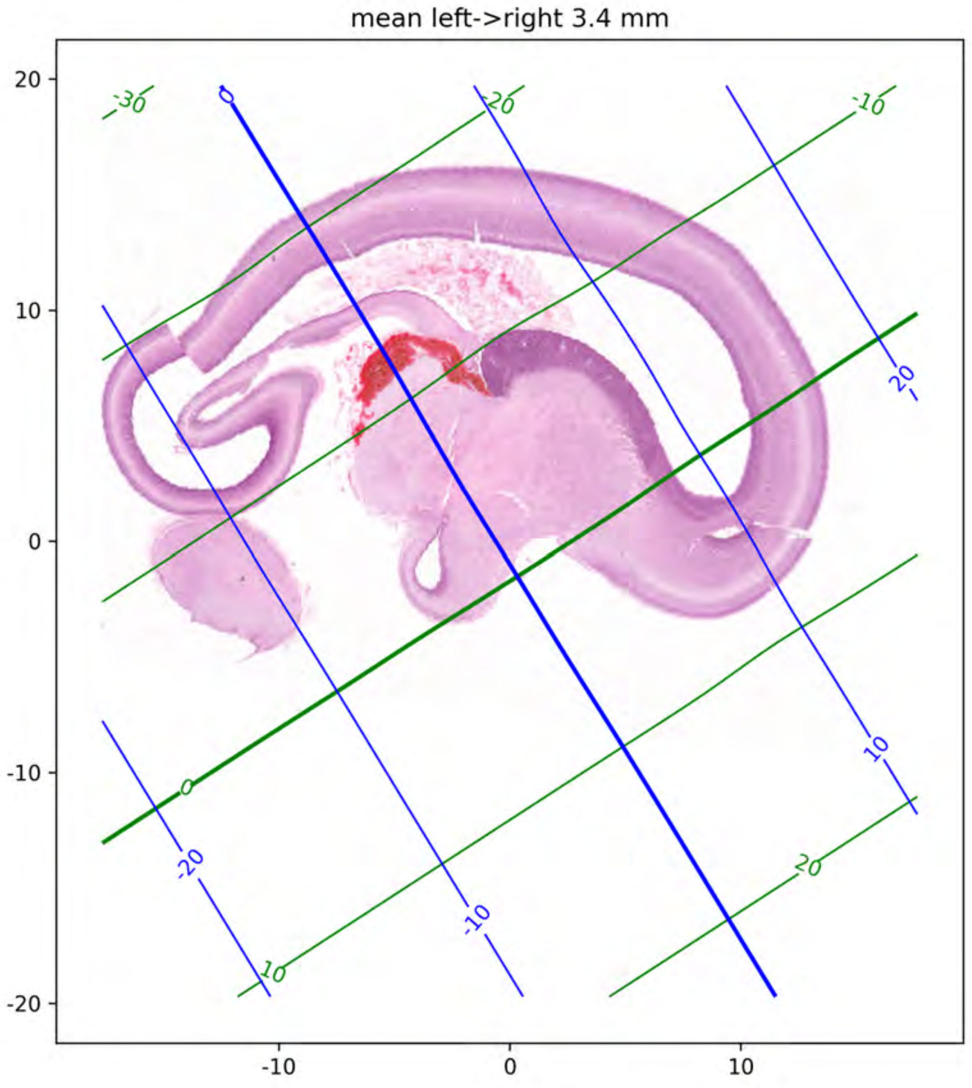
mean left->right 3.4 mm
20
10
0
-10
-20
-10
0
10
-30
-20
-10
0
10
20
0
-20
-10
10
20

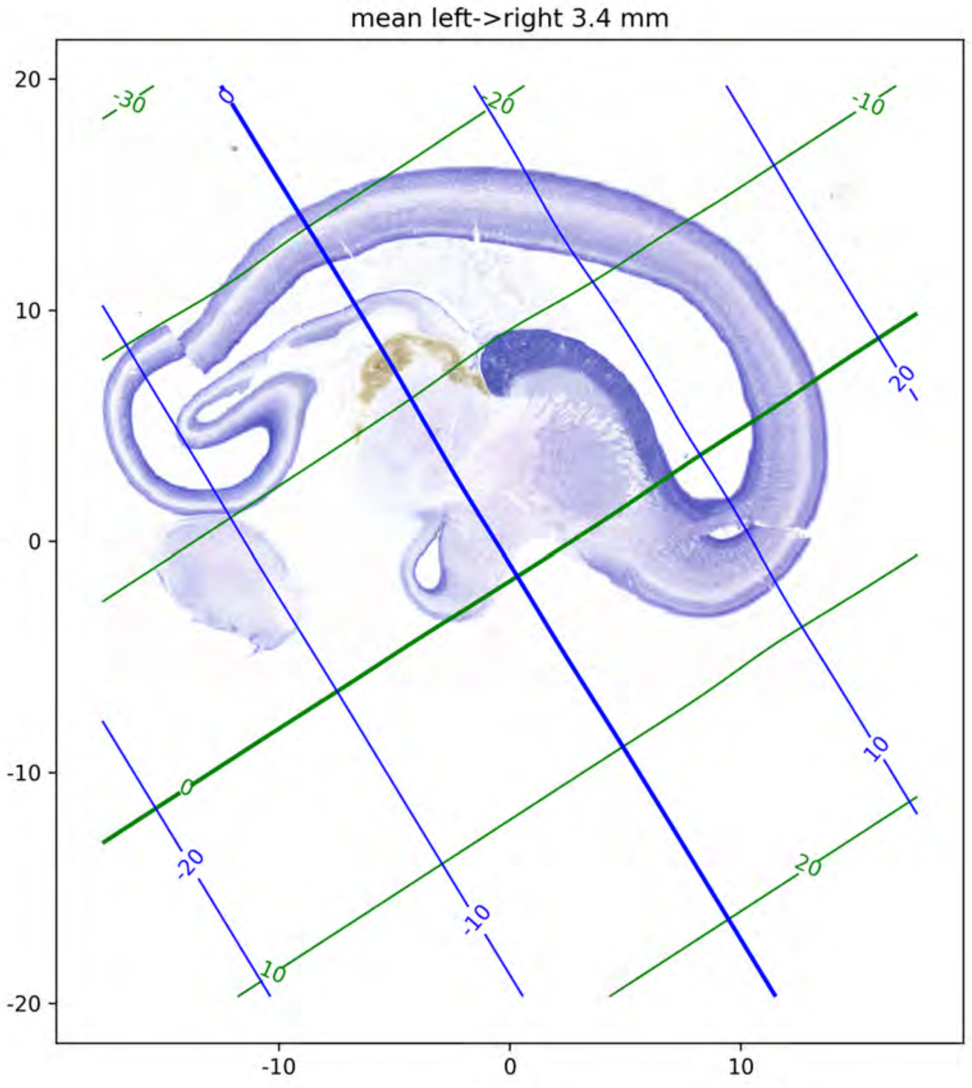
mean left->right 3.4 mm
20
10
0
-10
-20
-10
0
10
-30
-20
-10
0
10
20
0
20
10
-20
-10

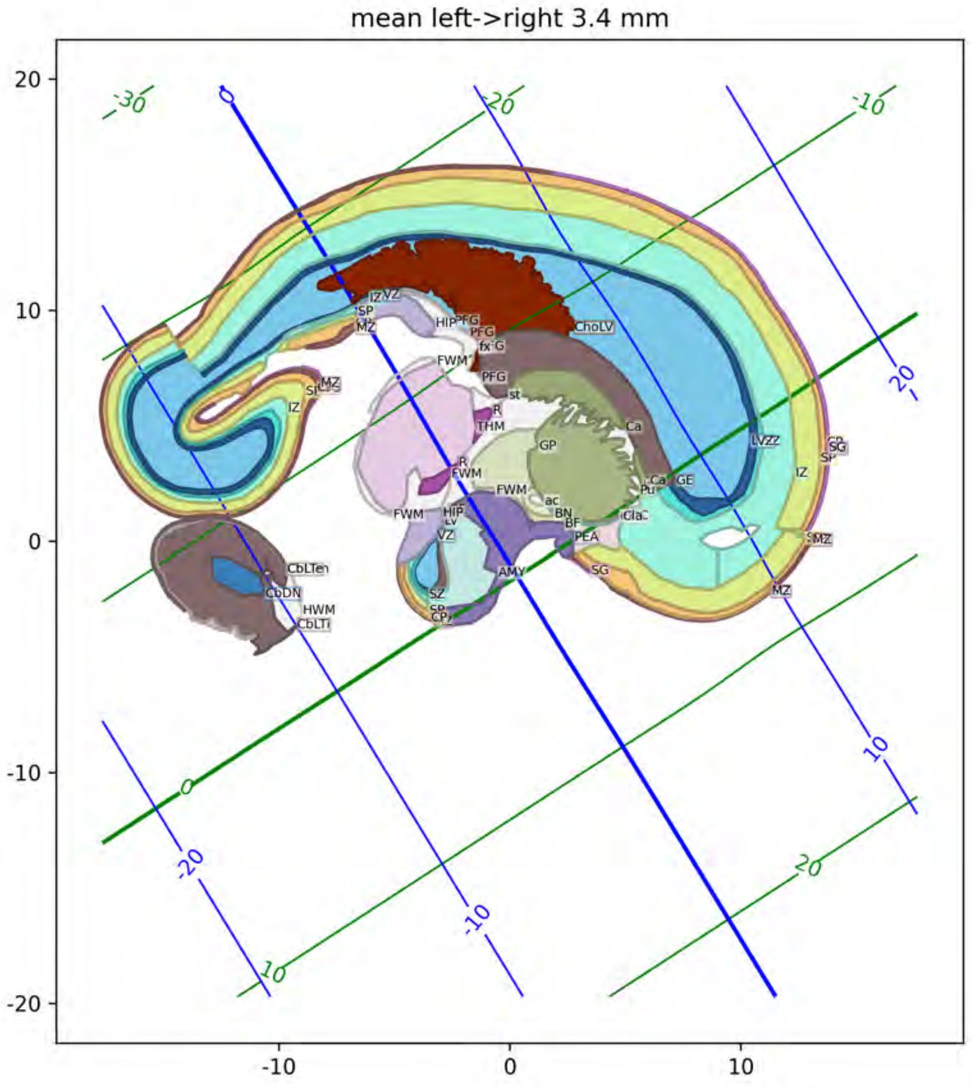
mean left->right 3.4 mm

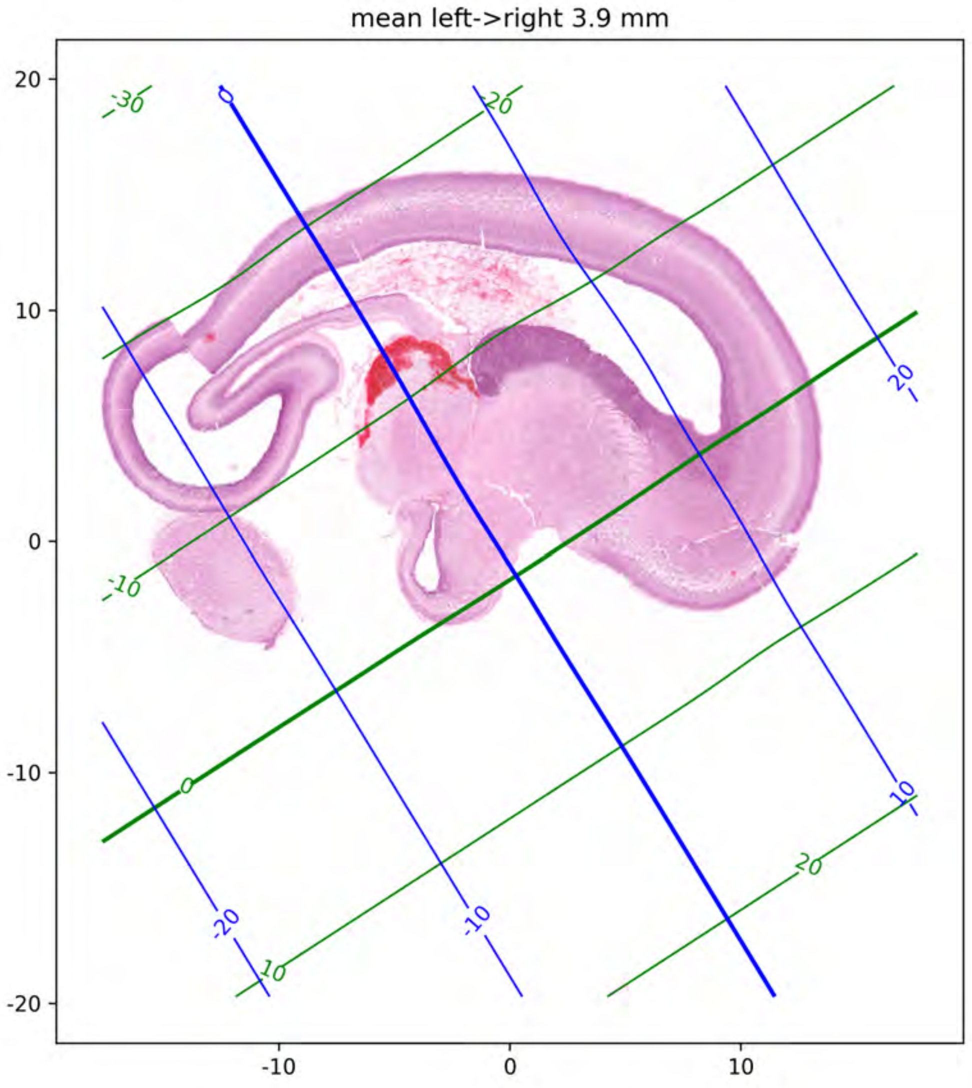

mean left->right 3.9 mm
20
10
0
-10
-20
-10
0
10
-30
-20
-10
0
10
20
0
20
20
10
-20
-10

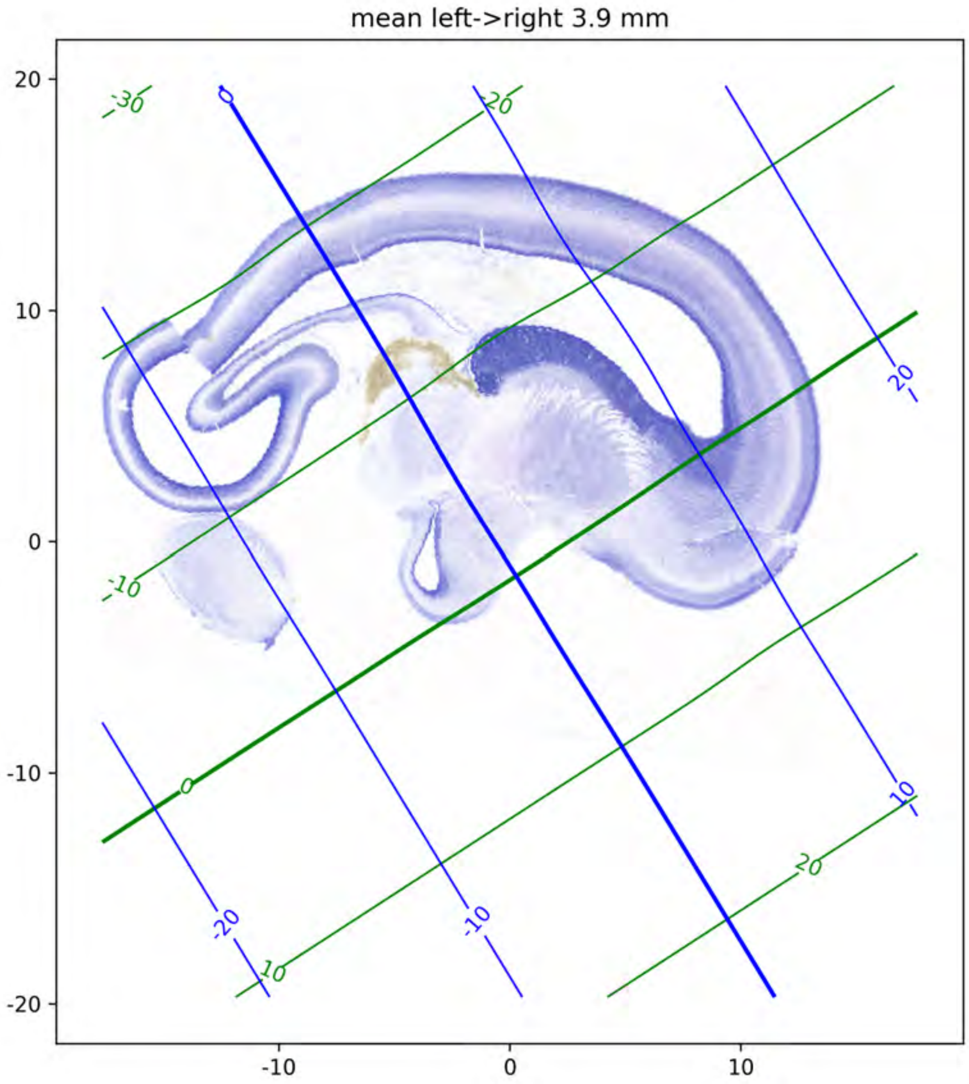
mean left->right 3.9 mm
20
10
0
-10
-20
-10
0
10
-30
-20
-10
0
10
20

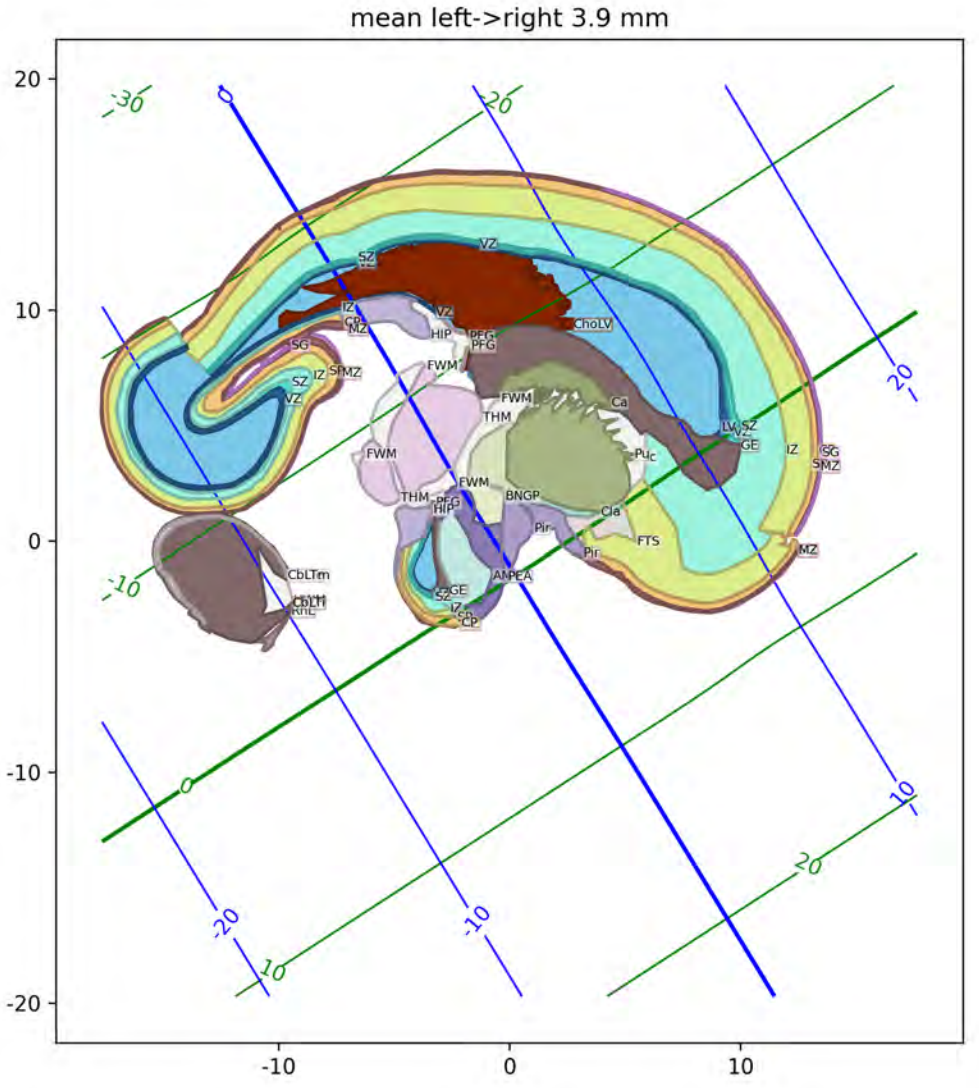
mean left->right 3.9 mm
20
10
0
-10
-20
-10
0
10
-30
-20
-10
0
10
20
VZ
SZ
IZ
CP
MZ
SG
HIP
PFG
FWM
THM
ChoLV
Ca
Puc
LV
GE
BNGP
Cla
Pir
FTS
CbLTm

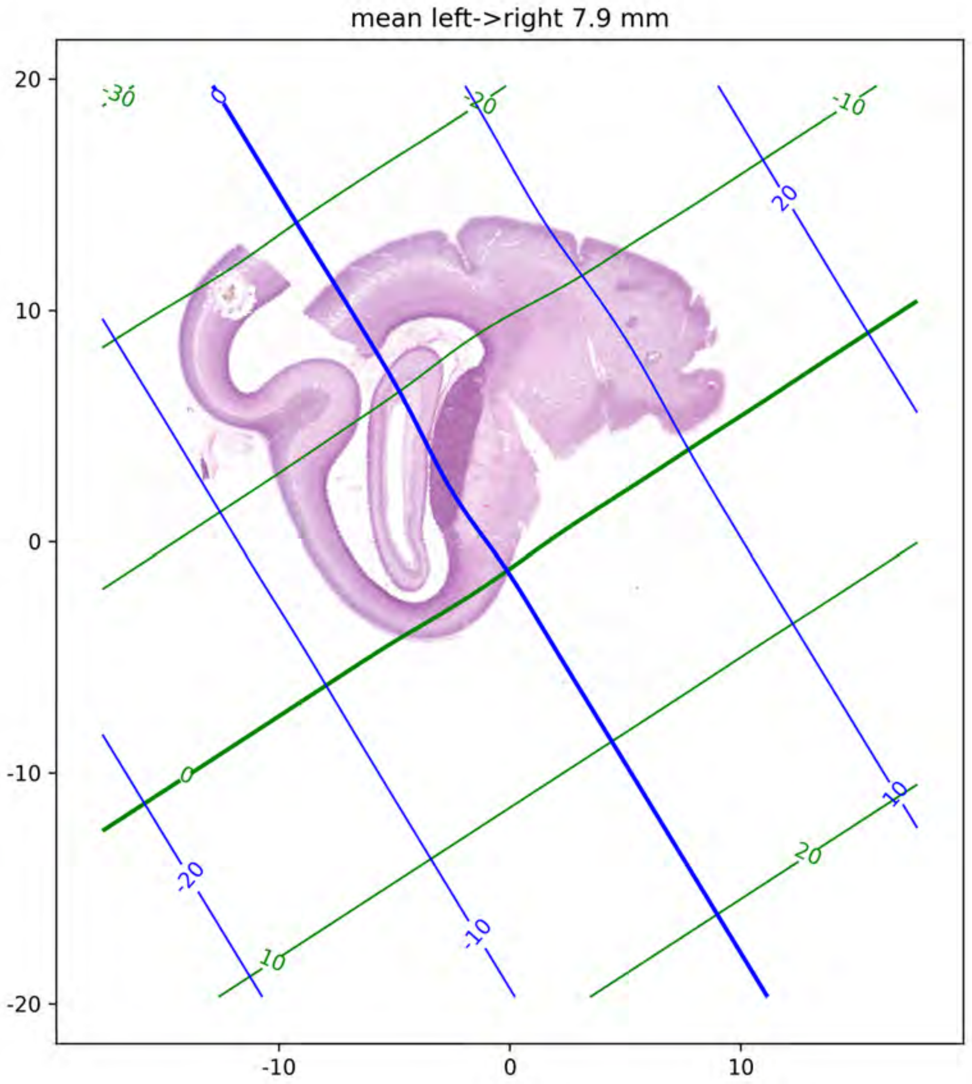

mean left->right 7.9 mm
20
10
0
-10
-20
-10
0
10
-30
0
-20
-10
20
0
-20
-10
10
20
10
-10
-20

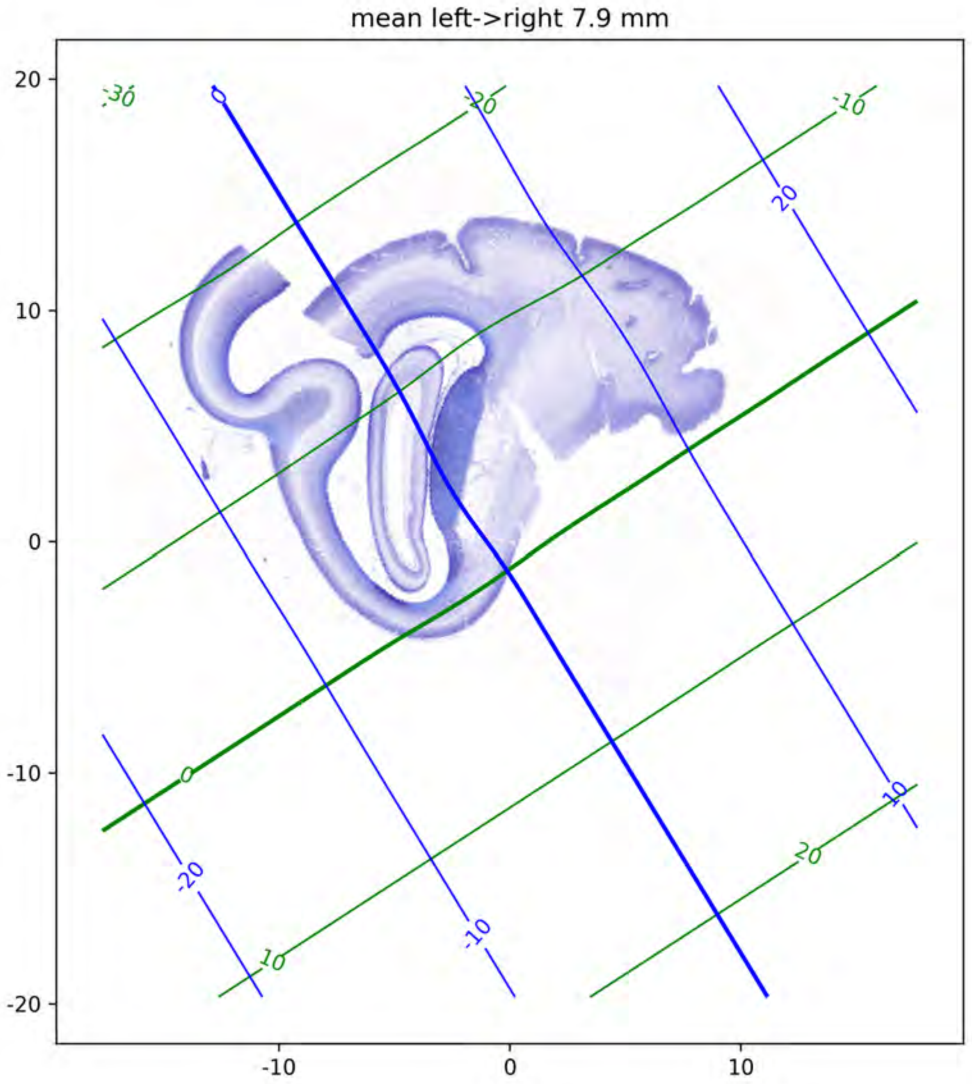
mean left->right 7.9 mm
20
10
0
-10
-20
-10
0
10
-30
0
-20
-10
20
10
0
-20
-10
10
20
10

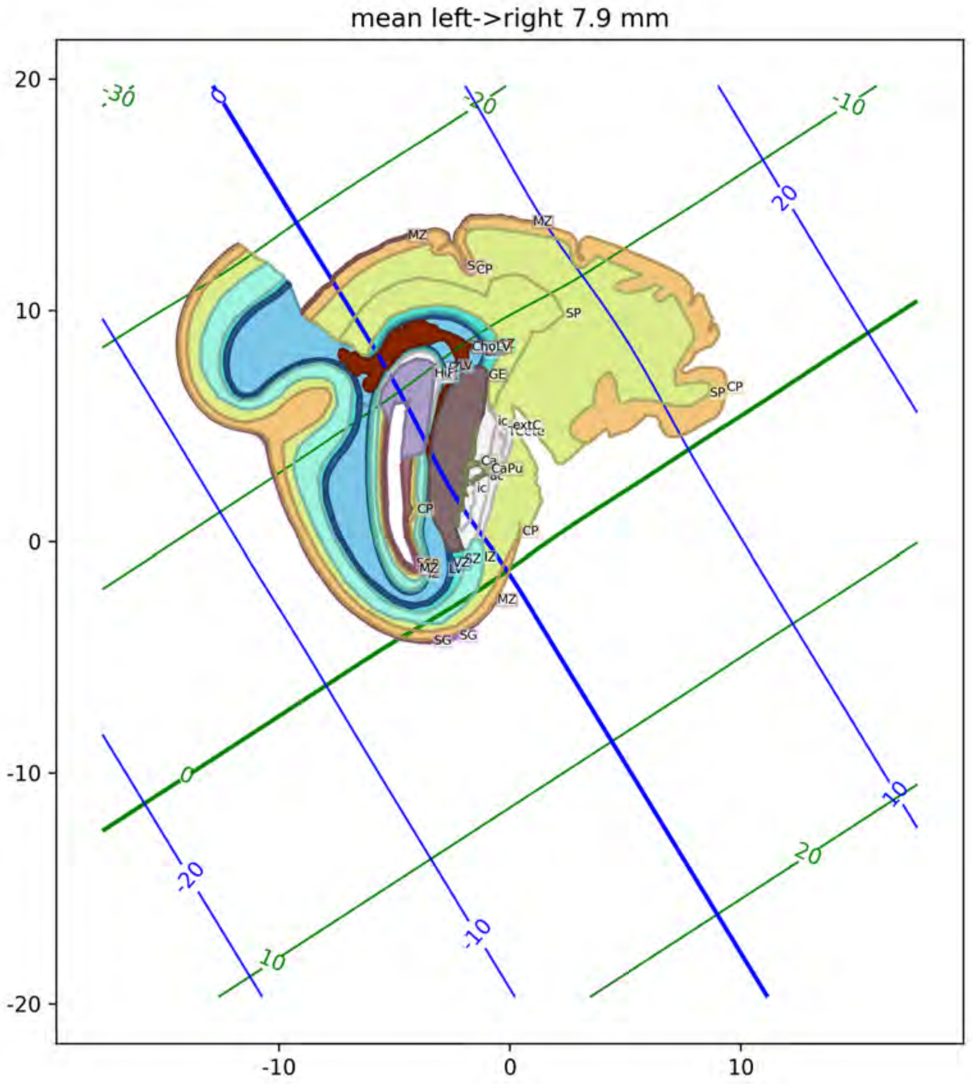
mean left->right 7.9 mm
20
10
0
-10
-20
-10
0
10
MZ
SCP
SP
CP
GE
ic
extC
Ca
CaPu
CP
MZ
SG

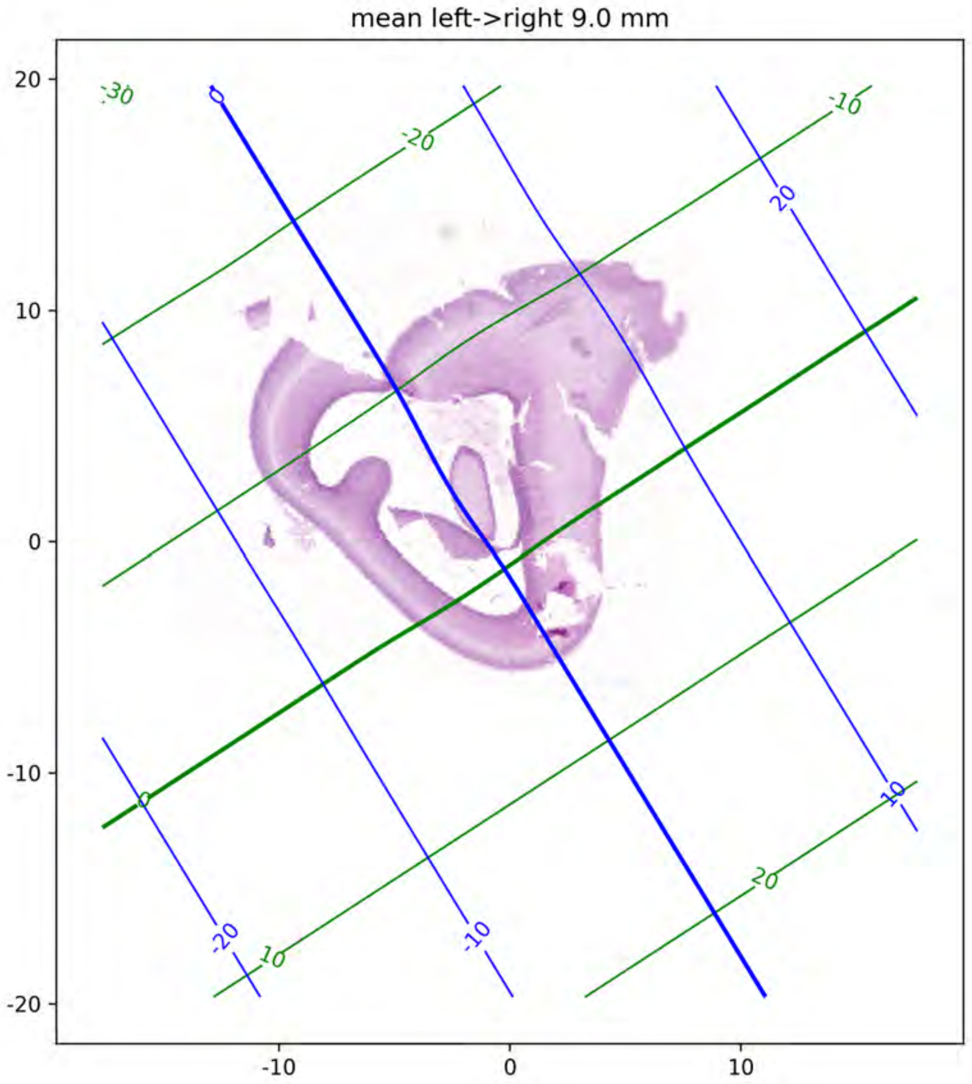
mean left->right 9.0 mm
20
10
0
-10
-20
-10
0
10
-30
-20
-10
0
10
20
0
20
10
-10
-20

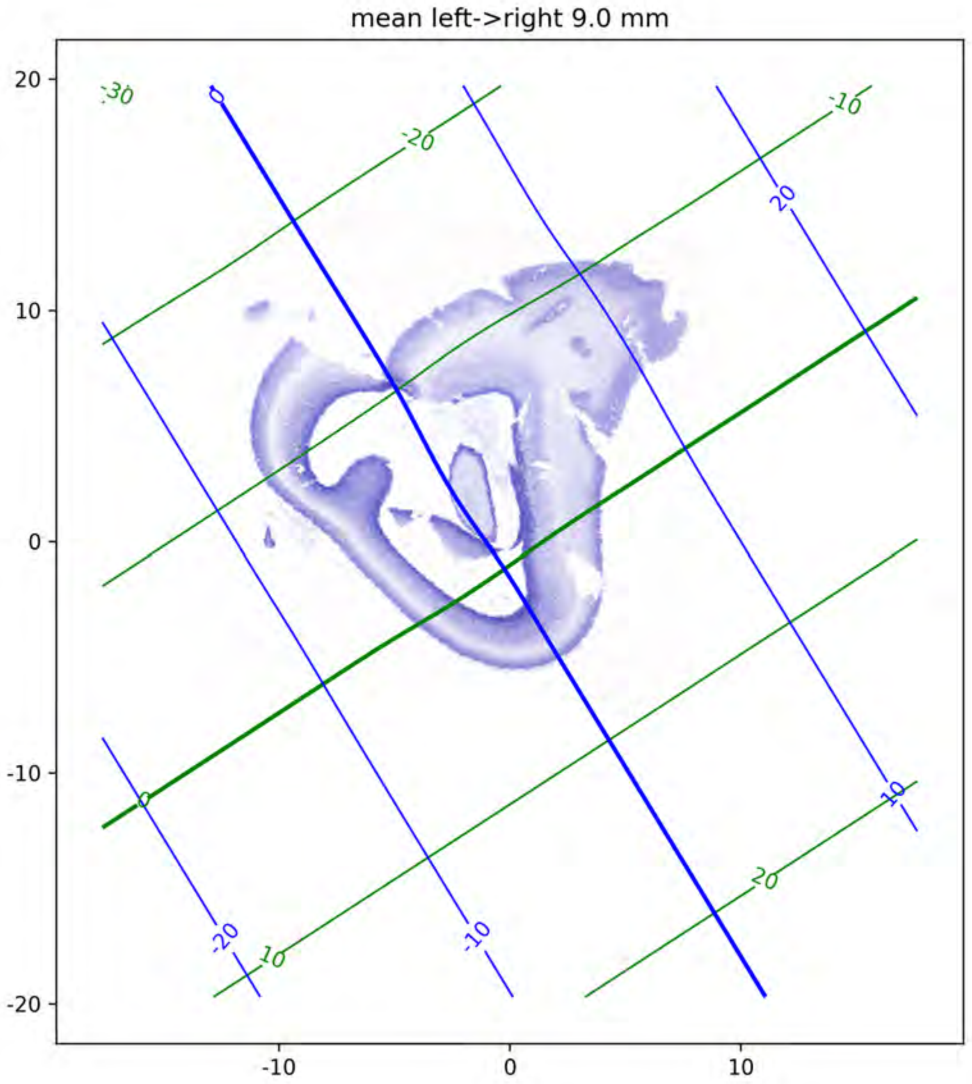
mean left->right 9.0 mm
20
10
0
-10
-20
-10
0
10
-30
-20
-10
0
10
20
0
20
10
-20
-10

mean left->right 9.0 mm

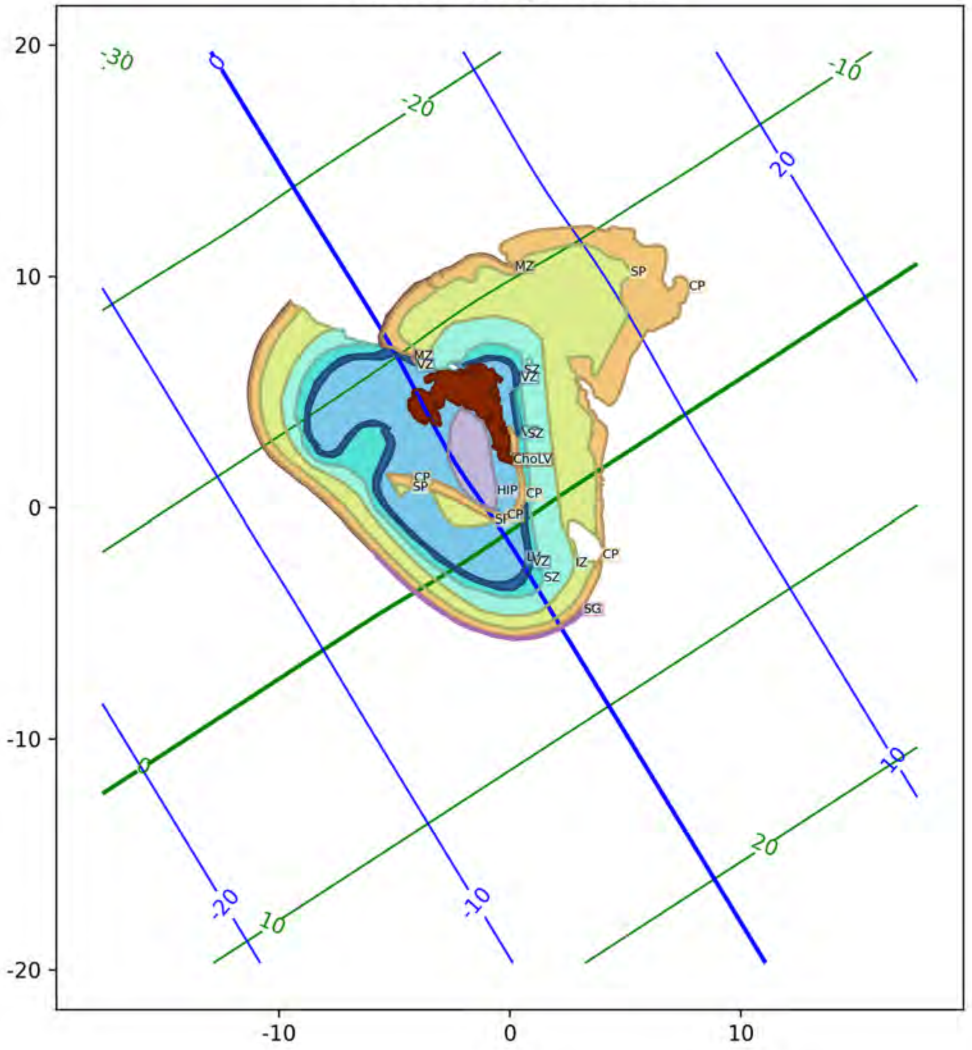
