## Supplementary material for "A three-dimensional histological cell atlas of the developing human brain": GW21 atlas plates

### Histological reference atlas and coordinate system for a 21 gestational week fetal brain

December 9, 2024

Corresponding manuscript: Jai Jayakumar, et al. “A three-dimensional histological cell atlas of the developing human brain”

Demographic information: Specimen ID: S1/ FB10 AP (cms): 6.6 Bi temporal diameter (cms): 5.1 Brain weight (gms): 56.24 Age (GW): 21 Sex: M

This atlas includes serial section histology data, with neighoring sections aligned. Sections are 20um thick and collected in a series of three, alternating between Nissl, H&E, and various antibody stains (not shown), so that sections corresponding to a individual stain type are 60 um apart.

We have developed a standard RAS (x=Right, y=Anterior, z=Superior) coordinate system described as shown below. Please refer to figure 2 in the manuscript.

The data below shows this coordinate system as a curvilinear grid, with 10mm spaced isocontours, labeled in mm. Red lines correspond to the left-right axis with right positive. Green lines correspond to the posterior-anterior axis with anterior positive. Blue lines correspond to the inferior-superior axis with superior positive. The x=0, y=0, and z=0 lines are drawn thicker. Note that our sagittal histology sections are nearly perpendicular to the left right axis, and so red lines appear more curved.

Below we show alternating pages with manually drawn annotations with median spacing between annotations of 0.66mm. Corresponding Nissl stained sections on which annotations were drawn are also included, here downsampled to 64 micron in plane resolution. Finally where neighboring tissue sections are available with H&E stain, these sections are also shown also at 64 micron resolution.

The horizontal and vertical axes of the image in each section also labeled with tick marks in mm, with 10 mm spacing to help with understanding scale.

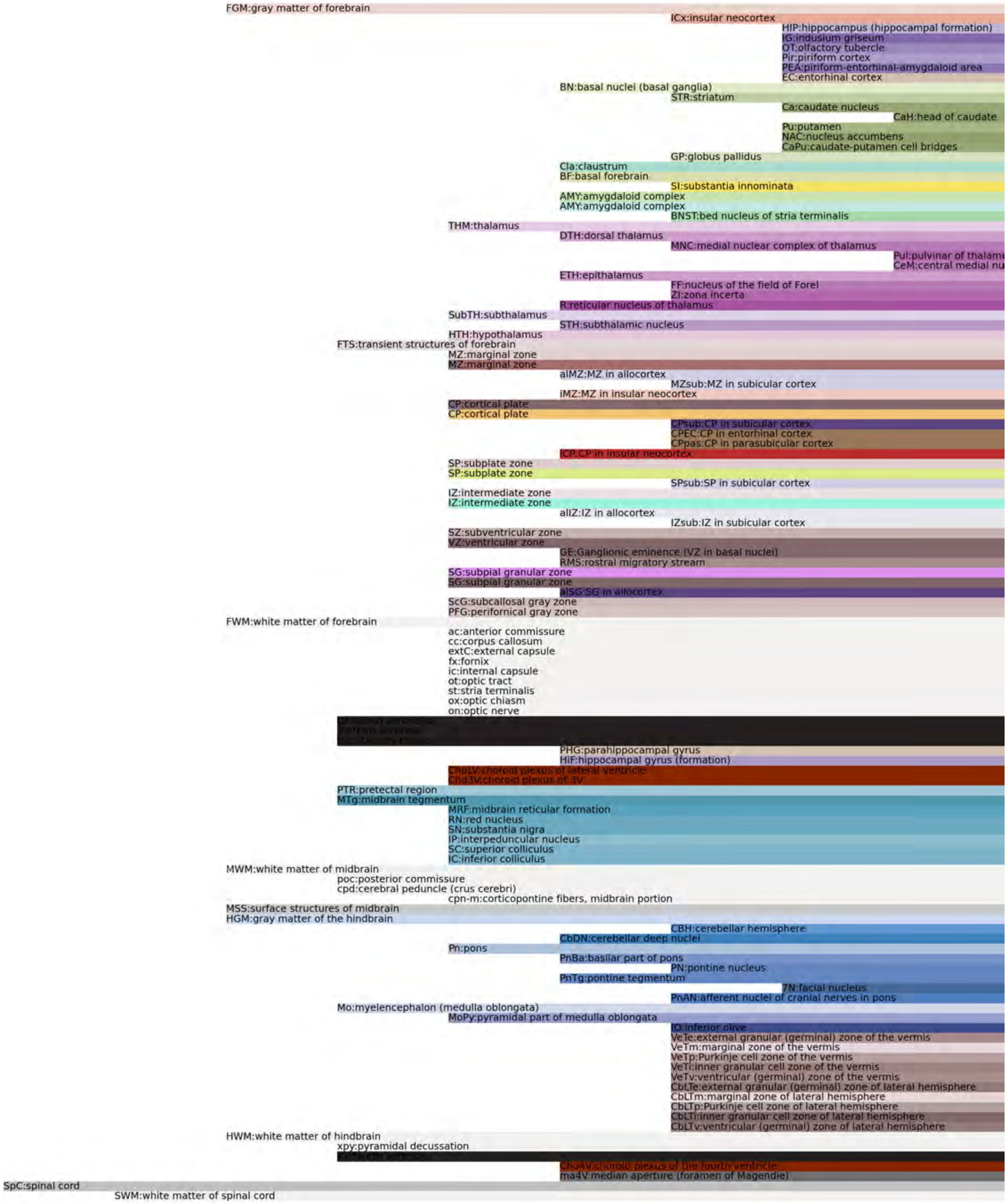
FGM:gray matter of forebrain
ICx:insular neocortex
HIP:hippocampus (hippocampal formation)
IG:indusium griseum
OT:olfactory tubercle
Pir:piriform cortex
PEA:piriform-entorhinal-amygdaloid area
EC:entorhinal cortex
BN:basal nuclei (basal ganglia)
STR:striatum
Ca:caudate nucleus
CaH:head of caudate
Pu:putamen
NAC:nucleus accumbens
CaPu:caudate-putamen cell bridges
GP:globus pallidus
Cla:claustrum
BF:basal forebrain
SI:substantia innominata
AMY:amygdaloid complex
AMY:amygdaloid complex
BNST:bed nucleus of stria terminalis
THM:thalamus
DTH:dorsal thalamus
MNC:medial nuclear complex of thalamus
Pul:pulvinar of thalam
CeM:central medial nu
ETH:epithalamus
FF:nucleus of the field of Forel
ZI:zona incerta
R:reticular nucleus of thalamus
SubTH:subthalamus
STH:subthalamic nucleus
HTH:hypothalamus
FTS:transient structures of forebrain
MZ:marginal zone
MZ:marginal zone
aIMZ:MZ in allocortex
MZsub:MZ in subicular cortex
iMZ:MZ in insular neocortex
CP:cortical plate
CP:cortical plate
CPsub:CP in subicular cortex
CPEC:CP in entorhinal cortex
CPpas:CP in parasubicular cortex
ICP:CP in insular neocortex
SP:subplate zone
SP:subplate zone
SPsub:SP in subicular cortex
IZ:intermediate zone
IZ:intermediate zone
aIIZ:IZ in allocortex
IZsub:IZ in subicular cortex
SZ:subventricular zone
VZ:ventricular zone
GE:Ganglionic eminence (VZ in basal nuclei)
RMS:rostral migratory stream
SG:subpial granular zone
SG:subpial granular zone
aISG:SG in allocortex
ScG:subcallosal gray zone
PFG:perifornical gray zone
FWM:white matter of forebrain
ac:anterior commissure
cc:corpus callosum
extC:external capsule
fx:fornix
ic:internal capsule
ot:optic tract
st:stria terminalis
ox:optic chiasm
on:optic nerve
PHG:parahippocampal gyrus
HiF:hippocampal gyrus (formation)
ChoLV:choroid plexus of lateral ventricle
Cho3V:choroid plexus of 3V
PTR:pretectal region
MTg:midbrain tegmentum
MRF:midbrain reticular formation
RN:red nucleus
SN:substantia nigra
IP:interpeduncular nucleus
SC:superior colliculus
IC:inferior colliculus
MWM:white matter of midbrain
poc:posterior commissure
cpd:cerebral peduncle (crus cerebri)
cpn-m:corticopontine fibers, midbrain portion
MSS:surface structures of midbrain
HGM:gray matter of the hindbrain
CBH:cerebellar hemisphere
CbDN:cerebellar deep nuclei
Pn:pons
PnBa:basilar part of pons
PN:pontine nucleus
PnTg:pontine tegmentum
7N:facial nucleus
PnAN:afferent nuclei of cranial nerves in pons
Mo:myelencephalon (medulla oblongata)
MoPy:pyramidal part of medulla oblongata
IO:inferior olive
VeTe:external granular (germinal) zone of the vermis
VeTm:marginal zone of the vermis
VeTp:Purkinje cell zone of the vermis
VeTi:inner granular cell zone of the vermis
VeTv:ventricular (germinal) zone of the vermis
CbLTe:external granular (germinal) zone of lateral hemisphere
CbLTm:marginal zone of lateral hemisphere
CbLTp:Purkinje cell zone of lateral hemisphere
CbLTi:inner granular cell zone of lateral hemisphere
CbLTv:ventricular (germinal) zone of lateral hemisphere
HWM:white matter of hindbrain
xpy:pyramidal decussation
Cho4V:choroid plexus of the fourth ventricle
ma4V:median aperture (foramen of Magendie)
SpC:spinal cord
SWM:white matter of spinal cord

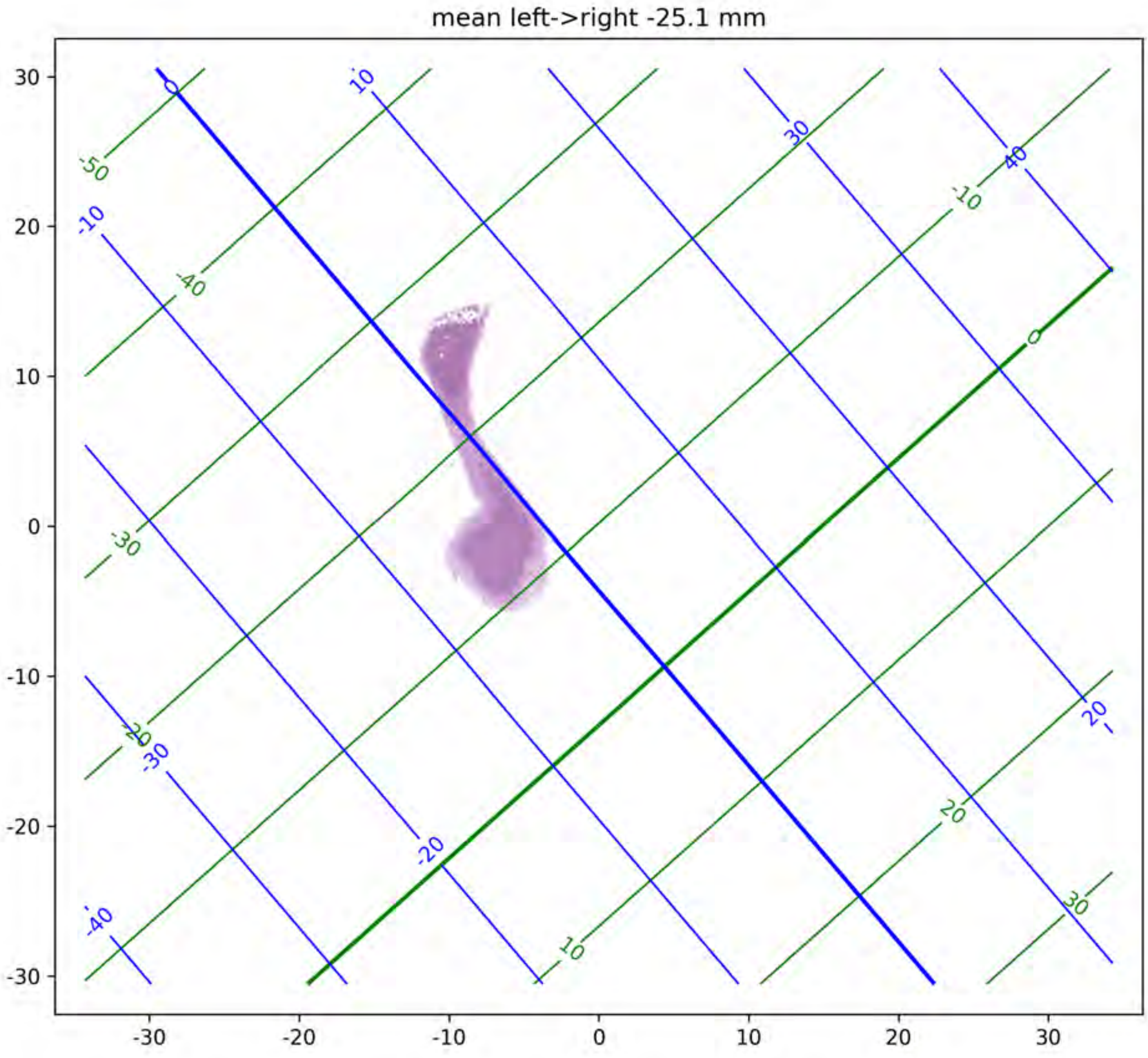

mean left->right -25.1 mm

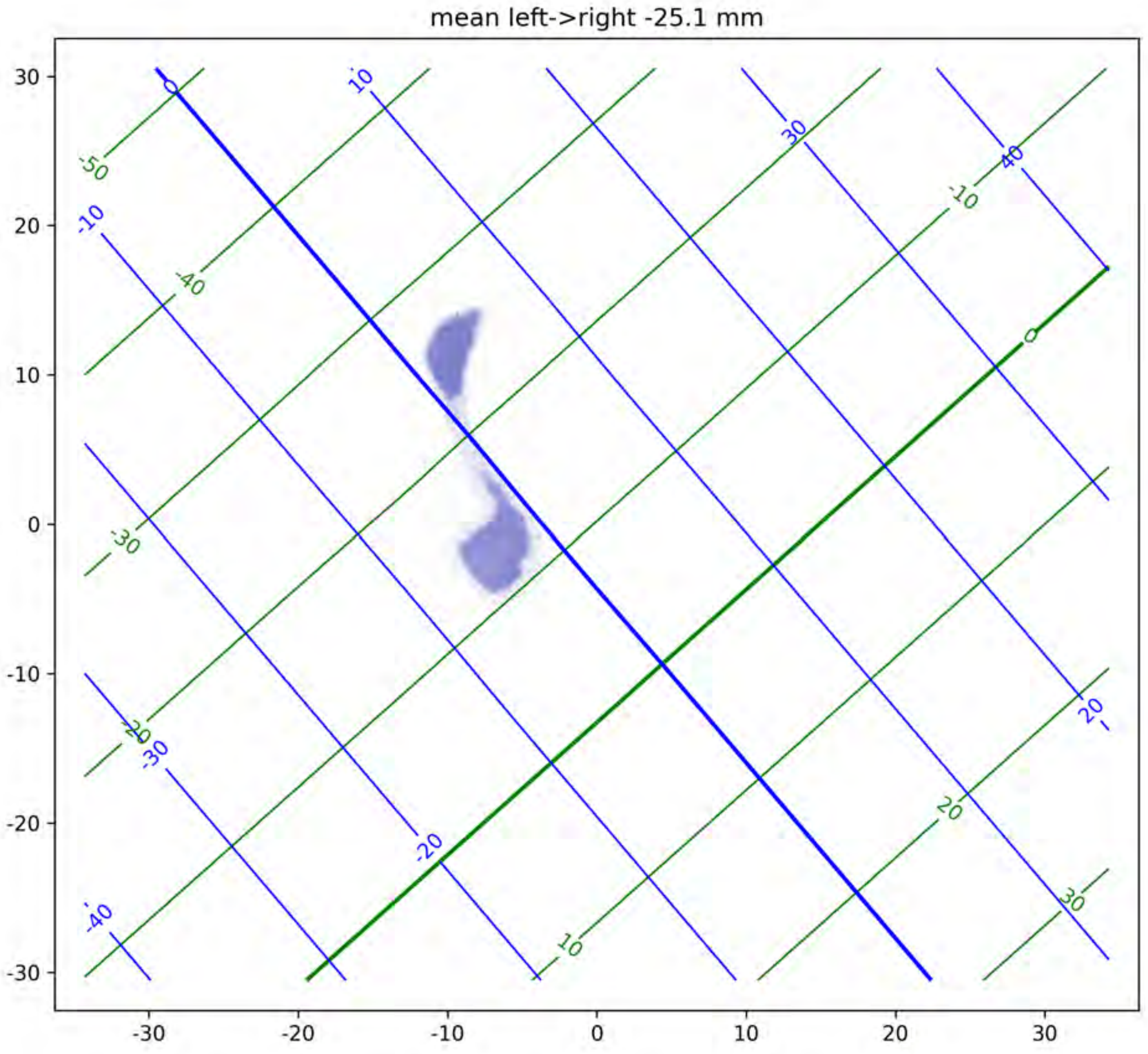

mean left->right -25.1 mm
30
20
10
0
-10
-20
-30
-30
-20
-10
0
10
20
30

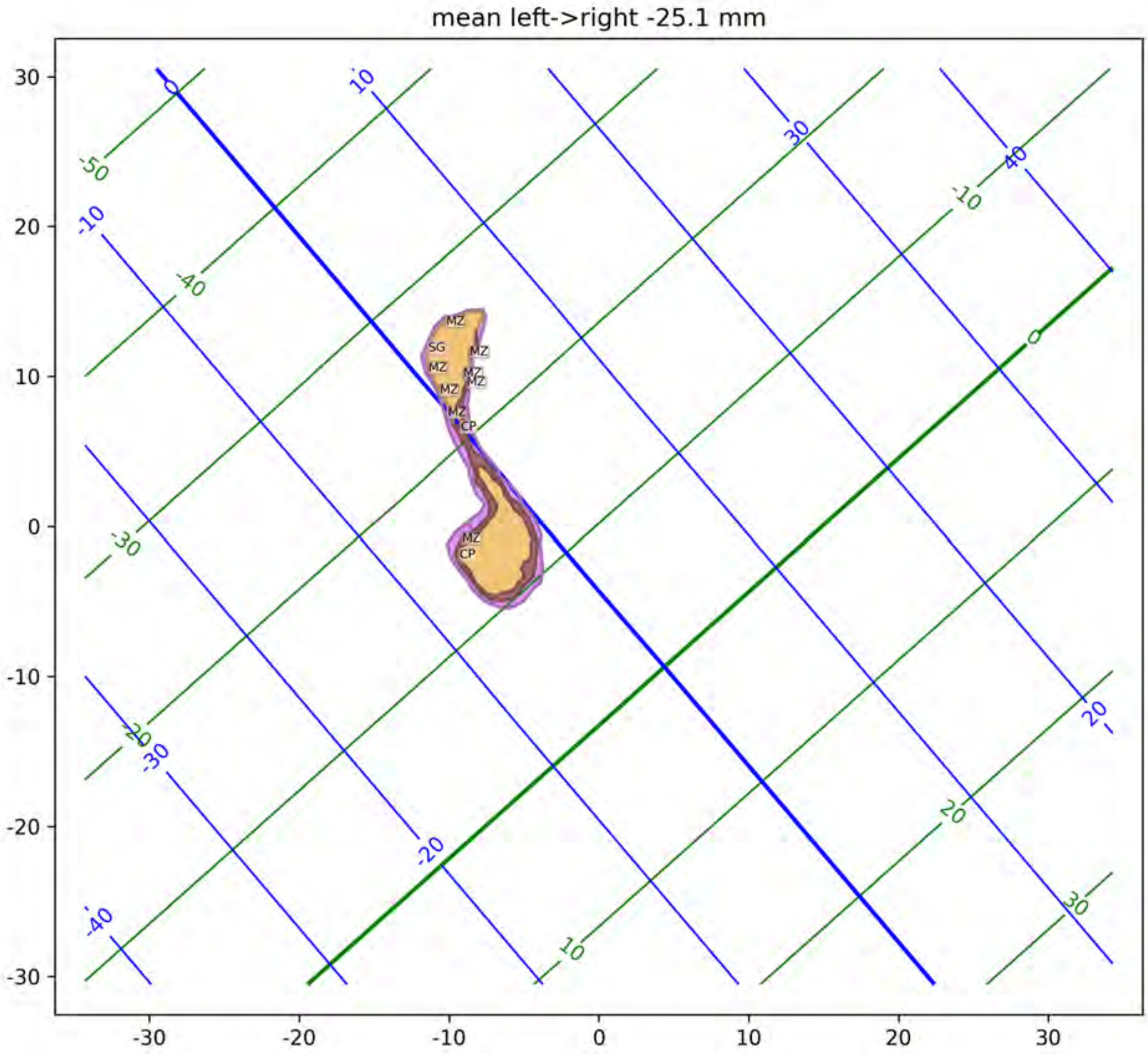

mean left->right -25.1 mm
MZ
SG
MZ
MZ
MZ
MZ
MZ
MZ
CP
MZ
CP

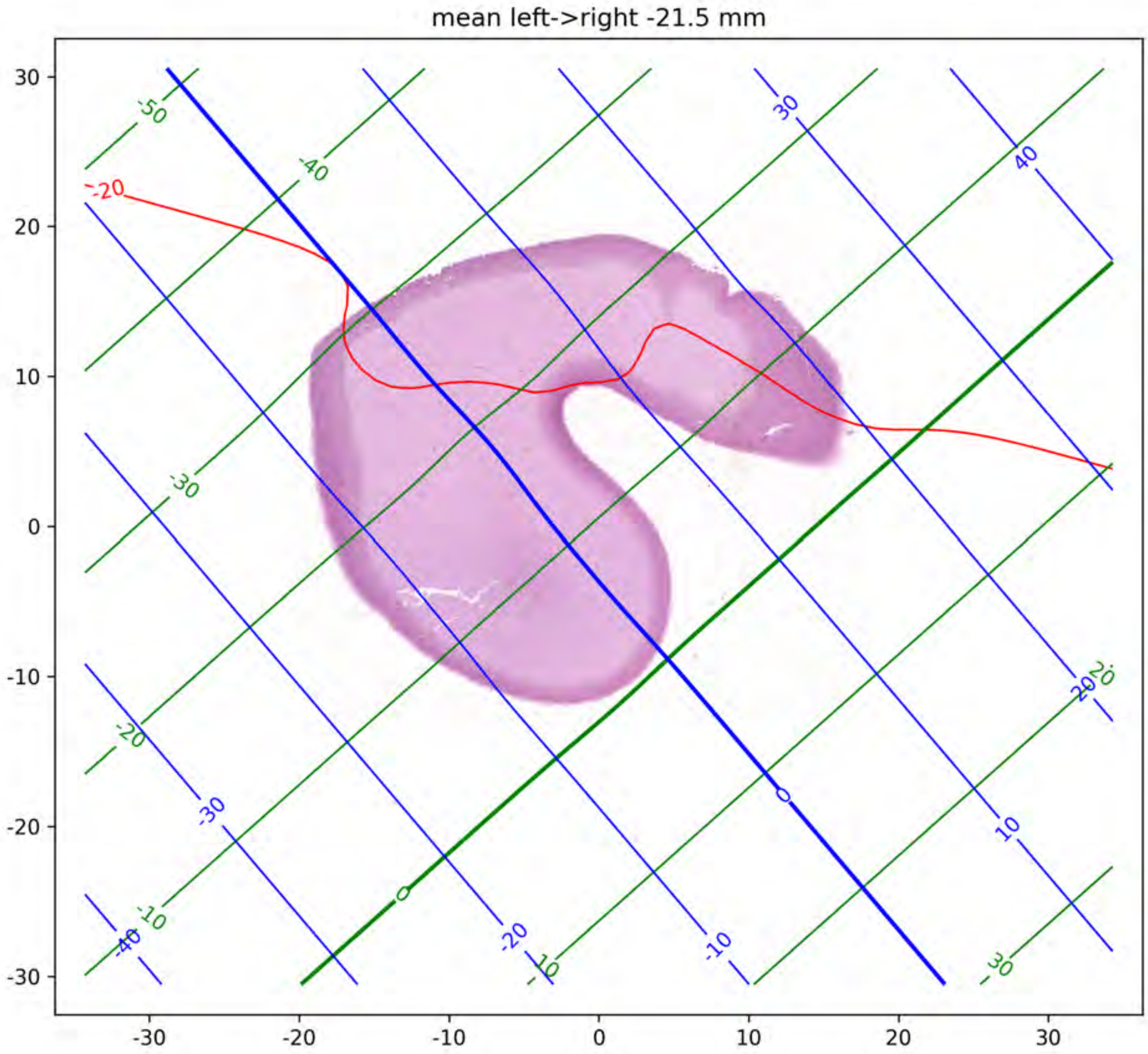

mean left->right -21.5 mm
30
20
10
0
-10
-20
-30
-30
-20
-10
0
10
20
30
-50
-40
-30
-20
-10
0
10
20
30
-20
30
40
-30
-40
-20
-10
0
10
20

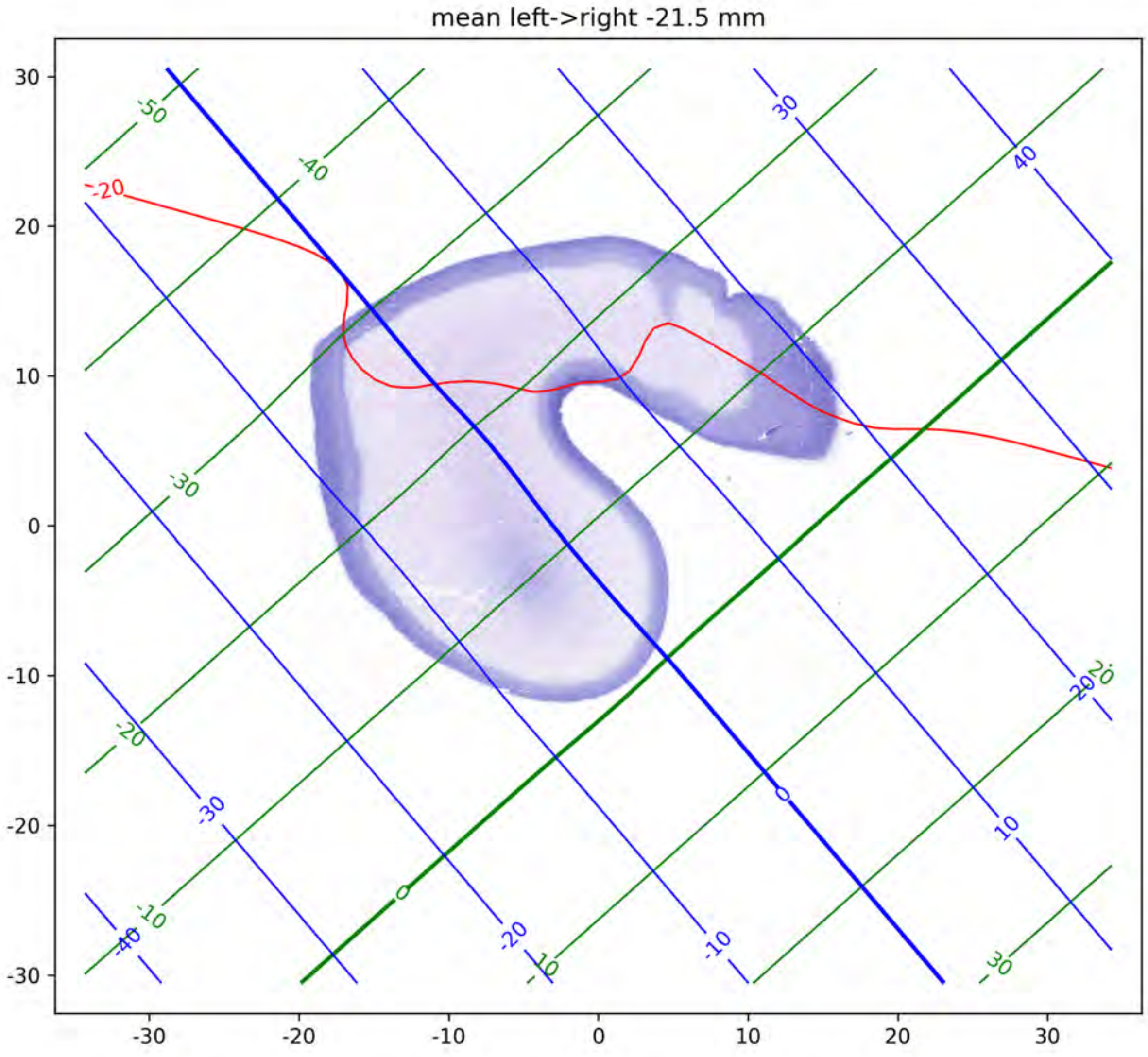

mean left->right -21.5 mm
-50
-40
-30
-20
-10
0
10
20
30
-40
-30
-20
-10
0
10
20
30
40
-20

mean left->right -21.5 mm

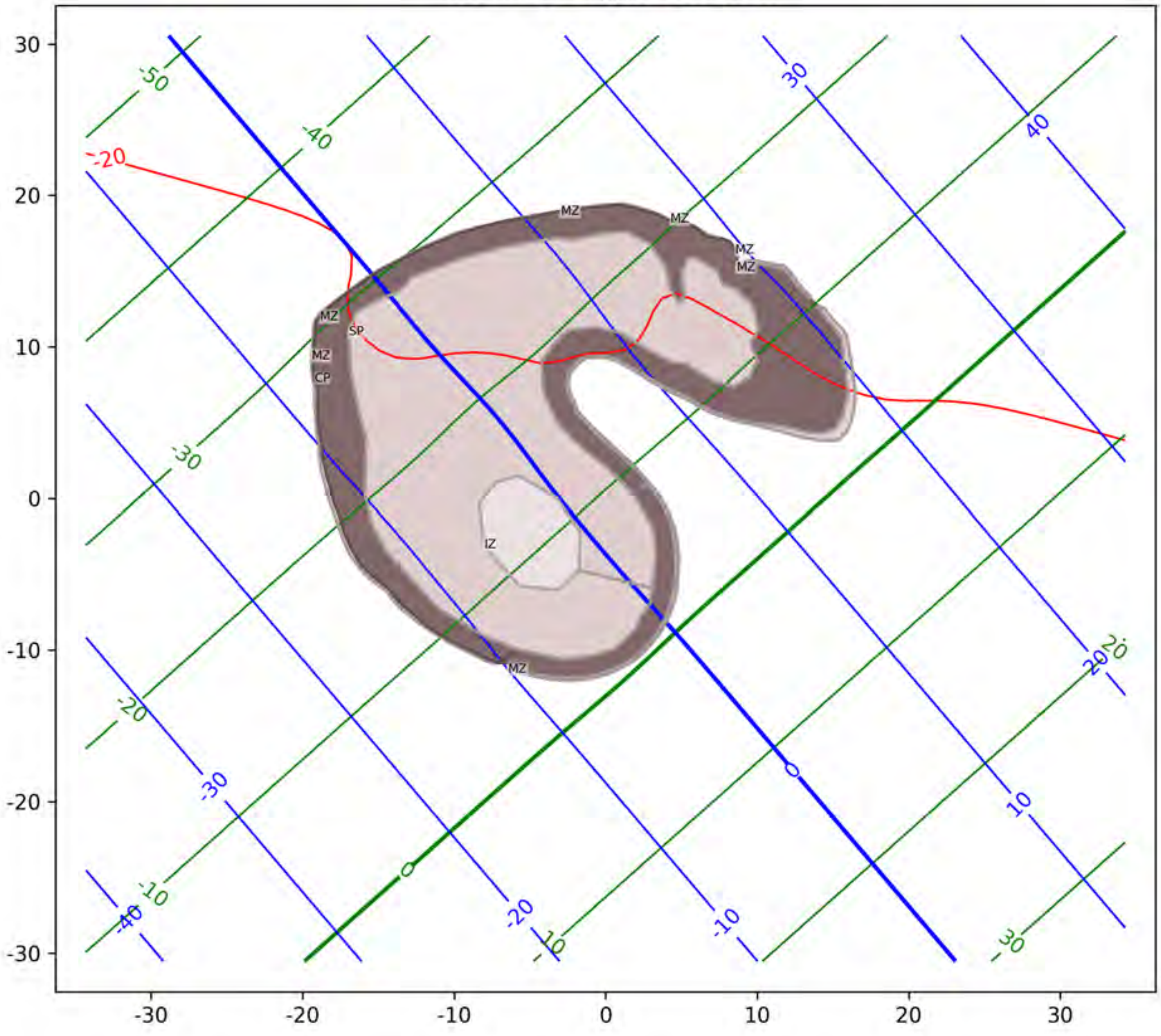

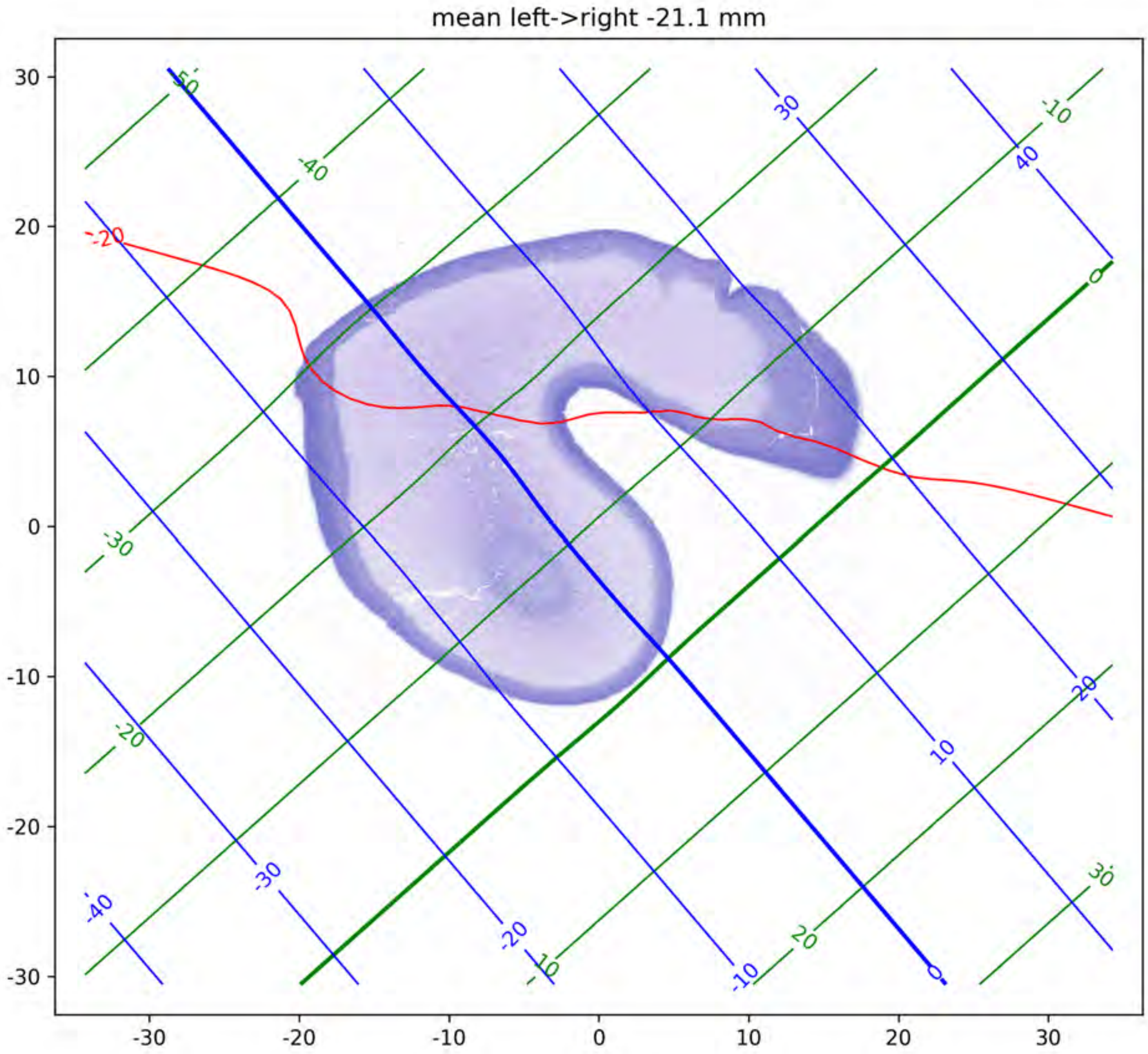

mean left->right -21.1 mm

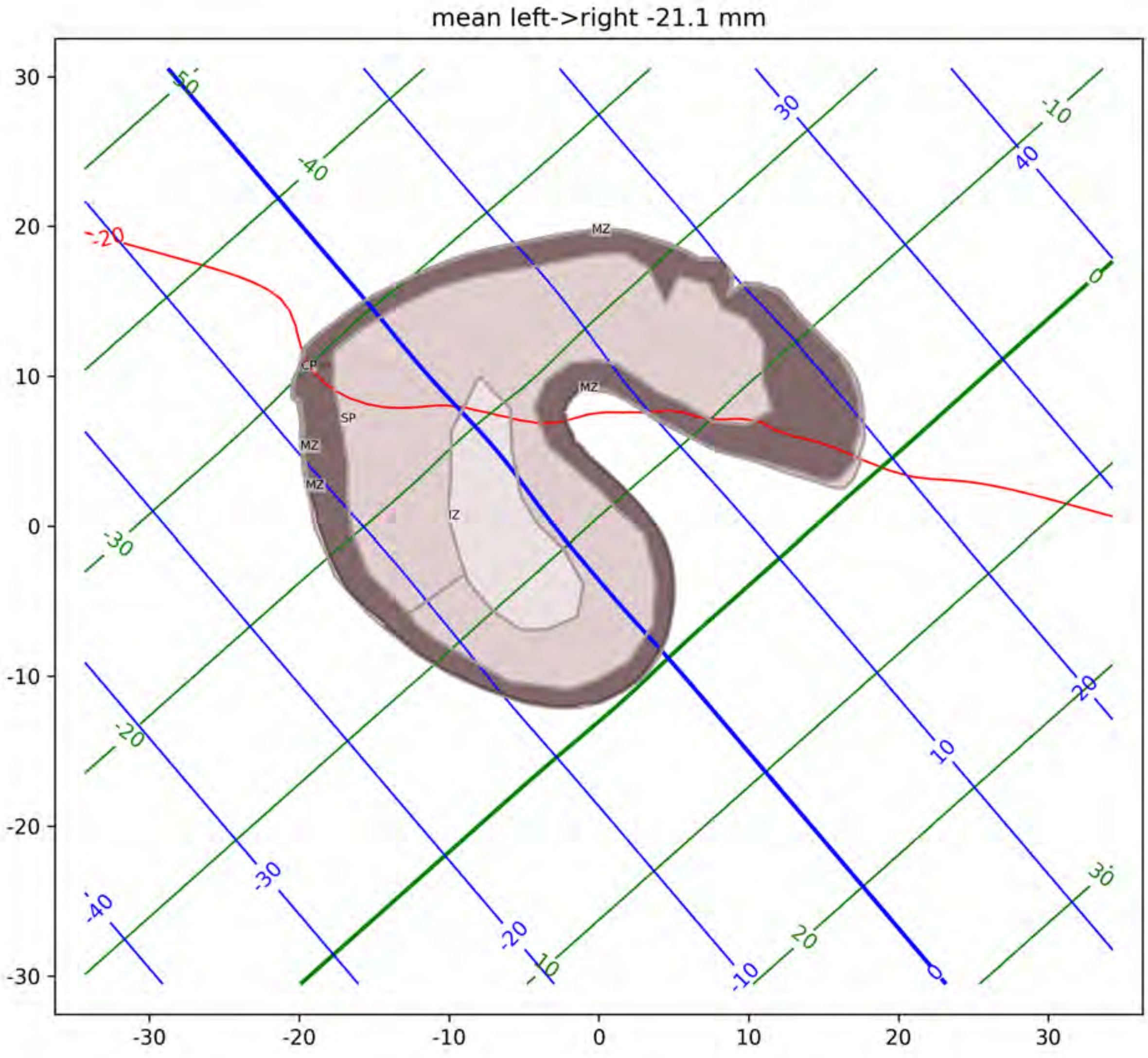
mean left->right -21.1 mm
MZ
CP
SP
IZ

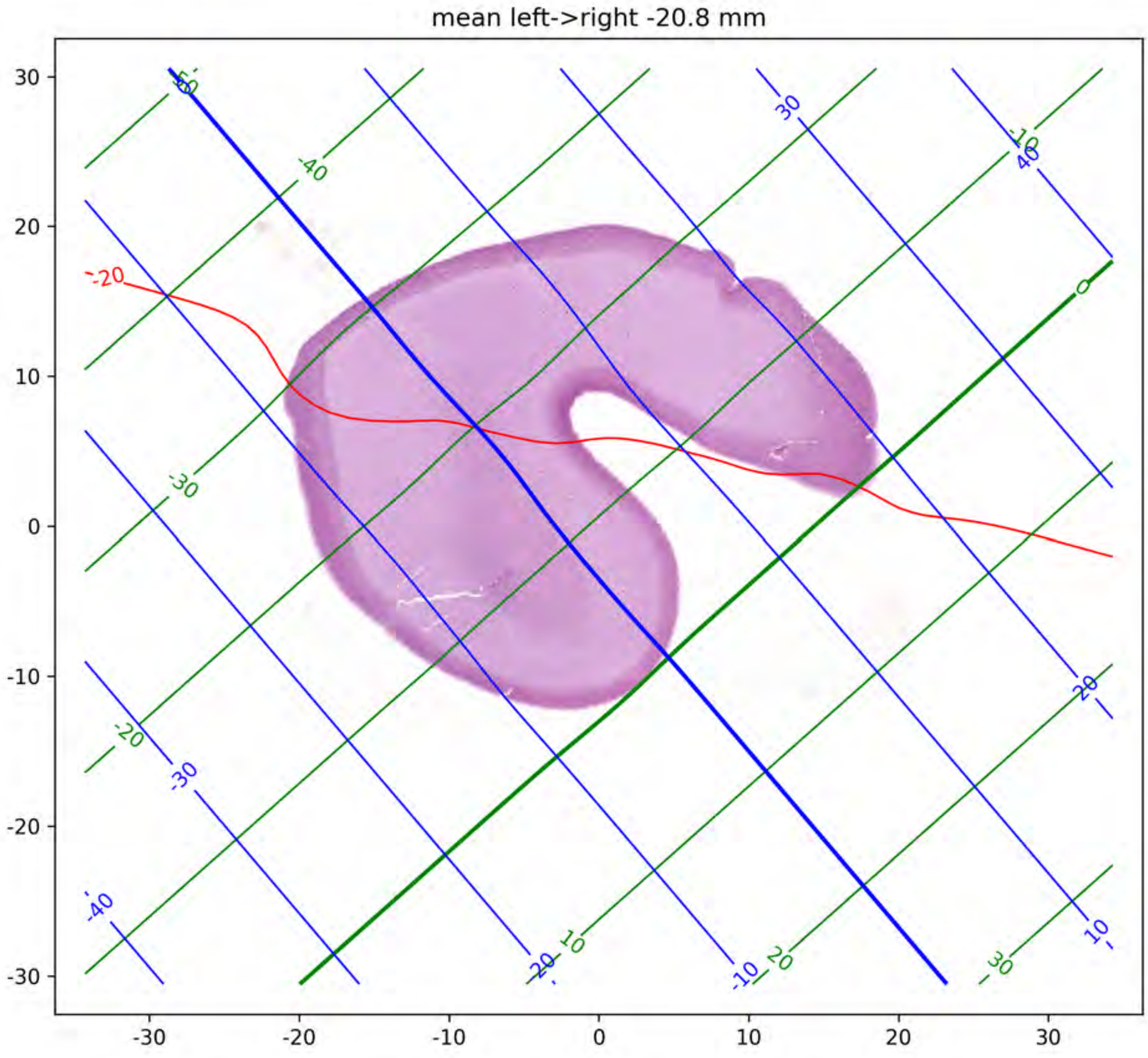

mean left->right -20.8 mm

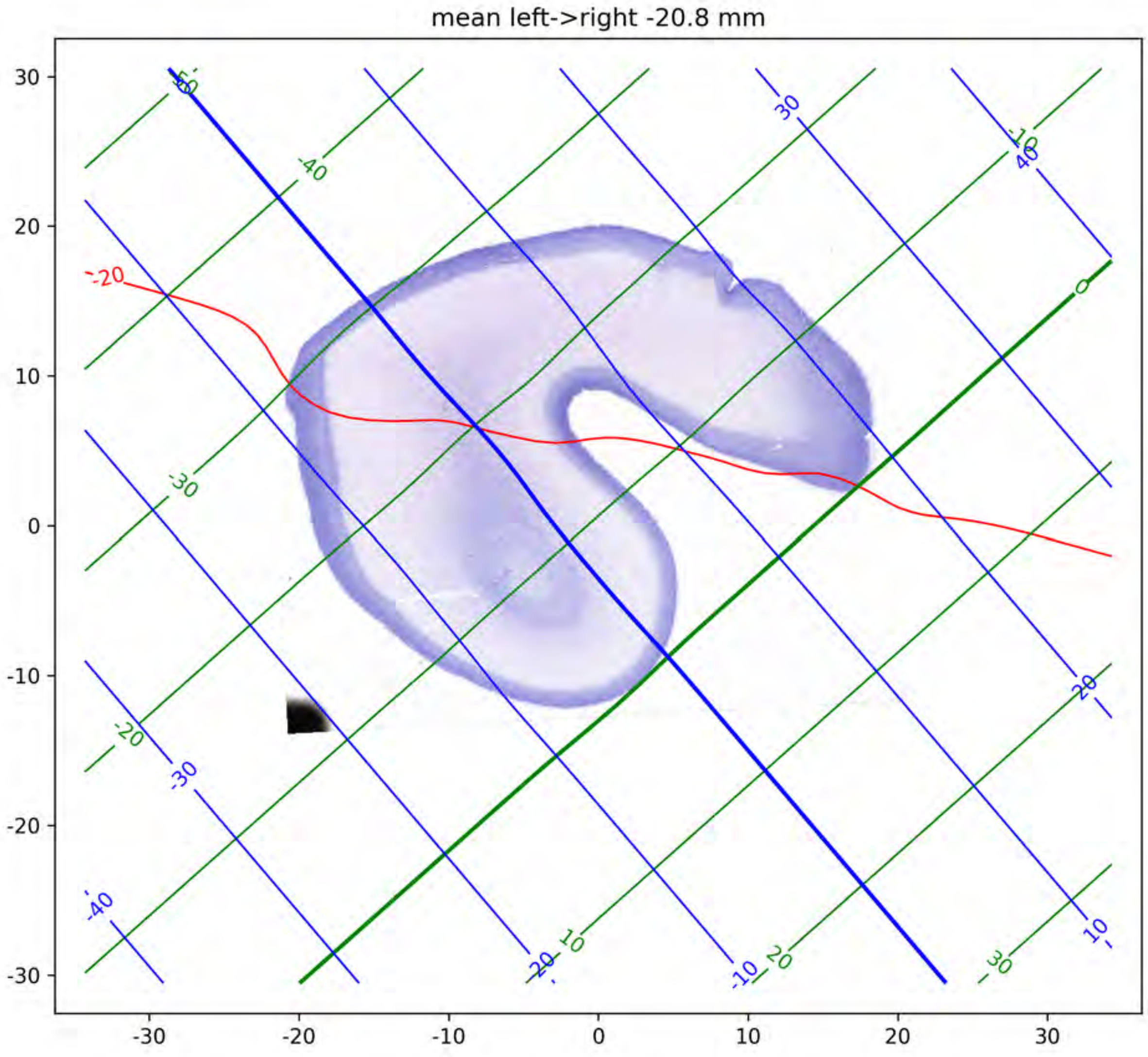
mean left->right -20.8 mm
-20
-50
-40
-30
-20
-10
0
10
20
30
40
-40
-30
-20
-10
10
20
30

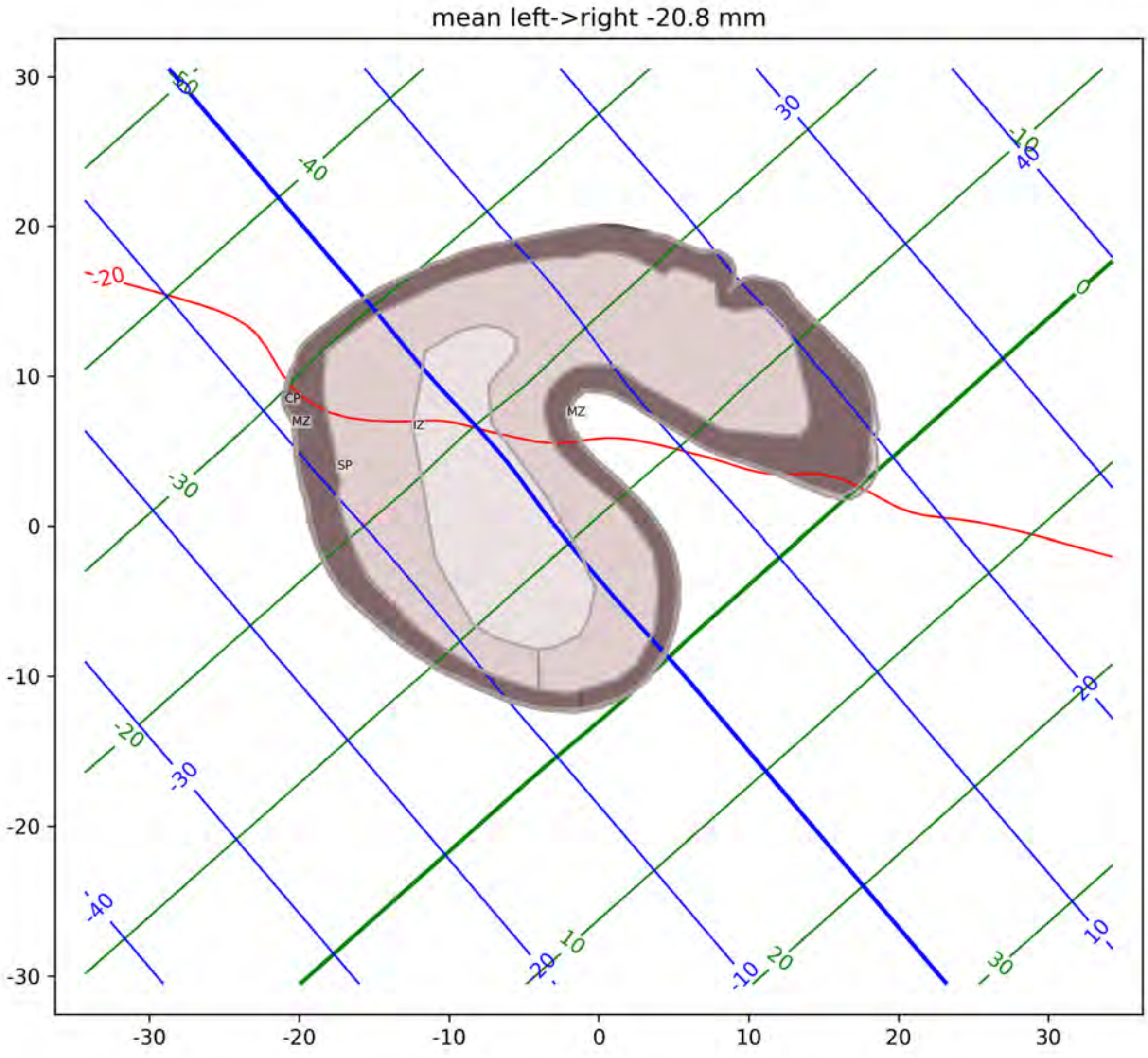
mean left->right -20.8 mm
CP
MZ
IZ
MZ
SP

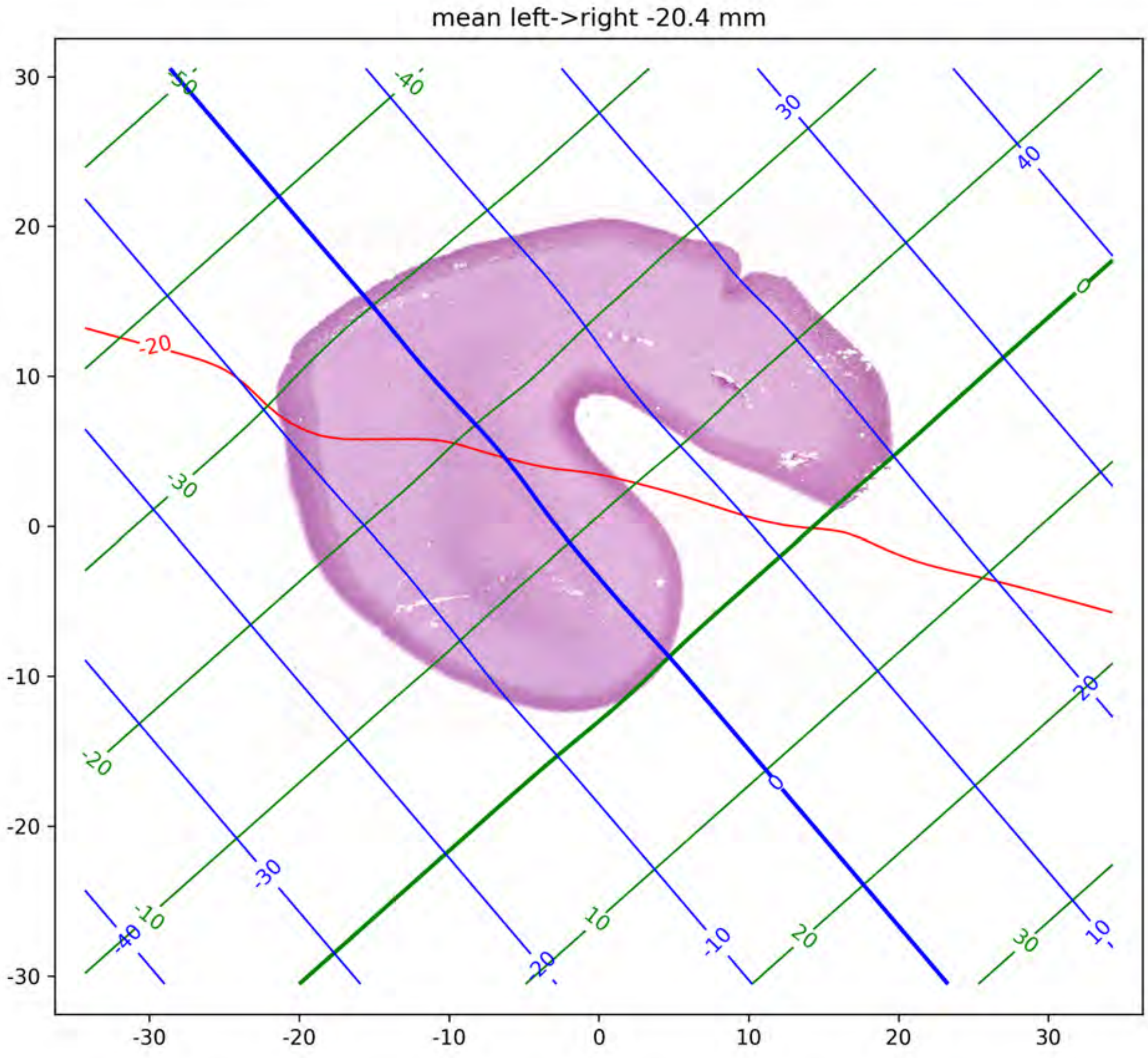
mean left->right -20.4 mm
30
20
10
0
-10
-20
-30
-30
-20
-10
0
10
20
30
-50
-40
30
40
-20
-30
0
-20
20
-30
-40
-10
10
-20
-10
20
30
10
0

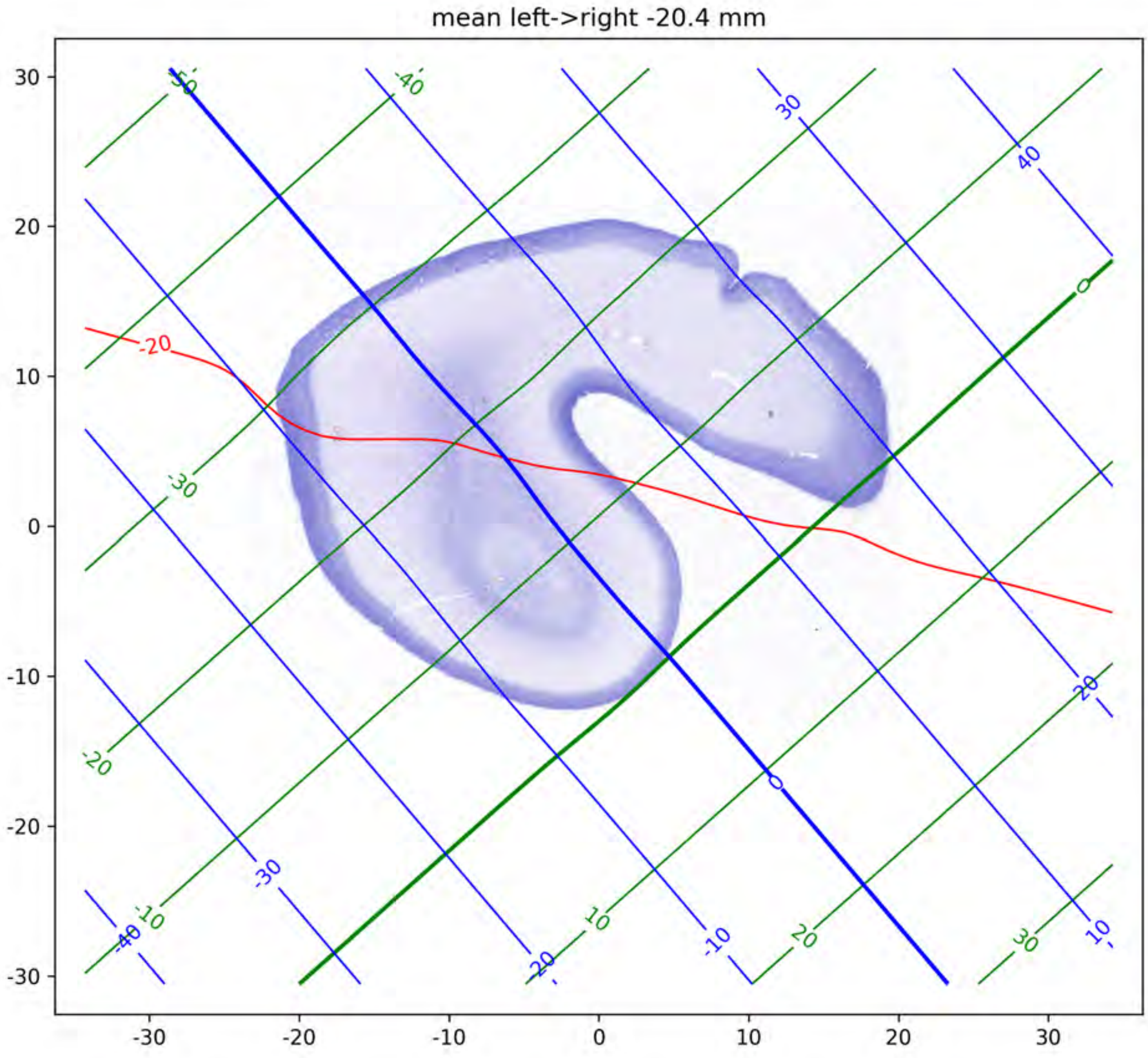
mean left->right -20.4 mm
-20
-50
-40
30
40
0
-30
-20
20
-10
10
-40
-30
-20
-10
0
10
20
30

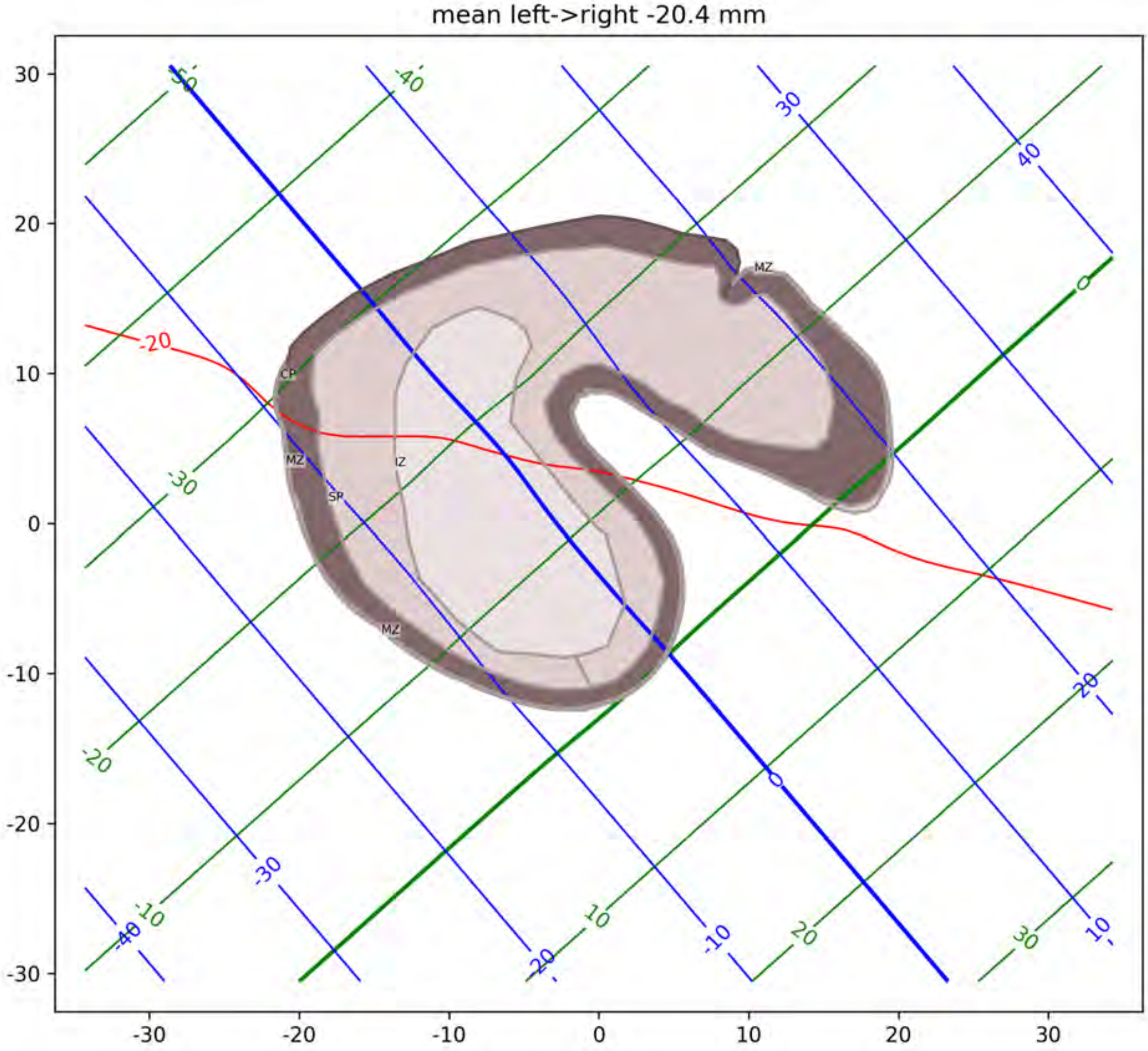
mean left->right -20.4 mm
MZ
CP
MZ
IZ
SP
MZ
-20
-50
-40
-30
-20
-10
10
20
30
0
30
40
20
10
0
-10
-20
-30
-40
30
20
10
0
-10
-20
-30
-30
-20
-10
0
10
20
30

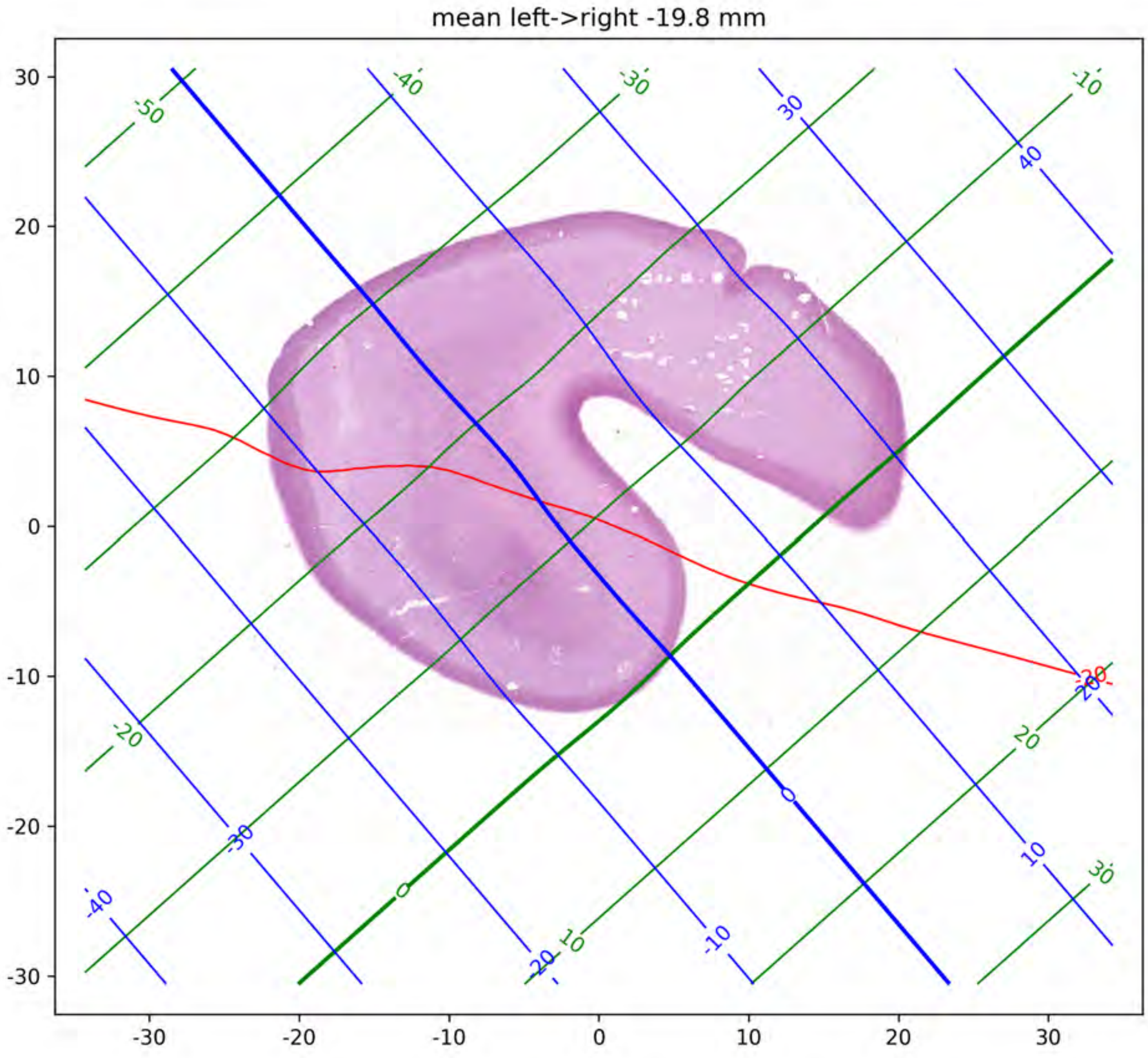
mean left->right -19.8 mm
30
20
10
0
-10
-20
-30
-30
-20
-10
0
10
20
30
-50
-40
-30
-10
-20
0
10
20
30
30
40
-30
-40
-20
-10
0
10
20
-20

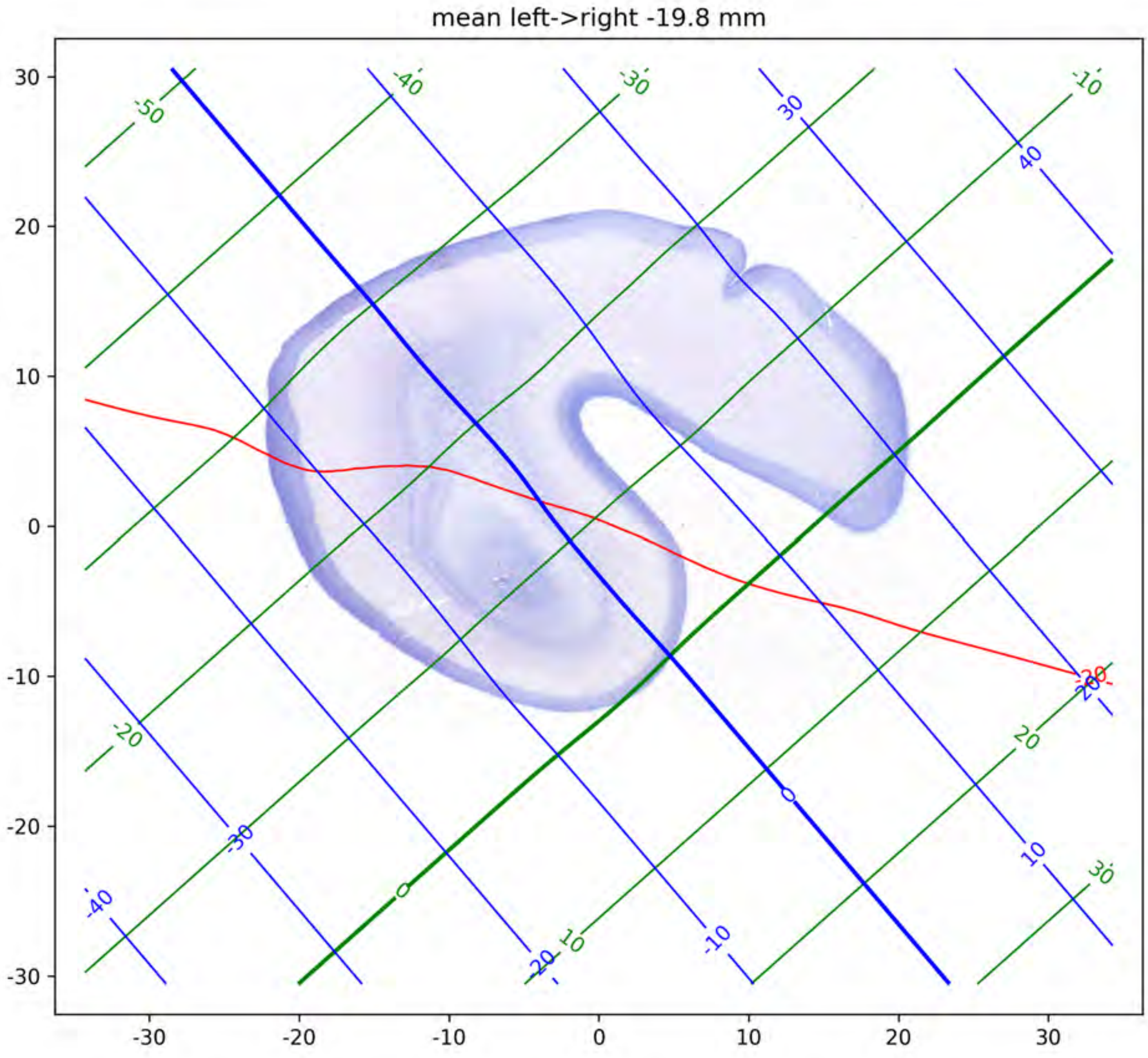

mean left->right -19.8 mm

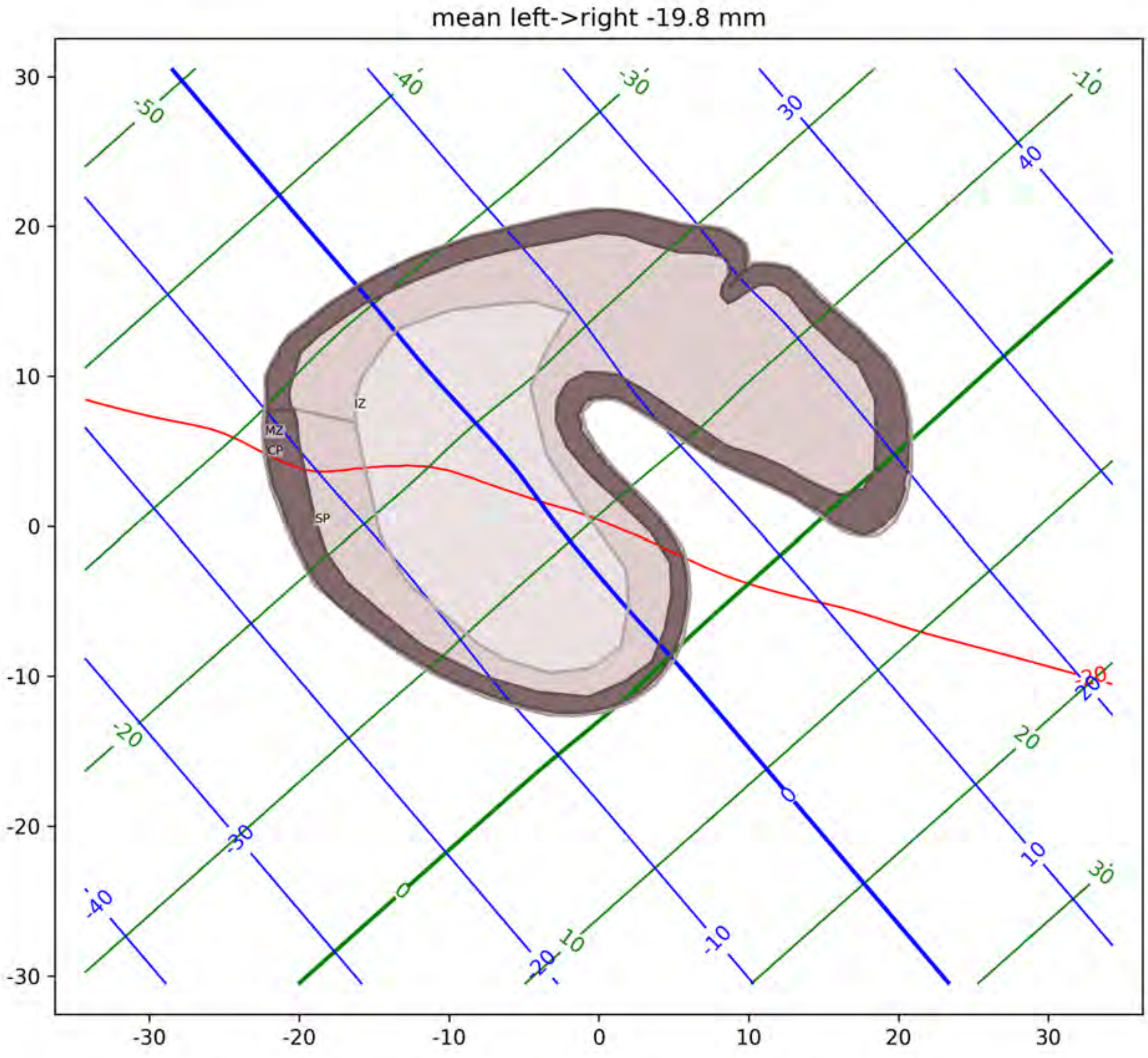

mean left->right -19.8 mm
MZ
CP
SP
IZ

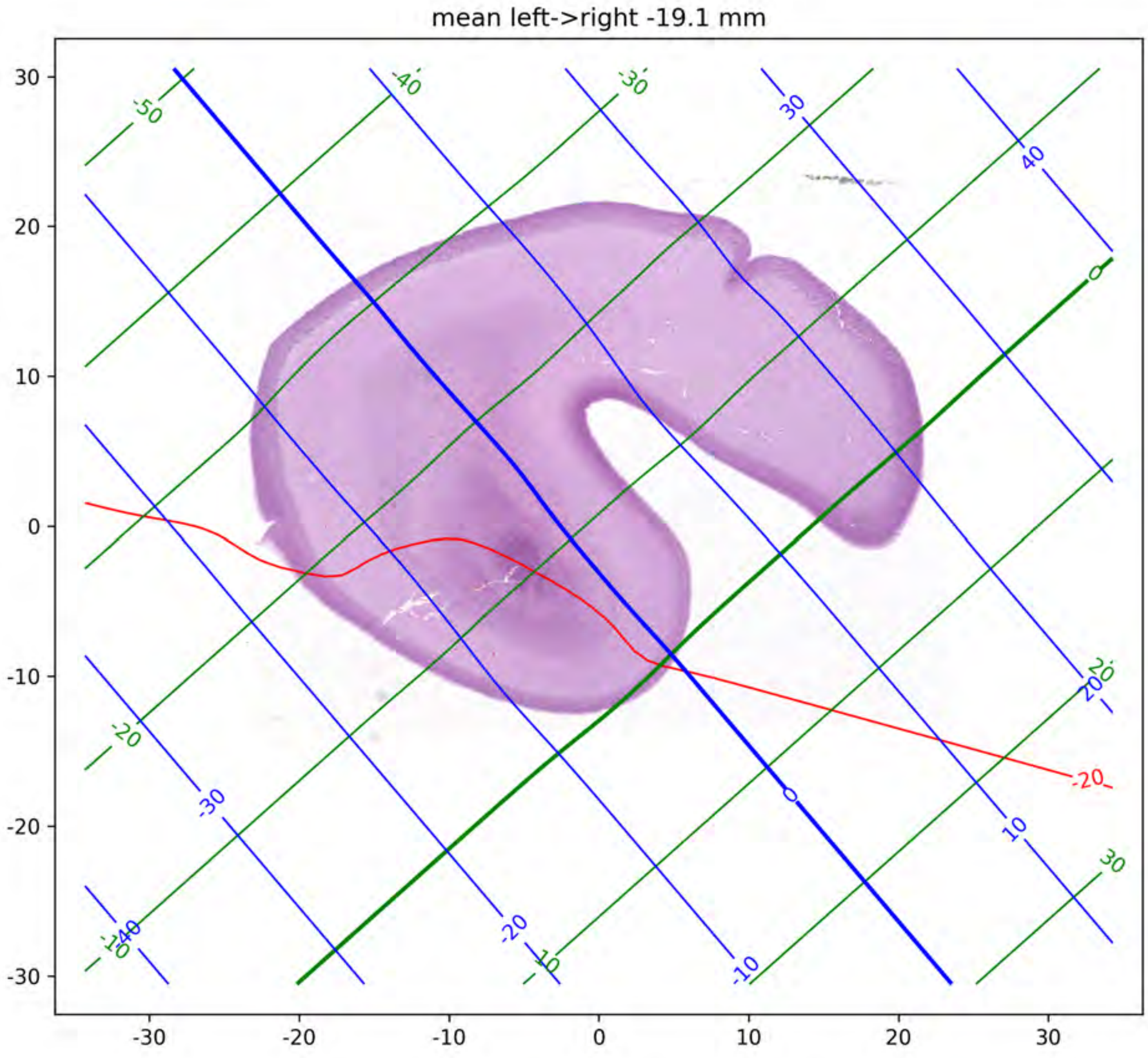

mean left->right -19.1 mm
30
20
10
0
-10
-20
-30
-30
-20
-10
0
10
20
30
-50
-40
-30
-20
-10
0
10
20
30
-20

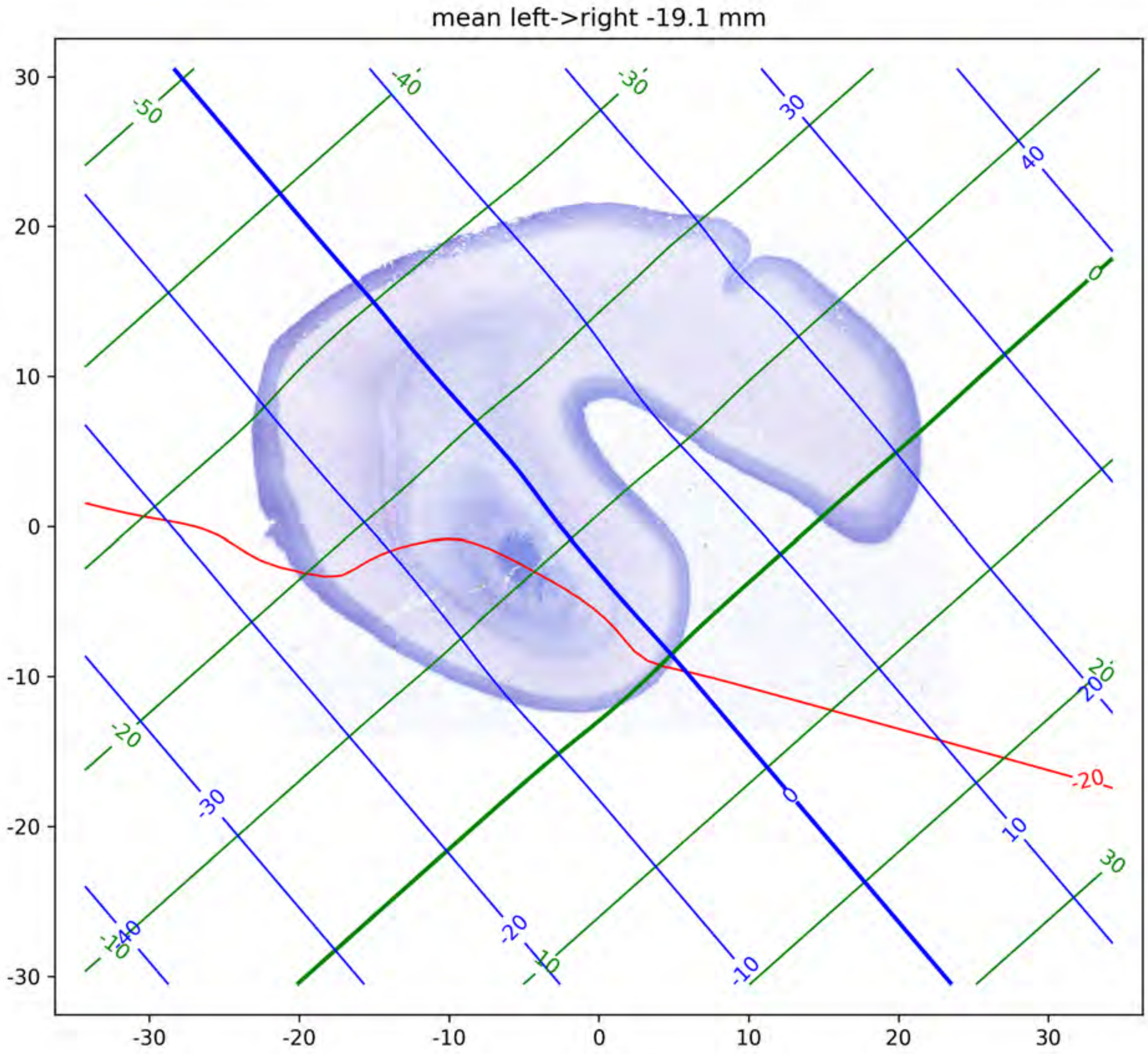

mean left->right -19.1 mm
30
20
10
0
-10
-20
-30
-30
-20
-10
0
10
20
30
-50
-40
-30
-20
-10
0
10
20
30
40
-20

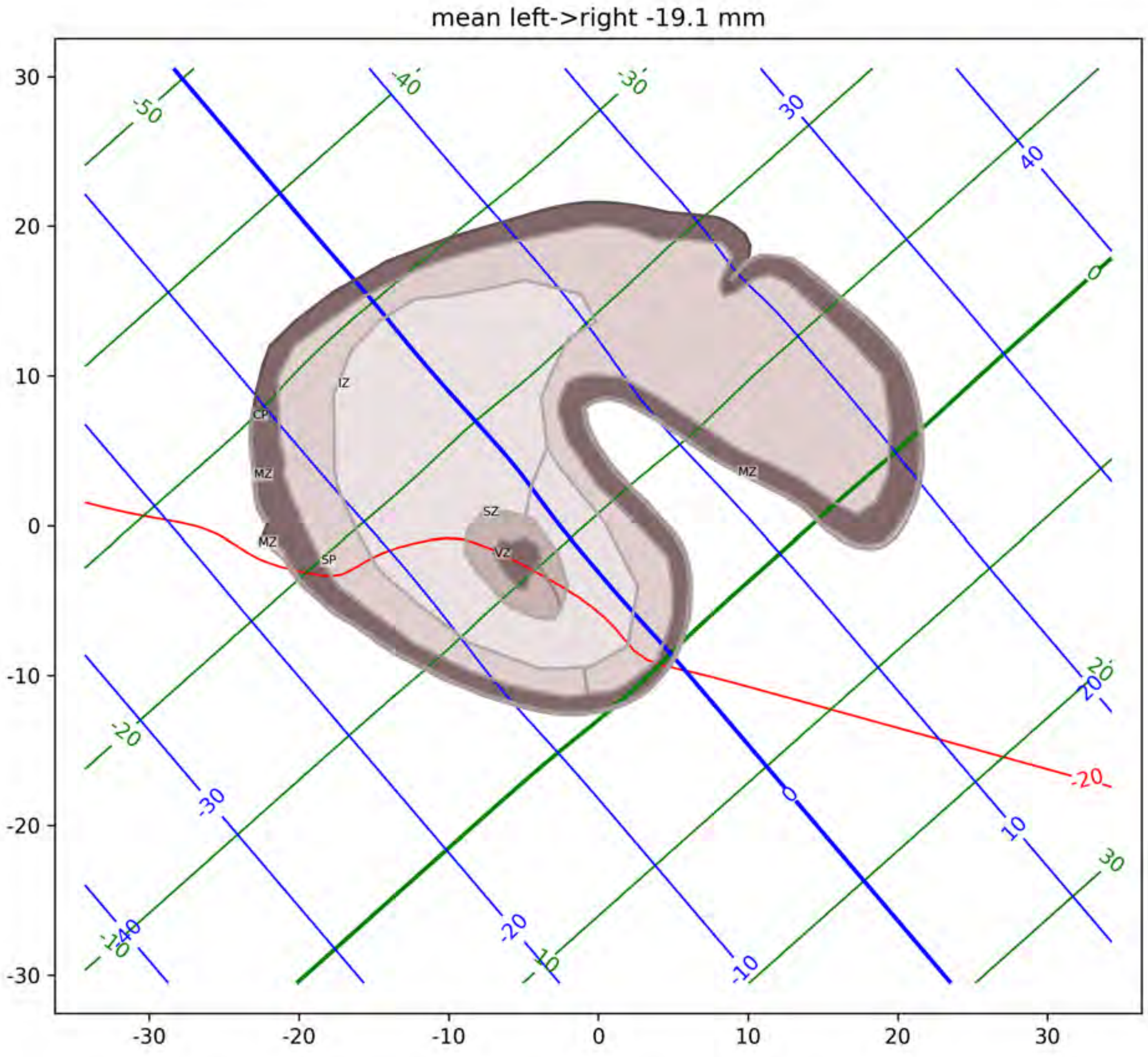

mean left->right -19.1 mm
IZ
CP
MZ
SZ
VZ
SP

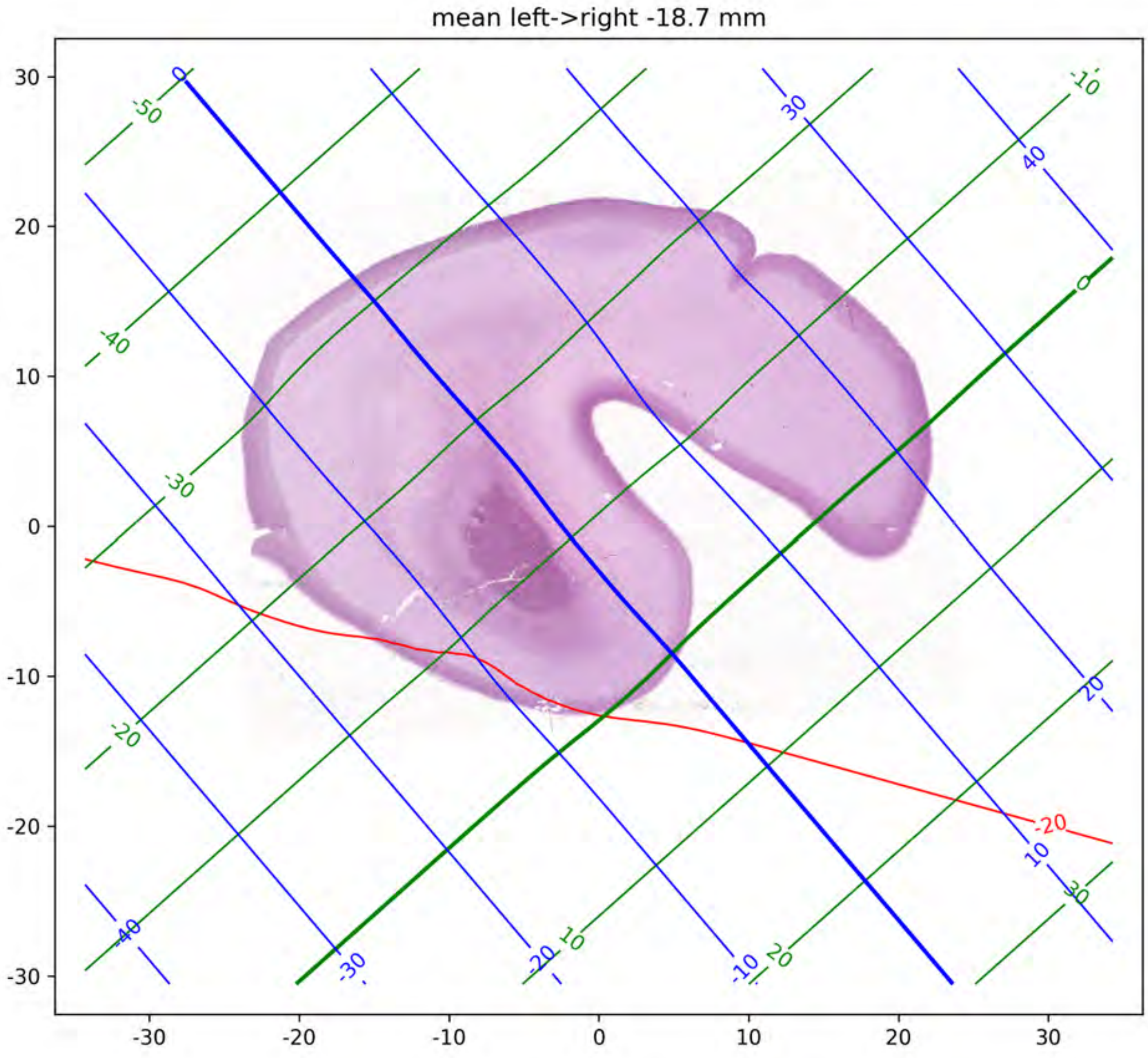
mean left->right -18.7 mm

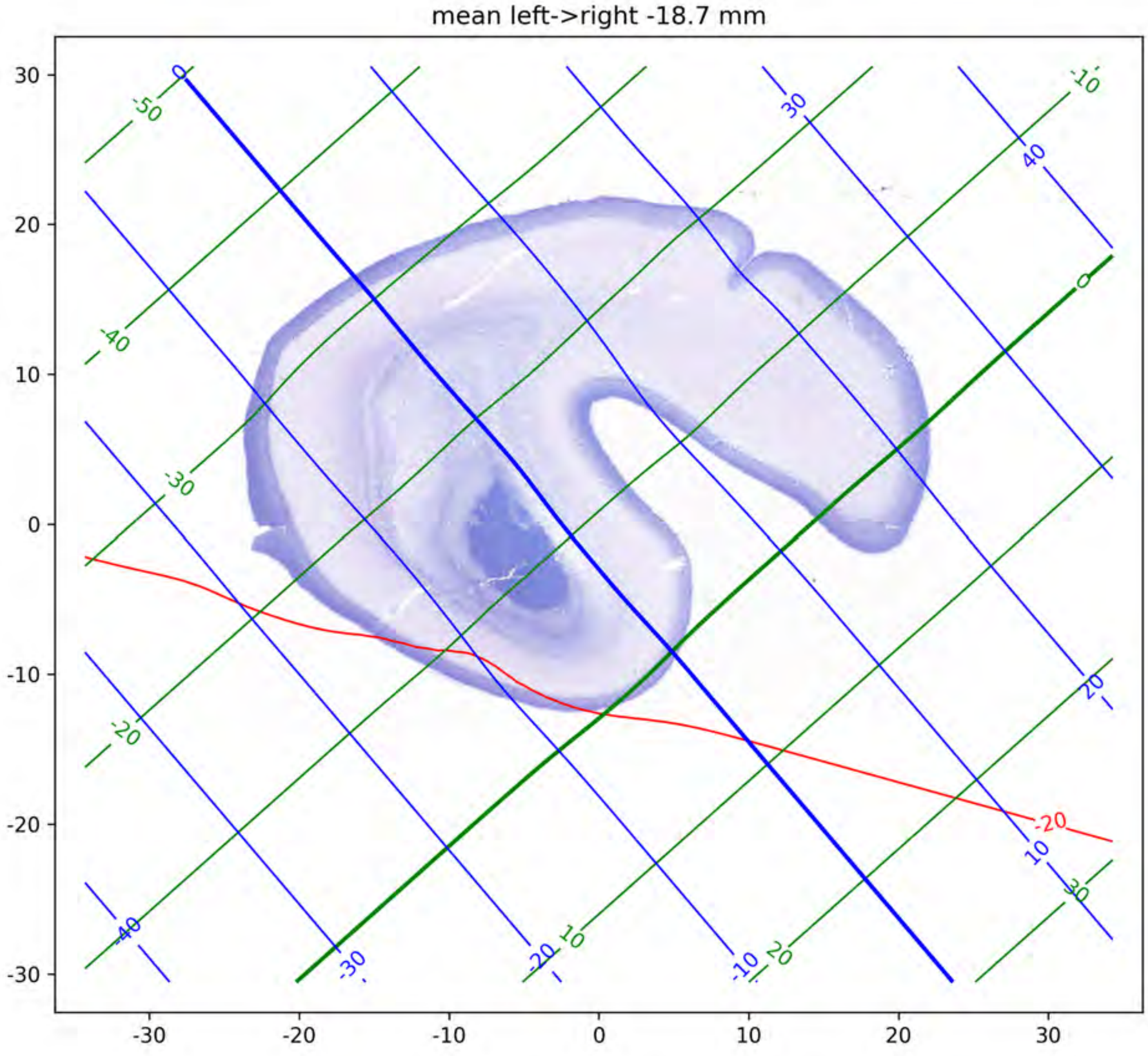
mean left->right -18.7 mm
30
20
10
0
-10
-20
-30
-30
-20
-10
0
10
20
30
-50
-40
-30
-20
-10
0
10
20
30
0
30
40
20
10
-10
-20
-30
-40
-20

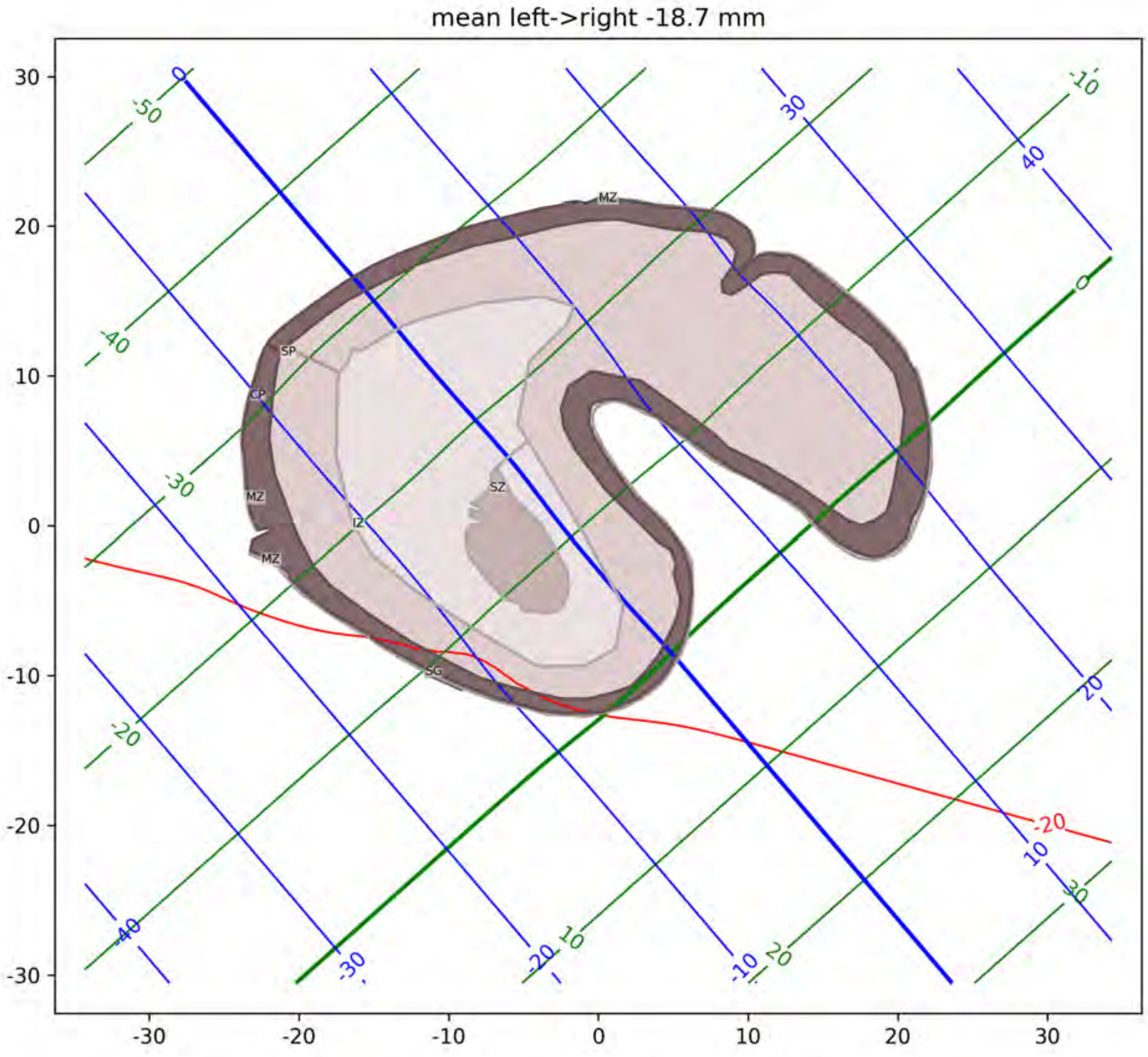
mean left->right -18.7 mm
MZ
SP
CP
SZ
IZ
SG
-20
30
20
10
0
-10
-20
-30

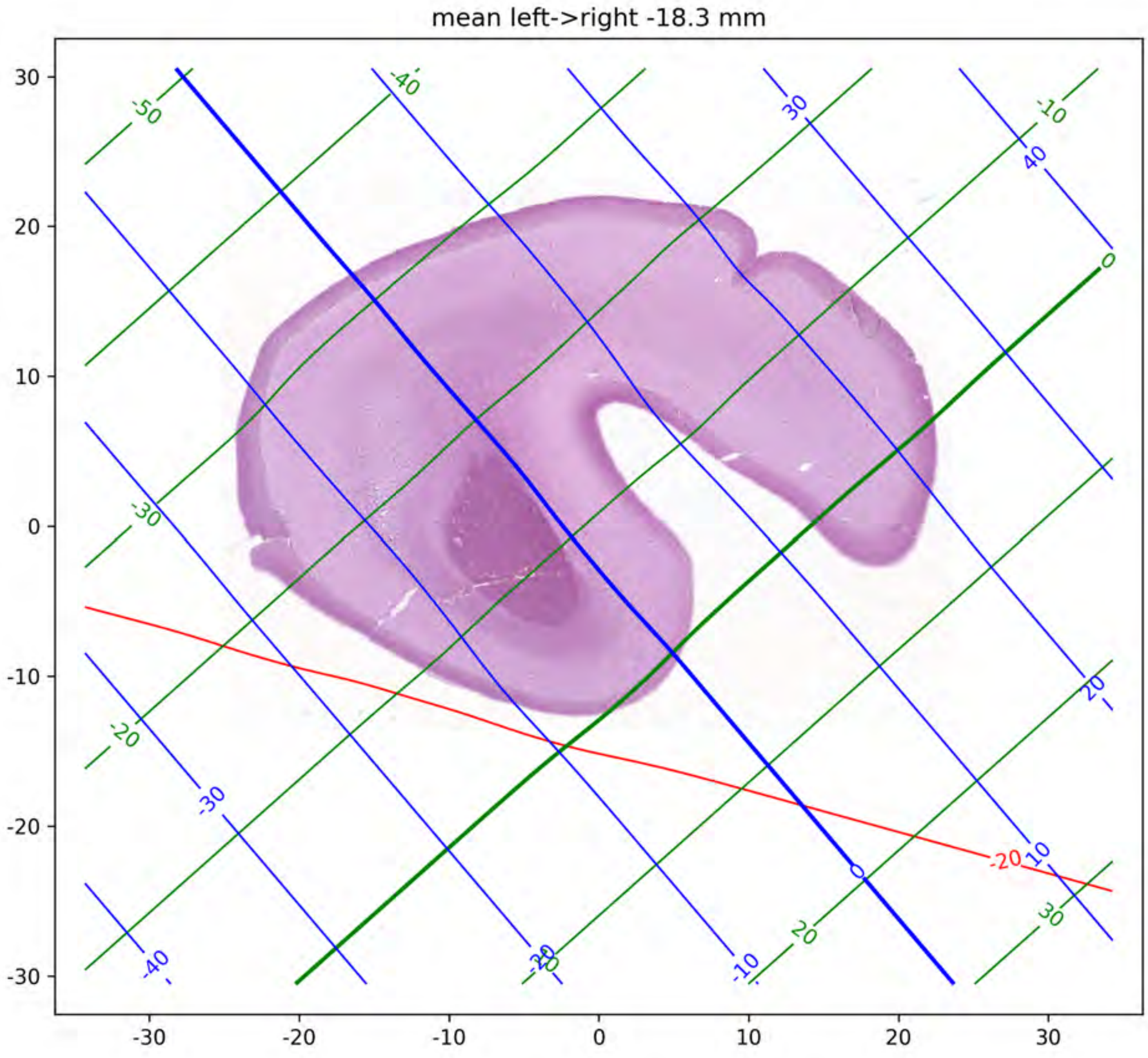
mean left->right -18.3 mm
30
20
10
0
-10
-20
-30
-30
-20
-10
0
10
20
30
-50
-40
30
-10
40
0
-30
20
-20
-30
-20
10
0
20
30
-40
-20
-10

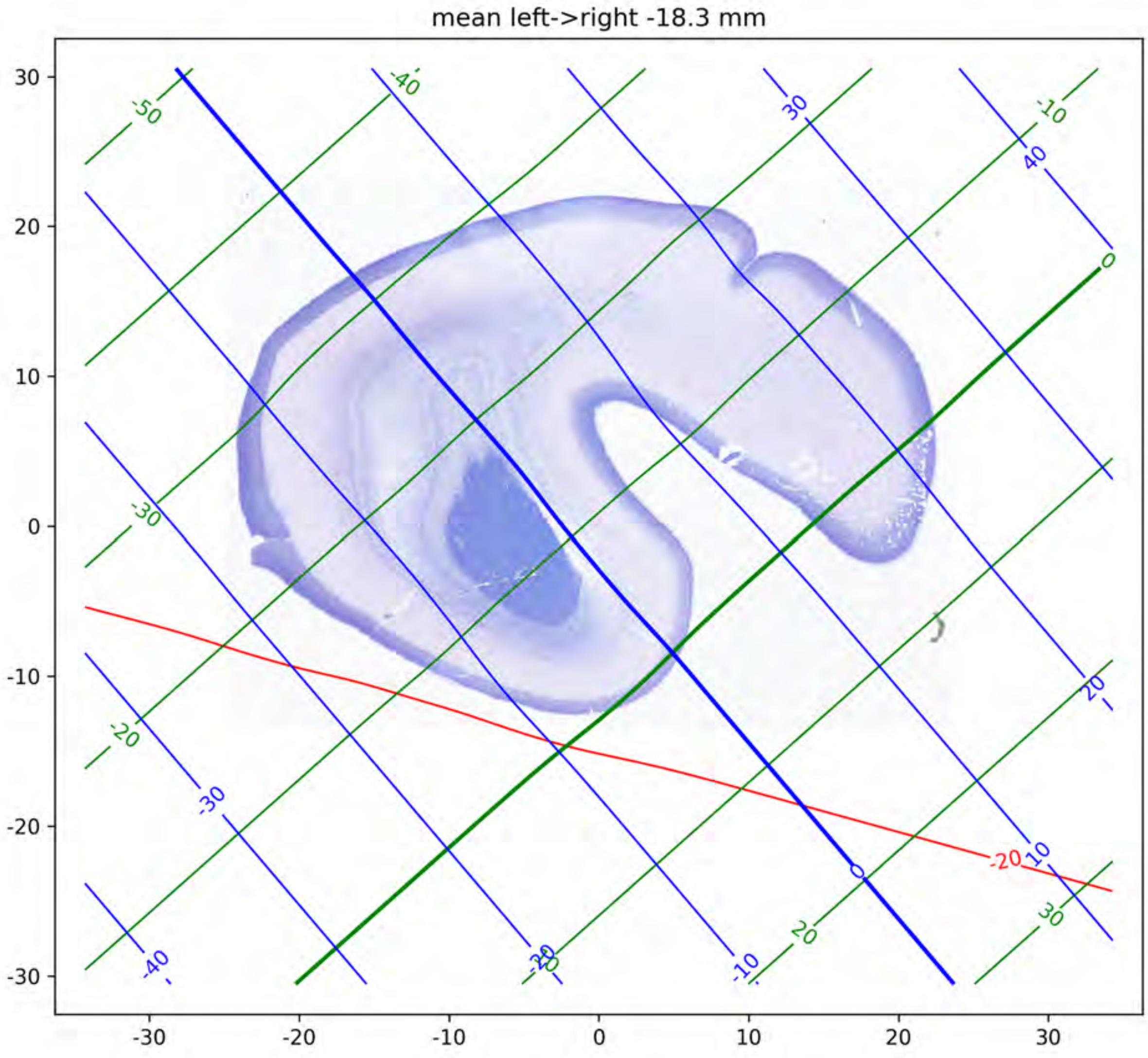

mean left->right -18.3 mm

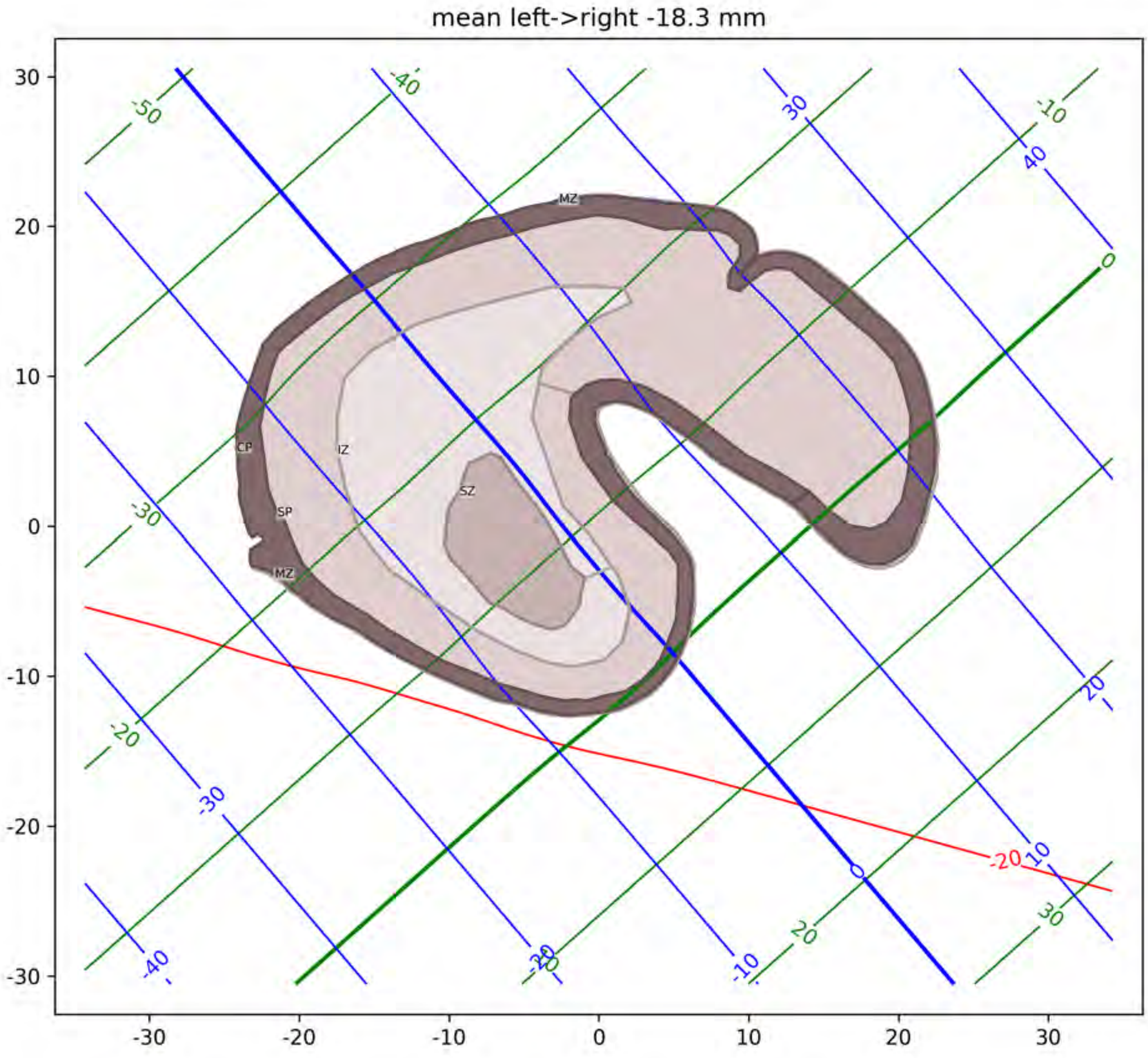
mean left->right -18.3 mm
MZ
CP
IZ
SZ
SP
MZ
-50
-40
-30
-20
-10
0
10
20
30
40
-20

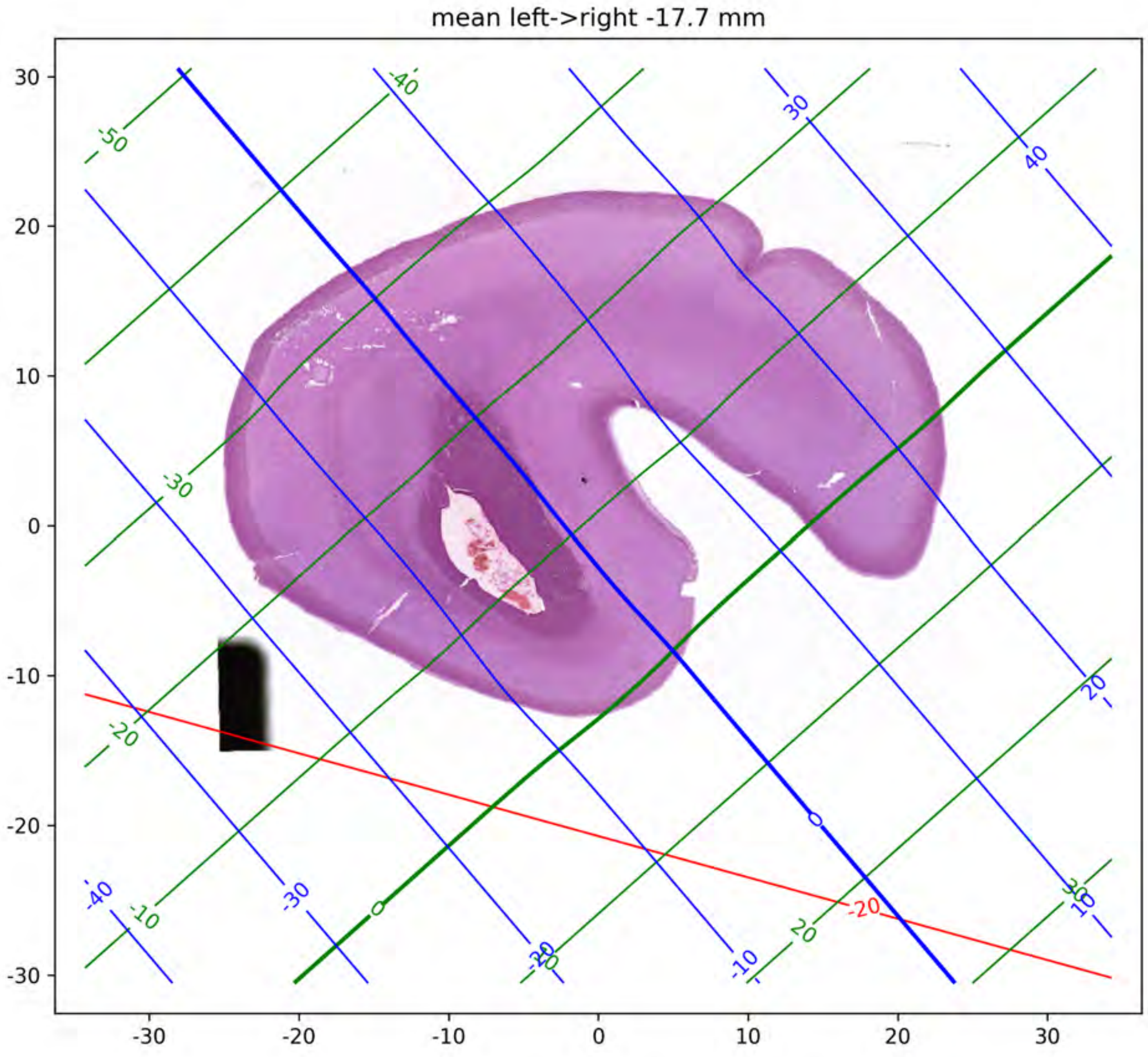
mean left->right -17.7 mm
30
20
10
0
-10
-20
-30
-30
-20
-10
0
10
20
30
-50
-40
-30
-20
-10
0
10
20
30
40
-20

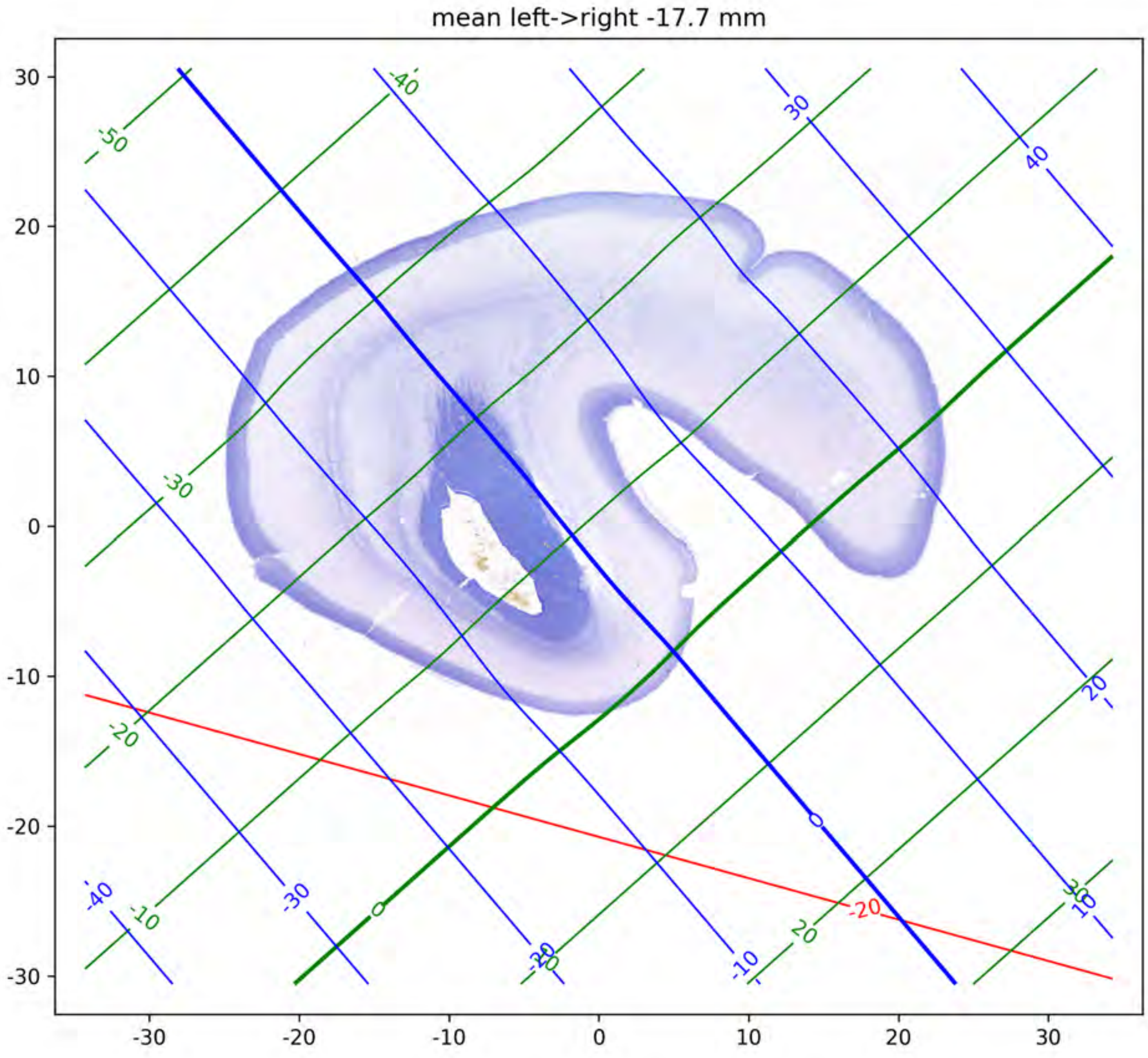
mean left->right -17.7 mm
30
20
10
0
-10
-20
-30
-30
-20
-10
0
10
20
30
-50
-40
30
40
-30
-20
-10
0
20
30
-40
-30
-20
-10
0
10
20
-20

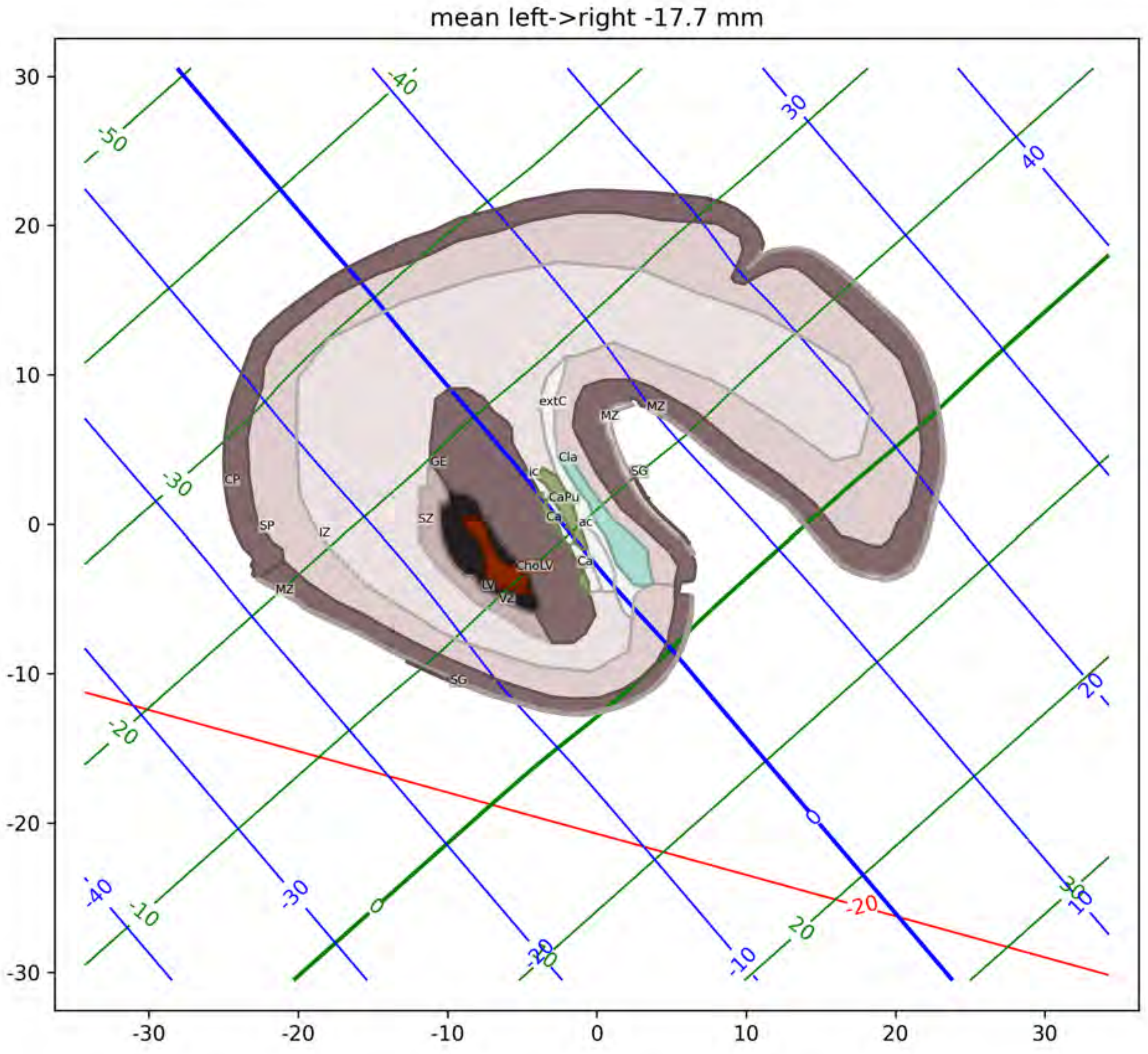
mean left->right -17.7 mm
extC
MZ
MZ
Cla
ic
GE
SG
CaPu
Ca
ac
SZ
CP
SP
IZ
Ca
ChoLV
LV
VZ
MZ
SG

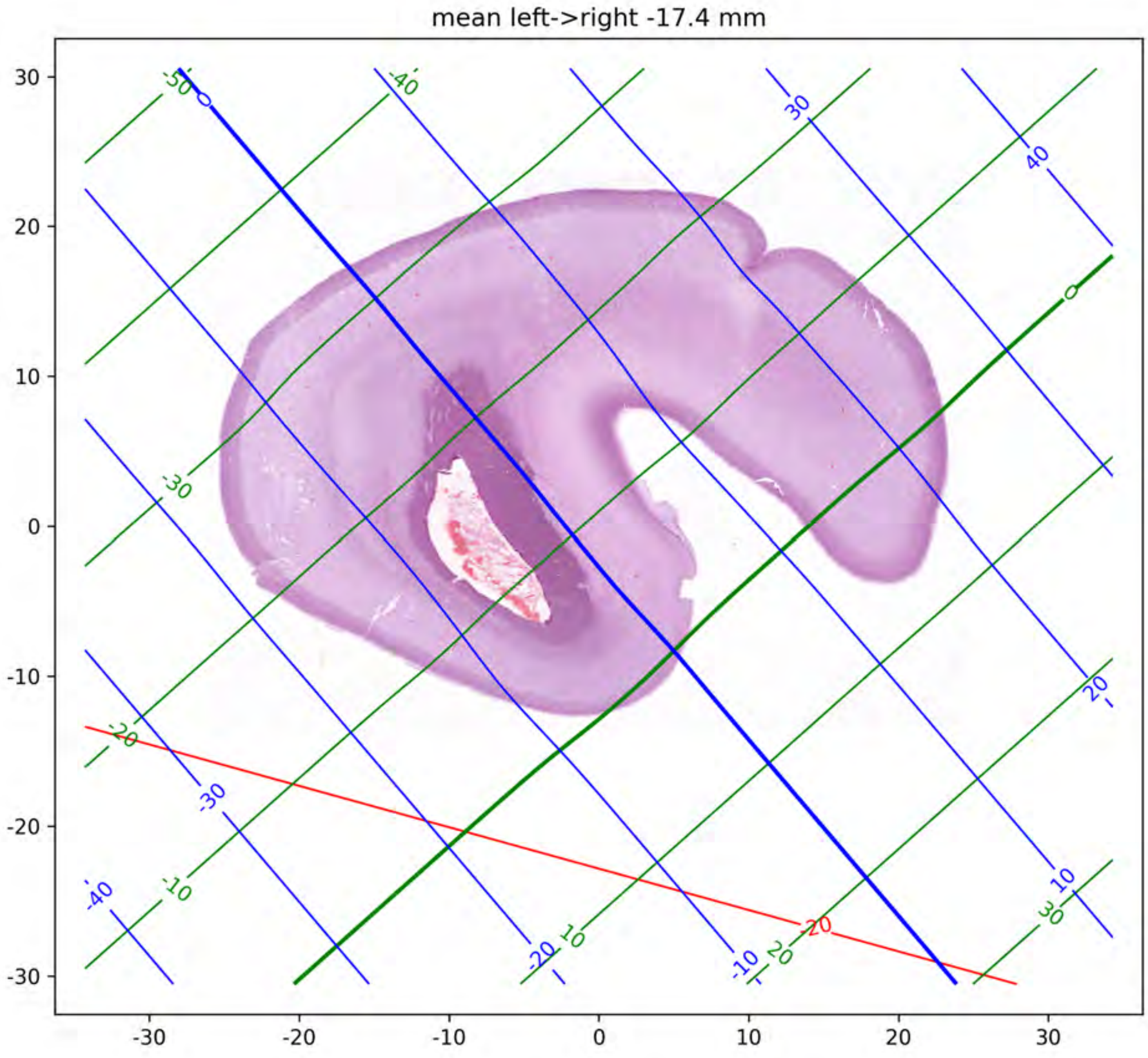
mean left->right -17.4 mm
30
20
10
0
-10
-20
-30
-30
-20
-10
0
10
20
30
-50
0
-40
30
40
0
-30
-20
-30
-40
-10
-20
10
-10
20
-20
20
10
30

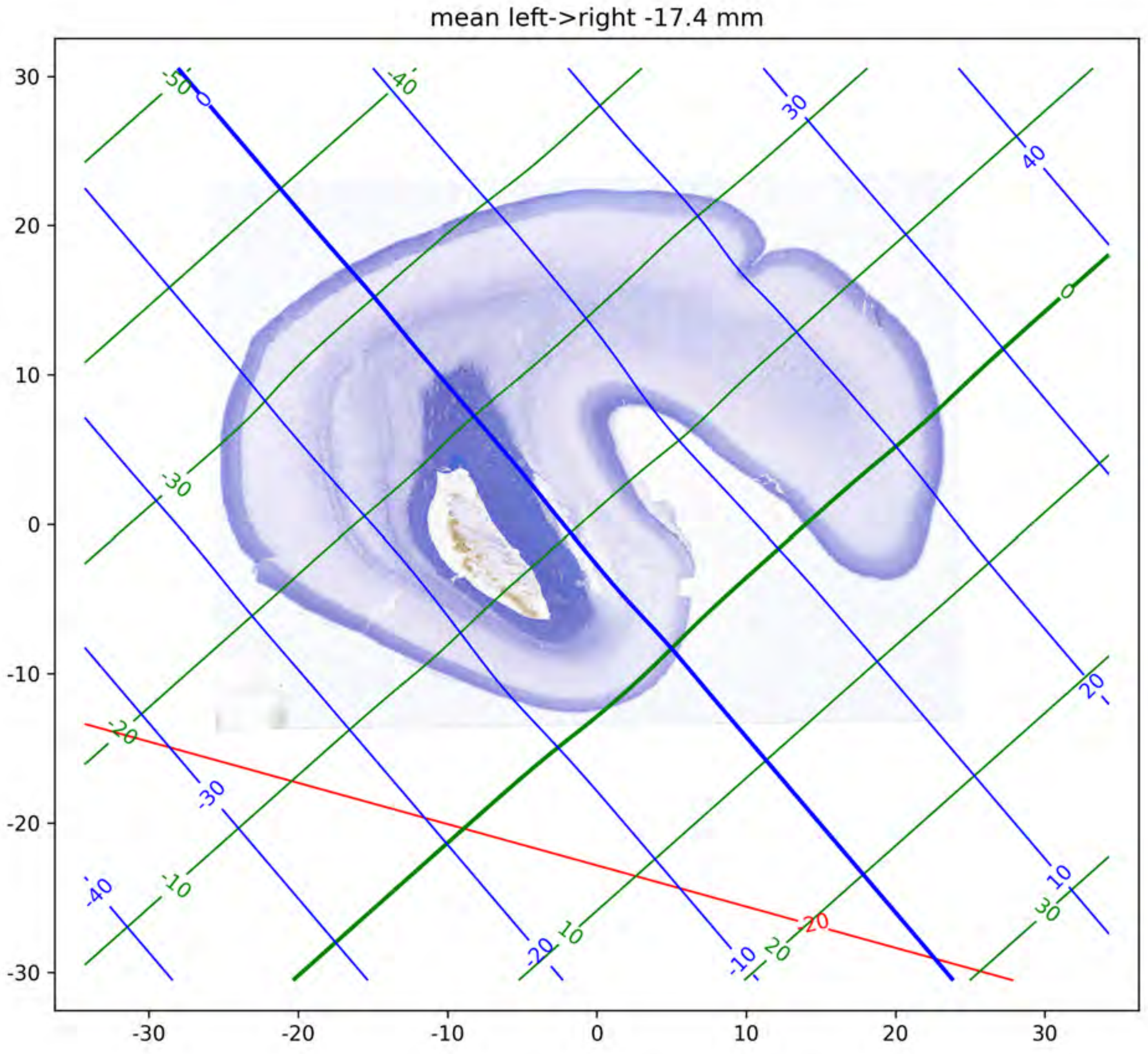

mean left->right -17.4 mm

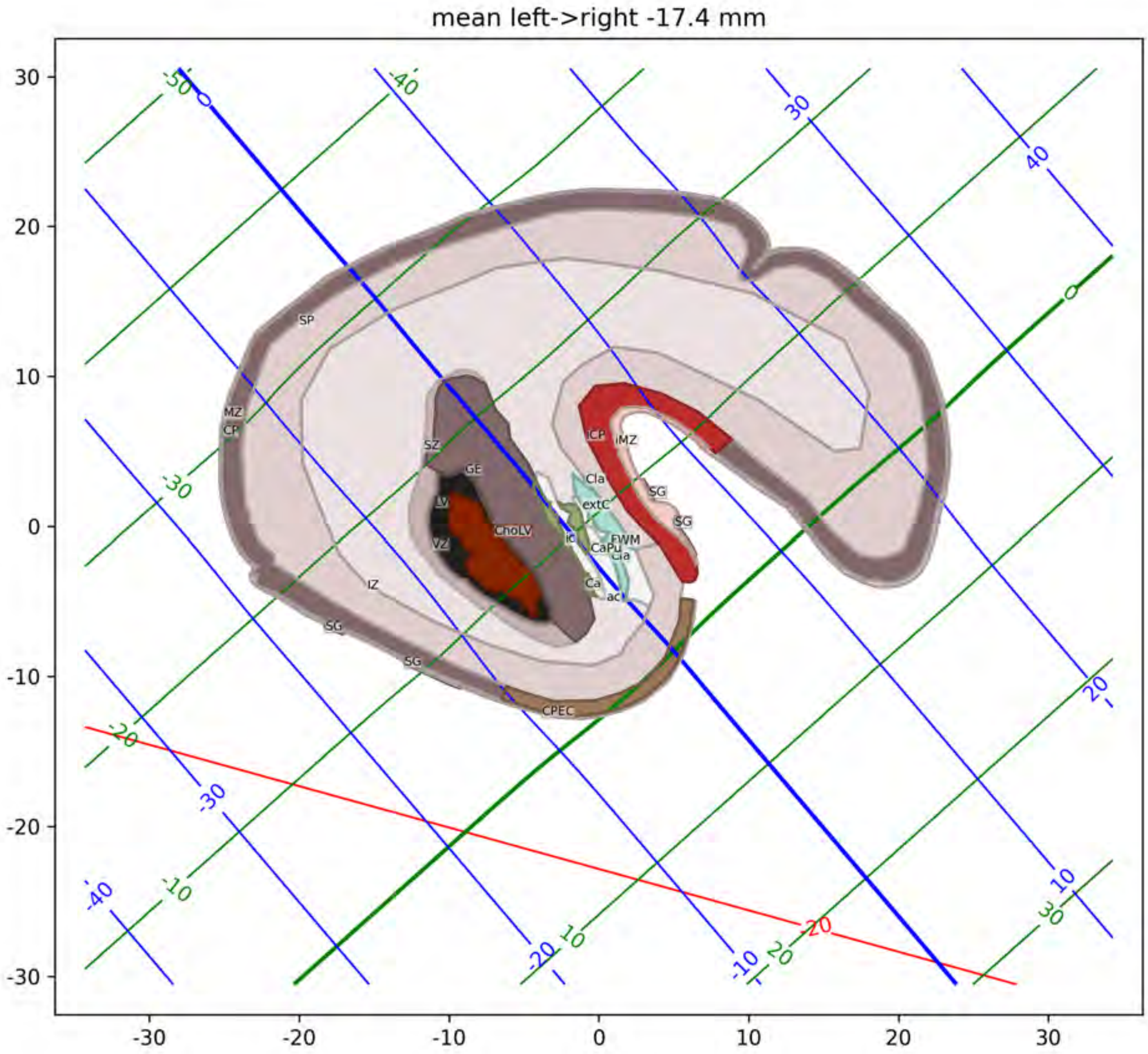

mean left->right -17.4 mm
SP
MZ
CP
SZ
GE
LV
VZ
ChoLV
IZ
SG
CPEC
Cla
extC
ic
CaPu
FWM
Ca
ac
iMZ

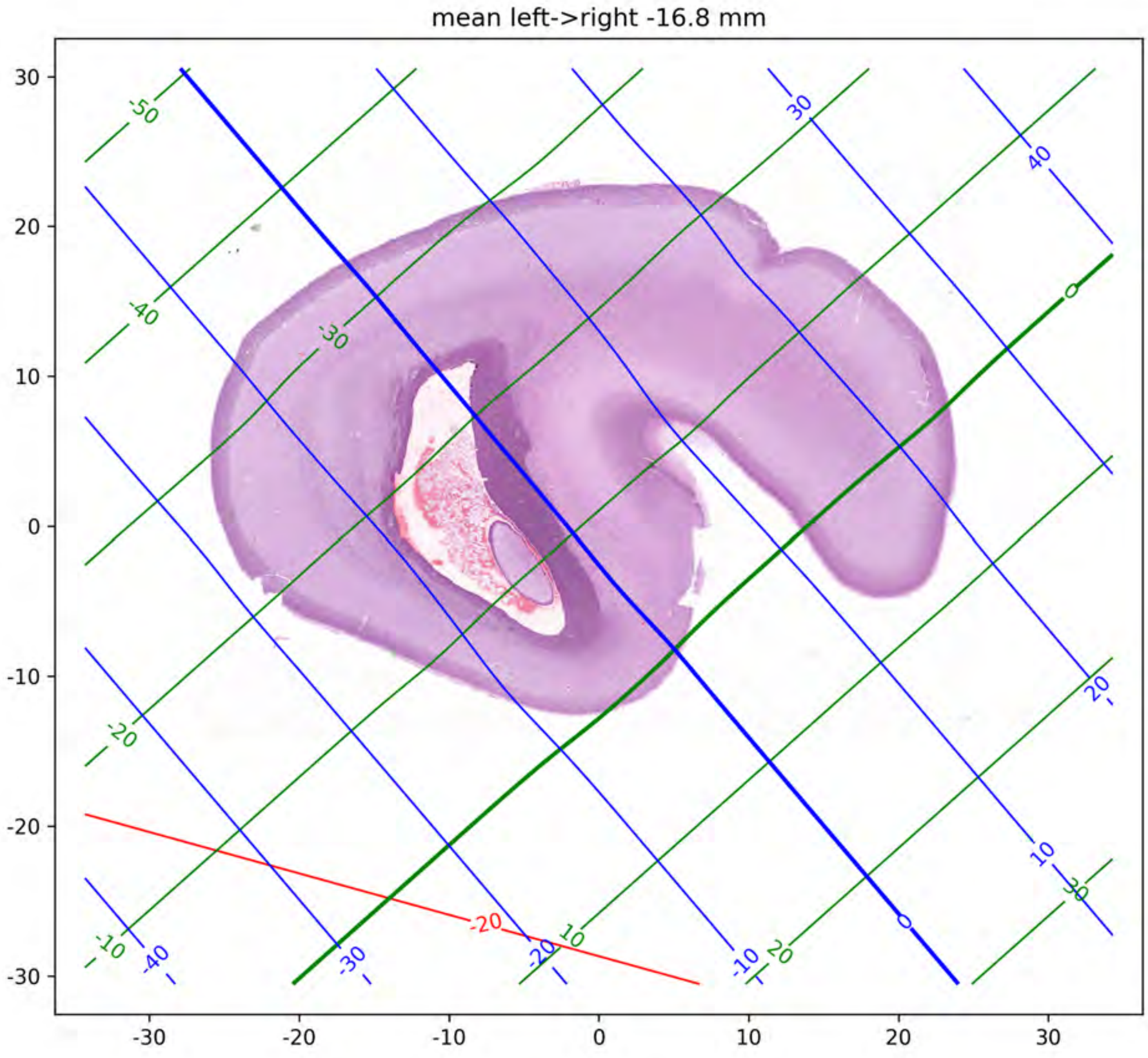

mean left->right -16.8 mm

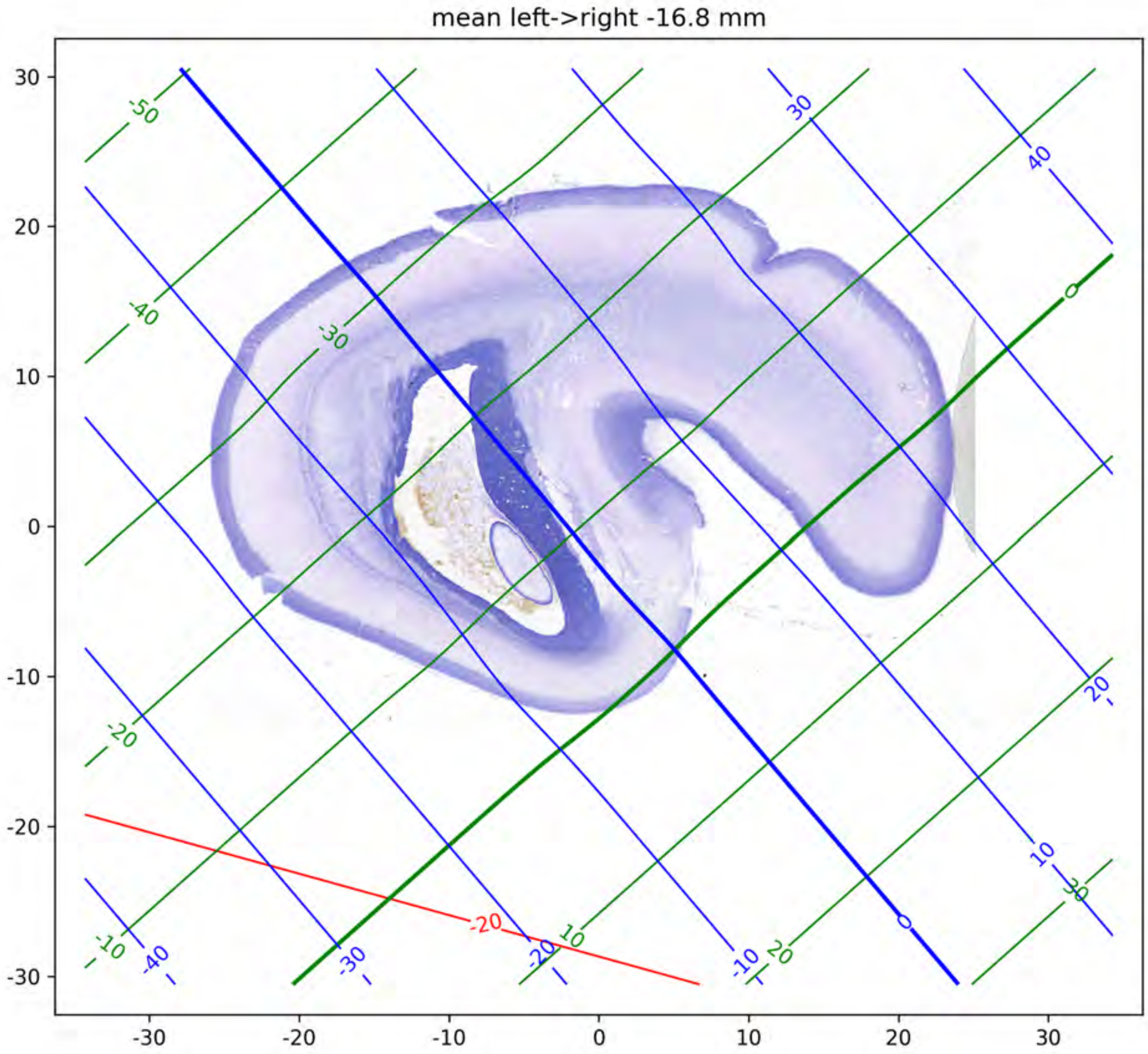
mean left->right -16.8 mm
30
20
10
0
-10
-20
-30
-30
-20
-10
0
10
20
30

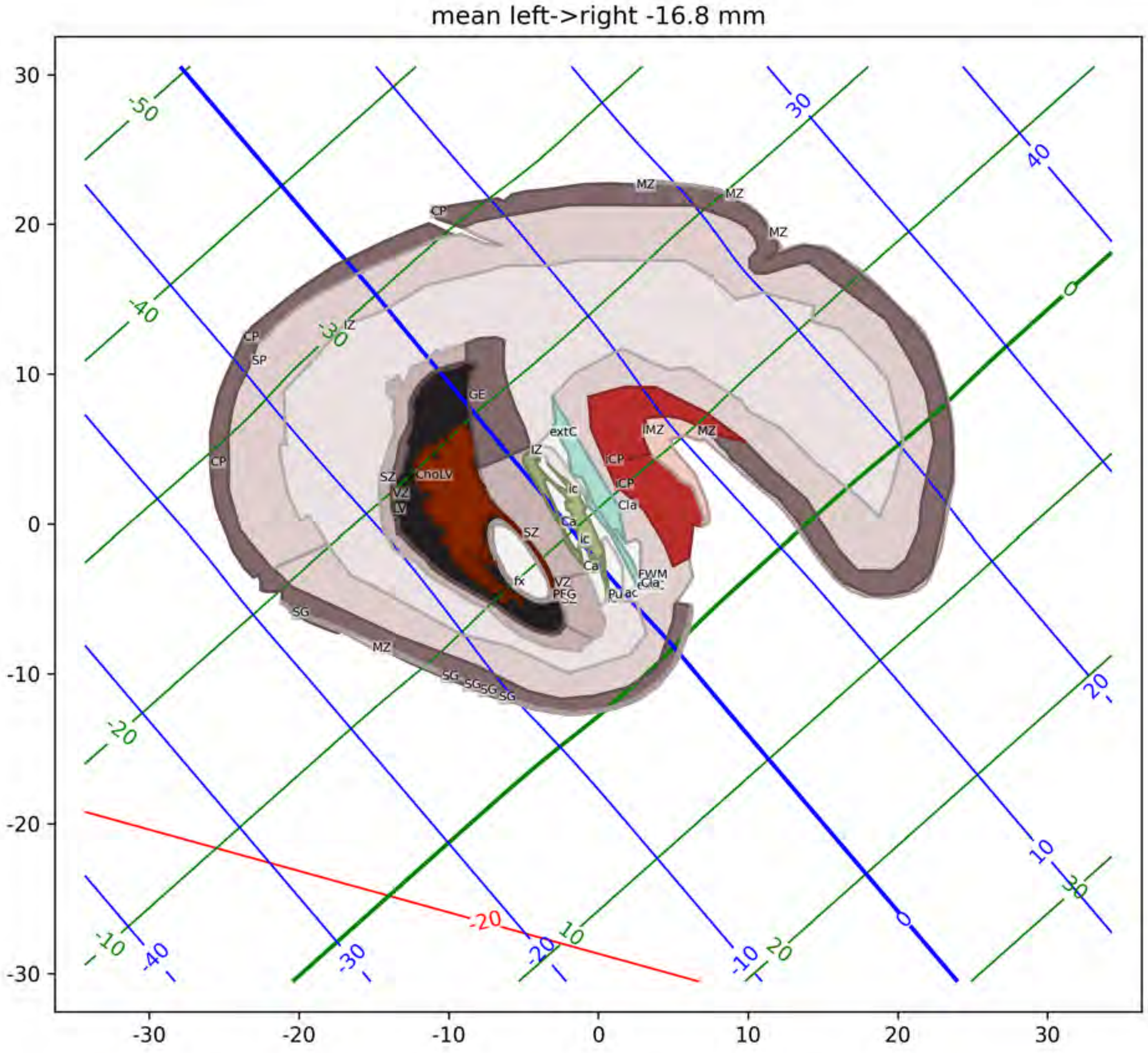

mean left->right -16.8 mm
MZ
CP
SP
IZ
GE
extC
SZ
ChoLV
VZ
LV
Ca
ic
Cla
FWM
Pu
ac
fx
PFG
SG
-50
-40
-30
-20
-10
0
10
20
30
40

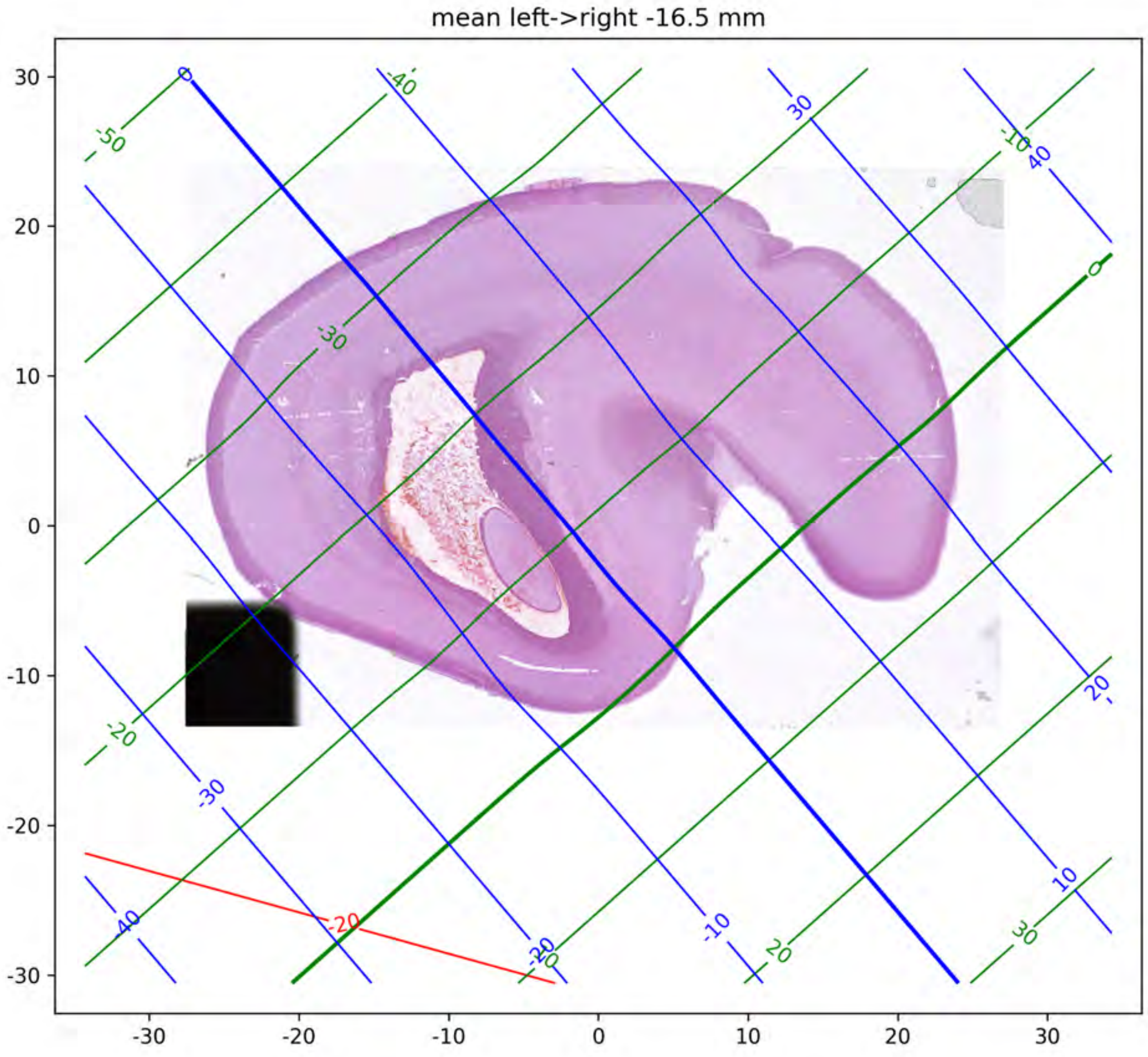
mean left->right -16.5 mm
30
20
10
0
-10
-20
-30
-30
-20
-10
0
10
20
30

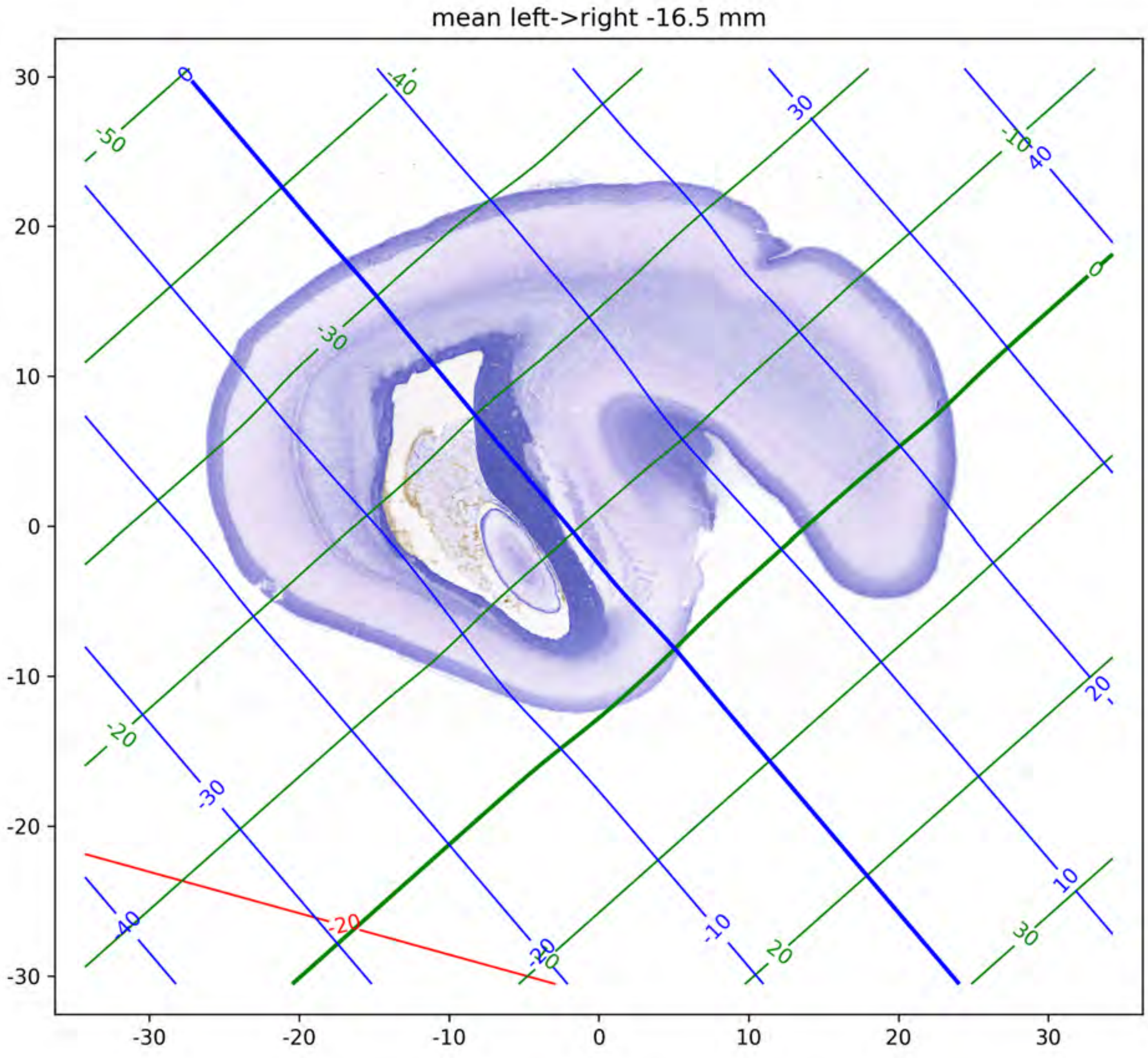
mean left->right -16.5 mm

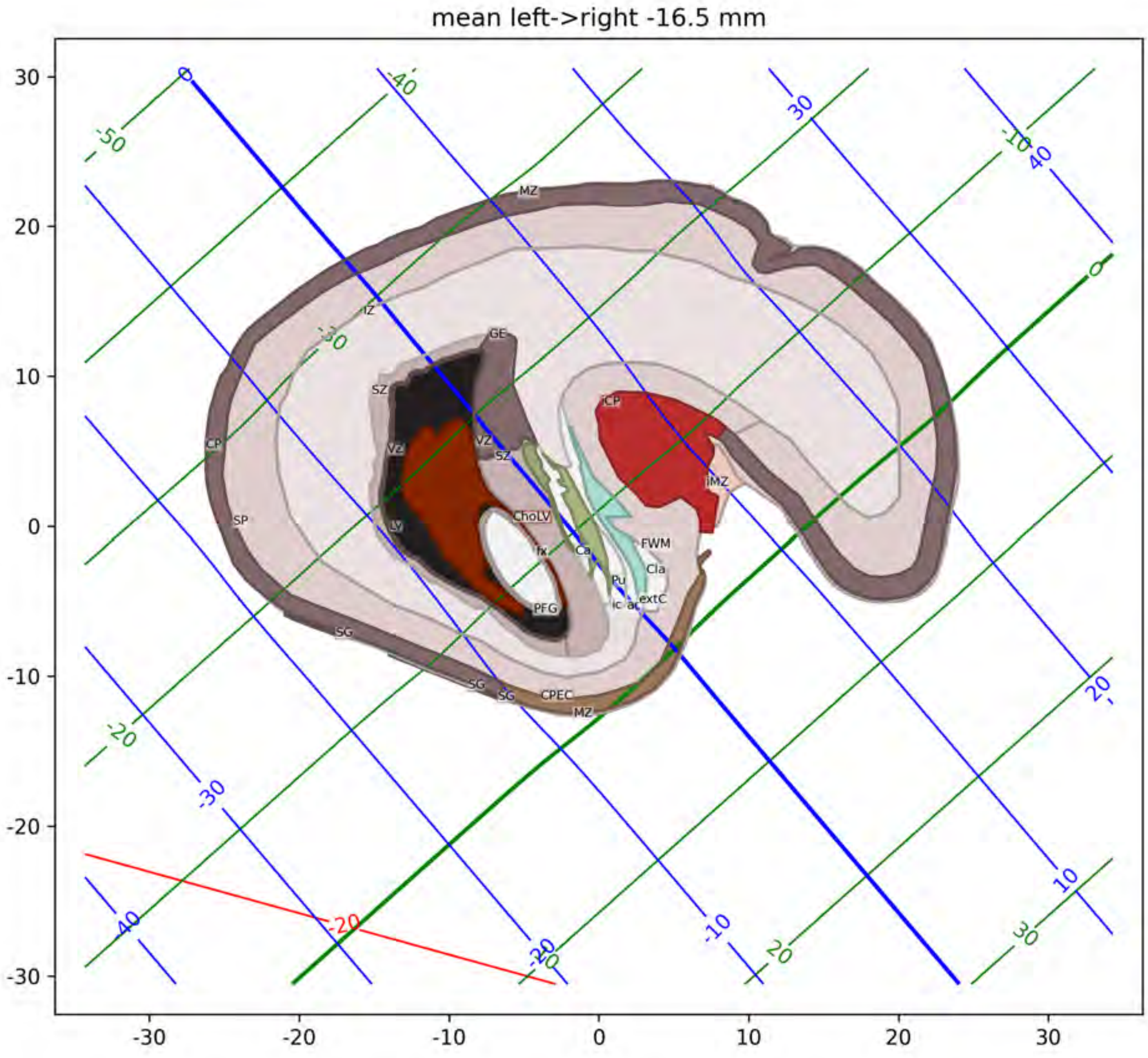
mean left->right -16.5 mm
MZ
IZ
GE
SZ
CP
VZ
SP
LV
SZ
ChoLV
fx
Ca
PFG
FWM
Cla
Pu
ic
ac
extC
iMZ
SG
CPEC

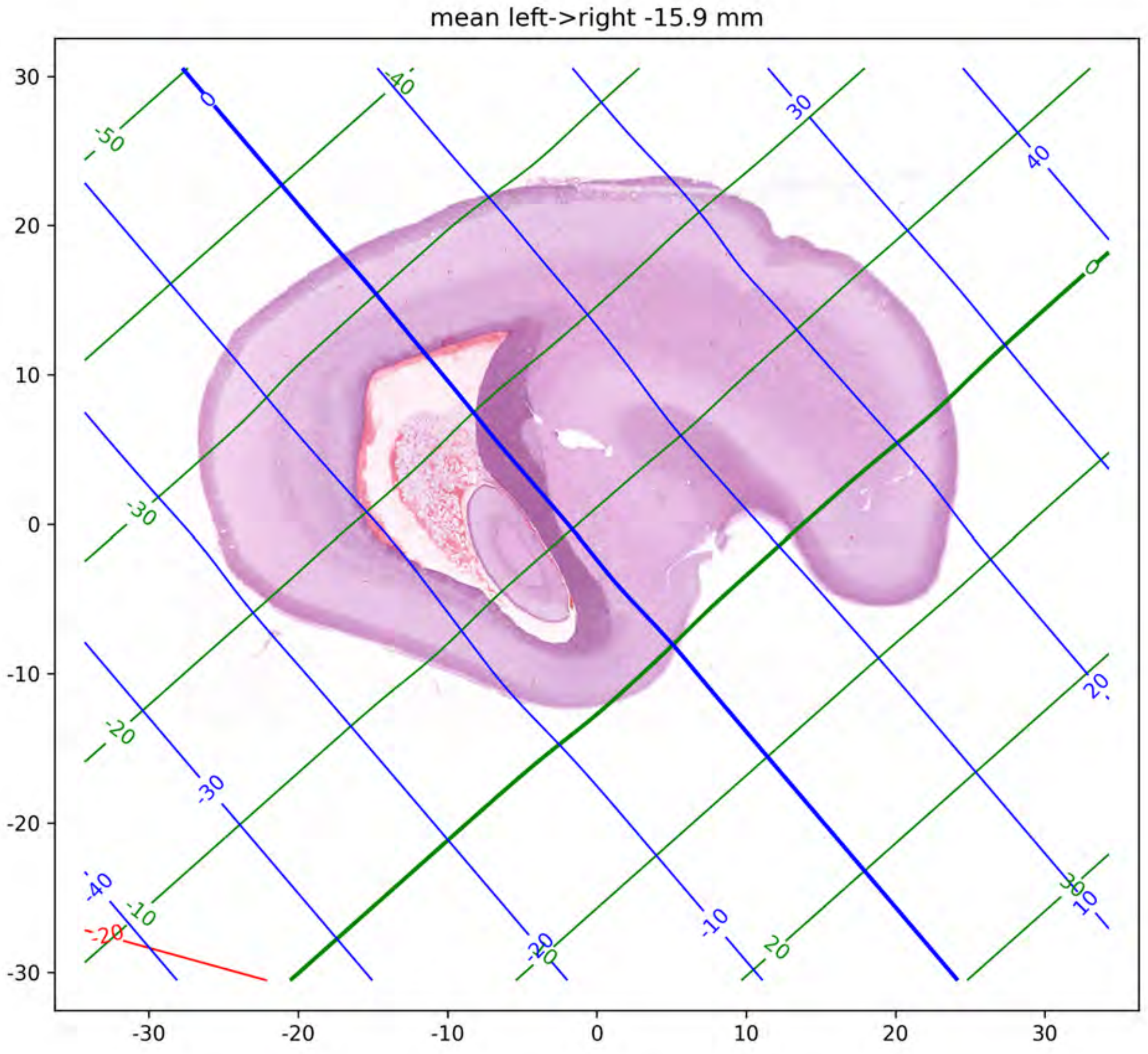
mean left->right -15.9 mm

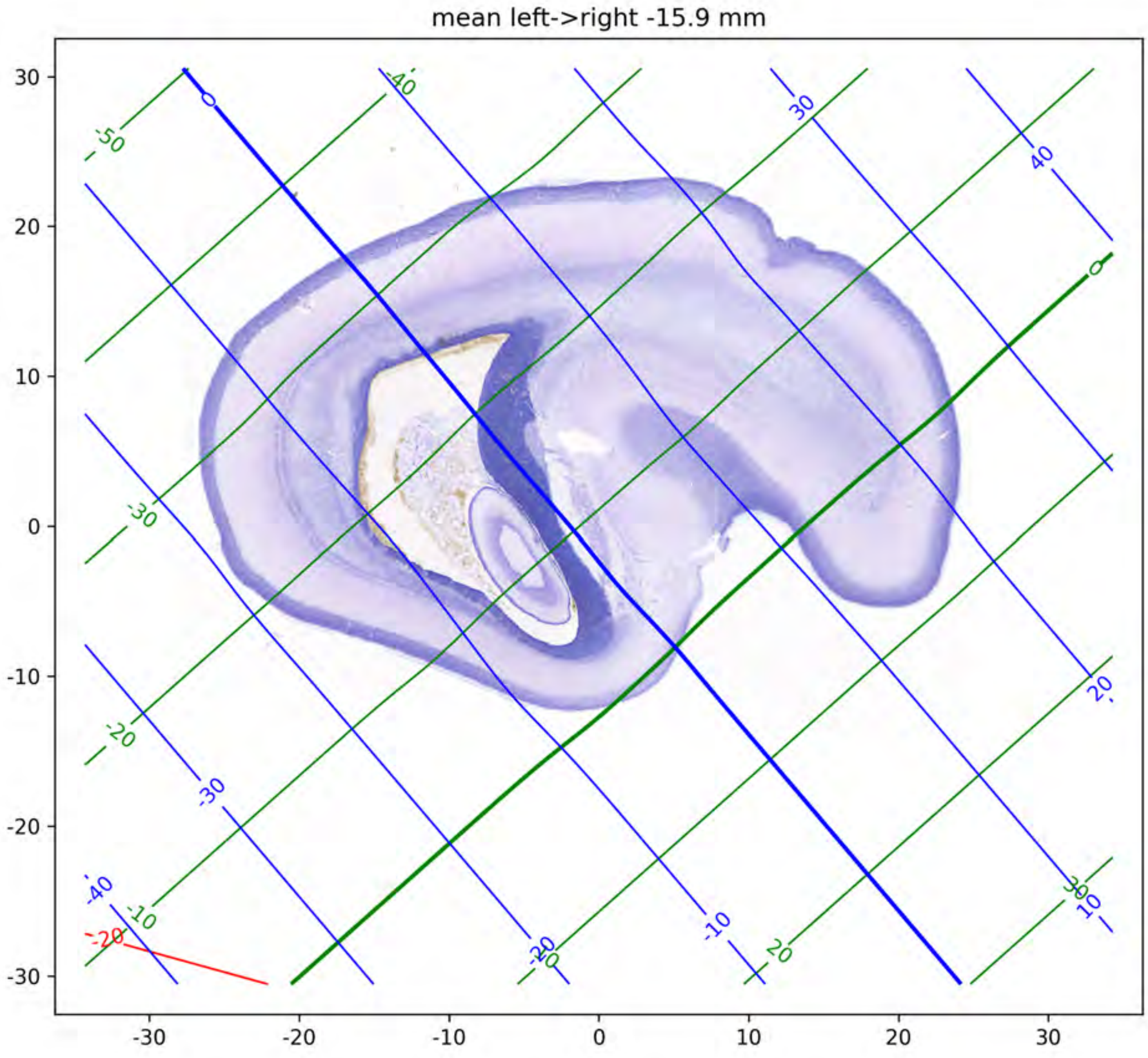

mean left->right -15.9 mm

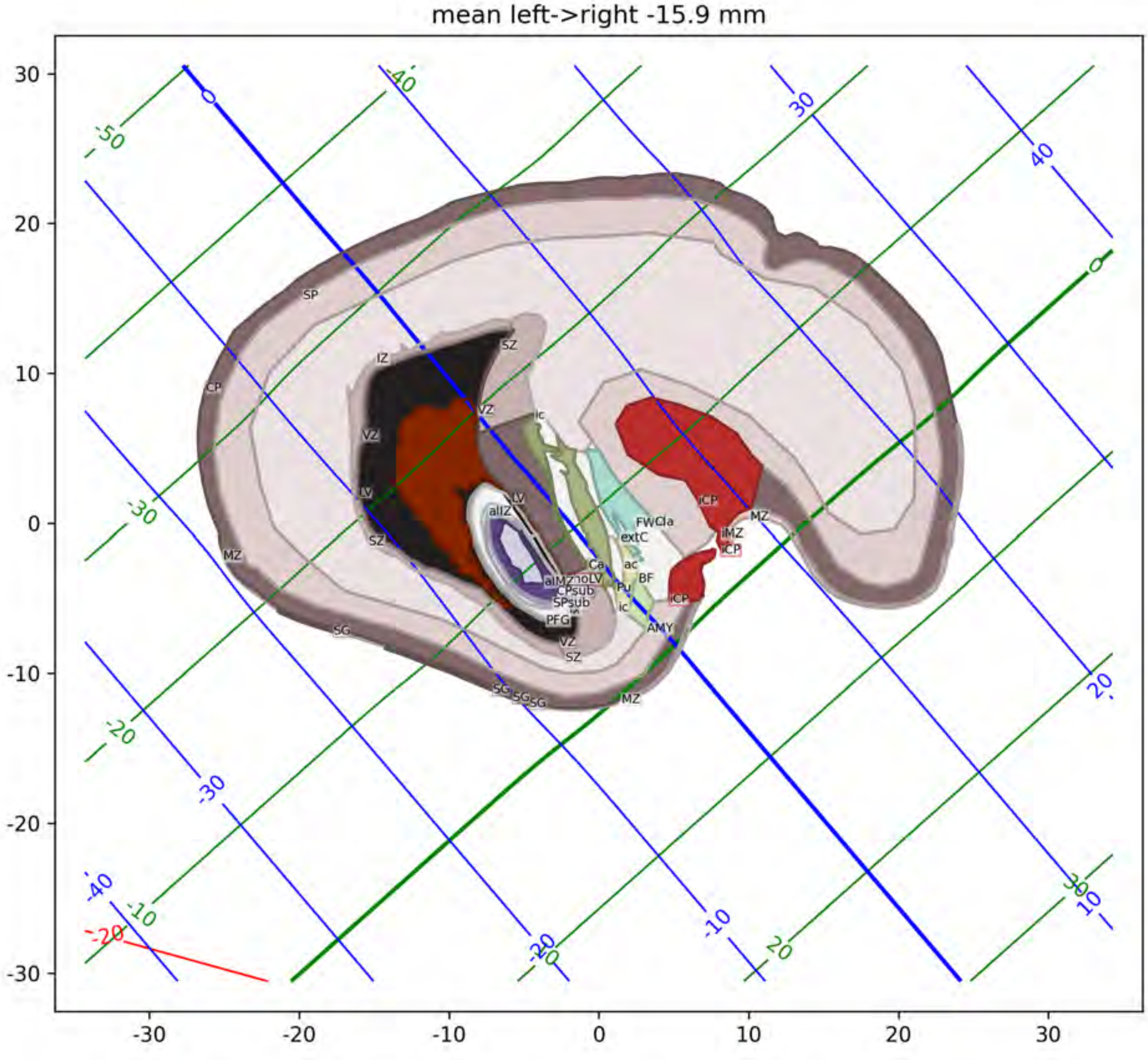
mean left->right -15.9 mm
SP
IZ
SZ
CP
VZ
ic
LV
aIIZ
MZ
iCP
FWCla
extC
iMZ
Ca
ac
BF
Pu
AMY
PFG
SG

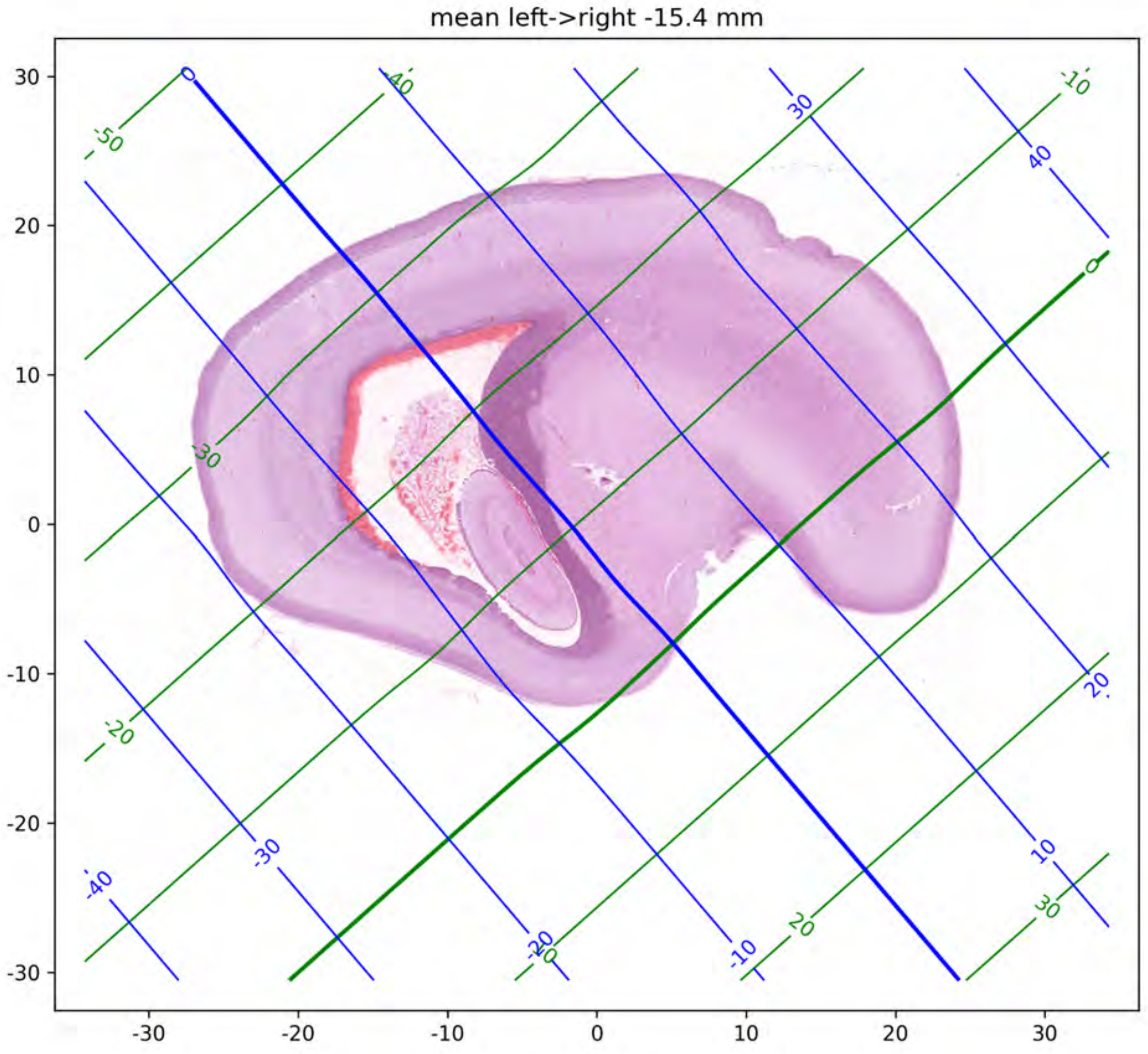
mean left->right -15.4 mm
30
20
10
0
-10
-20
-30
-30
-20
-10
0
10
20
30

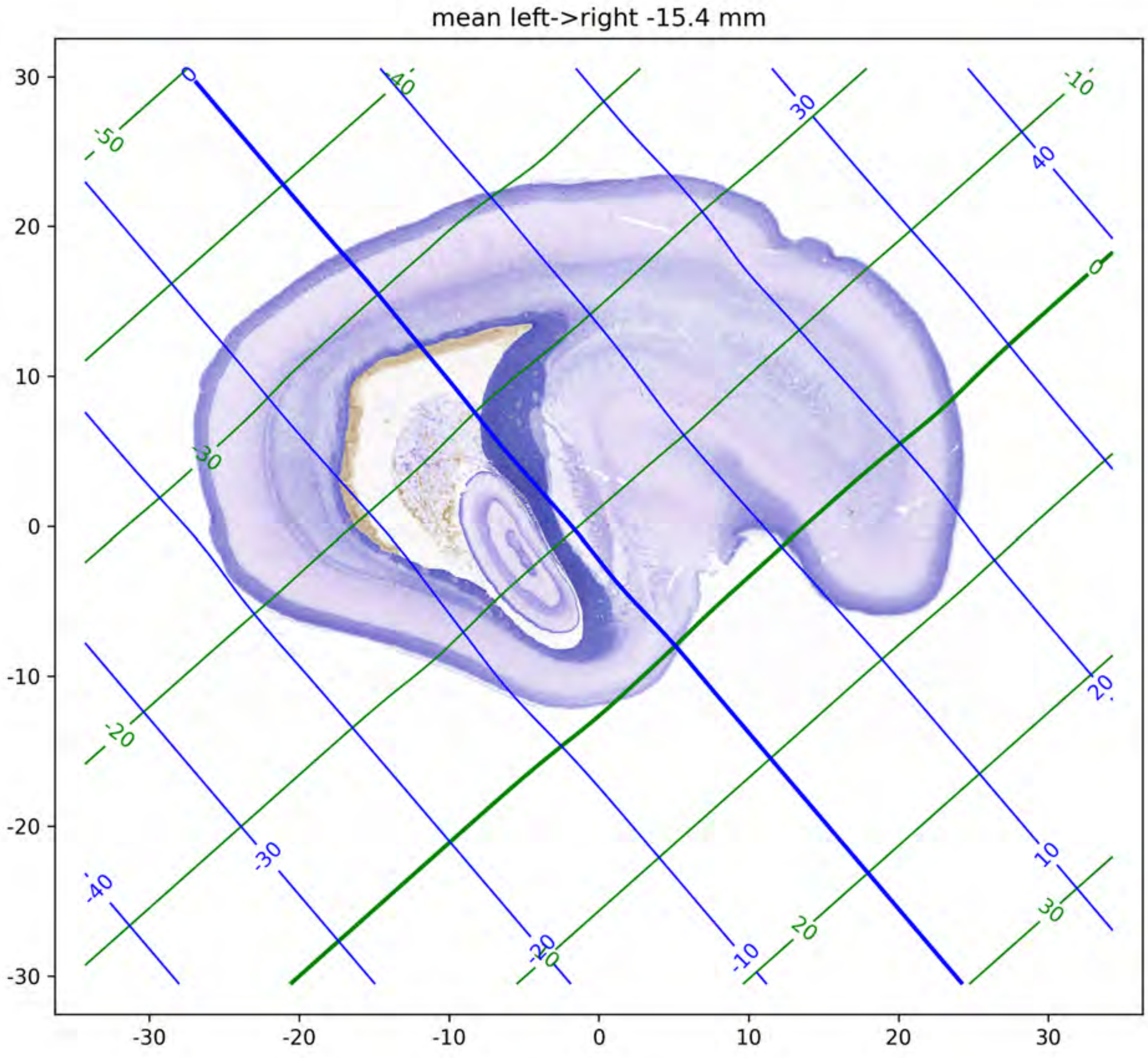

mean left->right -15.4 mm
30
20
10
0
-10
-20
-30
-30
-20
-10
0
10
20
30

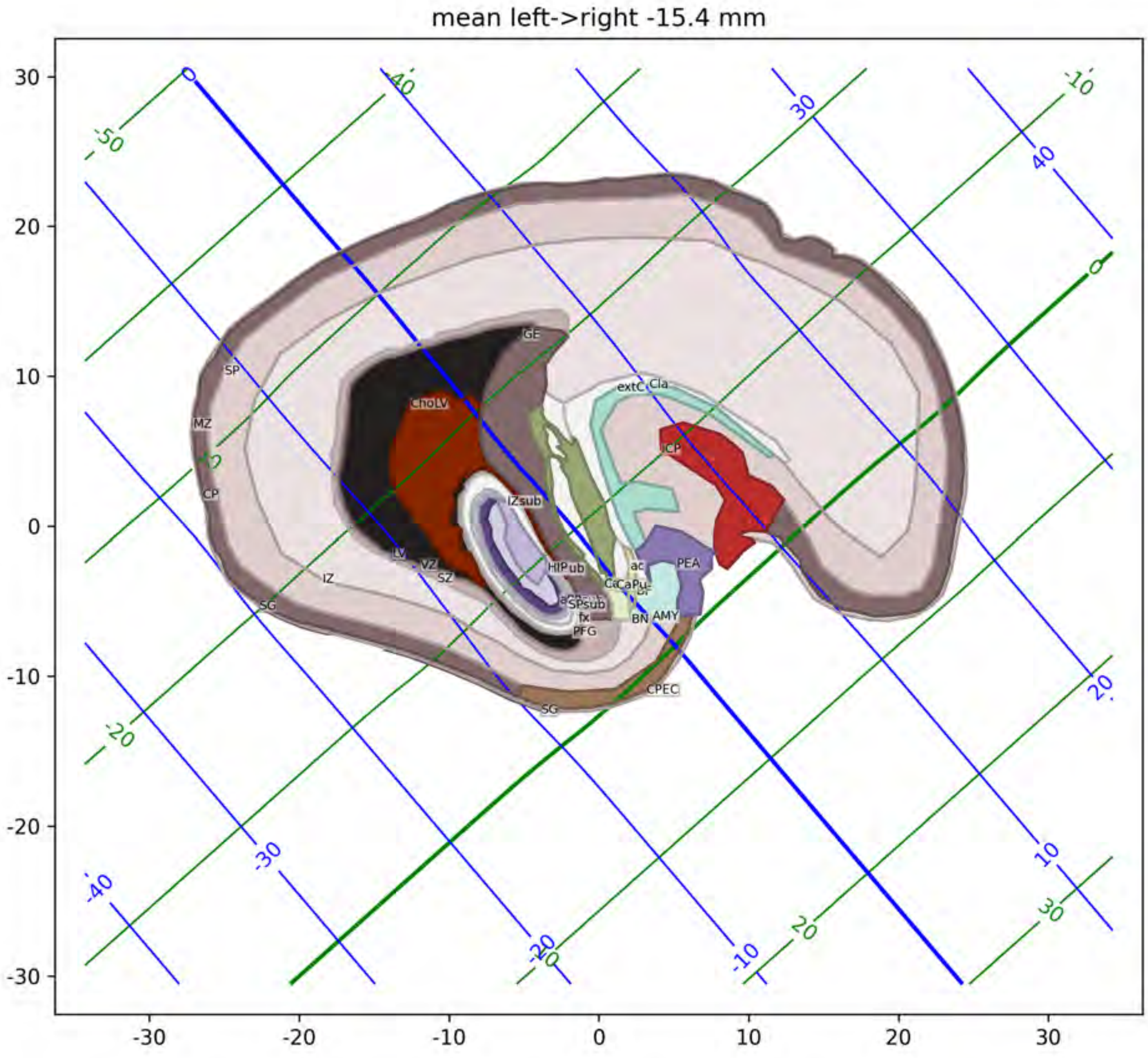
mean left->right -15.4 mm
GE
SP
MZ
CP
ChoLV
IZsub
LV
VZ
SZ
IZ
SG
extC
Cla
ICP
HIP
PEA
ac
BN
AMY
fx
PFG
CPEC

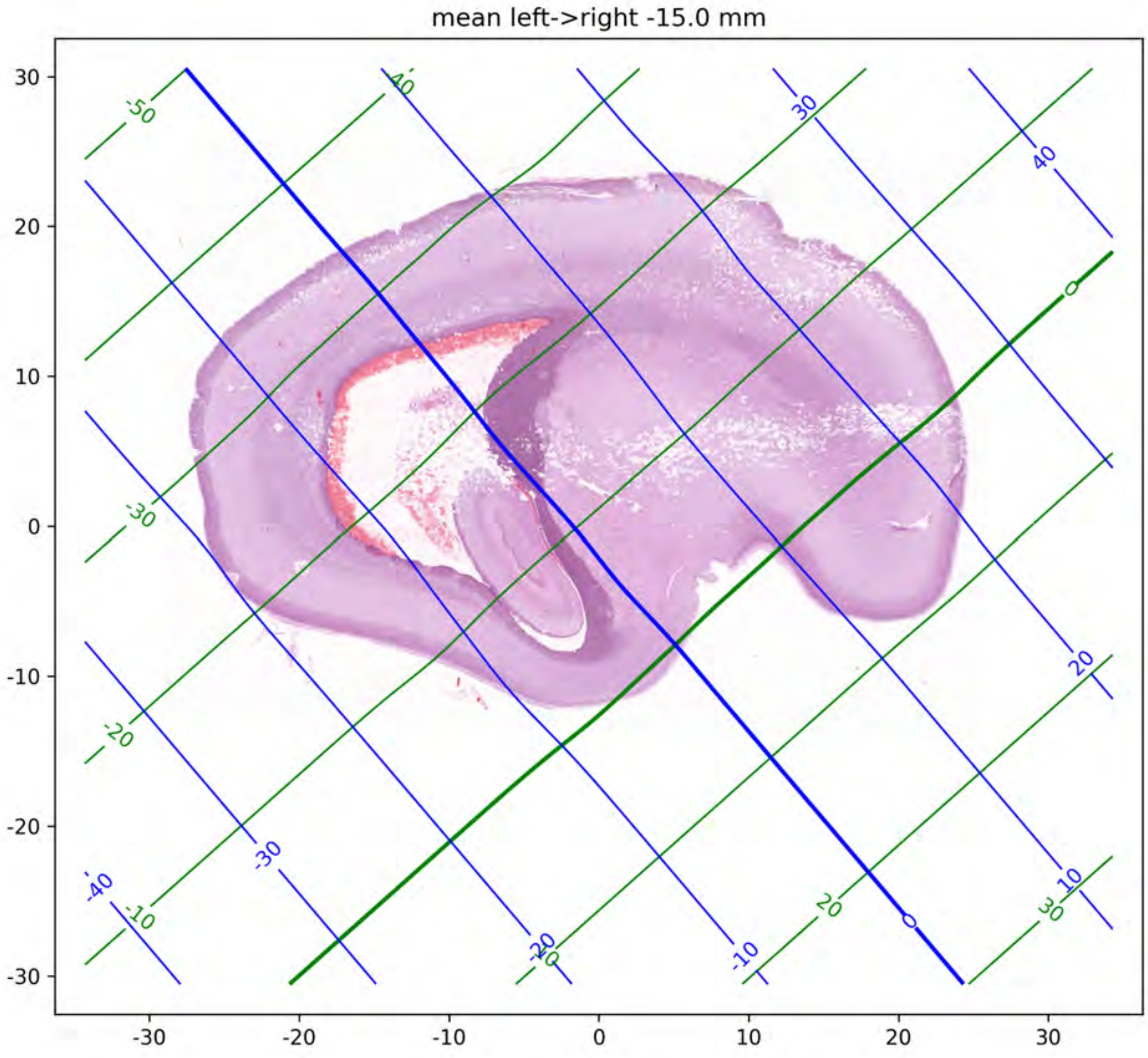

mean left->right -15.0 mm
30
20
10
0
-10
-20
-30
-30
-20
-10
0
10
20
30

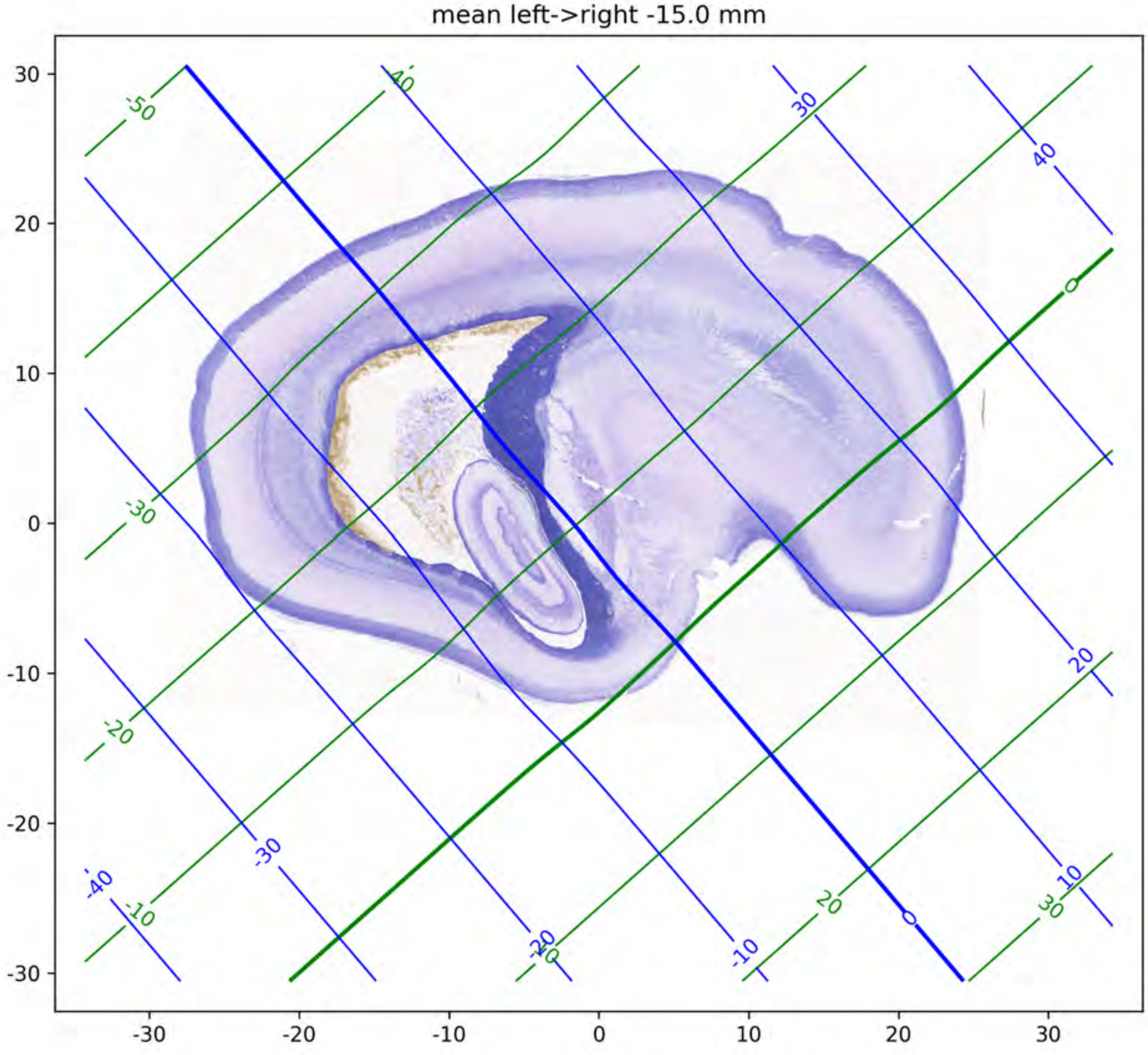
mean left->right -15.0 mm
30
20
10
0
-10
-20
-30
-30
-20
-10
0
10
20
30
-50
-40
-30
-20
-10
0
10
20
30
40
30
-30
-20
-10
0
10
20

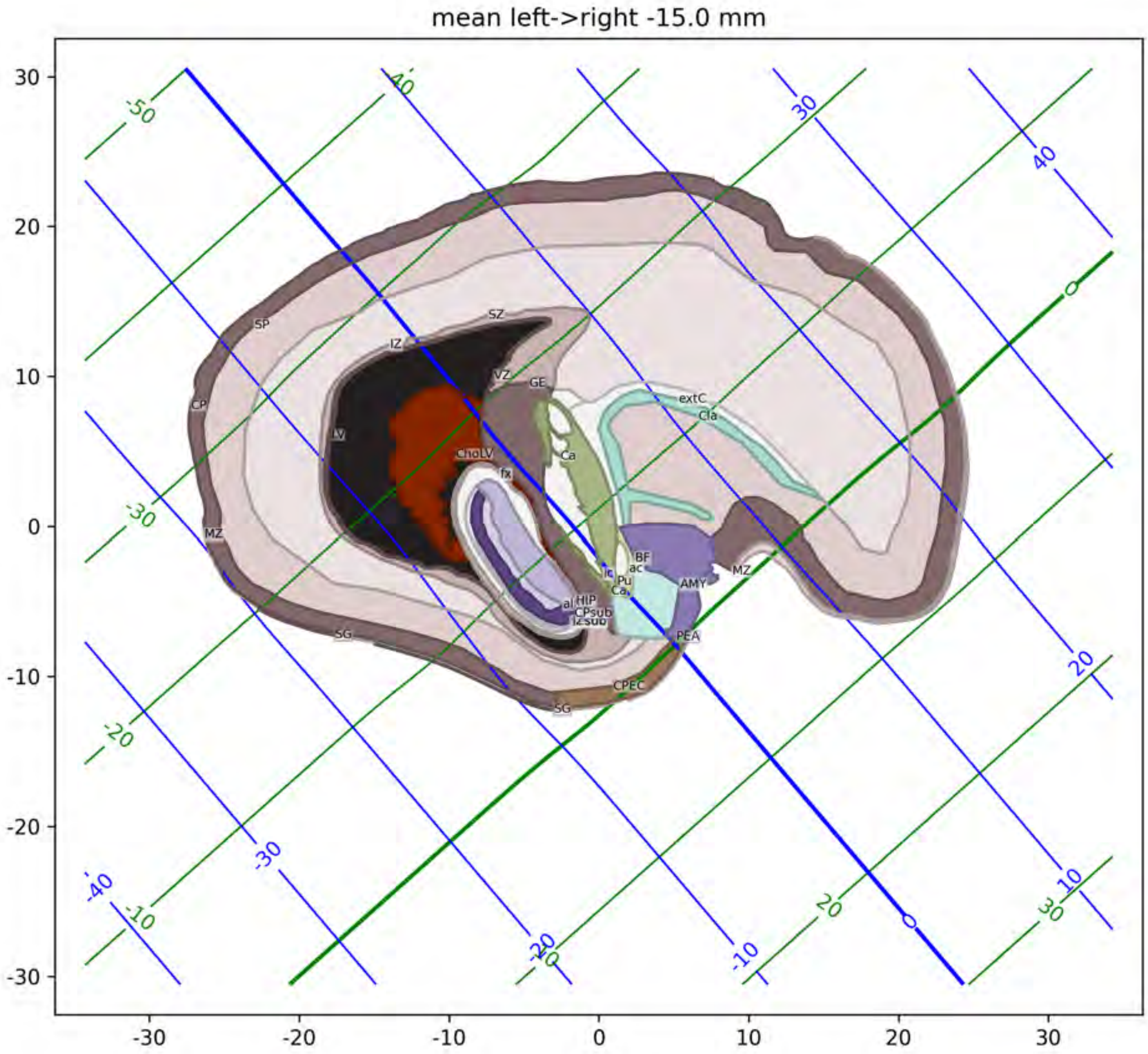
mean left->right -15.0 mm
SZ
SP
IZ
VZ
GE
CP
extC
Cla
LV
CholV
fx
Ca
MZ
BF
ac
Pu
AMY
HIP
CPsub
PEA
SG
CPEC
-30
-20
-10
0
10
20
30

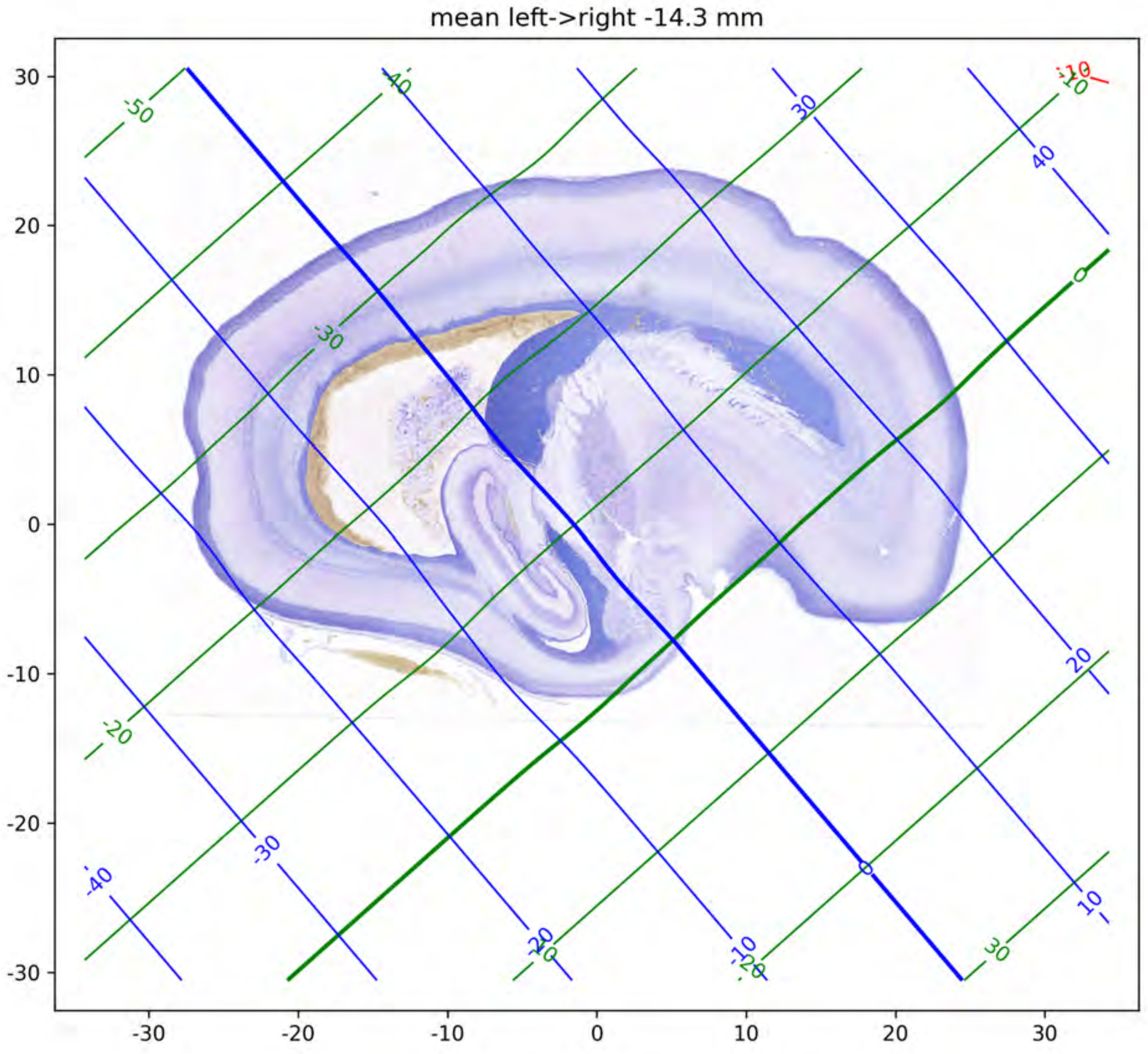

mean left->right -14.3 mm

mean left->right -14.3 mm

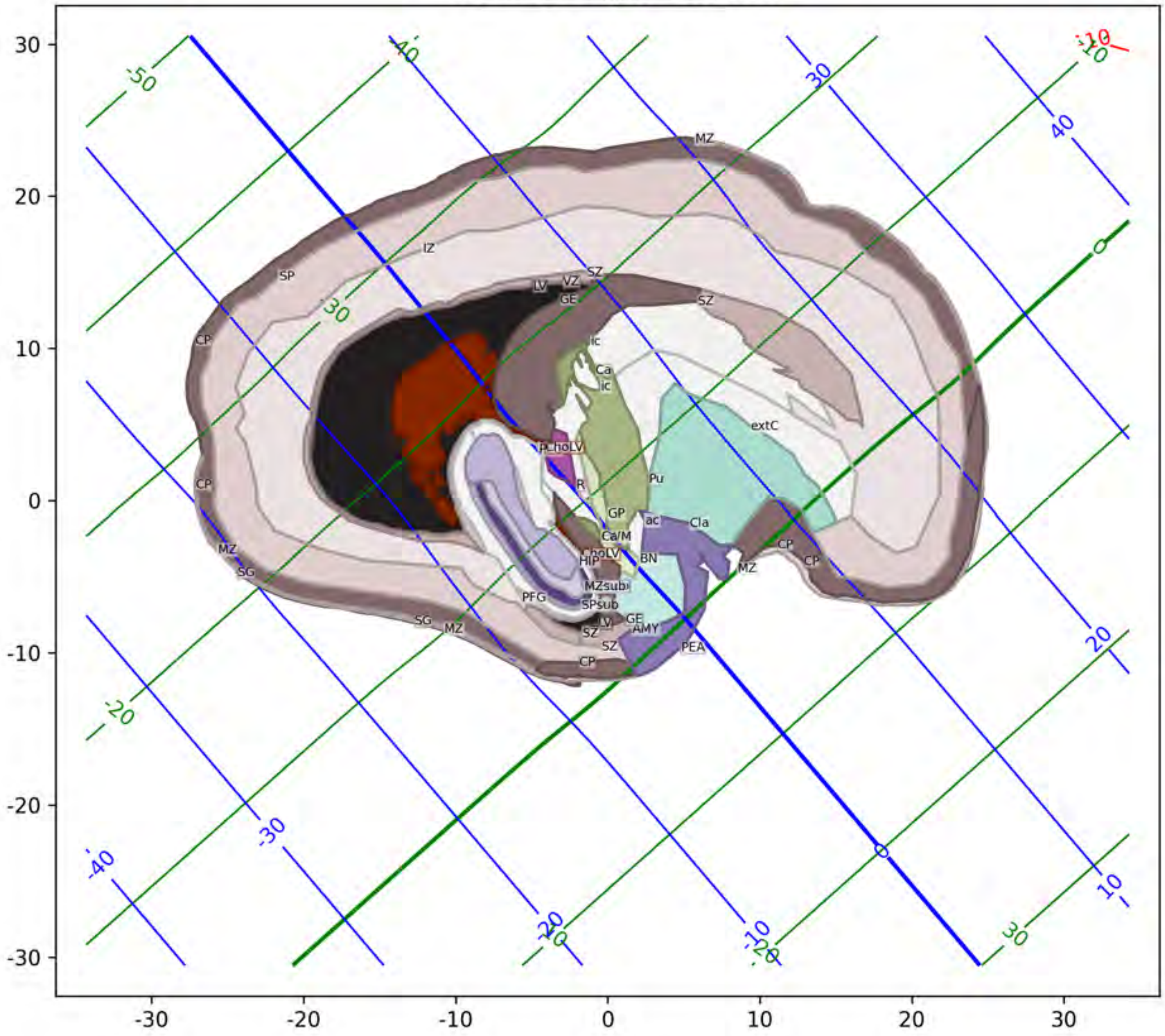

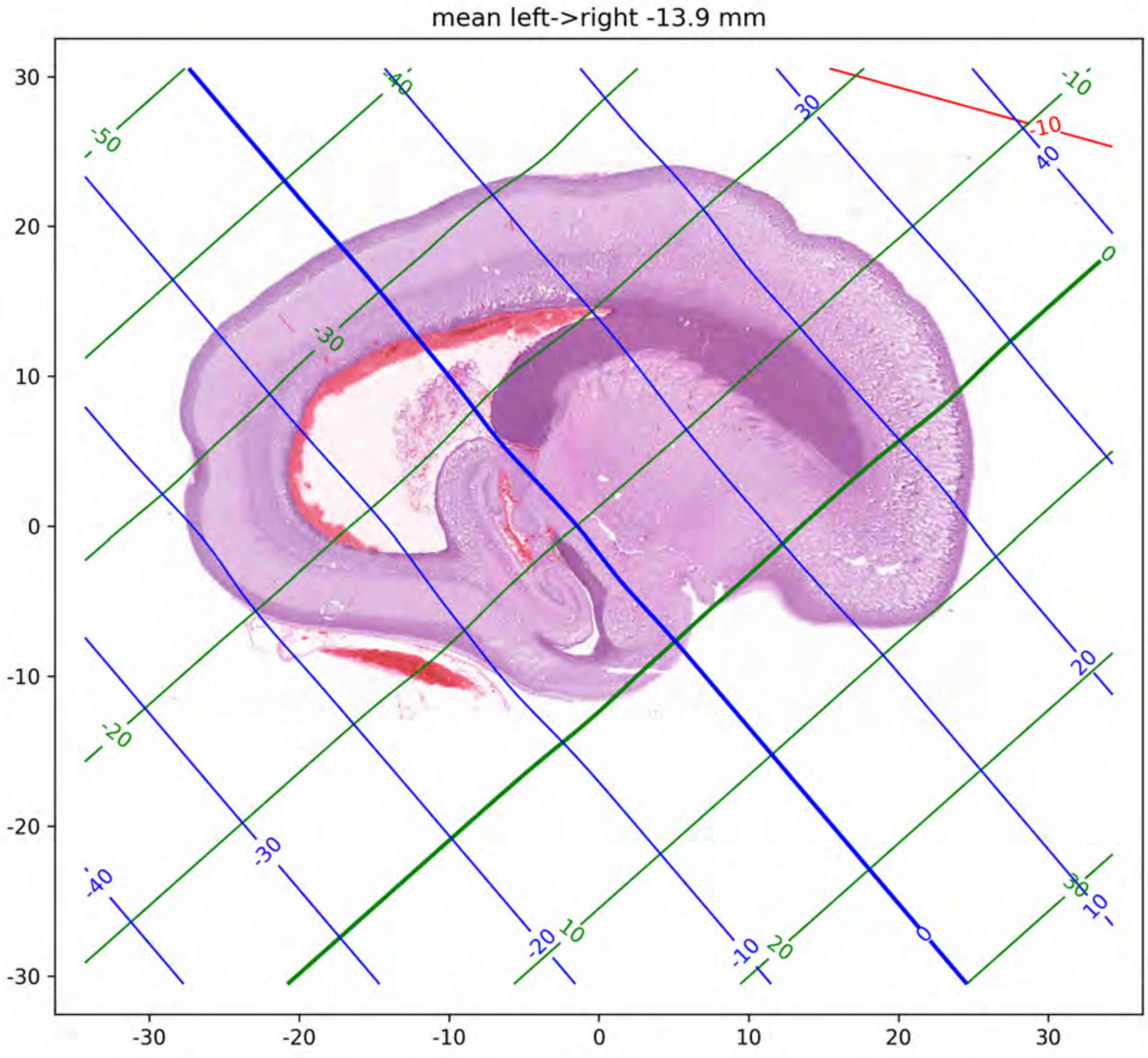
mean left->right -13.9 mm
30
20
10
0
-10
-20
-30
-30
-20
-10
0
10
20
30

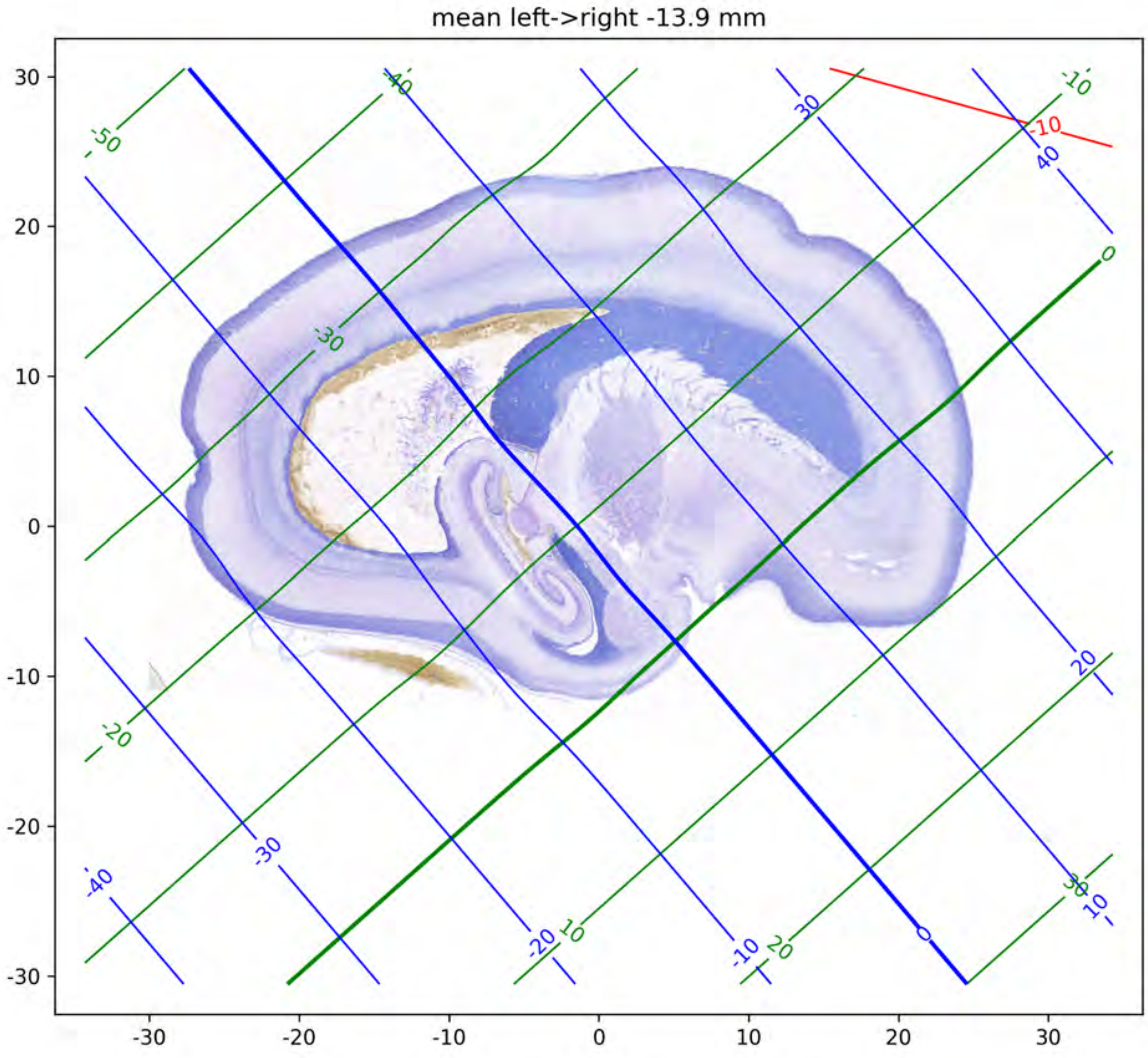

mean left->right -13.9 mm
30
20
10
0
-10
-20
-30
-30
-20
-10
0
10
20
30
-50
-40
-30
-20
-10
0
10
20
30
40
-10

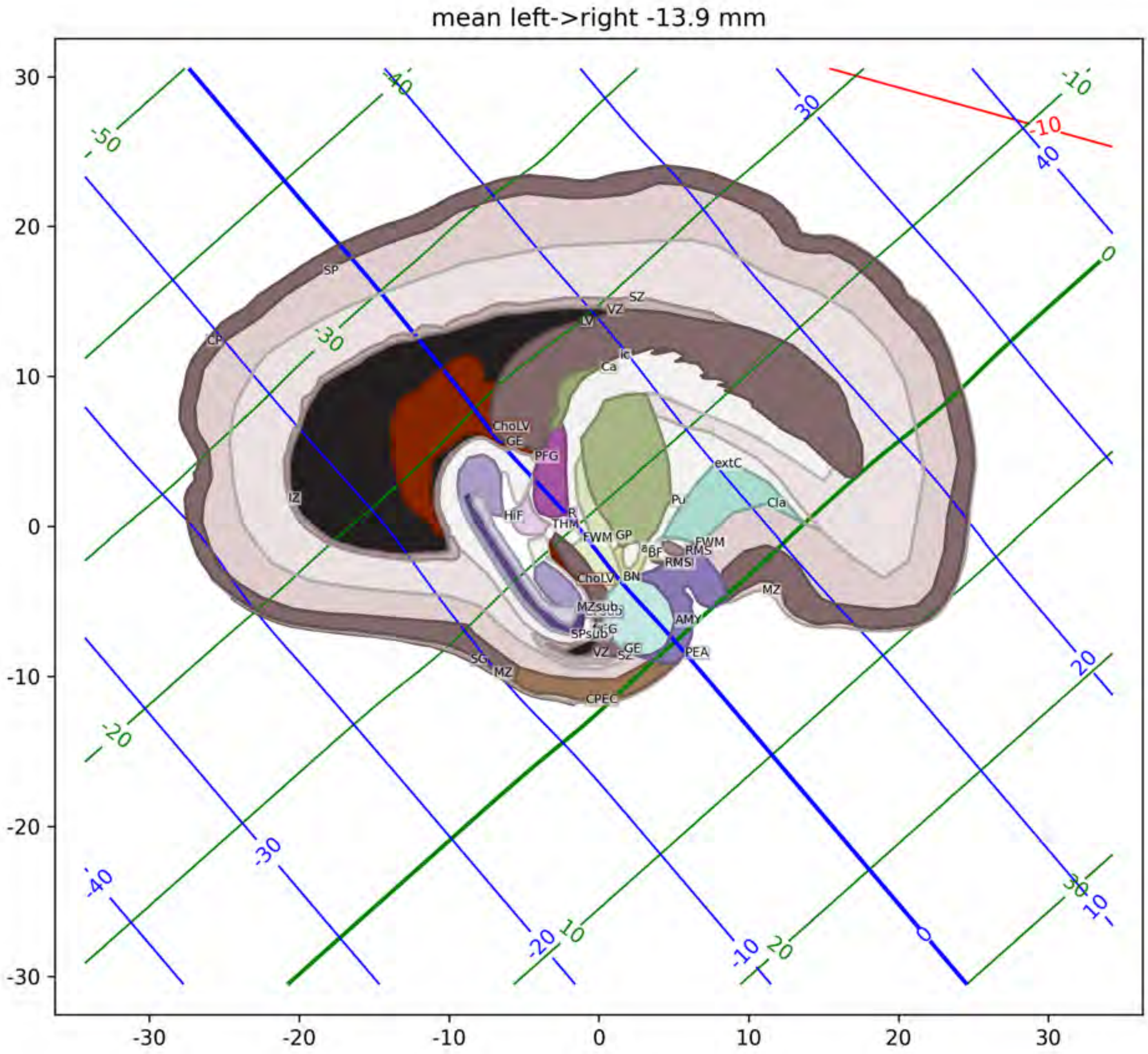
mean left->right -13.9 mm
SP
CP
IZ
SZ
VZ
LV
Ic
Ca
CholV
GE
PFG
HiF
R
THM
FWM
GP
Pu
extC
Cla
BF
RMS
BN
MZsub
SPsub
AMY
PEA
VZ
SZ
SG
MZ
CPEC

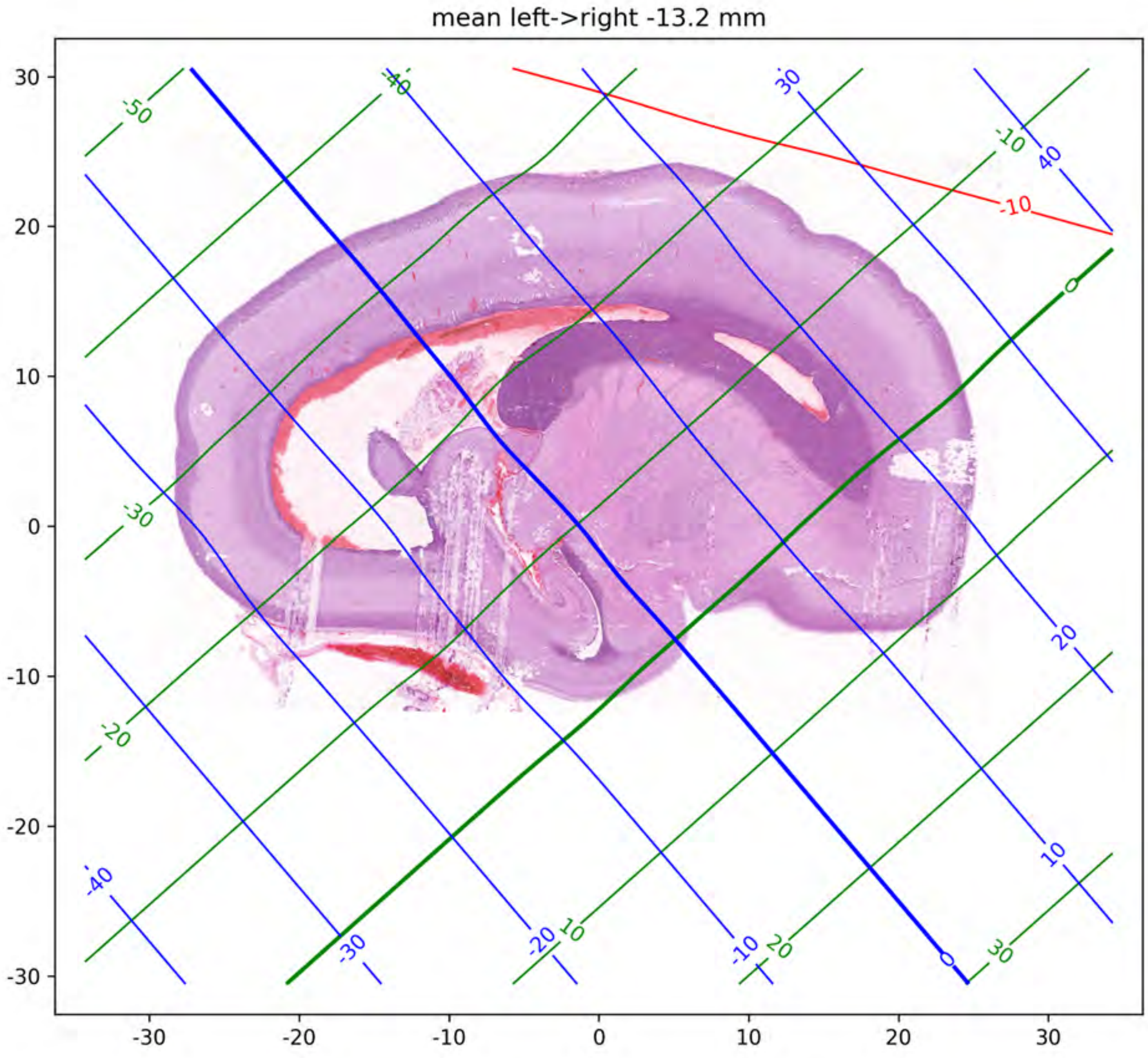
mean left->right -13.2 mm
30
20
10
0
-10
-20
-30
-30
-20
-10
0
10
20
30

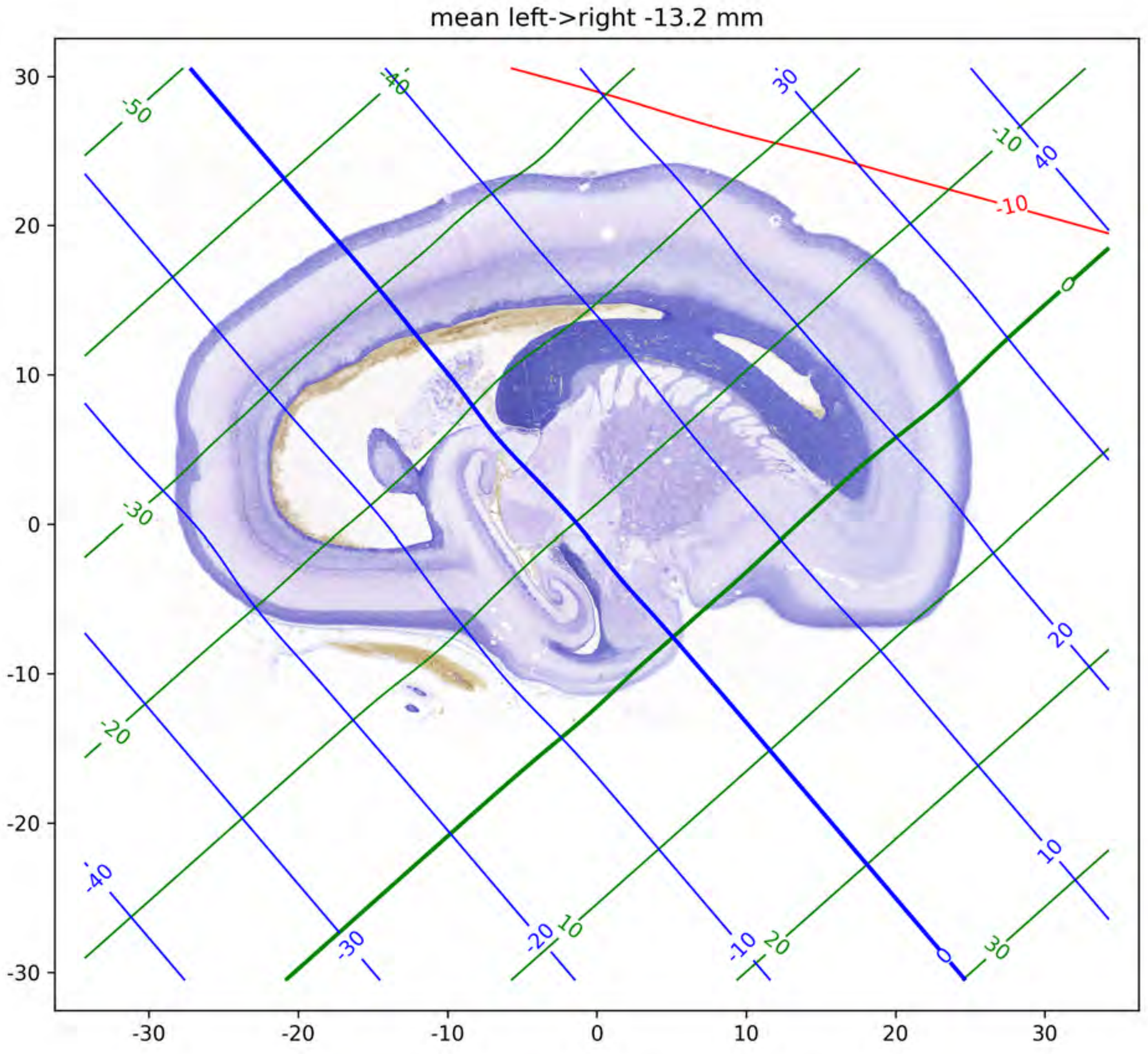
mean left->right -13.2 mm
-50
-40
30
40
-10
-10
0
-30
-20
-40
-30
-20
10
-10
20
0
30
10
20
30
20
10
0
-10
-20
-30
-30
-20
-10
0
10
20
30

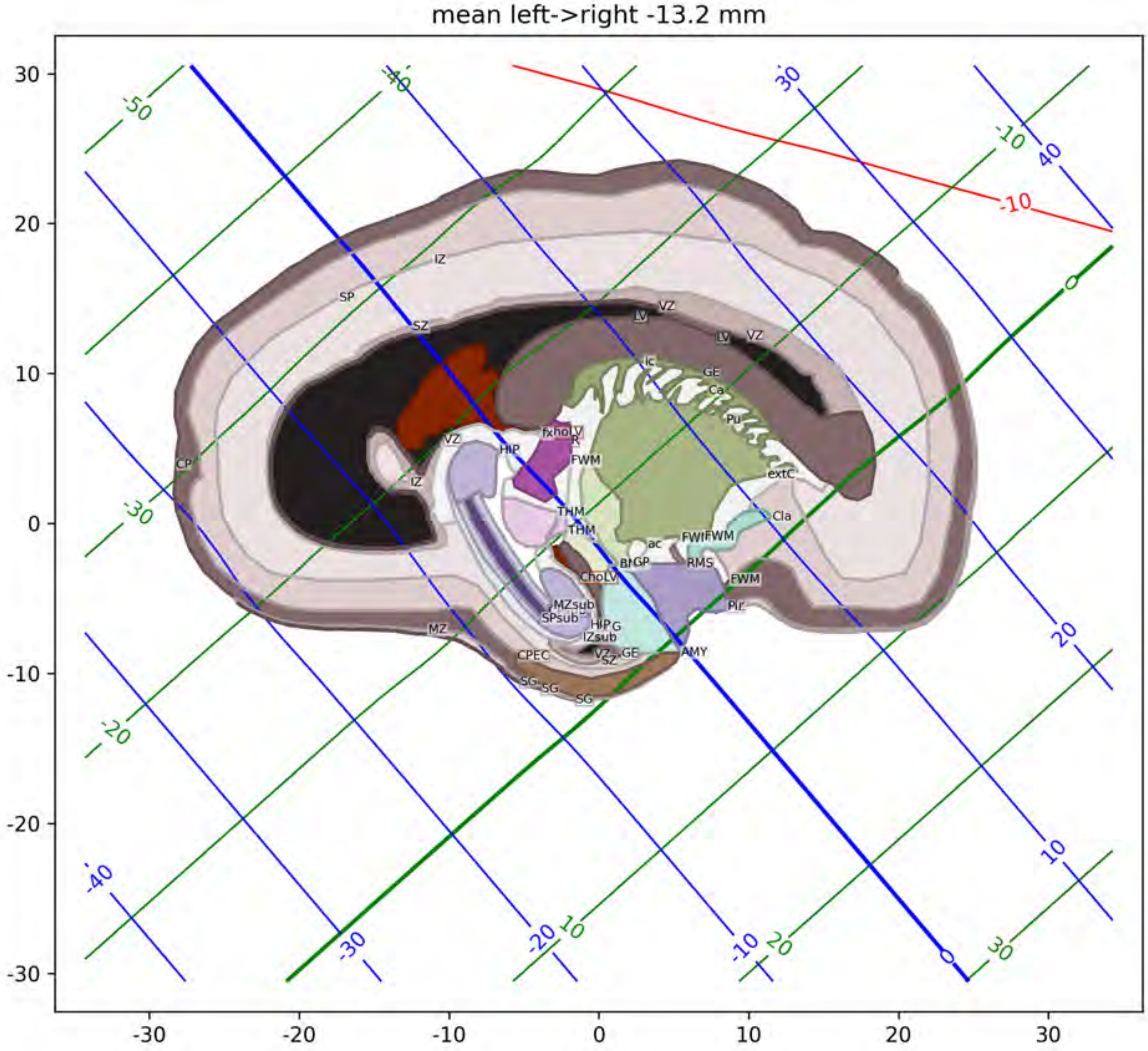
mean left->right -13.2 mm
30
20
10
0
-10
-20
-30
-30
-20
-10
0
10
20
30
-50
-40
-30
-20
-10
0
10
20
30
40
-10
IZ
SP
SZ
VZ
LV
Ic
GE
Ca
Pu
extC
Cla
CP
HIP
fx
FWM
THM
ac
GP
RMS
Pir
ChoLV
MZsub
SPsub
IZsub
MZ
CPEC
SG
AMY

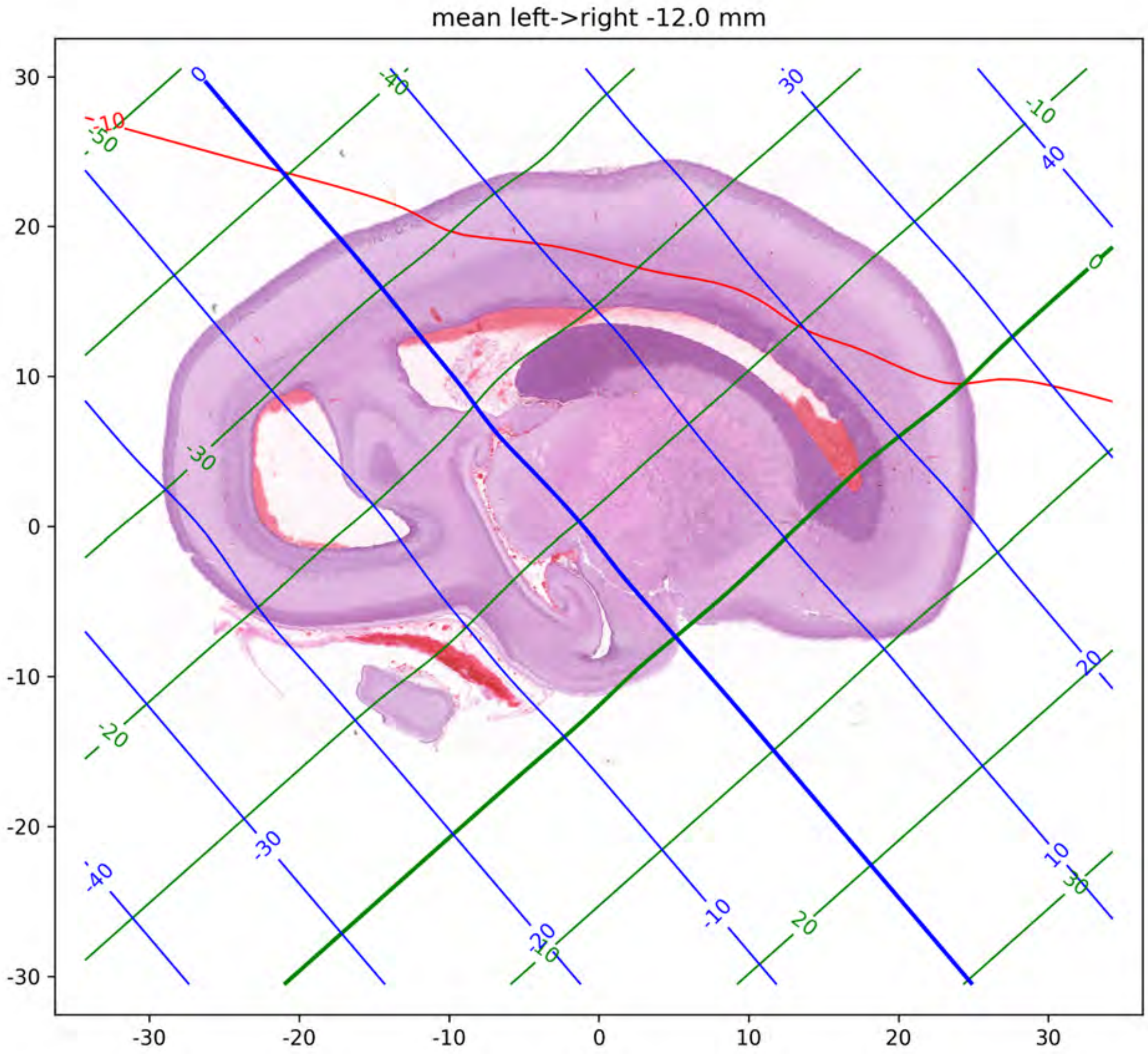
mean left->right -12.0 mm
30
20
10
0
-10
-20
-30
-30
-20
-10
0
10
20
30

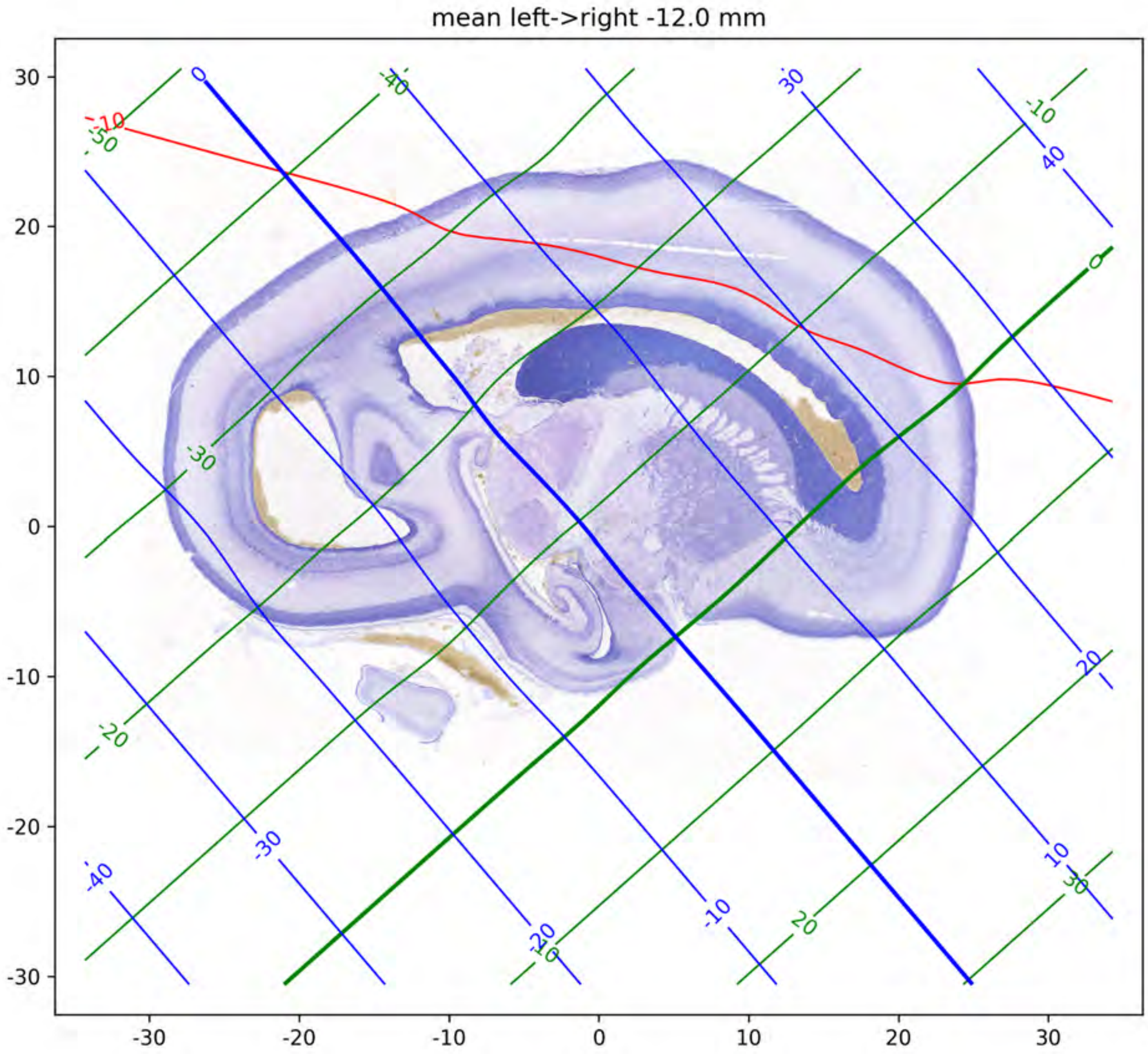

mean left->right -12.0 mm
30
20
10
0
-10
-20
-30
-30
-20
-10
0
10
20
30

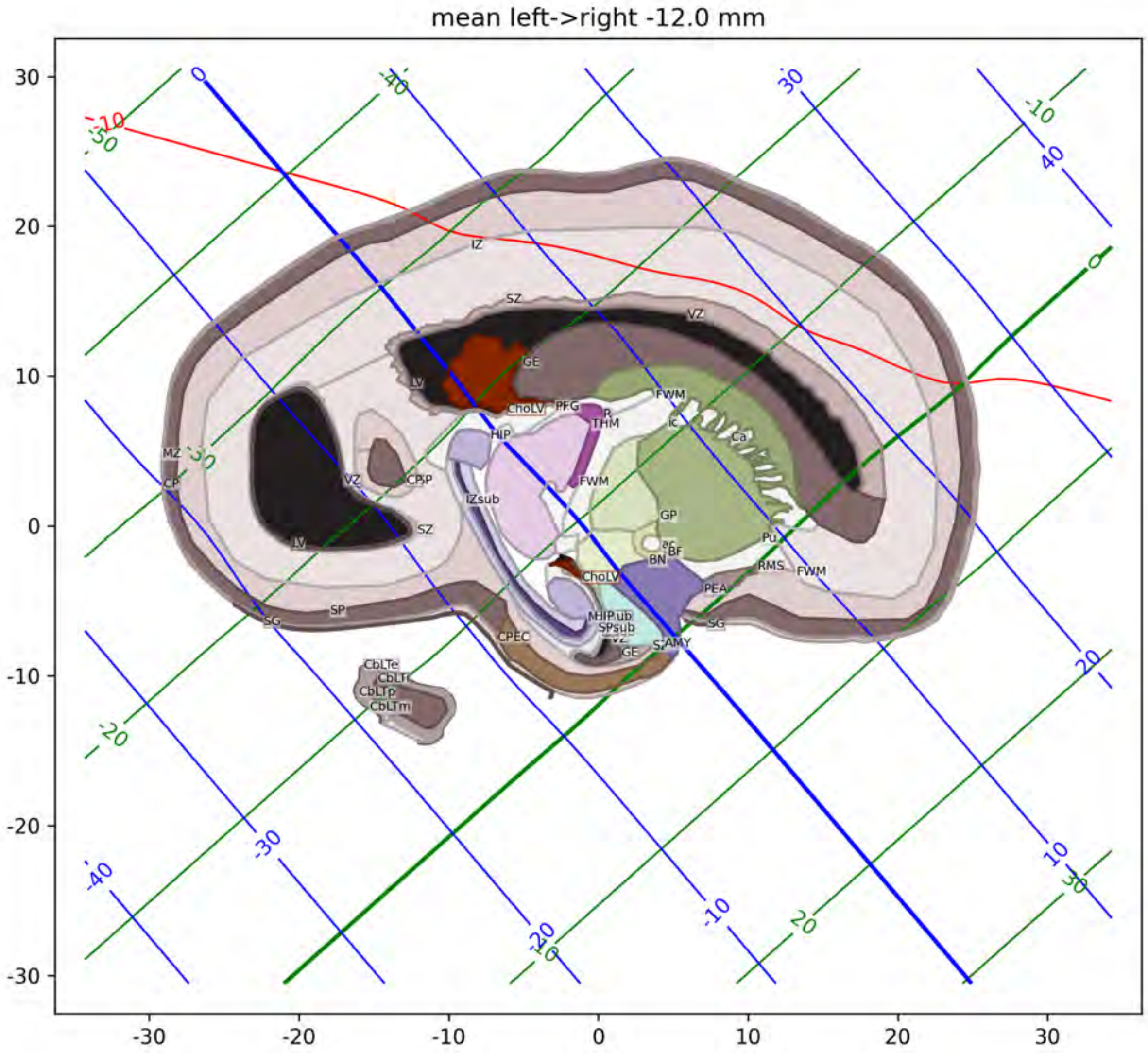

mean left->right -12.0 mm
IZ
SZ
VZ
GE
LV
FWM
ChoLV
PFG
R
THM
ic
Ca
HIP
MZ
CP
VZ
CP
SP
IZsub
FWM
GP
SZ
LV
ec
BF
BN
Pu
RMS
FWM
ChoLV
PEA
SP
SG
MHIPsub
SPsub
SG
CPEC
VZ
AMY
GE
CbLTe
CbLTi
CbLTp
CbLTm
30
20
10
0
-10
-20
-30
-30
-20
-10
0
10
20
30

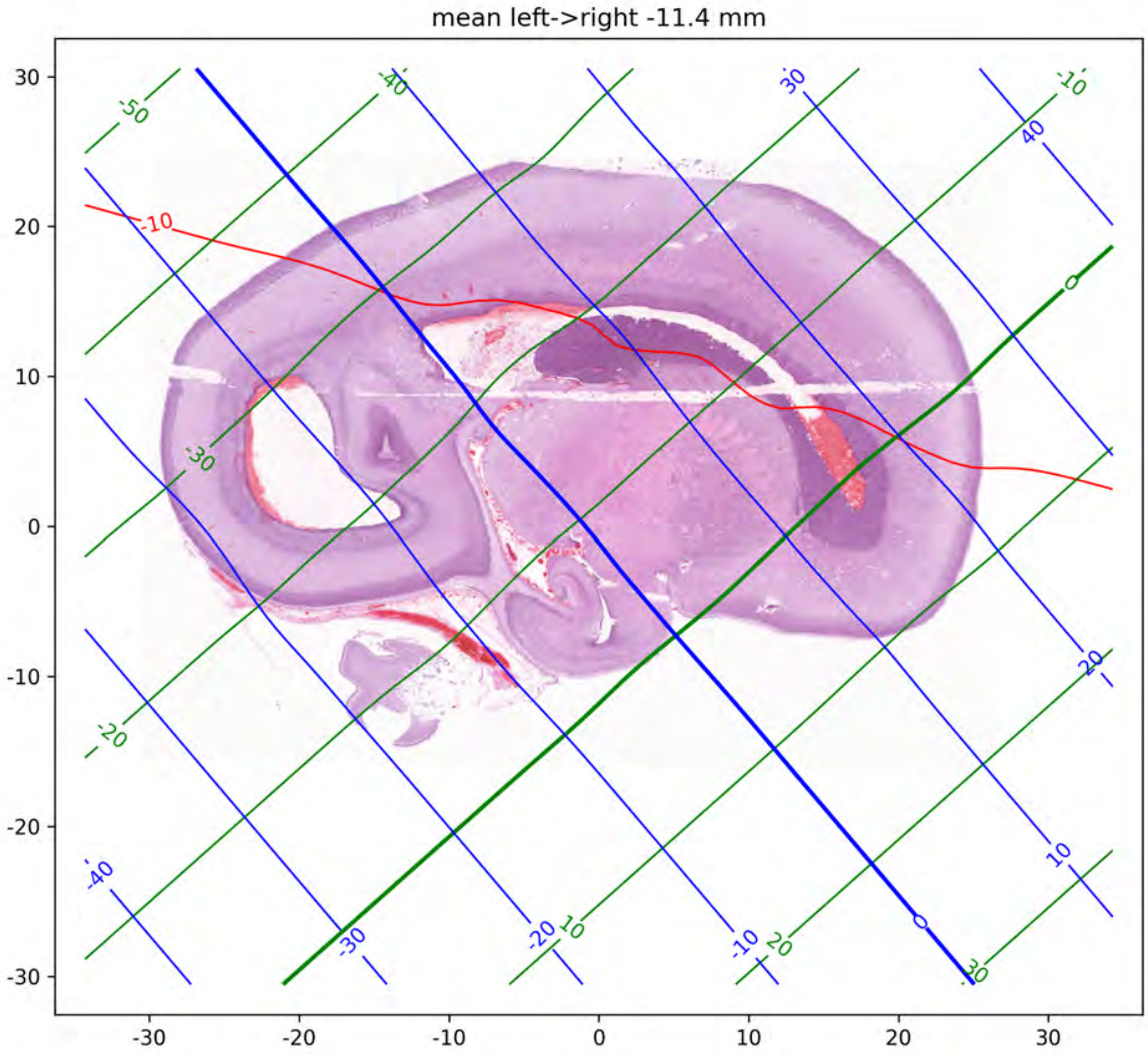
mean left->right -11.4 mm
30
20
10
0
-10
-20
-30
-30
-20
-10
0
10
20
30

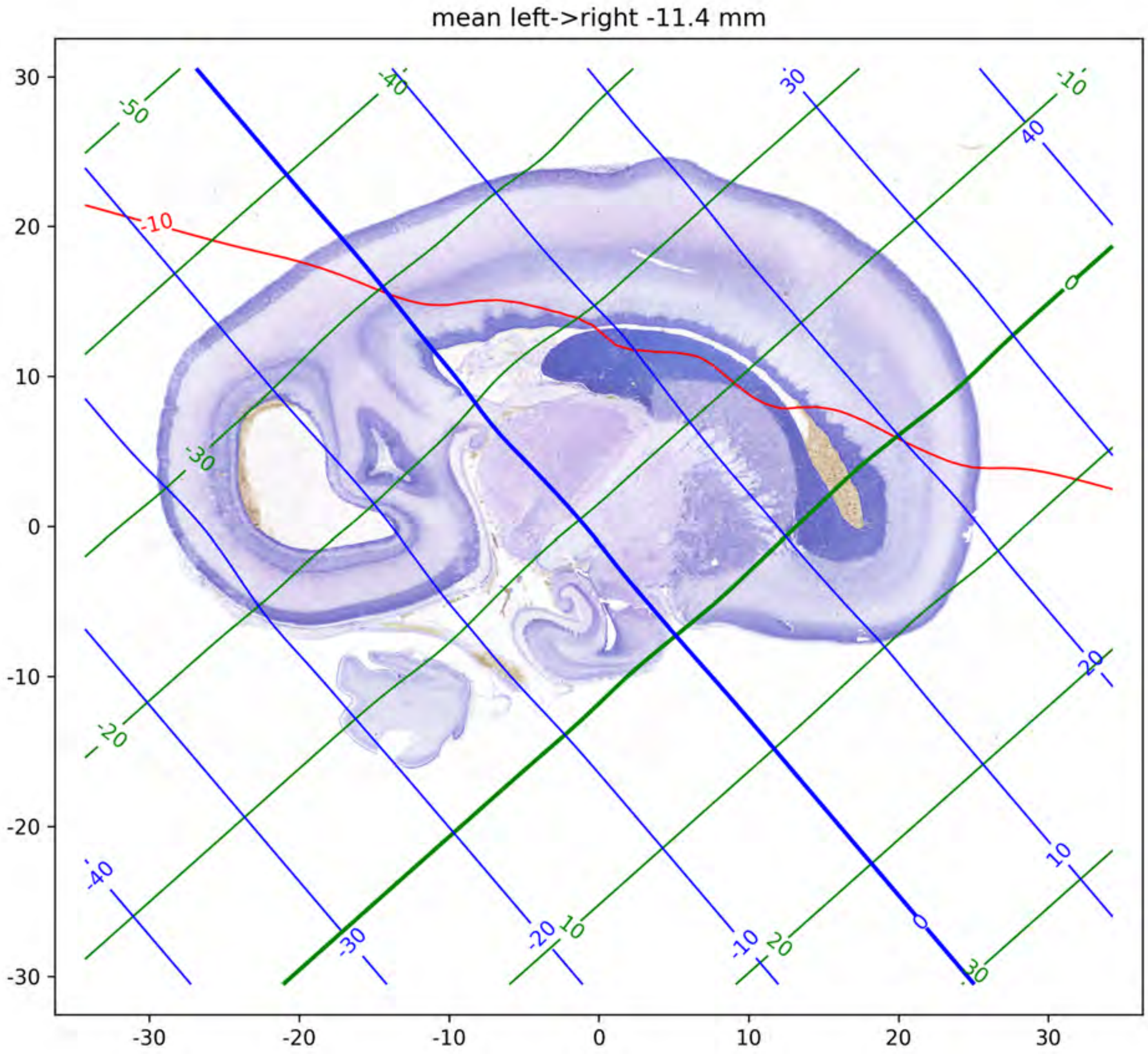

mean left->right -11.4 mm
30
20
10
0
-10
-20
-30
-30
-20
-10
0
10
20
30
-50
-40
-30
-20
-10
0
10
20
30
40
-10

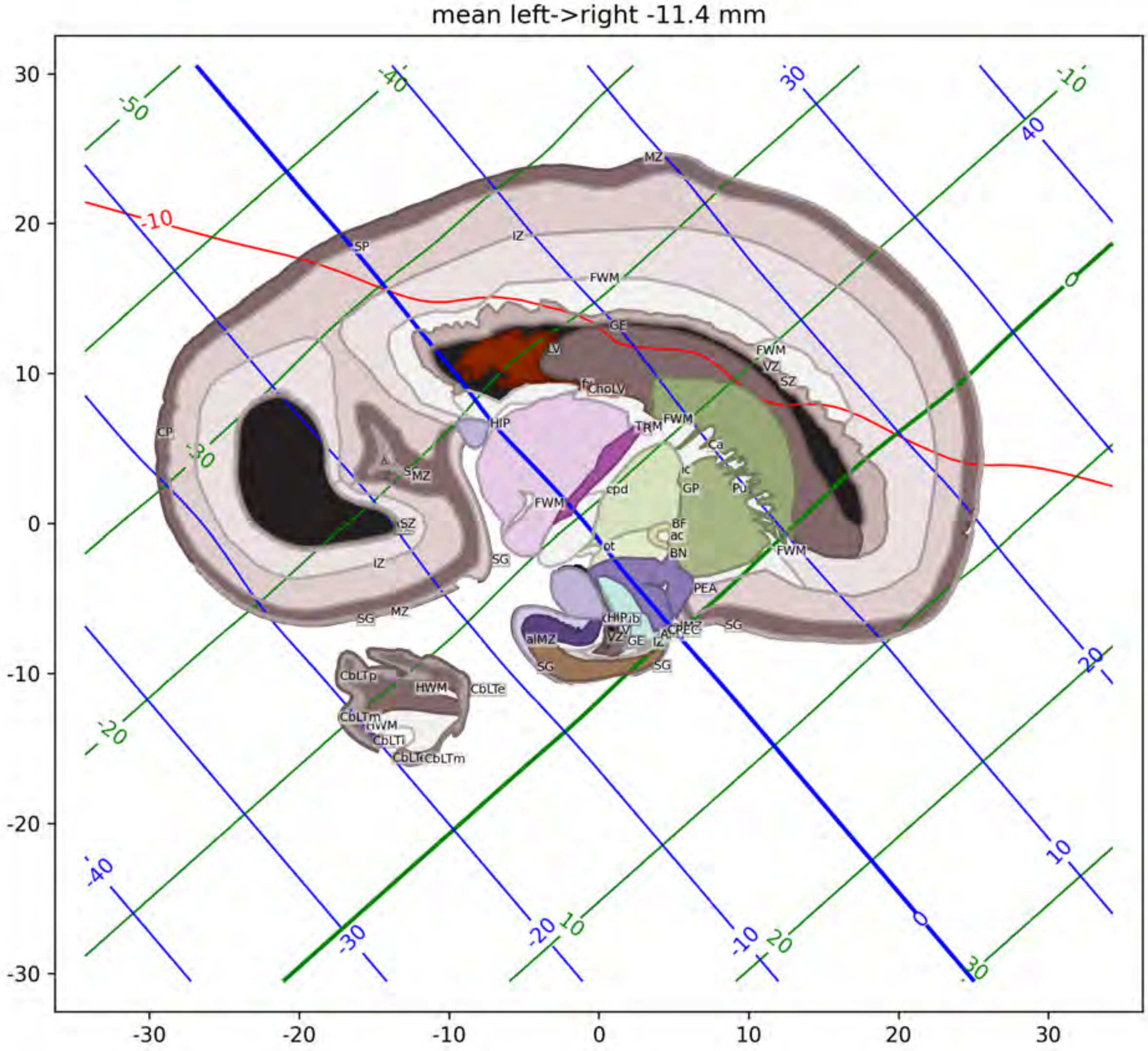
mean left->right -11.4 mm
MZ
IZ
SP
FWM
GE
LV
fr
CholV
HIP
TRM
FWM
VZ
SZ
Ca
ic
GP
Pu
cpd
BF
ac
BN
ot
PEA
SG
aIMZ
CbLTp
HWM
CbLTe
CbLTm
CbLTi
CP
-10
30
20
10
0
-10
-20
-30

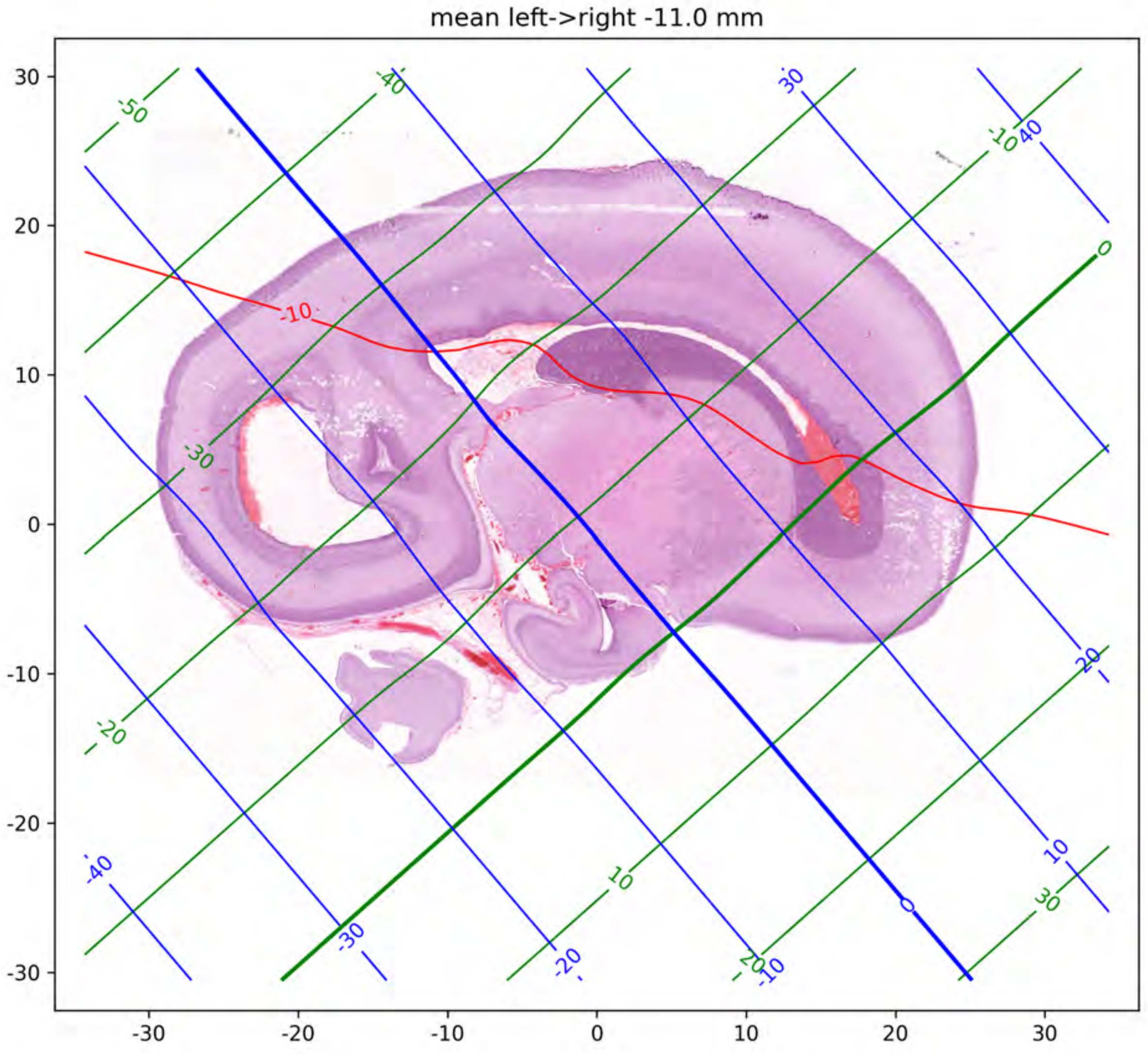
mean left->right -11.0 mm
30
20
10
0
-10
-20
-30
-30
-20
-10
0
10
20
30

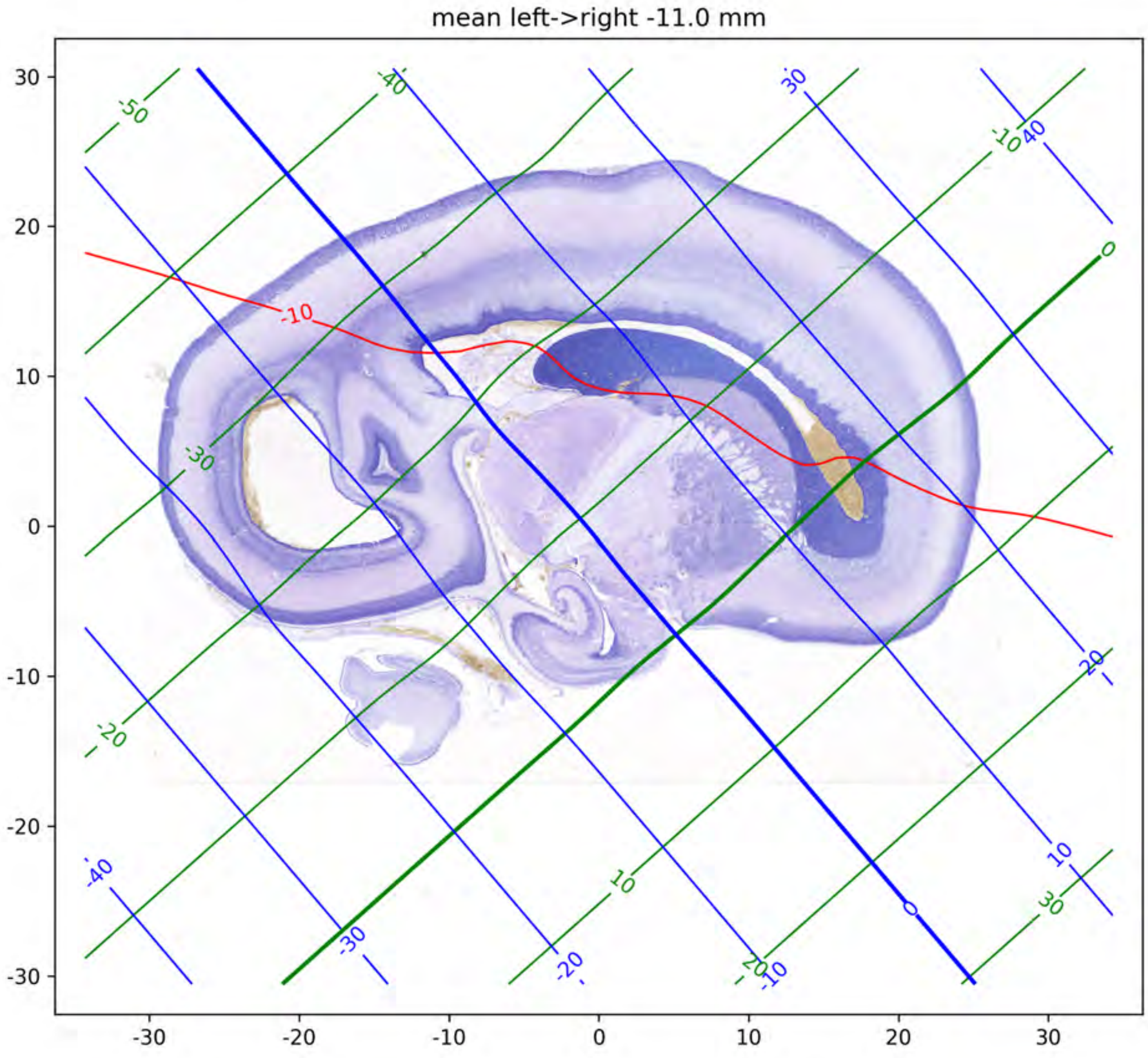
mean left->right -11.0 mm
30
20
10
0
-10
-20
-30
-30
-20
-10
0
10
20
30
-50
-40
30
-10
40
0
-10
-30
-20
20
-40
-30
-20
10
20
-10
0
10
30

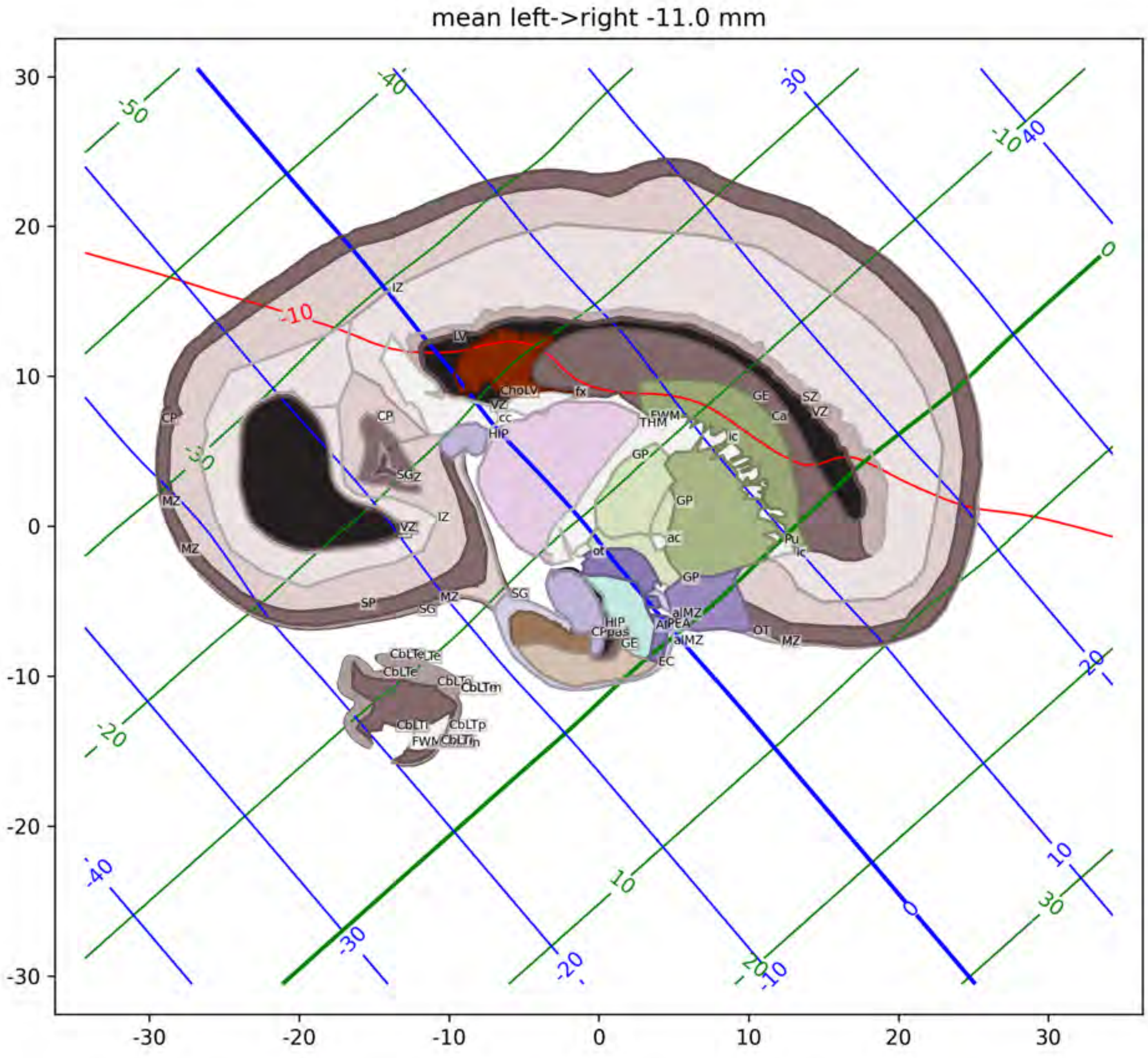

mean left->right -11.0 mm

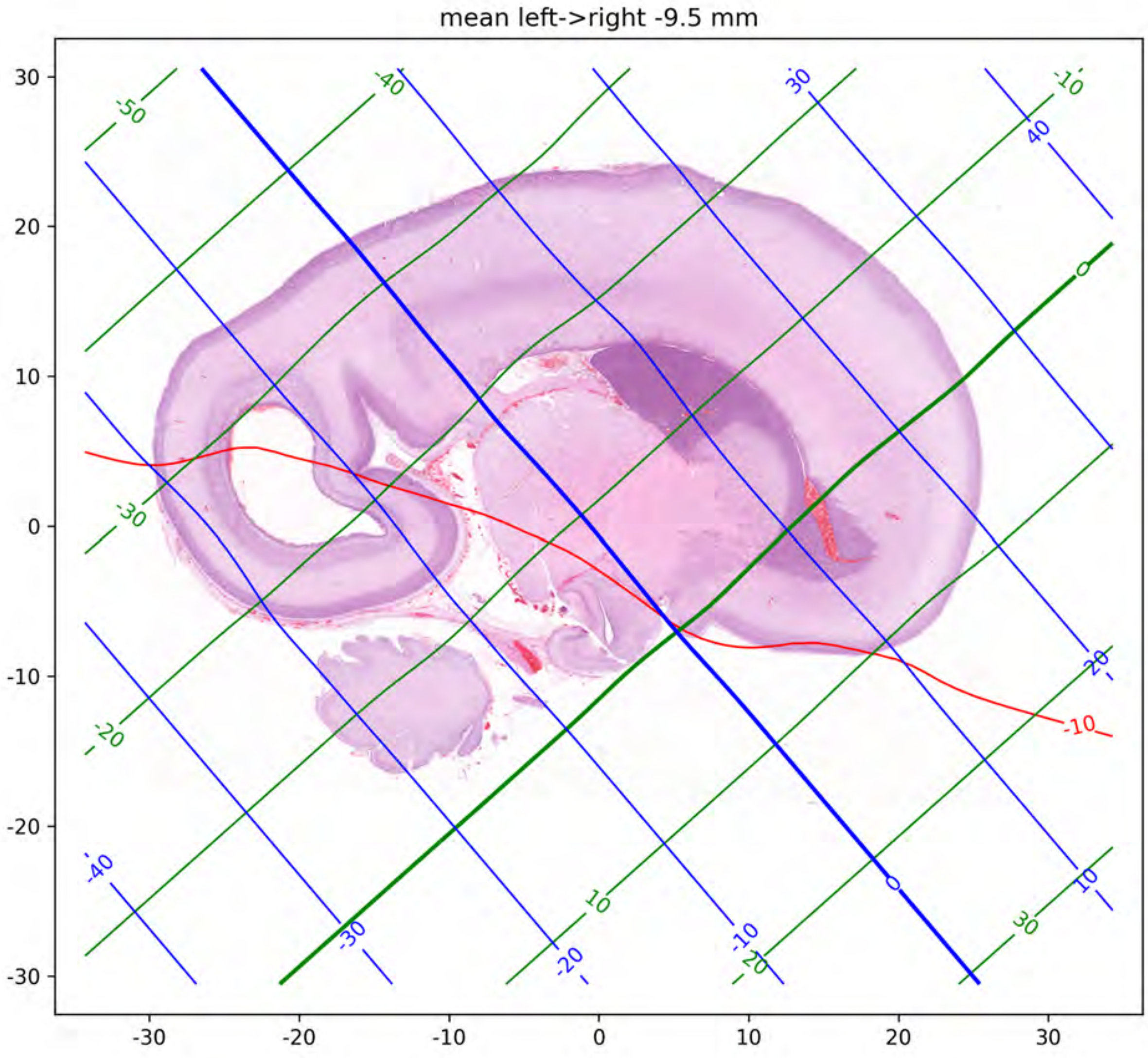
mean left->right -9.5 mm
30
20
10
0
-10
-20
-30
-30
-20
-10
0
10
20
30
-50
-40
-30
-20
-10
0
10
20
30
-40
-30
-20
-10
0
10
20
30
40
-10

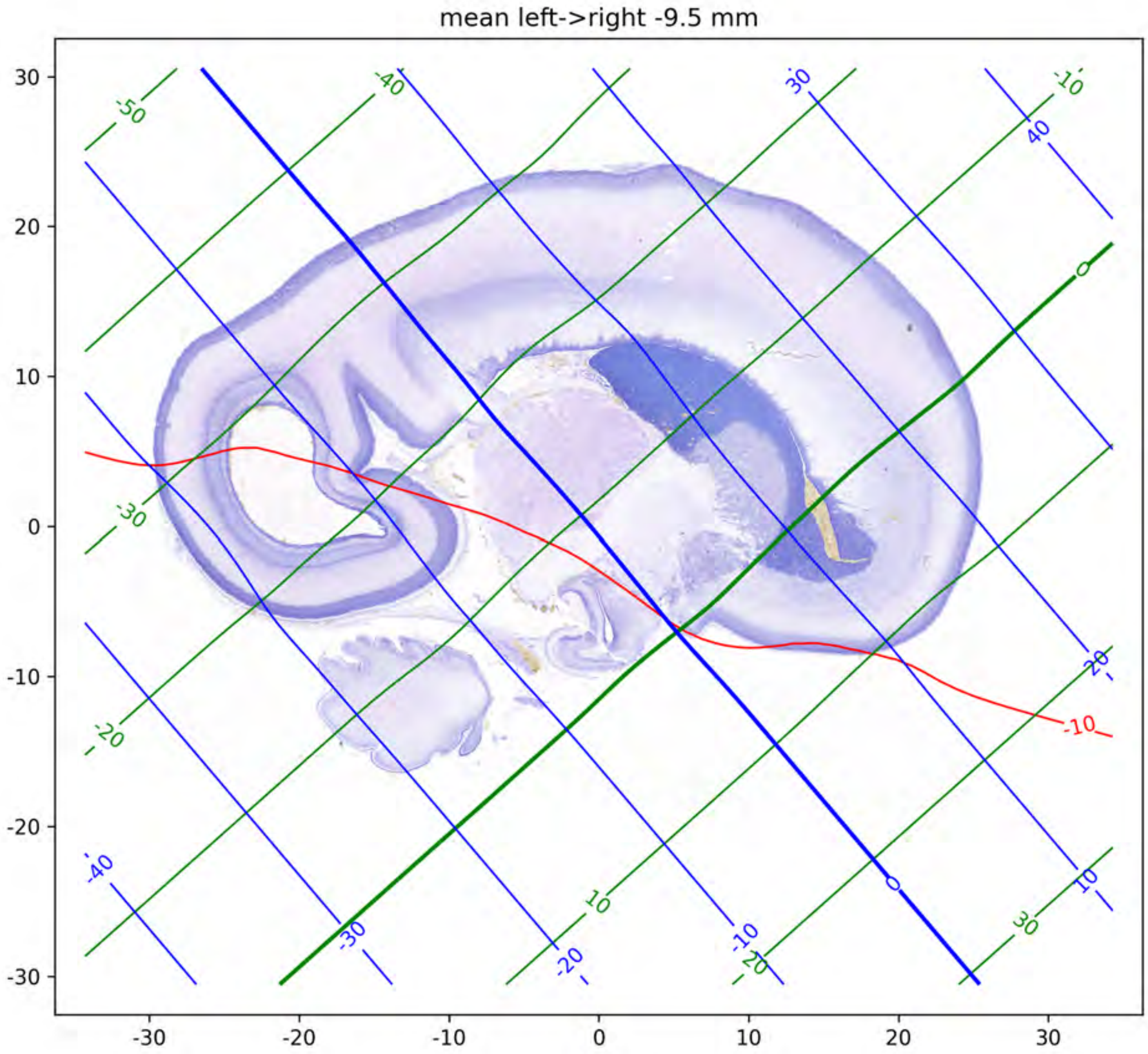
mean left->right -9.5 mm
30
20
10
0
-10
-20
-30
-30
-20
-10
0
10
20
30
-50
-40
-10
30
40
0
-30
-20
-10
-20
-40
-30
10
-20
-10
-20
0
10
30

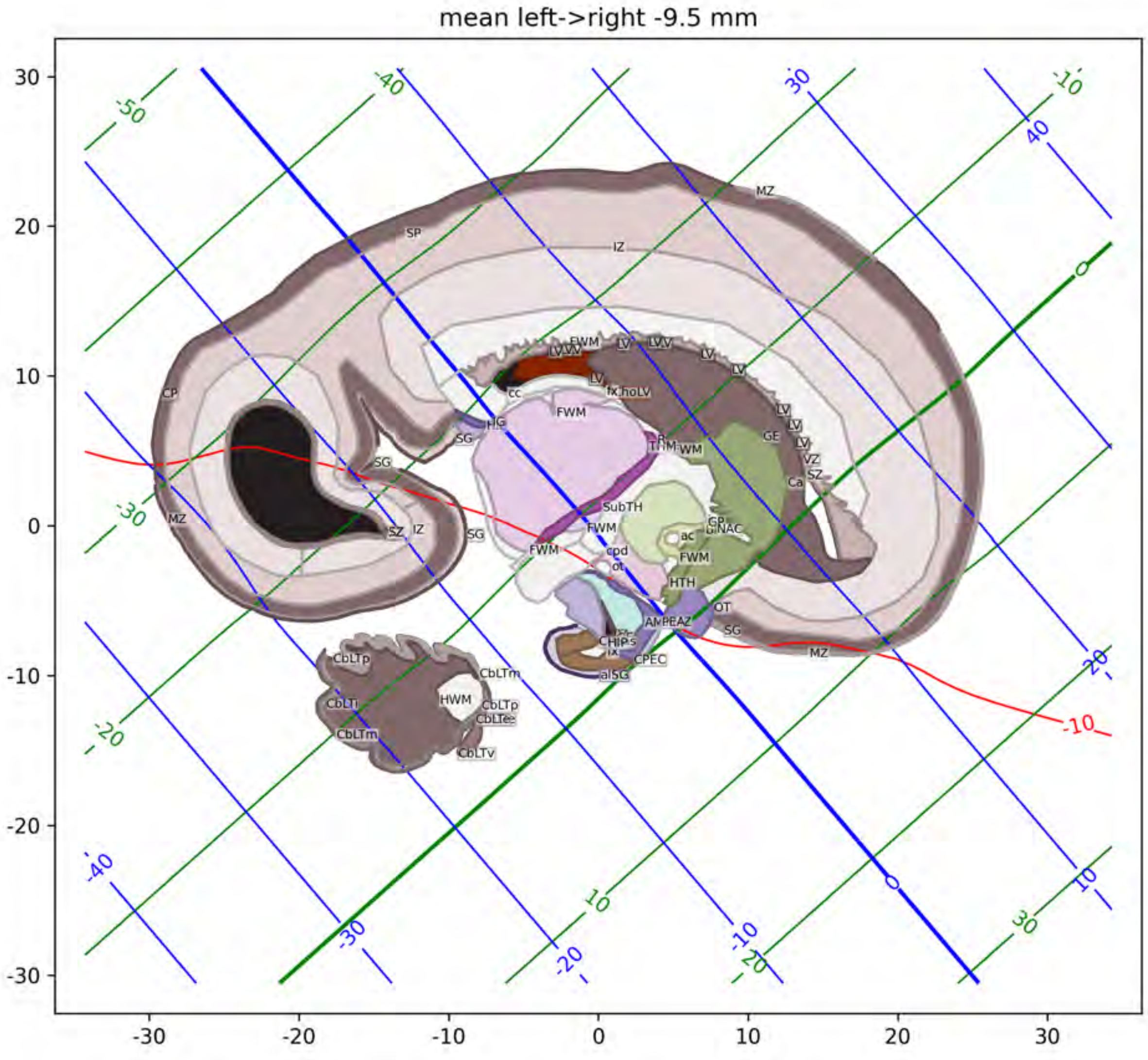
mean left->right -9.5 mm
MZ
SP
IZ
CP
SG
SZ
FWM
LV
cc
fx
GE
VZ
Ca
SubTH
ac
cpd
ot
HTH
OT
CPEC
HWM
CbLTp
CbLTm
CbLTi
CbLTe
CbLTv
-10

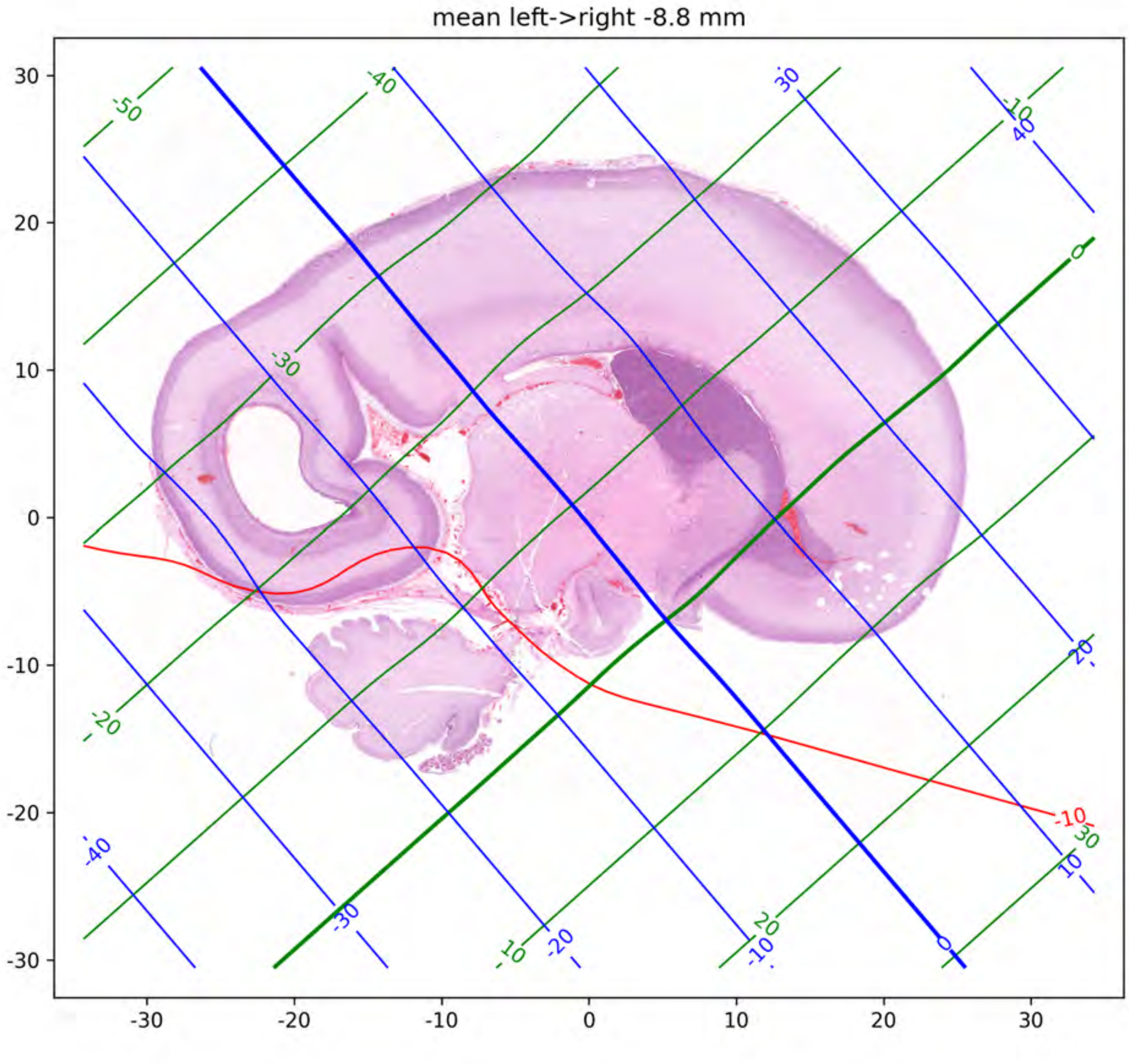
mean left->right -8.8 mm

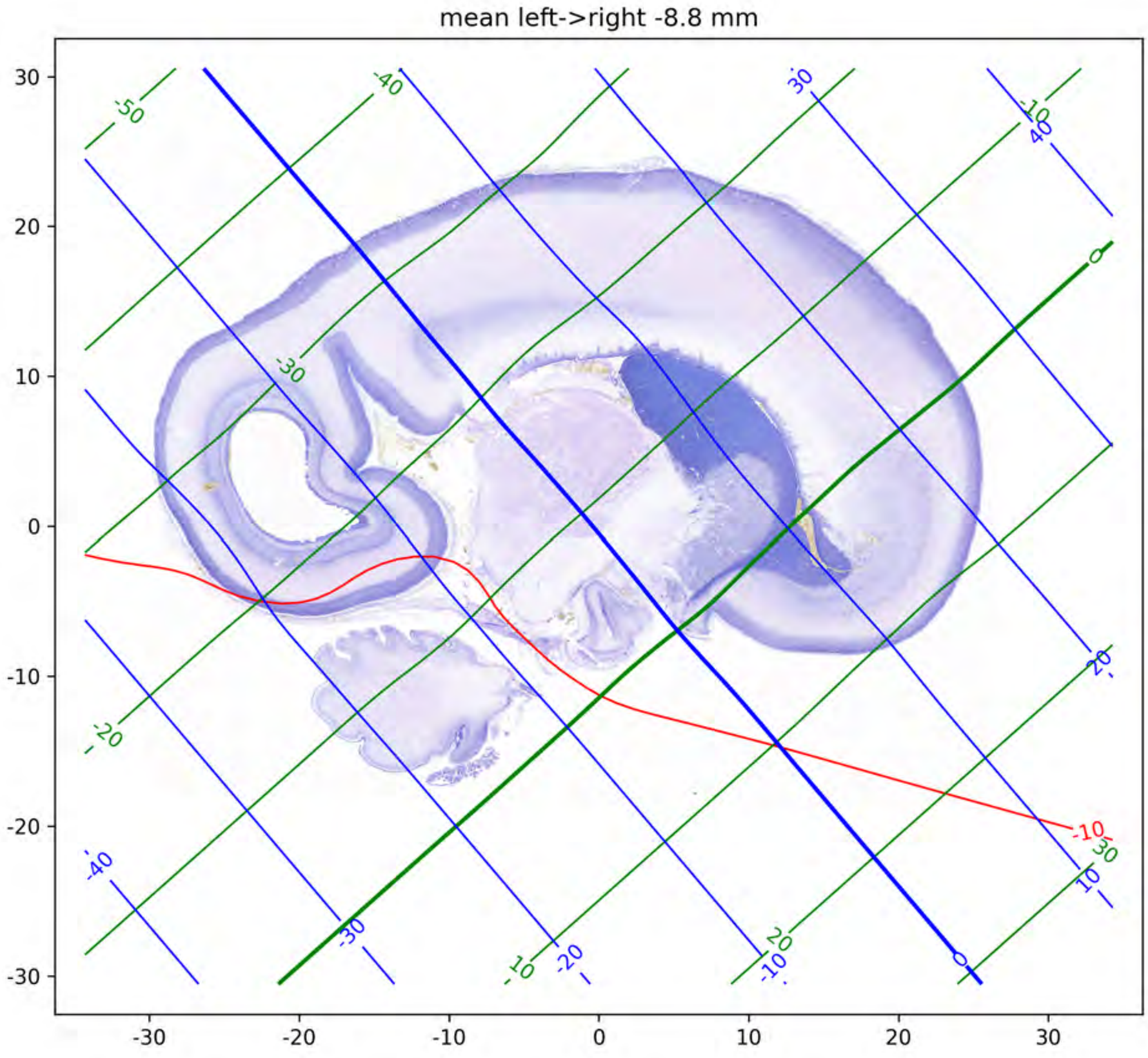
mean left->right -8.8 mm
30
20
10
0
-10
-20
-30
-30
-20
-10
0
10
20
30
-50
-40
-30
-20
-10
0
10
20
30
-10
30
40
-40
-30
-20
-10
0
10
20
-10

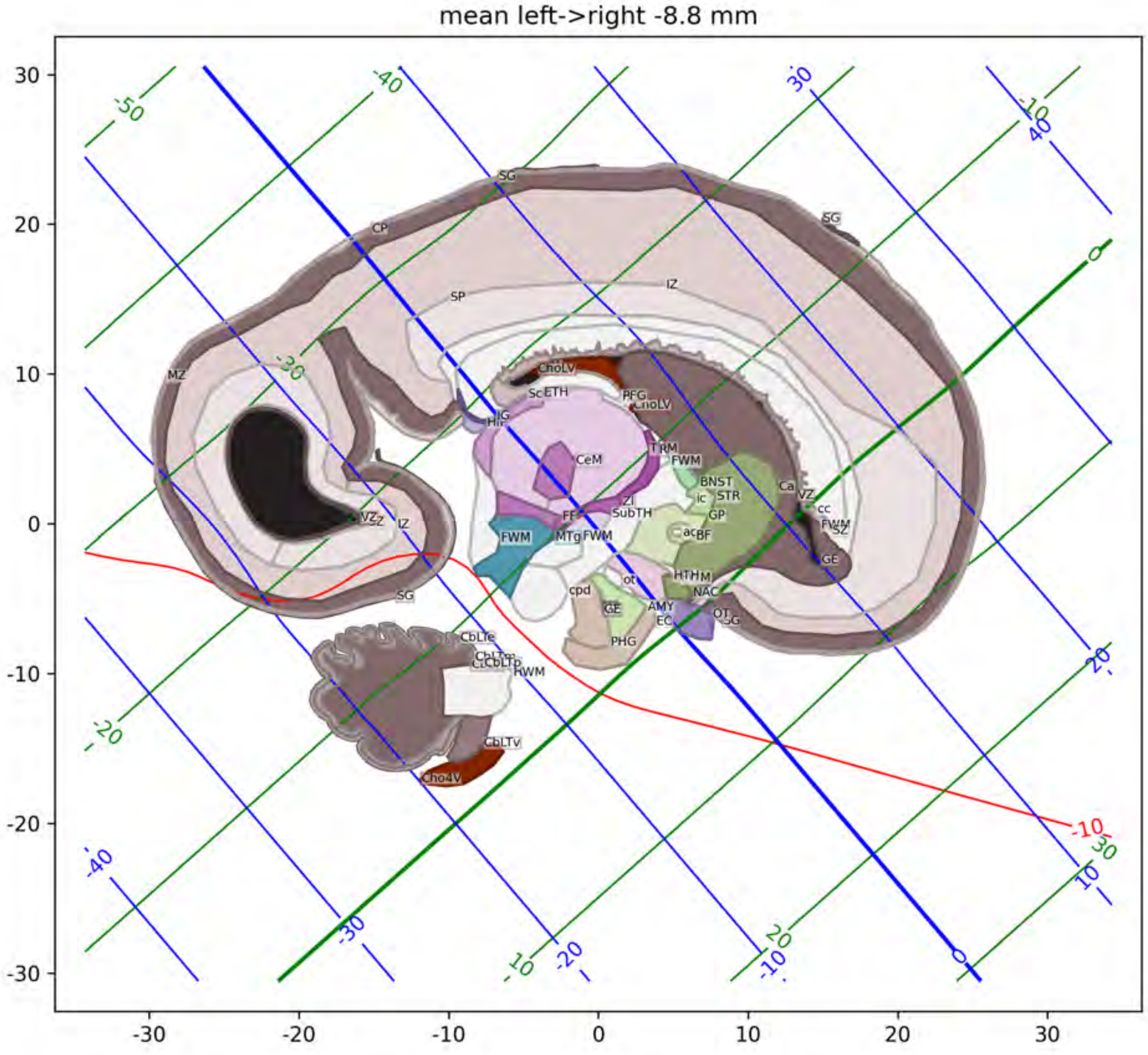
mean left->right -8.8 mm
SG
CP
SP
IZ
MZ
VZ
ChoLV
ScETH
PFG
CeM
TRM
FWM
BNST
STR
GP
Ca
cc
ZI
SubTH
MTg
BF
GE
HTHM
NAC
cpd
ot
AMY
EC
OT
PHG
CbLTe
CbLTv
Cho4V
-10
-50
-40
-30
-20
0
10
20
30
40
-30
-20
-10
0
10
20
30

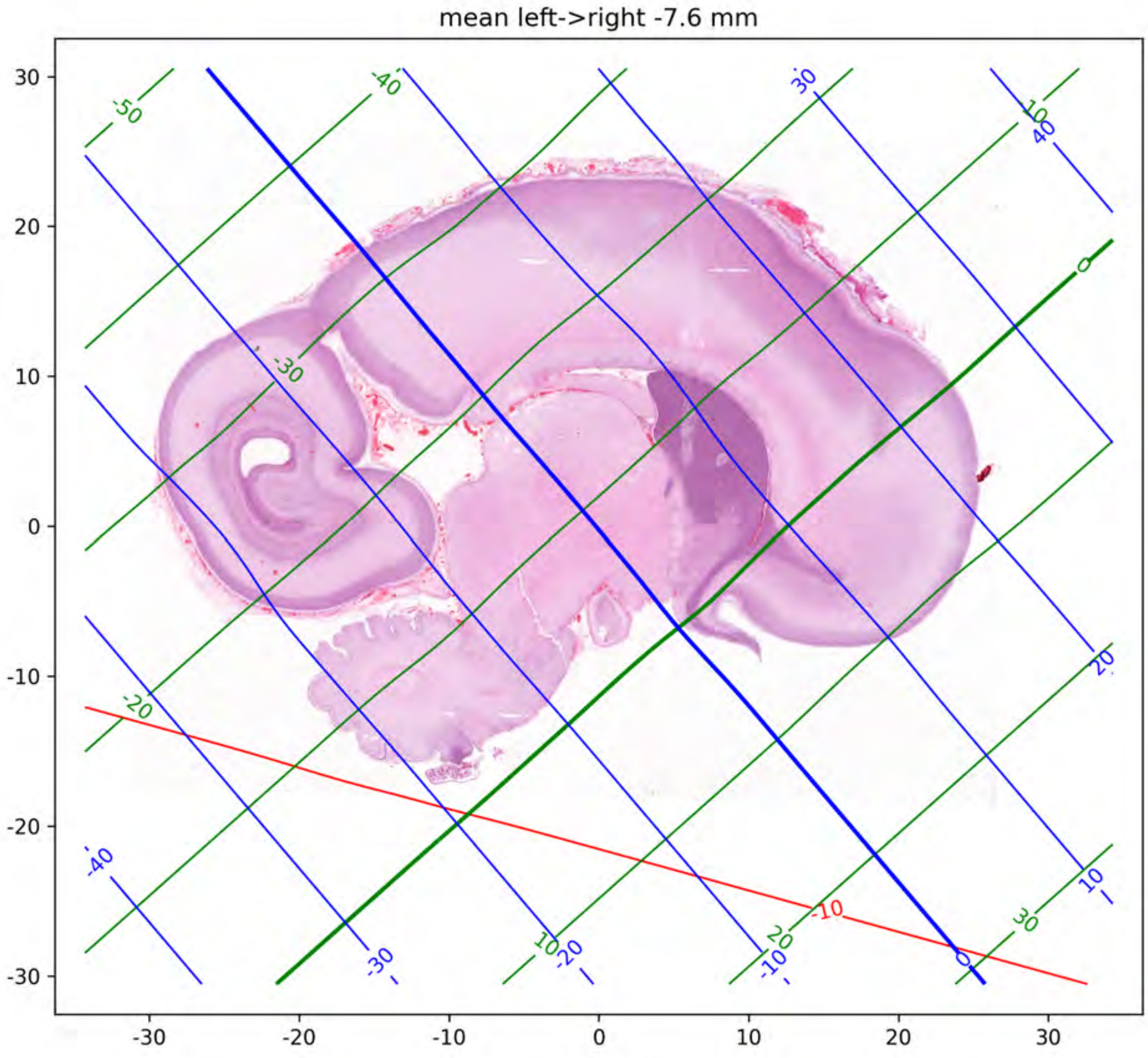
mean left->right -7.6 mm

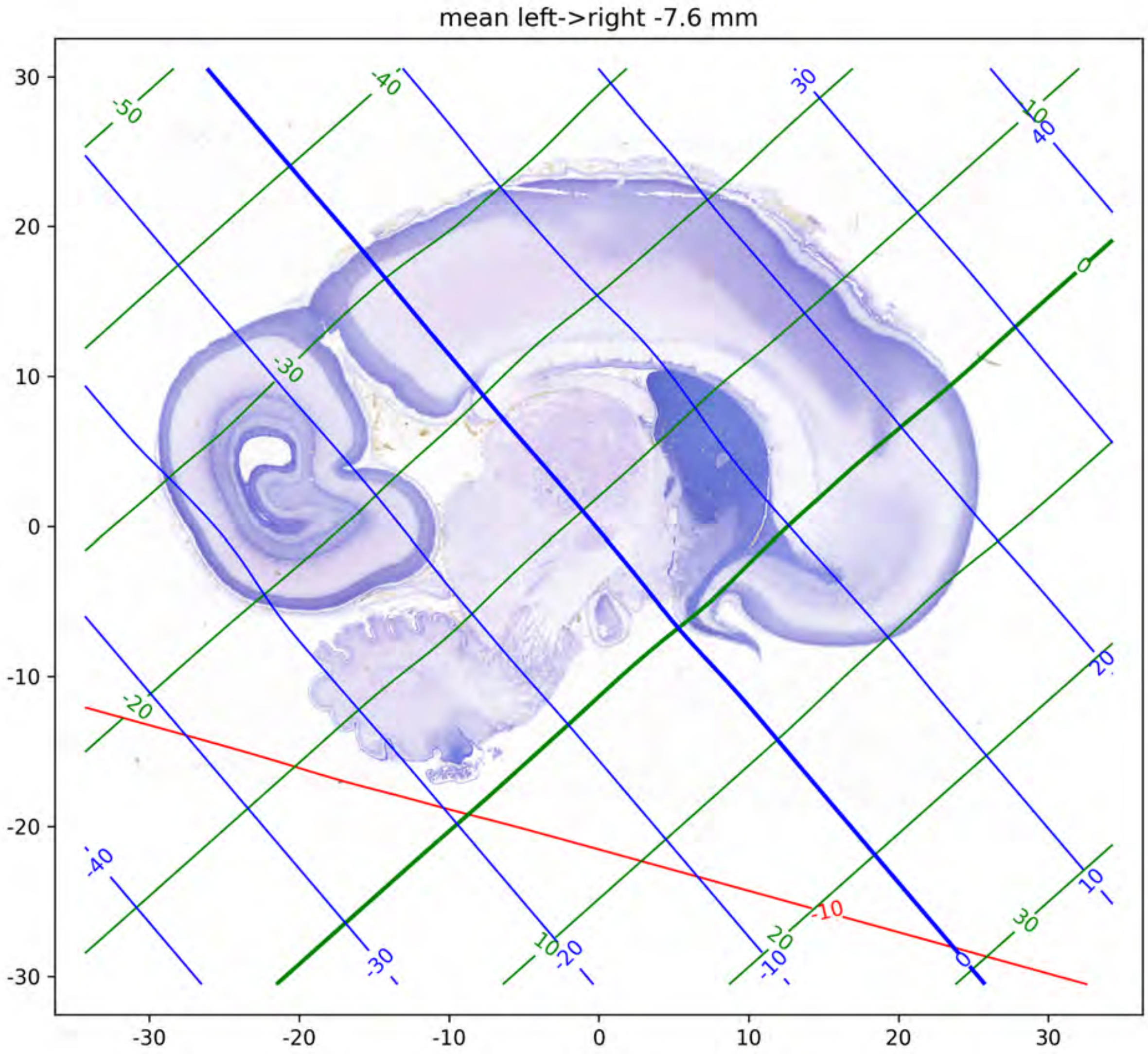

mean left->right -7.6 mm
30
20
10
0
-10
-20
-30
-30
-20
-10
0
10
20
30

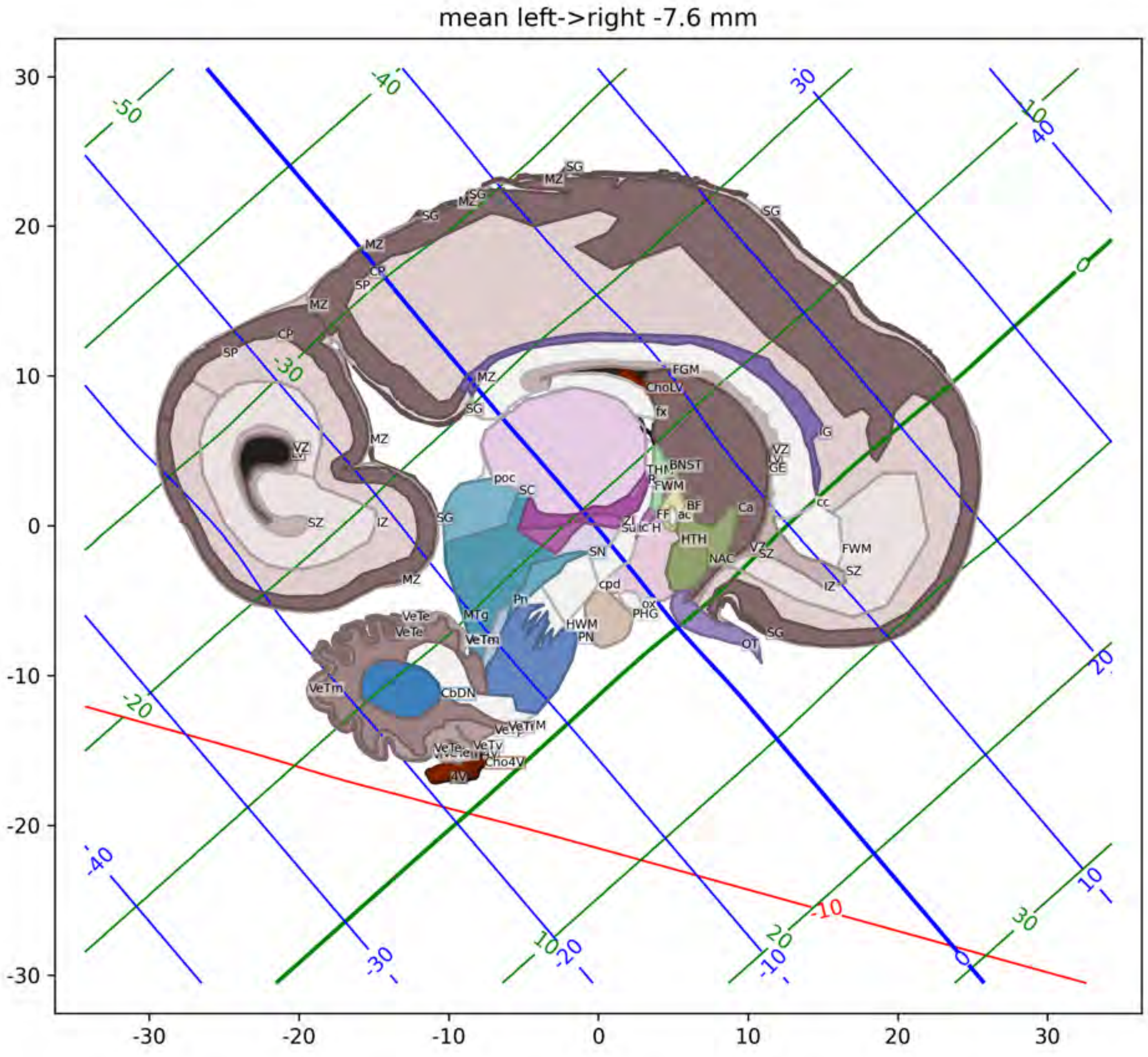
mean left->right -7.6 mm
30
20
10
0
-10
-20
-30
-30
-20
-10
0
10
20
30
-50
-40
-30
-20
-40
-30
-20
-10
0
10
20
30
30
40
10
0
20
10
-10
SG
MZ
CP
SP
FGM
ChoLV
fx
IG
VZ
LV
GE
THN
BNST
FWM
poc
SC
BF
ac
Ca
cc
SZ
IZ
FF
Su
H
HTH
SN
NAC
cpd
Pn
ox
PHG
VeTe
MTg
VeTm
HWM
PN
OT
CbDN
VeTv
Cho4V
4V

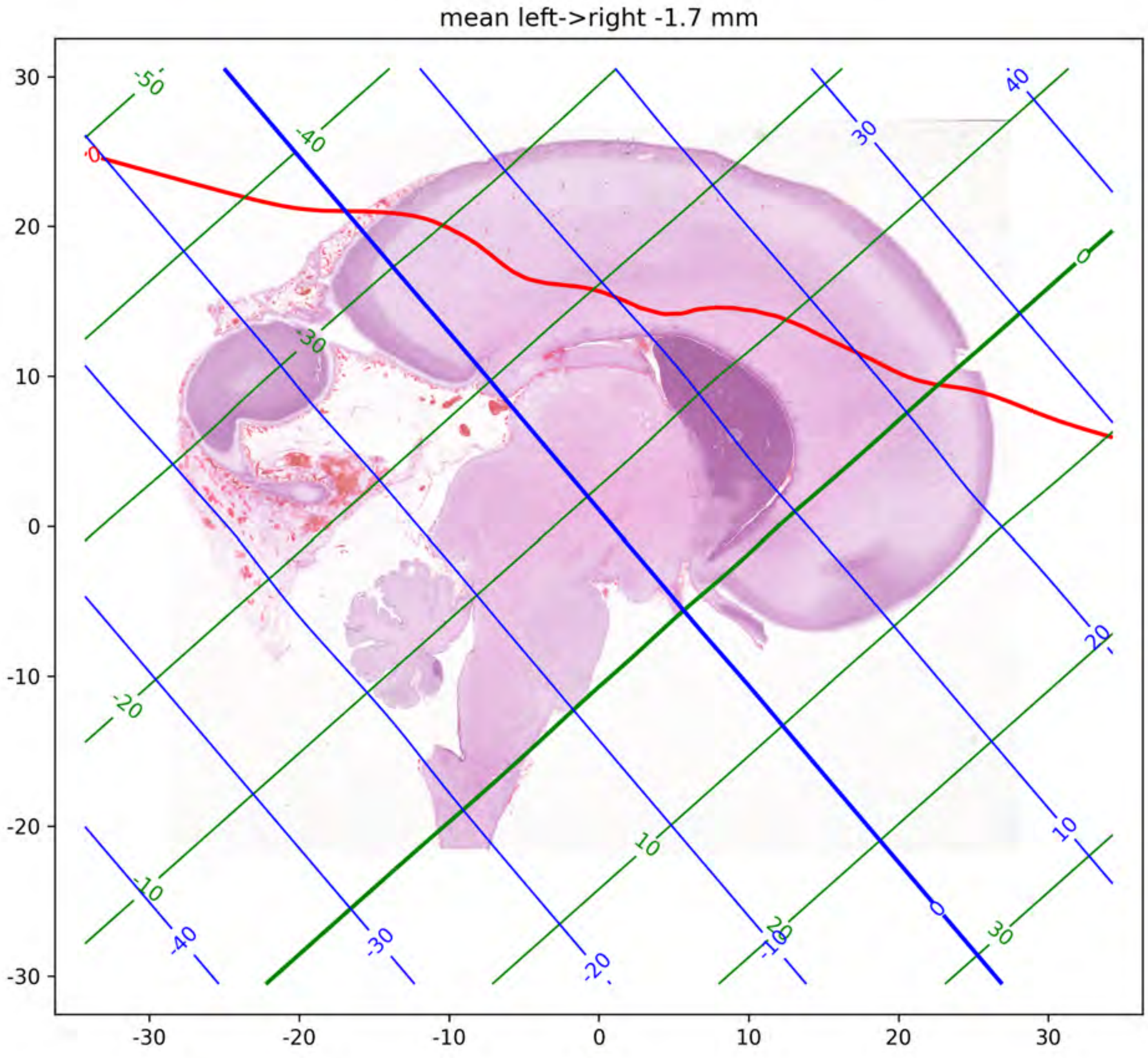

mean left->right -1.7 mm
30
20
10
0
-10
-20
-30
-30
-20
-10
0
10
20
30

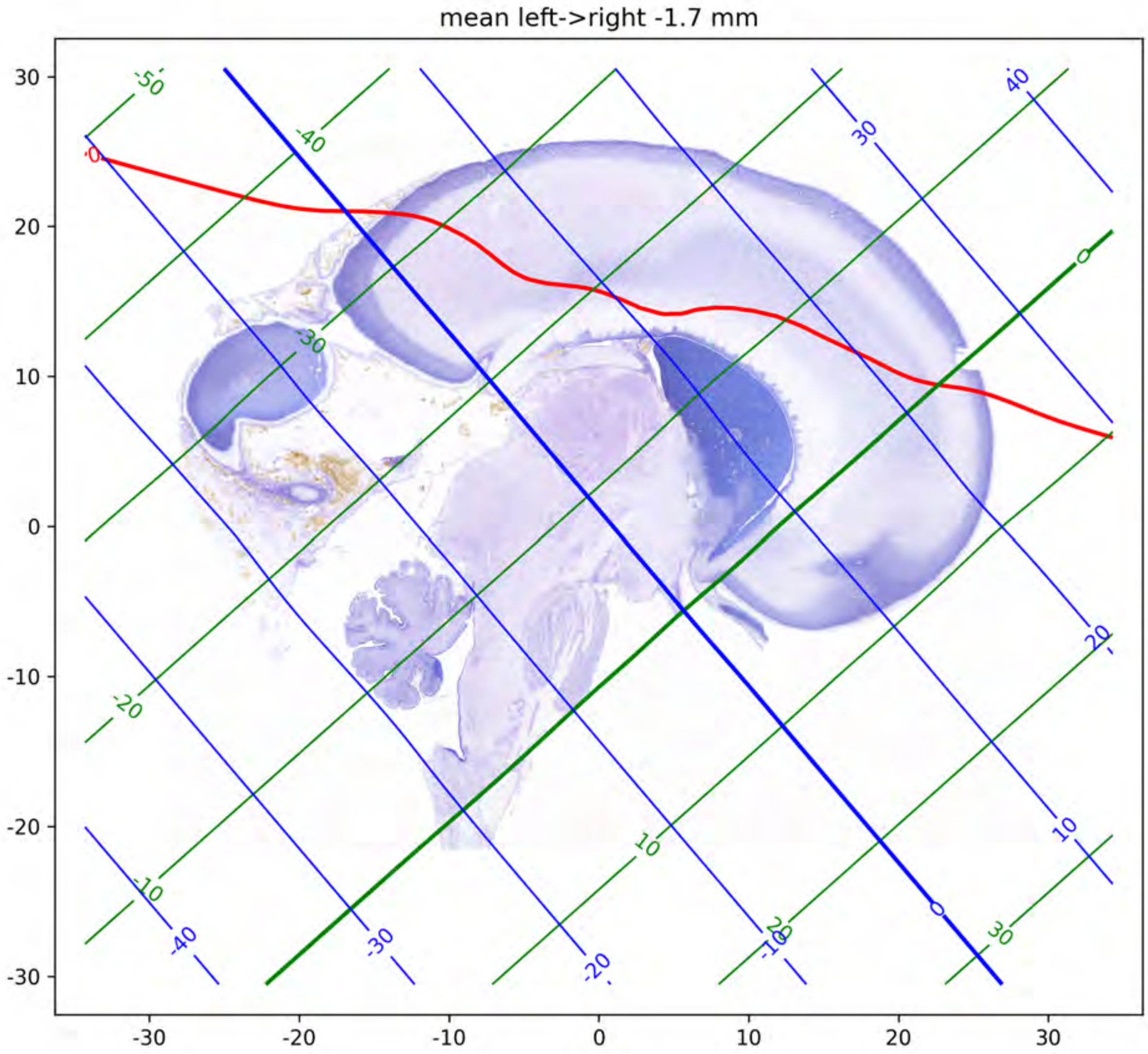
mean left->right -1.7 mm

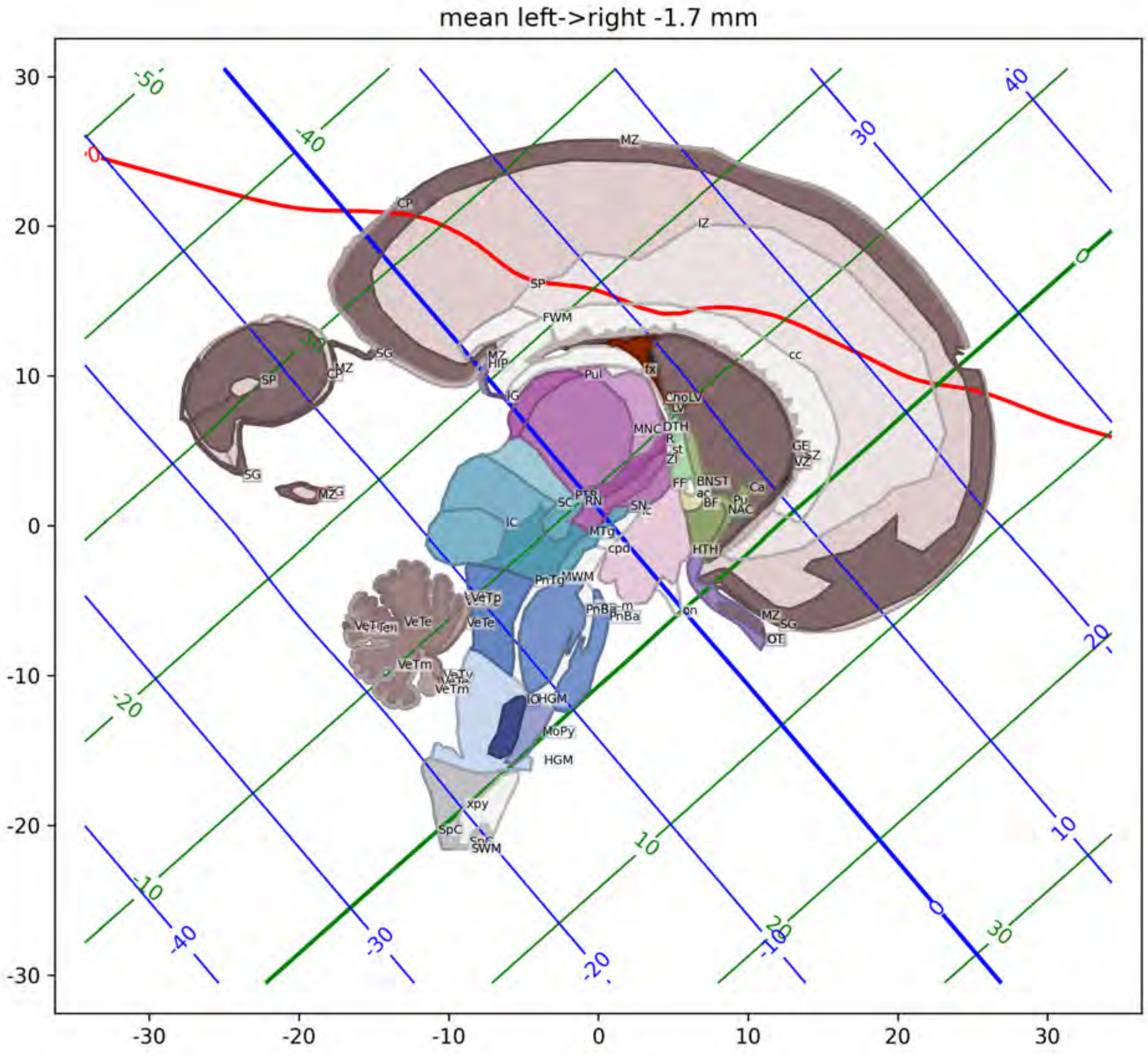
mean left->right -1.7 mm
MZ
CP
IZ
SP
FWM
cc
SG
HIP
Pul
fx
LV
DTH
MNC
ZI
GE
VZ
FF
BNST
ac
BF
Pu
NAC
Ca
SC
RN
SN
IC
MTg
cpd
HTH
PnTg
MWM
VeTp
VeTe
VeTm
PnBa
on
OT
MoPy
HGM
xpy
SpC
SWM

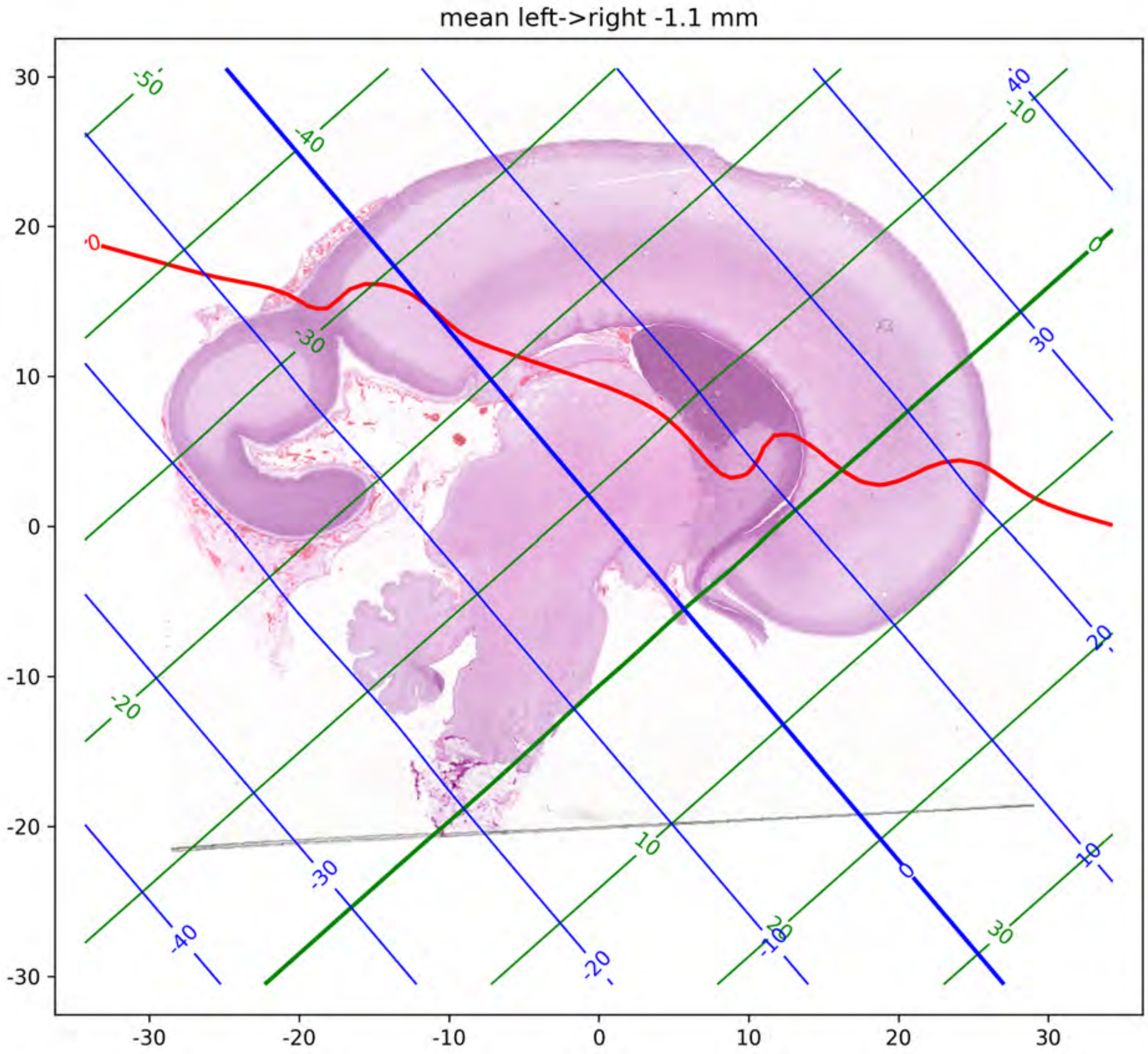
mean left->right -1.1 mm

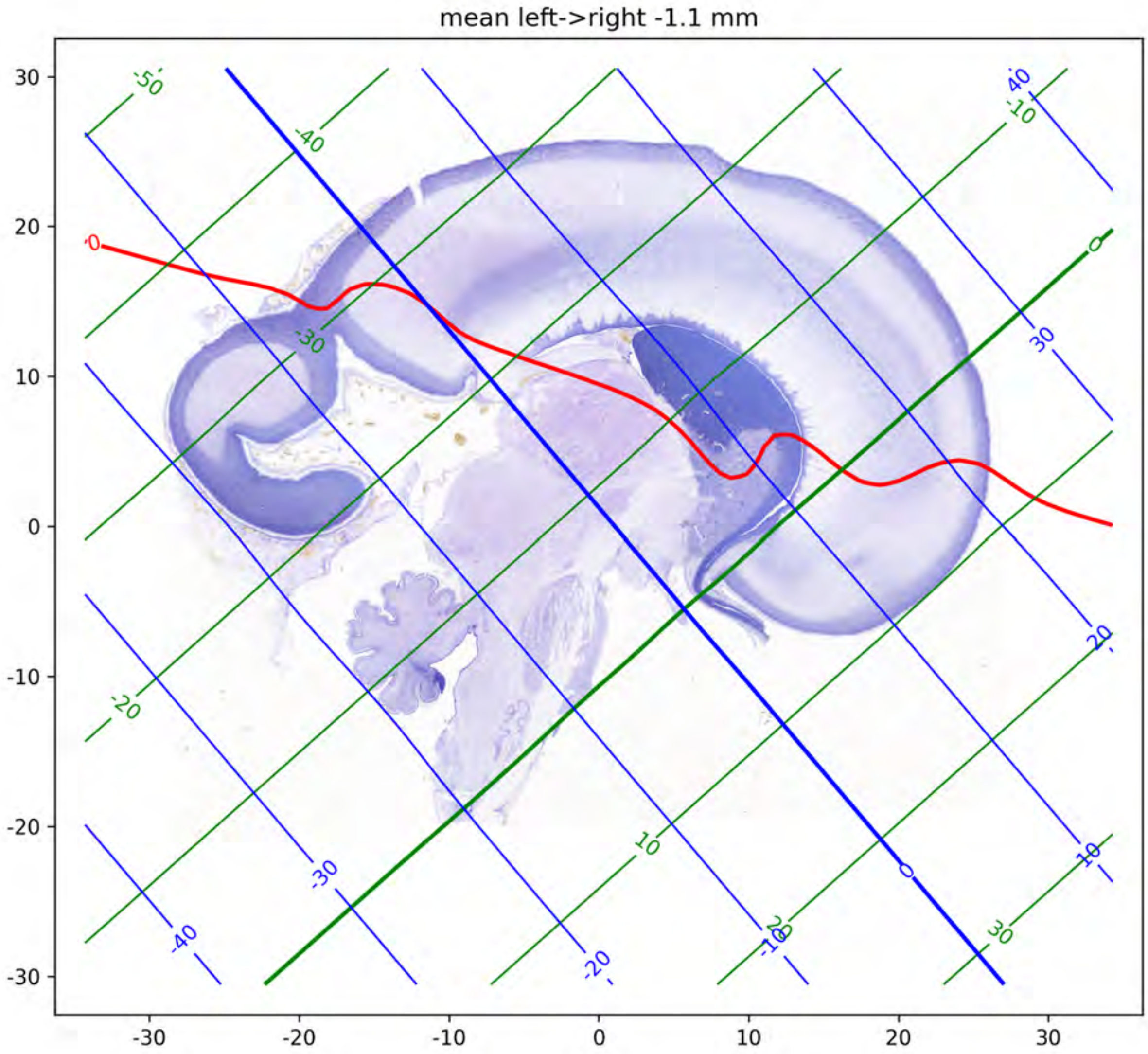
mean left->right -1.1 mm
30
20
10
0
-10
-20
-30
-30
-20
-10
0
10
20
30

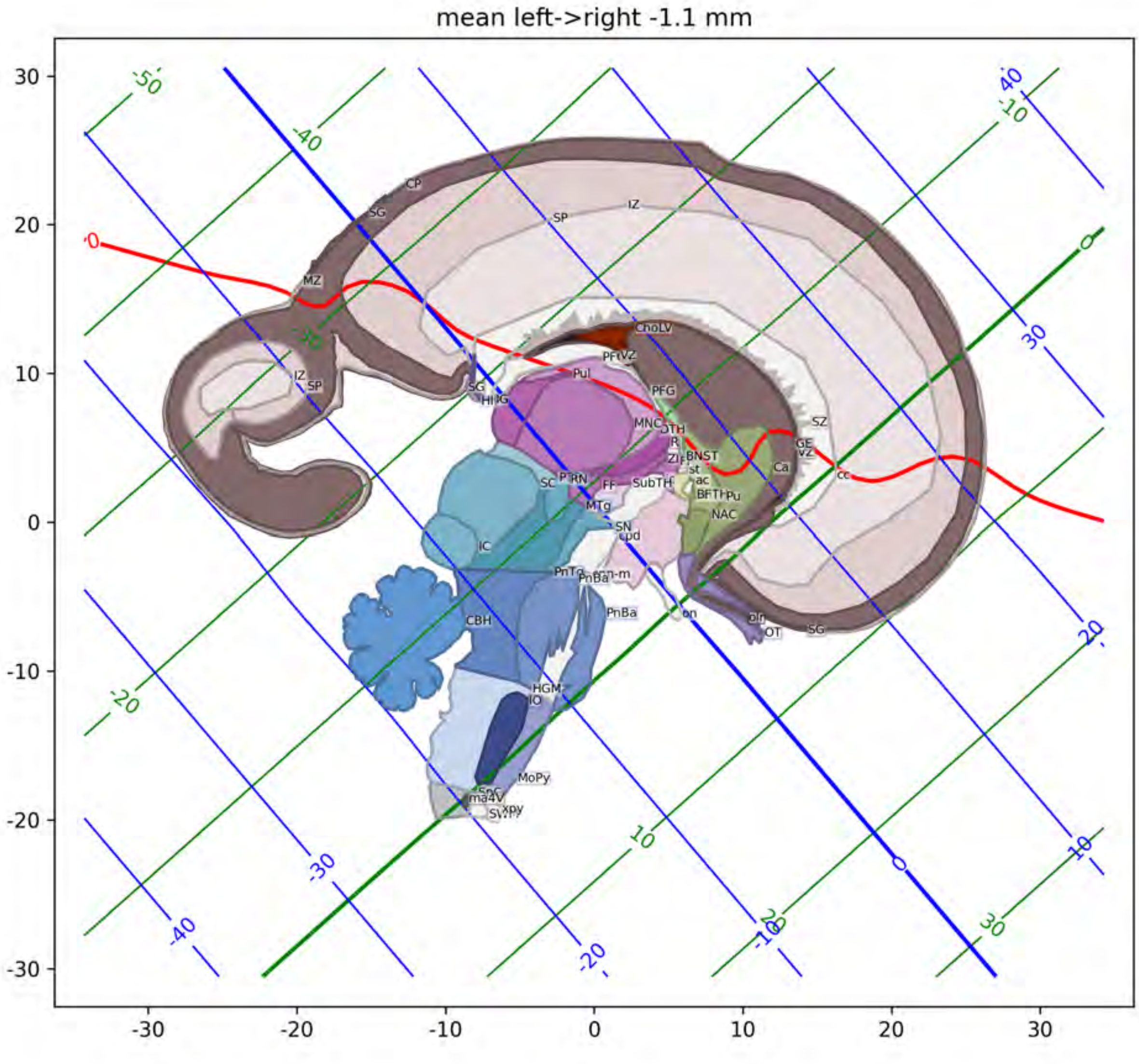

mean left->right -1.1 mm
CP
SG
SP
IZ
MZ
ChoLV
PFG
Pul
MNC
BNST
SZ
GE
VZ
Ca
cc
NAC
SN
IC
CBH
PnBa
HGM
IO
MoPy
OT

mean left->right 0.1 mm

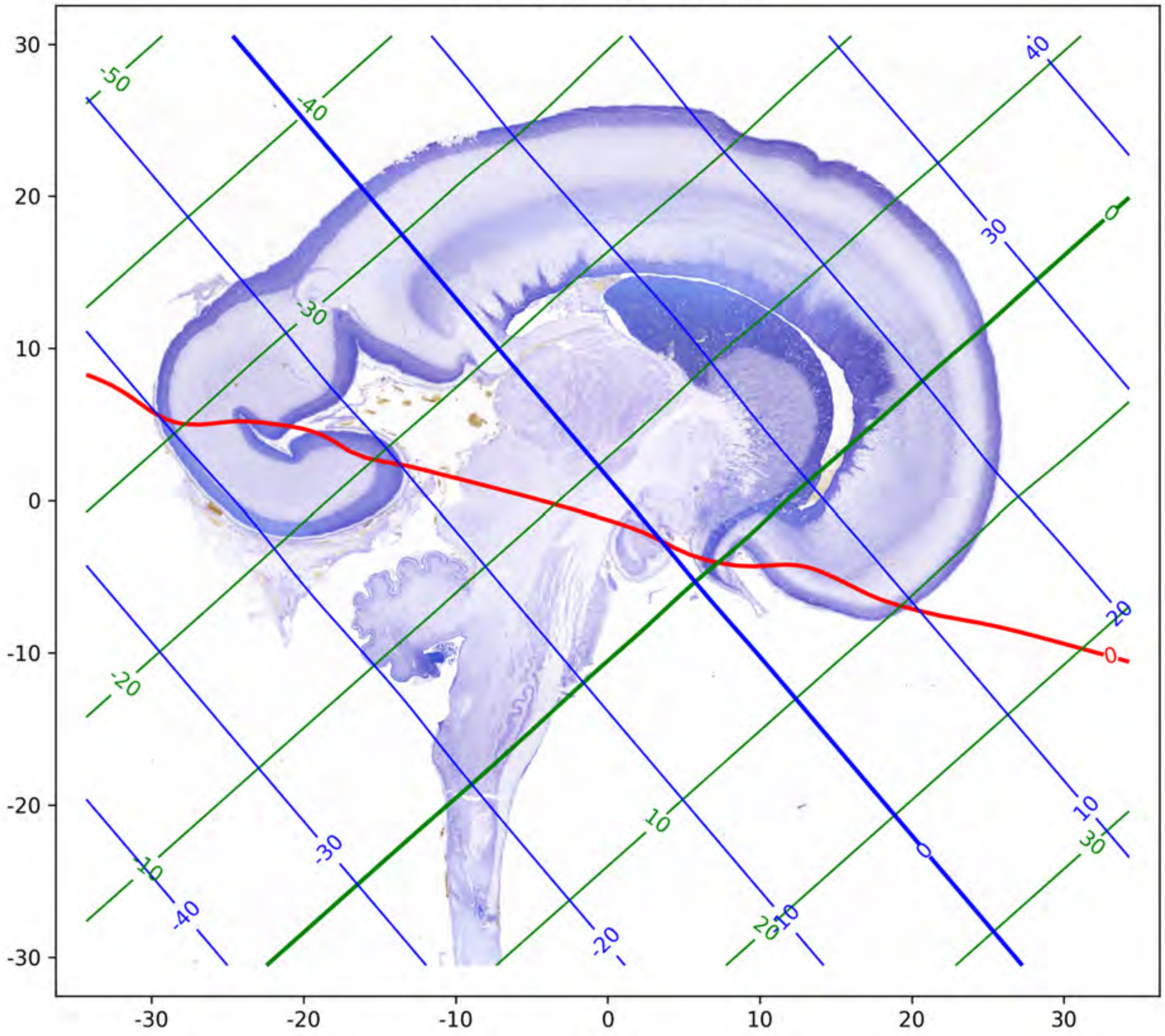

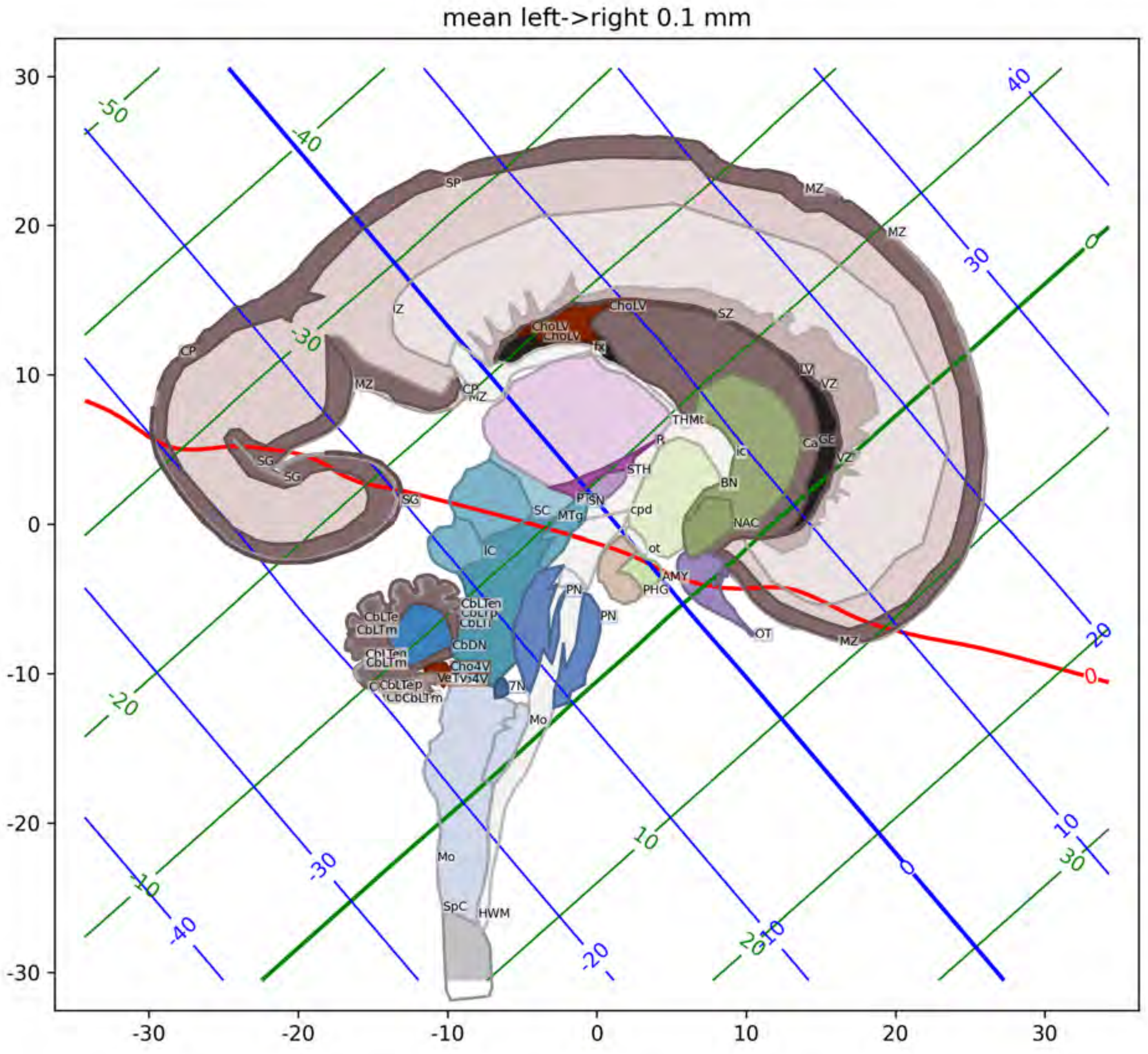
mean left->right 0.1 mm
SP
MZ
IZ
CP
ChoLV
SZ
SG
LV
VZ
THMt
IC
Ca
STH
BN
SN
cpd
NAC
SC
MTg
ot
AMY
PHG
PN
OT
CbLTen
CbDN
Cho4V
7N
Mo
SpC
HWM

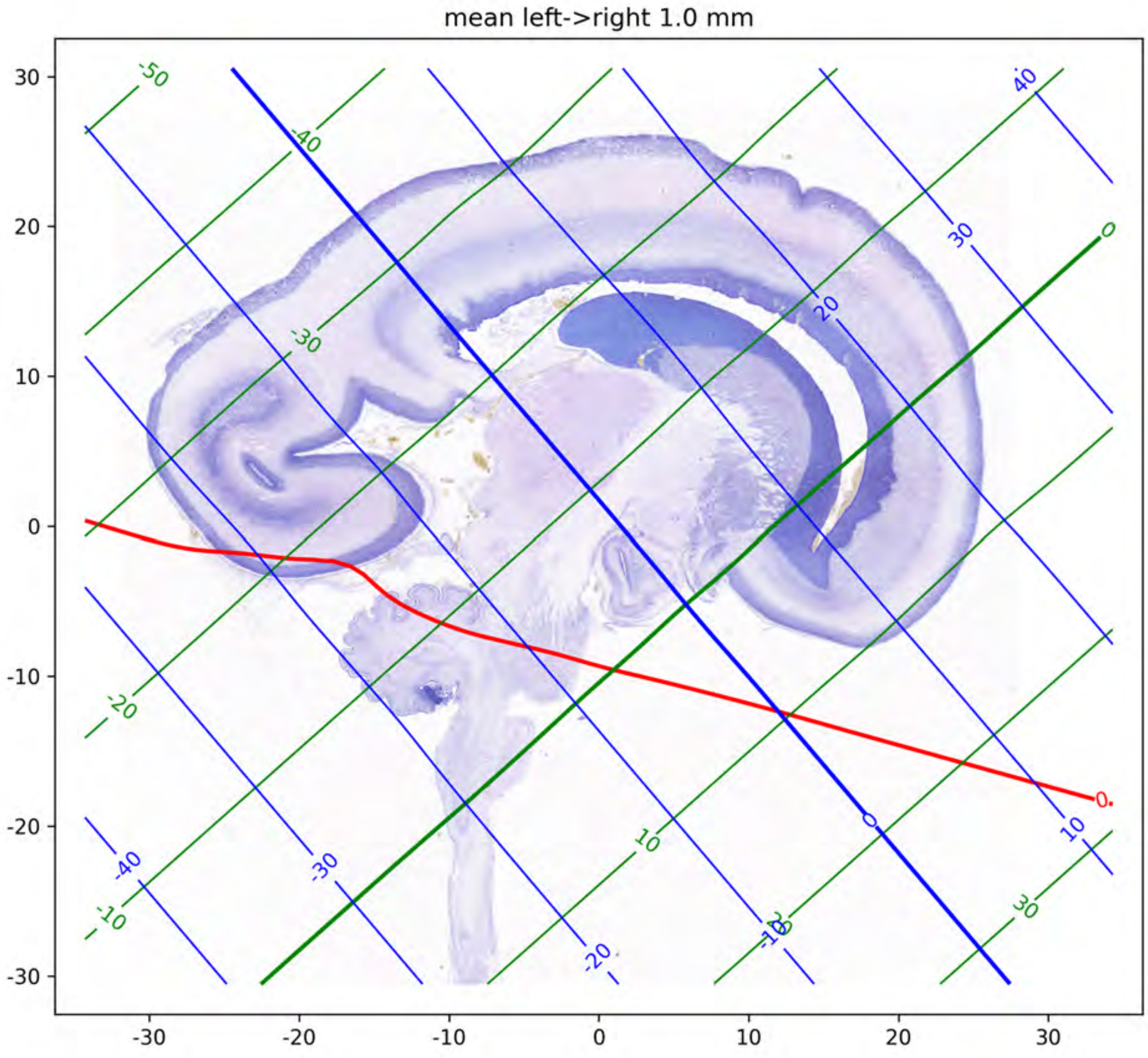

mean left->right 1.0 mm

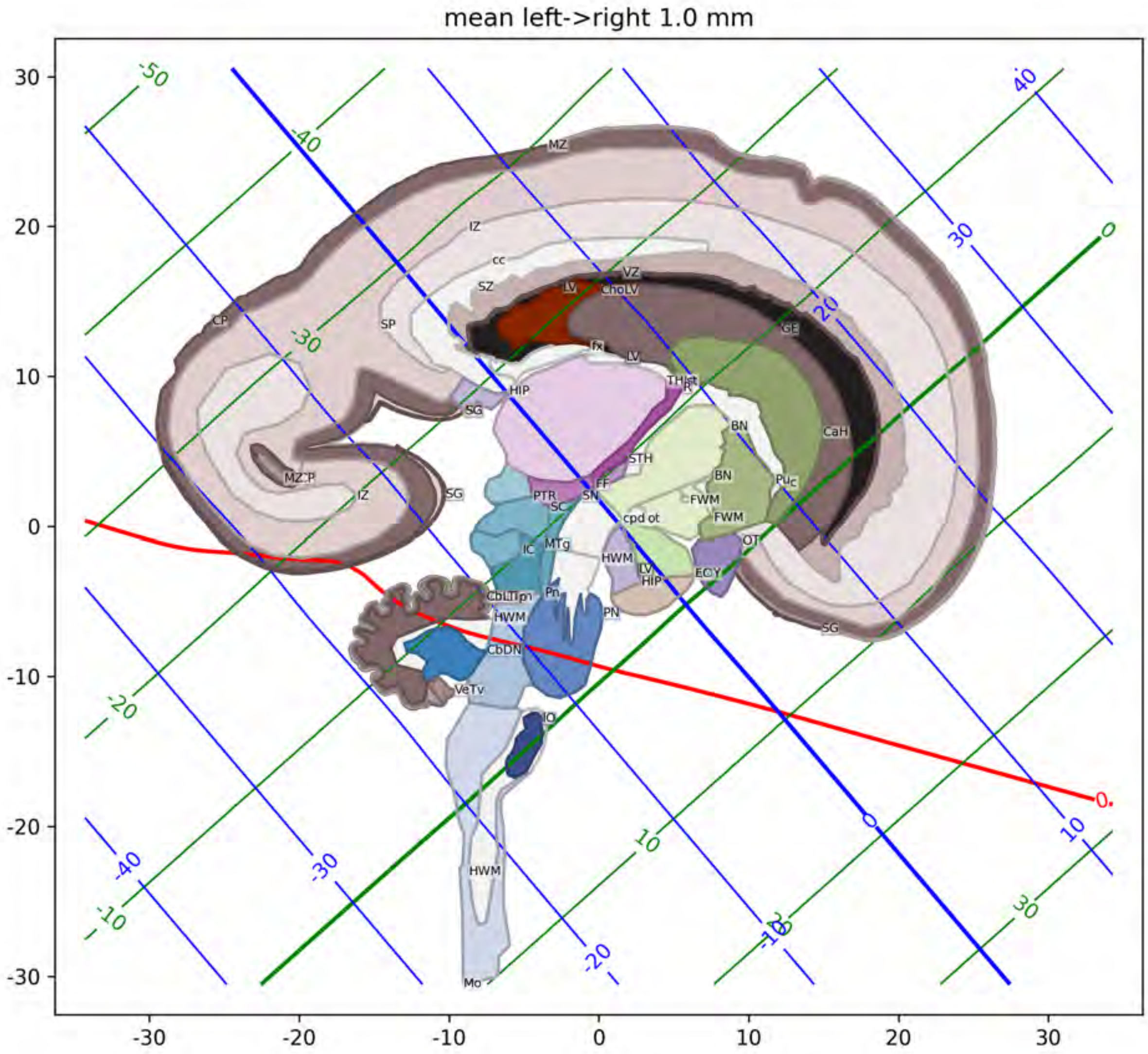
mean left->right 1.0 mm
MZ
IZ
cc
SZ
LV
VZ
ChoLV
CP
SP
fx
LV
GE
HIP
THlat
SG
BN
CaH
STH
BN
MZ:P
IZ
SG
PTR
SN
FF
FWM
Puc
SC
cpd ot
FWM
IC
MTg
HWM
OT
LV
HIP
ECY
CbLTipn
Pn
HWM
PN
SG
CbDN
VeTv
IO
HWM
Mo

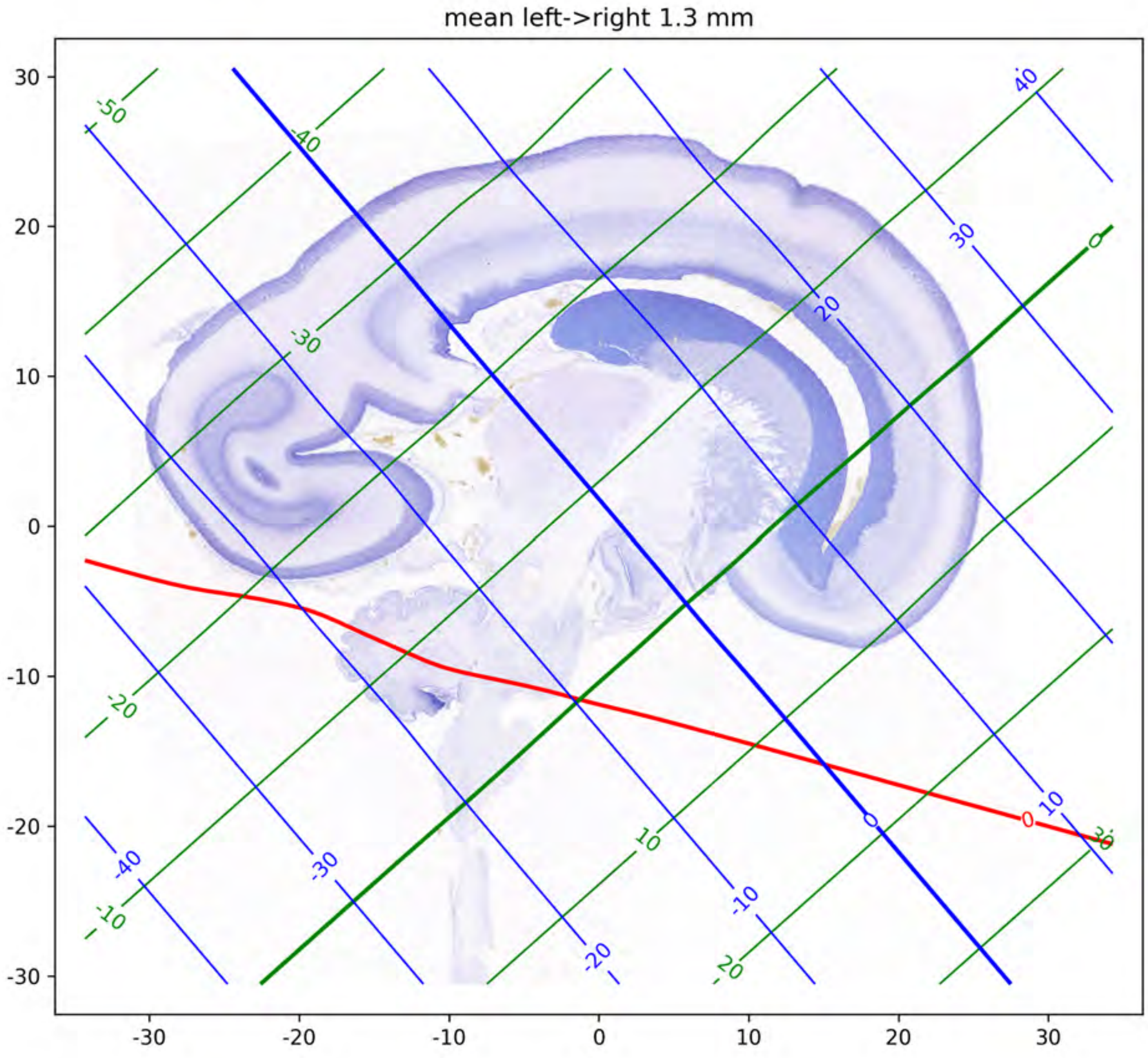
mean left->right 1.3 mm
30
20
10
0
-10
-20
-30
-30
-20
-10
0
10
20
30
-50
-40
-30
-20
-10
0
10
20
30
-40
-30
-20
-10
0
10
20
30
40
0

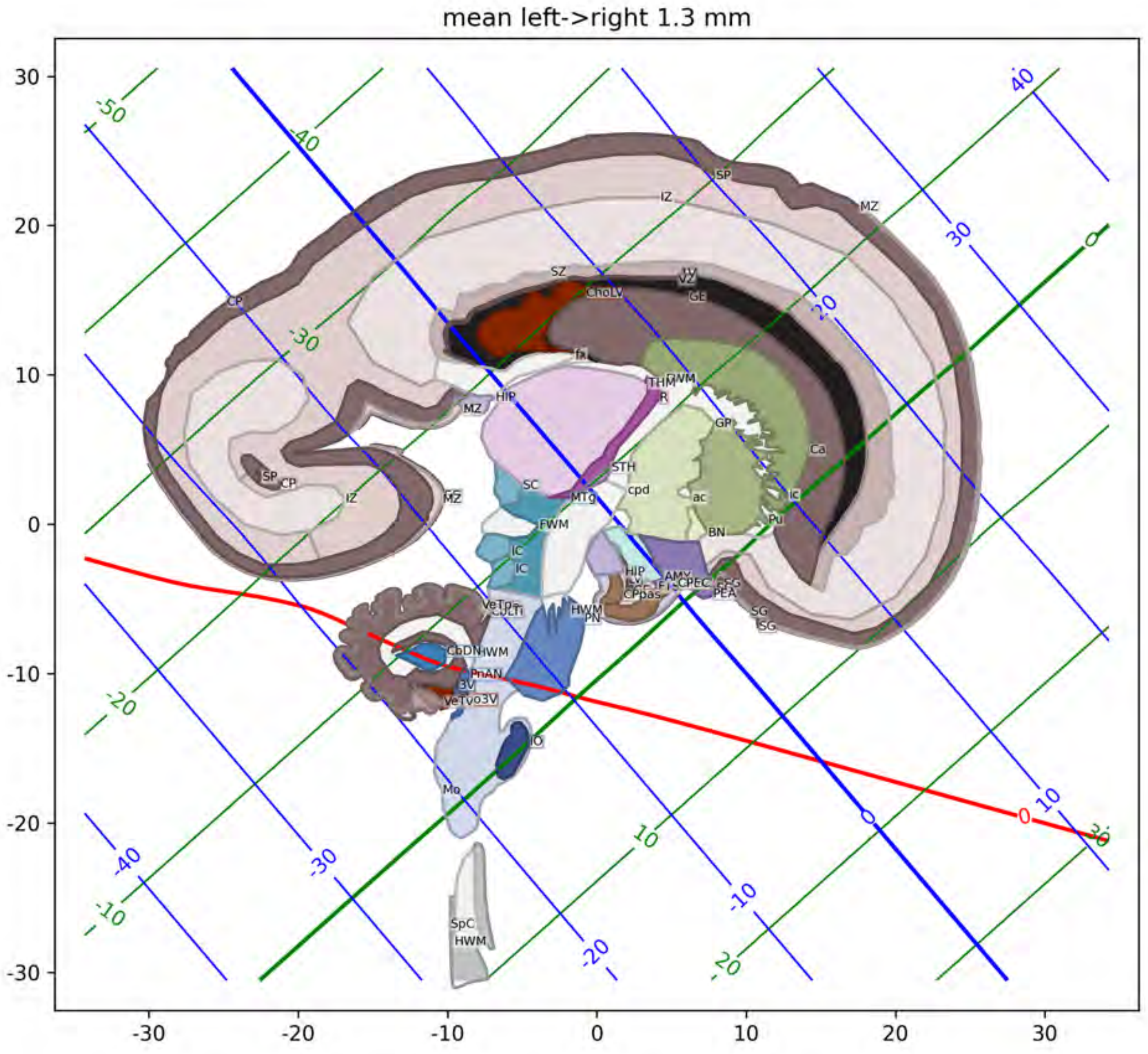
mean left->right 1.3 mm
30
20
10
0
-10
-20
-30
-30
-20
-10
0
10
20
30
SP
IZ
MZ
CP
SZ
ChoLV
GE
fx
HIP
MZ
THM
R
GP
Ca
STH
cpd
ac
ic
Pu
BN
SP
CP
IZ
MZ
SC
MTg
FWM
IC
IC
HIP
AMY
PEA
SG
SG
HWM
PN
CbDN
HWM
3V
IO
Mo
SpC
HWM

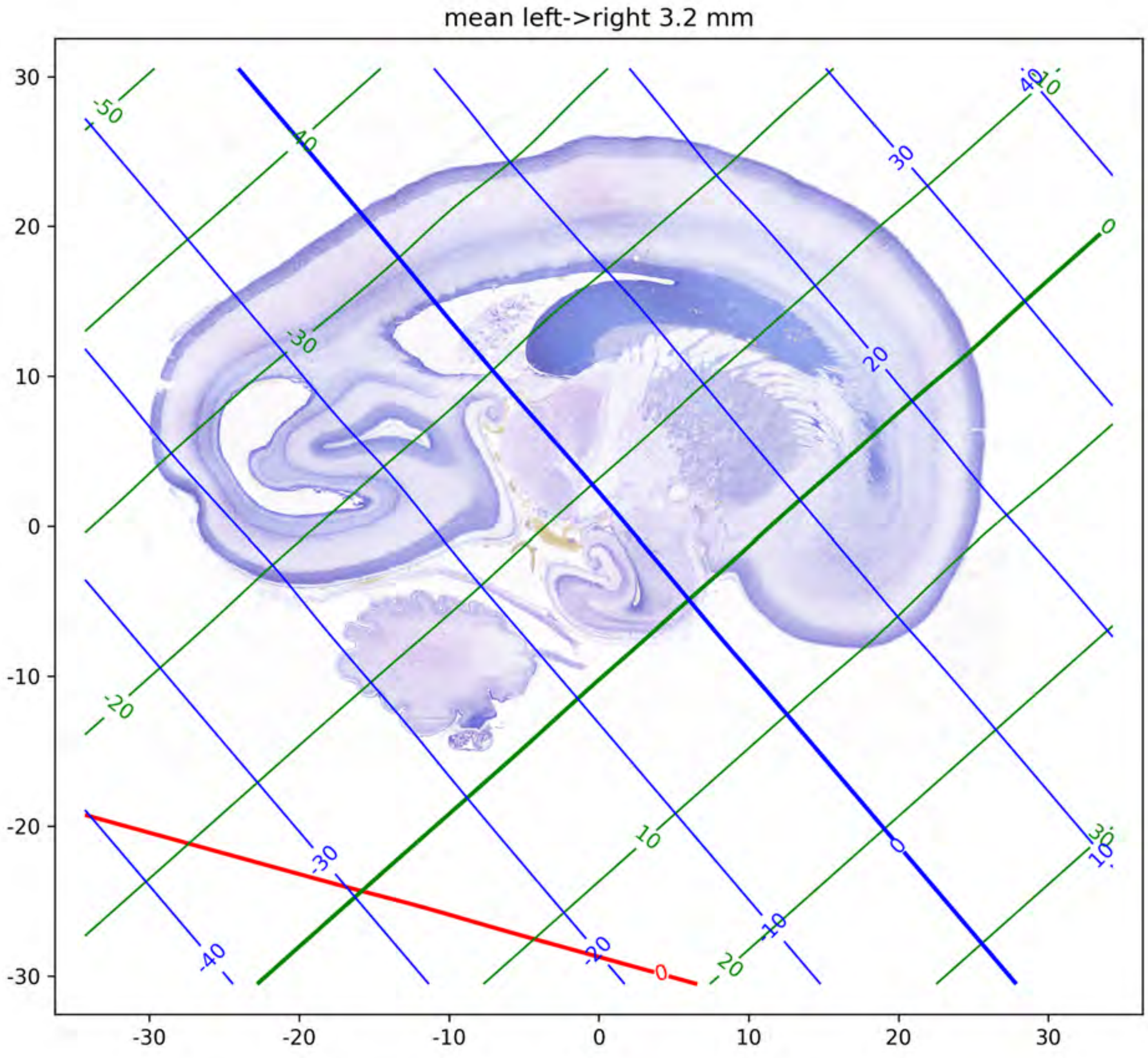

mean left->right 3.2 mm

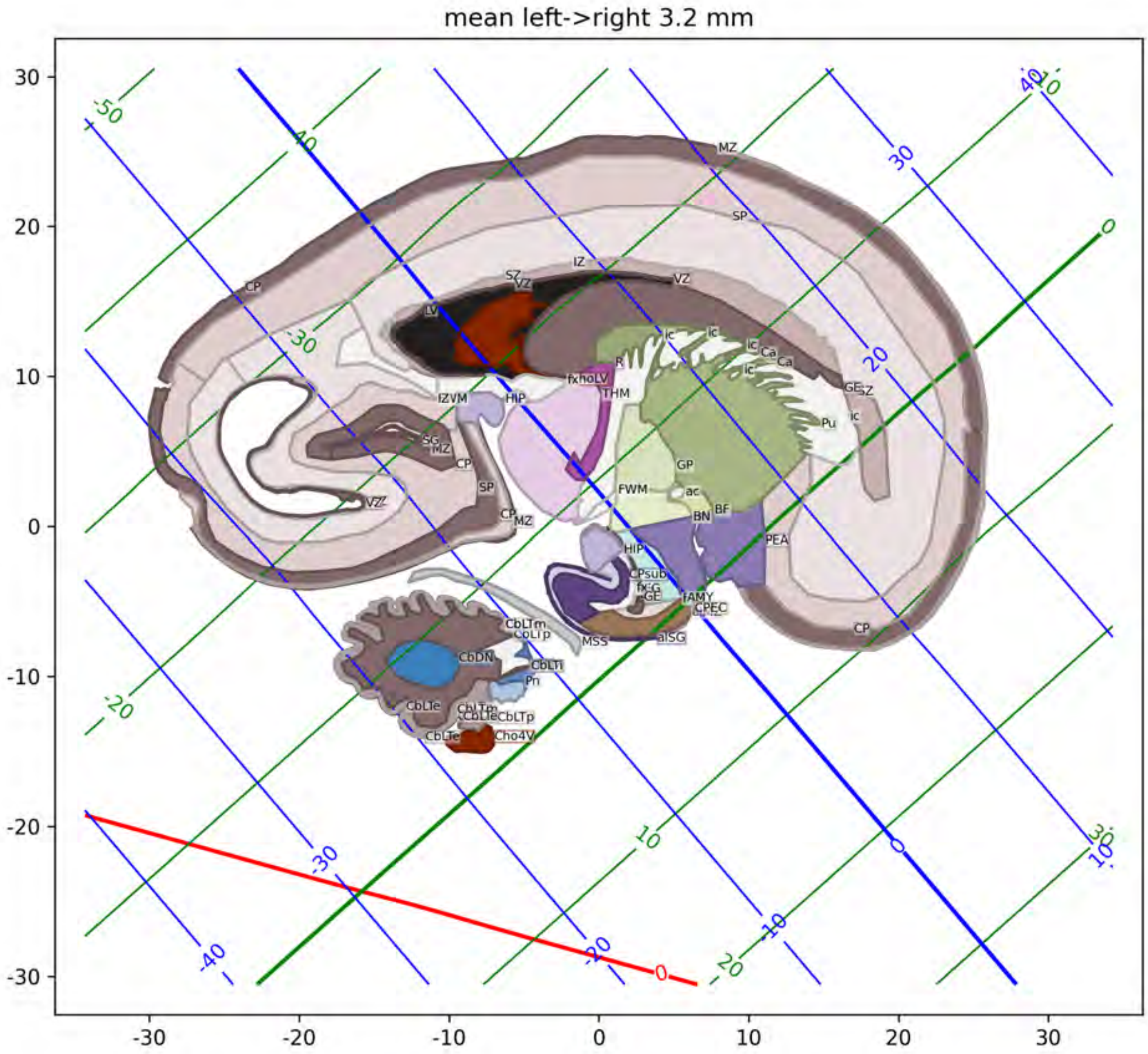

mean left->right 3.2 mm
MZ
SP
CP
IZ
SZ
VZ
LV
ic
Ca
R
fxhoLV
THM
IZVM
HIP
GE
SZ
Pu
SG
MZ
CP
SP
GP
FWM
ac
BN
BF
PEA
VZ
CP
MZ
HIP
CPsub
fAMY
CPEC
MSS
aISG
CbLTm
CbDN
CbLTi
Pn
CbLTe
CbLTp
Cho4V

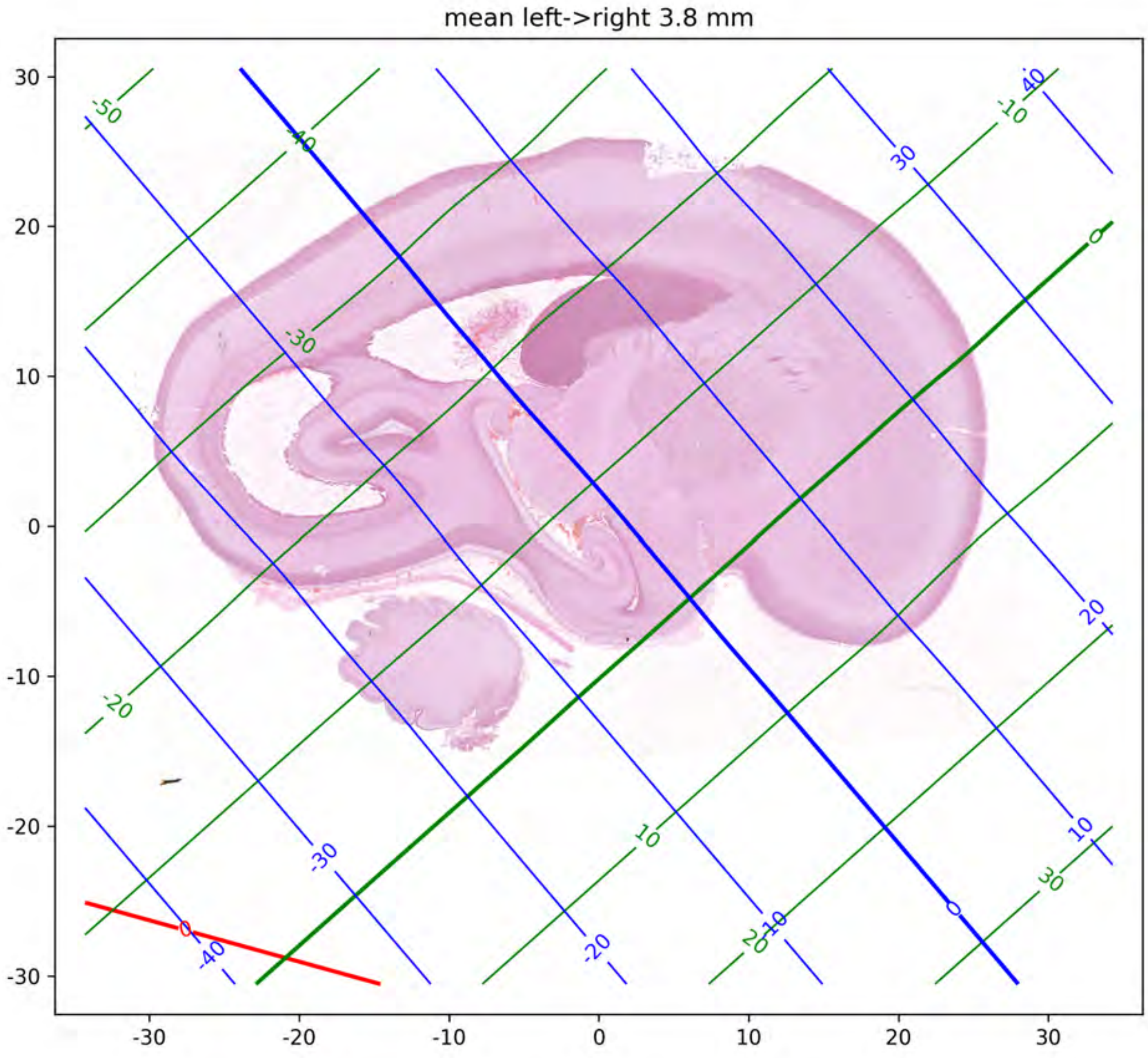

mean left->right 3.8 mm

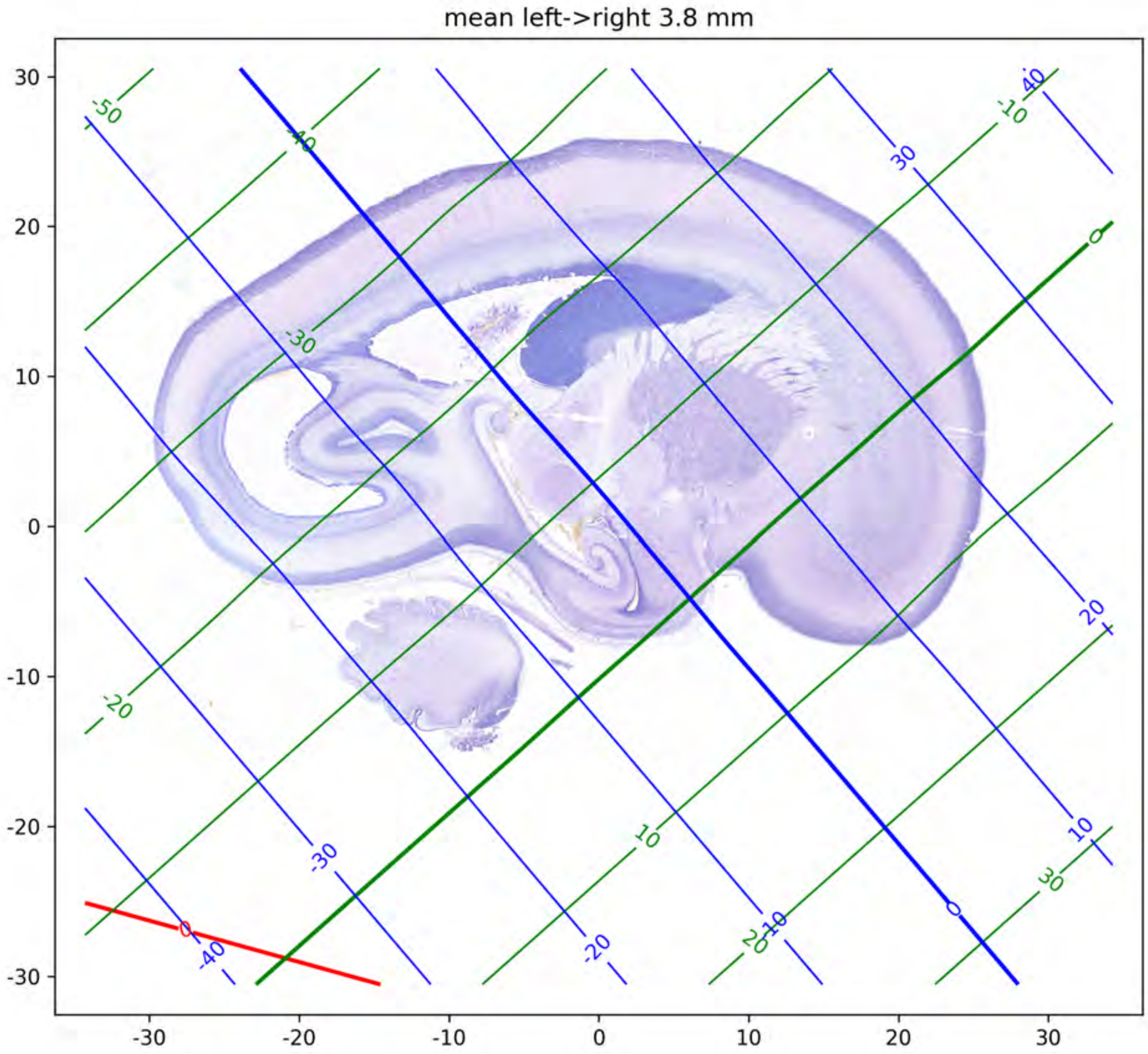
mean left->right 3.8 mm

mean left->right 3.8 mm

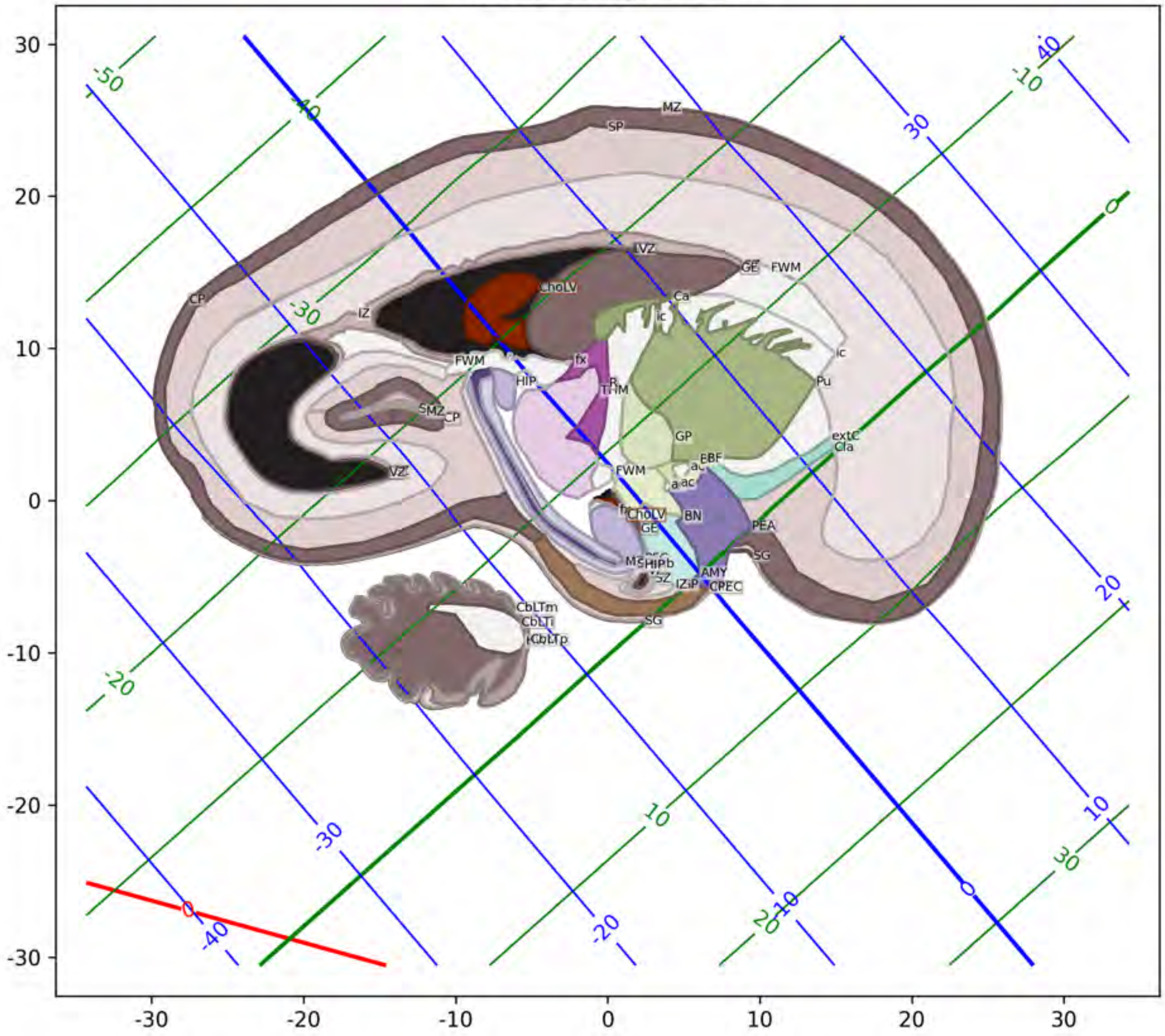

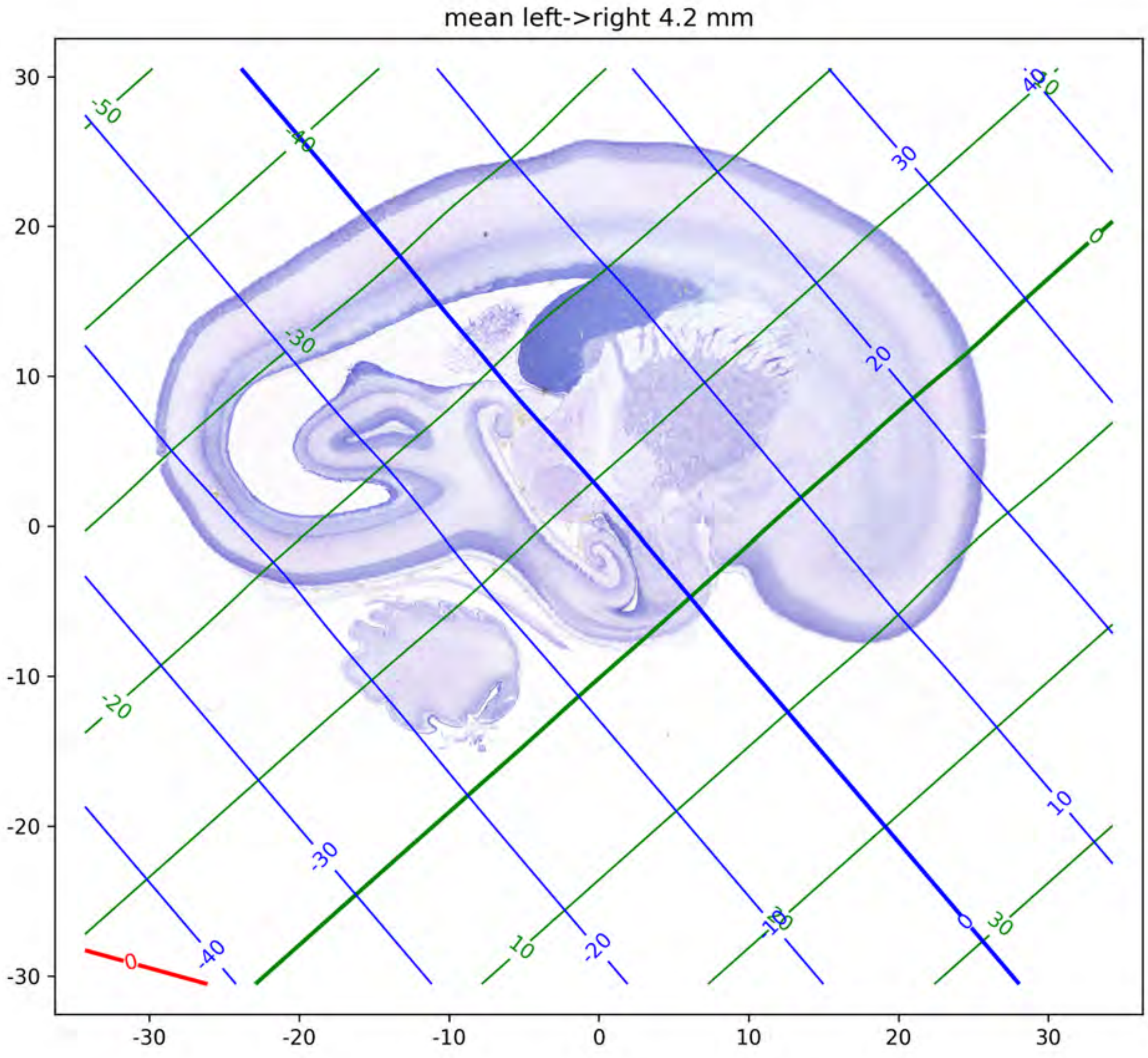
mean left->right 4.2 mm
30
20
10
0
-10
-20
-30
-30
-20
-10
0
10
20
30

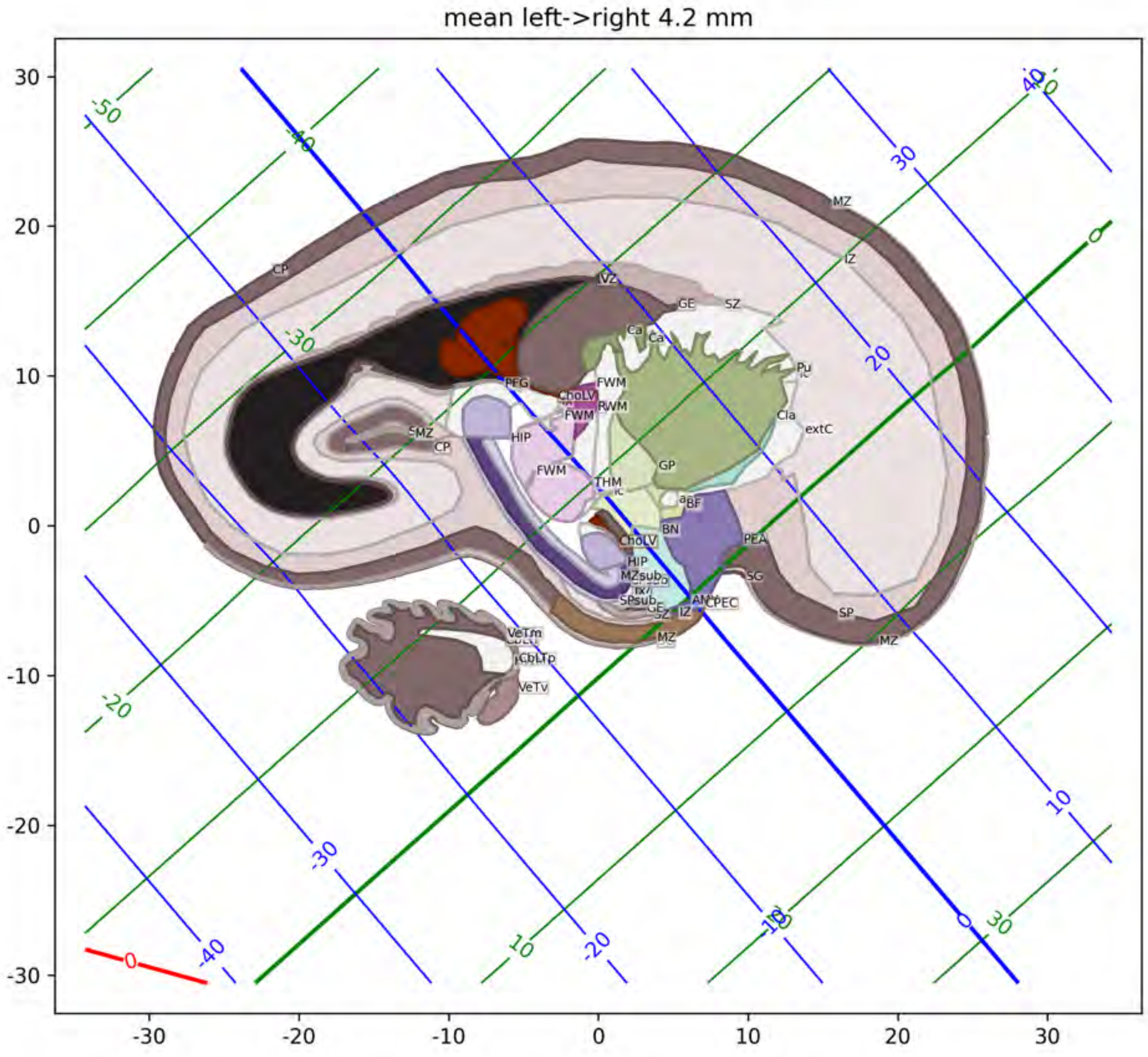

mean left->right 4.2 mm
MZ
IZ
CP
VZ
GE
SZ
Ca
Pu
PFG
ChoLV
FWM
RWM
Cla
extC
SMZ
CP
HIP
GP
THM
BF
BN
PEA
SG
SP
ChoLV
HIP
MZsub
SPsub
IZ
AN
PEC
VeTm
CbLTp
VeTv

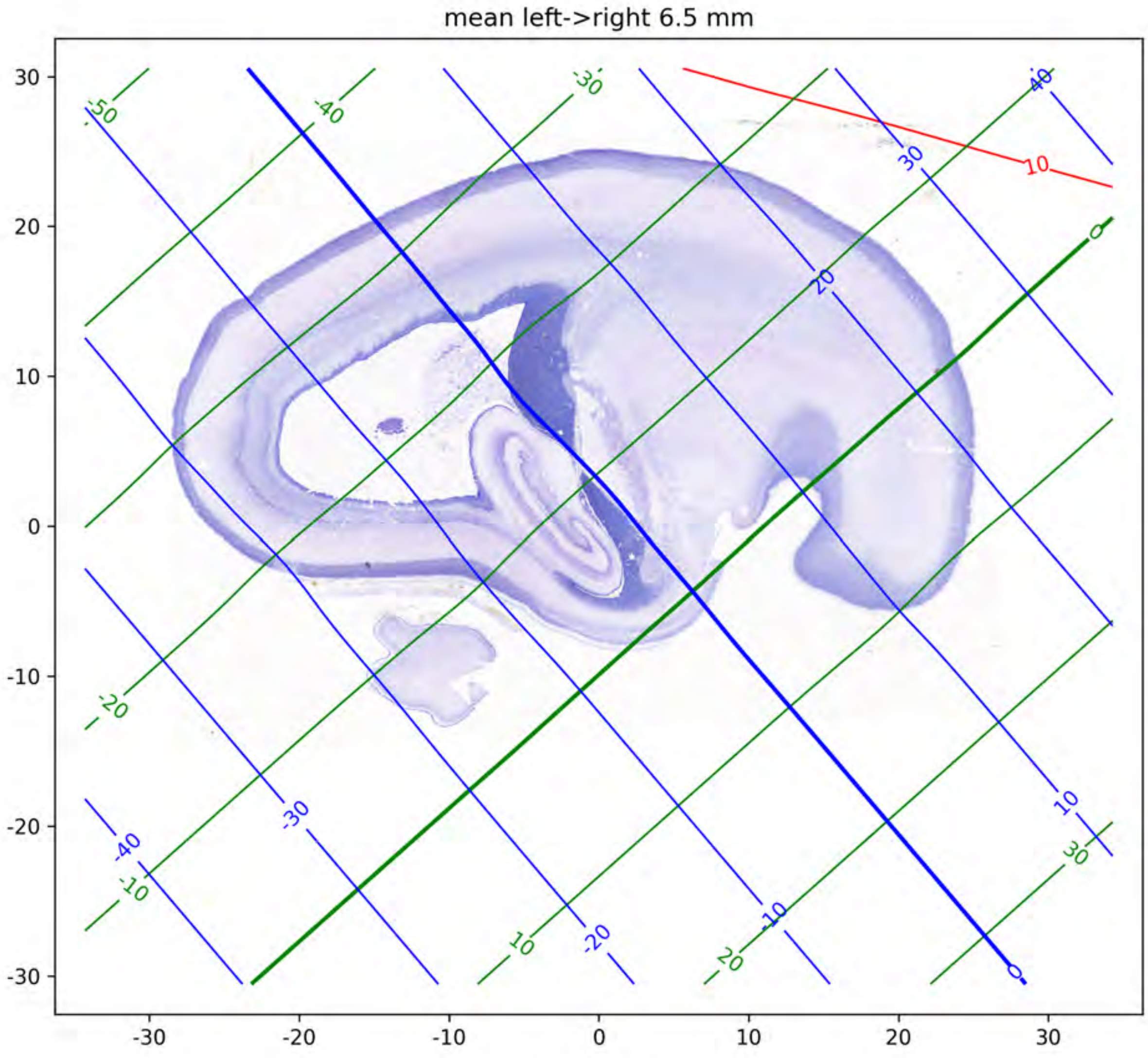

mean left->right 6.5 mm

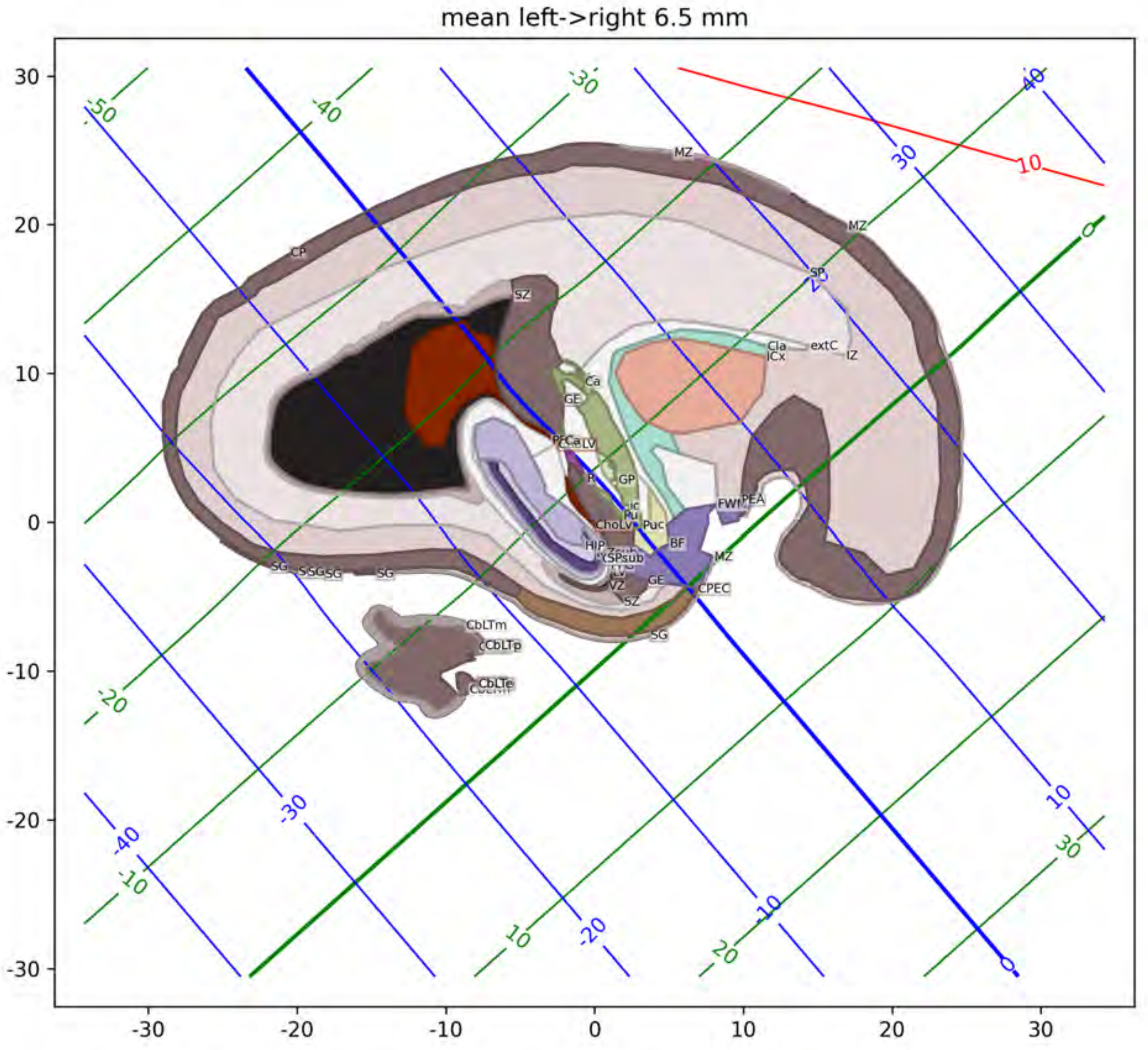
mean left->right 6.5 mm
30
20
10
0
-10
-20
-30
-30
-20
-10
0
10
20
30
MZ
CP
SZ
SP
Cla
ICx
extC
IZ
Ca
GE
LV
GP
Pu
ChoLv
Puc
BF
FWf
PEA
HIP
SPsub
MZ
VZ
GE
CPEC
SZ
SG
CbLTm
CbLTp
CbLTe
10

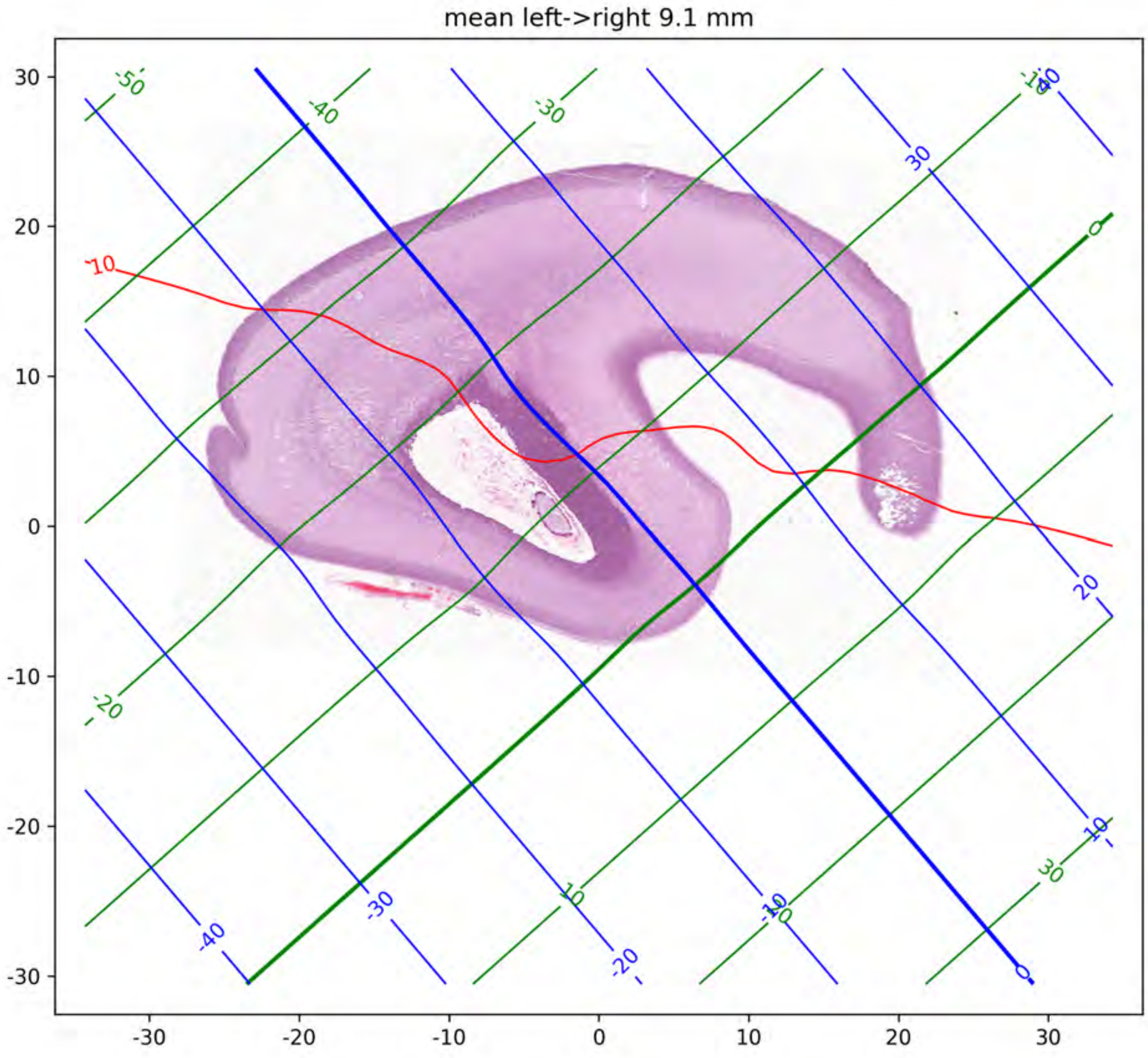

mean left->right 9.1 mm

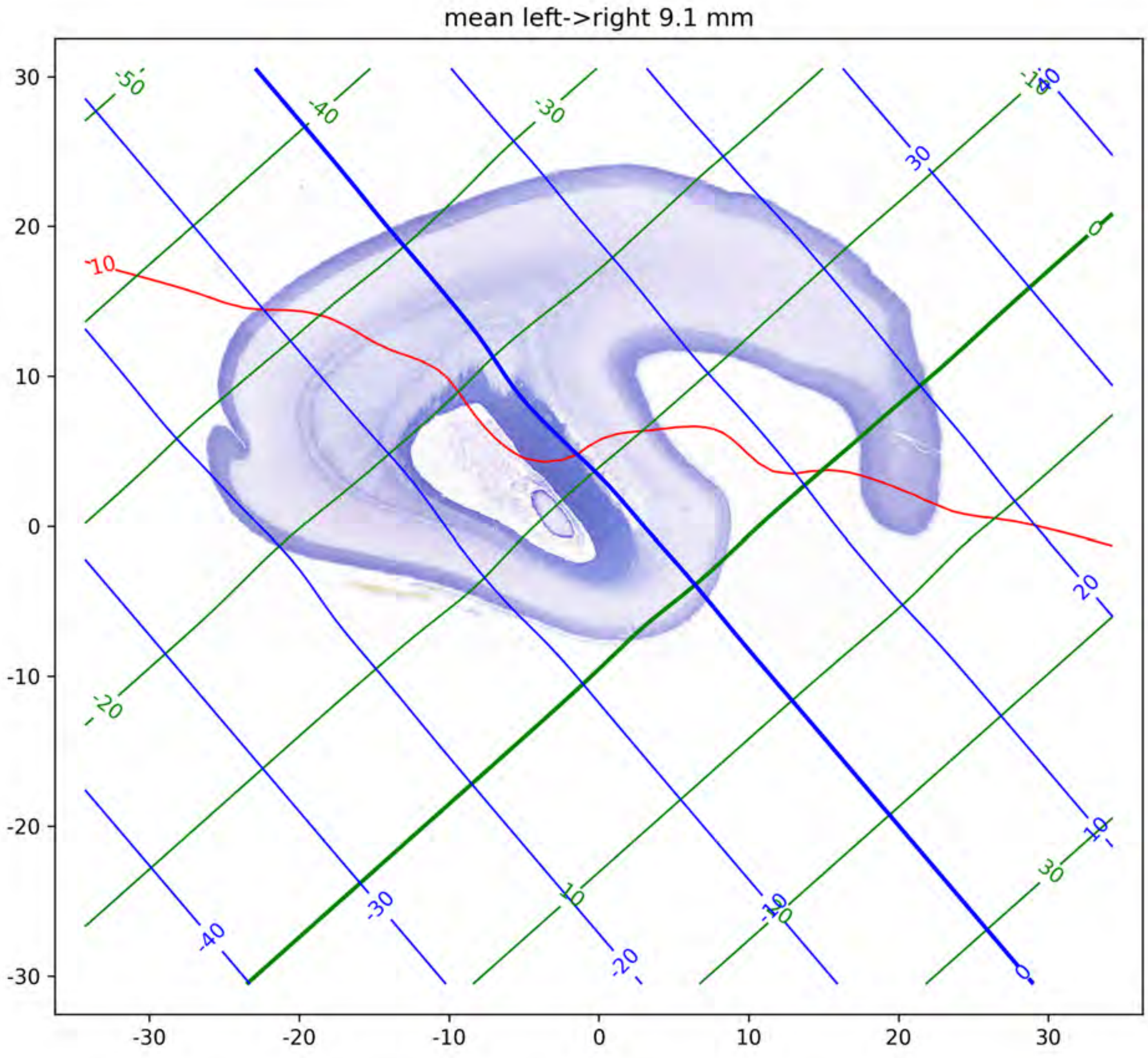
mean left->right 9.1 mm

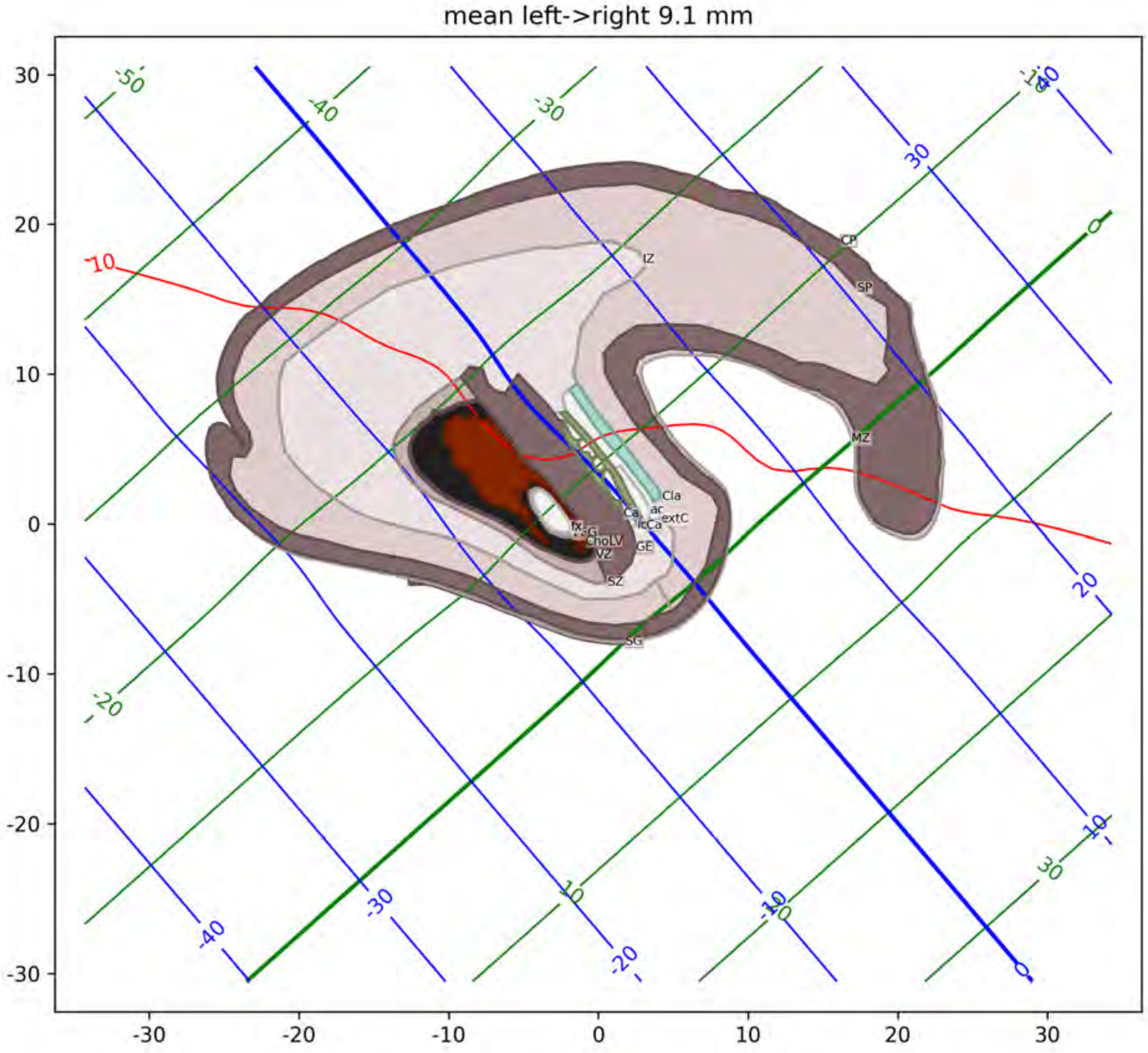
mean left->right 9.1 mm
IZ
CP
SP
MZ
Cla
ac
extC
Ca
icCa
GE
fx
ChoLV
VZ
SZ
SG

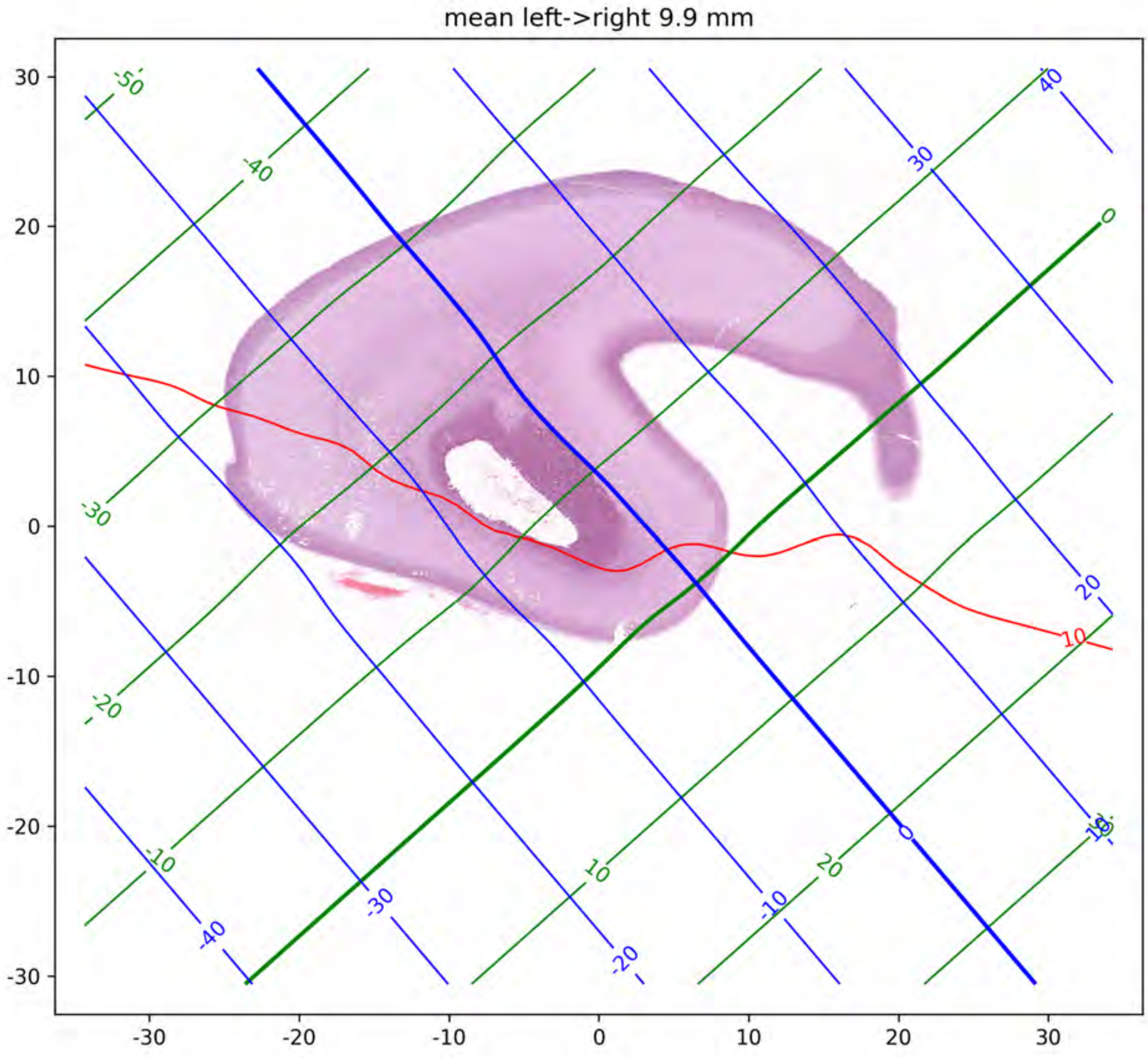
mean left->right 9.9 mm
-50
-40
-30
-20
-10
0
10
20
40
30
20
10
0
-10
-20
-30
-40
30
20
10
0
-10
-20
-30
-30
-20
-10
0
10
20
30

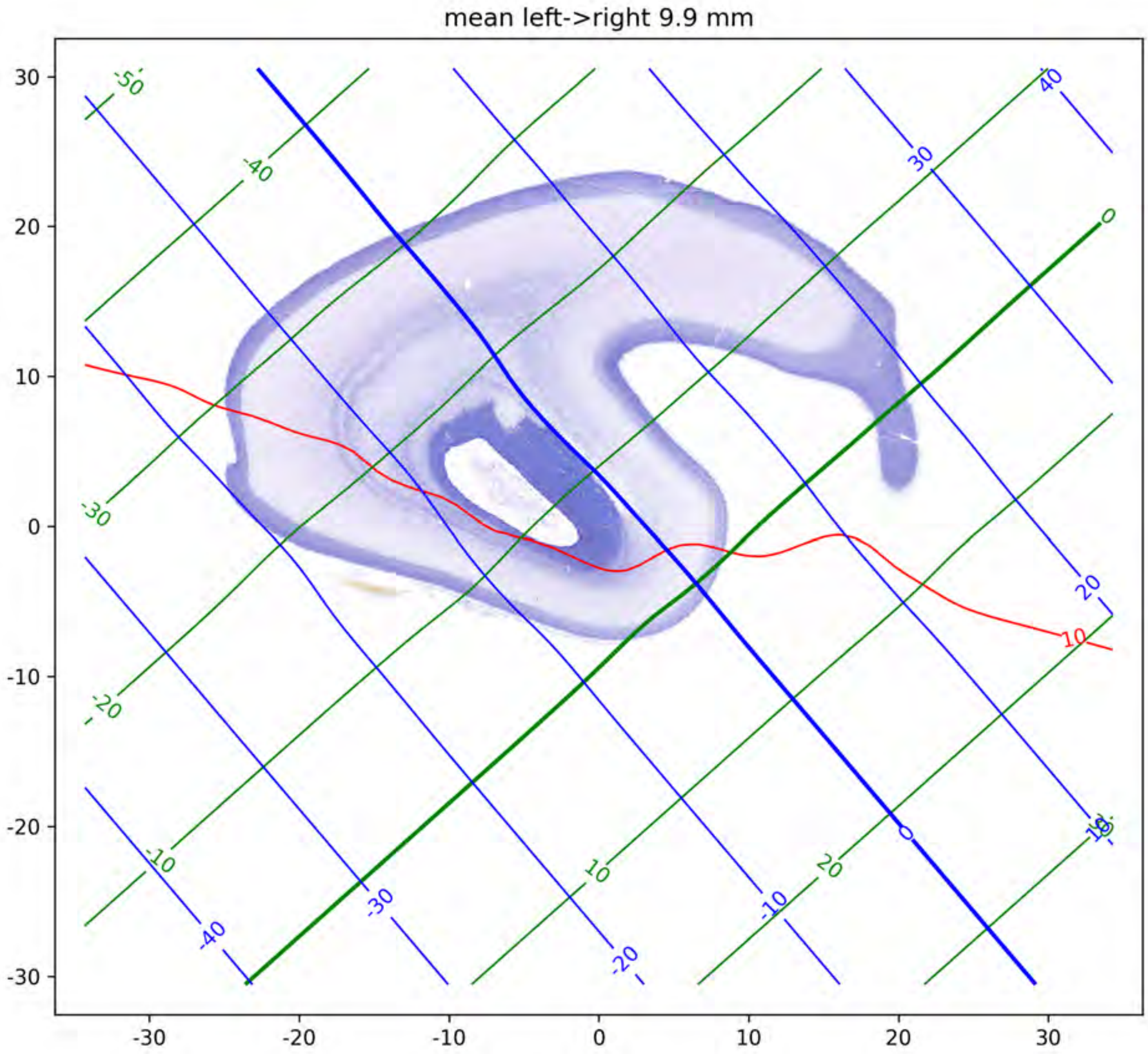
mean left->right 9.9 mm

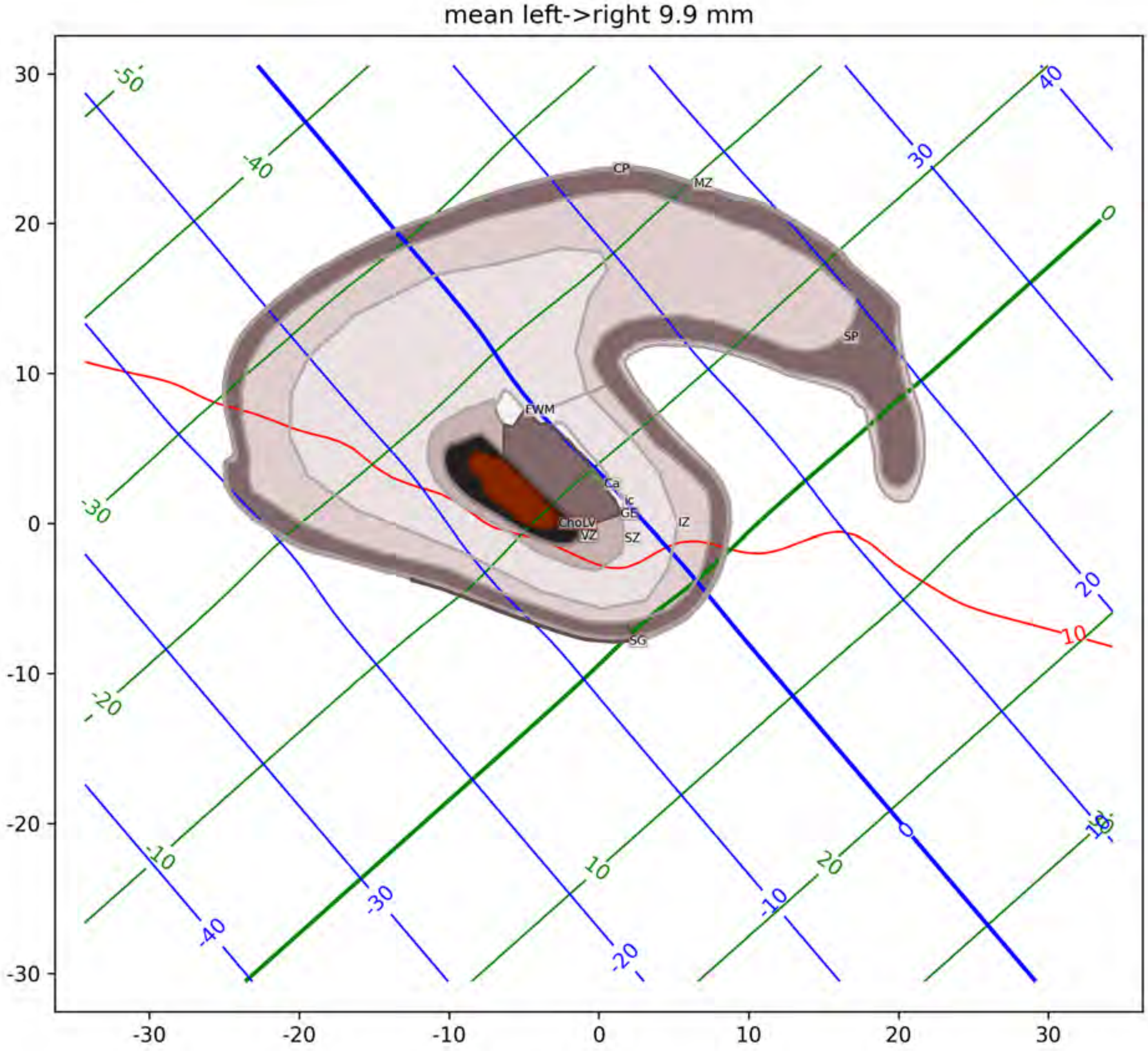

mean left->right 9.9 mm
CP
MZ
SP
FWM
Ca
Ic
GE
ChoLV
VZ
SZ
IZ
SG

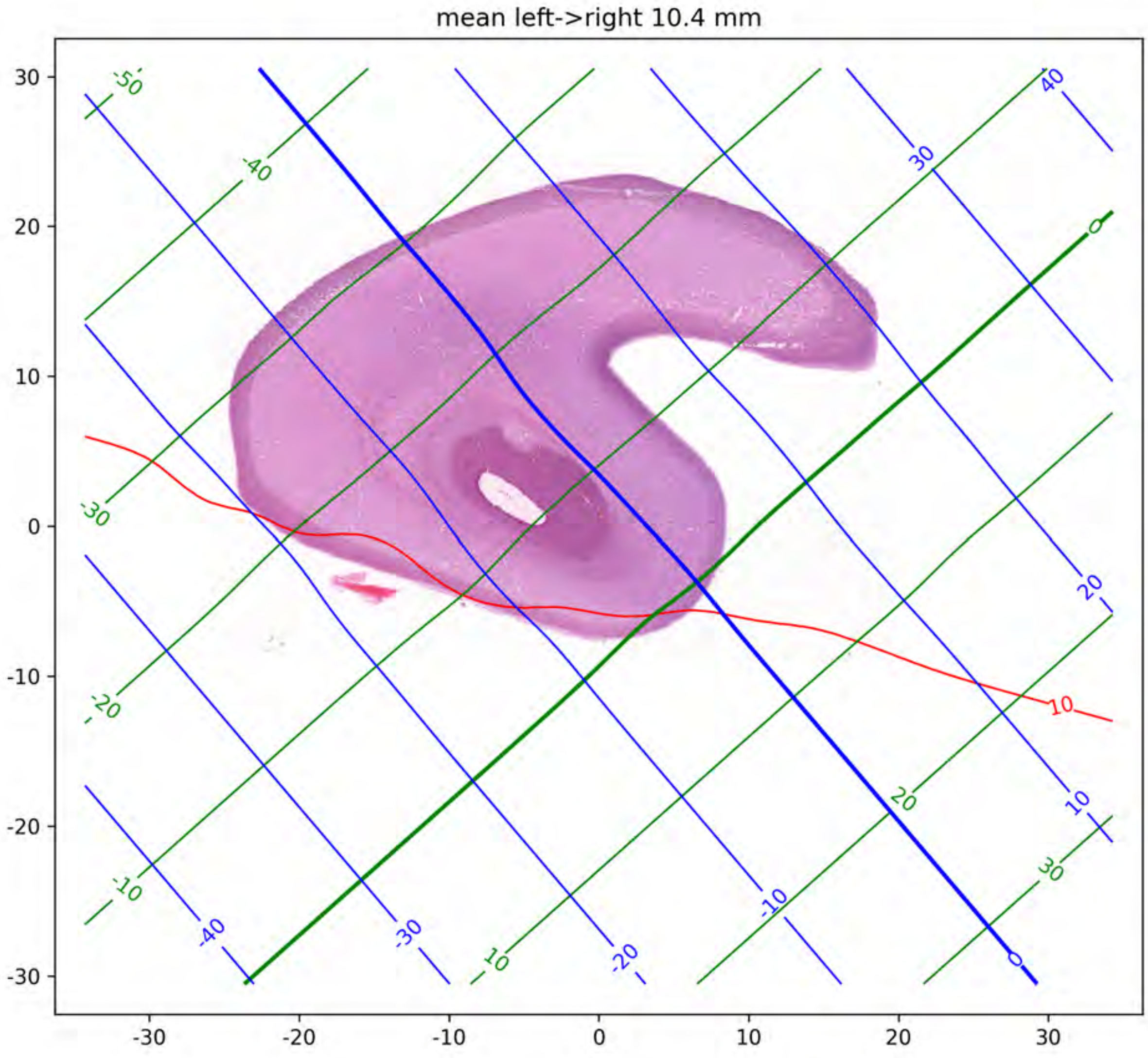

mean left->right 10.4 mm
30
20
10
0
-10
-20
-30
-30
-20
-10
0
10
20
30
-50
-40
-30
-20
-10
0
10
20
30
40
30
20
10
0
-10
-20
-30
-40
10

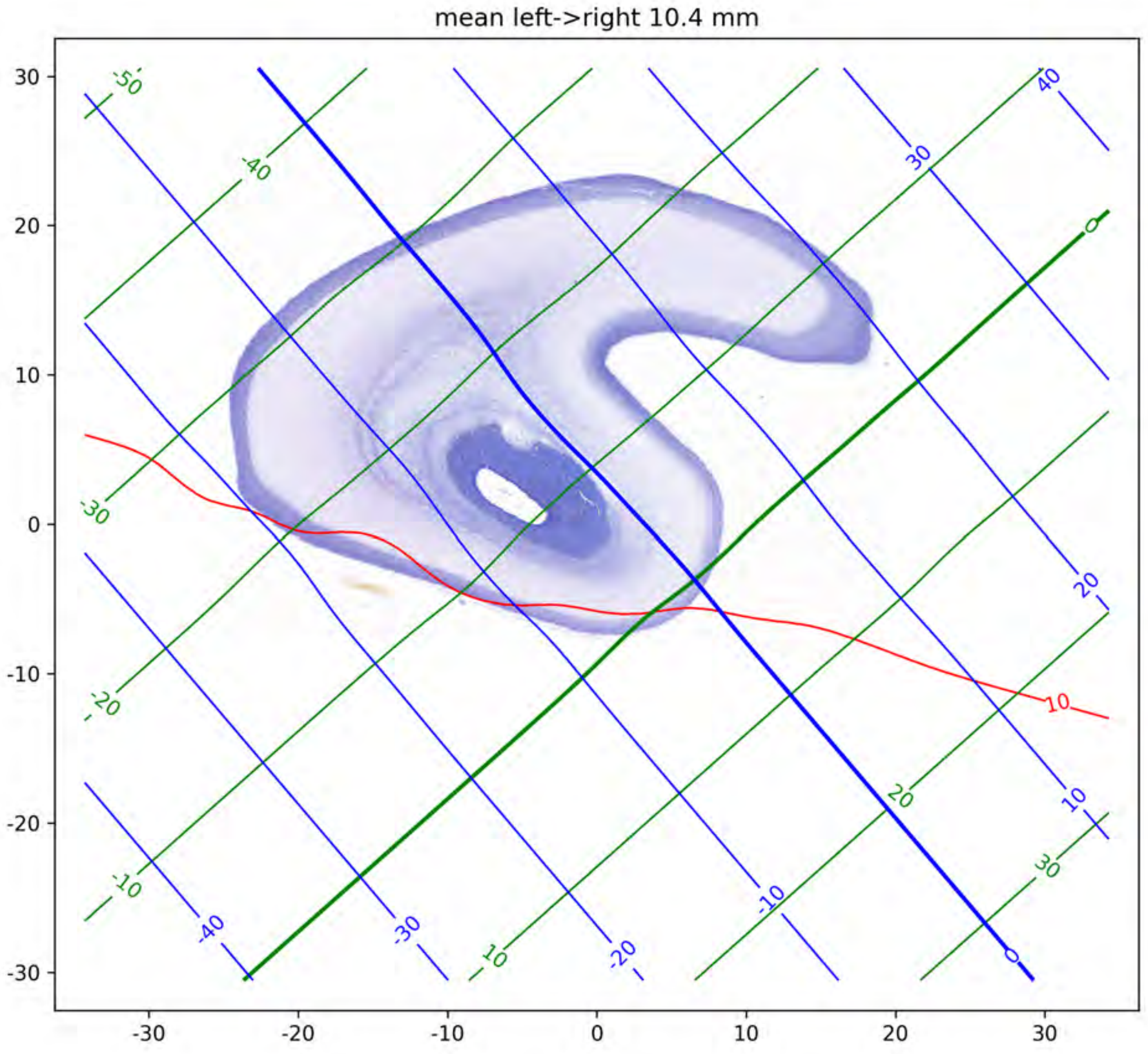
mean left->right 10.4 mm
30
20
10
0
-10
-20
-30
-30
-20
-10
0
10
20
30
-50
-40
-30
-20
-10
0
10
20
30
40
30
20
10
0
-10
-20
-30
-40
10

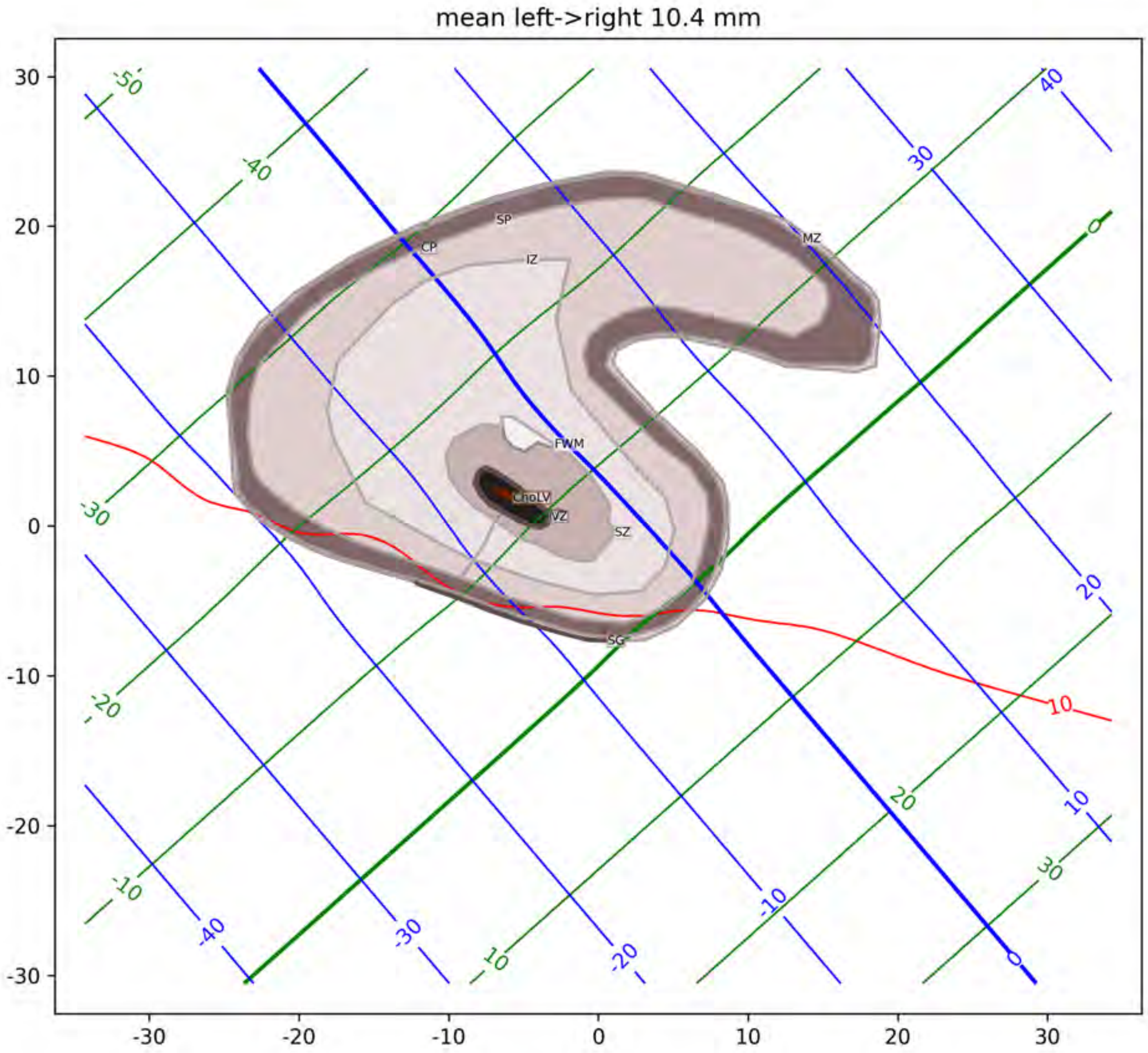

mean left->right 10.4 mm
SP
CP
IZ
MZ
FWM
ChoLV
VZ
SZ
SG

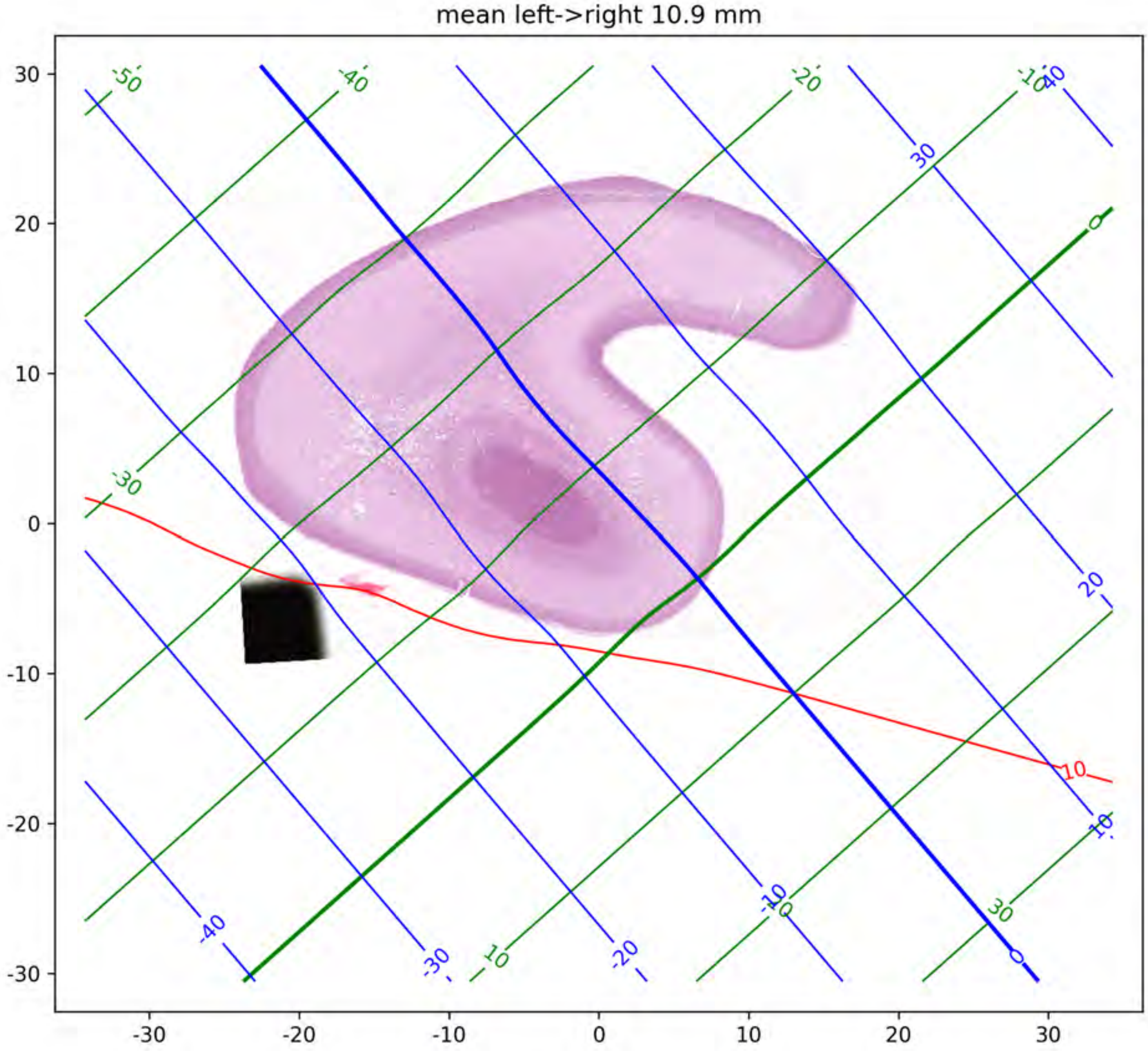
mean left->right 10.9 mm
30
20
10
0
-10
-20
-30
-30
-20
-10
0
10
20
30

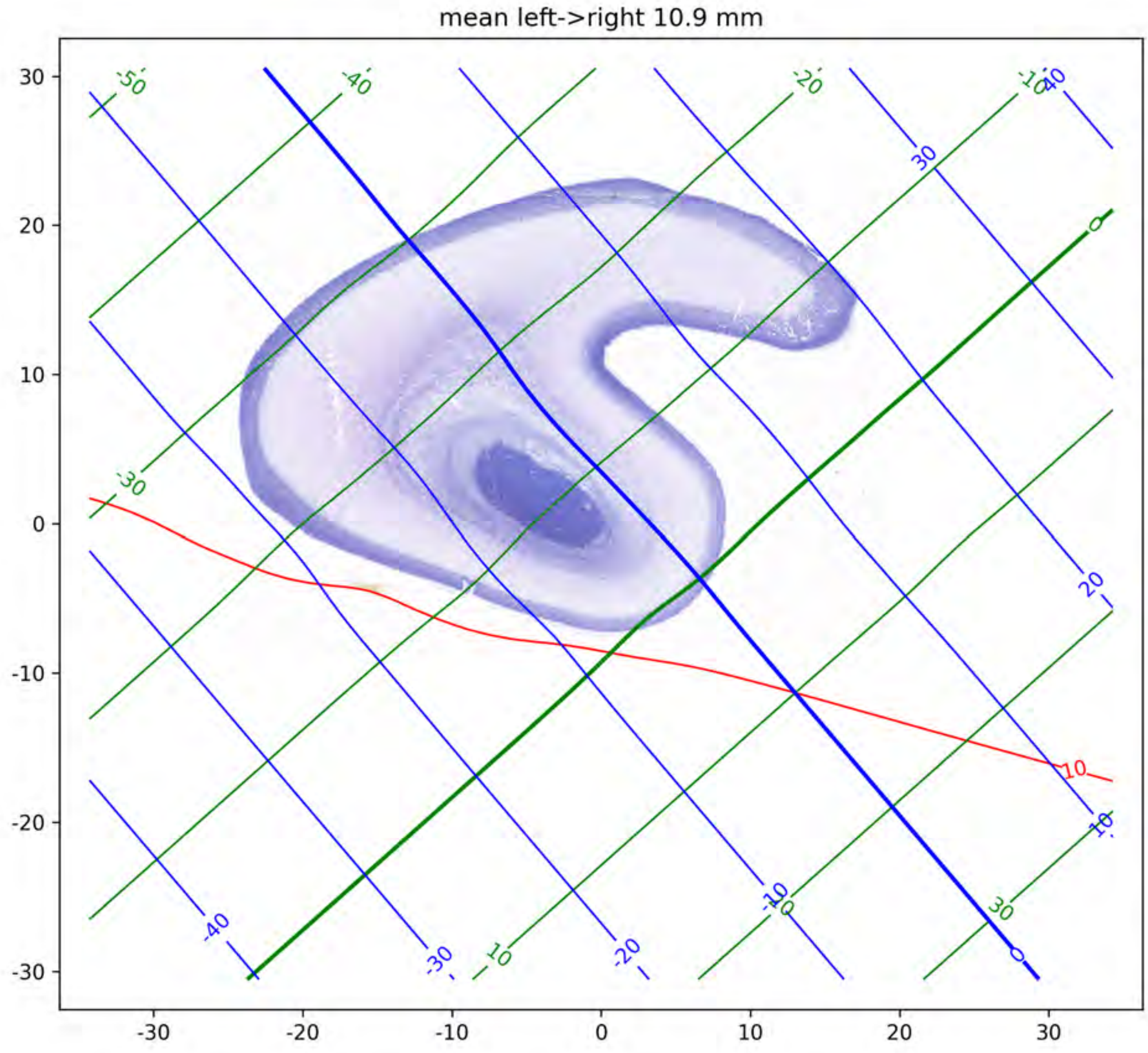

mean left->right 10.9 mm

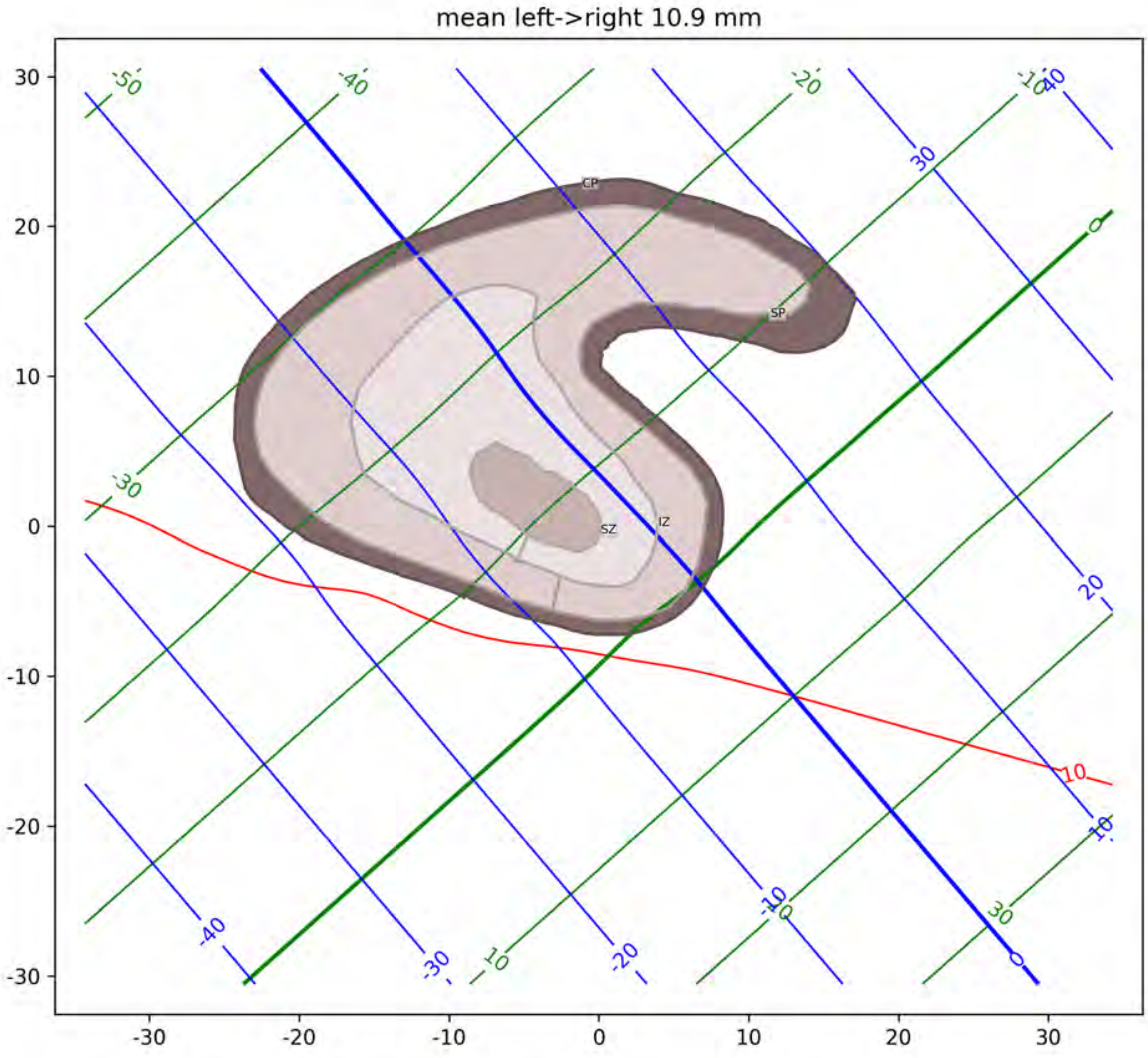
mean left->right 10.9 mm
CP
SP
SZ
IZ

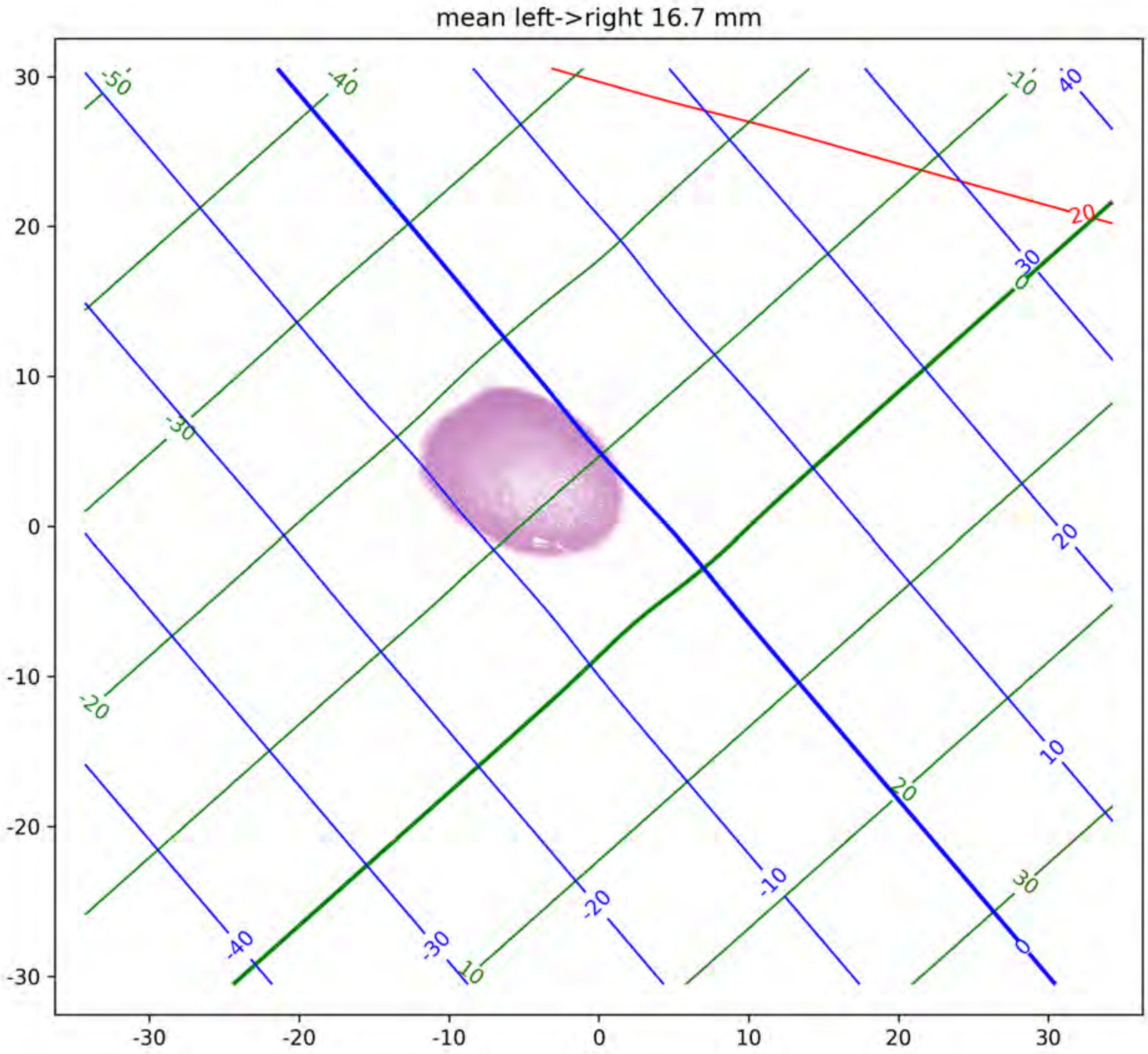
mean left->right 16.7 mm
30
20
10
0
-10
-20
-30
-30
-20
-10
0
10
20
30
-50
-40
-30
-20
-10
0
10
20
30
40
20
-40
-30
-20
-10
0
10
20
30

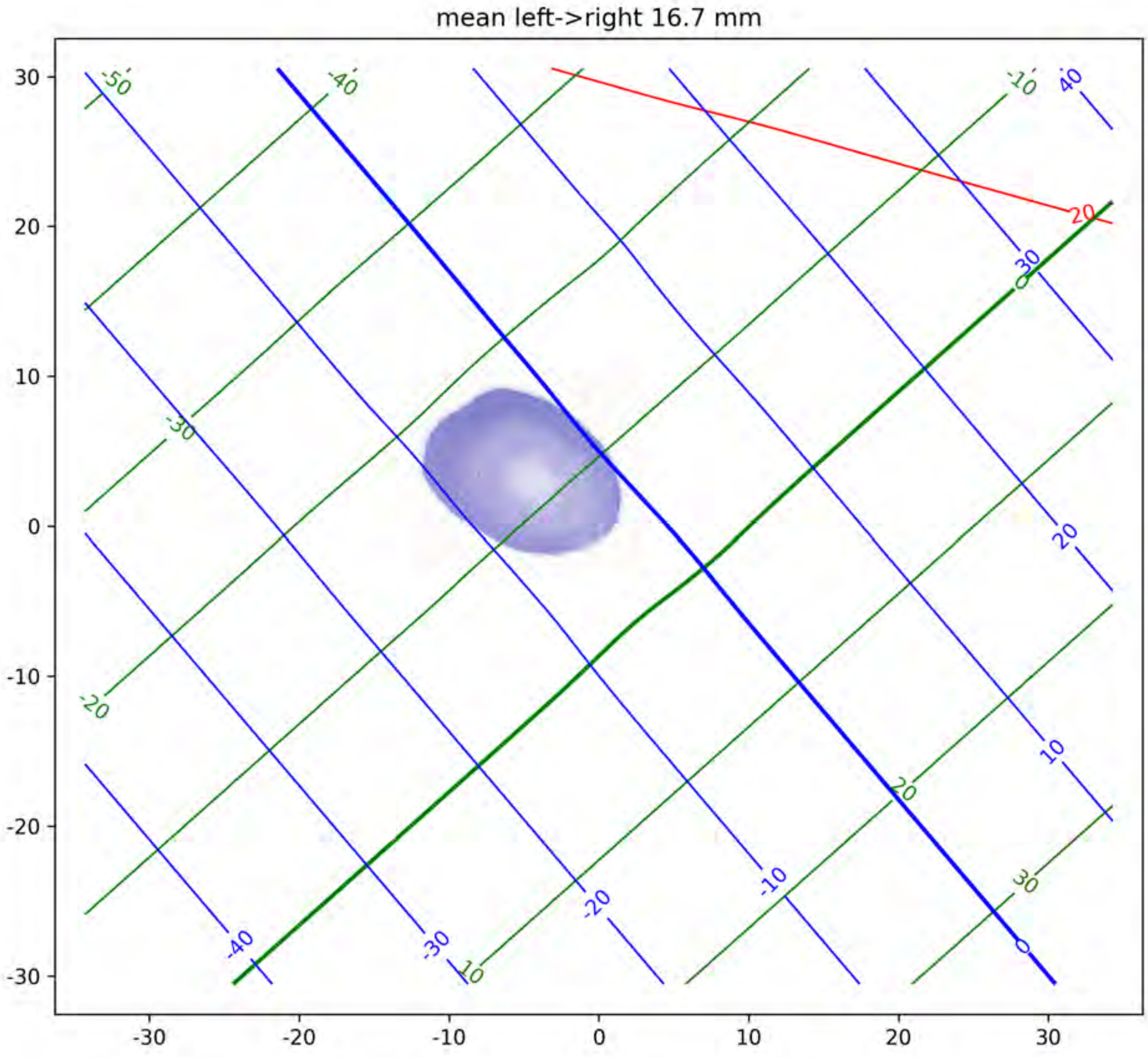
mean left->right 16.7 mm
30
20
10
0
-10
-20
-30
-30
-20
-10
0
10
20
30
-50
-40
-10
40
20
30
0
-30
20
-20
10
20
-10
30
-40
-30
10
-20
0

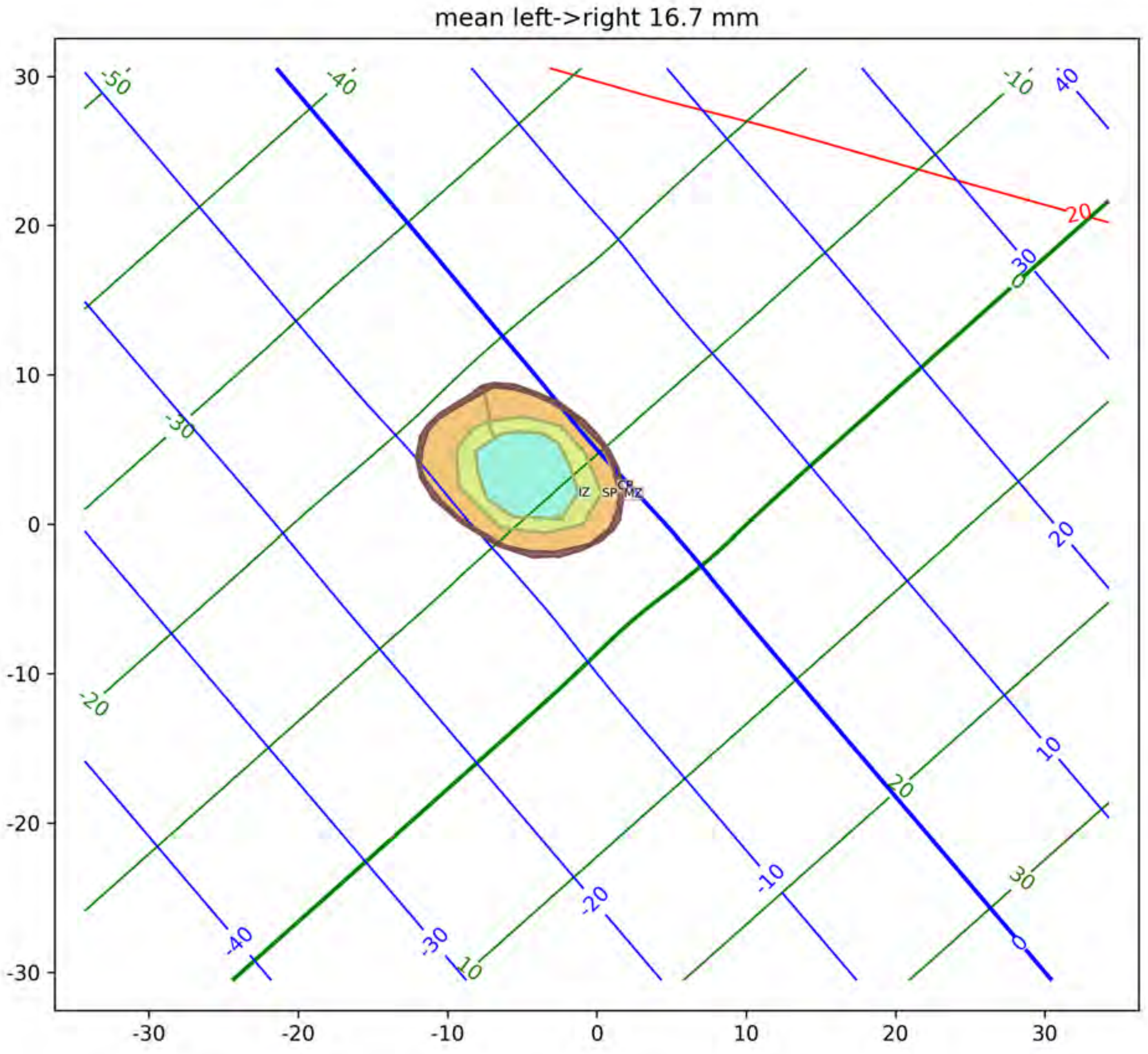

mean left->right 16.7 mm
IZ
SP
CP
MZ
