## Supplementary material for "A three-dimensional histological cell atlas of the developing human brain": GW14_MitoticFiguresPlates

### Mitotic figure reference atlas and coordinate system for a 14 gestational week fetal brain

December 15, 2024

Corresponding manuscript: Jai Jayakumar, et al. "A three-dimensional histological cell atlas of the developing human brain"

Demographic information: Specimen ID: S5/ FB40 AP (cms): 4.4 Bi temporal diameter (cms): 2.8 Brain weight (gms): 8.62 Age (GW): 14 Sex: M

This atlas includes serial section histology data, with neighoring sections aligned. Sections are 20um thick and collected in a series of three, alternating between Nissl, H&E, and various antibody stains (not shown), so that sections corresponding to a individual stain type are 60 um apart.

We have developed a standard RAS (x=Right, y=Anterior, z=Superior) coordinate system described as shown below. Please refer to figure 2 in the manuscript.

The data below shows this coordinate system as a curvilinear grid, with 10mm spaced isocontours, labeled in mm. Red lines correspond to the left-right axis with right positive. Green lines correspond to the posterior-anterior axis with anterior positive. Blue lines correspond to the inferior-superior axis with superior positive. The x=0, y=0, and z=0 lines are drawn thicker. Note that our sagittal histology sections are nearly perpendicular to the left right axis, and so red lines appear more curved.

Below we show H&E stained sections, here downsampled to 64 micron in plane resolution. Mitotic figures, detected as described in the manuscript, are illustrated as blue dots.

The horizontal and vertical axes of the image in each section also labeled with tick marks in mm, with 10 mm spacing to help with understanding scale.

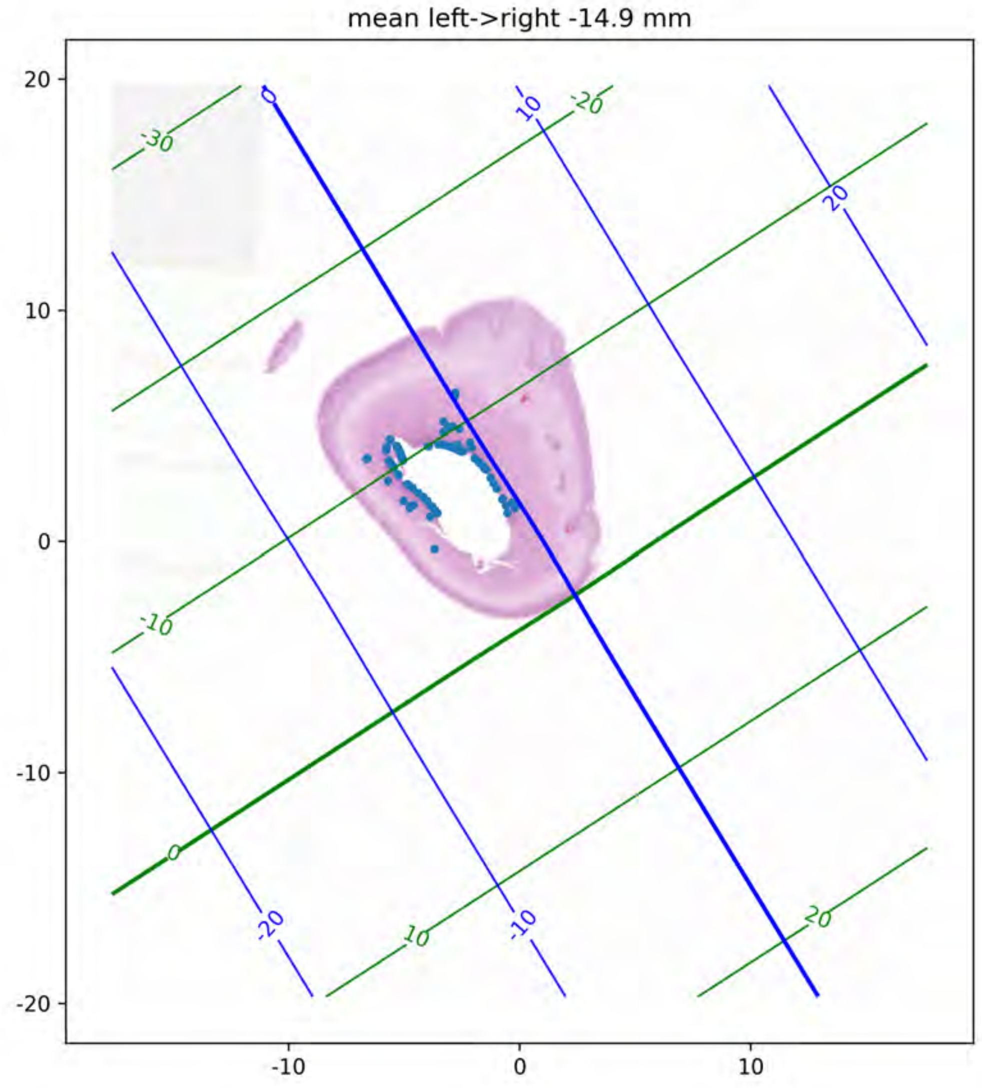

mean left->right -14.9 mm
20
10
0
-10
-20
-10
0
10
-30
-20
-10
0
10
20
0
10
20
-20
-10

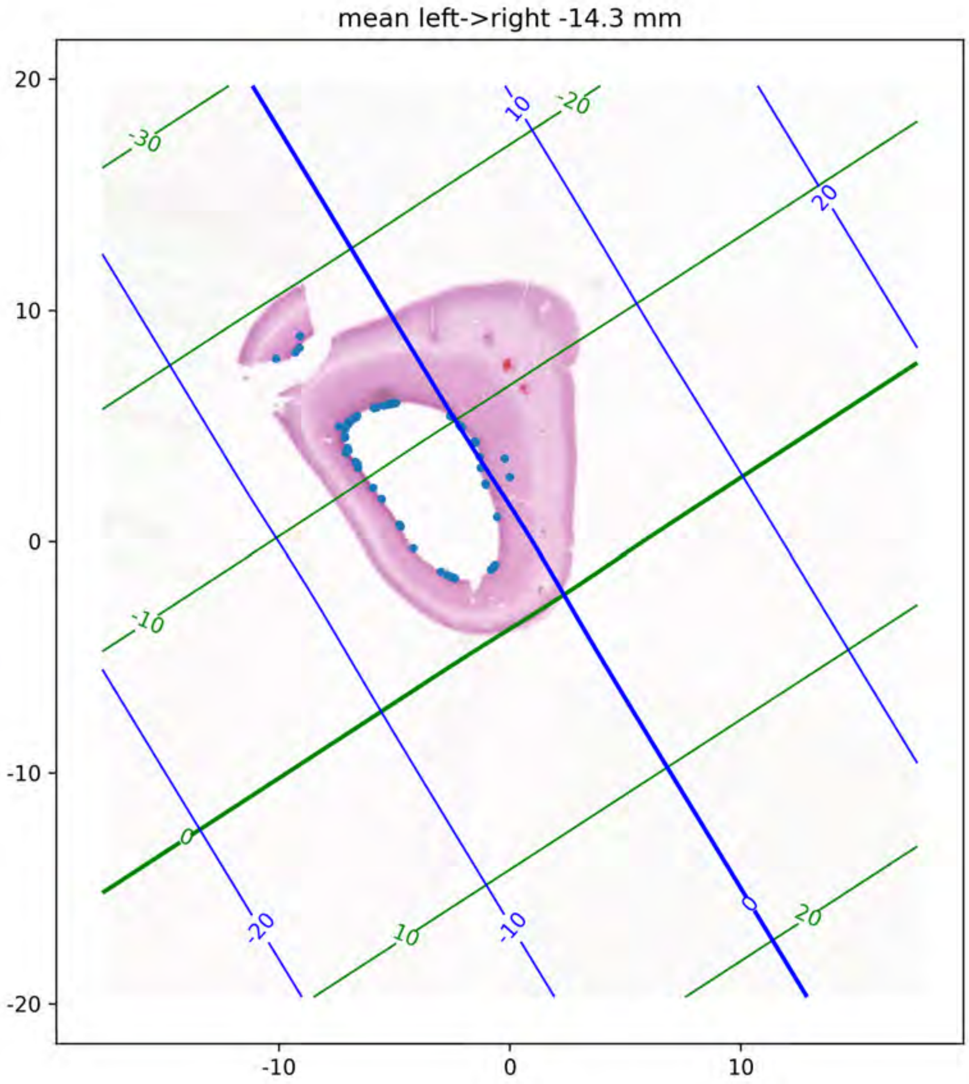
mean left->right -14.3 mm
20
10
0
-10
-20
-10
0
10
-30
-20
-10
0
10
20
10
20
0
-10
-20

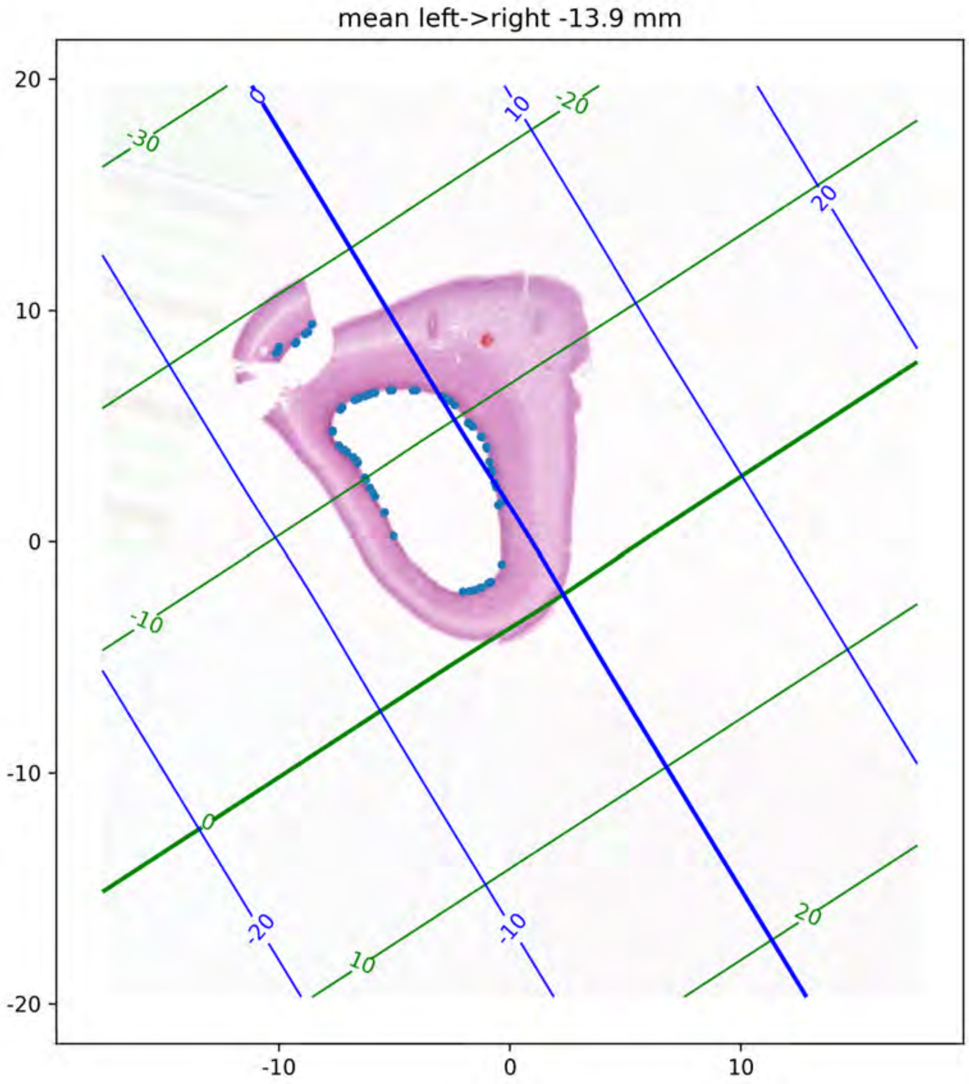
mean left->right -13.9 mm
20
10
0
-10
-20
-10
0
10
-30
-20
-10
0
10
20
0
10
20
-20
-10

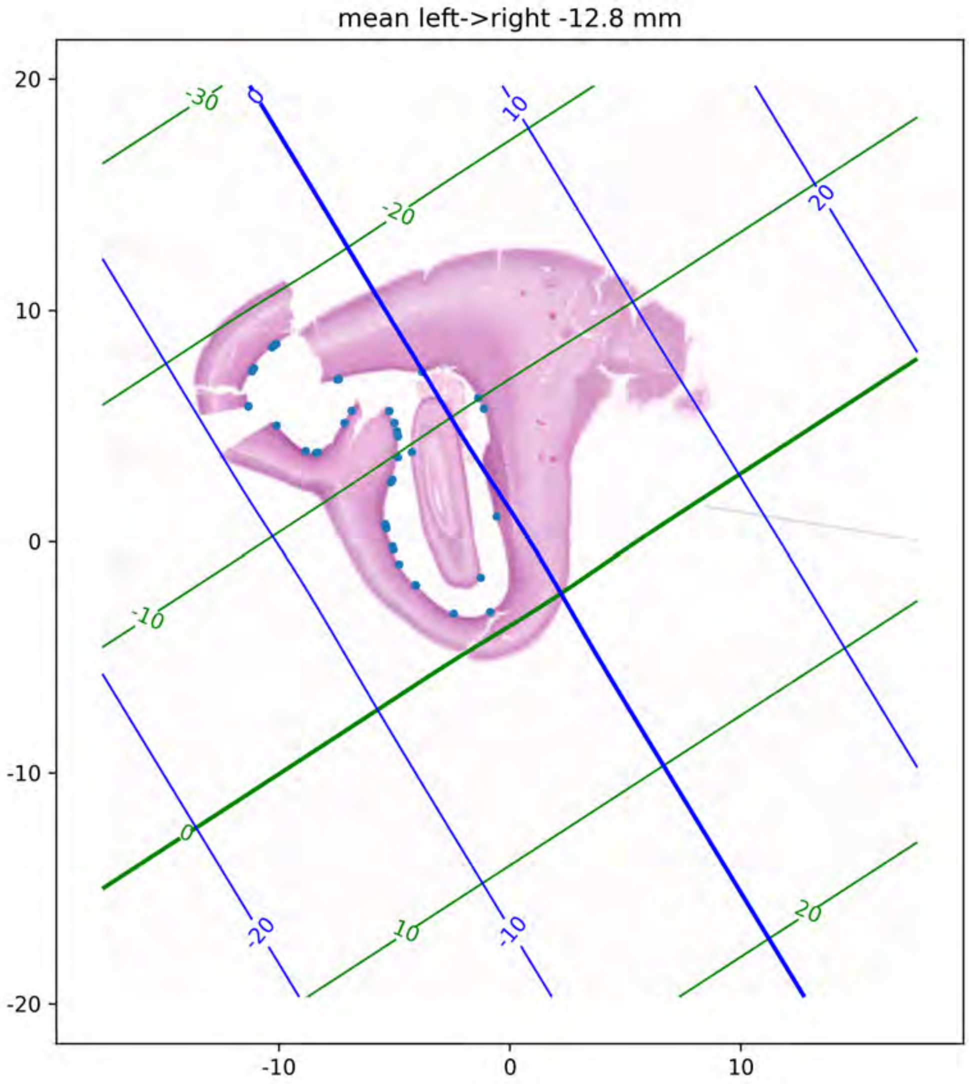

mean left->right -12.8 mm
20
10
0
-10
-20
-10
0
10
-30
-20
-10
0
10
20
0
10
20
-20
-10

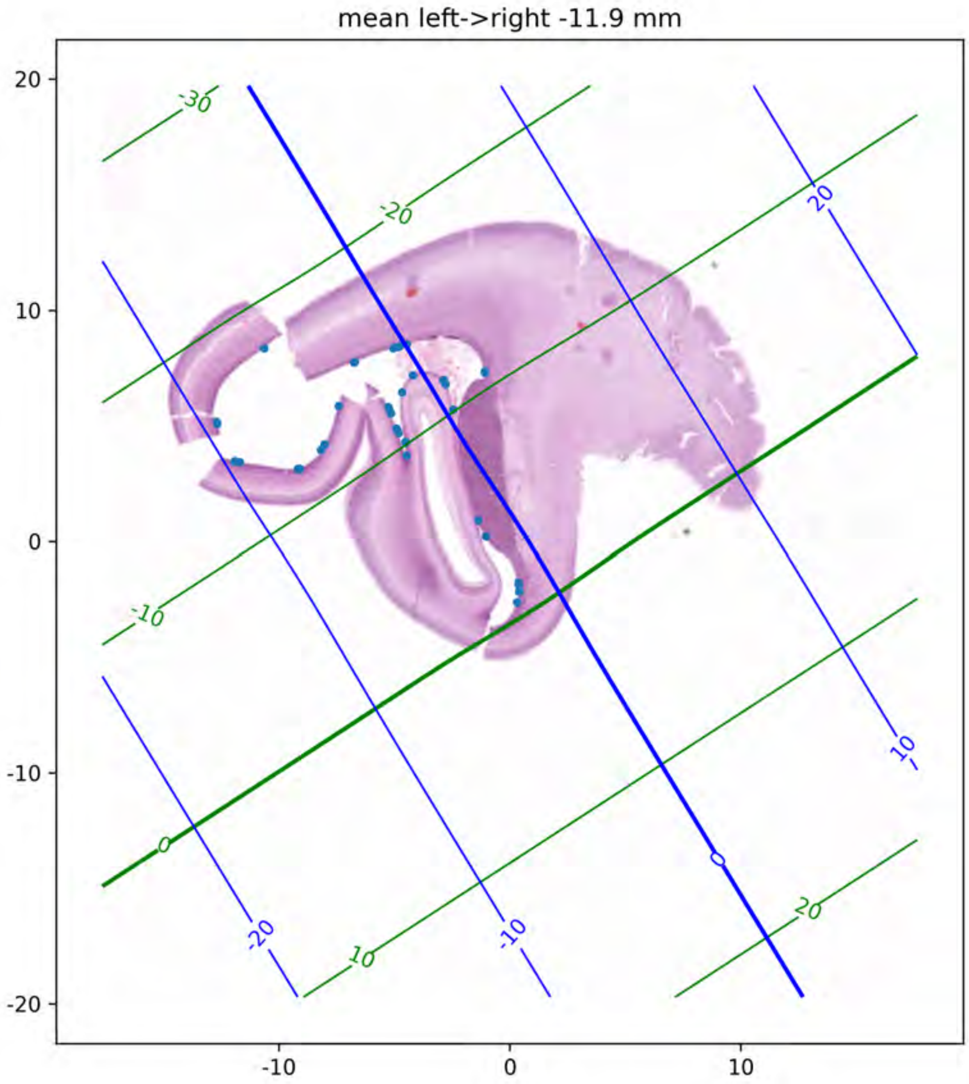
mean left->right -11.9 mm
20
10
0
-10
-20
-10
0
10
-30
-20
-10
0
10
20
20
10
0
-20
-10

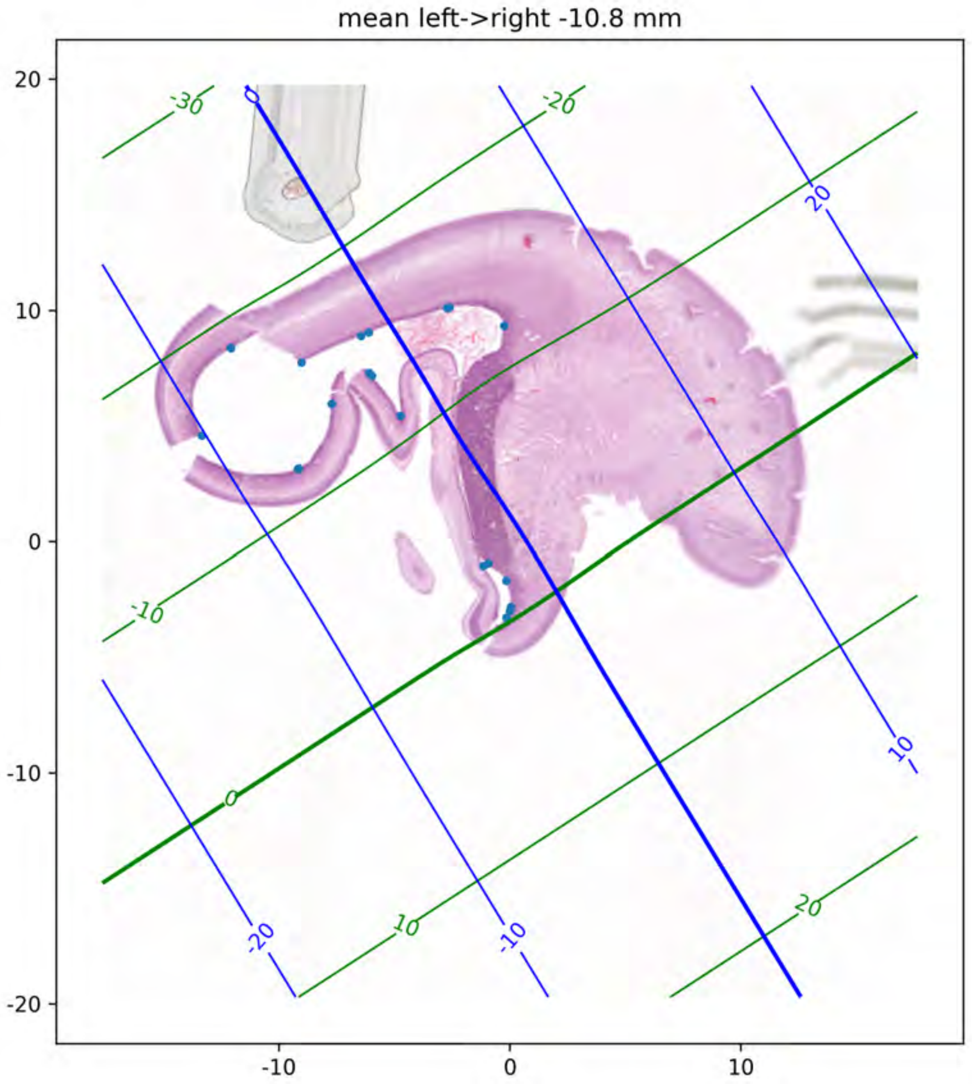
mean left->right -10.8 mm
20
10
0
-10
-20
-10
0
10
-30
-20
-10
0
10
20
0
20
10
-20
-10

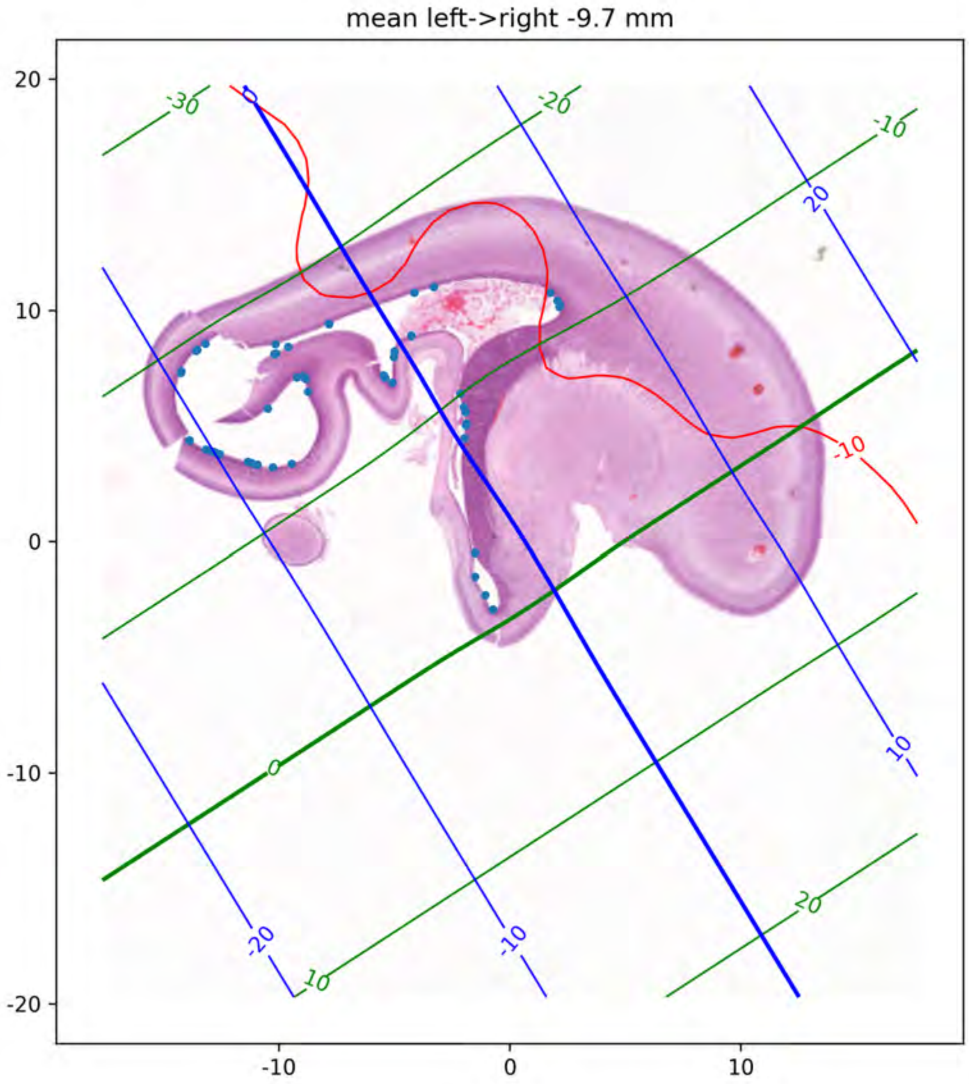

mean left->right -9.7 mm
20
10
0
-10
-20
-10
0
10
-30
-20
-10
0
10
20
20
10
-20
-10
-10

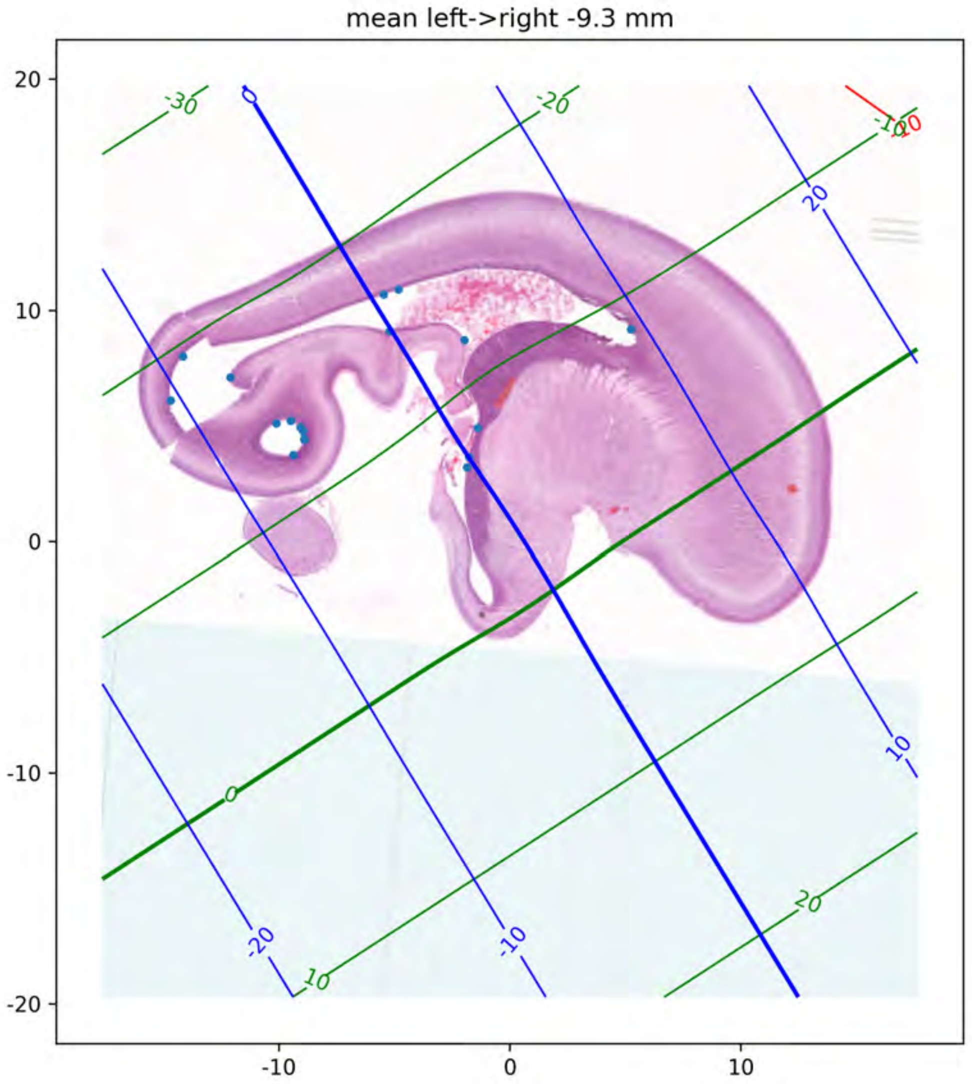

mean left->right -9.3 mm
20
10
0
-10
-20
-10
0
10
-30
-20
-10
0
10
20
-20
-10
10
20
-10

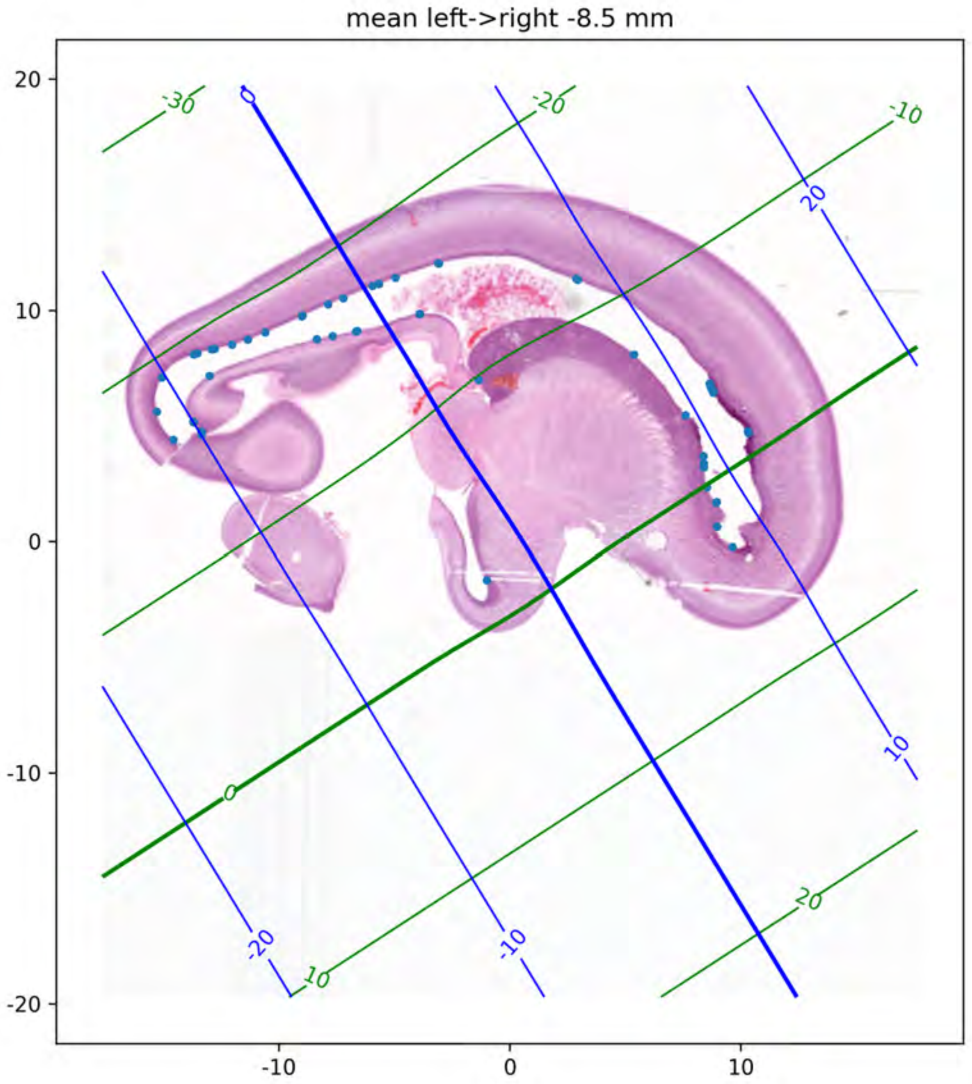

mean left->right -8.5 mm
20
10
0
-10
-20
-10
0
10
-30
-20
-10
0
10
20
0
20
10
-20
-10

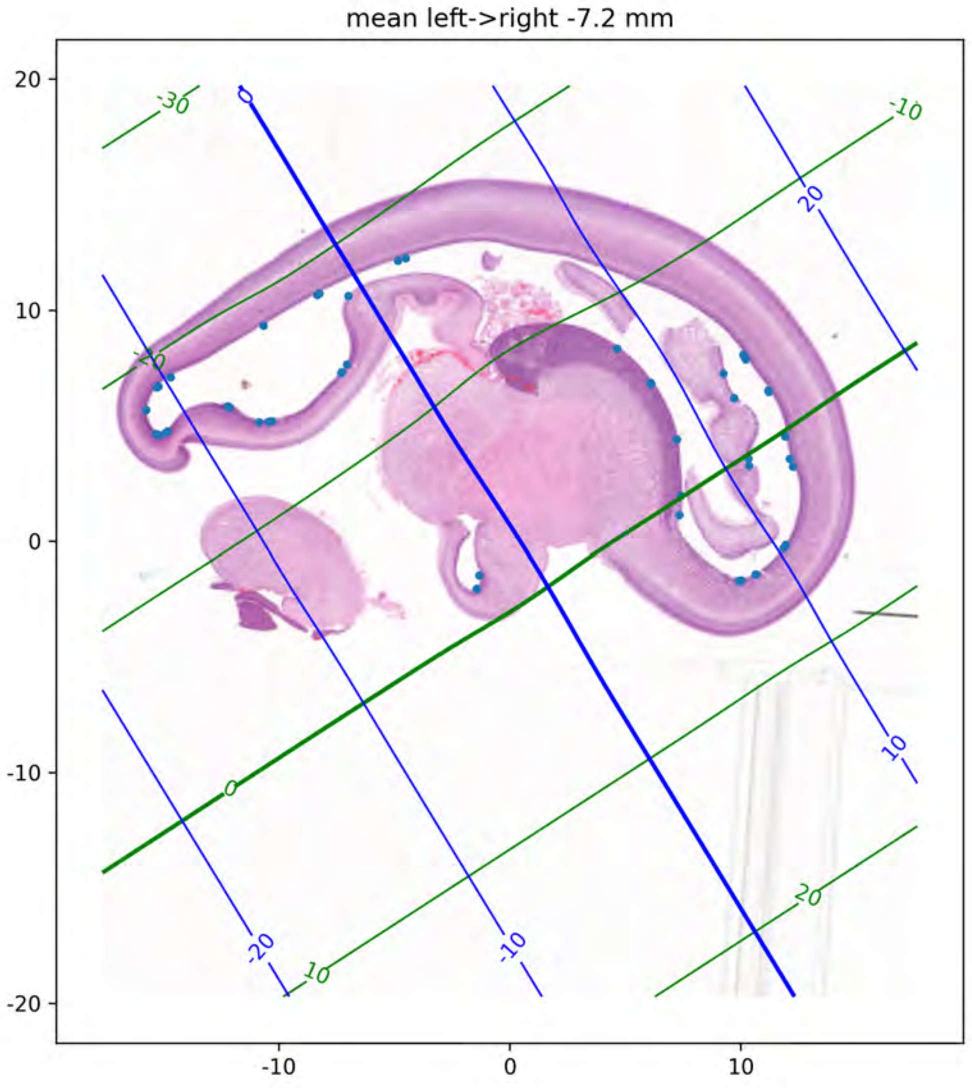
mean left->right -7.2 mm
20
10
0
-10
-20
-10
0
10
-30
-20
-10
0
10
20
0
20
10
-20
-10

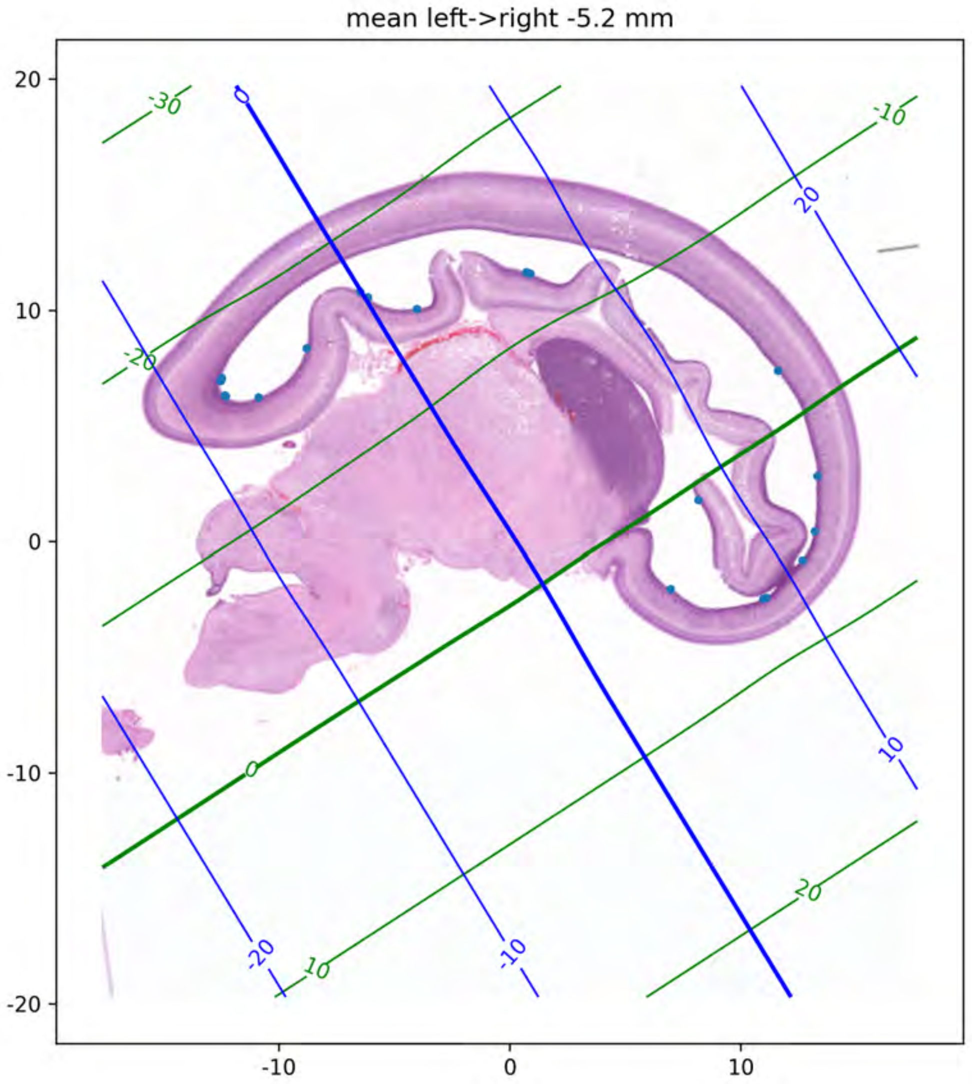
mean left->right -5.2 mm
20
10
0
-10
-20
-10
0
10
-30
-20
0
10
20
-10
0
20
10
-20
-10

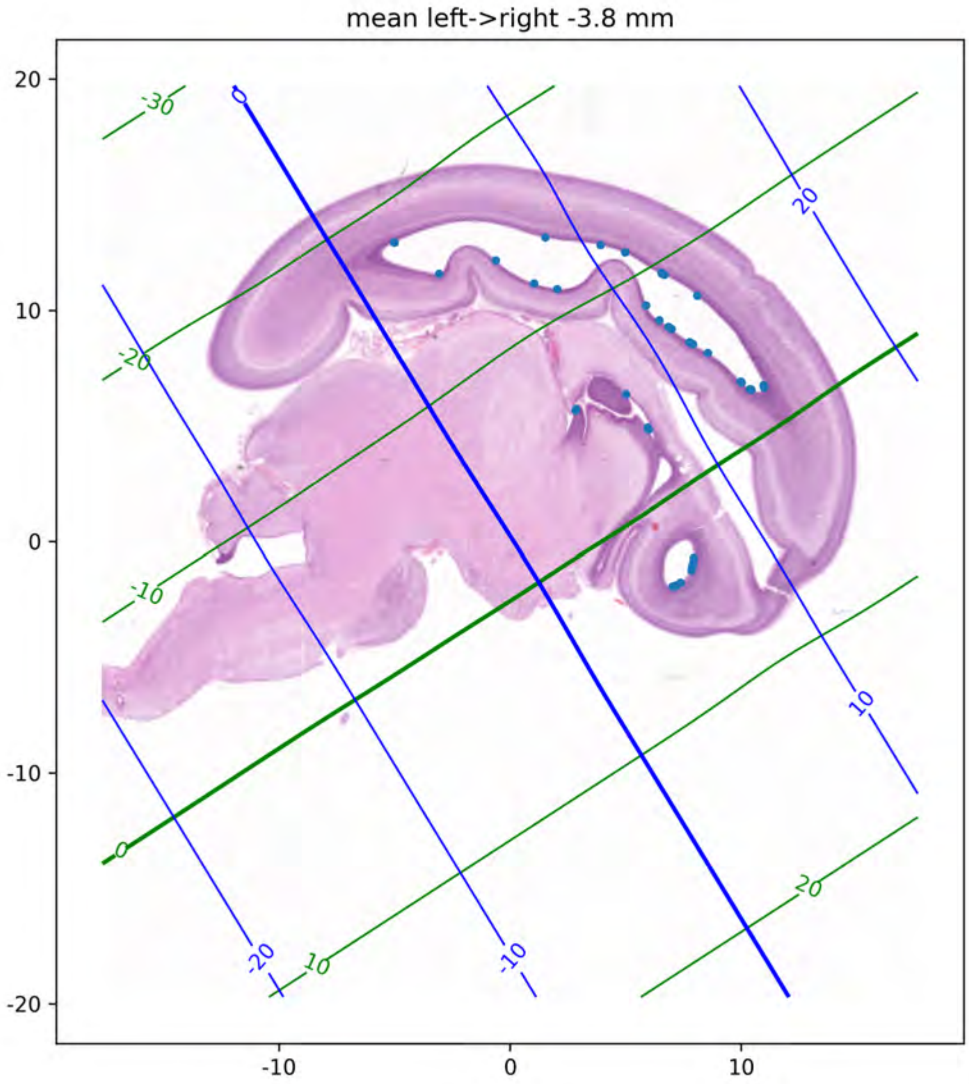

mean left->right -3.8 mm
20
10
0
-10
-20
-10
0
10
-30
-20
-10
0
10
20
0
20
10
-20
-10

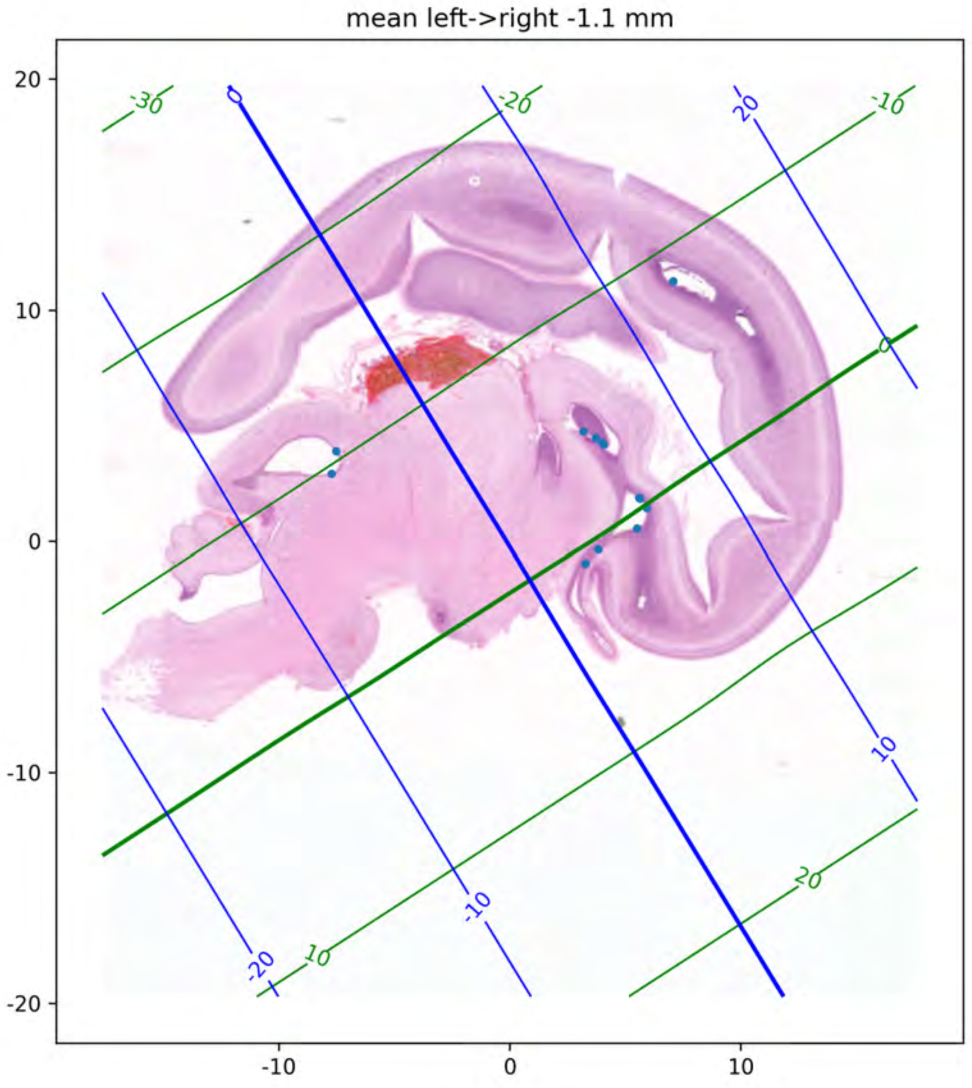
mean left->right -1.1 mm
20
10
0
-10
-20
-10
0
10
-30
-20
-10
0
10
20
0
-20
-10
10
20

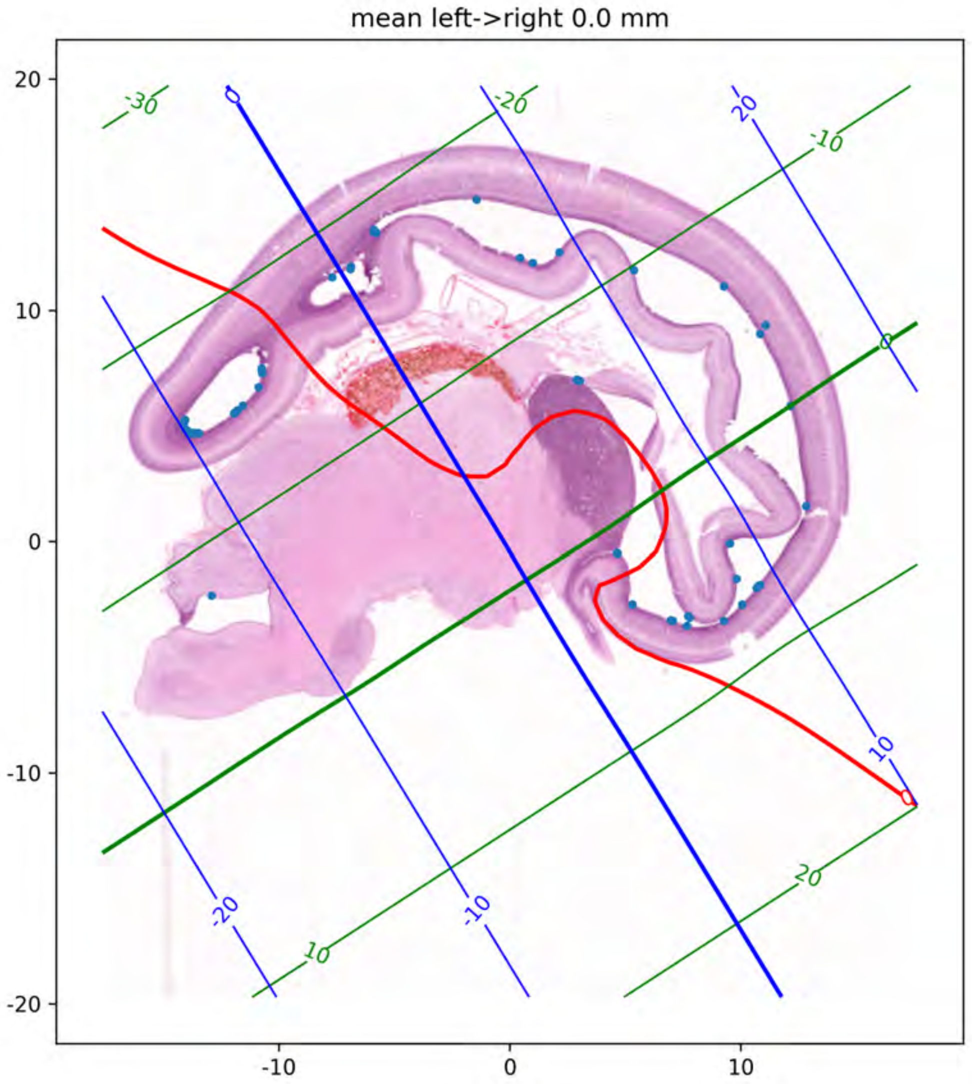

mean left->right 0.0 mm
20
10
0
-10
-20
-10
0
10
-30
-20
-10
0
10
20
0
-20
20
10
-20
-10
0

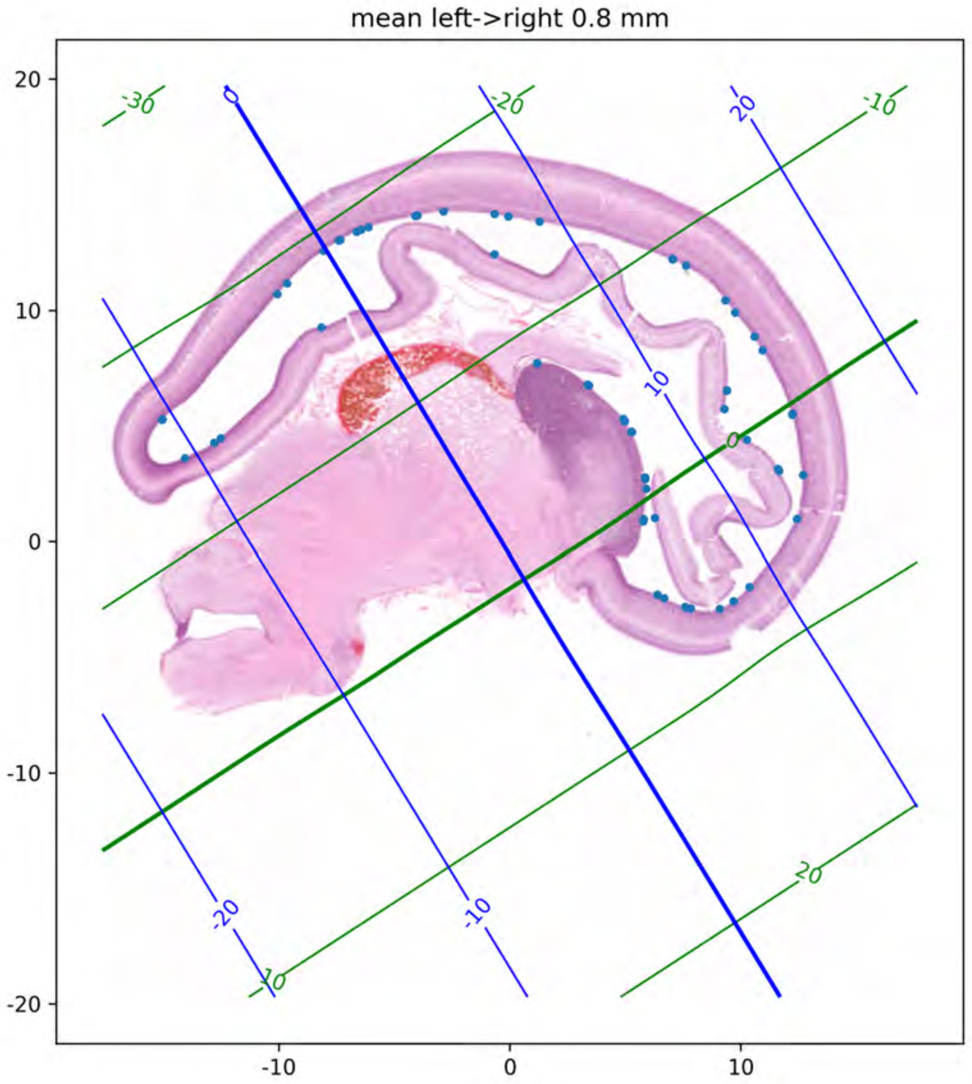
mean left->right 0.8 mm
20
10
0
-10
-20
-10
0
10
-30
-20
-10
0
10
20
0
10
20
-20
-10

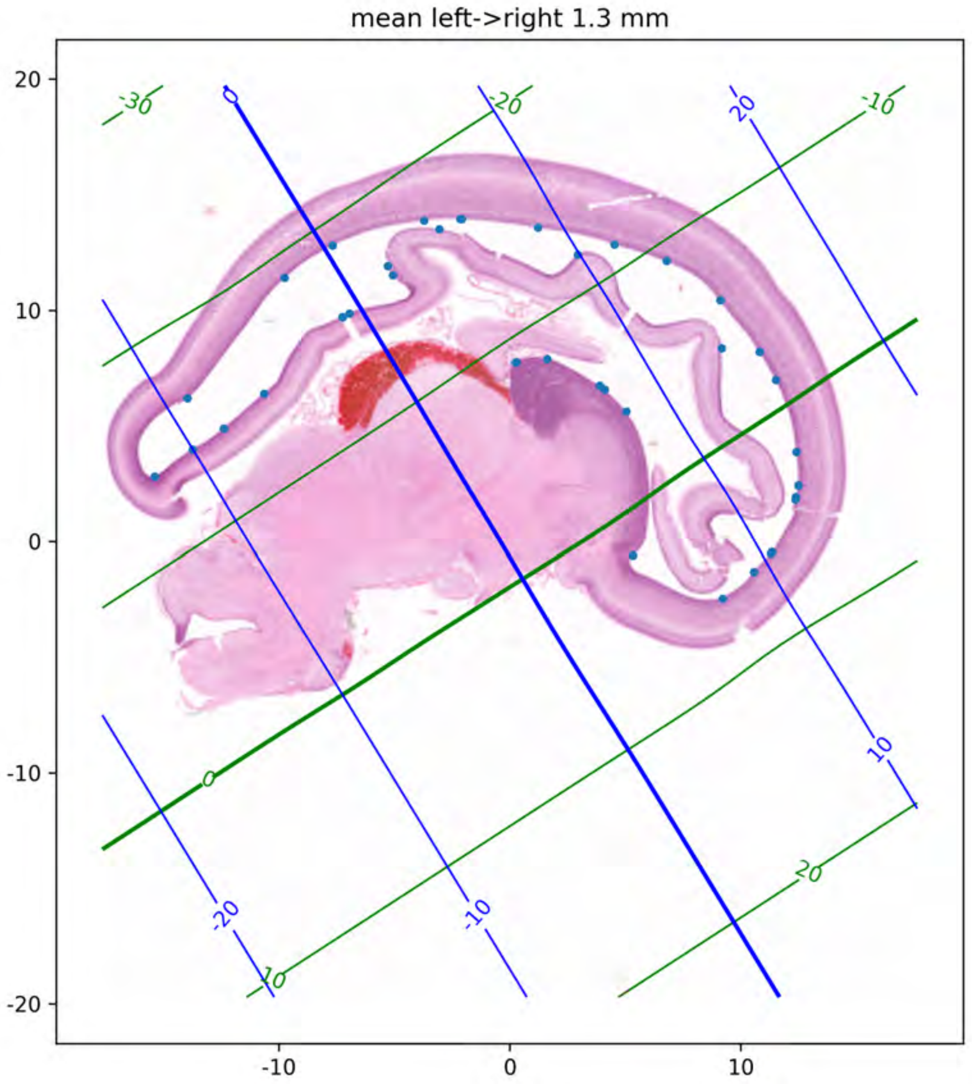
mean left->right 1.3 mm
20
10
0
-10
-20
-10
0
10
-30
-20
-10
0
10
20
0
20
10
-20
-10

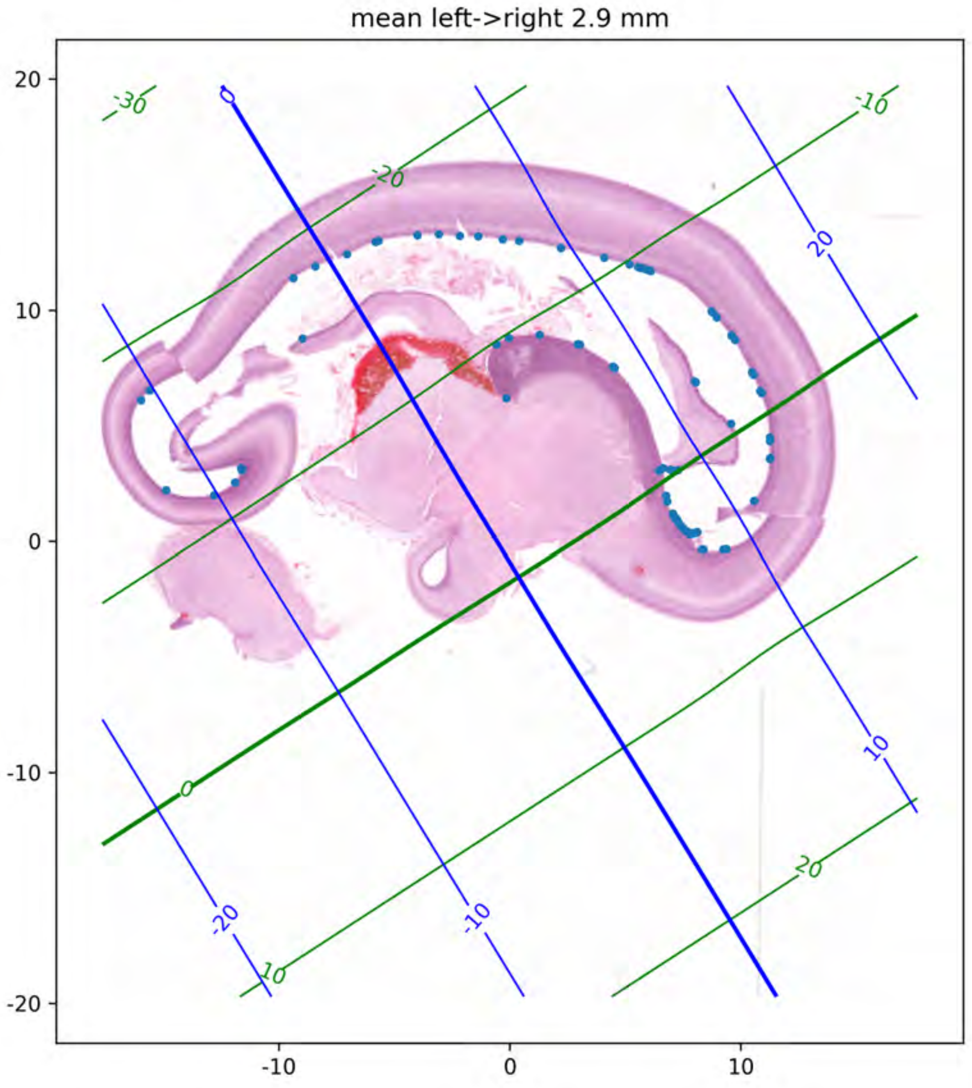
mean left->right 2.9 mm
20
10
0
-10
-20
-10
0
10
-30
-20
-10
0
10
20
0
20
10
-20
-10

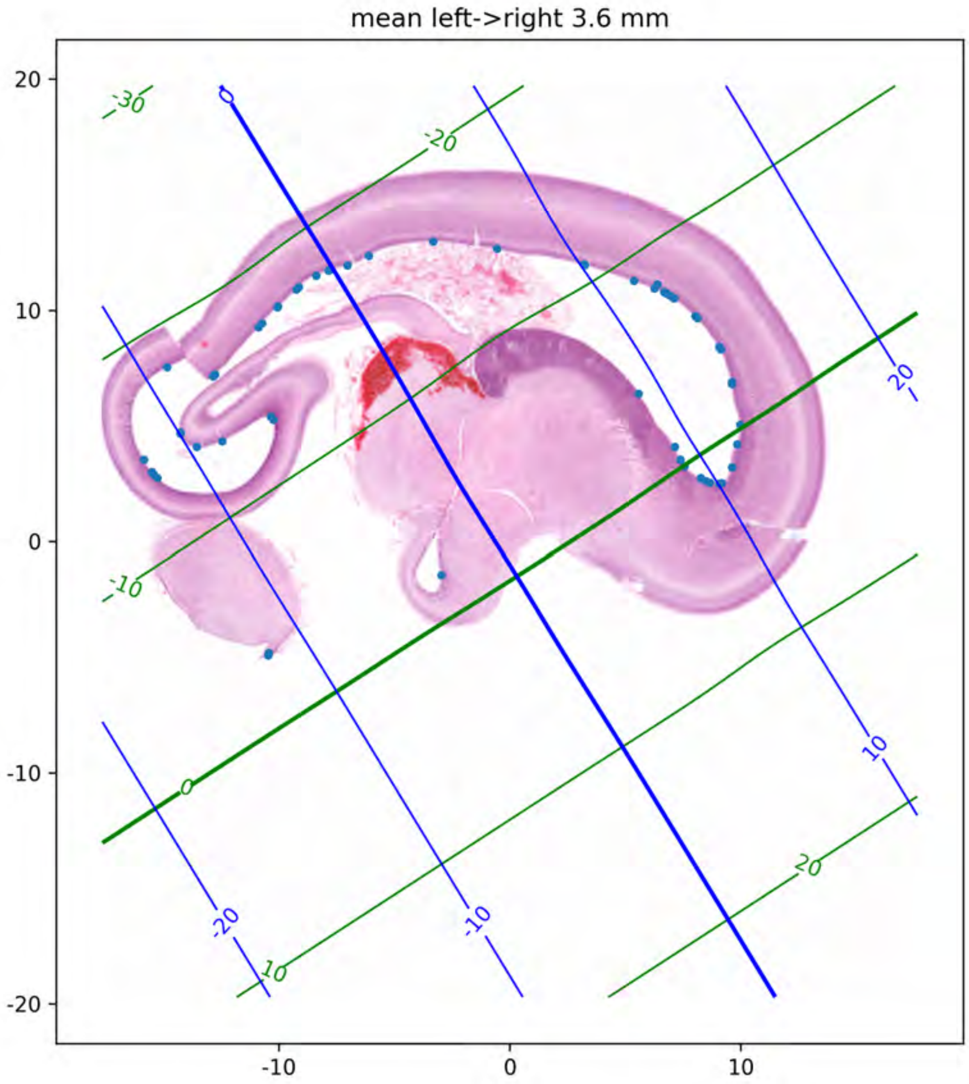

mean left->right 3.6 mm
20
10
0
-10
-20
-10
0
10
-30
-20
-10
0
10
20
0
20
10
-20
-10

mean left->right 4.2 mm

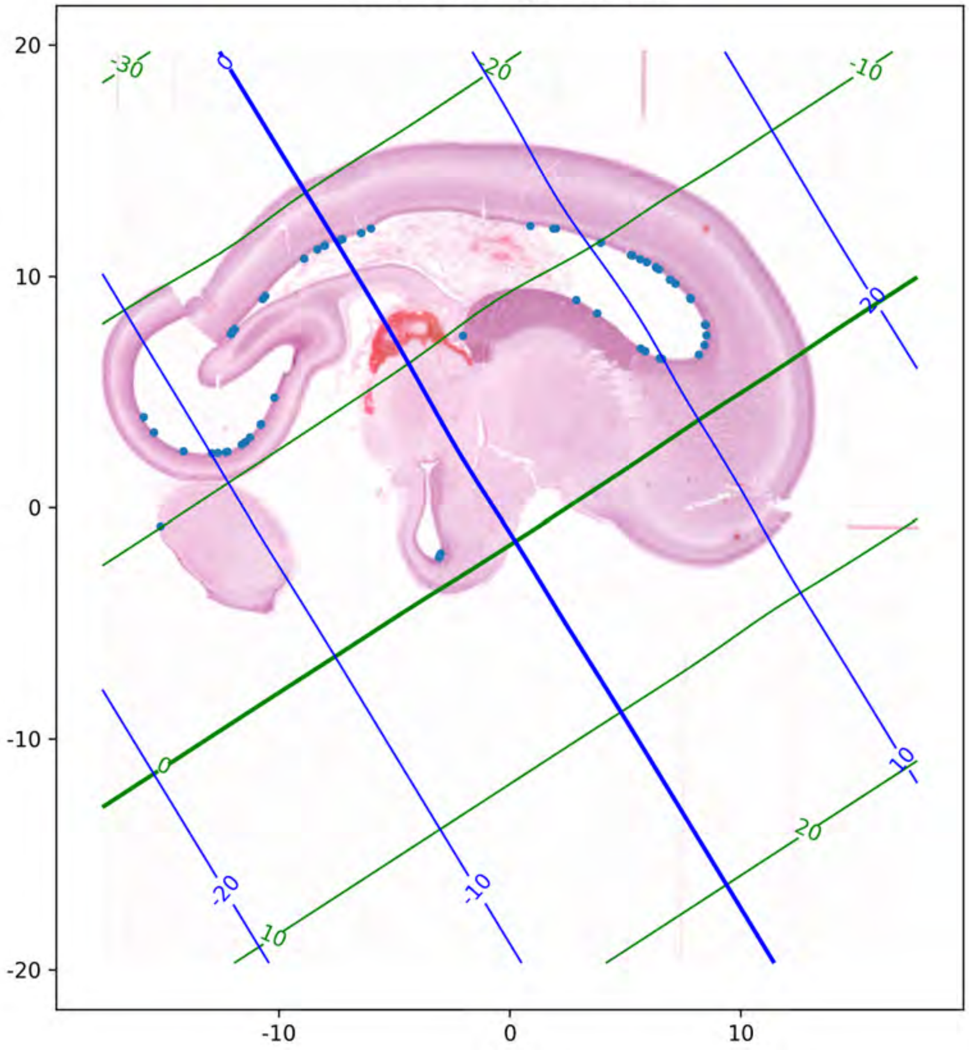

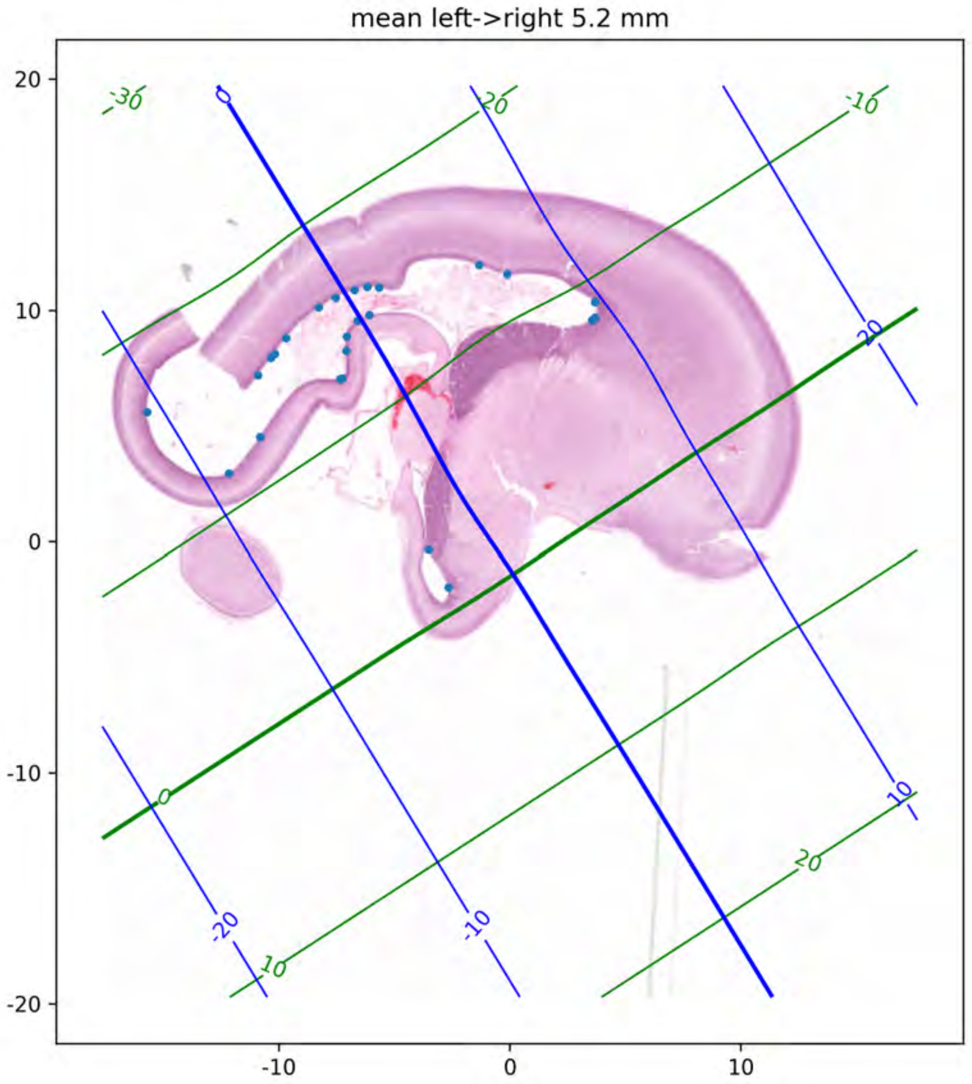

mean left->right 5.2 mm
20
10
0
-10
-20
-10
0
10
-30
-20
-10
0
10
20
-20
-10
0
10
20

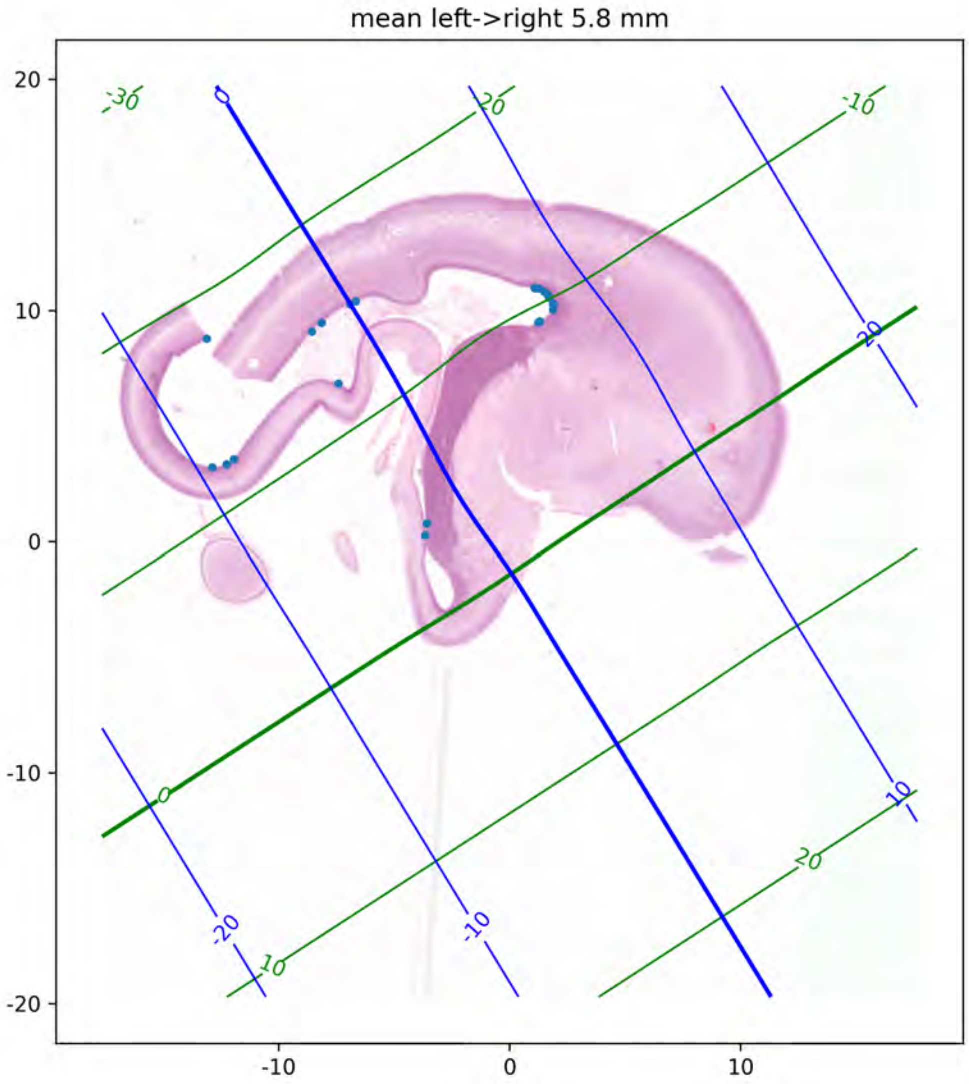

mean left->right 5.8 mm
20
10
0
-10
-20
-10
0
10
-30
-20
-10
0
10
20
-20
-10
0
10
20

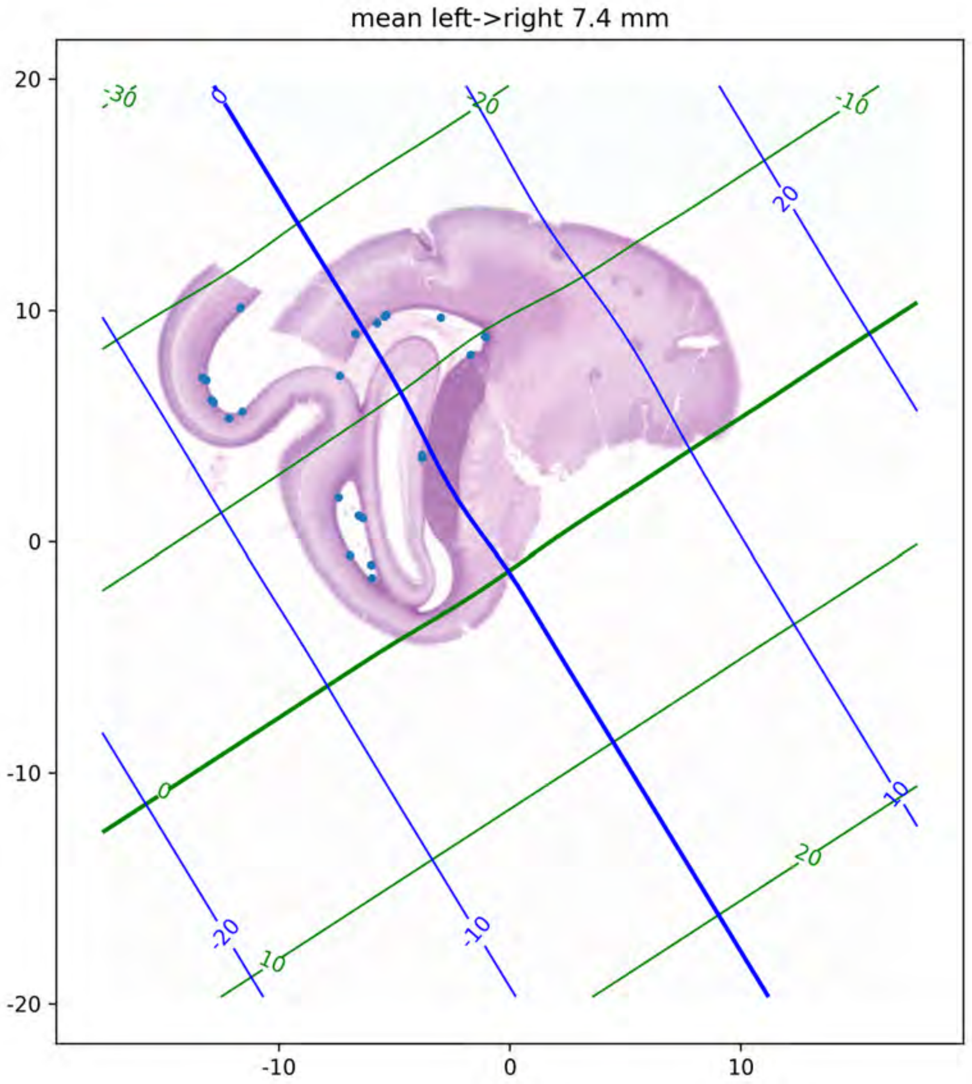
mean left->right 7.4 mm
20
10
0
-10
-20
-10
0
10
-30
-20
-10
0
10
20
0
20
10
-20
-10

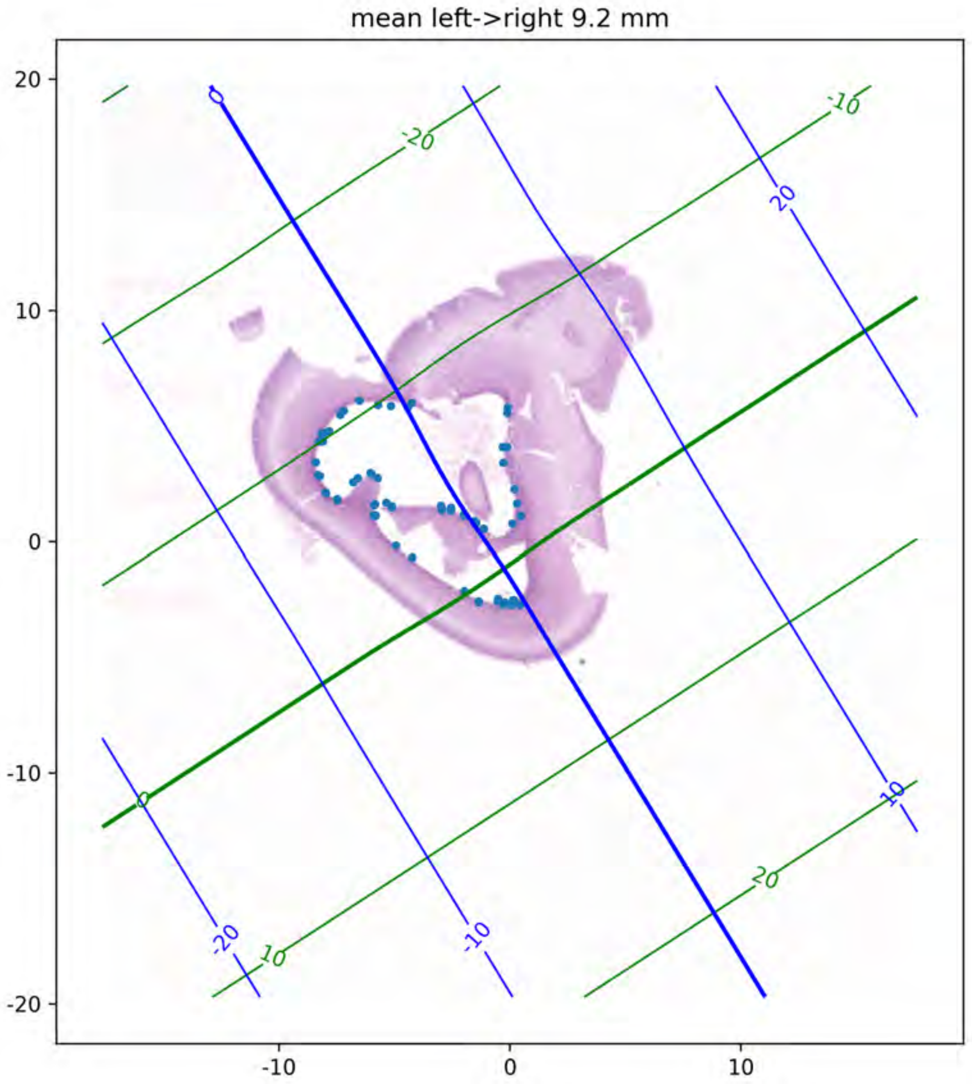
mean left->right 9.2 mm
20
10
0
-10
-20
-10
0
10
-20
-10
0
10
20
